# Resveratrol targets G-quadruplexes to exert its pharmacological effects

**DOI:** 10.1101/2024.07.29.605564

**Authors:** Ana Soriano-Lerma, Victoria Sánchez-Martín, Javier Murciano-Calles, Matilde Ortiz-González, María J Tello-López, Virginia Pérez-Carrasco, Ángel Linde-Rodríguez, Inmaculada Ramírez-Macías, Irene Gómez-Pìnto, Inmaculada López-Aliaga, Miguel Soriano, Jose A. Garcia Salcedo

## Abstract

Resveratrol (RSV) is one of the most studied and used biomolecules, for which many pharmacological effects targeting multiple tissues have been described. However, a common underlying mechanism driving its full pharmacological activity has not been elucidated to date. G-quadruplexes (G4s) are non-canonical nucleic acid structures found in promoters and involved in controlling gene transcription. This study demonstrates a G4-dependent mode of action for RSV, explaining its multi-target traits. RSV was shown to stabilise cellular G4s, which accumulate around double strand breaks (DSBs) in the promoters of differentially expressed genes (DEGs). G4 targeting triggers DNA damage and controls gene expression. Unravelling the main mode of action of RSV will be helpful to improve its therapeutic potential in a wide variety of health scenarios.

## Main Text

Resveratrol (RSV), or 3,4,5-trihydroxy-trans-stilbene, was first discovered in the roots of *Veratum grandiflorum* in 1939(*1*), and first isolated from *Polygonum cupsidatum*(*2*), a traditional chinese medicine. As a natural compound present in food (grapes, groundnuts, several species of berries) and plants(*3*), RSV has drawn a considerable amount of attention due to its numerous beneficial effects. Along the last twenty years, the number of studies involving RSV has considerably increased. This sparked interest might stem from its high accessibility at low cost, both from plants and chemical synthesis, and its wide range of pharmacological activities(*2*).

In human clinical trials, RSV has been related to enhanced cognitive function and reduced neuroinflammation in Alzheimer disease(*4*). Two studies reported reduced levels of beta amyloid, IL-12p40 and IL-12p70 in the cerebrospinal fluid of patients undertaking RSV treatment for 52 weeks. RSV also plays a role in diabetes since it exerts vascular protective functions, promotes fatty acid oxidation in the liver and attenuates oxidative stress, therefore enhancing insulin sensitivity(*4*). Its ability to activate sirtuin 1 (*SIRT1*) and AMP-activated protein kinase (*AMPK*), both involved in the maintenance of cell homeostasis and coordination of metabolic reactions, makes it suitable to treat disorders such as metabolic syndrome or obesity(*5*). RSV also exerts calorie restriction-mimetic effects, inducing the secretion of insulin-like growth factors (IGF), which leads to a reduction in cancer risk(*4*). Notably, RSV is involved in the maintenance of telomere functionality and is considered an antiaging compound(*6*). Lastly, antibacterial, antifungal and antiviral properties have also been reported for RSV(*7, 8*), picturing a broad landscape of applications against a wide range of infectious and non-infectious diseases of the 21st century.

In light of the above, RSV can be considered a multi-target compound, and its biological effects resemble those of a biological tsunami(*9*). Most of research has been focused on studying single targets or isolated pathways, not considering the whole set of biological events orchestrated by RSV when it enters the biological milieu. A common underlying mode of action driving these pharmacological effects has not been discovered to date.

In this sense, non-canonical secondary DNA structures arise as important regulators of gene expression and cell metabolism and represent an open gate for drugs to target multiple tissues and genes(*10*). Among these, G-quadruplexes (G4s) are found in key regulatory sites of highly transcribed genes allowing their regulation at the transcriptional level(*11*). G4s are formed in nucleic acids in physiological conditions through the self-stacking of guanine tetrads. They typically consist of four tracts of two or more guanines separated by loop sequences of variable composition and length. Such tracts self-associate and are further stabilised by cations such as Ca^2+^, Na^+^, and K^+^ (*12*). As a result of the stabilisation of G4s in a physiological setting or by ligands, mainly in promoter regions, gene expression can be either upregulated or downregulated depending on whether these structures block the progression of RNA polymerases or act as docking sites to recruit transcription factors. Therefore, DNA G4s and their ligands are considered key regulators of gene transcription.

In this study, we hypothesized that a G4-dependent mode of action is driving the biological effects of RSV.

### RSV shows G4 ligand effects in a cellular environment

Due to the high GC content in the ribosomal DNA and its ability to form G4s(*13*), analysis of the nucleolus is often used to screen G4s ligands.

The A375 melanoma cell line was chosen as a model to perform the screening due to its nucleolar traits. Nucleolin (NCL) was used as a biomarker for nucleolar disassembly; being naturally accumulated in the nucleolus, a nucleolar segregation of this protein occurs upon nucleolar stressing conditions, such as G4 binding-derived ribosomal DNA (rDNA) damage or inhibition of RNA polymerase I (RNA Pol-I)(*13*).

The 50% inhibitory concentration (IC50) of RSV was initially calculated in A375 cell line at 24h and 6h, yielding a value of 256 *μ*M and 740 *μ*M (Supplementary figure S1, Supplementary figure S2). The IC50 24h dosage (256 *μ*M) was used as a reference for the rest of the experiments in this cell line, using a treatment period of 6h to exclude bias by other RSV targets and to ensure the observation of pharmacological effects. Nucleolar disassembly was produced as a consequence of RSV treatment (IC50 dosage for 6h), with NCL being segregated from the nucleolus to the nucleoplasm in comparison with the control DMSO-treated cells (Figure 1A).

**Fig. 1.**
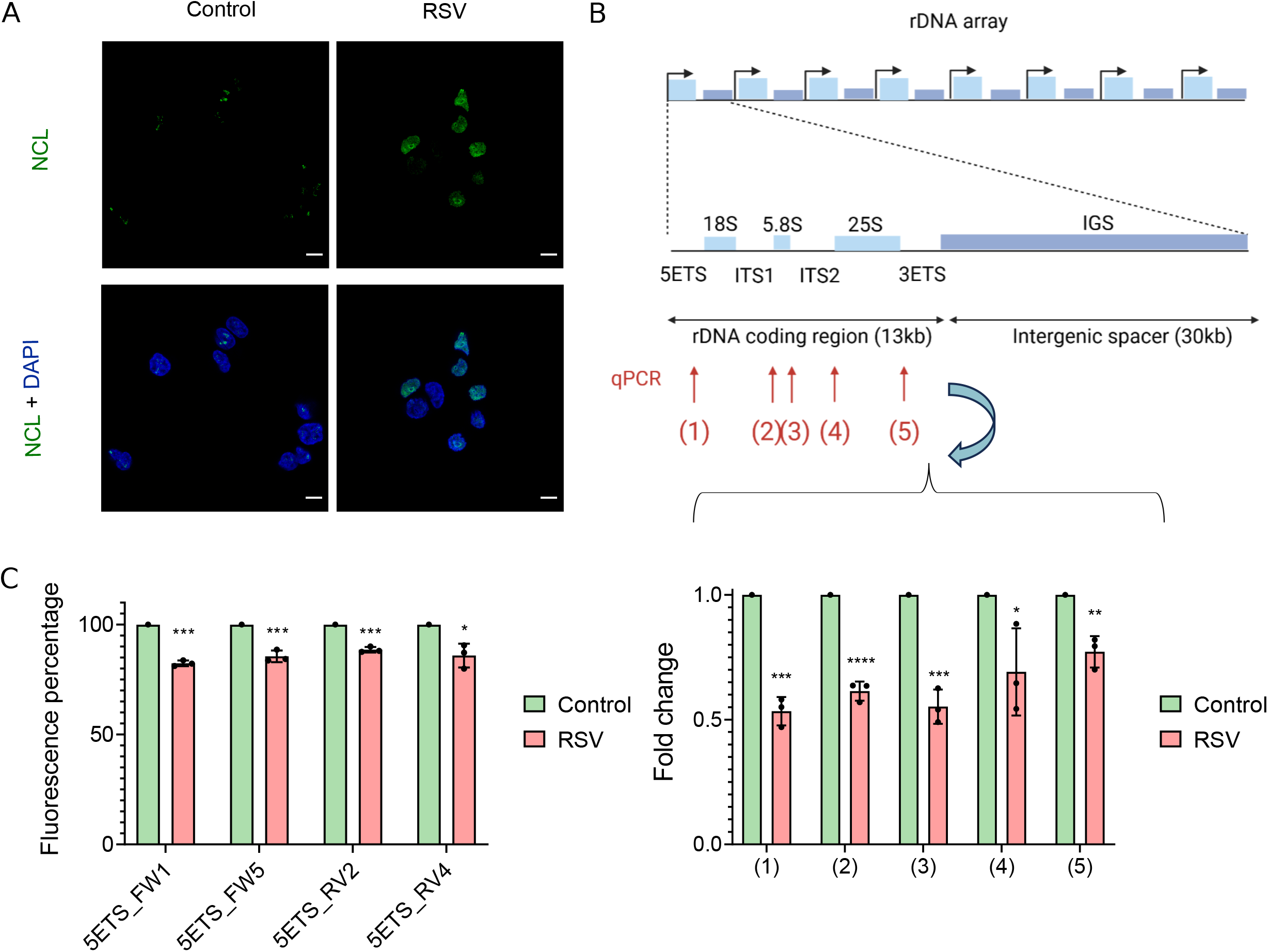
Effect of resveratrol on the nucleolus and RNA polymerase I in A375 cell line and rDNA G4 targeting properties. **(A)** Nucleolin (NCL) staining in the control and treated group (RSV IC50 6h); image shown are representative of three independent experiments (n=3). Counterstaining was performed with DAPI. Scale bar = 10 *μ*m. (B) A schematic view of the rDNA organization in the genomic DNA is represented in the top part. Primers targeting 5’ETS, ITS1, ITS2 and 3’ETS to analyse rDNA transcription are shown with red arrows. A bar diagram representing rRNA expression levels (fold change) in the control and treated group (RSV IC50 6h) is shown in the bottom part. (C) FID assay, bar diagram showing decreasing values of TOPRO3+DNA fluorescence in response to RSV in four G4-containing primers from the 5’ETS region. Error bars indicate standard deviations for three independent experiments (n=3).

Since the inhibition of RNA Pol-I transcription can produce nucleolar disorganization(*13*), rDNA transcription was checked through quantitative PCR (qPCR) using primers from the 5’-external transcribed spacer (ETS) region, intergenic spacers 1 and 2 (ITS1 and ITS2), and 3’ETS region. A 25-50% inhibition of Pol-I mediated transcription was shown upon treatment with RSV (IC50 6h) in relation to the control DMSO-treated cells (Figure 1B).

To check whether RSV was targeting G4s found in the rDNA, the QGRS software was used to find G4 forming sequences in the human ribosomal 5’ETS rDNA region with at least 3 guanines per tract (Supplementary table S1). TOPRO3 fluorescent intercalator displacement (FID) assays were performed using rDNA G4-forming oligonucleotides. RSV exhibited potential G4 binding capacity against four out of twelve tested rDNA G4s-containing primers, decreasing the TOPRO3+DNA fluorescence by 10-20% compared to the control condition (Figure 1C). G4 conformation was checked for positive candidates (5ETS_FW1, 5ETS_FW5, 5ETS_RV2, 5ETS_RV4) in the conditions of FID assay (5 *μ*M TOPRO3, 10 *μ*M DNA in G4s buffer) through circular dichroism (CD), and compared to standard folding conditions (10 *μ*M DNA in G4s buffer). Although TOPRO3 induced changes in the CD spectra, indicating intercalation with the G4, all quadruplexes showed G4 characteristic spectra under the standard and the FID experimental conditions (Supplementary figure S3), suggesting the quadruplexes were correctly folded after TOPRO3 addition.

Additional biophysical binding assays (CD and differential scanning calorimetry [DSC]) were performed on two rDNA G4s, one from the forward and one from the reverse strand, to confirm RSV interaction, and are described further below.

G4 stabilising properties of RSV were investigated *in vivo* through immunofluorescence using BG4 antibody, which recognizes G4 structures in a cellular context(*14*). BG4 immunofluorescence was performed using the IC50 dose (256 *μ*M) for A375 cell line and 6h as experimental conditions. A 10-fold increase in BG4 signal upon treatment was noticed in comparison with the control DMSO-treated cells in the A375 cell line (Figures 2A and 2B).

**Fig. 2.**
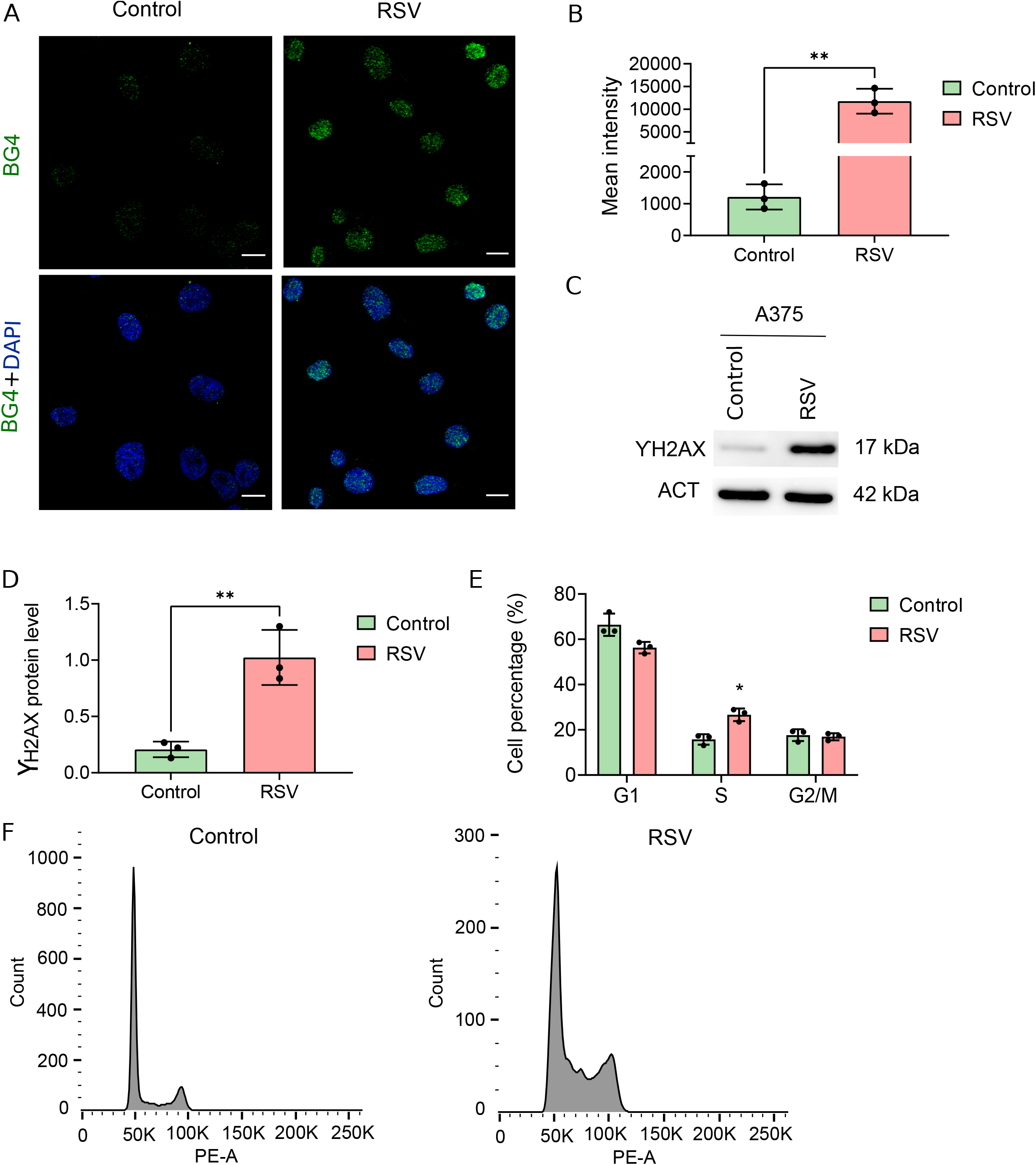
Genome-wide G4 targeting and derived effects. **(A)** Nuclear G4 staining in control and treated conditions in A375 cell line (RSV IC50 6h); images are representative of three independent experiments (n=3). (B) Quantification of one hundred nuclei per condition was performed using ImageJ in the three replicates. (C) ƔH2AX protein levels indicating DNA damage in the control and treated groups in the A375 cell line (RSV IC50 6h). Images are representative of three independent experiments (n=3). (D) Bar plot showing quantified and normalized protein levels relative to actin (ACT) in the three replicates. (E) Bar diagram for cell cycle distribution in the control and treated groups in the A375 cell line (n=3) (RSV IC50 24h). (F) Representative histograms of experiments in (E). Error bars indicate standard deviations.

A common effect of G4 ligands is the production of DNA damage and genome instability(*15*). Consequently, cells have developed different protective mechanisms, including DNA repair pathways, cell cycle checkpoints and break detection methods(*13*). Therefore, as a direct consequence of DNA damage, cell cycle arrest is another shared trait among G4 ligands(*15*).

DNA damage was assessed through Western-Blot using the ƔH2AX antibody and the A375 cell line treated with the IC50 dose for 6h. ƔH2AX signal increased four times upon treatment compared to the control condition (Figures 2C and 2D). Consequently, RSV treatment also triggered cell cycle arrest in the A375 cell line, which was arrested at S phase compared to control DMSO-treated cells when treated with the IC50 dose for 24h (Figures 2E and 2F).

Therefore, RSV exhibited cellular G4 binding properties and shared common traits of other G4 ligands, such as nucleolar disassembly, DNA damage and cell cycle arrest.

### Changes in gene expression produced by RSV are mediated by genomic G4 binding

RNA sequencing (RNA-seq) was performed to characterise changes at the transcriptional level in the A375 cell line (IC50 dose for 6h). 15,000 differentially expressed genes (DEGs) were found (False Discovery Rate, FDR < 0.05) (Figure 3A) (Supplementary File S1). Seven genes were selected along the whole spectrum of log_2_ Fold Change (log_2_FC) values for qPCR validation of mRNA levels. Therefore, qPCR validation was performed for the following downregulated genes: *Aurora Kinase A (AURKA)*, DNA helicase *PIF1, AMMECR1* and *phosphatidylinositol transfer protein beta gene* (*PITPNB)*, and the following upregulated ones: *cadherin 1* (*CDH1), coagulation factor 13 B chain* (*F13B)* and *histone H1*.*4* (*H1*.*4)* (Supplementary figure S4). Functional enrichment analysis was performed using gprofiler2 in upregulated (Supplementary File S2) and downregulated genes (Supplementary File S3) to analyse the changes in functional metabolic pathways upon treatment. Terms related to RNA polymerase II transcription, sequence specific DNA binding, small molecule binding, collagen metabolism, developmental processes and energy obtention (Supplementary figure S5) were shown to be upregulated while downregulated terms included also those related to RNA polymerase II transcription, as well as rRNA metabolic processes, ribosome biogenesis, the nucleolus, DNA synthesis and cell cycle progression (Supplementary figure S6).

**Fig. 3.**
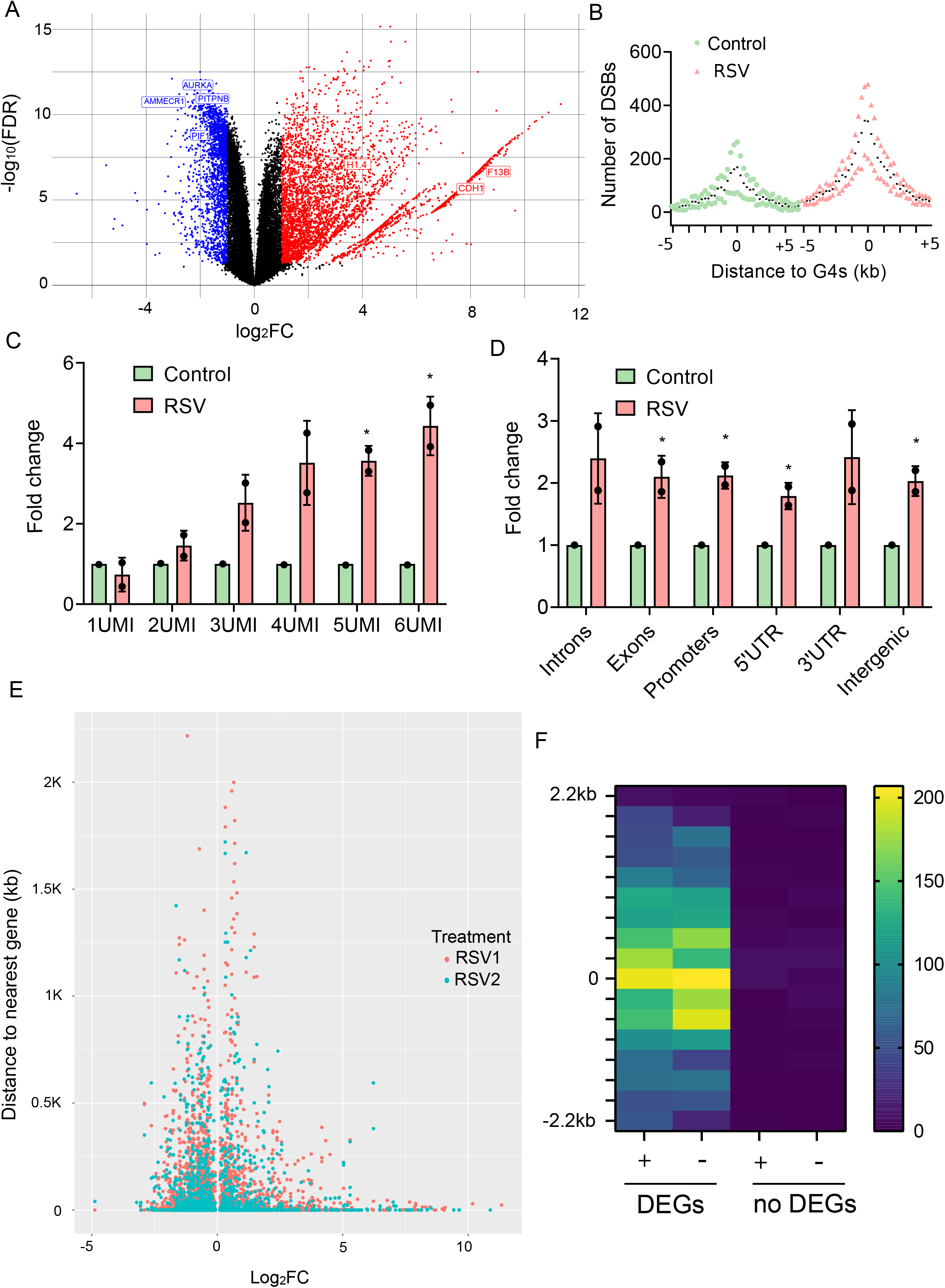
Changes in gene expression in response to RSV treatment are related to G4 targeting. **(A)** Vulcano plot showing differentially expressed genes (DEGs) (FDR<0.05) with log_2_FC < - 1 in blue and log_2_FC > 1 in red (n=2); qPCR validated genes have been highlighted with blue and red labels. (B) Scatter plot showing the number of double strand breaks (DSBs) per distance range (kb) in relation to the nearest G4(*17*), denoted by 0. Black dots indicate the median for each experimental group (n=2). (C) Bar plot for the number of DSBs per UMI slot in response to RSV treatment and expressed as fold change. (D) Bar plot for the number of DSBs in each genomic region in response to RSV treatment and expressed as fold change. (E) Scatter plot showing the relationship between the distance from each DSB to its nearest gene and its log_2_FC (n=2); all DEGs were considered (F) Heatmap showing the number of described G4s(*17*) in the forward (+) and reverse (−) strand per distance range (kb) in relation to the nearest DSB (denoted by 0) in treated samples; all DEGs and genes with no differential expression were considered. Error bars indicate standard deviations.

Aiming to identify G4s targeted by RSV, Breaks Labelling *In Situ* and Sequencing (BLISS)(*16*) was next performed in the A375 cell line to map double strand breaks (DSBs) produced in response to treatment (IC50 dose for 6h). DSBs tended to accumulate around previously described G4 regions(*17*) in the treated and control conditions, although RSV treatment doubled the total number of DSBs (Figure 3B). BLISS adapters include unique molecular identifiers (UMIs), 8 nucleotide long random sequences that label DSBs at a single cell and nucleotide resolution; they are indicative of recurrent DSBs that occurs repeatedly in the same position in different cells(*16*); DSBs with 5 and 6 UMIs increased after RSV exposure around four times compared to the control condition (Figure 3C), which suggests RSV treatment induces DNA damage in highly specific locations. DSBs were mapped onto genomic features (introns, exons, promoters, 5’UTR region, 3’UTR region and intergenic regions), showing all of them around twice more DSBs in the treated compared to the control condition except for introns and 3’UTR regions, where differences were not statistically significant (Figure 3D).

To relate G4 targeting by RSV to changes in gene transcription analysed by RNA-seq, the distance from each DSB to its nearest gene was calculated and represented against the log_2_FC value obtained for that gene from the RNA-seq analysis, considering only DEGs. An inverse correlation was found (Figure 3E), since shorter distances from DSBs to their nearest genes conditioned grater changes in log_2_FC. This result suggests that G4 targeting by RSV influences gene expression.

To investigate the potential of DSBs used as a mapping tool to find RSV targets involved in the control of gene expression, the distribution of described G4s(*17*) was analysed in relation to DSBs found in the promoter of DEGs (Supplementary File S1) and genes without differential expression. G4s clustered around DSBs only in the case of DEGs, both in the forward and reverse strand (Figure 3F), suggesting DSBs occur as a consequence of G4 targeting and the modulation of gene expression.

Therefore, *in vitro* binding assays were next performed using candidate G4s obtained from the proximity of DSBs located in the promoters of DEGs. In particular, validated DEGs showing recurrent DSBs in their promoters were selected for the identification of candidate G4s (*AMMECR1, PITPNB, F13B* and *H1*.*4*). Putative G4 forming sequences (PFS) were analysed using QGRS software (Supplementary table S1), 100bp upstream and downstream the DSB. No threshold was established in relation to the G-Score because it’s been shown that the number of G4s arising from ligand interactions are often non-canonic and display alternative structures (G4s with 2 guanine tracts)(*18*). Non-overlapping PFS showing the highest G-Score per gene were selected and screened for G4 formation by CD; PITPNB showed the spectra most similar to G4s (Supplementary figure S7). Therefore, binding assays were performed using PITPNB G4.

Circular dichroism (CD) and ultraviolet-visible (UV-vis) titration assays were carried out with the pre-folded oligonucleotide PITPNB. CD spectra of 10 *μ*M DNA showed a positive band around 260 nm with a negative band around 240 nm, which are indicative of parallel G4 structures(*19*) (Figure 4A). Changes in the CD spectra were noticed upon addition of RSV 100 *μ*M, with the maximum of the positive band being sightly displaced towards shorter wavelengths (Figure 4A). This result indicates an interaction between RSV and the G4 structure. UV-vis titration assays were performed to confirm the observed binding. The presence of at least one isosbestic point in the UV-vis spectra upon addition of RSV confirmed RSV interaction with PITPNB G4 (Figure 4B).

**Fig. 4.**
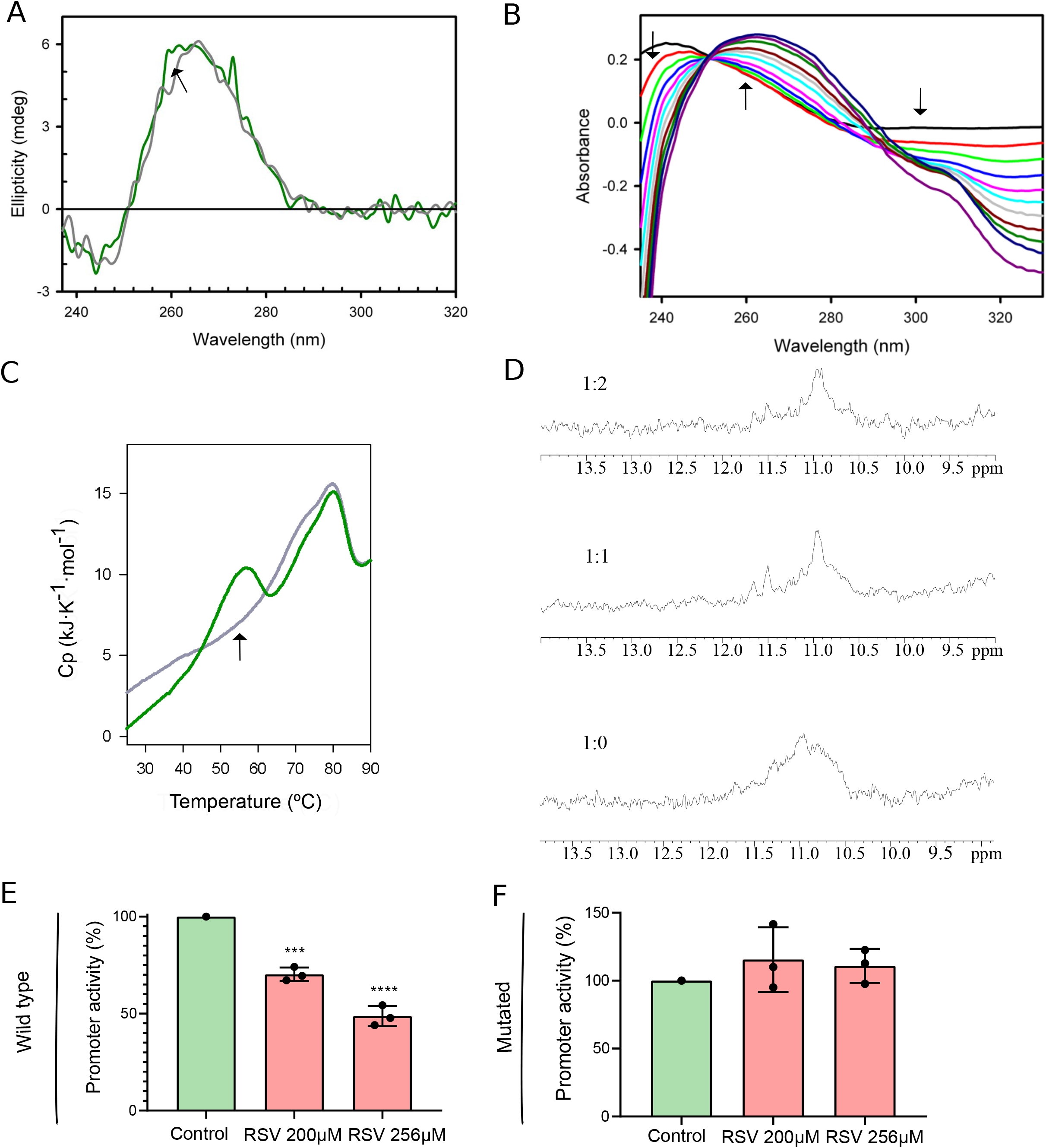
Binding assays and luciferase-based transcription assays confirm that RSV targets a G4 in the promoters of a DEG, affecting gene expression. **(A)** CD spectra of PITPNB G4 in the presence (green line) and absence (grey line) of RSV. Arrows indicate the direction of the movement of CD peaks upon addition of RSV. (B) UV-vis spectra resulting from the titration of PITPNB G4 with RSV. Arrows indicate the direction in which the absorption peak moves after each addition of RSV. (C) Thermal melting profiles showing excess heat capacity as a function of temperature for PITPNB with (green line) and without RSV (grey line); the arrow indicates the ligand-dependent conformational state (D) Exchangeable proton region of the NMR spectra of PITPNB with RSV at different DNA:RSV ratios. (E) Bar plot showing the inhibition of the promoter activity of *PITPNB* gene as a consequence of RSV treatment in the A375 cell line. (F) Bar plot showing changes in the activity of the mutated promoter of *PITPNB* gene as a consequence of RSV treatment in the A375 cell line. Error bars indicate the standard deviation of three independent experiments (n=3).

DSC and nuclear magnetic resonance (NMR) were performed to confirm the interaction of RSV with PITPNB G4. DSC was used as a melting high-sensitivity method to gain insight into DNA-ligand interactions. The thermogram of PITPNB shows a main transition of the folded to the unfolded state at 80ºC (Figure 4C, grey line). Upon the addition of RSV, an additional transition is shown at 56ºC (Figure 4C, green line), implying the presence of another state that is generated in presence of the drug, hence confirming interaction.

The NMR analysis of PITPNB confirmed the G4 folding with guanine imino signals appearing between 10.5 and 12.5 ppm (Figure 4D). In this case, sharp signals were not appreciated probably due to the formation of high-order structures as already described with other G4s with similar primary sequence(*20*). After the addition of RSV, imino NMR signals changed due to interaction between RSV and PITPNB G4 (Figure 4D). The narrowing of the signals upon RSV addition to PITPNB is indicative of the formation of lower-order structures resulting from the ligand-induced individual separation of the G4. The effect of DMSO on PITPNB G4 using the concentration equivalent to the 1:1 ratio was excluded (Supplementary figure S8). RSV hydroxyl (−OH) signals also flattened by the addition of PITPNB (Supplementary figure S9), confirming interaction.

To demonstrate that RSV binding is structure-specific, a DNA duplex structure was designed and tested through DSC and NMR. The thermogram of the D1 oligonucleotide is unfolded at 53ºC (Supplementary figure S10, grey line). Upon RSV addition, no additional calorimetric transitions appeared in the thermogram (Supplementary figure S10, green line), confirming the lack of interaction between the drug and the oligonucleotide. The duplex structure was confirmed by NMR with A-T and C-G signals appearing between 12 and 14 ppm (Supplementary figure S11). Upon addition of RSV, signals were mostly unchanged, confirming the absence of a significant interaction (Supplementary figure S11).

Two rDNA G4s, one from the forward (5ETS_FW1) and one from the reverse strand (5ETS_RV4) (Figure 1C) were also tested through CD and DSC to confirm RSV binding. CD spectra also showed parallel G4 structures (Supplementary Figure S12A and S12B); upon RSV addition, the 260 nm maximum was displaced towards longer wavelengths in 5ETS_FW1 (Supplementary Figure S12A) and decreased in the case of 5ETS_RV4 (Supplementary Figure S12B). The thermogram of 5ETS_FW1 shows a folded-to-unfolded transition at 71ºC (Supplementary Figure S12C, grey line). Moreover, and similar to PITPNB DSC analysis, an additional transition not shown in the oligonucleotide alone is shown at 42ºC (Supplementary Figure S12C, grey line vs green line), implying the presence of an additional state that appears as a consequence of the drug, hence confirming interaction. A similar thermogram to PITPNB and 5ETS_FW1 was obtained for 5ETS_RV4, with the main transition occurring at 83ºC (Supplementary Figure S12D, grey line) and the ligand-dependent one at 50ºC (Supplementary Figure S12D, green line).

To prove the direct consequences of G4 targeting on gene expression, a luciferase-based transcription assay was performed using the A375 cell line. *PITPNB* gene was chosen due to the already characterised interaction of RSV with the PITPNB G4 found in its promoter. As the *PITPNB* promoter (−500 to +100bp) is found on the reverse strand of the genome, it was cloned as the forward strand into a promoter testing vector, leading the transcription of the firefly luciferase gene. A *Renilla* luciferase normalizing gene was also included in the same vector under the *Tyrosin Kinase (TK)* promoter. Following cell transfection, doses of 200 *μ*M and 256 *μ*M of RSV were applied for 6h and the luminescence measured for firefly and *Renilla* luciferase. The promoter activity was progressively inhibited by around 25% and 50% (Figure 4E). A mutated plasmid where PITPNB G4 was unable to form was included in the experiment as a control and used in the same conditions. No inhibition of the promoter activity in the mutated plasmid could be observed upon addition of RSV 200 *μ*M and 256 *μ*M (Figure 4F), confirming that RSV binds to PITPNB G4 *in vivo* and inhibits gene transcription.

### G4 targeting by RSV is independent of SIRT1 activation

SIRT1, a histone deacetylase, has long been considered a classic target of RSV(*9*). Several studies have reported an increase in the transcription and activity of *SIRT1* gene in presence of RSV(*21, 22*). To confirm that the G4-dependent mode of action is independent of SIRT1 activation, mRNA levels of *SIRT1* were measured upon treatment with RSV (IC50 dosage 6h) using qPCR. *SIRT1* gene expression did not vary between the treated and control conditions in the A375 cell line (Supplementary figure S13A). The enzymatic activity of SIRT1 was evaluated through the analysis of the acetylation mark in histone 3 lysine 9 (H3K9ac). Similar levels of acetylation were found in the treated and control conditions (Supplementary figure S13B and 13C), confirming that SIRT1 is not mediating the observed mode of action, as already reported(*23*).

## Discussion

This study reveals, for the first time, the underlying mode of action of RSV, targeting specific DNA G4s throughout the genome, which explains its multi-target traits. RSV targets rDNA G4s, resulting in nucleolar disassembly and inhibition of RNA Pol I-mediated transcription. This action could explain the anti-cancer effects of RSV in different human cancers, such as breast, uterine, blood or kidney(*24*). Cancer cells show a higher ribogenesis to meet an increased demand for protein synthesis; in fact, the inhibition of rRNA synthesis has been described as a mode of action of anticancer drugs with a G4-dependent mode of action(*25*). RSV was also shown to target promoter-located G4s, such as PITPNB, directly influencing gene expression. Stabilised G4s in promoter regions can either block RNA polymerases movement, inhibiting gene transcription, or act as docking sites for the recruitment of transcription factors, promoting gene transcription(*26*). Both mechanisms justify the pleiotropic effects of RSV on gene expression.

An overall increase in BG4 signal was reported in response to treatment, confirming RSV is targeting G4s at the genomic level. Other studies have also described RSV as a G4 ligand, but only for specific G4s in an *in vitro* setting; moreover, in these studies G4 targeting was not associated with a direct modulation of gene expression(*27-29*). As a consequence of G4 targeting, G4 ligands exert DNA damage and cell cycle arrest(*30*), both of which have been reported in this study. These results are in line with a recent study showing that replicative stress and cell cycle arrest are triggered as a consequence of RSV treatment in NALM6 and U2OS cell lines (*23*).

Upregulated processes from RNA-seq in response to treatment included those involved in RNA polymerase II transcription, sequence specific DNA binding and small molecule binding, all of them in support of G4 targeting by RSV. As an antiaging compound that enhances bone morphogenesis(*31*), RSV also upregulates pathways related to collagen metabolism. Downregulated processes upon treatment included those related to the cell cycle, rRNA processing and ribosome biogenesis, which is also supports our hypothesis.

BLISS analysis revealed a higher number of DSBs with 5 and 6 UMIs as a consequence of RSV treatment, which confirms recurrent breaks at specific regions. The exact mechanism by which RSV exerts DNA damage still remains unknown, as recently reported(*32*). Our results show that DSBs accumulates around G4s upon treatment, and that G4s cluster around DSBs only in the promoter of DEGs, which shows that RSV binds G4s present in the promoters to control gene expression. G4s are therefore DNA damage hotspots, which highlights the utility of BLISS to map target G4s at single-nucleotide resolution. The use of DNA damage as a mapping tool to identify target G4s has already been described for the well characterised G4 ligand pyridostatin, but using a ƔH2AX chromatin immunoprecipitation (Chip) – sequencing protocol instead(*30*). However, BLISS allows a higher resolution to map DSBs compared to ƔH2AX Chip-seq (*16*).

RSV was found to interact with PITPNB, a G4 found in the promoter of the *PITPNB* gene. Phosphatidylinositol transfer proteins (PITP) are the main regulators of the phosphoinositide metabolism, whose main substrate is phosphatidylinositol(*33*). PITPs regulate the transport and distribution of phosphatidylinositol in different organelles to join the phosphoinositide synthesis and metabolism, which is known to be involved in biological processes relevant to RSV effects, such as the regulation of tumourigenesis, neuronal development or immune cell migration(*34, 35*).

This study demonstrates that RSV regulates gene expression through G4 binding at promoters, which is the main mode of action underlying its pleiotropic effects. We also provide a detailed characterisation of genomic spots where the interaction occurs. The described mechanisms will allow a more rational use of RSV and a better assessment of its therapeutic applications and side effects.

## Acknowledgments

The authors would like to thank the Genomics Unit at the Centre for Genomic Regulation (CRG, Barcelona, Spain) for their assistance in the sequencing service. We acknowledge the ‘‘Manuel Rico’’ NMR laboratory (LMR), a node of the ICTS R-LRB.

## Funding

Provide complete funding information, including grant numbers, complete funding agency names, and recipient’s initials. Each funding source should be listed in a separate paragraph.

Junta de Andalucía: PAIDI research group BIO-344

Ministry of Science and Innovation of Spain grant PID2020-120481RB-I00/AEI/10.13039/50110001103 (MS)

Instituto de Salud Carlos III (ISCIII), grant PI21/00497, co-funded by the European Union (JAGS)

University of Granada grant CONTRATOS PUENTE (ASL)

Ministry of Universities of Spain grant FPU 20/03952 (VPC)

## Author contributions

Conceptualization: MILA, MS, JAGS

Cell assays: VSM, MOG, ALR, VPC, MJTL, ASL

Biphysical assays: JMC, IGP, ASL

Bioinformatic analysis: ASL

Funding acquisition: IRM, MILA, MS, JAGS

Project administration: ASL

Supervision: MS, JAGS

Writing – original draft: ASL

Writing – review & editing: MILA, MS, JAGS

## Data and materials availability

Sequencing data derived from this study have been uploaded to the Sequence Read Archive (SRA) under the accession number PRJNA986468.

R codes used in bioinformatic analyses will be provided in the auxiliary supplementary material. Linux code to process BLISS fastq files is available at https://github.com/garner1/BLISS.

## Supplementary Materials

Materials and Methods

Figs. S1 to S13

Tables S1 to S5

## Auxiliary supplementary materials

Supplementary file S1, S2 and S3

