## Supplementary material for "Resveratrol targets G-quadruplexes to exert its pharmacological effects"

**The PDF file includes:**

Materials and Methods
Figs. S1 to S13
Tables S1 to S5

**Other Supplementary Materials for this manuscript include the following:**

Supplementary file S1, S2 and S3

### Materials and Methods

Cell lines and treatment

A375 cell line was acquired from the American Type Culture Collection (ATCC). A375 cell line was originally isolated from the skin of a female patient with malignant melanoma and grown in DMEM medium supplemented with 10% fetal bovine serum, penicillin (5000 U/mL), streptomycin (5 mg/mL) and amphotericin (25 mg/mL). Culture conditions were 37ºC and 5% $CO_2$ atmosphere.

Resveratrol was acquired from Merck (R5010, Merck), dissolved in dimethyl sulfoxide (DMSO) (D2650, Sigma Aldrich) and stored at -20ºC protected from light. Treatment was applied in a proportional volume relative to the number of cells treated (100 µL per $8 \cdot 10^3$ cells treated).

Cell proliferation assays

Cells were seeded into 96-well plates ($8 \cdot 10^3$ cells/well) and treated with RSV at concentrations ranging from 1 mM to 976 nM in the case of the A375 cell line (24h) and 2 mM to 1.95 µM for the A375 cell line (6h). After 24h or 6h (A375), Resarzurin (R7017, Sigma Aldrich) was added at a final concentration of 0.025 mg/mL and incubated for 4h. Fluorescence was determined using Nanoquant Infinite M200 Pro multi-plate reader (Tecan). IC50 values were determined by non-linear regression with GraphPad Prism.

Cell cycle analysis

A375 cells ($10^6$ cells/condition) were seeded into 10 cm culture dishes and treated with RSV 256 µM for 24h. An appropriate vehicle (DMSO) was added in equivalent concentration to the treated condition as a control. Cells were then fixed on ice with ice-cold 70% ethanol and stained with 0.04 mg/mL propidium iodide (P4864, Sigma Aldrich) and 0.1 mg/mL ribonuclease A (19101, Qiagen). Cell cycle distribution was determined by an analytical DNA flow cytometer (FACSCanto II, BD Biosciences) and the FlowJo software. Experiments were performed in biological triplicates.

Immunofluorescence

A375 cells ($8 \cdot 10^4$ cells/condition) were seeded into 13 mm circular coverslips and placed in 24-well plates; RSV treatment was added at appropriate concentration following incubation for 6h. Fixation was performed with 4% (v/v) paraformaldehyde (P6148, Sigma Aldrich) for 10 min at room temperature (RT), permeabilization with 0.1% (v/v) Triton-X100 (T8787, Sigma Aldrich) for 10 min and blocking with 10% bovine serum albumin (A7906, Sigma Aldrich), 0.5% (v/v) Triton-X100 for 30 min at RT. Primary antibodies were subsequently incubated for 1h at RT for NCL staining or following the standard protocol in case of the BG4 antibody(*1*). Secondary antibodies were incubated for 30 min at 4ºC in darkness. All coverslips were mounted with Vectashield (H-1200, Vector Laboratories) including DAPI (4',6-diamidino-2-phenylindole) for nuclear counterstain. Images were acquired through a Confocal Zeiss LSM 710 inverted microscope with a 63x immersion objective.

Antibodies used included NCL (Invitrogen, Cat #39-6400) and synthetic BG4(*1*) (Supplementary table S2). Standard immunofluorescence procedures were used for BG4 immunofluorescence(*1*). Dilutions of the antibodies used are summarised in Supplementary table S3.

Histone isolation

For histone isolation, $2.5 \cdot 10^6$ cells/condition were seeded and incubated overnight for adhesion. After RSV treatment, cells were harvested and washed in ice-cold PBS and resuspended in Triton Extraction Buffer (TEB: PBS containing 0.5% Triton X 100 (v/v), 2 mM phenylmethylsulfonyl fluoride PMSF, 0.02% (w/v) sodium azide) at a cell density of $10^7$ cells/mL. Cells were lysed on ice for 10 min with gentle stirring. After centrifugation at 650g for 10 min (4ºC), supernatant was discarded and nuclei resuspended in half the volume of TEB. After another centrifugation round using the same conditions, the pellet was resuspended in one quarter of the original volume of HCl 0.2N. Histones were acid extracted overnight (4ºC), samples centrifuged again at 650g for 10 min (4ºC) to pellet debris and the supernatant recovered and neutralised using 2M NaOH (1:10 relative to the recovered volume). Protein content was determined using Bradford assay (5000006, Bio-Rad).

Western-Blot

$4 \cdot 10^5$ cells/condition were seeded in a 6-well plate. Cells were treated with RSV at appropriate concentration for 6h. Total protein was extracted using RIPA lysis buffer containing 1% PMSF (P7626, Merck) and 1% protease inhibitor cocktail (P8340, Merck). Total protein content (12 µg) for ɎH2AX determination or histone extracts (1 µg) for analysis of acetylation marks (H3K9ac) were loaded on 15% SDS-polyacrylamide gels for electrophoresis and semi-dry transferred to nitrocellulose membranes (66485, Pall corporation) using Trans-Blot (Bio-Rad). Membranes were blocked with 5% semi-skimmed milk and incubated overnight at 4ºC with primary antibodies. Next, they were incubated with Horseradish Peroxidase (HRP)-labelled secondary antibodies for 1h at RT. After incubation with HRP substrate (1705060, Bio-Rad), chemiluminescence signals were measured using Image Quant LAS 4000 (GE Healthcare Life Sciences). Protein levels were quantified and normalised in relation to actin (ACT) or histone 3 (H3) levels by ImageJ. Experiments were performed in biological triplicates.

Antibodies used are listed in Supplementary table S2; conditions used are summarised in Supplementary table S3.

Quantitative PCR (qPCR)

A375 cells were seeded at a density of $4 \cdot 10^6$ cell/condition in a 6-well plate, grown overnight and treatment applied the following morning. Total cellular RNA was isolated from cells under different experimental conditions with Trizol Reagent (15596, Invitrogen). Reverse transcription was performed using RevertAid First Strand cDNA Synthesis Kit (K1622, Thermo Scientific) with random primers and 1 µg of RNA, according to the manufacturer's protocol. A previous treatment with TURBO Dnase (AM2238, Thermo Scientific), according to the manufacturer's protocol, was applied before reverse transcription when analysing rDNA transcription. Quantitative PCR was conducted on QuantStudio 6 (Thermo Scientific) with SYBR Green (4309155, Thermo Scientific), a final primer concentration of 500 nM and using 2 µL of previously diluted cDNA (1:10). Target mRNA levels were normalized in relation to actin and fold change was subsequently determined. Experiments were conducted in biological triplicates. Primers used for this study are listed in Supplementary table S4.

Image acquisition

Images were obtained with a confocal microscope (Confocal Zeiss LSM 710). Mean nuclear fluorescence intensity was measured using ImageJ (n≥100). To allow comparability, instrument settings were equally adjusted across treatment conditions.

Fluorescent intercalator displacement (FID) assay

For FID assays, 5 µM TOPRO3 (T3605, Thermo Scientific) was used as a fluorescent intercalator, being saturated with 10 µM pre-folded G4s and titrated with RSV 100 µM in G4s buffer in 96-well plates. After a 30min incubation, TOPRO3 was excited at 642 nm and the emission profile was monitored between 650–800 nm with a multi-plate reader (Nanoquant Infinite M200, Tecan). All assays were conducted in biological triplicates. Fluorescence values were calculated as follows: %Fluorescence = A/B·100; where (A) is the fluorescence value in presence of RSV and (B) corresponds to the fluorescence value in RSV-free controls. Primers used in FID assay are described in Supplementary table S5.

G4s pre-folding

G4 containing oligonucleotides listed in Supplementary table S1 and S5 were purchased from Integrated DNA Technologies (IDT). Upon arrival from the supplier, all oligonucleotides were dissolved in G4s buffer (10 mM potassium phosphate buffer containing 100 mM potassium chloride at pH 7.0), heated at 95ºC for 10 min, slowly cooled to RT and stored at 4ºC until analysis.

RNA sequencing

RNA sequencing was performed in biological duplicates. Total RNA was isolated using Trizol reagent (15596, Invitrogen), and its quality was assessed through the determination of the RNA Integrity Number (RIN) via 2100 Bioanalyzer Instrument (Agilent Technologies, Santa Clara, CA, USA). mRNA libraries were prepared using 500-1000 ng of total RNA as an input and the polyA selection protocol (TruSeq stranded mRNA Library Prep, 20020594, Illumina). Libraries were then sequenced using the Hiseq2500 platform (Illumina, San Diego) and the paired-end 50bp format.

A total of 25-30 million reads per sample were obtained and mapped using *Rsubread*. Reads were aligned and annotated using the human genome hg38 as a reference, downloaded from http://hgdownload.soe.ucsc.edu/goldenPath/hg38/. Further analyses in R software were performed using *edgeR* package. Differentially expressed genes were obtained using p-adjusted values < 0.1. Functional analysis to determine differentially upregulated and downregulated metabolic pathways were performed and plotted using *gprofiler2* package and the Reactome and Gene onthology (GO) databases. The R code used in the bioinformatic and statistical analysis of RNA-seq data is provided as an auxiliary supplementary file.

Candidate genes were validated using qPCR as described earlier. Primers used are listed in Supplementary table S4.

Breaks Labelling In situ and Sequencing (BLISS)

Library preparation for BLISS was performed as previously described, in biological duplicates(*2*). Size selection was done using 6% Novex TBE PAGE Gels (ref. EC6265BOX) and the final pool was quantified by qPCR using the KAPA Library Quantification Kit (KK4835, Kapa Biosystems). Libraries were sequenced via NextSeq 500 platform (Illumina, San Diego), using the 75 + 8 bp, single-ended format kit (Illumina, San Diego) and yielded a total of 122683723 reads (9.2 Gb), between 18-40 million reads per sample to ensure enough coverage. DSBs were identified following the already described pipeline(*2*). FASTQ files were quality filtered using a Phred score > 30 for every base. The filtered reads were scanned on the basis of the presence of 8 nucleotide UMIs and sample barcode. After removal of the prefix, the reads were aligned against the human reference genome GRCh37/hg19 retaining reads mapping with a quality score > 5. Next, a further filtering step based on UMI sequences was performed to filter out PCR duplicates. Finally, BED files containing a list of genomic locations with unique UMIs were generated to be used in downstream analyses. Processing of BED files was performed in R software; UMI values were transformed into UMIs per million (UPMs) according to the following formula: Number of UMIs/Number of mapped reads * $10^6$. Since DSBs with 3 or more UMIs showed a statistical tendency upon treatment ($p<0.1$), BED files were filtered using an UPM cut off of 0.15 for all samples. This value allowed us to keep DSBs with 2 UMIs onwards in samples with lower sequencing depth or 3 UMIs onwards in samples with higher sequencing depth. DSBs were then mapped to genomic features (introns, exons, 5'UTR regions, 3'UTR regions, promoters and intergenic regions for the remainder) using the *TxDb.Hsapiens.UCSC.hg19.knownGene* and *GenomicRanges* packages, having previously eliminated overlapping regions with the *reduce* function. Distances of each DSB to its nearest described G-quadruplex (*3*) were calculated using the *GenomicDistributions* R package. Bed files containing sequenced G-quadruplexes under potassium stabilizing conditions were downloaded from https://www.ncbi.nlm.nih.gov/geo/query/acc.cgi?acc=GSE63874:GSE63874_Na_K_12_minus.bedGraph and GSE63874_Na_K_12_plus.bedGraph, and processed using the *plyranges* R package. G-quadruplexes were filtered using 18% mismatch as the cut off value(*3*). The study of the relationship between the distance from each DSB to its nearest gene and $log_2$FC values was performed using the *plyranges* and the *TxDb.Hsapiens.UCSC.hg19.knownGene* R packages. The analysis of the distance from described G4s(*3*) to DSBs in the promoter DEGs and genes without differential expression was performed using the *GenomicRanges*, *GenomicDistributions* and *TxDb.Hsapiens.UCSC.hg19.knownGene* R packages. First, gene promoters were downloaded from a txdb object. After merging the two RSV replicates, DSBs and described G4s(*3*) contained in these promoters were subset using the *GenomicRanges* package. Distance from G4s to DSBs was calculated via the *GenomicDistributions* package.

R Code used to analyse BLISS data is supplied as auxiliary supplementary files. Linux code to process BLISS fastq files is available at https://github.com/garner1/BLISS.

Promoters of validated DEGs with recurrent DSBs were obtained along with their genomic coordinates using the *TxDb.Hsapiens.UCSC.hg19.knownGene* and *GenomicDistributions* packages. BLISS-derived SORTED.BAM files (generated during the bioinformatic processing of FASTQ files) were loaded into the Integrative Genomics Viewer (IGV). Files for treated samples were checked using the obtained genomic coordinates. G4 containing sequences were obtained from the proximity of DSBs (100bp up and downstream the DSB) and the G4 forming potential checked using QGRS mapper (Supplementary table S1). Candidate rDNA G4s (5ETS_FW1, 5ETS_RV4) and observed genomic G4s (PITPNB) were screened for RSV binding in the biophysical assays (CD, DSC, NMR) (Supplementary table S5).

<u>Circular dichroism (CD)</u>

CD spectra were recorded at 25ºC on a JASCO 715 CD spectropolarimeter in G4s buffer conditions or FID conditions (5 µM TOPRO3). The concentration of DNA was 10 µM and RSV was added at 100 µM to obtain the new spectra. DNA and RSV were mixed and heated at 95ºC for 10 min; mixtures were left to cool down at RT overnight before the recording of the spectra. The wavelength range was 240–320 nm with 100 nm/min as scan speed. The cuvette path length was 0.1 cm, and 3-5 accumulations were averaged for each measurement. G4 oligonucleotides used in this analysis are listed in Supplementary table S1and Supplementary table S5.

Ultraviolet-visible (UV-vis) titration assay

A Varian Cary 50 UV-vis spectrophotometer was used to obtain the UV-vis absorption spectra at 25ºC. Pre-folded G4s were added at 5 µM in G4s buffer and the spectrum recorded. A concentrated solution of RSV (1 mM) was subsequently added, 1 µL each time, with a Hamilton syringe, the mixture homogenised and the UV-vis spectrum recorded after each addition. A total volume of 10 µL of RSV were added, with a final ratio of 1:20 DNA:RSV. One blank containing all reagents in the assay except for the DNA was included and subtracted from each spectrum. The cuvette path length was 0.3 cm and the wavelength range used was 235-320 nm. Experiments were conducted in triplicates. Spectra were plotted representing the differential absorbance against the recorded wavelength range.

Differential scanning calorimetry (DSC)

To explore the thermal transition from folded G4s to the unfolded state, an automated VP-Capillary DSC instrument from Microcal INC (Northhampton, MA, USA) was used to estimate the excess heat capacity as a function of temperature. DNA quadruplexes were used at 40-70 µM in G4s buffer and RSV was added using a 1:10 molar ratio. DNA and RSV were mixed and heated at 95ºC for 10 min; mixtures were left to cool down at RT overnight before the recording of the thermograms. Thermograms were analysed using Origin 7.0 software.

Nuclear magnetic resonance (NMR)

DNA oligonucleotides including PITPNB and D1 sequence (Supplementary table S5) were resuspended in $H_2O/D_2O$ 9:1 in G4s buffer. NMR spectra were acquired in a Bruker Avance spectrometer operating at 600 MHz equipped with cryoprobes, and processed with TopSpin software. Water suppression was achieved by including a WATERGATE module in the pulse sequence prior to acquisition. NMR titrations were performed by adding increasing amounts of RSV to the oligonucleotide solution at 100 µM. Different ratios (R) = [DNA] : [RSV] were considered (R= 0, 1, 2). In the first addition, DNA and RSV were mixed and heated at 95ºC for 10 min; mixtures were left to cool down at RT overnight before the recording of the NMR spectra.

In vitro transcription assay

A375 cells were seeded at a cell density of $6 \cdot 10^3$ cells/well and grown overnight. The experimental plasmids were produced and delivered by VectorBuilder GmbH (Germany). Cell transfection was carried out using 0.5 µL/well of Lipofectamine 2000 (11668030, Thermo Scientific) and 0.2 µg/well of plasmid DNA, according to the manufacturer's protocol. After a 24h incubation, transfection medium was removed and RSV treatment applied (200 µM and 256 µM, 6h). RSV treatment was performed independently and after transfection to avoid biases and perturbations of cell transfection by the drug. Luminiscence was measured for firefly and *Renilla* luciferase using the Dual-Glo(R) Luciferase Assay System (E2920, Promega) according to the manufacturer's instructions. Non-transfected cell signal was removed from their respective positive condition for treated and non-treated cells and firefly/*Renilla* luciferase ratios were calculated. Promotor activity was normalized relative to 100% of activity in control DMSO-treated cells. Experiments were performed in biological triplicates.

Statistics

Statistical analysis was performed using GraphPad Prism 9, or R software in case of next generation sequencing experiments. Statistical significance was assessed using Student's two-tailed t-test or a non-parametric alternative (Mann-Whitney test) if data were not normally distributed. In next generation sequencing data, p-adjusted values were used. For all tests, p values below 0.05 were considered significant (unless otherwise indicated) and expressed as follows: *$p < 0.05$; **$p < 0.01$, *** $p < 0.001$ and **** $p<0.0001$.

#### Reagents

All reagents, oligonucleotides and experimental conditions used in this study are listed in Supplementary table S1-S5

#### Data and code availability

Sequencing data derived from this study have been uploaded to the Sequence Read Archive (SRA) under the accession number PRJNA986468.

R codes used in bioinformatic analyses will be provided as auxiliary supplementary files. Linux code to process BLISS fastq files is available at https://github.com/garner1/BLISS.

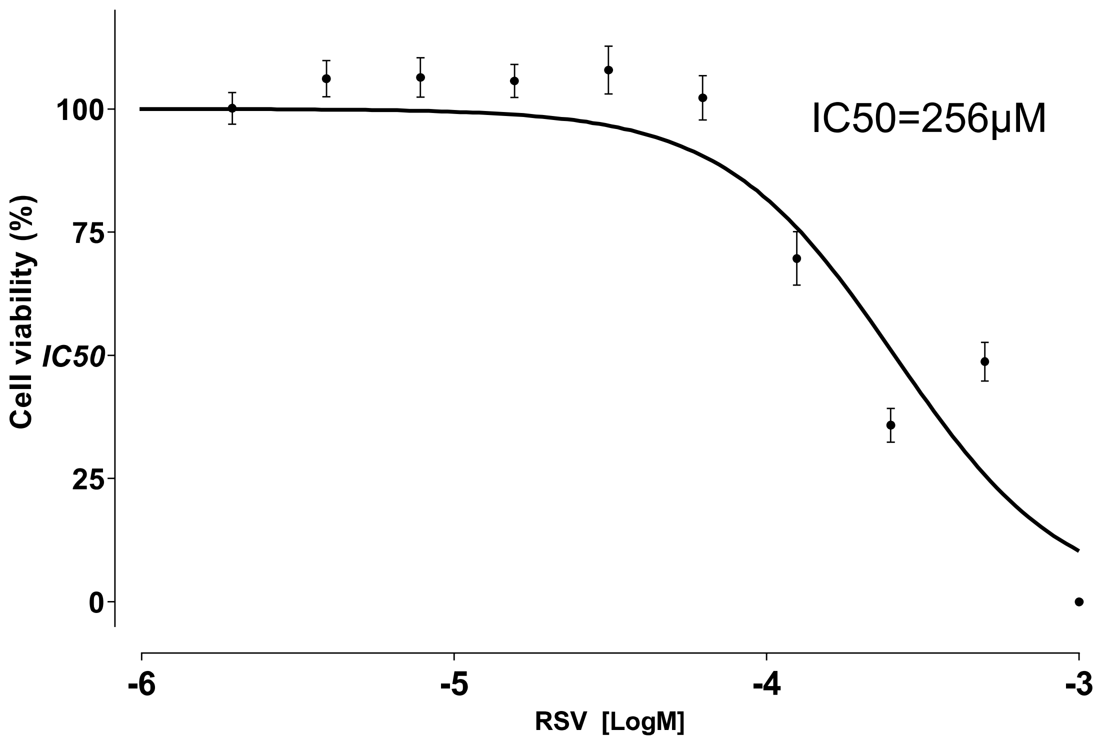


**Fig. S1.** IC50 value of RSV in the A375 cell line at 24h (n=3)

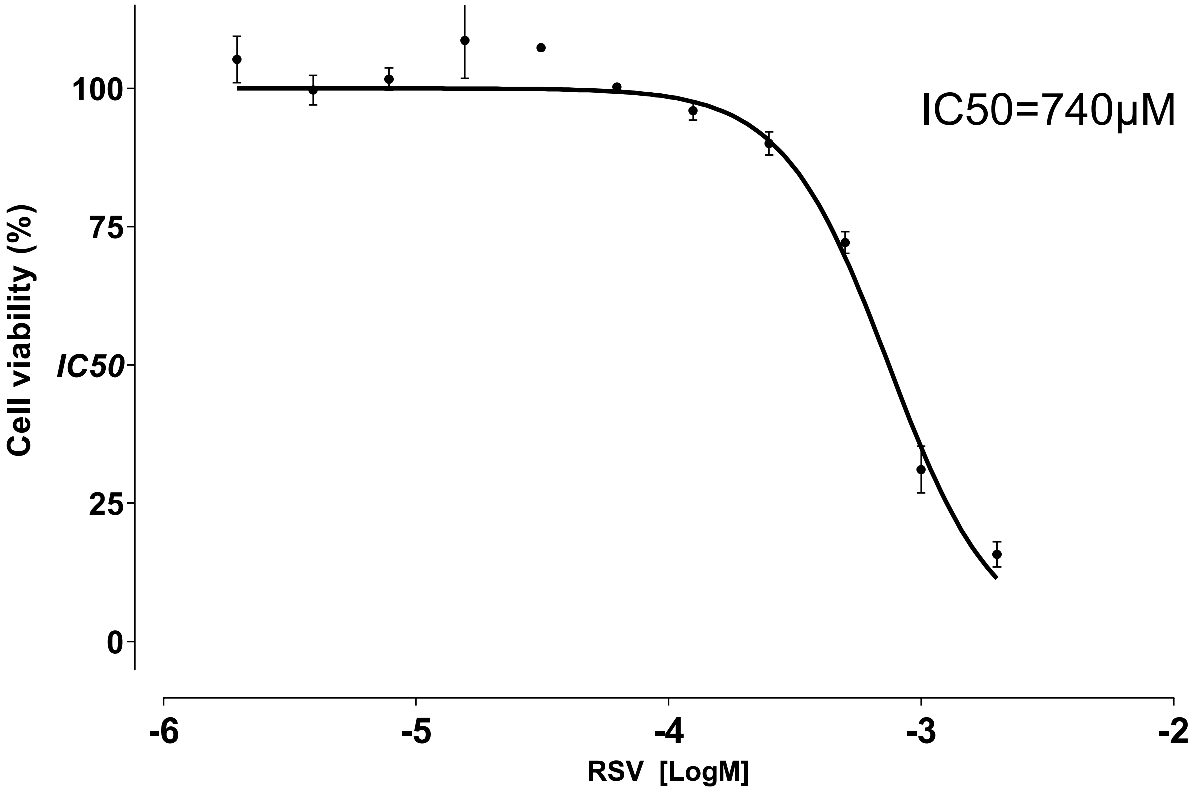


**Fig. S2.** IC50 value of RSV in the A375 cell line at 6h (n=3)

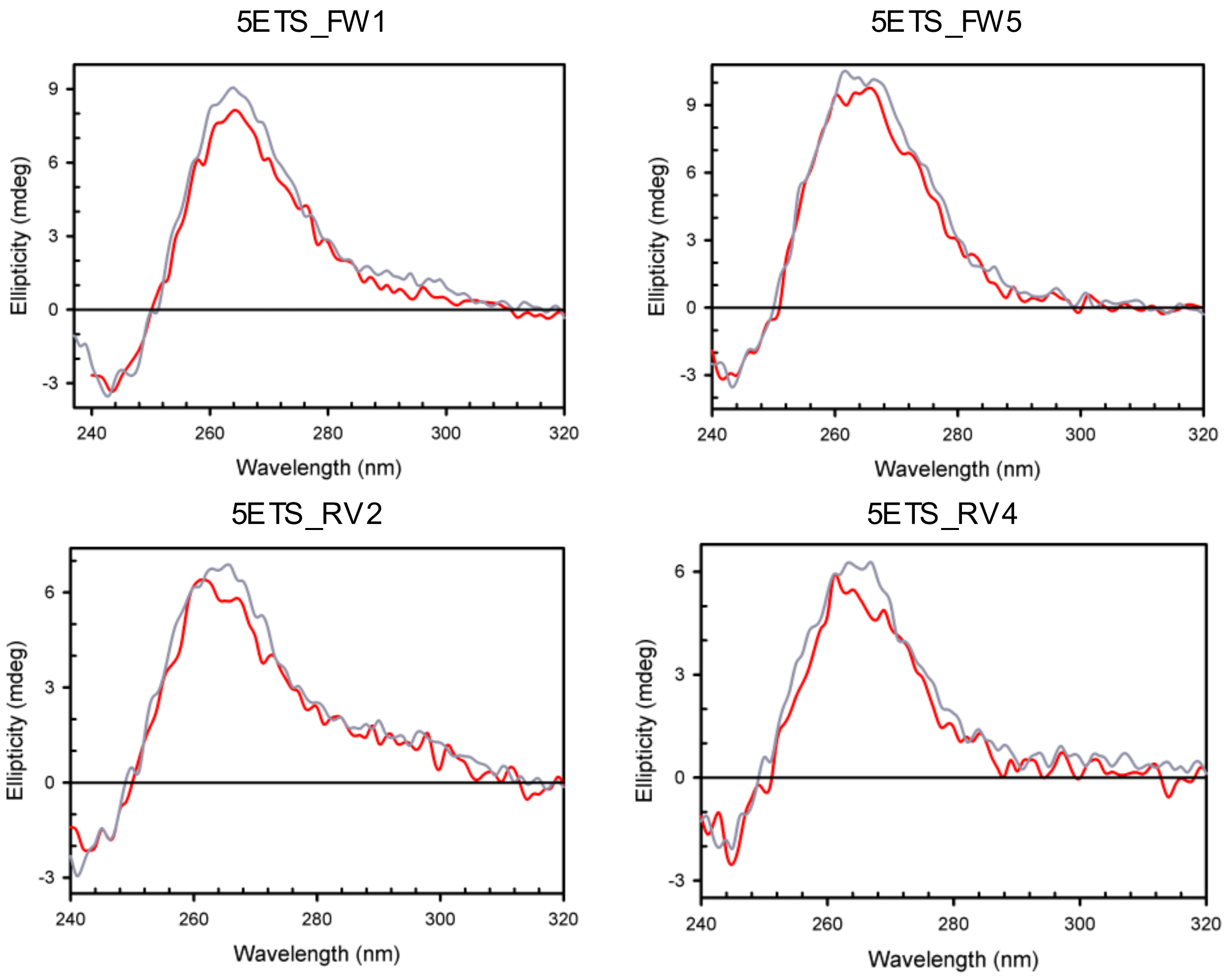


**Fig. S3.** CD spectra of 10 μM 5ETS_FW1, 5ETS_FW5, 5ETS_RV2 and 5ETS_RV4 using FID experimental conditions (5 μM TOPRO3 in G4s buffer) (red line) and standard folding conditions (G4s buffer) (grey line)

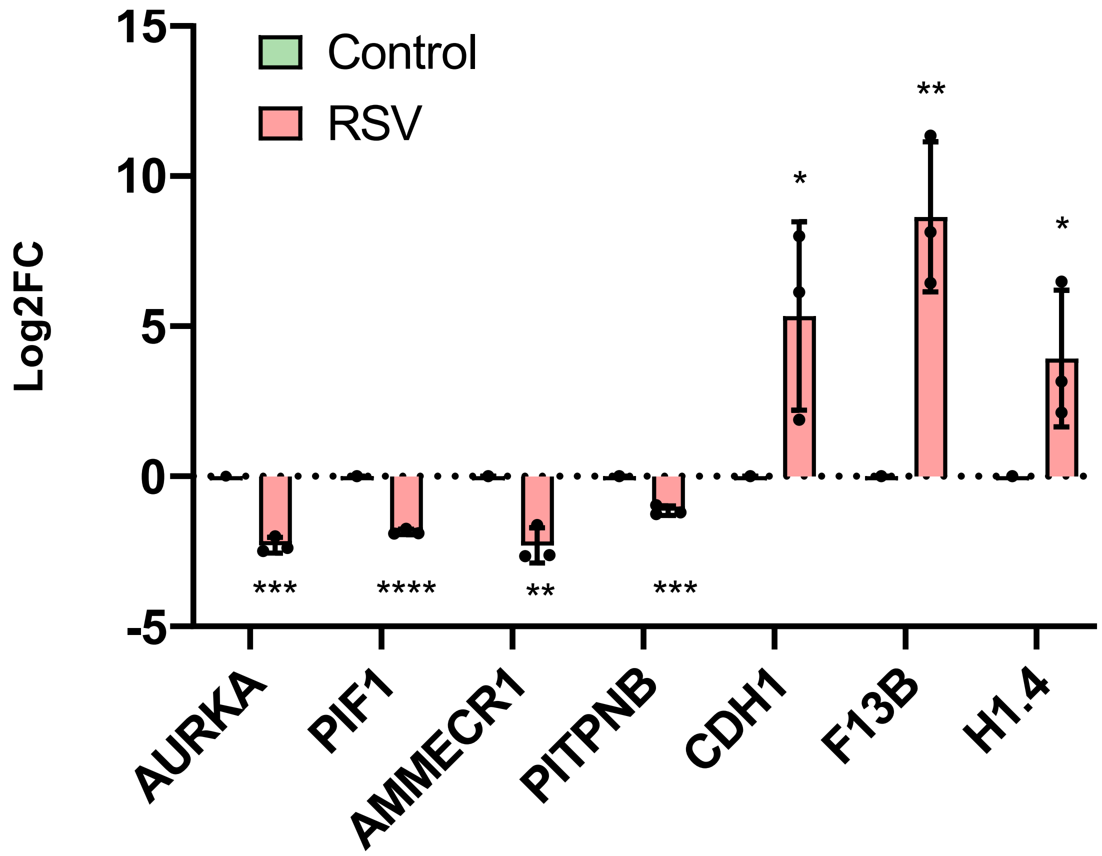


**Fig. S4.** qPCR validation of differentially expressed genes obtained from RNA-seq. Bar plot expressing mean log2FC value for seven genes in three biological replicates; error bars indicate standard deviations.

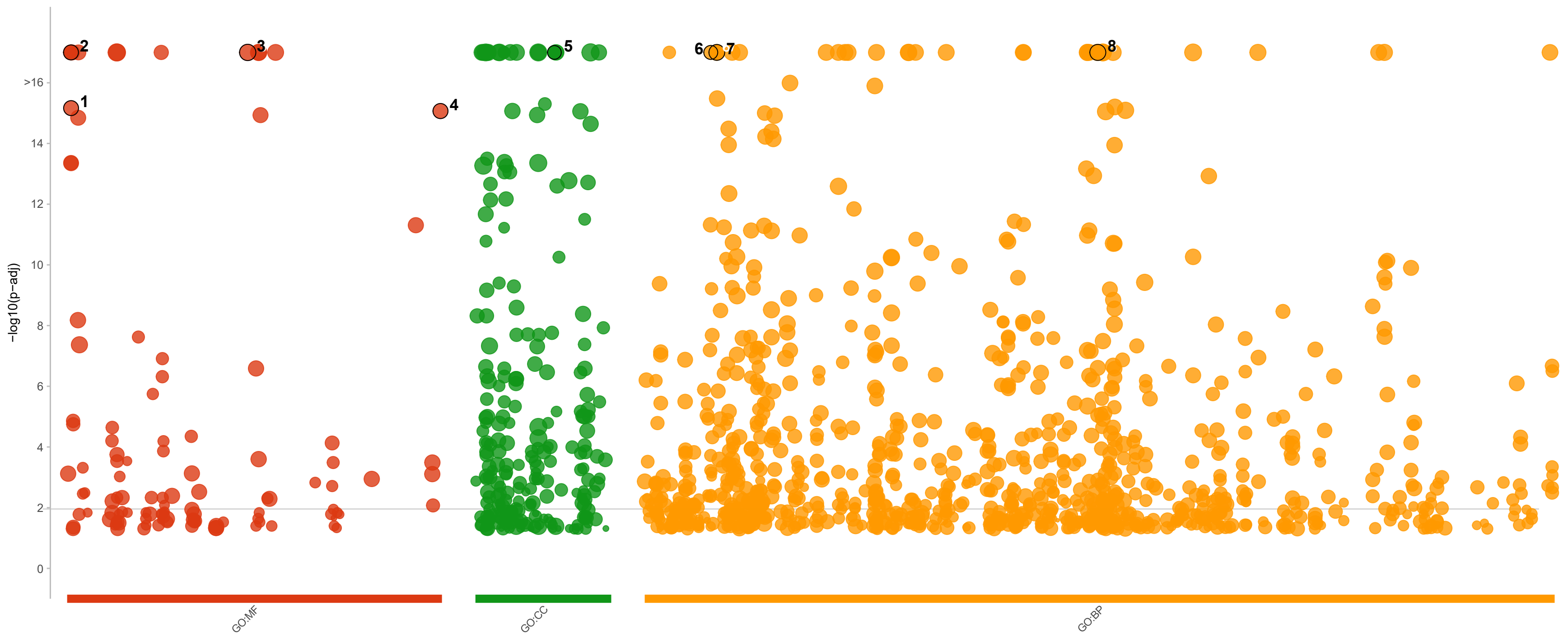


| id | source | term_id | term_name | term_size | | p_value |
|---|---|---|---|---|---|---|
| 1 | GO:MF | GO:0000981 | DNA−binding transcription factor activity, RNA polymerase II−specific | 1351 | | 6.8e−16 |
| 2 | GO:MF | GO:0000977 | RNA polymerase II transcription regulatory region sequence−specific DNA binding | 1383 | | 1.7e−17 |
| 3 | GO:MF | GO:0036094 | small molecule binding | 6357 | | 7.7e−29 |
| 4 | GO:MF | GO:1990837 | sequence−specific double−stranded DNA binding | 1541 | | 8.5e−16 |
| 5 | GO:CC | GO:0062023 | collagen−containing extracellular matrix | 425 | | 1.9e−17 |
| 6 | GO:BP | GO:0006091 | generation of precursor metabolites and energy | 515 | | 9.2e−18 |
| 7 | GO:BP | GO:0006355 | regulation of DNA−templated transcription | 3446 | | 7.2e−18 |
| 8 | GO:BP | GO:0048869 | cellular developmental process | 4367 | | 9.3e−20 |

g:Profiler (biit.cs.ut.ee/gprofiler)

**Fig. S5**. Scatter plot for functional analysis of significantly upregulated genes upon RSV treatment. Gene onthology (GO): molecular function (MF), cellular component (CC) and biological process (BP) databases were highlighted in the graph along with some significant terms of interest in relation to RSV actions.

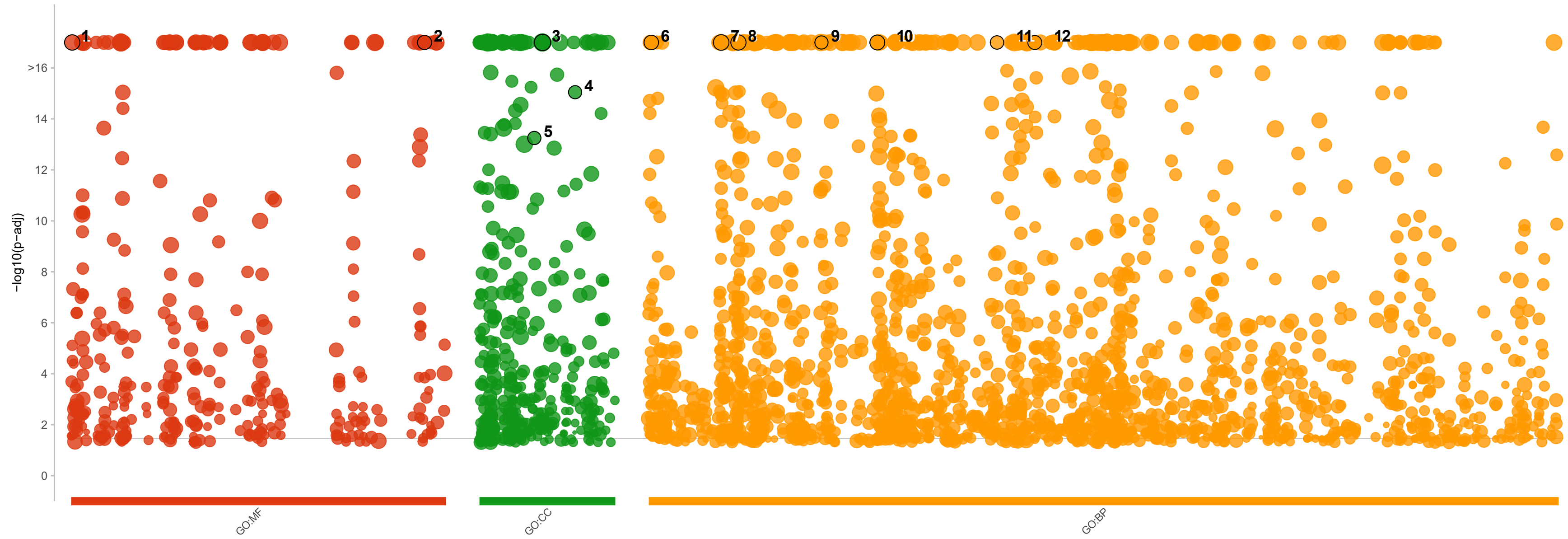


| id | source | term_id | term_name | term_size | | p_value |
|---|---|---|---|---|---|---|
| 1 | GO:MF | GO:0000166 | nucleotide binding | 2187 | | 6.0e−48 |
| 2 | GO:MF | GO:0140640 | catalytic activity, acting on a nucleic acid | 623 | | 2.3e−42 |
| 3 | GO:CC | GO:0043232 | intracellular non−membrane−bounded organelle | 6591 | | 2.3e−54 |
| 4 | GO:CC | GO:0090575 | RNA polymerase II transcription regulator complex | 258 | | 8.9e−16 |
| 5 | GO:CC | GO:0035770 | ribonucleoprotein granule | 272 | | 5.6e−14 |
| 6 | GO:BP | GO:0000278 | mitotic cell cycle | 895 | | 1.4e−83 |
| 7 | GO:BP | GO:0006366 | transcription by RNA polymerase II | 2691 | | 4.5e−47 |
| 8 | GO:BP | GO:0007049 | cell cycle | 1819 | | 3.4e−113 |
| 9 | GO:BP | GO:0016072 | rRNA metabolic process | 263 | | 5.7e−39 |
| 10 | GO:BP | GO:0022402 | cell cycle process | 1276 | | 3.8e−93 |
| 11 | GO:BP | GO:0042254 | ribosome biogenesis | 322 | | 1.9e−42 |
| 12 | GO:BP | GO:0044770 | cell cycle phase transition | 537 | | 2.2e−48 |

g:Profiler (biit.cs.ut.ee/gprofiler)

**Fig. S6.** Scatter plot for functional analysis of significantly downregulated genes upon RSV treatment. Gene onthology (GO): molecular function (MF), cellular component (CC) and biological process (BP) databases were highlighted in the graph along with some significant terms of interest in relation to RSV actions.

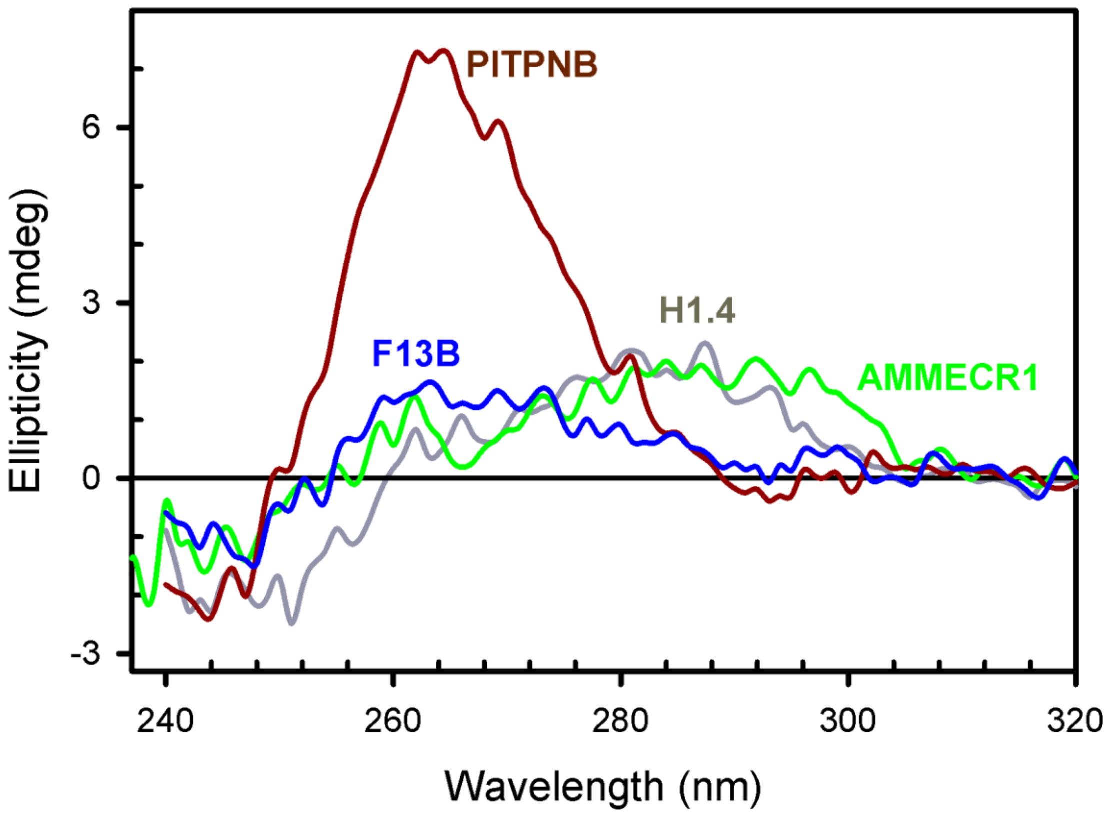

**Fig. S7.** CD spectra of genomic putative G4 forming sequences obtained from the proximity of recurrent DSBs in the promoter of DEGs.

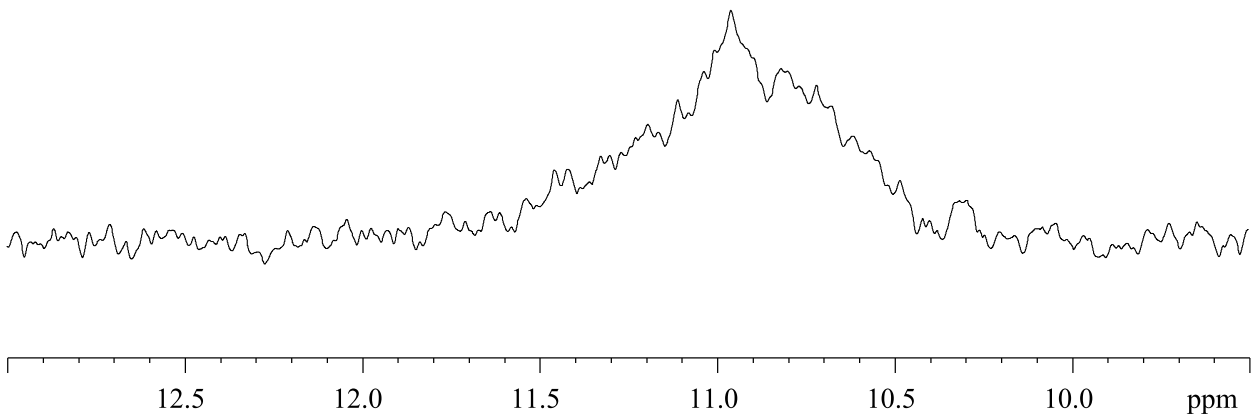


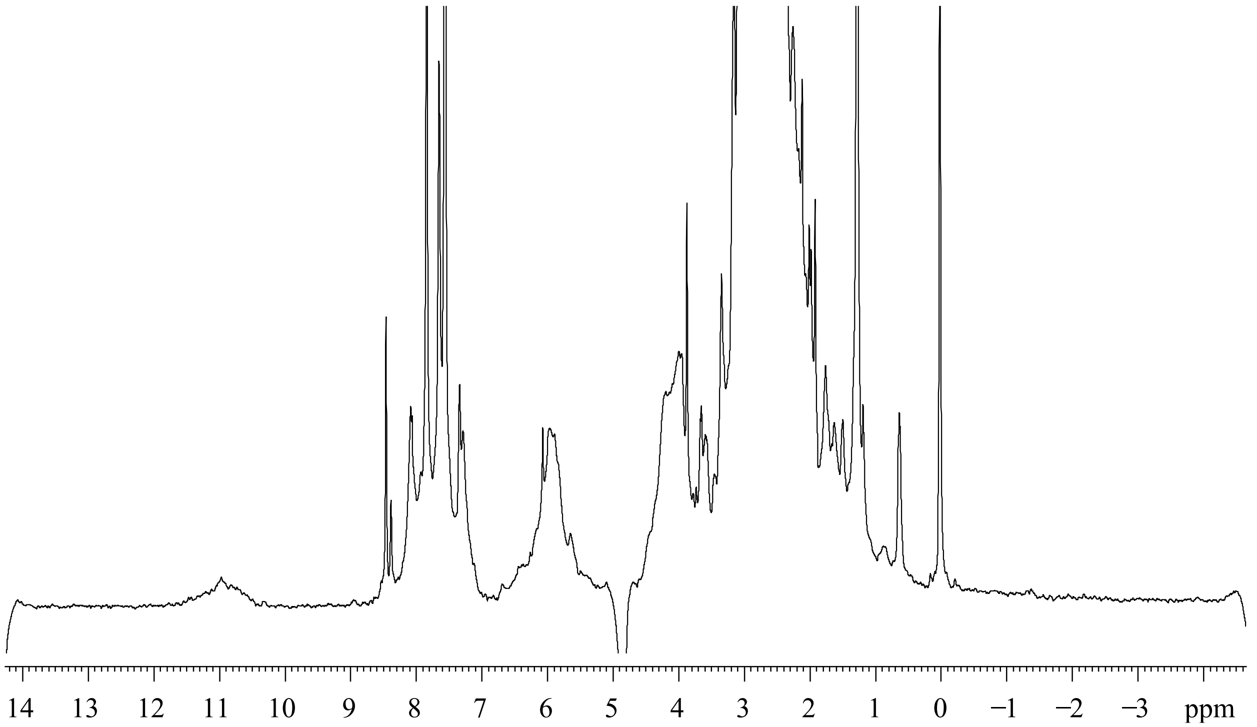


**Fig. S8.** Exchangeable proton region of the NMR spectra of PITPNB G4 with DMSO using the concentration equivalent to R DNA:RSV 1:1

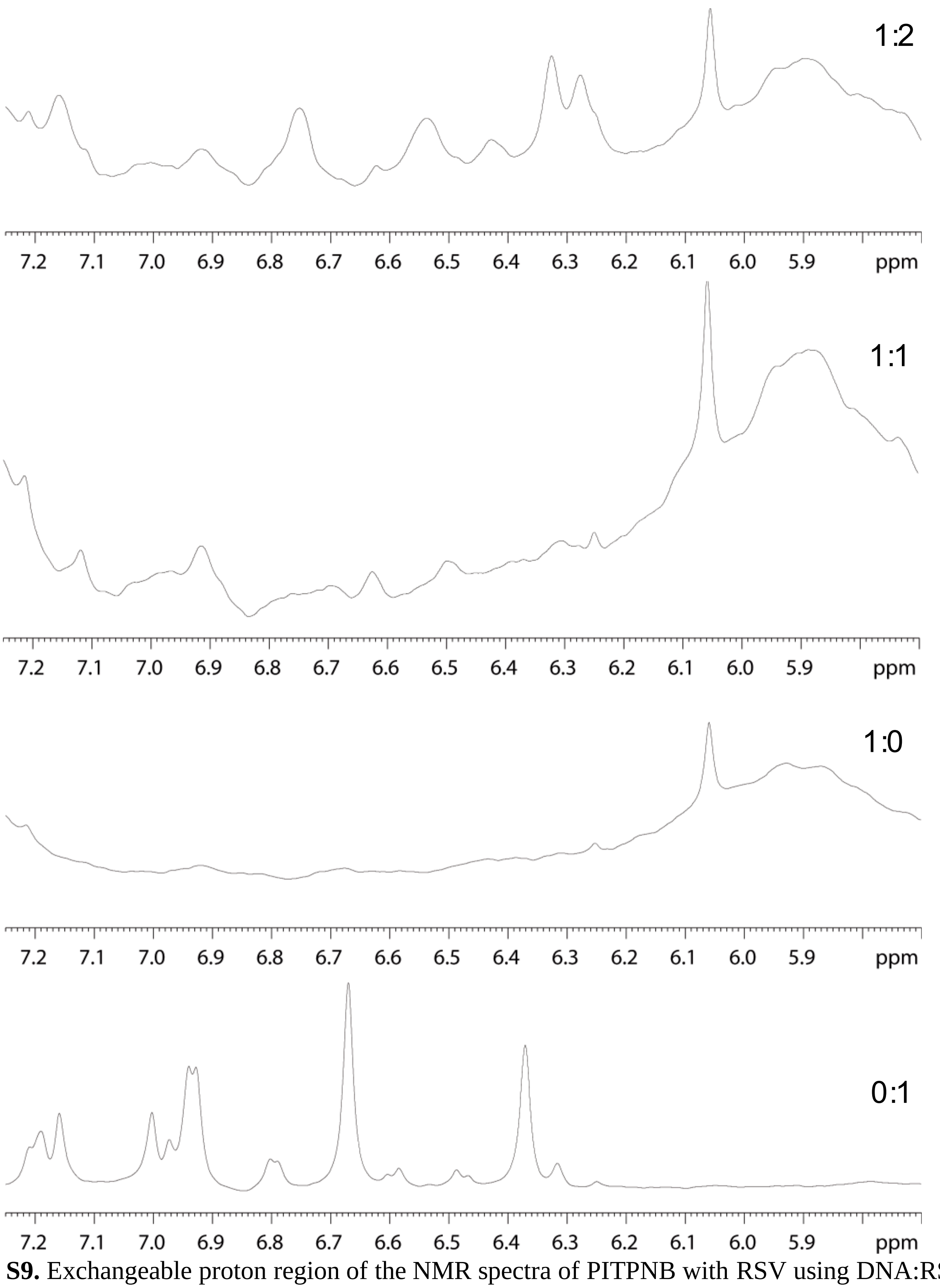


**Fig. S9.** Exchangeable proton region of the NMR spectra of PITPNB with RSV using DNA:RSV ratios 0:1, 1:0, 1:1 and 1:2.

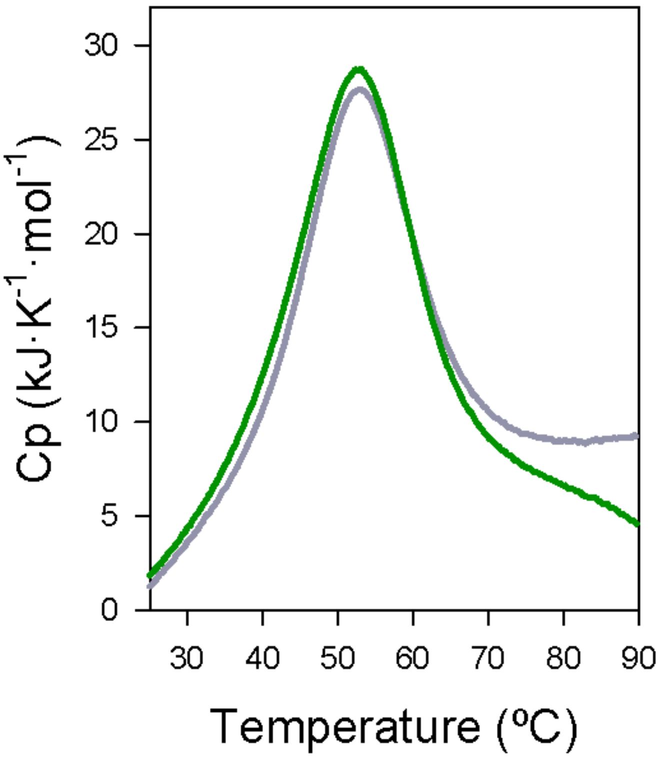


**Fig. S10.** Thermal melting profiles showing excess heat capacity as a function of temperature for D1 sequence with (green line) and without RSV (grey line)

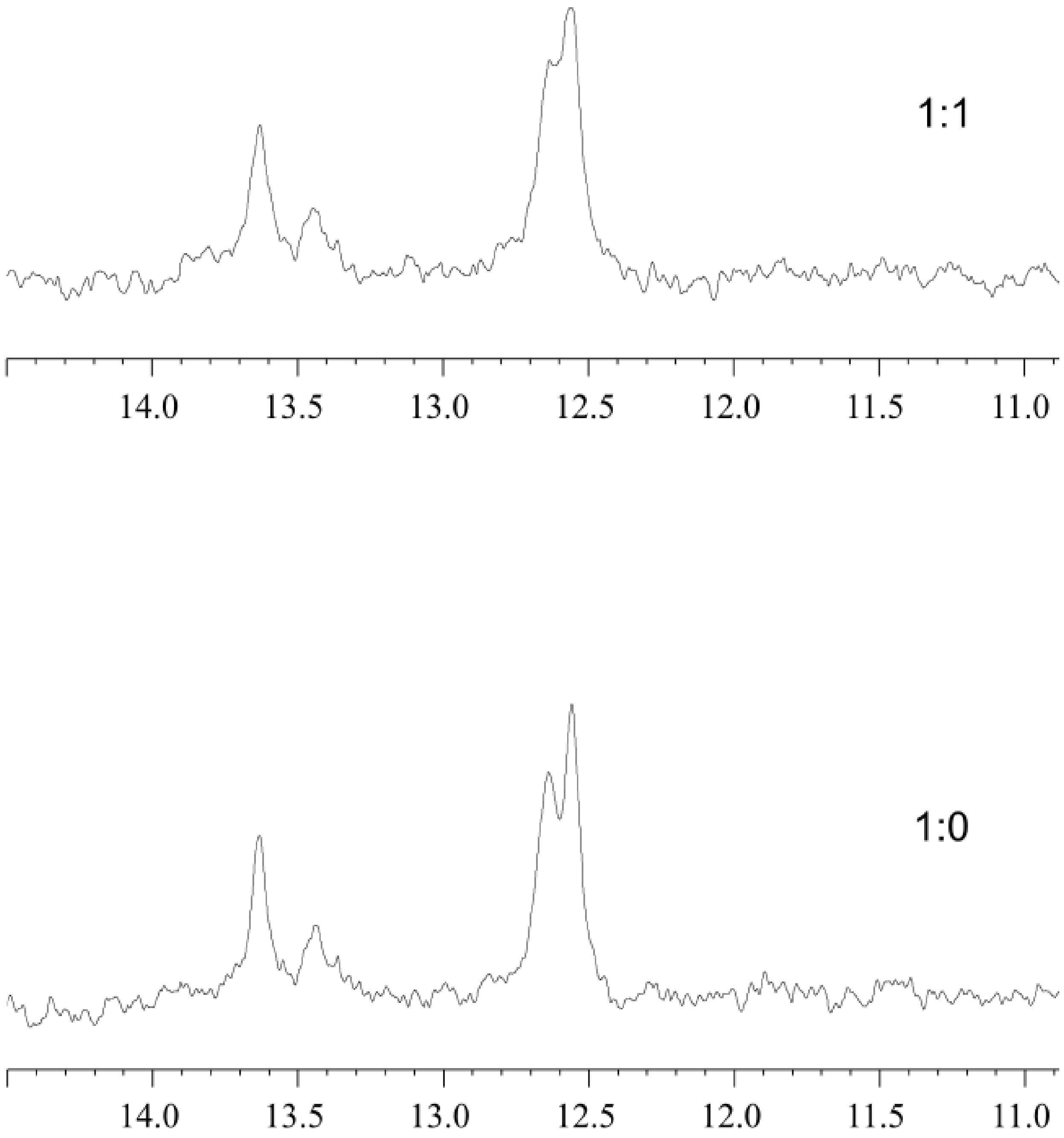


**Fig S11**. Exchangeable proton region of the NMR spectra of D1 oligonucleotide with RSV at different DNA:RSV ratios

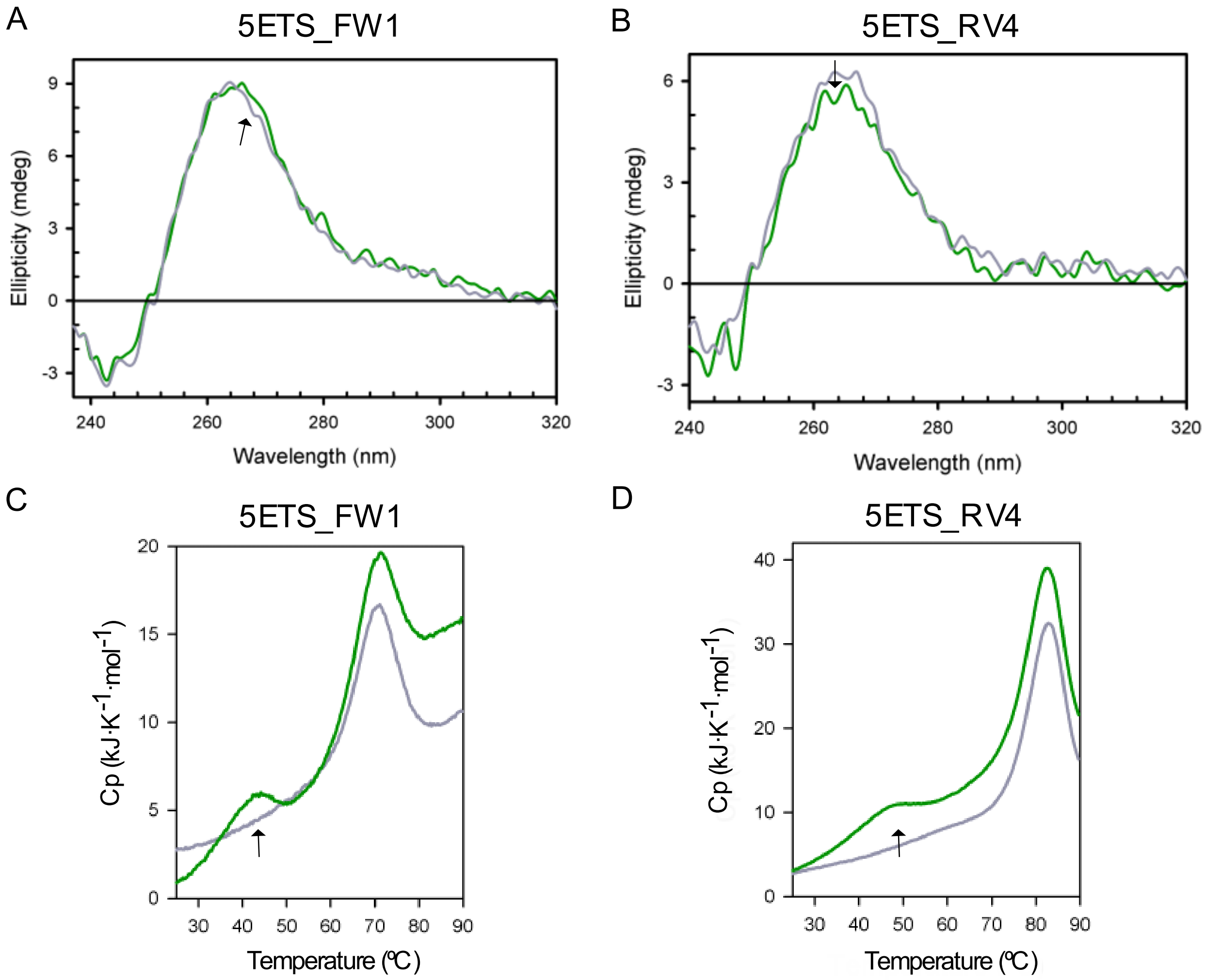


**Fig. S12.** Binding assays (CD and DSC) characterising RSV interaction with 5ETS_FW1 and 5ETS_RV4. (A) CD spectra of 10 µM 5ETS_FW1 in presence and absence of RSV 100 µM. (B) CD spectra of 10 µM 5ETS_RV4 in presence and absence of RSV 100 µM. (C) Thermal melting profiles showing excess heat capacity as a function of temperature for 5ETS_FW1 and without RSV; the arrow indicates the ligand-dependent conformational state (C) Thermal melting profiles showing excess heat capacity as a function of temperature for 5ETS_RV4 with and without RSV; the arrow indicates the ligand-dependent conformational state. Grey lines represent the control condition in absence of RSV and green lines represent test conditions with the addition of RSV.

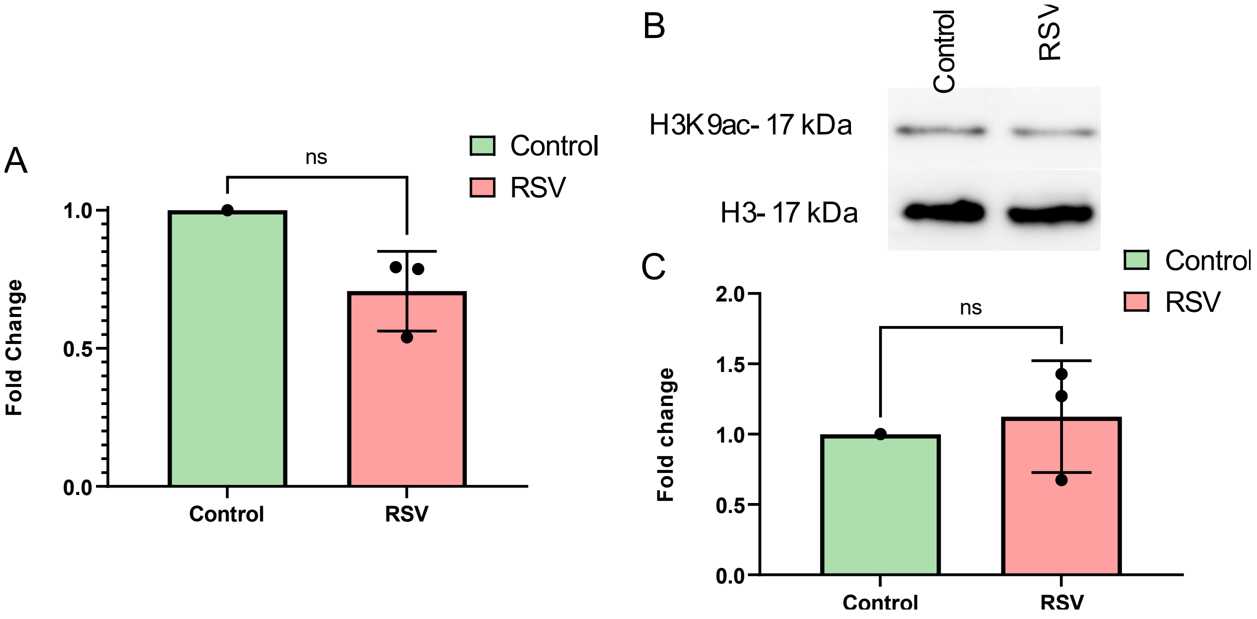


**Fig. S13.** SIRT1 analysis upon RSV exposure. (A) Bar plot representing mean expression levels (fold change) calculated in relation to actin of SIRT1 in A375 cell line treated with RSV IC50 for 6h. Error bars show standard deviations for three biological replicates (n=3). (B) Western-Blot analysis of H3K9 acetylation in A375 cell line treated with RSV IC50 for 6h; representative image of three independent experiments (n=3). (C) Bar plot showing the quantification of protein levels expressed as fold change in three independent experiments (n=3); normalization was performed in relation to total H3. Ns: non-significant.

**Table S1**. Putative G4 forming sequences screened in this study from 5'ETS rDNA region and promoters of DEGs

| Oligonucleotide | Strand | Sequence | Gscore |
|---|---|---|---|
| 5ETS_FW1 | forward | GGGTGGACGGGGGGGCCTGGTGGG | 37 |
| 5ETS_FW2 | forward | GGGTTGGGG GGGAGAAGCG AGGG | 35 |
| 5ETS_FW3 | forward | GGGGGGAGC CGCGGGGATC GCCGAGGG | 34 |
| 5ETS_FW4 | forward | GGGGTGGG GCCCGGGCCG GGG | 42 |
| 5ETS_FW5 | forward | GGGGGGCGGG TGGTTGGG | 37 |
| 5ETS_RV1 | reverse | GGGAGGGA GCGAGCGGGC GCGGG | 36 |
| 5ETS_RV2 | reverse | GGGACCGGTGGGGCCGGGGCGGGG | 39 |
| 5ETS_RV3 | reverse | GGGA GGGACCACCG GGCCGCGCTC GGG | 35 |
| 5ETS_RV4 | reverse | GGGCGGCGGGCGGGGAAGAGGG | 40 |
| 5ETS_RV5 | reverse | GG GCGAGGGCCG GGGACCGCGA GGG | 38 |
| 5ETS_RV6 | reverse | GGGT GGGAGCGCCG GGCCCGGCCC GGCGGG | 32 |
| 5ETS_RV7 | reverse | GGGT GGGAGCGCCG GGCCCGGCCC GGCGGG | 32 |
| AMMECR1 | forward | AGGCAGGGG TTTGGAAAGA GGTCCTAAGG ATGC | 20 |
| PITPNB | reverse | CAAG GGCTGGGGAG GGGAAC | 19 |
| F13B | forward | ACAGGGGCTGCCGGAGGAACTGGGAAAAG | 17 |
| H1.4 | reverse | AGCCGGAAGCAGAGGCACTGGCCGGGACTC | 17 |

**Table S2**. Resource table

| Resource | Source | Identifier |
|---|---|---|
| **Antibodies** | | |
| Actin | Sigma Aldrich | Cat #A5441 |
| BG4 | Shankar Balasubramanian's lab (Cambridge, UK) *(14)* | N/A |
| FLAG | Sigma Aldrich | Cat #F1804 |
| H3 | Abcam | Cat #ab1791 |
| H3K9ac | Abcam | Cat #ab4441 |
| Mouse Alexa Fluor 466 | Invitrogen | Cat #A-11001 |
| Mouse HRP conjugated | Promega | Cat #W4021 |
| Nucleolin | Invitrogen | Cat #39-6400 |
| Rabbit Alexa Fluor 555 | Invitrogen | Cat #A-31572 |
| Rabbit HRP conjugated | Promega | Cat #W4011 |
| **Reagents** | | |
| Bovine serum albumin | Sigma Aldrich | Cat #A7906 |
| Bradford assay | Bio-Rad | Cat #5000006 |
| DMSO | Sigma Aldrich | Cat #D2650 |
| Dual-Glo(R) Luciferase Assay System | Promega | Cat #E2920 |
| HRP substrate (Clarity Western ECL Substrate) | Bio-Rad | Cat #1705060 |
| Lipofectamin 2000 | Thermo Scientific | Cat #11668030 |
| Nitrocellulose membranes | Pall corporation | Cat #66485 |
| Paraformaldehyde | Sigma Aldrich | Cat #P6148 |
| Phenylmethylsulfonyl fluoride (PMSF) | Merck | Cat #P7626 |
| Propidium iodide | Sigma Aldrich | Cat #P4864 |
| Protease inhibitor cocktail | Merck | Cat #P8340 |
| Resarzurin | Sigma Aldrich | Cat #R7017 |
| Resveratrol | Merck | Cat #R5010 |
| RevertAid First Strand cDNA Synthesis Kit | Thermo Scientific | Cat #K1622 |
| Ribonuclease A | Qiagen | Cat #19101 |
| Sodium azide | Acros organics | Cat #447810250 |
| SYBR Green | Thermo Scientific | Cat #4309155 |
| TOPRO3 | Thermo Scientific | Cat #T3605 |
| Triton-X100 | Sigma Aldrich | Cat #T8787 |
| Trizol Reagent | Invitrogen | Cat #15596 |
| TruSeq stranded mRNA Library Prep | Illumina | Cat #20020594 |
| TURBO Dnase | Thermo Scientific | Cat #AM2238 |
| Vectashield mounting medium | Vector Laboratories | Cat #H-1200 |
| **Oligonucleotides** | | |
| Supplementary table S1, S4, S5 | Integrated DNA Techonologies (IDT) | N/A |
| **Software** | | |
| Fiji | ImageJ | https://imagej.nih.gov/ij/ |

| | | |
|---|---|---|
| FlowJo, LLC | Becton Dickinson | https://www.flowjo.com/ |
| Prism | GraphPad | https://www.graphpad.com/ |
| QGRS mapper | Ramapo College Bioinformatics | http://bioinformatics.ramapo.edu/QGRS/index.php |
| R software | R project | https://www.r-project.org/ |
| Integrative Genomics Viewer | Broad Institute, University of California | https://software.broadinstitute.org/software/igv/ |

**Table S3**. Conditions used for each antibody

| Antibody | Conditions | Experiments |
|---|---|---|
| ƔH2AX | 1:800 | Western Blot |
| ACT | 1:10000 | Western Blot |
| Anti-mouse Alexa 488 | 1:1000 | Immunofluorescence |
| Anti-mouse HRP | 1:2500 | Western Blot |
| Anti-rabbit HRP | 1:2500 | Western Blot |
| H3 | 1:50000 | Western Blot |
| H3K9ac | 1:1600 | Western Blot |
| NCL | 1:100 | Immunofluorescence |

**Table S4.** qPCR primers used in this study

| Gene | Abbreviation | Experiment | Sequence |
|---|---|---|---|
| 5'ETS | (1) | rRNA transcription | F: GTGCGTGTCAGGCGTTCT; R: GGGAGAGGAGCAGACGAG |
| ITS1-5.8S | (2) | rRNA transcription | F: CCCGTTTGCTGTCTCGTCT; R: GCAAGTGCGTTCGAAGTGTC |
| 5.8S-ITS2 | (3) | rRNA transcription | F: CTTAGCGGTGGATCACTCGG; R: GTCTGCGCTTAGGGGGAC |
| ITS2_25S | (4) | rRNA transcription | F: CTCTCTCCCGTCGCCTCT; R: CCTGTTCACTCGCCGTTACT |
| 25S-3'ETS | (5) | rRNA transcription | F: GGAACCTGGCGCTAAACCAT; R: GGAGGAAGACGAACGGAAGG |
| Aurora kinase A | AURKA | RNAseq validation | F: GGTTCCTCCGTCCCTGA, R: GCAGTTTTCTTTAGATCGGTCC |
| PIF-1 | PIF-1 | RNAseq validation | F: CCCTGGATTGTGTGGAGATT; R: ACTCCAGACTGAGGCTCCTG |
| AMMECR1 | AMMECR1 | RNAseq validation | F: TGCATTCAGGACTCAGGGAG; R: CACTGAGCAGAAAAGCCGTG |
| Phosphatidylinositol transfer protein beta gene | PITPNB | RNAseq validation | F: ATTCCGTGTGGTTTTGCCAT; R: TTCCTTCTCCACCACCAGTC |
| Cadherin 1 | CDH1 | RNAseq validation | F: CCCGGGACAACGTTTATTAC; R: GCTGGCTCAAGTCAAAGTCC |
| Coagulation factor 13 B chain | F13B | RNAseq validation | F: AGATGGAATGTGGACTACAC;R:TTGCATTCTAAGTATAGATCCAG |
| Histone H1.4 | H1.4 | RNAseq validation | F: AGCGAAGGCCAAAGCAGTTA; R: CTGAAGTCCCTCACTGCCTG |
| Sirtuin 1 | SIRT1 | SIRT1 analysis | F: GCAGATTAGTAGGCGGCTTG; R: TCTGGCATGTCCCACTATCA *(36)* |

**Table S5.** G4-contaning oligonucleotides used in the binding assays

| Gene | Abbreviation (manuscript) | Strand | Sequence |
|---|---|---|---|
| 5ETS rDNA | 5ETS_FW1 | forward | GGGTGGACGGGGGGGCCTGGTGGG |
| 5ETS rDNA | 5ETS_FW5 | forward | GGGGGGCGGG TGGTTGGG |
| 5ETS rDNA | 5ETS_RV2 | reverse | GGGACCGGTGGGGCCGGGGCGGGG |
| 5ETS rDNA | 5ETS_RV4 | reverse | GGGCGGCGGGCGGGGAAGAGGG |
| PITPNB | PITPNB | reverse | CAAG GGCTGGGGAG GGGAAC |
| DNA Duplex | D1 | NA | AGCTTAGGAGTGACATGAGGTCACTTGGGTCACGGCCTGC |
