## Supplementary file S1 for "Resveratrol targets G-quadruplexes to exert its pharmacological effects"

**Supplementary file 1. Differentially expressed genes (DEGs)**

| external_gene_name | logFC | FDR |
|---|---|---|
| TSPAN6 | 0.246875464 | 0.016610949 |
| SCYL3 | 0.774950889 | 6.70E-07 |
| CFH | 2.131903226 | 5.45E-05 |
| GCLC | -0.673192066 | 2.95E-07 |
| NFYA | -1.049650453 | 2.04E-09 |
| STPG1 | 1.022554033 | 1.76E-06 |
| NIPAL3 | 0.325339759 | 9.39E-05 |
| LAS1L | -0.20075996 | 0.002757218 |
| ENPP4 | -0.653939882 | 0.000245905 |
| SEMA3F | 0.490400325 | 0.004201374 |
| ANKIB1 | -0.899141713 | 5.20E-09 |
| CYP51A1 | 0.731391687 | 5.77E-08 |
| KRIT1 | -0.800981335 | 1.46E-07 |
| RAD52 | 1.299282924 | 8.02E-08 |
| MYH16 | 1.923097104 | 0.000793028 |
| BAD | 0.376535371 | 0.001289215 |
| CD99 | 0.358795659 | 0.031878456 |
| MAD1L1 | -0.571988345 | 0.00023563 |
| SNX11 | -1.868339042 | 4.30E-09 |
| M6PR | -0.496388417 | 6.39E-07 |
| KLHL13 | -1.114631053 | 0.000163817 |
| ICA1 | 2.307917995 | 3.17E-07 |
| DBNDD1 | 0.585250298 | 0.000351536 |
| ALS2 | -0.131931943 | 0.04662204 |
| CFLAR | -0.703497343 | 5.71E-05 |
| NDUFAF7 | -0.652282795 | 8.61E-06 |
| MTMR7 | -1.548271184 | 0.000330087 |
| SLC7A2 | 0.301604523 | 0.000276252 |
| ARF5 | 0.298628778 | 0.000844142 |
| PLXND1 | -0.374166273 | 8.05E-05 |
| AK2 | -0.271760226 | 4.95E-05 |
| CD38 | 5.188396468 | 0.000249629 |
| FKBP4 | 0.195258889 | 0.001105763 |
| KDM1A | -0.20859584 | 0.001453215 |
| RBM6 | -0.469817748 | 4.05E-06 |
| CAMKK1 | 0.423258265 | 0.048792816 |
| RECQL | 0.44450665 | 1.16E-06 |
| VPS50 | -0.947916404 | 8.05E-06 |
| HSPB6 | 1.433679751 | 0.006232466 |
| ARHGAP33 | -0.733080773 | 0.003285735 |
| PDK4 | 1.803828897 | 2.52E-09 |
| ZMYND10 | 1.865500251 | 3.65E-05 |
| SLC25A13 | -0.480800395 | 1.00E-05 |
| ST7 | -0.918541958 | 5.28E-06 |
| CDC27 | -1.777802368 | 1.25E-11 |
| HCCS | -0.784455336 | 2.08E-07 |

| | | |
|---|---|---|
| DVL2 | -0.629878053 | 5.86E-07 |
| UPF1 | -0.554024884 | 1.48E-07 |
| SLC25A5 | 0.379586744 | 8.96E-07 |
| MCUB | -0.448345883 | 0.00267262 |
| HOXA11 | -1.802792045 | 0.001360272 |
| DHX33 | -1.389794829 | 7.43E-11 |
| MEOX1 | 4.546303878 | 0.000528002 |
| LIG3 | -0.744598808 | 1.70E-07 |
| RPAP3 | -0.620386446 | 3.45E-06 |
| REXO5 | 1.400012918 | 1.33E-06 |
| CIAPIN1 | -0.322539772 | 0.000423298 |
| SPPL2B | 0.366970906 | 0.001597131 |
| COPZ2 | 1.984370885 | 4.97E-07 |
| PRKAR2B | -1.122988322 | 4.62E-09 |
| MSL3 | 0.289106681 | 0.000939893 |
| CREBBP | -0.906723687 | 1.35E-08 |
| GCFC2 | -0.415960402 | 0.000432261 |
| WDR54 | 0.620789555 | 2.71E-06 |
| KMT2E | -0.977083622 | 2.28E-08 |
| RHBDD2 | 0.409430489 | 5.15E-05 |
| SOX8 | 3.264029237 | 4.78E-07 |
| IBTK | 0.16468355 | 0.022985904 |
| MYCBP2 | -0.697245696 | 1.24E-07 |
| ITGAL | 4.455806736 | 0.000673117 |
| PDK2 | 0.924407703 | 3.10E-06 |
| ITGA3 | 0.232952402 | 0.00014444 |
| ITGA2B | 5.082612714 | 1.66E-08 |
| OSBPL7 | 0.433550072 | 0.00344099 |
| MAP3K14 | -0.401022426 | 1.40E-05 |
| TMEM132A | 0.197742037 | 0.021554363 |
| TAC1 | 2.930772983 | 1.10E-09 |
| ZNF263 | -0.452706734 | 7.49E-06 |
| CX3CL1 | 2.189944108 | 0.001020981 |
| SPATA20 | 0.35081933 | 0.000167472 |
| TNFRSF12A | 0.292709607 | 6.64E-05 |
| DLX6 | -1.583427055 | 0.005841783 |
| MAP3K9 | -0.60860923 | 2.60E-05 |
| RALA | -0.431558034 | 4.52E-06 |
| KDM7A | -0.945804816 | 3.17E-06 |
| ETV1 | -0.301440409 | 0.000220637 |
| ALDH3B1 | 0.78062088 | 0.000261415 |
| TTC22 | 8.714679541 | 4.21E-08 |
| PHTF2 | -1.233692019 | 6.63E-09 |
| FARP2 | -1.383830679 | 1.80E-10 |
| GGCT | -0.346054921 | 3.32E-05 |
| DBF4 | -0.859622741 | 6.57E-08 |
| TBXA2R | -1.941162699 | 3.90E-05 |
| IFRD1 | 0.557322599 | 9.30E-07 |

| | | |
|---|---|---|
| COX10 | -1.565142872 | 3.03E-09 |
| GTF2IRD1 | 0.736444433 | 0.000808377 |
| PAF1 | -0.317803805 | 0.000114325 |
| ELAC2 | -0.250893184 | 0.001008138 |
| ARSD | 0.930944582 | 1.36E-06 |
| ADIPOR2 | -0.165370517 | 0.005357646 |
| CDKL3 | 3.078895278 | 5.77E-09 |
| CCDC124 | 0.360303697 | 0.000243626 |
| PAFAH1B1 | -0.999608194 | 1.19E-09 |
| DNAH9 | 0.633569592 | 0.00241682 |
| KIAA0100 | -0.123602985 | 0.011859324 |
| AC005747.1 | 0.106509362 | 0.048901378 |
| TRAPPC6A | 1.545725469 | 0.006913562 |
| RPUSD1 | -0.161208421 | 0.021042349 |
| RHBDF1 | 0.461047592 | 8.50E-05 |
| CACNA2D2 | 0.917978965 | 0.023976325 |
| BAIAP3 | 1.903835957 | 4.08E-05 |
| PIGQ | 0.283076358 | 0.004304187 |
| CRAMP1 | 0.374398214 | 4.90E-05 |
| TEAD3 | -0.581619392 | 0.003748435 |
| SELE | 5.092298736 | 8.87E-05 |
| DNAJC11 | -0.579123927 | 4.61E-07 |
| MYLIP | 0.397004978 | 0.000967012 |
| E2F2 | -2.523900022 | 1.11E-09 |
| PSMB1 | 0.166489845 | 0.007364924 |
| CDKL5 | 0.797371479 | 1.62E-06 |
| CDK11A | -0.206978287 | 0.011261842 |
| NADK | -0.367952345 | 0.000101486 |
| TFAP2B | 0.573156628 | 5.15E-05 |
| DLEC1 | 0.485822334 | 0.001674842 |
| CYTH3 | -0.247808524 | 0.001039911 |
| ADAM22 | -1.031988533 | 2.34E-07 |
| SYPL1 | -0.513107529 | 1.01E-06 |
| CYB561 | 0.160380627 | 0.028950678 |
| SPAG9 | -0.154027835 | 0.00634722 |
| CELSR3 | 1.580945878 | 1.18E-10 |
| SS18L2 | -0.241893457 | 0.007720173 |
| MPND | 0.690094717 | 0.018952161 |
| MGST1 | 0.164970427 | 0.029812283 |
| CRY1 | -0.613591585 | 1.22E-06 |
| NFIX | -0.418947082 | 0.000447332 |
| ST3GAL1 | 0.227195645 | 0.001188007 |
| PKD1 | 0.598724825 | 1.01E-07 |
| MAPK8IP2 | 2.842121051 | 4.44E-06 |
| MED24 | -0.804338587 | 4.44E-08 |
| RHOBTB2 | -0.869073586 | 2.52E-08 |
| HEATR5B | 0.554783568 | 9.32E-06 |
| SEC62 | -0.733989301 | 3.02E-08 |

| | | |
|---|---|---|
| RPS20 | 0.345097743 | 1.09E-06 |
| UBE3C | -1.206993656 | 2.20E-10 |
| REV3L | -1.124921195 | 8.46E-09 |
| TENM1 | -1.651532446 | 2.05E-07 |
| MASP2 | 2.948674507 | 1.88E-08 |
| FAM76A | 2.234491613 | 7.98E-10 |
| TRAF3IP3 | 1.093902225 | 0.000278084 |
| POMT2 | -0.809416149 | 3.99E-06 |
| VTA1 | -0.356409933 | 3.08E-05 |
| BAZ1B | -1.1587773 | 1.77E-10 |
| RANBP9 | -0.772789943 | 2.57E-07 |
| SPRTN | -0.631211677 | 4.49E-07 |
| EEF1AKNMT | -1.169497555 | 1.32E-09 |
| ZNF207 | -1.598051118 | 6.29E-12 |
| UQCRC1 | 0.236639392 | 0.000455775 |
| CD9 | 0.361986856 | 0.000284147 |
| HHATL | 7.234104704 | 8.03E-06 |
| NCAPD2 | -0.10688313 | 0.011108236 |
| IFFO1 | -0.730487583 | 9.21E-06 |
| GIPR | 2.383142302 | 3.38E-05 |
| PHF7 | 2.232415127 | 2.34E-08 |
| NISCH | -0.843000578 | 2.32E-08 |
| STAB1 | 4.142673507 | 4.36E-06 |
| FUZ | 0.70115198 | 6.02E-06 |
| SLC6A13 | 8.649695571 | 5.49E-08 |
| ZNF200 | -0.33736174 | 0.000390443 |
| CD4 | 1.83221051 | 0.009259653 |
| LRRC23 | 1.46269577 | 2.14E-06 |
| BTK | 8.210165997 | 3.61E-07 |
| ELOA | -1.633418007 | 1.35E-11 |
| LYPLA2 | -0.684310195 | 1.72E-07 |
| CLCN6 | 0.363448684 | 0.00150618 |
| MRC2 | 0.512037441 | 7.21E-06 |
| NME1-NME2 | 0.340109509 | 1.93E-05 |
| TSPAN9 | -0.739492079 | 2.82E-05 |
| BTBD7 | -0.849011351 | 7.45E-08 |
| APBA3 | 0.753677311 | 2.16E-05 |
| ABHD5 | -1.177865104 | 1.62E-07 |
| AKAP8L | -0.413589669 | 3.86E-05 |
| UTP18 | -0.713152443 | 1.33E-06 |
| RNF216 | -0.536091892 | 1.22E-06 |
| TTC19 | -0.493773878 | 7.88E-06 |
| PTBP1 | -0.307625298 | 6.41E-06 |
| DPF1 | 0.371830395 | 0.017685886 |
| LARS2 | -0.581082383 | 4.09E-06 |
| PIK3C2A | -0.435422646 | 6.42E-06 |
| PLAUR | -1.234774378 | 1.71E-09 |
| ANLN | -0.995661824 | 4.77E-10 |

| | | |
|---|---|---|
| WIZ | -0.232295212 | 0.00298888 |
| QPCTL | 0.966010731 | 1.67E-06 |
| PPP5C | -0.428594563 | 1.31E-05 |
| CEP68 | 0.382197627 | 0.000544571 |
| MAP4K3 | -0.684220719 | 1.28E-05 |
| ZBTB32 | 2.736046935 | 7.29E-06 |
| GABRA3 | -0.50831931 | 0.00278419 |
| BRCA1 | -0.866774654 | 7.58E-09 |
| ERCC1 | -0.510271769 | 6.94E-07 |
| CD22 | 3.074027311 | 2.55E-10 |
| SEMA3B | 0.972463481 | 2.81E-09 |
| MBTPS2 | -0.308100393 | 9.00E-05 |
| PRICKLE3 | 1.123263163 | 1.81E-05 |
| EXTL3 | -1.31716184 | 1.69E-09 |
| NR1H4 | 4.50585024 | 0.000625696 |
| ELOVL5 | -0.424930013 | 1.07E-05 |
| CALCOCO1 | 1.357838822 | 8.01E-09 |
| UBR7 | -0.530102621 | 5.43E-07 |
| MAP4K5 | -1.354362356 | 5.53E-09 |
| EHD3 | -0.917858979 | 1.11E-08 |
| MAN2B2 | -0.523952578 | 0.001145009 |
| SLC25A39 | 0.161063075 | 0.023009668 |
| MVP | 0.609104256 | 4.43E-06 |
| NUB1 | -0.468297265 | 1.11E-05 |
| RWDD2A | 1.635354046 | 2.41E-07 |
| CLK1 | 1.096874633 | 1.88E-08 |
| POLR3B | -0.22782953 | 0.031865999 |
| RNF14 | -0.840452973 | 2.35E-08 |
| DNASE1L1 | 1.160546284 | 1.21E-07 |
| DDX11 | -0.874767332 | 5.63E-08 |
| HEBP1 | 0.82166267 | 3.09E-05 |
| GPRC5A | -0.706162266 | 7.16E-10 |
| MAMLD1 | -1.304170722 | 5.71E-06 |
| CD6 | 7.403498125 | 4.53E-06 |
| TACC3 | -1.058068836 | 2.00E-09 |
| UFL1 | 0.537501375 | 7.06E-05 |
| POLA2 | -1.136527634 | 7.83E-08 |
| ZC3H3 | -0.437169697 | 0.01101015 |
| CAPN1 | 0.337396098 | 0.000484204 |
| MDH1 | 0.497779577 | 4.71E-06 |
| SLC30A9 | -0.320001811 | 0.000184998 |
| MTMR11 | 2.10178564 | 6.98E-09 |
| COX15 | -0.175069341 | 0.009438099 |
| ZMYND11 | -0.995068361 | 9.91E-09 |
| WAS | 4.794196183 | 3.48E-07 |
| MATR3 | 0.111510247 | 0.004022285 |
| NPC1L1 | 4.351760224 | 7.86E-09 |
| XYLT2 | -1.017716513 | 4.28E-08 |

| | | |
|---|---|---|
| NUDCD3 | -0.699434507 | 3.97E-08 |
| ISL1 | -0.570989151 | 0.000771077 |
| CHDH | 4.863982114 | 0.005349376 |
| GLT8D1 | -0.351145975 | 4.27E-05 |
| ATP2C1 | -0.301841599 | 0.00040064 |
| RALBP1 | -0.452382629 | 1.35E-05 |
| RUFY3 | -0.781868254 | 0.000184021 |
| WWTR1 | -0.343776316 | 7.20E-07 |
| AGPS | -0.510163612 | 9.18E-07 |
| CXorf56 | -0.781372742 | 2.69E-06 |
| TTC27 | -0.92200439 | 7.06E-05 |
| ZNF582 | -0.850244001 | 0.00020105 |
| VSIG2 | -0.887502978 | 0.001326173 |
| PHLDB1 | -0.404846424 | 1.28E-05 |
| PRDM11 | 0.782236438 | 8.92E-05 |
| SNAI2 | 3.667943964 | 4.12E-10 |
| CD74 | 0.38907223 | 3.68E-07 |
| ZRANB1 | -0.717923379 | 1.22E-07 |
| NCDN | -0.228617306 | 0.000734619 |
| ZFP64 | -1.049788379 | 1.81E-06 |
| MNAT1 | -0.433505374 | 0.00590844 |
| SAMD4A | -2.728328809 | 7.81E-09 |
| RUNX3 | -1.526436541 | 3.70E-10 |
| MRE11 | -0.385067945 | 0.000114472 |
| PLEKHB1 | 0.40995602 | 0.022130823 |
| SERPINB1 | 0.286126905 | 0.04556514 |
| CYP3A43 | 4.307090806 | 0.001033285 |
| SPAST | -1.493808052 | 1.43E-09 |
| NRXN3 | -1.157823564 | 5.19E-05 |
| AQR | -0.624419746 | 2.29E-07 |
| RTF2 | -0.154344225 | 0.014187948 |
| SLC45A4 | -0.187730403 | 0.040204218 |
| RNF10 | -0.5037281 | 1.81E-06 |
| ZDHHC6 | -1.406084221 | 1.19E-09 |
| GRAMD1B | 1.224295728 | 1.58E-10 |
| NDUFS1 | -0.19358239 | 0.005374656 |
| RB1CC1 | -0.325899241 | 4.13E-05 |
| ALAS1 | 0.746126603 | 1.02E-07 |
| BIRC3 | 0.917882993 | 2.19E-07 |
| AKAP11 | -0.170573231 | 0.002950087 |
| SNAPC1 | -1.170392443 | 5.02E-06 |
| STRAP | -0.185714123 | 0.000629208 |
| ABCC2 | 0.624156568 | 1.66E-05 |
| DEF6 | 1.713012698 | 0.003999008 |
| PLEKHO1 | -0.613499589 | 0.000306485 |
| GCLM | -1.312564528 | 2.13E-09 |
| UBR2 | -0.578575675 | 4.59E-05 |
| EHD2 | 0.301634598 | 0.000255305 |

| | | |
|---|---|---|
| DEPDC1 | -1.792914259 | 3.70E-10 |
| CCDC28A | 0.472662931 | 0.001038492 |
| HSF2 | -0.860762906 | 6.78E-05 |
| PHF20 | 0.239018352 | 0.000630288 |
| HSD17B6 | 1.66415503 | 0.000110351 |
| NR1H3 | 1.637004037 | 1.18E-07 |
| NCAPH2 | -0.497112476 | 2.03E-05 |
| TOMM34 | 0.262822834 | 0.000218512 |
| KPNA6 | -0.348592126 | 1.05E-05 |
| VIM | 0.445548257 | 1.35E-06 |
| RTEL1-TNFRSF6B | -0.624085797 | 1.29E-05 |
| FAS | 1.077520069 | 1.66E-08 |
| CD44 | -0.209022755 | 2.21E-05 |
| KCNG1 | 1.454979703 | 3.72E-10 |
| AGPAT4 | 0.820125078 | 0.000386263 |
| SLAMF7 | -1.120986197 | 0.037462433 |
| MIPEP | 0.636552755 | 0.001324077 |
| PRKCH | -0.351671202 | 0.021668746 |
| IFNGR1 | 0.674428027 | 8.60E-06 |
| B4GALT7 | 0.296348848 | 0.009830053 |
| SH2D2A | 1.806118344 | 0.001098943 |
| VRK2 | -0.962160508 | 7.22E-06 |
| VEZT | 0.436184004 | 6.94E-06 |
| POU2F2 | 3.844726335 | 5.92E-10 |
| BRD9 | -0.766726766 | 7.45E-08 |
| SNX1 | 0.534234183 | 3.29E-07 |
| ARNTL2 | -1.887085511 | 7.73E-09 |
| BCLAF1 | -1.39313406 | 4.12E-11 |
| SLC39A9 | -0.653268146 | 5.06E-08 |
| TFB1M | 0.374721652 | 0.006399518 |
| RABEP1 | -0.434581555 | 3.70E-06 |
| HMGB3 | -0.825723835 | 1.35E-08 |
| NUP160 | -0.225686588 | 0.000463668 |
| BAK1 | -0.371690441 | 0.000184315 |
| GRN | 0.642247709 | 1.60E-07 |
| FAM13B | -1.832178884 | 1.08E-10 |
| ARHGAP31 | -0.887023855 | 7.66E-05 |
| CENPQ | -0.387163919 | 0.002002245 |
| SARS1 | 0.25605095 | 0.001577851 |
| RANBP3 | -0.889777322 | 6.13E-08 |
| ARID4A | 0.709938503 | 3.86E-06 |
| PNPLA6 | 0.373917859 | 5.13E-05 |
| IFT88 | 0.626661757 | 0.00121831 |
| ZCCHC8 | -1.31815061 | 3.42E-09 |
| ABCF2 | -0.493828052 | 8.55E-07 |
| CHPF2 | -0.552054193 | 9.96E-06 |
| FUT8 | -0.338767123 | 0.002299442 |
| UBA6 | -0.588282302 | 9.75E-07 |

| | | |
|---|---|---|
| PIAS1 | -1.422179088 | 7.10E-10 |
| SLC4A7 | -0.646763865 | 1.09E-07 |
| APBA2 | -0.727325045 | 1.81E-05 |
| MAP2K3 | 0.175476828 | 0.003749271 |
| TMSB10 | 0.170402629 | 0.000422405 |
| ASTE1 | -0.361376011 | 0.005860795 |
| SH3YL1 | 0.509587634 | 0.002229588 |
| FAM136A | -0.465074064 | 1.62E-06 |
| VCL | -0.801530637 | 1.22E-08 |
| DEPDC1B | -1.264577059 | 2.00E-09 |
| DAPK2 | 1.271318784 | 0.007327022 |
| NSMAF | -1.191280706 | 1.08E-07 |
| ADSS2 | -1.120414124 | 3.35E-09 |
| TIMP2 | -0.179674193 | 0.003677818 |
| RFC1 | -0.603492629 | 2.68E-07 |
| TBC1D23 | -0.408230795 | 4.78E-05 |
| CUL3 | -1.088106365 | 1.91E-08 |
| MYOM2 | 4.620117788 | 0.001429091 |
| CYP46A1 | 4.23406747 | 4.81E-06 |
| AC118549.1 | -0.721659998 | 1.99E-08 |
| NSUN2 | -0.81094997 | 4.23E-08 |
| MRI1 | -0.202974716 | 0.004909585 |
| HOXC8 | -1.701696316 | 6.32E-08 |
| AGA | 1.121995228 | 1.82E-07 |
| PI4K2B | -0.323113539 | 0.002380581 |
| BOD1L1 | -0.274525682 | 7.15E-05 |
| MAT2B | 0.318154331 | 4.30E-05 |
| EDC4 | -0.636177594 | 1.38E-07 |
| TRIO | -1.316773882 | 8.90E-10 |
| CLEC16A | -1.331430843 | 2.71E-09 |
| MSR1 | -0.600275579 | 0.005265573 |
| CDH1 | 7.771008004 | 1.58E-06 |
| MTREX | -0.497586333 | 2.19E-06 |
| DNAH5 | 1.801105263 | 2.17E-05 |
| ZFYVE16 | -0.218305928 | 0.002435011 |
| RIPOR1 | -0.449628075 | 6.54E-06 |
| RAI14 | -0.535121654 | 1.12E-06 |
| SOX30 | 7.709867965 | 1.76E-08 |
| PNKP | 0.216764803 | 0.036705231 |
| BEST2 | 6.822795279 | 2.73E-05 |
| PHLPP2 | -0.508811082 | 5.02E-06 |
| SPDL1 | -1.077834184 | 4.82E-09 |
| CTNS | -0.404935268 | 0.00124692 |
| RTN4R | 1.056450937 | 0.0001499 |
| PHF23 | -1.043335687 | 1.43E-08 |
| CDH10 | 2.48623849 | 2.96E-05 |
| INPP4A | -0.479919218 | 4.46E-05 |
| RAB27B | -0.905416293 | 0.000151896 |

| | | |
|---|---|---|
| MYO16 | 2.678092098 | 4.58E-07 |
| LSG1 | -0.58540692 | 9.27E-07 |
| PARP3 | 0.550553929 | 0.000493436 |
| TNC | 0.793730872 | 2.05E-10 |
| THAP3 | -0.334687787 | 0.000194999 |
| RIPOR3 | 5.355790773 | 2.69E-05 |
| AIFM2 | -0.248566222 | 0.04035249 |
| AL049834.1 | 1.300279914 | 0.000161856 |
| MED17 | -0.754043794 | 2.21E-07 |
| RETSAT | 1.018778215 | 4.99E-09 |
| CAPG | 0.378285824 | 6.39E-05 |
| AP2S1 | 0.299554408 | 8.33E-05 |
| TG | 8.431921979 | 1.19E-07 |
| DCUN1D1 | -0.880749421 | 1.71E-08 |
| JADE2 | -0.281785467 | 0.001396316 |
| ZIC2 | -1.764925951 | 5.16E-08 |
| TRIT1 | -0.190442496 | 0.036681849 |
| CUL7 | 0.285635924 | 0.003643565 |
| CTNNA1 | -0.247831187 | 1.69E-05 |
| PHKA2 | -0.888193424 | 8.51E-06 |
| CNTLN | -0.917649722 | 2.87E-07 |
| HSPA5 | 0.83943527 | 2.08E-11 |
| GEMIN8 | -0.376427057 | 0.038555122 |
| OFD1 | -0.498686534 | 0.000140805 |
| GPM6B | 1.099905521 | 0.003372637 |
| MAGEC2 | -0.100895022 | 0.035169746 |
| WDR37 | -0.840664051 | 1.67E-06 |
| YTHDC2 | -0.403148304 | 4.08E-05 |
| CTPS2 | -0.370870291 | 0.007819331 |
| POLR2B | -0.501386978 | 6.77E-07 |
| FAM214A | 1.493497622 | 1.83E-10 |
| ARAP2 | -1.193032099 | 4.84E-07 |
| TPR | 0.221017607 | 0.000179716 |
| KIAA0556 | 0.603033407 | 1.64E-05 |
| XK | -1.188879766 | 0.000495983 |
| C12orf4 | -0.717429657 | 2.80E-06 |
| WWC3 | -0.550702612 | 1.70E-05 |
| FAM184B | -2.408643976 | 2.32E-07 |
| GOPC | -0.263237158 | 0.000635875 |
| USP28 | -0.465947623 | 1.81E-05 |
| HDAC9 | 1.556743253 | 1.45E-08 |
| TSPAN17 | -0.293622999 | 0.002474542 |
| NOP16 | -1.199843594 | 5.23E-09 |
| RRM2B | -0.231772971 | 0.000691293 |
| ZNF800 | -0.134562928 | 0.030132947 |
| SNX29 | -0.849886411 | 8.25E-08 |
| MRPS10 | -0.579707986 | 2.23E-05 |
| RSF1 | -0.447483486 | 5.64E-06 |

| | | |
|---|---|---|
| VPS13D | -0.522860059 | 5.02E-06 |
| FAM120A | -0.430987997 | 1.20E-06 |
| R3HDM1 | -0.180737174 | 0.012295595 |
| ERCC8 | -0.361529468 | 0.010685185 |
| H6PD | 0.282993306 | 0.000873961 |
| VAMP3 | -0.133929988 | 0.0116576 |
| PER3 | 1.254979976 | 4.76E-05 |
| UTS2 | 1.315713817 | 0.001744072 |
| TNFRSF9 | 4.771127444 | 1.79E-08 |
| EPN3 | 3.914693528 | 0.002726178 |
| ELN | 4.865357194 | 0.000189092 |
| RFC2 | -0.288417517 | 0.005657935 |
| ARID1B | -0.979878309 | 8.10E-09 |
| CLPTM1L | -0.277565913 | 0.000696586 |
| NEDD4L | -0.749234285 | 4.13E-07 |
| FOXP3 | 2.101549718 | 0.005696355 |
| PPP1R3F | 1.885073873 | 1.69E-06 |
| JKAMP | -0.385799236 | 0.000270689 |
| DKK3 | 0.938562088 | 0.000233086 |
| NFE2L3 | -0.134844105 | 0.025730911 |
| LIMA1 | -1.140844616 | 1.99E-10 |
| LETMD1 | 0.508048641 | 5.47E-06 |
| SLC4A8 | -1.00928545 | 2.57E-05 |
| PTGER3 | 5.088113147 | 1.92E-05 |
| BCAR1 | -0.817267356 | 6.40E-08 |
| FAM160A2 | 0.341350851 | 0.000942889 |
| HERPUD1 | 1.680579504 | 3.55E-11 |
| RAD51 | -1.147393489 | 1.97E-06 |
| POLQ | -0.559098187 | 2.88E-06 |
| PIK3CB | -1.879252189 | 1.73E-10 |
| CYBA | 0.524610759 | 0.00077926 |
| MPHOSPH9 | -1.032375814 | 6.84E-09 |
| SIKE1 | 0.184461954 | 0.003291243 |
| RRP12 | -0.173722431 | 0.003759416 |
| FNIP2 | -1.186666735 | 1.43E-07 |
| MSMO1 | 3.045957589 | 9.00E-13 |
| TTC17 | -0.24603058 | 0.00053815 |
| ALX4 | -1.449958041 | 0.001977229 |
| FOXN3 | -0.370564332 | 0.000124177 |
| AKR7A2 | 0.323975648 | 0.000302107 |
| MRTO4 | -0.702103383 | 5.71E-08 |
| NNAT | -2.169379108 | 0.002561928 |
| USE1 | 0.384334345 | 0.000849816 |
| MCF2L2 | -0.513605517 | 0.019885001 |
| NRIP2 | -1.733010109 | 0.000119011 |
| AP5M1 | -0.266305848 | 0.004565815 |
| ANAPC4 | -0.430154426 | 0.000139789 |
| TRAPPC3 | -0.347716332 | 9.90E-05 |

| | | |
|---|---|---|
| THRAP3 | -0.474215749 | 1.96E-06 |
| PHPT1 | 0.556664743 | 8.77E-05 |
| ARID4B | -0.510170513 | 1.91E-06 |
| OPN3 | 0.570594763 | 0.000377675 |
| SDCCAG8 | 1.427651778 | 3.30E-06 |
| PTPRN | 3.072309457 | 1.15E-09 |
| HHAT | -1.405710017 | 1.87E-08 |
| KIF1B | -0.49624126 | 4.55E-07 |
| FOXC1 | -0.488817505 | 7.60E-05 |
| SYNE2 | -0.279395793 | 0.012793279 |
| PLEKHH1 | -0.79413611 | 4.68E-06 |
| ATP9A | -0.411836291 | 5.94E-05 |
| FAM168A | -0.355482534 | 1.07E-05 |
| RELT | -1.228461381 | 7.71E-09 |
| NOP58 | -0.685284471 | 9.53E-08 |
| SZRD1 | -0.988624378 | 1.17E-09 |
| CUL1 | -0.772427731 | 4.50E-07 |
| FAM114A2 | 0.562209795 | 1.92E-05 |
| CYFIP2 | 0.479931715 | 0.001809805 |
| TAB2 | -1.006706883 | 2.55E-09 |
| EIF2AK2 | -0.680999635 | 3.76E-08 |
| KMT2C | -0.158764018 | 0.005266242 |
| MCOLN3 | 0.44335844 | 0.001113554 |
| PUM2 | -1.249926836 | 1.43E-10 |
| MRPL43 | -0.473791898 | 1.26E-05 |
| ITIH4 | 2.875569388 | 8.72E-08 |
| ITIH1 | 5.094330671 | 8.92E-05 |
| ZFR | -0.190760702 | 0.000660993 |
| ZNF280C | 1.345770156 | 2.06E-07 |
| TRAF1 | 0.562874019 | 0.012871273 |
| RC3H2 | -0.842376545 | 4.31E-09 |
| IL17RB | -0.521726429 | 0.036888193 |
| TRAF3IP2 | -0.47589407 | 0.002060088 |
| DCBLD2 | -0.410326114 | 5.89E-08 |
| SOAT1 | -0.187741587 | 0.002668306 |
| MSH4 | 3.734125839 | 3.62E-07 |
| GDI2 | -0.3089447 | 2.73E-05 |
| PRDM1 | -0.809840934 | 0.02366404 |
| ATG5 | -1.07669378 | 1.62E-07 |
| PITHD1 | -0.225746684 | 0.002465671 |
| MTA3 | -0.373126318 | 0.011451317 |
| USP13 | -1.021597968 | 5.37E-08 |
| ATP11B | -0.753567114 | 6.10E-08 |
| CDK14 | -1.118036428 | 1.24E-08 |
| SEC61A1 | -0.463158669 | 1.79E-07 |
| PPP1R12A | -0.797183837 | 7.96E-09 |
| CAMK2B | 4.248104474 | 1.24E-05 |
| CROCC | -0.515073838 | 0.00056081 |

| | | |
|---|---|---|
| POLR3E | -0.81599515 | 2.71E-08 |
| ZC3H11A | -0.825101673 | 4.69E-09 |
| RIOK2 | 0.583173228 | 1.05E-06 |
| NDC1 | -0.698120767 | 3.88E-08 |
| DGKG | -1.735627562 | 8.33E-07 |
| UNKL | -0.356497497 | 0.000548413 |
| PARP12 | -0.987041505 | 7.13E-07 |
| ALDH18A1 | -0.157437638 | 0.02156346 |
| TARBP1 | -0.497066152 | 1.85E-05 |
| GATB | 0.426125991 | 0.0068284 |
| MXD1 | 1.627636815 | 7.79E-11 |
| CDK17 | -0.656520414 | 1.13E-05 |
| SLC2A3 | 2.764367445 | 1.44E-08 |
| PSD | 0.61579349 | 0.039310082 |
| CTDP1 | -0.882984601 | 0.000108143 |
| YBX3 | 0.307256293 | 2.88E-05 |
| STYK1 | -1.512681896 | 0.008089109 |
| WNK1 | -0.267940681 | 1.92E-05 |
| CCAR1 | -0.176635886 | 0.007340523 |
| OGFR | -0.452976997 | 5.91E-05 |
| PIGV | 1.01020038 | 9.79E-07 |
| SNRNP40 | -0.221494971 | 0.015746426 |
| QSER1 | -1.552482665 | 1.69E-11 |
| BCAT1 | -0.522145562 | 2.38E-07 |
| HDAC7 | -0.880975326 | 8.67E-08 |
| LZTS1 | -0.840350178 | 6.56E-06 |
| SPAG4 | 1.43985248 | 9.70E-05 |
| NCKAP1 | -0.18281799 | 0.006440292 |
| MRPS35 | -0.571144161 | 4.22E-07 |
| SFSWAP | -0.202983293 | 0.005709017 |
| TNK2 | -0.564995861 | 5.13E-05 |
| CDH3 | 2.577396248 | 2.45E-05 |
| GPBP1 | -0.552494927 | 3.80E-07 |
| LTK | 5.274313179 | 0.018820769 |
| WAPL | -0.672238997 | 4.89E-08 |
| VMP1 | -0.102130426 | 0.034711013 |
| APPBP2 | -0.413633645 | 0.00010198 |
| SEZ6 | 6.095546923 | 5.91E-06 |
| EIF4B | 0.107651606 | 0.014731288 |
| SLC6A16 | 3.764385406 | 6.38E-06 |
| SPHK2 | -0.616757603 | 1.44E-05 |
| RPL18 | 0.330850061 | 2.10E-06 |
| CA11 | 0.83579356 | 0.007500291 |
| ISOC2 | 0.467405915 | 0.003859133 |
| U2AF2 | -0.302053816 | 1.61E-05 |
| MED29 | -0.493107428 | 2.23E-06 |
| AHRR | -0.580039359 | 2.16E-06 |
| MTMR1 | -1.100224726 | 3.20E-09 |

| | | |
|---|---|---|
| GPC1 | 0.434765505 | 1.39E-05 |
| ADCK1 | 2.175530265 | 1.87E-07 |
| HAGH | 0.474476004 | 0.000978325 |
| RNF4 | -0.701926515 | 1.78E-07 |
| CASP8 | -0.401048489 | 9.29E-05 |
| LIMCH1 | -0.427092171 | 1.97E-05 |
| INTS13 | -0.758208889 | 9.76E-08 |
| TM7SF3 | -0.132088823 | 0.0326506 |
| DLX3 | 2.507959353 | 2.37E-05 |
| SPA17 | 1.004520443 | 0.000110921 |
| ST3GAL6 | -0.764701815 | 0.000709013 |
| ATP2C2 | 3.072110089 | 4.15E-08 |
| NGFR | 0.451065858 | 9.53E-06 |
| CDON | -1.170454821 | 1.47E-08 |
| TAF2 | -0.79758266 | 5.45E-08 |
| HIPK2 | -1.46036809 | 1.42E-11 |
| TNPO3 | -0.773186821 | 5.36E-08 |
| BORCS8-MEF2B | 1.465584229 | 1.68E-05 |
| TMEM161A | 0.210794627 | 0.0203069 |
| LPAR2 | 1.413771987 | 8.35E-06 |
| SUGP2 | -1.048787001 | 2.00E-09 |
| SLC12A2 | -1.100496766 | 3.07E-09 |
| SNX24 | -0.963413954 | 5.05E-08 |
| ABCA7 | 1.095723606 | 1.10E-05 |
| DDX20 | -0.655467498 | 1.93E-07 |
| BTBD1 | -0.274830545 | 0.000161661 |
| FAR2 | -0.652438307 | 0.000304625 |
| SBNO2 | -0.895797482 | 2.32E-07 |
| PMS1 | 0.391646743 | 0.003538647 |
| TAF11 | -1.181953678 | 2.21E-08 |
| ANKS1A | -0.631946818 | 4.61E-06 |
| ZNF76 | -0.374882998 | 0.017378586 |
| SLC9A3R2 | 0.239848455 | 0.001049213 |
| UHRF1BP1 | -0.88689971 | 5.25E-08 |
| GNAI3 | -0.289451141 | 6.39E-05 |
| IPO5 | -0.084803141 | 0.049301344 |
| OAT | -0.384885158 | 5.50E-06 |
| WDR3 | -0.856642263 | 3.89E-09 |
| PKN2 | -0.813444668 | 1.06E-08 |
| WDR18 | 0.214577185 | 0.017246424 |
| TRAM2 | -1.095361457 | 3.68E-10 |
| NTN1 | -0.529897015 | 0.015019872 |
| MCM10 | -1.529654402 | 4.22E-10 |
| DGKA | 0.653896171 | 4.21E-05 |
| ERBB3 | 0.66859868 | 1.93E-07 |
| ANKRD44 | -1.75376976 | 5.76E-06 |
| KARS1 | 0.118321765 | 0.0204557 |
| ADAT1 | -0.602890741 | 6.44E-07 |

| | | |
|---|---|---|
| SPEN | -0.884229128 | 1.42E-09 |
| ZC3H15 | -0.805880319 | 2.06E-08 |
| MAP2K4 | -1.398199586 | 9.77E-10 |
| PACC1 | -0.905242368 | 1.01E-05 |
| SLK | -0.416342797 | 3.71E-06 |
| GSTO2 | 5.405367103 | 1.36E-12 |
| SEC61A2 | 0.753317044 | 2.94E-05 |
| PRKCQ | -0.415986575 | 0.043018391 |
| ASB1 | -1.603035027 | 7.08E-09 |
| CDK13 | -1.83536728 | 2.72E-10 |
| SLC9A7 | -0.682712124 | 2.52E-06 |
| FOXJ2 | -0.224982621 | 0.002839737 |
| YBX1 | 0.210769423 | 0.001045256 |
| PDE4A | -1.527481588 | 4.25E-09 |
| PPP2R5A | -1.102019065 | 1.69E-08 |
| ELAVL1 | -0.423559193 | 1.53E-06 |
| TIE1 | 1.87513866 | 6.36E-06 |
| DIP2B | -0.970709437 | 5.55E-09 |
| SMARCD1 | -1.109903712 | 3.98E-09 |
| KDM4A | -1.35361812 | 9.14E-10 |
| NFYC | -0.635904833 | 1.43E-06 |
| ZMYND12 | 8.316381629 | 2.05E-07 |
| NGEF | 1.832217292 | 9.81E-05 |
| ASPM | -1.787959886 | 1.21E-11 |
| ELOVL1 | -0.241432317 | 0.000659589 |
| ZNRD1 | -0.764648398 | 0.01309801 |
| ZBTB11 | -0.28558458 | 9.80E-05 |
| ATXN3 | -0.400225368 | 0.00305296 |
| GOLGA5 | 0.599855329 | 1.13E-05 |
| LRRC40 | -0.695855573 | 2.79E-05 |
| ISOC1 | -0.428562678 | 3.04E-05 |
| EML1 | -1.338517825 | 4.16E-06 |
| TRMT11 | -0.284399602 | 0.020507487 |
| THUMPD1 | -1.201160966 | 5.53E-10 |
| MSANTD3 | -0.782109059 | 5.58E-07 |
| ATG2B | -0.553370337 | 1.48E-05 |
| ARFGEF1 | -0.539799599 | 3.89E-06 |
| ZFAT | -0.730943135 | 0.000272411 |
| MTFR1 | -0.621292126 | 2.48E-06 |
| STAG3 | 0.382758991 | 0.034451786 |
| FECH | -0.401721586 | 0.000764163 |
| MYO9A | -0.96942355 | 1.92E-08 |
| IDI1 | 1.797957611 | 9.07E-12 |
| SP100 | -0.374357629 | 0.002928343 |
| KLF6 | -0.596253124 | 9.02E-07 |
| PLPP1 | -0.319843654 | 0.012373111 |
| NEO1 | -0.706045623 | 3.53E-06 |
| TRAM1 | -0.208561139 | 0.002977024 |

| | | |
|---|---|---|
| PHKA1 | -0.366356676 | 0.006425709 |
| TNFRSF1A | -0.241311393 | 0.000551377 |
| EVI5 | -1.031266994 | 1.93E-07 |
| STOML1 | 2.497598153 | 2.58E-10 |
| PKM | 0.124690072 | 0.000441922 |
| DHX29 | -0.265995543 | 0.000422906 |
| DNTTIP2 | -1.219232588 | 3.99E-10 |
| METTL22 | 0.336836852 | 0.006379336 |
| TP53BP1 | -0.762607267 | 1.34E-08 |
| RRP15 | -0.754019836 | 1.32E-07 |
| RHOA | 0.142452688 | 0.001798991 |
| DHX8 | -0.416757659 | 2.45E-06 |
| PRKCZ | 0.573384292 | 0.001964102 |
| SYT1 | -0.66122427 | 0.000774559 |
| NAV3 | -1.874732826 | 0.0001086 |
| ROCK1 | -0.830201886 | 2.39E-08 |
| CBFB | -1.585910846 | 4.69E-11 |
| HYAL2 | 0.510249952 | 1.10E-05 |
| HDAC4 | -0.422195298 | 0.00108206 |
| RASSF1 | 0.714030593 | 1.16E-06 |
| FGFR3 | -0.42175955 | 0.000408529 |
| IFI35 | 0.321359123 | 0.02155302 |
| HEATR6 | -0.401648468 | 0.000103539 |
| COASY | -0.275541014 | 8.26E-05 |
| PLEKHH3 | 0.580949666 | 0.00047469 |
| MEF2A | -0.682847578 | 4.44E-07 |
| OTUD5 | -0.317092245 | 0.000123111 |
| TFE3 | -0.199191788 | 0.008406779 |
| TBC1D25 | -0.344576706 | 0.002178722 |
| ACSL4 | 0.465648445 | 4.65E-06 |
| INPP5A | -1.337234395 | 1.85E-08 |
| GPKOW | -0.209494441 | 0.043498366 |
| GRIPAP1 | 0.142678981 | 0.049040429 |
| FTSJ1 | -0.190087851 | 0.004045075 |
| PRR11 | -0.451704138 | 7.47E-06 |
| ATP11A | -0.850522974 | 2.46E-08 |
| POLR1A | -1.232898639 | 2.45E-10 |
| TTC7A | -0.828758071 | 4.15E-06 |
| IP6K2 | 0.553908085 | 1.07E-05 |
| STON1-GTF2A1L | 1.639892206 | 0.000934903 |
| SRBD1 | 0.893411849 | 4.90E-06 |
| KIF2A | -0.555553954 | 1.71E-06 |
| RASGRP2 | 8.195340173 | 2.95E-07 |
| IFT80 | -0.181634324 | 0.031422989 |
| SIRT2 | 0.225707656 | 0.033747358 |
| ERLEC1 | -0.354822386 | 3.02E-05 |
| PPP2R5B | 0.771997426 | 9.89E-05 |
| PYGM | 3.900266989 | 5.89E-10 |

| | | |
|---|---|---|
| MAST4 | 0.582221549 | 0.002769215 |
| SDK2 | 2.18394466 | 0.005879426 |
| NUP133 | -0.719417822 | 1.89E-07 |
| NUCKS1 | -0.333186754 | 4.91E-07 |
| DNAJA2 | -0.562778159 | 4.31E-07 |
| BCL3 | -1.132051941 | 4.37E-05 |
| KCNAB2 | 1.180866807 | 2.64E-06 |
| CLEC2D | 0.590679826 | 0.000153133 |
| FUNDC1 | -0.262134984 | 0.030646429 |
| RORA | 1.10720157 | 0.000511915 |
| TGFBR3 | -0.808340517 | 0.000351505 |
| PLA2G10 | 3.852114453 | 2.91E-06 |
| HES2 | 1.365381045 | 0.000355102 |
| ATP1B3 | -0.109493301 | 0.023784179 |
| NEDD4 | -1.285905312 | 2.08E-10 |
| MAPK6 | -0.752157135 | 8.04E-09 |
| GNB5 | -0.75952121 | 1.16E-06 |
| PHRF1 | -0.786424855 | 0.00425102 |
| ELP1 | -0.220302584 | 0.002231079 |
| NUCB2 | 0.496090666 | 0.004436017 |
| PTPN3 | -2.275666558 | 6.46E-10 |
| DAPP1 | 3.412535166 | 0.000225993 |
| SLC44A1 | -0.221846397 | 0.000547977 |
| TMEM260 | 1.887654243 | 2.90E-09 |
| SMG6 | -0.282257879 | 0.000963505 |
| EXOC5 | -0.681086387 | 1.03E-07 |
| CLTCL1 | 0.650789081 | 0.00079363 |
| DGCR2 | -0.476551287 | 1.62E-05 |
| RNF126 | -0.36992005 | 0.000441514 |
| MNT | 1.038930642 | 8.24E-09 |
| ZXDC | -0.543224953 | 2.89E-06 |
| JMJD6 | -0.703831284 | 1.47E-06 |
| WIPI1 | 1.378904296 | 2.98E-06 |
| FRMPD1 | 4.290934621 | 1.30E-05 |
| GBA2 | -1.175814158 | 3.38E-09 |
| NDST1 | -0.715165655 | 1.30E-07 |
| ASNS | 1.086920629 | 1.73E-09 |
| AP3M2 | -1.28404418 | 3.45E-08 |
| ST6GALNAC2 | -2.262484398 | 4.26E-07 |
| PABPC1 | 0.131300787 | 0.00239482 |
| TESK2 | 0.678931558 | 6.01E-05 |
| CSNK2A2 | -0.718382754 | 1.25E-08 |
| PTPN21 | -0.962702339 | 9.16E-06 |
| TCOF1 | -0.218877475 | 0.000916654 |
| CDC42 | -0.751031265 | 6.77E-09 |
| OSBPL3 | -1.062116885 | 1.51E-08 |
| EPHA8 | 10.37278074 | 2.63E-10 |
| RAD18 | -1.208967511 | 1.98E-09 |

| | | |
|---|---|---|
| ATP2B1 | -0.490223374 | 2.10E-06 |
| MAP4K4 | -0.337010048 | 4.16E-05 |
| RPL31 | 0.897416031 | 5.57E-09 |
| WDR1 | -0.539420279 | 7.48E-08 |
| SNX13 | -0.366962796 | 0.000416709 |
| ARHGAP10 | -1.276514656 | 2.64E-05 |
| RPS6KA2 | 0.765703953 | 2.46E-05 |
| VASH1 | -0.427151299 | 0.00073359 |
| LMCD1 | 0.983256314 | 1.27E-06 |
| TRIP13 | -0.85603732 | 1.01E-08 |
| ATP6AP1 | 0.204887824 | 0.008844055 |
| TRIB2 | -1.22994588 | 4.14E-09 |
| DAZAP1 | -0.328976873 | 7.95E-05 |
| FAM50A | 0.292833587 | 0.001044261 |
| CYBRD1 | 0.403002597 | 0.000170427 |
| CDH19 | 0.273083661 | 0.003285303 |
| SLC6A15 | -0.395193204 | 1.50E-05 |
| RDH11 | 0.517886962 | 5.98E-07 |
| ADGRL1 | 0.282293954 | 0.00097222 |
| ACTN1 | -0.48070513 | 5.84E-07 |
| ZFYVE26 | -0.631833603 | 2.79E-05 |
| EPN2 | -0.32460094 | 0.000423207 |
| PTPN18 | -0.66399046 | 6.06E-06 |
| LIMS2 | 1.667216395 | 0.002603017 |
| ASIC4 | 2.393682283 | 0.009197675 |
| SPEG | 0.422279456 | 0.016649514 |
| LNX1 | -0.53323093 | 0.001572187 |
| ALDH3A2 | -1.197161269 | 1.50E-09 |
| TFRC | -1.110396271 | 8.82E-11 |
| SREBF1 | 0.420784183 | 9.93E-06 |
| AFF4 | 0.177549687 | 0.000600009 |
| UBE2D1 | -0.355802199 | 0.018104606 |
| MPP5 | -1.440859977 | 1.23E-09 |
| RHOBTB1 | -1.238775444 | 0.000408606 |
| SMC1A | -0.237606635 | 4.17E-05 |
| MARK2 | -0.304954893 | 0.000175176 |
| HMMR | 0.246198793 | 0.001942209 |
| TRHDE | -0.341458943 | 0.000101425 |
| P4HA2 | 0.337826372 | 0.002466479 |
| NFATC3 | -1.29955861 | 2.33E-10 |
| TRNT1 | -1.240259008 | 4.62E-09 |
| ACADVL | 0.503734085 | 1.01E-07 |
| STK10 | -0.520191097 | 8.55E-07 |
| FBXW11 | -0.677836396 | 9.41E-08 |
| ACAP1 | 3.053667561 | 6.84E-08 |
| CRMP1 | -0.262828115 | 0.023167167 |
| EVC | -0.236823341 | 0.002505122 |
| DERL2 | -0.239747423 | 0.002529498 |

| | | |
|---|---|---|
| NDE1 | -0.875166192 | 1.49E-05 |
| TMEM38A | 0.714383951 | 0.016947245 |
| AP1M1 | -0.163577921 | 0.008913149 |
| PVR | -0.735299187 | 5.00E-08 |
| XRCC1 | 0.503872805 | 0.000177662 |
| SCARB1 | -0.125525933 | 0.034451786 |
| MCM2 | -0.16512691 | 0.002285698 |
| SELENOO | -0.405488902 | 0.004298712 |
| ALPK1 | 1.019175363 | 3.71E-05 |
| PDE8A | -0.453809071 | 4.60E-05 |
| CLCN4 | -1.016208302 | 4.86E-05 |
| NLE1 | -0.350424449 | 0.000308297 |
| SDHA | 0.228575342 | 0.002511287 |
| SMARCE1 | -0.234313933 | 0.00069073 |
| GSDMB | 2.299485643 | 1.16E-09 |
| KDM5A | -1.125785728 | 1.04E-09 |
| ADAM11 | 0.868884979 | 0.00029818 |
| PPP2R3A | -1.238588187 | 1.86E-09 |
| FERMT2 | -0.212349271 | 0.002759589 |
| PTGS2 | 1.118247423 | 0.000377675 |
| IGF2BP2 | -0.64404935 | 1.57E-07 |
| FRY | 0.962418877 | 0.002365383 |
| PICALM | -0.638957193 | 3.58E-08 |
| NSF | -0.466987652 | 1.82E-05 |
| GLI2 | -1.450177141 | 7.82E-08 |
| CLASP1 | -1.318470562 | 8.65E-10 |
| NOTCH3 | -0.447598467 | 0.001120686 |
| CLNS1A | -0.146304797 | 0.008265089 |
| PPP2R2C | 2.937887556 | 1.92E-05 |
| EED | -0.362549638 | 0.000312235 |
| CDHR2 | 1.042338724 | 0.00447969 |
| NCBP3 | -1.534109357 | 1.34E-09 |
| ATP2A3 | 1.628207805 | 0.029370221 |
| NTN4 | -0.558100498 | 0.003562211 |
| NUAK1 | -2.037648593 | 4.95E-08 |
| DPP8 | -0.571979397 | 2.69E-06 |
| SLC24A1 | 0.554282576 | 7.60E-05 |
| ZNF532 | -0.495994879 | 2.95E-06 |
| HACD3 | -0.529890764 | 1.46E-07 |
| ZZEF1 | -0.258125156 | 0.002044961 |
| ENO1 | 0.159458772 | 0.003467082 |
| SLC12A1 | 9.639851358 | 2.18E-09 |
| ANO8 | 0.305024328 | 0.008564927 |
| TUBE1 | -0.25141792 | 0.005186597 |
| ARHGEF10L | -1.089779134 | 0.007806804 |
| TXK | -0.894744693 | 0.000467251 |
| TACR2 | 2.116344171 | 5.74E-05 |
| ACTR6 | -0.215547715 | 0.035891489 |

| | | |
|---|---|---|
| EIF4G3 | -0.509223938 | 5.57E-07 |
| SEMA3A | -0.612698319 | 4.26E-05 |
| GTSE1 | -1.514313511 | 7.72E-09 |
| SEMA3C | -0.123472803 | 0.045742917 |
| ACAT1 | 0.228490615 | 0.007277646 |
| GRAMD4 | -0.57761125 | 0.000759048 |
| CELSR1 | -0.421816021 | 4.64E-05 |
| WNT8B | 3.467825129 | 5.68E-07 |
| ZNF638 | -0.639456218 | 9.31E-08 |
| SLC25A40 | -1.295200138 | 1.13E-08 |
| TIMM21 | -1.303958777 | 1.47E-08 |
| ADD2 | -0.374948257 | 0.001002786 |
| RASAL2 | -1.048660474 | 2.95E-09 |
| ZNF37A | -0.812576627 | 3.94E-08 |
| MARK3 | -0.738740419 | 8.17E-08 |
| SLC25A3 | 0.181556273 | 0.000544867 |
| FOSL2 | -0.420108928 | 3.67E-05 |
| FRYL | -1.000032076 | 1.40E-08 |
| TMEM131 | -0.600311076 | 2.06E-06 |
| FSCN1 | -0.423022703 | 9.83E-06 |
| ACTB | -0.367647489 | 3.71E-09 |
| MOCOS | -0.329826937 | 0.000110655 |
| PLD1 | 0.414230926 | 0.025530699 |
| WDR62 | -1.24847348 | 1.28E-08 |
| DLG1 | -1.55995273 | 9.66E-10 |
| RAB7A | -0.236890878 | 0.000104488 |
| BCAP29 | -0.411526767 | 2.53E-05 |
| SEC31B | 1.837779377 | 2.71E-08 |
| SART3 | -1.197832484 | 6.62E-10 |
| EXOSC7 | -0.380810369 | 0.003164224 |
| MKRN2 | -0.400287464 | 2.17E-05 |
| REXO2 | -0.136353032 | 0.040720082 |
| RBMS2 | -1.198212991 | 2.99E-09 |
| BAZ2A | -0.505567858 | 2.21E-07 |
| PTPN23 | -0.493883361 | 1.35E-06 |
| MLH1 | 0.709590639 | 6.15E-07 |
| UNG | -0.678547057 | 4.49E-08 |
| FMO4 | 1.32626299 | 0.003369493 |
| KLHL20 | -2.019138884 | 1.80E-10 |
| RGS11 | 5.920741351 | 2.63E-05 |
| SLC46A1 | -0.18951482 | 0.00642591 |
| PLXNA2 | -0.773319744 | 0.00018134 |
| SPAG5 | -0.514043756 | 4.10E-07 |
| TRAF4 | -0.301502318 | 0.000184128 |
| PAG1 | -0.198202685 | 0.007912557 |
| GPATCH1 | -0.534372224 | 0.001475782 |
| ICAM3 | 0.436202629 | 0.001524685 |
| NT5C2 | -1.148263491 | 4.74E-09 |

| | | |
|---|---|---|
| MCAM | 0.396351237 | 9.78E-07 |
| RAP1GAP | 0.952341656 | 0.003073669 |
| ARHGEF1 | 0.221260357 | 0.001701385 |
| STXBP2 | 4.072672752 | 3.22E-05 |
| MAP2K7 | -0.503771094 | 4.91E-06 |
| DGKD | -0.787774209 | 3.60E-07 |
| RARB | 0.80409285 | 0.033599228 |
| TOP2B | -0.16063029 | 0.005541547 |
| TM9SF3 | -0.723812511 | 4.79E-09 |
| NFKB2 | 1.277054168 | 1.84E-10 |
| UBE2T | -0.288209124 | 0.000435498 |
| PPP1R12B | -0.833445005 | 2.88E-07 |
| GTF3C1 | -0.880523146 | 5.12E-09 |
| IL4R | 0.236167159 | 0.016957218 |
| USP33 | -0.251383656 | 0.000773145 |
| PAK3 | 1.620085245 | 0.000163069 |
| SNRPA | 0.14728066 | 0.014955824 |
| EXOSC5 | 0.426001621 | 0.000357068 |
| FAM76B | -0.753305784 | 1.50E-06 |
| POLD3 | -2.172898317 | 1.32E-10 |
| ACTN2 | 2.702029762 | 1.19E-07 |
| GPR137B | -0.533689097 | 0.00132275 |
| JADE1 | -1.308709184 | 1.79E-09 |
| FGFR1 | -0.226451256 | 0.004531047 |
| FKBP6 | 2.410462973 | 0.001053772 |
| FBLN1 | 0.733650791 | 0.023543949 |
| MAP2 | 1.768715358 | 0.008183946 |
| PIAS2 | -1.345699851 | 2.28E-08 |
| ARAF | -0.229436029 | 0.001637615 |
| ACER3 | -1.134486729 | 5.21E-09 |
| UBE2K | -1.099662581 | 4.84E-10 |
| N4BP2 | -0.877723758 | 5.25E-08 |
| TIGAR | -0.18594575 | 0.016708785 |
| TULP3 | -0.252544303 | 0.00171221 |
| SYNJ2 | -1.154023991 | 2.64E-09 |
| PPP2R5C | 0.426278668 | 5.88E-05 |
| MLLT10 | -1.295019972 | 3.38E-09 |
| ZCWPW1 | 1.053320256 | 7.53E-05 |
| NRDC | 0.193466624 | 0.002931654 |
| VDAC3 | -0.138046584 | 0.044281304 |
| PCM1 | -0.235885689 | 0.001413063 |
| TNRC6C | -0.16590203 | 0.035963314 |
| ITCH | -1.220731178 | 4.79E-10 |
| PKD2L2 | -0.538645432 | 0.002158684 |
| TP53INP2 | 1.200034119 | 2.55E-06 |
| SDF4 | 0.205677164 | 0.009441003 |
| MYH7B | 3.674717878 | 1.42E-06 |
| TP73 | -1.462777276 | 3.50E-07 |

| | | |
|---|---|---|
| TOLLIP | -1.259281861 | 1.08E-08 |
| UBE2D4 | -0.854581458 | 4.48E-06 |
| RUNX1T1 | -2.440568636 | 5.74E-07 |
| THOC1 | -0.459733445 | 0.00029134 |
| FKBP7 | 0.703523076 | 0.002243625 |
| OSBPL6 | -0.661548416 | 8.56E-05 |
| SLC1A3 | -0.937979137 | 4.64E-06 |
| XRCC5 | -0.182533423 | 0.00291052 |
| SP140 | -0.944459929 | 0.015097533 |
| MKNK1 | 0.187902966 | 0.03871093 |
| TNS1 | -0.863779318 | 0.013726086 |
| REXO1 | -1.158911318 | 4.74E-08 |
| SAR1A | -1.281123273 | 2.58E-10 |
| CDC14A | -0.888004719 | 6.55E-06 |
| RAPGEF3 | 1.537241479 | 3.62E-06 |
| CEACAM1 | 1.372128338 | 0.008185126 |
| SENP1 | -0.963053835 | 5.06E-08 |
| FDFT1 | 0.77759753 | 1.21E-05 |
| OPHN1 | -0.866124282 | 0.000188433 |
| KIF22 | -0.611314414 | 1.30E-07 |
| CARMIL1 | -1.331251173 | 2.53E-05 |
| PGM1 | 0.522203131 | 4.19E-06 |
| DNM2 | -0.11395296 | 0.027293361 |
| MOXD1 | 0.421802625 | 0.024045465 |
| STX7 | -0.621057799 | 0.000153401 |
| RABL2B | 0.613844853 | 2.90E-05 |
| KEAP1 | -1.203736692 | 5.59E-09 |
| PTPRH | 1.838192566 | 1.62E-07 |
| CRYBG3 | -1.050444675 | 5.00E-09 |
| EPHA6 | 1.629177226 | 0.010215025 |
| RFX3 | -1.241123464 | 5.66E-07 |
| RIF1 | -0.868367272 | 2.29E-08 |
| RAB21 | -0.526979514 | 1.51E-06 |
| SMARCA2 | 0.186488446 | 0.009090813 |
| RDH8 | 4.752049801 | 0.000405365 |
| SESN1 | 1.066620873 | 5.43E-07 |
| MID2 | -1.425455293 | 0.014767265 |
| COL5A3 | 1.086131189 | 3.13E-05 |
| SRCAP | -0.960242852 | 3.75E-10 |
| PUM3 | -0.732615199 | 3.30E-08 |
| KCNN2 | 5.288004761 | 5.05E-05 |
| CNOT4 | -0.475771942 | 2.37E-05 |
| PSEN1 | -0.316125922 | 0.0002987 |
| CPOX | -0.476615325 | 8.58E-07 |
| CLDND1 | -0.328611132 | 1.77E-05 |
| MOK | -1.041817108 | 3.42E-09 |
| HSP90AA1 | 0.161515625 | 1.20E-06 |
| RBL1 | -0.994932907 | 1.22E-08 |

| | | |
|---|---|---|
| DLGAP4 | -0.785361481 | 2.45E-08 |
| NDC80 | -1.677084975 | 1.25E-08 |
| AP4E1 | -1.047026982 | 1.85E-08 |
| RSBN1 | -0.890922074 | 1.61E-07 |
| MAGI3 | -1.185186765 | 2.81E-06 |
| CXCL2 | 0.930416149 | 0.001342065 |
| COL4A4 | 1.969067547 | 0.00038061 |
| OSTM1 | -0.52583153 | 3.35E-05 |
| EXD2 | -0.865860732 | 2.05E-06 |
| UBA5 | -0.820450847 | 2.45E-08 |
| STK17B | -0.743066167 | 2.09E-05 |
| CDC14B | -0.878046017 | 2.90E-07 |
| ZNF510 | 0.403663033 | 0.000112809 |
| ZNF506 | 0.186510268 | 0.032493155 |
| JMJD4 | 0.48937153 | 1.12E-05 |
| DUSP12 | 0.248585731 | 0.002013288 |
| DELE1 | -0.479905693 | 5.02E-06 |
| PCDHA6 | -0.901148888 | 3.47E-08 |
| PHLPP1 | -0.971920631 | 4.30E-06 |
| SMARCD3 | 0.631316129 | 0.000254281 |
| MPP4 | 0.714683887 | 0.005235127 |
| BZW1 | -0.378072232 | 1.48E-06 |
| ME2 | 0.291529362 | 0.000575115 |
| C5orf22 | -1.063814736 | 3.32E-09 |
| CCNT2 | 0.2809643 | 0.000804777 |
| FAM135A | -0.877283578 | 3.03E-06 |
| COBLL1 | -1.167154415 | 1.09E-05 |
| DLG3 | -0.75255119 | 0.000152738 |
| KCNK2 | -1.304748616 | 0.03497479 |
| SERTAD4 | -3.046159647 | 7.83E-13 |
| TRAF5 | -2.155786561 | 3.62E-10 |
| GEMIN5 | -0.352465278 | 2.73E-05 |
| NFE2L1 | -0.09206582 | 0.028462899 |
| GSK3B | -0.73268798 | 2.20E-08 |
| ERC1 | -1.337585025 | 3.35E-10 |
| XPO1 | -1.284329965 | 2.12E-11 |
| RNF13 | -0.292502608 | 0.001082061 |
| PALB2 | 0.84789492 | 7.60E-08 |
| TUT7 | -0.423408958 | 0.000146799 |
| TNPO1 | -0.829712904 | 1.73E-09 |
| PLOD1 | 0.537981717 | 8.53E-07 |
| P2RX5 | -0.485050016 | 0.004856806 |
| ITGAE | -0.716450012 | 0.000163343 |
| DIS3 | -0.764130287 | 2.73E-08 |
| PIBF1 | 0.318697875 | 0.008162148 |
| TDRD3 | -0.45025208 | 0.0032465 |
| NUFIP1 | -1.042586335 | 1.31E-06 |
| PDS5B | -1.258836943 | 4.67E-09 |

| | | |
|---|---|---|
| OXCT1 | -0.15882259 | 0.021047818 |
| RRAGB | 0.624678012 | 0.001154691 |
| SLC27A5 | 0.732670794 | 1.38E-05 |
| ZNF324 | 1.271907498 | 9.80E-10 |
| ZNF671 | 0.473234927 | 0.017200551 |
| ZNF586 | -1.458835884 | 2.33E-08 |
| ZNF446 | 1.060866916 | 1.01E-07 |
| ZNF264 | 1.018537784 | 5.45E-09 |
| RPS5 | 0.51302991 | 2.02E-07 |
| FAT1 | -0.735154278 | 5.90E-10 |
| SMAP2 | -0.782917189 | 2.48E-07 |
| PPIE | 0.140236208 | 0.049836417 |
| ZMPSTE24 | -0.856494563 | 8.44E-09 |
| STARD7 | -1.236523858 | 1.03E-10 |
| NOA1 | 0.213906669 | 0.009221691 |
| REST | -0.74282915 | 3.98E-08 |
| HAL | 4.562672061 | 4.48E-08 |
| SSH1 | -1.012607072 | 4.99E-10 |
| GSTP1 | 0.283647317 | 6.74E-05 |
| FAM234B | -0.857891863 | 4.07E-05 |
| WBP11 | -0.531483115 | 7.02E-07 |
| EIF3I | 0.296819795 | 0.000528516 |
| TXLNA | -1.064526763 | 5.47E-10 |
| NCOA1 | -1.2747229 | 1.38E-08 |
| AGBL5 | 0.244409161 | 0.003914321 |
| EFR3B | 1.262319092 | 1.41E-07 |
| KIF3C | 0.176459229 | 0.007758522 |
| RAB10 | -1.055034098 | 4.08E-10 |
| HADHA | 0.218750951 | 0.001078106 |
| MAPRE3 | 0.411021549 | 0.003975709 |
| CAD | -0.318490854 | 1.79E-05 |
| CD59 | -0.584349801 | 4.46E-08 |
| BCORL1 | 0.407767279 | 0.00011576 |
| ATRX | -0.667936516 | 3.14E-08 |
| AK6 | -0.321067685 | 0.017207636 |
| MYNN | -0.396154322 | 0.000813661 |
| MECOM | -0.984866068 | 1.57E-05 |
| SCAMP1 | -0.600167712 | 3.72E-06 |
| PREP | -0.616692194 | 5.02E-06 |
| HACE1 | 0.410461059 | 0.003546624 |
| SEH1L | -0.331279189 | 7.60E-05 |
| WDFY1 | -0.599409849 | 3.57E-06 |
| OVGP1 | 2.223656593 | 7.63E-07 |
| SLC25A24 | -0.754443677 | 1.03E-07 |
| MAP3K4 | -1.478814191 | 7.71E-09 |
| PILRA | 1.919281082 | 0.000463426 |
| IGSF9 | 1.251390966 | 0.000343134 |
| ABCB1 | -0.539436345 | 0.001753005 |

| | | |
|---|---|---|
| AKR1B1 | 0.249592788 | 0.001222676 |
| CPNE3 | -0.33639408 | 6.91E-06 |
| RRN3 | -1.211361855 | 3.06E-07 |
| CTTN | -0.437378351 | 2.18E-06 |
| MTIF2 | 0.173184709 | 0.011595453 |
| DDHD2 | -0.551704878 | 1.31E-05 |
| TTC39A | 1.42837522 | 0.019962039 |
| EPS15 | -0.736970538 | 3.08E-07 |
| ORC1 | -0.812261001 | 3.19E-07 |
| CHERP | -0.726007207 | 3.19E-08 |
| ATG16L1 | -0.885358364 | 1.60E-07 |
| USP40 | 0.859639954 | 3.81E-06 |
| RAD54L | -1.120405201 | 8.64E-09 |
| MAST2 | -0.667588555 | 4.91E-07 |
| DNAJA1 | -0.399419281 | 3.38E-07 |
| B4GALT1 | -0.148344924 | 0.011630738 |
| CHMP5 | 0.193814482 | 0.002945071 |
| NFX1 | -0.518849068 | 3.83E-06 |
| DIMT1 | -0.890500095 | 1.04E-08 |
| IPO11 | -0.41962501 | 3.38E-05 |
| EIF2AK1 | -0.40378426 | 5.21E-06 |
| EPDR1 | -1.057014133 | 1.39E-08 |
| SNX10 | 0.203392626 | 0.009456185 |
| SEPHS1 | -0.911986959 | 3.52E-07 |
| MRPL28 | 0.186023437 | 0.008135205 |
| ITPKC | 0.69954001 | 6.63E-06 |
| FAT2 | 3.60063454 | 1.04E-08 |
| RBM22 | -1.123505135 | 8.99E-10 |
| TMED2 | -0.079138251 | 0.046849161 |
| ERO1B | -0.208871098 | 0.00989614 |
| ZFAND6 | 0.469335729 | 4.82E-06 |
| TXLNG | -1.115295398 | 6.21E-10 |
| PPEF1 | 1.49331989 | 0.010379886 |
| LAT2 | 2.228095966 | 1.55E-07 |
| HUWE1 | -0.175585332 | 0.000689812 |
| ZW10 | -0.601319495 | 3.90E-06 |
| ALG9 | -0.788081406 | 9.56E-05 |
| MTMR2 | -1.080298531 | 3.02E-09 |
| PPP1R15A | 1.435257873 | 3.52E-11 |
| TRIP6 | 0.265267546 | 0.001444981 |
| FTL | 0.248456356 | 5.36E-05 |
| SRRT | -0.940268349 | 3.73E-09 |
| BAX | 0.219544205 | 0.014292404 |
| NLK | -0.170901361 | 0.021918886 |
| ATXN7L3 | -0.294470771 | 0.000102686 |
| PGS1 | -0.596403201 | 3.46E-05 |
| PSMC5 | 0.226800748 | 0.000162078 |
| MMP2 | 0.420549731 | 9.03E-06 |

| | | |
|---|---|---|
| LPCAT2 | -0.506420122 | 4.22E-06 |
| GNAO1 | -0.501910799 | 0.006816744 |
| OGFOD1 | -0.747636264 | 4.23E-08 |
| SH3BP2 | -0.686170329 | 1.89E-07 |
| NOP14 | -0.503010429 | 1.75E-05 |
| ADD1 | -0.287702867 | 6.60E-05 |
| L2HGDH | -1.019176473 | 5.39E-07 |
| RTRAF | 0.165040173 | 0.012697445 |
| NID2 | 1.780330887 | 7.25E-06 |
| GMCL1 | -1.089509162 | 4.58E-09 |
| KLHL42 | -2.23075376 | 6.47E-12 |
| GNAS | 0.48317061 | 8.82E-09 |
| DNM1L | -0.725390705 | 1.64E-08 |
| PTHLH | -0.370705814 | 0.04837589 |
| ERGIC2 | -1.217824863 | 1.71E-09 |
| TFAP2C | -1.437602646 | 5.93E-10 |
| AURKA | -2.296536653 | 2.60E-12 |
| CASS4 | 6.72236136 | 1.09E-09 |
| PIR | 0.533880282 | 7.21E-05 |
| AAMDC | 1.169906567 | 0.006408723 |
| RFX2 | 1.003360986 | 1.42E-05 |
| METTL2A | -0.419354812 | 0.000169063 |
| ALG6 | -0.619303398 | 0.000539592 |
| CNOT3 | -0.314357299 | 0.041321635 |
| PTPN4 | -0.981013037 | 5.56E-06 |
| DDX18 | -0.573773254 | 8.60E-08 |
| KHSRP | -0.315837057 | 1.54E-05 |
| GNA11 | -0.625768728 | 1.09E-06 |
| ASAP3 | 0.584934662 | 0.002176192 |
| DNMT3B | -0.798557697 | 0.000246068 |
| TPX2 | -1.117756136 | 3.05E-10 |
| FER1L4 | 2.763447026 | 1.20E-08 |
| PDRG1 | -0.307674828 | 0.00260598 |
| EPB41L1 | -0.220547785 | 0.019785283 |
| ANKRD10 | 0.815304227 | 4.30E-09 |
| TGDS | -0.372773664 | 0.001524915 |
| DOCK3 | 1.391591399 | 6.06E-06 |
| C3orf18 | 1.277279778 | 2.84E-05 |
| COQ9 | -0.526705215 | 5.73E-05 |
| CRLS1 | -0.940329943 | 2.15E-07 |
| ATRN | -0.498231725 | 2.68E-06 |
| SMOX | 0.618670359 | 1.14E-05 |
| SIGLEC1 | 4.864732899 | 0.000373477 |
| NSFL1C | -0.168500691 | 0.01399195 |
| SLC4A11 | 0.479029366 | 0.022815038 |
| ZNF343 | -1.000609698 | 1.80E-06 |
| MAVS | -0.786019828 | 7.58E-09 |
| LZTS3 | 0.412815539 | 0.004038834 |

| | | |
|---|---|---|
| DYNLL1 | -0.319621773 | 3.37E-06 |
| SNX5 | -0.732386659 | 2.80E-08 |
| RPL6 | 0.136732064 | 0.008106483 |
| MAPKAPK5 | -0.786812242 | 3.13E-05 |
| P2RX7 | 2.450679787 | 0.000974881 |
| ESF1 | -0.653147362 | 1.69E-06 |
| ANAPC5 | 0.521583758 | 7.07E-07 |
| SLC23A2 | -0.717258878 | 3.62E-08 |
| TMEM230 | -0.126923604 | 0.037617451 |
| DZANK1 | 1.048683005 | 7.73E-05 |
| KDM2B | -1.263930477 | 1.07E-09 |
| GCN1 | -0.165547496 | 0.005288044 |
| RPLP0 | 0.373314004 | 9.69E-07 |
| PXN | -0.302353726 | 5.39E-06 |
| SIRT4 | 1.6946907 | 9.55E-06 |
| KIF16B | -0.206282207 | 0.029562913 |
| TRMT6 | -0.565823398 | 1.75E-06 |
| CHGB | 2.653059234 | 9.88E-10 |
| PEBP1 | 0.221029234 | 0.000757437 |
| BRAP | -0.851072495 | 1.49E-07 |
| ERP29 | 0.434828913 | 4.20E-06 |
| NOS1 | 3.663249516 | 0.004436017 |
| FUS | -0.553720666 | 2.57E-08 |
| IGBP1 | 0.549462692 | 6.04E-06 |
| GRAMD1A | 0.621136528 | 9.38E-06 |
| FXYD3 | 0.919727531 | 1.57E-07 |
| CMTM1 | 0.950586467 | 0.004436017 |
| GANAB | 0.280642273 | 1.11E-05 |
| GMIP | 0.356726151 | 0.011655106 |
| RBM41 | -1.237788339 | 1.61E-06 |
| BIRC5 | -0.872783774 | 1.52E-08 |
| LAG3 | 3.031150021 | 1.97E-09 |
| OTUB2 | -1.698099022 | 3.52E-05 |
| DDX24 | -0.598066579 | 5.08E-08 |
| ZBTB25 | 0.829895819 | 6.72E-07 |
| NECAP1 | -0.183876459 | 0.005176737 |
| ARHGAP4 | 0.410685215 | 8.17E-05 |
| ANKRD24 | 4.624078266 | 3.34E-09 |
| DHX32 | -0.290545479 | 0.003162058 |
| RCOR1 | -0.756036953 | 6.92E-08 |
| GPATCH2L | -0.6877952 | 1.59E-07 |
| LTBP4 | 0.506819202 | 2.03E-05 |
| BLVRB | 1.139665628 | 3.91E-05 |
| SLC9A1 | -1.212784767 | 3.99E-09 |
| SPTLC1 | -0.396305836 | 0.000249244 |
| PAPOLA | -0.180906572 | 0.001388699 |
| CCNK | -0.576106792 | 5.02E-06 |
| PCBP4 | 0.544942347 | 4.28E-07 |

| | | |
|---|---|---|
| YPEL3 | 1.143361745 | 2.85E-05 |
| MRPS33 | 0.899780735 | 3.36E-06 |
| MAEA | -0.670997571 | 1.78E-07 |
| ICAM1 | 1.048095169 | 3.54E-09 |
| STRN4 | -0.627698916 | 3.50E-07 |
| IRAK3 | 5.493832467 | 8.48E-05 |
| MUL1 | 0.386961859 | 8.67E-05 |
| TFAP4 | -0.415814185 | 0.000386115 |
| PDCD7 | -0.415134117 | 1.15E-05 |
| DNAJB11 | 0.142541258 | 0.025835402 |
| P3H2 | -0.968704645 | 0.000149243 |
| RAB11FIP3 | -1.209628604 | 2.41E-07 |
| ZNF268 | -0.512337871 | 1.84E-06 |
| GOLGA3 | -0.354795938 | 1.28E-05 |
| CERS4 | 1.169888268 | 0.006258178 |
| USP48 | -0.801214654 | 1.46E-07 |
| AARS1 | 0.866845934 | 2.38E-09 |
| KIF4A | -1.056166755 | 4.02E-09 |
| TNRC6A | -1.408151365 | 6.82E-11 |
| PLEKHG2 | -0.320570151 | 0.002705843 |
| NAT14 | 0.589779353 | 0.000147215 |
| PITPNM2 | -1.161878249 | 1.06E-07 |
| RBM27 | -0.123254654 | 0.031096063 |
| OSBPL8 | -0.679582828 | 1.14E-07 |
| DTX2 | -0.418061124 | 0.012969114 |
| NLRC4 | 1.654089452 | 0.000332243 |
| PUS7 | -0.388481413 | 0.000117905 |
| SLC26A4 | 4.142811087 | 1.88E-09 |
| DLD | -0.444506609 | 2.19E-06 |
| WDR7 | -0.884248656 | 1.21E-06 |
| TXNL1 | 0.180935127 | 0.013358437 |
| IL5RA | 6.892710482 | 2.00E-06 |
| CMTM6 | -0.158620723 | 0.001745427 |
| ITGA6 | 0.29338162 | 5.02E-06 |
| MAP3K20 | -0.488234165 | 1.07E-05 |
| SEL1L3 | 0.449348624 | 0.0014102 |
| TF | 4.630384467 | 9.44E-10 |
| CDV3 | -1.03542739 | 2.43E-10 |
| MYO15A | 5.281475834 | 0.000105952 |
| ALKBH5 | -0.916140182 | 6.08E-09 |
| NLRP1 | 2.357309989 | 1.44E-10 |
| PITPNM3 | 2.575414138 | 1.55E-07 |
| ORC6 | -1.232054719 | 2.61E-09 |
| ZFHX4 | -1.538545827 | 3.90E-10 |
| ZC3HC1 | -0.631924363 | 1.69E-05 |
| RGS17 | -0.755377692 | 0.000367743 |
| CCDC80 | 0.978844723 | 2.28E-05 |
| PSME1 | 0.333845983 | 9.66E-05 |

| | | |
|---|---|---|
| PPP2R3C | -0.412578853 | 0.011718585 |
| HAUS4 | 0.953493243 | 1.31E-07 |
| SLC22A17 | 0.997377974 | 0.002252084 |
| RNF31 | -0.453117679 | 0.000111034 |
| G2E3 | -2.162744007 | 3.14E-11 |
| HECTD1 | -1.2952929 | 1.71E-10 |
| HNRNPC | 0.228984761 | 7.36E-05 |
| SUPT16H | -0.445575963 | 3.34E-07 |
| TOX4 | -0.51202528 | 1.59E-06 |
| GEMIN2 | -0.623476949 | 0.001659152 |
| TGM1 | 0.976863429 | 0.025477926 |
| TINF2 | 0.217643325 | 0.034205807 |
| TRPM7 | -0.576673893 | 2.96E-07 |
| TYRO3 | -0.304554518 | 0.002305605 |
| WDR76 | -1.701582024 | 1.09E-09 |
| CAPN3 | 4.570921035 | 9.34E-08 |
| TBX15 | 5.034425173 | 4.12E-07 |
| PHGDH | 0.27239821 | 0.000279215 |
| COL9A3 | 0.401469186 | 0.000550707 |
| EZR | -0.727804899 | 8.99E-08 |
| MYL6 | 0.181167371 | 0.002009052 |
| ago-01 | -0.698707697 | 2.91E-08 |
| TEKT2 | 8.384779425 | 1.45E-07 |
| CLSPN | -0.592062913 | 2.84E-07 |
| RFFL | -0.665129156 | 6.03E-07 |
| UNC13D | 3.206880413 | 7.43E-09 |
| MFSD11 | -1.486529379 | 4.46E-10 |
| DPYSL2 | -0.822585212 | 6.52E-08 |
| TGFB2 | -2.044334955 | 8.66E-05 |
| GPATCH2 | 1.046480374 | 8.83E-09 |
| NUP50 | -1.38654691 | 8.52E-11 |
| COMT | 0.411508997 | 0.000598605 |
| AC016026.1 | -1.290532088 | 7.10E-10 |
| ECHDC1 | -0.629204899 | 4.91E-07 |
| LRRFIP2 | -0.617026787 | 2.18E-06 |
| SEC22C | -1.282000099 | 1.53E-09 |
| XYLB | -0.501717384 | 0.000275853 |
| HDAC6 | 0.959288602 | 2.64E-07 |
| CDC6 | -0.542547178 | 2.68E-06 |
| UPRT | -0.513806507 | 0.00182952 |
| CDC23 | -0.992441229 | 3.12E-09 |
| CBX5 | 0.273658299 | 0.000117865 |
| SUCO | -0.302568634 | 0.000100489 |
| MSH2 | -0.567599315 | 3.96E-07 |
| MAP3K1 | -0.371311486 | 0.000107595 |
| HOOK2 | 1.712640532 | 8.36E-08 |
| EPB41L4B | -1.65704486 | 3.76E-06 |
| TMEM38B | -0.422952617 | 3.31E-05 |

| | | |
|---|---|---|
| PSMD5 | -0.72018478 | 4.97E-08 |
| PTGS1 | 1.752051469 | 3.79E-06 |
| NUP188 | -0.366272217 | 9.23E-06 |
| CRAT | 0.528368554 | 0.001035315 |
| SH2D3C | 4.503813915 | 0.000582386 |
| NANS | -0.320400297 | 0.000474311 |
| TBC1D2 | -1.229692599 | 0.000149186 |
| WHRN | -0.273285315 | 0.042621884 |
| PDE6C | 1.618408465 | 0.005829307 |
| CWF19L1 | -0.579210853 | 3.18E-05 |
| SEMA4G | 1.343983179 | 5.82E-07 |
| BTAF1 | -0.938279232 | 4.82E-09 |
| IKZF5 | 0.897041845 | 1.88E-08 |
| TLL2 | 3.050423206 | 8.38E-07 |
| SORBS1 | -0.525626552 | 9.08E-05 |
| IL11 | 4.656265979 | 6.97E-16 |
| WAC | -1.043434877 | 1.73E-09 |
| CREM | 2.156853585 | 5.03E-12 |
| NUBP2 | -0.215381111 | 0.006078702 |
| SMIM24 | 5.119203467 | 0.000114146 |
| HIVEP1 | -0.671229793 | 5.08E-06 |
| FKBP5 | -0.336598858 | 0.000243728 |
| SRPK1 | -0.594481144 | 7.19E-07 |
| BRPF3 | -0.666097196 | 1.78E-07 |
| TMEM14A | -0.500549625 | 0.00360126 |
| EFHC1 | 0.356191582 | 0.00363188 |
| RPS18 | 0.334659216 | 0.023497291 |
| HSP90AB1 | 0.417380295 | 1.19E-09 |
| CDC5L | -1.132863731 | 2.76E-09 |
| ITPR3 | 0.262645569 | 0.000295368 |
| ZNF184 | 0.866425865 | 0.000244791 |
| SIRT1 | -0.24125799 | 0.001825876 |
| IFT74 | 0.381151355 | 0.021765289 |
| JAK2 | -1.288734737 | 9.94E-08 |
| ABL1 | -0.618179778 | 1.03E-07 |
| ACOT7 | 0.297551491 | 0.000115081 |
| SH3GLB1 | -0.215648687 | 0.00449303 |
| CDC7 | -0.268778664 | 0.010782336 |
| SYDE2 | -0.615844873 | 0.001049213 |
| SCD | 1.712367361 | 3.02E-13 |
| ABLIM1 | -0.809865262 | 4.24E-06 |
| ERMP1 | -0.53146878 | 6.32E-06 |
| RAB18 | -0.60361007 | 3.69E-07 |
| HSD17B7P2 | 3.351001651 | 3.79E-08 |
| WASHC2A | -0.458580316 | 0.000242609 |
| MAST3 | 0.206741914 | 0.031418573 |
| MZF1 | -0.485931982 | 1.06E-05 |
| OCEL1 | 1.213116123 | 2.21E-07 |

| | | |
|---|---|---|
| MYO9B | -0.145024546 | 0.012353948 |
| CATSPERG | 4.294657547 | 8.95E-06 |
| FBXL19 | -0.672370607 | 9.91E-06 |
| STX1B | 2.868124445 | 4.30E-07 |
| HSD3B7 | 1.042503262 | 0.001587611 |
| SETD1A | -0.40014013 | 1.26E-05 |
| BCL7C | 1.005545471 | 1.34E-07 |
| EFNA2 | 3.859591937 | 9.23E-05 |
| CIRBP | 1.007196122 | 3.02E-09 |
| CBARP | 2.245992121 | 4.11E-07 |
| PRKY | -2.921450238 | 8.52E-05 |
| IGFALS | 3.860120422 | 2.28E-05 |
| MARCHF2 | 1.132850653 | 0.002008964 |
| TECR | 0.177265791 | 0.016283473 |
| TIMM13 | -0.221543381 | 0.003065788 |
| CDC34 | -0.350612101 | 0.00050444 |
| MTAP | -0.847948906 | 9.76E-10 |
| CEP170B | 0.436664484 | 1.13E-05 |
| POLRMT | -0.190867402 | 0.016649683 |
| RASSF7 | 0.646732151 | 0.001953606 |
| GADD45B | 3.103469294 | 3.02E-13 |
| MADCAM1 | 1.661340047 | 0.014291729 |
| MKNK2 | -0.645065588 | 9.55E-07 |
| ARVCF | 0.945116884 | 0.006189248 |
| TRMT2A | -0.608992768 | 1.55E-06 |
| RANBP1 | -0.135779963 | 0.027108775 |
| KLHL22 | 1.042462309 | 3.22E-05 |
| SNAP29 | -0.201057027 | 0.041513142 |
| CRKL | -1.33559119 | 1.66E-10 |
| LZTR1 | -0.170051976 | 0.041726631 |
| CECR2 | -0.970705676 | 0.00015762 |
| P2RX6 | 2.399253552 | 3.20E-06 |
| DERL3 | 3.22029237 | 0.000561783 |
| SLC7A4 | 7.402069451 | 5.15E-06 |
| DDTL | 0.445987588 | 0.00336983 |
| DDT | 0.408395508 | 0.005229294 |
| GSTT2 | 2.257793107 | 0.000783452 |
| OSM | 7.996346514 | 5.87E-07 |
| CABIN1 | -0.162037201 | 0.023741954 |
| TBC1D10A | 0.483273404 | 0.007240426 |
| SUSD2 | 0.620221534 | 0.000522653 |
| RNF215 | 0.71242695 | 0.000227428 |
| SEC14L2 | -0.396482802 | 0.002729438 |
| SEC14L3 | 7.273051107 | 7.59E-06 |
| SPECC1L | -0.600550983 | 2.00E-07 |
| PPIL2 | -0.743781871 | 2.19E-07 |
| UPB1 | 7.667375571 | 1.95E-06 |
| YPEL1 | 0.703415078 | 0.006307665 |

| | | |
|---|---|---|
| SNRPD3 | -0.267561826 | 0.000993907 |
| PES1 | -0.271431438 | 0.000281198 |
| MAPK1 | -0.455107258 | 1.65E-06 |
| GGT1 | 3.946067827 | 4.75E-05 |
| PPM1F | -0.382334897 | 2.08E-05 |
| SLC35E4 | -1.266244887 | 2.22E-07 |
| TOP3B | -1.148758467 | 2.32E-05 |
| ESS2 | -0.266287691 | 0.017221553 |
| CRYBB2P1 | -0.453310575 | 0.024432511 |
| MFNG | -1.366623927 | 0.000623019 |
| CARD10 | -2.342750805 | 8.25E-10 |
| SLC25A1 | 1.410429151 | 4.09E-10 |
| GRK3 | 1.365532995 | 0.001405024 |
| GGA1 | -0.578513427 | 1.34E-05 |
| HIRA | -0.579551422 | 8.49E-06 |
| LGALS1 | 0.578564918 | 0.000122313 |
| PIK3IP1 | 5.908688584 | 0.0005541 |
| SRRD | -0.467717902 | 1.04E-05 |
| PATZ1 | -1.063358029 | 2.22E-08 |
| TFIP11 | -0.317102872 | 0.000253581 |
| GCAT | 0.93999968 | 3.83E-05 |
| EIF3L | 0.205458195 | 0.000535713 |
| SNU13 | -0.245383935 | 0.00082134 |
| SOX10 | -0.977719052 | 5.52E-10 |
| DEPDC5 | -0.680877635 | 0.002758676 |
| PICK1 | -0.664388989 | 0.001484098 |
| TTC28 | -1.058684821 | 6.74E-05 |
| SEPTIN3 | 0.815797018 | 0.000298673 |
| DDX17 | -1.526482538 | 5.34E-12 |
| HSCB | 0.657188603 | 0.013875602 |
| CBY1 | 1.30312919 | 5.35E-06 |
| TOMM22 | -0.560206828 | 0.00015731 |
| XBP1 | 0.72020878 | 3.45E-08 |
| RTCB | -0.189459438 | 0.021385571 |
| JOSD1 | 1.528572361 | 4.31E-11 |
| GTPBP1 | -0.342835483 | 0.001414569 |
| POLDIP3 | -0.744156104 | 3.43E-08 |
| RAB36 | -0.590261048 | 0.000636811 |
| TIMP3 | 0.652821784 | 2.60E-10 |
| PPP6R2 | -0.743771931 | 8.33E-06 |
| SBF1 | -0.230259509 | 0.000617452 |
| SUN2 | 0.143162416 | 0.043152735 |
| CYB5R3 | 0.175870341 | 0.007237428 |
| DNAL4 | 1.091784491 | 1.74E-05 |
| LMF2 | 0.154553459 | 0.048153031 |
| PACSIN2 | -0.821317137 | 1.35E-07 |
| TTLL1 | 1.401077748 | 0.000877088 |
| RASL10A | 2.364551541 | 2.76E-05 |

| | | |
|---|---|---|
| AP1B1 | -0.333553909 | 0.000223091 |
| HMGXB4 | -1.086219152 | 2.91E-09 |
| TOM1 | 0.615816514 | 0.000673729 |
| CHKB | -0.494741779 | 0.008452197 |
| HMOX1 | -0.525059879 | 0.002184131 |
| MCM5 | -0.448740008 | 8.44E-05 |
| ARSA | 0.530173987 | 0.009084653 |
| TSPO | 0.427976731 | 0.000446808 |
| RASD2 | 1.177126659 | 0.00015629 |
| TTLL12 | -0.731177161 | 6.86E-08 |
| CBX7 | -0.715273327 | 0.000413577 |
| PDGFB | -2.279877352 | 0.001329754 |
| CABP7 | 0.643222556 | 0.009120934 |
| RPL3 | 0.530382153 | 1.16E-07 |
| ZMAT5 | 0.583253986 | 0.012961361 |
| RBFOX2 | -1.035130988 | 1.75E-09 |
| SYNGR1 | 0.735215816 | 0.002380868 |
| TAB1 | -0.744157276 | 9.55E-06 |
| ASCC2 | -0.236029092 | 0.025236074 |
| MTMR3 | -1.003125497 | 1.36E-07 |
| MIEF1 | -1.013025173 | 1.84E-09 |
| APOL4 | 4.82744295 | 0.000213797 |
| MYH9 | -0.279458233 | 3.61E-05 |
| CACNA1I | 3.135308052 | 0.000690679 |
| FOXRED2 | -0.76459021 | 1.89E-07 |
| EIF3D | 0.232409878 | 0.002346297 |
| TNRC6B | 1.050123848 | 2.38E-09 |
| SGSM3 | 0.369774767 | 0.000655089 |
| IFT27 | 1.204337575 | 4.68E-05 |
| KIAA0930 | -0.16774649 | 0.022126159 |
| SLC25A17 | -1.069339738 | 3.43E-07 |
| FAM118A | -1.14243015 | 3.66E-07 |
| IL2RB | 6.480758263 | 9.35E-10 |
| EP300 | -1.000064725 | 9.19E-09 |
| L3MBTL2 | -1.378960884 | 1.05E-08 |
| RANGAP1 | -0.65396947 | 5.42E-08 |
| ZC3H7B | -0.4571665 | 3.82E-06 |
| PHF5A | -1.054844169 | 1.55E-08 |
| POLR3H | -0.78330373 | 5.02E-08 |
| TRMU | -0.873409851 | 8.94E-08 |
| PMM1 | 0.737197549 | 0.000106976 |
| DESI1 | -0.305217216 | 0.000412642 |
| CERK | -1.365298612 | 1.58E-09 |
| BRD1 | -0.949661358 | 1.08E-06 |
| HDAC10 | -0.369522473 | 0.009207748 |
| ABHD4 | 0.419086777 | 0.001393239 |
| KHNYN | 0.284414628 | 0.000107855 |
| RBM23 | -0.561411239 | 9.89E-07 |

| | | |
|---|---|---|
| PRMT5 | 0.150764636 | 0.009064948 |
| AP4S1 | -0.522416449 | 0.011075081 |
| POLE2 | -0.888928556 | 0.00089529 |
| SOS2 | -0.3829692 | 0.0017025 |
| NIN | -0.396265645 | 6.55E-06 |
| PYGL | 0.151015485 | 0.028581609 |
| PSMC6 | -0.411907201 | 8.87E-05 |
| GNPNAT1 | -1.332301673 | 1.81E-10 |
| DDHD1 | -1.976595414 | 3.50E-09 |
| CNIH1 | -0.301453106 | 0.000117591 |
| CGRRF1 | 0.656919255 | 0.000513112 |
| ATP6V1D | -0.824472392 | 7.27E-07 |
| PIGH | -0.256304973 | 0.010715499 |
| GSTZ1 | 2.000004751 | 9.42E-08 |
| TMED8 | -0.672284407 | 1.30E-07 |
| SAMD15 | -0.730488578 | 0.004109518 |
| SPTLC2 | -1.060787289 | 5.90E-09 |
| AL159141.1 | -0.593212881 | 0.000328579 |
| LGMN | 0.275065937 | 0.017000696 |
| ALKBH1 | -0.384804882 | 0.001975911 |
| SNW1 | -0.971741629 | 1.55E-08 |
| PPM1A | -0.698423378 | 6.09E-07 |
| SIX4 | 0.21030925 | 0.023217592 |
| GALNT16 | 2.385802417 | 0.008416564 |
| CEP128 | -0.47690471 | 0.002095852 |
| ERH | -0.232411232 | 0.006865155 |
| HIF1A | -0.66788027 | 1.44E-08 |
| SUSD6 | 0.181383754 | 0.03580952 |
| SRSF5 | -0.128866251 | 0.020196365 |
| SLC10A1 | 3.643272263 | 0.010448035 |
| EIF5 | -0.892057714 | 3.29E-10 |
| DICER1 | -1.949861595 | 5.17E-11 |
| ZC3H14 | -0.831300177 | 2.41E-08 |
| TELO2 | -1.020872587 | 2.19E-07 |
| PCNX1 | -0.232955827 | 0.001097186 |
| GSKIP | 0.579352496 | 6.80E-07 |
| PAPLN | 1.306674217 | 4.34E-06 |
| RPS6KA5 | -0.996955573 | 1.99E-05 |
| PPP4R3A | -0.994047726 | 3.33E-09 |
| C14orf93 | 1.537412153 | 2.41E-07 |
| YY1 | -0.472947478 | 1.35E-06 |
| ACIN1 | -0.605836228 | 3.74E-08 |
| CCNB1IP1 | 0.704091608 | 1.21E-06 |
| TRIP11 | -0.52464118 | 4.09E-06 |
| APEX1 | 0.1723359 | 0.002367462 |
| PABPN1 | -0.37032588 | 1.82E-05 |
| ARHGAP5 | -0.44683161 | 2.44E-06 |
| CINP | -0.413113752 | 0.000407693 |

| | | |
|---|---|---|
| DHRS2 | -0.474718448 | 0.00017846 |
| CHD8 | -0.638152318 | 1.41E-07 |
| PCK2 | 1.285445402 | 2.31E-08 |
| PRORP | -0.869619839 | 5.75E-07 |
| DCAF11 | 0.187612322 | 0.017591453 |
| NFKBIA | 0.989863303 | 2.40E-08 |
| PSME2 | 0.221461733 | 0.012034907 |
| BRMS1L | -1.284210222 | 2.29E-07 |
| REC8 | 1.186064346 | 0.000634755 |
| TM9SF1 | -0.152938239 | 0.019199404 |
| PNN | -0.526578944 | 5.69E-07 |
| PLTP | 0.877938106 | 1.27E-07 |
| PCIF1 | 0.516474576 | 6.84E-05 |
| GSS | 0.403471583 | 0.000111341 |
| MMP9 | 1.005630236 | 0.014649654 |
| TRPC4AP | -0.428095184 | 1.34E-05 |
| PYGB | 0.195977599 | 0.000243543 |
| ABHD12 | 0.239682012 | 0.011794301 |
| GINS1 | -1.759640426 | 1.41E-09 |
| NINL | 1.689805159 | 0.000113488 |
| CD40 | 1.258189781 | 0.000196488 |
| UQCC1 | -0.383572104 | 6.21E-05 |
| ZMYND8 | -0.26374543 | 0.000321484 |
| SGK2 | 8.412581001 | 2.09E-06 |
| IFT52 | 0.799253212 | 5.97E-05 |
| MYBL2 | -0.464622287 | 5.40E-06 |
| RAB5IF | -0.476908239 | 1.06E-06 |
| NFATC2 | -0.521160137 | 1.06E-05 |
| PABPC1L | -0.198729573 | 0.006219285 |
| STK4 | -0.965091558 | 1.34E-09 |
| ADNP | -1.225177167 | 6.76E-11 |
| PFDN4 | 0.267558552 | 0.010132158 |
| CSTF1 | -0.750248426 | 1.65E-07 |
| RAE1 | -0.811228433 | 2.08E-08 |
| TPD52L2 | -0.303939302 | 2.29E-05 |
| DNAJC5 | -1.170451894 | 3.61E-10 |
| NELFCD | -0.225855818 | 0.000624443 |
| CTSZ | 0.44248576 | 9.94E-07 |
| PRPF6 | -0.492229389 | 2.59E-06 |
| TUBB1 | 0.606436454 | 0.001773345 |
| PRELID3B | 0.16529326 | 0.0027536 |
| MTG2 | -0.159771988 | 0.024195841 |
| PSMA7 | 0.282658512 | 0.000913637 |
| SLCO4A1 | -0.461067229 | 0.000202572 |
| NTSR1 | 0.685986383 | 0.037793742 |
| MRGBP | -0.471608971 | 1.25E-05 |
| TCFL5 | 0.865910711 | 1.28E-07 |
| DIDO1 | -0.398288975 | 9.28E-06 |

| | | |
|---|---|---|
| GID8 | -0.325515384 | 4.16E-05 |
| ARFGAP1 | -0.597489075 | 1.95E-07 |
| COL20A1 | 4.554833343 | 1.04E-06 |
| EEF1A2 | 0.806007389 | 1.63E-05 |
| GMEB2 | -1.311931652 | 1.29E-08 |
| SPEF1 | 1.124326196 | 0.007229197 |
| CDC25B | -0.741063084 | 1.70E-08 |
| RNF24 | 0.180629336 | 0.003494069 |
| ARFRP1 | -0.467164224 | 0.00031645 |
| RASSF2 | 2.332203312 | 3.34E-07 |
| CSNK2A1 | -1.070850675 | 1.12E-09 |
| RSPO4 | 3.857866598 | 8.25E-05 |
| CDS2 | -1.433064509 | 1.88E-09 |
| SIRPB1 | -1.529870757 | 3.48E-07 |
| FERMT1 | -0.747293338 | 1.70E-07 |
| PLCB4 | -0.582126005 | 0.013311768 |
| CRNKL1 | -0.381042953 | 0.000108372 |
| POFUT1 | -0.446521674 | 5.79E-07 |
| SAMHD1 | -0.54419602 | 3.31E-06 |
| KIF3B | -1.881785384 | 6.46E-12 |
| MROH8 | 1.456021842 | 0.000228623 |
| NOP56 | -0.819556668 | 7.80E-10 |
| MANBAL | -0.401281561 | 0.000132315 |
| MAPRE1 | -0.845683293 | 2.38E-09 |
| JAG1 | 0.790263212 | 1.46E-07 |
| CDK5RAP1 | -0.29352913 | 0.006509263 |
| TTI1 | -0.734041053 | 1.24E-08 |
| E2F1 | -0.315461838 | 0.000193125 |
| RPRD1B | -1.544824713 | 1.30E-10 |
| CHMP4B | -0.136726579 | 0.049994082 |
| CST3 | 0.715709644 | 1.27E-07 |
| ACTR5 | -0.442707145 | 0.000778651 |
| WFDC2 | 2.951983263 | 0.001109667 |
| PPP1R16B | 1.251333331 | 0.014907233 |
| FAM83D | -1.62791439 | 4.36E-11 |
| DHX35 | -1.00204702 | 1.84E-06 |
| MAP1LC3A | 0.735832534 | 0.000284851 |
| PIGU | -0.325944622 | 0.025294334 |
| ACOT8 | 0.332925859 | 0.002084855 |
| APMAP | -0.387649956 | 5.10E-06 |
| ADNP2 | 0.120168849 | 0.039924488 |
| USP14 | -0.332230332 | 3.29E-05 |
| VAPA | -0.883766475 | 2.88E-08 |
| METTL4 | -2.119514416 | 1.00E-08 |
| LPIN2 | -0.814355218 | 6.19E-08 |
| SMCHD1 | -0.572081975 | 1.45E-06 |
| MYOM1 | 4.199737967 | 1.87E-05 |
| MYL12A | 0.229321293 | 0.00199467 |

| | | |
|---|---|---|
| CEP76 | -0.451812035 | 0.00387191 |
| CEP192 | 0.206523222 | 0.01532248 |
| RNMT | -0.85284247 | 1.10E-08 |
| SMAD7 | -0.530656524 | 0.001117876 |
| RNF125 | -0.842367125 | 4.30E-06 |
| MIB1 | -0.543438601 | 1.74E-06 |
| RBBP8 | -0.950809188 | 2.33E-08 |
| RIOK3 | -0.291607538 | 0.000384234 |
| CSTF2 | -0.417558241 | 4.86E-05 |
| PSMD10 | 0.41368181 | 9.20E-06 |
| TBL1X | -0.808854664 | 1.28E-07 |
| PGRMC1 | -0.122304371 | 0.037787139 |
| POLA1 | -0.340119438 | 8.23E-05 |
| MID1 | -0.992956235 | 2.33E-08 |
| MCTS2P | -1.24467883 | 3.40E-06 |
| ALG13 | -0.263607622 | 0.020577466 |
| PRPS2 | -0.654952912 | 1.55E-07 |
| TLR8 | 6.722959852 | 3.84E-05 |
| MOSPD1 | 0.337387492 | 0.001093913 |
| AMMECR1 | -3.02761286 | 1.38E-11 |
| CHRDL1 | 0.518376444 | 6.86E-05 |
| WDR13 | 0.519281488 | 9.13E-06 |
| SUV39H1 | -1.548211023 | 9.55E-09 |
| SRPX | 0.586101837 | 3.85E-07 |
| XIAP | -0.292129595 | 0.000318743 |
| STAG2 | -0.197252038 | 0.005491092 |
| ATP11C | -0.573442557 | 0.000117886 |
| ABCD1 | -0.553996189 | 1.42E-05 |
| CACNA1F | 5.439460706 | 2.75E-05 |
| SYP | 1.271310464 | 5.14E-05 |
| PLP2 | 0.226112609 | 0.001980304 |
| NAA10 | 0.264290796 | 0.000650299 |
| ELF4 | -0.88885313 | 3.30E-06 |
| SMARCA1 | 0.2663208 | 0.003248392 |
| RBBP7 | -0.189494976 | 0.002111268 |
| KCND1 | 1.143134903 | 0.000673729 |
| FMR1 | -0.471376282 | 6.72E-06 |
| PIM2 | 0.139380172 | 0.038958862 |
| SCML2 | -1.61320408 | 1.54E-09 |
| EMD | 0.38263455 | 9.61E-05 |
| RAB40AL | 1.376959482 | 0.032330118 |
| PGK1 | 0.084528253 | 0.031768368 |
| MAGT1 | -0.448832697 | 3.29E-06 |
| PHEX | 0.724421191 | 0.013065494 |
| UBL4A | -0.431606505 | 3.88E-06 |
| CD99L2 | -0.200885004 | 0.013248069 |
| EEA1 | -0.790543253 | 3.85E-07 |
| GPR50 | 7.085654925 | 1.26E-05 |

| | | |
|---|---|---|
| JADE3 | -0.597664565 | 5.67E-06 |
| USP11 | 0.639578322 | 2.45E-08 |
| TIMP1 | 1.21980563 | 2.83E-06 |
| GABRE | 1.409249985 | 0.00136186 |
| FGD1 | -0.40605374 | 0.002809476 |
| PIN4 | 0.409160615 | 0.010196278 |
| ITIH6 | 0.83173964 | 0.021428724 |
| MAGED2 | 0.382476108 | 6.44E-05 |
| RBM3 | -0.149304606 | 0.00580017 |
| SRPX2 | 0.499235287 | 0.020974248 |
| CENPI | -1.40198727 | 2.56E-06 |
| DRP2 | 3.341780235 | 1.85E-06 |
| PBDC1 | -1.120312944 | 1.17E-08 |
| GLA | -0.282068231 | 0.002005312 |
| ARMCX3 | -0.649713143 | 0.000202184 |
| BEX4 | 1.003344444 | 1.40E-05 |
| RUBCNL | -0.912269951 | 0.031012296 |
| NDFIP2 | -0.437836383 | 1.49E-05 |
| FNDC3A | -0.621545789 | 1.41E-07 |
| CAB39L | -0.865365599 | 0.007282961 |
| KLF5 | 1.320436752 | 1.73E-10 |
| STK24 | -0.386804684 | 1.63E-05 |
| DNAJC3 | -0.794287654 | 7.22E-09 |
| UGGT2 | 0.202650365 | 0.045627143 |
| ARHGEF7 | -1.726446759 | 3.94E-11 |
| PARP4 | -0.42728327 | 1.37E-05 |
| SUPT20H | 0.462178677 | 7.85E-06 |
| SLC25A15 | -0.557875285 | 2.17E-05 |
| KPNA3 | -1.430551911 | 6.51E-11 |
| VWA8 | -0.272008272 | 0.004695851 |
| DGKH | -1.269769955 | 5.16E-09 |
| KATNAL1 | -1.807506571 | 3.19E-09 |
| INTS6 | -2.301847604 | 1.69E-11 |
| DHRS12 | 1.804431538 | 8.87E-05 |
| TSC22D1 | 0.75314021 | 7.21E-09 |
| CLN5 | -0.484021895 | 2.49E-05 |
| MSLN | 0.671681759 | 0.030811984 |
| MGRN1 | -0.302168977 | 0.000355717 |
| ZNF629 | 0.46851581 | 0.000276252 |
| TRADD | 0.344378916 | 0.049238365 |
| HSF4 | 0.860447575 | 0.000397899 |
| CORO1A | 0.433214637 | 0.002154308 |
| MAPK3 | 0.279299876 | 0.004436017 |
| GDPD3 | 1.32726054 | 0.004809434 |
| LYRM1 | 0.948021568 | 4.26E-08 |
| NUTF2 | 0.45358466 | 3.43E-06 |
| NUP93 | -0.159395102 | 0.038750466 |
| CENPT | -0.546242633 | 5.98E-06 |

| | | |
|---|---|---|
| TSNAXIP1 | 1.591413623 | 1.82E-05 |
| NFAT5 | -0.726655758 | 1.38E-08 |
| LONP2 | -0.626141578 | 1.34E-07 |
| ARL2BP | -0.403078363 | 1.68E-05 |
| PLLP | -0.383716043 | 2.20E-05 |
| DHODH | 0.247793564 | 0.02290562 |
| CTCF | -1.00860709 | 2.00E-09 |
| ACD | 0.550796282 | 3.46E-06 |
| POLR2C | -0.231746248 | 0.000421585 |
| PARD6A | 1.619480195 | 4.13E-06 |
| USB1 | -0.159422414 | 0.023525291 |
| CYB5B | -0.287410802 | 0.000140312 |
| CCDC113 | 3.258832085 | 1.02E-09 |
| PRSS54 | 2.409760435 | 1.47E-05 |
| NME3 | 0.698124918 | 0.000177077 |
| PSMD7 | -0.254971767 | 0.000189713 |
| SETD6 | -0.648839784 | 3.39E-07 |
| SLC38A7 | -0.604742339 | 0.000186895 |
| VAC14 | -0.190425915 | 0.009890389 |
| HAS3 | -0.726023279 | 1.59E-06 |
| TANGO6 | -0.766759593 | 9.48E-06 |
| COG4 | 0.320055145 | 0.000311414 |
| SMPD3 | 1.20708505 | 0.00031823 |
| SLC7A6OS | -1.06546046 | 3.31E-09 |
| SLC7A6 | -1.182730573 | 3.06E-09 |
| PLA2G15 | 0.558908769 | 5.69E-06 |
| ESRP2 | 2.707843514 | 3.70E-06 |
| MON1B | -0.601580383 | 3.46E-07 |
| CMC2 | -0.229116562 | 0.015705624 |
| HCFC1R1 | 0.532965368 | 1.95E-05 |
| MPG | 0.581972364 | 9.34E-06 |
| NECAB2 | 3.102288428 | 1.00E-04 |
| HSDL1 | -0.577157177 | 9.16E-07 |
| TAF1C | -0.480192853 | 2.69E-06 |
| NAGPA | -0.929900632 | 5.84E-05 |
| SEC14L5 | 2.920874333 | 0.000207544 |
| COTL1 | 0.190813028 | 9.38E-05 |
| USP10 | -0.906073645 | 2.52E-09 |
| CRISPLD2 | 0.475826623 | 0.006936321 |
| NME4 | 0.28799234 | 0.000240365 |
| NOMO3 | 0.21521253 | 0.000983953 |
| LMF1 | 0.797667698 | 0.00338271 |
| FOXF1 | 0.724968193 | 0.019414367 |
| CIAO3 | -0.228596487 | 0.038137773 |
| MTHFSD | -0.231554226 | 0.02093875 |
| CLCN7 | -0.225100419 | 0.010404023 |
| HAGHL | 0.464526209 | 0.004819709 |
| ANTKMT | 0.286559239 | 0.034852227 |

| | | |
|---|---|---|
| SLC7A5 | -1.329052564 | 6.26E-12 |
| METRN | 0.291471832 | 0.000714257 |
| FBXO31 | -0.234669219 | 0.004023301 |
| STUB1 | 0.270252195 | 0.000182832 |
| RHBDL1 | 0.542090043 | 0.006877681 |
| NUBP1 | -0.288057308 | 0.026202785 |
| UBE2I | -0.305176795 | 5.50E-05 |
| MEFV | 9.124947972 | 1.11E-08 |
| EEF2K | -0.874624302 | 2.15E-08 |
| CAPN15 | -0.913398494 | 1.87E-08 |
| PIEZO1 | -0.415504867 | 2.34E-06 |
| GSPT1 | -0.269568564 | 4.68E-05 |
| ZNF174 | 0.538211653 | 3.84E-05 |
| CLUAP1 | -0.524897739 | 0.000105187 |
| UBFD1 | -0.361329772 | 6.24E-06 |
| PRSS33 | 0.835302379 | 0.00010077 |
| EARS2 | -0.239030066 | 0.000991488 |
| ELOB | 0.327900238 | 0.000459268 |
| GGA2 | -0.90883009 | 8.13E-09 |
| USP31 | -1.483712919 | 3.94E-09 |
| HMOX2 | -0.380608607 | 0.00055757 |
| DNAJA3 | -0.485788996 | 5.21E-06 |
| BFAR | -0.179316347 | 0.003450941 |
| SALL1 | -1.081033869 | 2.57E-08 |
| QPRT | 1.188432935 | 0.001431712 |
| XYLT1 | -0.406949856 | 0.000421873 |
| PYCARD | 2.906786642 | 0.000120446 |
| RPGRIP1L | 0.456000721 | 0.001139796 |
| MAZ | -0.443223913 | 1.76E-06 |
| STX4 | 0.372896521 | 0.000697383 |
| BCKDK | 0.370821495 | 0.000117075 |
| NOMO1 | 0.176659478 | 0.008698812 |
| IL21R | 3.630668334 | 1.18E-08 |
| SYT17 | 2.062055872 | 0.000395479 |
| TMC5 | 1.691811468 | 0.016302513 |
| CCP110 | -0.217445465 | 0.000563295 |
| VPS35L | -0.705182996 | 2.68E-06 |
| RNF40 | -0.316808451 | 1.43E-05 |
| KNOP1 | -1.166151863 | 1.42E-09 |
| AAGAB | -0.612523249 | 4.59E-07 |
| IQCH | 0.716448729 | 0.019160949 |
| LACTB | 0.319760142 | 0.012874813 |
| CSK | -0.609356386 | 1.19E-07 |
| HERC1 | -0.336769475 | 5.30E-05 |
| MTFMT | 0.637524148 | 0.00023923 |
| ACSBG1 | -0.511928899 | 2.12E-06 |
| FAH | -0.956002894 | 9.06E-05 |
| CEMIP | 0.641676778 | 1.19E-07 |

| | | |
|---|---|---|
| RPAP1 | 0.13520321 | 0.042161469 |
| HOMER2 | 0.480155257 | 0.000686357 |
| EHD4 | -0.681958497 | 3.07E-08 |
| TMEM87A | -0.641626173 | 3.74E-07 |
| ZNF106 | -0.39735613 | 2.38E-06 |
| CEP152 | -1.33815341 | 1.72E-09 |
| ATP8B4 | 4.691772447 | 0.000602259 |
| OCA2 | 1.025408425 | 0.028914468 |
| DTWD1 | 0.536582546 | 6.44E-05 |
| TJP1 | -0.550523981 | 2.09E-06 |
| DMXL2 | -0.898467293 | 1.11E-08 |
| DNAJC17 | 0.334690282 | 0.048800037 |
| EIF3J | -1.266272664 | 7.31E-11 |
| VPS18 | -0.793697254 | 8.46E-08 |
| OIP5 | -0.810025529 | 9.63E-06 |
| SLC30A4 | 1.089722655 | 1.33E-06 |
| BLOC1S6 | -0.836283081 | 2.66E-08 |
| MYEF2 | -0.334406396 | 0.015898657 |
| CSPP1 | 0.69670569 | 2.06E-06 |
| TRIM35 | -0.733252774 | 3.67E-06 |
| RP1 | 3.872690442 | 0.004897431 |
| FZD3 | -1.673078785 | 5.13E-09 |
| INTS9 | -0.348428819 | 0.034744803 |
| RIPK2 | -0.664307824 | 4.57E-05 |
| EYA1 | -1.118098351 | 5.54E-07 |
| NBN | -0.885902469 | 6.78E-09 |
| DECR1 | 0.509211418 | 0.018356333 |
| CALB1 | 1.896980637 | 0.000143857 |
| IMPAD1 | -0.762521007 | 3.06E-09 |
| SFRP1 | 0.141084835 | 0.003048582 |
| UBE2W | -0.544225127 | 1.46E-06 |
| POP1 | -1.186569699 | 2.28E-09 |
| NIPAL2 | -0.565172097 | 0.000933112 |
| IKBKB | -0.633863289 | 3.86E-05 |
| PLAT | 0.907238032 | 4.97E-09 |
| JPH1 | 0.621243408 | 7.69E-06 |
| STK3 | -1.272488505 | 3.33E-09 |
| GDAP1 | -0.163310459 | 0.0326506 |
| RAB2A | -0.513577553 | 7.09E-07 |
| EIF3E | 0.222784108 | 0.002704713 |
| NDRG1 | 0.228098556 | 0.001451881 |
| ZC2HC1A | -1.666557574 | 5.50E-08 |
| IL7 | -1.031576355 | 8.50E-05 |
| ARMC1 | -0.796462775 | 2.69E-08 |
| TRPS1 | -0.963415524 | 1.73E-07 |
| SPAG1 | -0.400955495 | 0.000541143 |
| CHRAC1 | -0.483667197 | 1.33E-05 |
| UBR5 | -0.585558105 | 2.13E-07 |

| | | |
|---|---|---|
| GSDMD | 0.358895843 | 0.012476157 |
| SQLE | 0.63932192 | 1.21E-07 |
| SH2D4A | -1.862198705 | 1.43E-08 |
| INTS10 | -0.218570205 | 0.017073717 |
| ERI1 | -0.841632272 | 4.04E-07 |
| SLC39A14 | -0.357228143 | 6.95E-06 |
| MTMR9 | -1.114904047 | 8.10E-09 |
| LEPROTL1 | -1.145891603 | 1.52E-09 |
| DCTN6 | -0.303345653 | 0.030432684 |
| R3HCC1 | -0.975703539 | 3.68E-06 |
| GSR | -0.485984167 | 4.99E-05 |
| TNFRSF10A | -0.461390576 | 8.40E-05 |
| UBXN8 | -0.389450951 | 0.00665134 |
| PPP2CB | -0.629897559 | 1.33E-06 |
| ERICH1 | -0.811790705 | 3.42E-05 |
| NEFM | -2.027079608 | 0.004076895 |
| KLHDC4 | -0.2476451 | 0.011634437 |
| MCM4 | -0.593310626 | 2.16E-08 |
| KCTD9 | -0.766862297 | 1.88E-07 |
| FGL1 | 8.301460104 | 1.94E-07 |
| ASAH1 | 0.300743312 | 0.001013188 |
| BNIP3L | 0.149576909 | 0.035701906 |
| MAN2B1 | 0.344561951 | 0.00016421 |
| KCNN4 | 0.58396268 | 1.36E-05 |
| NUCB1 | 0.324204388 | 0.000449984 |
| DHDH | 3.874381431 | 0.005538334 |
| MAP4K1 | 2.368470033 | 0.000504939 |
| ECH1 | 0.541174898 | 2.88E-06 |
| HNRNPL | 0.374568063 | 9.78E-06 |
| NFKBIB | 0.567158266 | 8.06E-05 |
| TUBB4A | 0.899350707 | 1.83E-05 |
| SNRNP70 | -0.513696117 | 1.10E-06 |
| CLPTM1 | -0.378579184 | 1.53E-05 |
| RELB | 1.989433436 | 1.27E-09 |
| LIN7B | 0.868646491 | 0.004922 |
| PPP1R37 | -0.203692198 | 0.048341075 |
| CKM | 4.146158147 | 0.003806858 |
| ARHGEF18 | -0.805885722 | 1.07E-07 |
| PPP1R13L | 2.27103769 | 1.93E-08 |
| ERCC2 | -0.589467201 | 2.89E-06 |
| DOT1L | -0.786346228 | 3.08E-07 |
| PLEKHJ1 | -0.27646027 | 0.008186475 |
| SLC17A7 | 1.551111372 | 0.000193209 |
| RNASEH2A | 0.475427358 | 4.84E-06 |
| KLC3 | 2.070370833 | 0.001146792 |
| CD37 | 1.262318721 | 0.000374059 |
| SF3A2 | -0.339668572 | 0.000792817 |
| AMH | 0.257574491 | 0.019421251 |

| | | |
|---|---|---|
| LYL1 | -0.958733162 | 0.005228411 |
| OAZ1 | 0.469352392 | 2.91E-07 |
| TRMT1 | -0.164118049 | 0.027458279 |
| DMPK | -0.938901499 | 5.15E-07 |
| RSPH6A | 4.422077965 | 0.000917626 |
| TBC1D17 | 0.925582146 | 5.37E-06 |
| IL4I1 | -1.344470125 | 0.000771975 |
| TLE6 | 1.444138539 | 0.011584797 |
| CCDC130 | 0.76793075 | 5.14E-07 |
| TLE5 | 0.392089302 | 2.29E-05 |
| SGTA | -0.357927145 | 6.48E-05 |
| MED25 | -0.162208328 | 0.046688285 |
| SNAPC2 | 0.392298513 | 9.35E-05 |
| CCDC61 | 0.907854803 | 0.002145612 |
| ASF1B | -0.18063136 | 0.004051216 |
| TNNT1 | 0.698293789 | 0.011862103 |
| FAM32A | -0.470764644 | 6.97E-07 |
| PPP6R1 | -0.464018634 | 2.22E-06 |
| C19orf44 | -0.579878509 | 5.84E-07 |
| MED26 | 0.702295051 | 3.61E-07 |
| OLFM2 | 2.888666508 | 5.80E-05 |
| AKAP8 | -2.213870718 | 3.51E-11 |
| EPHX3 | 0.393068558 | 0.04133572 |
| SYDE1 | -1.334989899 | 6.96E-10 |
| CASP14 | 7.043972366 | 1.39E-05 |
| URI1 | -0.138303352 | 0.023618435 |
| RPS16 | 0.367749994 | 3.67E-06 |
| TIMM50 | -0.296805318 | 0.000372109 |
| DYRK1B | 1.011279827 | 2.13E-05 |
| CNTD2 | 3.894307744 | 0.001168911 |
| AKT2 | -0.159059246 | 0.015877665 |
| PLD3 | 1.108528992 | 1.86E-08 |
| PRX | 0.603589345 | 0.011408407 |
| PIAS4 | -0.785133109 | 2.59E-05 |
| YJU2 | -0.550868653 | 0.003319121 |
| TBCB | 0.236366469 | 0.005663363 |
| FSD1 | 0.248908692 | 0.011888444 |
| OVOL3 | 3.970248563 | 2.38E-05 |
| CLIP3 | 0.444209165 | 0.000871867 |
| ZFR2 | 8.765756193 | 3.56E-08 |
| SLC1A5 | -1.173678128 | 1.05E-10 |
| PRKD2 | 0.514680279 | 2.53E-05 |
| APLP1 | 0.830346617 | 0.003100074 |
| CACTIN | -0.277275211 | 0.002416289 |
| CCDC9 | -0.916283034 | 6.80E-05 |
| HNRNPUL1 | -0.136997926 | 0.003695115 |
| FZR1 | -1.039814742 | 2.50E-08 |
| BBC3 | 3.413136644 | 3.86E-12 |

| | | |
|---|---|---|
| TGFB1 | 0.144560613 | 0.025730911 |
| DENND3 | 0.817808196 | 8.19E-05 |
| DMAC2 | -0.249202196 | 0.001929602 |
| PLIN3 | 0.194763251 | 0.008596765 |
| MYH14 | 0.485890941 | 0.000299975 |
| MRPL4 | -0.368090491 | 5.74E-05 |
| CD79A | 4.528922192 | 0.00177281 |
| RPS19 | 0.445215468 | 2.09E-07 |
| NOP53 | 0.219365784 | 0.013726054 |
| NKG7 | 1.839337819 | 0.003872978 |
| ICAM5 | 1.213739131 | 1.75E-07 |
| ETFB | 0.455439341 | 0.000123014 |
| TYK2 | -0.146867047 | 0.026835296 |
| CDC37 | 0.174234988 | 0.000867883 |
| RABAC1 | 0.548618056 | 0.000482811 |
| MEIS3 | 0.624015444 | 0.012542209 |
| PTPRS | 0.335442785 | 7.19E-05 |
| CNFN | 0.981069658 | 0.04320227 |
| KDELR1 | 0.15862165 | 0.00958726 |
| CYTH2 | 0.737363151 | 4.93E-07 |
| GRWD1 | -0.824614246 | 1.38E-08 |
| CCDC114 | 3.189778138 | 0.01873054 |
| CARD8 | -0.757150933 | 9.06E-06 |
| LIG1 | -0.325558423 | 0.001851955 |
| ZNF175 | 0.916355023 | 6.52E-08 |
| PLA2G4C | 4.715598685 | 0.00042424 |
| HAS1 | 3.090616187 | 0.000718678 |
| RAB3D | 0.312723071 | 0.030472647 |
| DBP | 1.302982528 | 2.63E-05 |
| TMEM205 | 0.558260675 | 0.034030008 |
| CAPS | 0.543128323 | 1.34E-06 |
| FAM83E | 3.729847137 | 0.006672996 |
| RASIP1 | 1.906750794 | 0.006888 |
| BCAT2 | -0.151028333 | 0.039675517 |
| MIER2 | -0.977238616 | 8.02E-07 |
| TNPO2 | -0.770184224 | 1.08E-08 |
| WDR83OS | 0.184006602 | 0.011560156 |
| CACNG7 | 1.479507229 | 0.006919595 |
| GCDH | 0.369820881 | 0.000561111 |
| DNASE2 | 0.304526434 | 0.001501088 |
| MAST1 | 2.662542553 | 1.29E-05 |
| JAK3 | 1.054632117 | 0.032463416 |
| ARRDC2 | 0.158854431 | 0.049668547 |
| RAB3A | 1.473628664 | 2.22E-07 |
| PDE4C | 6.629880063 | 0.031817537 |
| ISYNA1 | 1.103556574 | 1.52E-08 |
| CRTC1 | 0.349065403 | 0.001360226 |
| UPK1A | 5.65488602 | 2.40E-08 |

| | | |
|---|---|---|
| COPE | 0.442021868 | 5.15E-06 |
| DDX49 | 0.413027256 | 4.85E-06 |
| ARMC6 | -0.70618268 | 1.19E-07 |
| TMEM147 | 0.269883767 | 0.000281357 |
| GAPDHS | 2.361977491 | 0.013713363 |
| LSR | 0.529667767 | 0.049127808 |
| KXD1 | 0.123201069 | 0.032330118 |
| FKBP8 | 0.367579466 | 9.75E-05 |
| ZNF14 | 1.8144459 | 1.84E-10 |
| PBX4 | 1.12422794 | 0.000899936 |
| GSK3A | -0.590297303 | 1.56E-06 |
| ATP13A1 | -0.198578675 | 0.001575892 |
| ZNF574 | 0.552437624 | 4.92E-06 |
| GRIK5 | 3.705208315 | 3.25E-06 |
| SIPA1L3 | -0.57563299 | 3.68E-06 |
| ZNF85 | 0.354335266 | 0.00271647 |
| ETHE1 | 0.734528177 | 1.96E-05 |
| SMG9 | -0.464792007 | 2.40E-05 |
| AVL9 | -1.334466045 | 3.62E-10 |
| RUNDC3B | -2.09041501 | 0.000402469 |
| CFAP69 | 1.247523442 | 0.000200775 |
| GTPBP10 | -0.664676161 | 2.44E-06 |
| RASA4 | 2.281800782 | 6.92E-06 |
| CDK6 | -1.318599779 | 3.76E-11 |
| PMPCB | 0.151867078 | 0.016641128 |
| DNAJC2 | -0.7015094 | 9.54E-07 |
| TFPI2 | 1.47017499 | 7.86E-11 |
| TWISTNB | -0.23484502 | 0.000393406 |
| PON2 | -0.555193475 | 1.48E-06 |
| ITGB8 | -0.982942713 | 1.93E-08 |
| HBP1 | 1.962118912 | 4.86E-11 |
| DUS4L | 1.078979905 | 1.14E-06 |
| SP4 | -0.669303884 | 2.59E-07 |
| DNAH11 | -0.665407064 | 6.85E-06 |
| CBLL1 | -0.601486987 | 1.21E-06 |
| MTPN | -0.116532028 | 0.002998683 |
| STEAP1B | -0.437899944 | 0.001810793 |
| PTN | 0.428630161 | 0.036466782 |
| MPP6 | -1.700812803 | 2.06E-10 |
| GSDME | 0.970187341 | 7.41E-07 |
| ZC3HAV1 | 0.218921182 | 0.000553889 |
| TTC26 | -1.174661133 | 2.86E-07 |
| ADAP1 | 1.276559358 | 0.001452274 |
| H2AZ2 | -0.199498281 | 0.001880918 |
| CAV2 | -0.726369181 | 2.37E-05 |
| MET | -0.489643445 | 4.32E-07 |
| RNF32 | 0.44387302 | 0.039434389 |
| LMBR1 | -1.087407134 | 2.77E-08 |

| | | |
|---|---|---|
| HOXA1 | 0.763029984 | 4.84E-05 |
| DNAJB6 | 0.401735965 | 1.13E-06 |
| HOXA2 | -1.704851353 | 0.000139696 |
| HOXA3 | -2.416620906 | 1.11E-08 |
| HOXA5 | -1.422336371 | 1.90E-06 |
| HOXA6 | -1.507239355 | 0.00306898 |
| BRAT1 | -0.553866491 | 3.11E-06 |
| IQCE | -0.480489196 | 8.48E-05 |
| HOXA13 | -1.567684447 | 2.53E-05 |
| TAX1BP1 | 0.258207768 | 0.000780782 |
| CPVL | 0.554728971 | 0.017444651 |
| GRB10 | -0.672974905 | 6.01E-05 |
| ABHD11 | 0.409213918 | 0.007371392 |
| COBL | -0.527204179 | 0.000748473 |
| FKBP14 | -0.728548521 | 2.07E-07 |
| PLEKHA8 | -0.776615849 | 7.41E-08 |
| STX1A | -0.489900354 | 0.000140528 |
| NOD1 | -0.820911375 | 4.43E-05 |
| GARS1 | -0.32229885 | 3.69E-05 |
| CRHR2 | 1.765165462 | 0.003308549 |
| NSUN5P2 | -0.486727449 | 0.000171082 |
| CASP2 | -1.95070153 | 3.55E-11 |
| CHCHD2 | 0.35067459 | 5.86E-05 |
| HSPB1 | 0.604100265 | 1.01E-07 |
| ZKSCAN1 | 1.458718959 | 2.39E-11 |
| EIF3B | -0.46704753 | 3.68E-07 |
| SNX8 | -1.122852791 | 1.70E-08 |
| PTPRZ1 | 0.643211139 | 0.006392712 |
| TAF6 | 0.406136157 | 2.74E-05 |
| WASL | -2.012599793 | 1.39E-11 |
| AIMP2 | -0.291108757 | 0.005315161 |
| FSCN3 | 0.850835102 | 0.000315397 |
| PAX4 | 7.270706977 | 6.81E-06 |
| PCOLCE | 0.454422 | 6.31E-05 |
| FBXO24 | 2.981527939 | 3.13E-05 |
| RBM28 | -0.753135581 | 8.88E-08 |
| IMPDH1 | -0.642577009 | 2.72E-07 |
| SERPINE1 | 4.636714214 | 6.82E-12 |
| AP1S1 | 0.246209924 | 0.001515592 |
| C1GALT1 | -1.054122793 | 8.17E-08 |
| PLOD3 | 0.169876663 | 0.001758426 |
| RPA3 | 0.657787945 | 1.03E-05 |
| CLDN15 | 0.704229926 | 0.000104062 |
| GLCCI1 | -2.54691667 | 1.26E-09 |
| NRF1 | -0.755969738 | 7.13E-06 |
| TMEM106B | 0.185302597 | 0.027140081 |
| EZH2 | -0.202165378 | 0.011475381 |
| CEP41 | -0.722806705 | 2.14E-05 |

| | | |
|---|---|---|
| ZNF862 | 3.80221414 | 2.12E-11 |
| MEOX2 | -2.558775802 | 8.83E-11 |
| ANKMY2 | -0.420570653 | 0.00379938 |
| POU6F2 | 2.647686457 | 0.00210662 |
| AC004837.1 | 6.000931793 | 5.97E-06 |
| AHR | 0.397550304 | 6.64E-06 |
| CHCHD3 | 0.281137683 | 3.77E-05 |
| GLI3 | -0.452912737 | 0.000303509 |
| COA1 | -0.337528754 | 3.73E-05 |
| BLVRA | 0.682187496 | 0.000193829 |
| URGCP | 0.168713933 | 0.037966904 |
| TMEM248 | -0.796675322 | 1.76E-08 |
| RHEB | -0.148756713 | 0.020312308 |
| PRKAG2 | -1.583039189 | 4.07E-07 |
| AEBP1 | 0.432629413 | 4.21E-06 |
| POLD2 | 0.218473517 | 0.000880441 |
| GCK | 2.347384059 | 0.000780672 |
| BCL7B | 0.174580292 | 0.019627685 |
| YKT6 | -0.444455671 | 1.13E-06 |
| CLIP2 | 0.305344428 | 0.000367057 |
| EIF4H | -0.126458544 | 0.002147958 |
| LIMK1 | -0.399704734 | 0.000634107 |
| SPATA6L | 2.535096083 | 3.28E-06 |
| SLC1A1 | -0.514543235 | 1.44E-05 |
| FKTN | -1.292503162 | 6.13E-10 |
| FSD1L | 0.312605603 | 0.005279198 |
| CNTNAP3 | -0.302529404 | 0.018294181 |
| SPIN1 | -1.315358816 | 1.28E-10 |
| NMRK1 | 2.733285743 | 2.85E-08 |
| TMEM245 | -0.812584856 | 3.00E-08 |
| PRUNE2 | 1.844630811 | 4.09E-05 |
| MEGF9 | -0.307134566 | 0.000277981 |
| TRIM14 | -0.628783179 | 1.52E-06 |
| CORO2A | 0.294633694 | 0.010346746 |
| C5 | 3.749349744 | 0.000373626 |
| ECM2 | 2.781330867 | 1.45E-06 |
| TLE4 | -1.463067684 | 2.04E-09 |
| LHX6 | -0.786591998 | 0.005054652 |
| PTGR1 | 0.364980033 | 0.018011601 |
| DNM1 | 0.196389044 | 0.00726464 |
| ENG | -0.346995795 | 3.73E-05 |
| AK1 | 0.213131619 | 0.0273769 |
| CDC37L1 | -1.093918892 | 5.56E-08 |
| PLGRKT | 0.66172645 | 0.003936831 |
| TBC1D13 | -0.990340282 | 1.44E-08 |
| RIC1 | -1.462772396 | 8.42E-10 |
| KDM4C | -0.721742768 | 1.85E-06 |
| DOCK8 | -2.456736153 | 0.00719652 |

| | | |
|---|---|---|
| KANK1 | -1.755151579 | 6.83E-11 |
| NCS1 | -0.563815287 | 5.95E-07 |
| FUBP3 | -0.716751237 | 1.15E-07 |
| CREB3 | 0.301788611 | 0.000875596 |
| RGP1 | -0.523807537 | 3.68E-07 |
| MPDZ | -0.782737902 | 2.87E-07 |
| LHX3 | 5.637440054 | 1.59E-05 |
| EDF1 | 0.309944777 | 0.000195676 |
| GLIS3 | 0.274098531 | 0.00250454 |
| BAG1 | -0.210110897 | 0.037360729 |
| RAPGEF1 | -0.597762974 | 8.12E-06 |
| NPDC1 | 0.442459687 | 0.000303293 |
| SETX | 0.167897585 | 0.006567448 |
| ABCA2 | 0.703455226 | 2.21E-07 |
| SHB | -1.068453604 | 2.47E-06 |
| UBE2R2 | -0.995942202 | 5.01E-09 |
| ABHD17B | -0.254725766 | 0.001245668 |
| EXOSC3 | -1.198207297 | 1.53E-09 |
| ZFAND5 | -0.66686293 | 1.52E-08 |
| CCNJ | -0.752237736 | 1.18E-07 |
| GATA3 | 1.468922887 | 1.56E-06 |
| HPS1 | 0.37149338 | 0.002929014 |
| PHYH | 0.714542197 | 0.000476927 |
| DNMBP | -0.279863819 | 8.24E-05 |
| RAB11FIP2 | -1.812958982 | 7.09E-11 |
| ERLIN1 | -0.577663199 | 4.36E-07 |
| EIF3A | -0.148938737 | 0.001220094 |
| PKD2L1 | 8.740443859 | 3.85E-08 |
| CUBN | 1.530870503 | 0.003695115 |
| TRDMT1 | -1.289206199 | 3.53E-06 |
| DDX50 | -0.21537886 | 0.010802036 |
| MAPK8 | -0.705047831 | 1.35E-06 |
| SEC23IP | -1.494521083 | 1.19E-10 |
| ATE1 | -0.558537226 | 8.01E-07 |
| NSMCE4A | -0.610371488 | 8.58E-05 |
| PLEKHA1 | -0.381407456 | 0.000102459 |
| PALD1 | 0.536307216 | 0.017439738 |
| UNC5B | -0.560659775 | 0.004315461 |
| CDH23 | 4.51922711 | 3.45E-05 |
| MICU1 | -0.182443298 | 0.047063866 |
| CCSER2 | 0.278184374 | 0.000247946 |
| BMPR1A | -1.030228163 | 9.65E-08 |
| ACTA2 | 2.385034805 | 3.50E-08 |
| LIPA | 0.573951512 | 1.42E-06 |
| TWNK | -0.399148345 | 1.05E-05 |
| LZTS2 | -0.248204446 | 0.000734481 |
| KAZALD1 | 1.601783306 | 6.22E-08 |
| FBXW4 | 0.545427533 | 0.002543142 |

| | | |
|---|---|---|
| TNKS2 | -0.700788724 | 3.23E-08 |
| GBF1 | -0.259123885 | 0.001548691 |
| ARHGAP21 | -0.64655864 | 3.12E-07 |
| FBXL15 | 0.581020267 | 0.004912796 |
| CUEDC2 | 0.520629887 | 1.99E-05 |
| SUFU | -0.857500503 | 1.97E-07 |
| ACBD5 | -0.951769601 | 2.38E-08 |
| LHPP | 1.091237188 | 0.00200374 |
| LARP4B | -0.585709986 | 9.02E-07 |
| GTPBP4 | -0.665429649 | 1.61E-07 |
| EDRF1 | -1.472874663 | 5.55E-10 |
| BCCIP | -0.350420503 | 2.90E-05 |
| MTPAP | -0.808883496 | 9.03E-07 |
| NEURL1 | 1.354323668 | 0.003287417 |
| SH3PXD2A | -0.945213542 | 5.41E-10 |
| STN1 | 0.584618182 | 0.003242321 |
| DKK1 | 1.584750001 | 1.11E-11 |
| EBF3 | -0.748341164 | 2.67E-07 |
| SORCS1 | 0.340388553 | 9.09E-05 |
| TASOR2 | -0.869274163 | 8.48E-10 |
| SMC3 | -0.373858386 | 5.18E-06 |
| TFAM | -0.790052922 | 8.35E-09 |
| CCDC6 | -0.376232441 | 2.19E-05 |
| CUL2 | -0.43649439 | 7.15E-05 |
| CCNY | -0.927573281 | 1.85E-08 |
| UBE2S | -0.508683534 | 3.92E-06 |
| RPL28 | 0.644857016 | 1.35E-08 |
| ZMIZ1 | -0.713047678 | 3.81E-08 |
| PPIF | -1.212549634 | 1.95E-10 |
| TSPAN14 | -0.532023662 | 3.33E-07 |
| TBC1D12 | -0.898051064 | 1.80E-06 |
| NUFIP2 | -0.655408726 | 2.97E-09 |
| RPL19 | 0.451777425 | 1.34E-07 |
| FBXL20 | 0.256530107 | 0.006755749 |
| UBTF | -0.947557098 | 1.96E-09 |
| CASC3 | -0.639314893 | 4.82E-08 |
| RAPGEFL1 | 1.069661124 | 3.35E-05 |
| RNF43 | 2.937084324 | 0.000370932 |
| RAD51C | -0.201224965 | 0.033208184 |
| SEPTIN4 | 3.135357518 | 1.03E-06 |
| MTMR4 | 0.144939448 | 0.013613983 |
| TRIM37 | -0.989847773 | 4.54E-08 |
| DHX40 | -0.178521358 | 0.009954033 |
| KPNB1 | -0.241589485 | 1.25E-05 |
| GOSR2 | -1.0921875 | 1.17E-08 |
| PNPO | 0.519709628 | 2.76E-05 |
| RPS6KB1 | -0.276156123 | 0.001787546 |
| TRIM16L | -1.073756178 | 1.27E-06 |

| | | |
|---|---|---|
| CDK5RAP3 | 0.337455907 | 0.000166573 |
| CBX1 | -0.249550168 | 0.000158403 |
| RECQL5 | 0.445504422 | 0.000245786 |
| PIGL | 1.255064203 | 6.96E-07 |
| GALK1 | 0.463644263 | 0.012125111 |
| INTS2 | -0.50204627 | 6.01E-06 |
| CAMTA2 | -0.570225481 | 2.04E-05 |
| MED13 | -0.737301633 | 1.35E-08 |
| HOXB6 | -0.897187254 | 3.49E-05 |
| ENO3 | 0.573014596 | 0.002093626 |
| PFN1 | 0.274706473 | 4.33E-06 |
| RNF167 | -0.166955994 | 0.008582334 |
| SLC25A11 | 0.174586664 | 0.010931545 |
| RASD1 | 3.703675457 | 2.54E-08 |
| RAI1 | -1.054860404 | 4.58E-09 |
| NUP88 | -0.281309611 | 0.000504416 |
| SLC6A4 | 6.537521726 | 8.59E-10 |
| BLMH | -0.658456863 | 1.13E-07 |
| GOSR1 | -1.028029318 | 1.11E-08 |
| CCDC47 | -0.515973986 | 3.70E-07 |
| MED31 | 0.644897708 | 7.64E-06 |
| DRG2 | -0.38200269 | 0.000401406 |
| FTSJ3 | -0.860087684 | 8.12E-10 |
| AKAP10 | -0.537468113 | 5.51E-06 |
| SMARCD2 | -0.407405417 | 1.42E-05 |
| SYNGR2 | 0.328364399 | 0.000150235 |
| B9D1 | 0.862970176 | 0.000112809 |
| UTP6 | -0.453048611 | 1.23E-05 |
| DDX5 | -0.50641446 | 8.14E-09 |
| CYTH1 | -0.609188474 | 5.86E-06 |
| PSMD11 | -0.619637794 | 3.93E-08 |
| PEX12 | 1.065582476 | 9.47E-09 |
| DHX58 | 1.294472084 | 0.000123362 |
| KAT2A | -0.878493152 | 6.22E-08 |
| NAGLU | 0.561230041 | 6.48E-05 |
| HSD17B1P1 | 1.522780571 | 7.00E-05 |
| HSD17B1 | 0.254154845 | 0.034754975 |
| MLX | -0.658124092 | 8.69E-08 |
| CNTNAP1 | 0.383521792 | 6.60E-05 |
| EZH1 | -0.673536333 | 3.11E-06 |
| PPP1R9B | -0.800145249 | 2.35E-08 |
| COL1A1 | 1.224869301 | 0.00276894 |
| PTGES3L-AARSD1 | 0.43750638 | 0.006027834 |
| MRPL27 | -0.56827426 | 4.30E-06 |
| VAT1 | 0.296197684 | 4.22E-05 |
| LRRC59 | -0.425786799 | 2.42E-07 |
| RND2 | 2.688159897 | 0.000393928 |
| ALOX12 | 1.062839146 | 0.033110175 |

| | | |
|---|---|---|
| HDAC5 | 0.830546726 | 1.49E-07 |
| ABCC3 | 1.0254177 | 0.001160714 |
| LUC7L3 | -0.69963577 | 1.61E-08 |
| SMURF2 | -1.743591857 | 4.61E-10 |
| DUSP3 | -0.731347996 | 2.48E-07 |
| SLC16A6 | 0.50039806 | 0.022935266 |
| PRKAR1A | 0.619286163 | 2.98E-09 |
| EFNB3 | 0.75891097 | 0.000595297 |
| FAM20A | 3.842429347 | 1.14E-08 |
| MMD | -0.299387801 | 0.004723029 |
| DPH1 | -0.289281364 | 0.000234844 |
| DHRS7B | 0.658946181 | 9.73E-05 |
| WSB1 | 0.217547742 | 0.003642345 |
| MYH1 | 3.597700304 | 4.88E-09 |
| SLC9A3R1 | -0.171436101 | 0.027518706 |
| MYH3 | 2.705269407 | 6.85E-10 |
| NAT9 | 0.300222963 | 0.000405304 |
| TMEM104 | -0.55667588 | 1.27E-05 |
| IFT20 | 0.527482376 | 3.35E-05 |
| TMEM97 | -0.513142483 | 2.01E-06 |
| CDR2L | -1.229295819 | 3.17E-07 |
| PMP22 | -0.811587305 | 6.55E-09 |
| UNC119 | -0.253971152 | 0.003378725 |
| ALDOC | 1.005876449 | 3.05E-07 |
| SUPT6H | -0.692717382 | 1.26E-08 |
| PHF12 | -0.375682573 | 0.000118262 |
| TMEM33 | -1.108410928 | 6.51E-10 |
| GNRHR | 3.422787219 | 3.82E-05 |
| SLAIN2 | -1.122170676 | 3.45E-09 |
| DCUN1D4 | -0.916459337 | 3.68E-08 |
| USP46 | -1.636854808 | 5.41E-09 |
| CRACD | -0.450741503 | 0.013491803 |
| LAMTOR3 | -1.121578164 | 6.03E-08 |
| NFKB1 | -0.48266344 | 9.45E-06 |
| AREG | 7.10296337 | 0.002389341 |
| MANBA | 0.886657039 | 1.72E-05 |
| UBE2D3 | -0.491485017 | 1.21E-07 |
| NDUFC1 | 1.155845051 | 1.67E-06 |
| TBC1D9 | -1.623902725 | 1.04E-08 |
| INPP4B | -0.605081604 | 1.14E-06 |
| GAB1 | -0.581730497 | 0.000505516 |
| KLHL2 | -0.791677115 | 5.00E-05 |
| RPL34 | 0.318307714 | 2.48E-05 |
| WFS1 | 0.614430986 | 2.96E-05 |
| GRPEL1 | -0.478649921 | 1.51E-06 |
| GAR1 | -1.124065406 | 3.26E-08 |
| FRG1 | 0.354150397 | 0.011675596 |
| CLCN3 | -0.697772175 | 1.95E-07 |

| | | |
|---|---|---|
| AADAT | -0.766160753 | 1.46E-05 |
| GALNT7 | -0.78887018 | 3.02E-07 |
| DHX15 | -0.780023584 | 1.73E-09 |
| SOD3 | 0.816583053 | 0.000275064 |
| SEPSECS | 0.631474204 | 4.18E-05 |
| TRIM2 | 0.146773234 | 0.019478765 |
| SLC2A9 | 7.082101064 | 1.41E-05 |
| FBXW7 | 0.194967706 | 0.032748667 |
| NEIL3 | -0.282941769 | 0.004670146 |
| NSD2 | -0.655510612 | 2.27E-08 |
| STIM2 | -0.365853449 | 3.60E-05 |
| NKX3-2 | 1.214847378 | 0.023163055 |
| MFSD10 | 0.277076058 | 0.000811198 |
| BST1 | 0.657079405 | 0.005509355 |
| RAPGEF2 | -0.324979764 | 0.000346771 |
| HGFAC | 2.459077763 | 0.000257144 |
| SNX25 | -0.459780177 | 0.001209174 |
| LRP2BP | 2.347869357 | 2.81E-09 |
| UFSP2 | 0.547387545 | 0.000709225 |
| KLF3 | -0.852527275 | 2.53E-07 |
| NCAPG | -0.549710975 | 2.68E-06 |
| DDX25 | 6.774837438 | 3.09E-05 |
| CRYAB | 1.826408608 | 0.008908158 |
| HTATIP2 | 0.755554281 | 7.38E-07 |
| CTSC | 0.134280383 | 0.009329351 |
| ELP4 | -0.288610321 | 0.037683254 |
| ZPR1 | -0.166814478 | 0.008883999 |
| MTCH2 | 0.219784651 | 0.004203792 |
| FNBP4 | 0.233211375 | 0.000358027 |
| TECTA | 4.913120897 | 2.43E-10 |
| SC5D | 0.512074315 | 1.48E-05 |
| CRTAM | 5.355930411 | 7.84E-05 |
| JHY | 1.123152483 | 0.000959827 |
| HSPA8 | -0.242596401 | 2.11E-08 |
| VWA5A | 1.130927982 | 6.53E-06 |
| DNAJC4 | 0.855247104 | 5.65E-05 |
| SIAE | 0.857605247 | 3.29E-08 |
| SNX15 | 0.354907381 | 0.005410134 |
| LPXN | 0.393827619 | 0.000849461 |
| DTX4 | -0.715314912 | 0.005268547 |
| ATG2A | 0.458351889 | 1.31E-05 |
| EHD1 | -0.999366604 | 3.03E-09 |
| OSBP | 0.305206042 | 6.11E-05 |
| UNC93B1 | 0.83579515 | 0.000166035 |
| PUS3 | 0.332854656 | 0.000109449 |
| KMT5B | -1.049323483 | 2.48E-07 |
| FOXRED1 | 0.488993794 | 5.81E-05 |
| PPP6R3 | -0.80533232 | 1.06E-08 |

| | | |
|---|---|---|
| CPT1A | 0.187851359 | 0.015317282 |
| CCND1 | -2.413302786 | 3.81E-11 |
| CCDC86 | -0.696997916 | 2.94E-07 |
| PRPF19 | -0.511454353 | 1.02E-07 |
| TMEM109 | -0.672947878 | 2.98E-08 |
| HPX | 2.780328144 | 2.36E-05 |
| TRIM3 | -1.090602191 | 7.24E-07 |
| CHORDC1 | -1.154517433 | 2.72E-07 |
| PANX1 | -0.324066026 | 0.000710261 |
| ARHGEF17 | -0.65391527 | 9.24E-05 |
| CEP164 | -0.485433334 | 6.57E-06 |
| RNF141 | 1.184869729 | 2.78E-08 |
| EIF4G2 | -0.65186486 | 6.47E-10 |
| GALNT18 | -1.092575252 | 0.000254775 |
| BIRC2 | -0.593157482 | 6.15E-08 |
| UBE4A | -0.479924212 | 1.73E-06 |
| DDX6 | -0.807978678 | 3.12E-09 |
| CBL | -0.426207692 | 9.89E-07 |
| NECTIN1 | -1.022163311 | 5.17E-09 |
| HIPK3 | -1.599349218 | 4.49E-07 |
| KIAA1549L | -0.550124018 | 4.30E-07 |
| FBXO3 | -0.632509246 | 1.93E-06 |
| PDHX | -0.713701863 | 4.15E-07 |
| COMMD9 | -0.486311989 | 0.000180814 |
| SLC15A3 | 3.434123516 | 9.78E-06 |
| CD5 | 6.726363307 | 3.65E-05 |
| MDK | 0.611555112 | 3.70E-06 |
| AMBRA1 | -0.140669025 | 0.037069794 |
| MADD | 0.193705433 | 0.004639935 |
| PTPMT1 | -0.26265154 | 0.001177536 |
| NAA40 | -1.205938883 | 3.96E-09 |
| SLC35F2 | -0.246244721 | 0.0015731 |
| SOX6 | -0.669920845 | 0.038192113 |
| PITPNM1 | 0.710226846 | 3.41E-06 |
| RPS13 | 0.321520203 | 1.42E-05 |
| AIP | 0.241978893 | 0.001866418 |
| NUP98 | -0.285247781 | 0.000272411 |
| NDUFS8 | 0.285896248 | 0.001349324 |
| TCIRG1 | -0.207307044 | 0.01883871 |
| CHKA | -0.990057317 | 1.33E-07 |
| EXPH5 | 3.489571987 | 9.11E-07 |
| GTF2H1 | -0.533695309 | 4.92E-06 |
| PTPN5 | 4.511547404 | 0.000706496 |
| PSMD9 | 0.196989367 | 0.011850581 |
| PPFIBP1 | -0.374786442 | 7.88E-06 |
| PRDM4 | -0.271846248 | 0.000440505 |
| CLEC2B | -0.418830909 | 0.041495175 |
| COQ5 | 0.532071406 | 6.08E-05 |

| | | |
|---|---|---|
| SELPLG | 3.37812027 | 9.98E-09 |
| CORO1C | -1.135309389 | 3.48E-10 |
| DAO | 7.00011059 | 1.83E-05 |
| CAPRIN2 | 0.957369723 | 2.99E-09 |
| TSPAN11 | 0.252905464 | 0.012770615 |
| KCTD10 | -1.156368656 | 8.54E-10 |
| SLC11A2 | -1.361410352 | 1.64E-07 |
| MLEC | -0.595199614 | 5.33E-09 |
| MVK | 0.952864354 | 2.07E-06 |
| CSRNP2 | 0.57125131 | 5.88E-07 |
| CAMKK2 | -0.726716388 | 3.90E-07 |
| IL23A | 3.261740501 | 5.96E-08 |
| ATP5F1B | 0.286153527 | 2.06E-05 |
| PTGES3 | -0.248493862 | 4.71E-06 |
| BCL7A | -1.540641534 | 9.09E-07 |
| RSRC2 | 0.129381054 | 0.025353329 |
| LIN7A | -1.159635696 | 0.009773547 |
| KRT18 | 0.291984197 | 2.60E-05 |
| ACSS3 | 0.613360804 | 0.015980892 |
| GLI1 | 4.278710033 | 5.83E-06 |
| METAP2 | -0.173074873 | 0.001526488 |
| LTA4H | 1.339511519 | 1.71E-09 |
| ELK3 | -1.180589777 | 2.44E-09 |
| SLC6A12 | 8.174490314 | 3.28E-07 |
| MAGOHB | -0.693345693 | 1.03E-05 |
| TRPV4 | 0.478812621 | 0.00111042 |
| ITFG2 | -0.795008936 | 1.01E-07 |
| FOXM1 | -0.742645363 | 6.94E-09 |
| PARP11 | -1.868910644 | 3.54E-08 |
| ARPC3 | 0.379720138 | 1.84E-05 |
| RAD51AP1 | -0.571329486 | 6.28E-06 |
| CUX2 | 2.110684619 | 0.006735481 |
| SH2B3 | -0.49749565 | 2.21E-08 |
| AKAP3 | 7.309503727 | 0.010503575 |
| MANSC1 | -1.603140625 | 2.03E-06 |
| DUSP16 | -0.312082741 | 0.013429708 |
| CREBL2 | -0.665494991 | 1.42E-06 |
| ACAD10 | 0.402457821 | 0.001841095 |
| ALDH2 | 0.7920478 | 2.56E-05 |
| CDKN1B | -0.345000612 | 0.000128787 |
| GPRC5D | 4.383862569 | 7.79E-09 |
| NAA25 | -1.024787475 | 2.75E-09 |
| SCNN1A | 2.746635593 | 0.003166361 |
| LTBR | -0.577729528 | 4.44E-06 |
| CDK2AP1 | 0.123209415 | 0.011041885 |
| OAS3 | -0.33227311 | 4.80E-05 |
| MGP | 0.719952699 | 0.045743318 |
| RASAL1 | 1.336719178 | 0.038761796 |

| | | |
|---|---|---|
| GTF2H3 | 0.159536467 | 0.008788943 |
| EIF2B1 | -0.286776643 | 9.60E-05 |
| DDX55 | -0.132451517 | 0.044050312 |
| SLC38A1 | -0.939587832 | 7.88E-10 |
| C12orf49 | -1.190017993 | 3.76E-10 |
| VDR | -0.918527191 | 1.83E-08 |
| RFC5 | -0.670819482 | 2.49E-07 |
| STX2 | -0.998990806 | 1.70E-07 |
| ADGRD1 | 3.711219819 | 4.97E-05 |
| TBC1D30 | 0.463639636 | 0.025329397 |
| CAND1 | -0.986998504 | 4.67E-10 |
| RAB5B | 0.20714898 | 0.002199562 |
| MDM1 | -0.683237981 | 4.72E-05 |
| NUP107 | -0.443851702 | 1.47E-05 |
| CNOT2 | -0.910847623 | 2.75E-07 |
| CPSF6 | -0.583872559 | 1.28E-07 |
| KRR1 | -1.09385654 | 7.28E-10 |
| GAPDH | 0.248575352 | 0.000110499 |
| NOP2 | -0.566017144 | 3.12E-07 |
| CHD4 | 0.178407912 | 0.000310903 |
| ACRBP | 4.754965625 | 8.48E-10 |
| UHRF1BP1L | -0.875765751 | 1.32E-07 |
| COPS7A | 0.276760926 | 0.000344109 |
| ING4 | 1.0036034 | 2.17E-06 |
| GNB3 | -1.818087816 | 7.83E-07 |
| CDCA3 | -1.275015698 | 3.89E-09 |
| USP5 | -0.208818632 | 0.002122547 |
| TPI1 | 0.145086174 | 0.004204414 |
| GNPTAB | -0.946442428 | 1.04E-08 |
| SPSB2 | 0.929920801 | 5.18E-05 |
| ENO2 | 0.384976621 | 4.28E-05 |
| ATN1 | -0.112968419 | 0.020084954 |
| C12orf57 | 0.284533584 | 0.000459847 |
| PTPN6 | 1.6495589 | 0.00056866 |
| LPCAT3 | -0.481003226 | 2.80E-05 |
| NT5DC3 | -0.882627343 | 3.31E-07 |
| SUDS3 | -2.109708495 | 1.60E-11 |
| GOLT1B | -0.911988685 | 5.82E-09 |
| GYS2 | 4.293328728 | 0.00304926 |
| LDHB | 0.392520092 | 0.000133817 |
| PRKAB1 | -0.859207492 | 5.48E-07 |
| HCFC2 | -1.627603553 | 2.79E-07 |
| C2CD5 | -1.015556987 | 1.75E-08 |
| RAB35 | -1.451851269 | 2.76E-09 |
| PHC1 | -0.172342795 | 0.027453652 |
| COX6A1 | 0.36254626 | 3.02E-05 |
| RIC8B | -0.839519907 | 1.50E-07 |
| SRSF9 | -0.195822483 | 0.001001772 |

| | | |
|---|---|---|
| AC009533.1 | -0.532036177 | 0.000338526 |
| FGFR1OP2 | 0.257928743 | 0.000369359 |
| COL12A1 | -0.660109877 | 2.25E-07 |
| BTN3A3 | -0.497867356 | 0.013199107 |
| TDP2 | -1.411351617 | 1.47E-08 |
| FRK | 2.415711857 | 0.00026434 |
| DSE | 0.656341146 | 2.59E-06 |
| RSPH4A | 3.568315648 | 1.16E-06 |
| TMEM14C | 0.397736258 | 2.30E-05 |
| PAK1IP1 | -0.571958548 | 3.10E-06 |
| GCNT2 | -0.718670684 | 0.001442442 |
| NEDD9 | 2.302432643 | 2.42E-11 |
| CEP85L | -0.547938699 | 4.75E-05 |
| ADTRP | 2.08927014 | 0.007315606 |
| ASF1A | -0.689642787 | 2.61E-06 |
| MCM9 | -0.278537946 | 0.012498892 |
| FAM184A | -1.203900481 | 0.001141698 |
| RNGTT | -1.756874521 | 7.50E-10 |
| MAN1A1 | -0.859143628 | 2.64E-06 |
| SERINC1 | 0.26015502 | 0.000114062 |
| TPD52L1 | 0.319408547 | 0.034379939 |
| HINT3 | -0.929800411 | 3.28E-06 |
| NCOA7 | -0.853869927 | 5.84E-07 |
| RIPOR2 | 0.97778855 | 0.009193956 |
| SASH1 | -0.569668715 | 2.27E-06 |
| UST | -1.405799716 | 1.73E-06 |
| FBXO5 | -1.080897935 | 6.68E-09 |
| MTRF1L | -0.613883757 | 1.07E-05 |
| PPARD | 0.524447392 | 2.61E-05 |
| MAPK14 | -2.012447899 | 1.05E-10 |
| KCTD20 | -0.32848893 | 2.77E-05 |
| STK38 | -0.751242512 | 1.30E-06 |
| SRSF3 | -1.065137553 | 2.38E-10 |
| SOD2 | -1.007604326 | 5.23E-09 |
| MRPL18 | -0.281236052 | 0.000475448 |
| IL17F | 8.974834989 | 2.35E-08 |
| MCM3 | -0.645506667 | 7.21E-08 |
| RNF8 | -1.224851906 | 8.94E-08 |
| PHACTR1 | 0.872118488 | 0.009651057 |
| ICK | -1.150684767 | 4.30E-08 |
| FBXO9 | -0.633154337 | 1.00E-05 |
| CD83 | 0.230207997 | 0.021012493 |
| MDN1 | -0.460756968 | 8.87E-06 |
| SAYSD1 | -0.611615668 | 0.002028826 |
| BACH2 | -0.750538462 | 0.00790976 |
| RBM24 | -1.561916064 | 0.000707978 |
| TREML2 | 6.671503917 | 4.29E-05 |
| BAG2 | -1.714102289 | 1.11E-10 |

| | | |
|---|---|---|
| RAB23 | -0.486679962 | 0.000183391 |
| FHL5 | 1.649352052 | 0.000126192 |
| CCNC | -0.762221911 | 7.66E-08 |
| E2F3 | -0.608738566 | 2.73E-07 |
| PTP4A1 | 1.207113702 | 2.25E-11 |
| ASCC3 | -0.317252856 | 8.29E-05 |
| BVES | -0.71496516 | 8.40E-05 |
| COL9A1 | 3.273557373 | 0.013771252 |
| MED23 | -0.903708452 | 5.91E-07 |
| WASF1 | -0.495832057 | 5.99E-06 |
| ALDH5A1 | -0.651633054 | 0.000399629 |
| CRYBG1 | -0.520098871 | 3.87E-06 |
| SMAP1 | -0.312523557 | 0.002973208 |
| RPS12 | 0.479531449 | 3.92E-07 |
| C6orf62 | -0.162424014 | 0.004851354 |
| B3GAT2 | -0.445548989 | 0.000227481 |
| GMNN | -1.018702228 | 1.55E-08 |
| EYA4 | -0.327087855 | 0.000625198 |
| SOBP | -0.77278405 | 0.000896615 |
| SNX3 | 0.241722094 | 0.000505083 |
| HBS1L | -0.218667283 | 0.019327696 |
| TRIM38 | -1.589920647 | 1.71E-08 |
| ZBTB24 | -0.743814651 | 1.85E-06 |
| FIG4 | 1.362869089 | 0.000103039 |
| PERP | 0.17879518 | 0.003690348 |
| ARFGEF3 | -1.241222796 | 1.11E-05 |
| SLC16A10 | -2.020554492 | 3.31E-07 |
| HECA | 0.745095757 | 5.76E-07 |
| ADGRG6 | -0.179082208 | 0.010804715 |
| PHACTR2 | -1.14494955 | 6.39E-07 |
| HSD17B8 | 2.441784795 | 0.006932977 |
| CCR6 | 2.832905281 | 0.000126156 |
| PHF1 | 0.747989229 | 0.001193078 |
| CUTA | 0.388620158 | 0.000136949 |
| QKI | -1.180938869 | 3.72E-10 |
| PDE10A | -0.58865951 | 0.005395849 |
| CCND3 | -0.170851963 | 0.010709205 |
| BYSL | -0.869549503 | 8.47E-07 |
| TBP | -0.523608831 | 0.000262679 |
| GUCA1B | 4.222507834 | 0.001592965 |
| BICRAL | 1.131193609 | 3.08E-08 |
| PPP2R5D | -0.19608072 | 0.005541547 |
| MRPL2 | 0.297678541 | 0.007103375 |
| PTK7 | 0.30754369 | 0.000634344 |
| SRF | -0.632890231 | 2.20E-06 |
| CUL9 | 0.226474415 | 0.011179796 |
| DUSP22 | -1.019056695 | 5.59E-06 |
| EXOC2 | -0.735011304 | 4.42E-05 |

| | | |
|---|---|---|
| COX7A2 | 0.526004101 | 1.98E-05 |
| TMEM30A | -0.448849137 | 3.81E-07 |
| GMDS | 0.512631432 | 0.026326699 |
| SENP6 | -0.4870055 | 0.002030597 |
| VEGFA | -0.286233753 | 0.000562137 |
| PRPF4B | -1.940385972 | 4.69E-11 |
| TTK | -1.101849762 | 6.89E-09 |
| SLC29A1 | -0.66262693 | 1.42E-06 |
| BTN2A1 | -0.855369408 | 4.63E-07 |
| LAMA4 | 0.617348003 | 3.68E-07 |
| TENT5A | 1.472017207 | 2.68E-10 |
| FBRSL1 | -0.533177871 | 1.39E-05 |
| ENPP5 | 2.580694957 | 0.00281732 |
| PRSS16 | 1.910346823 | 0.00061428 |
| TBX18 | -0.904721302 | 1.56E-06 |
| ERBIN | -1.513586652 | 4.36E-11 |
| PCDHB2 | -0.301714595 | 0.002386435 |
| NUDT12 | 0.529474577 | 0.000149645 |
| CEP72 | 0.406363718 | 0.011518215 |
| TENT4A | -1.885106155 | 9.98E-11 |
| GHR | -1.316805974 | 1.92E-05 |
| HMGCS1 | 0.8494504 | 2.03E-08 |
| DAP | 0.657809281 | 2.24E-08 |
| BRD8 | -0.157401857 | 0.009273853 |
| KIF20A | -1.074878975 | 6.77E-10 |
| MRPS30 | -0.346282585 | 9.39E-05 |
| HSPA9 | -0.132649871 | 0.002249198 |
| MRPS27 | -0.411776397 | 3.99E-06 |
| PFDN1 | -0.621155894 | 5.56E-07 |
| HBEGF | 0.62336775 | 1.16E-06 |
| SLC4A9 | 4.311705309 | 0.001254075 |
| APBB3 | -0.885032613 | 1.36E-06 |
| SPARC | 0.396936835 | 1.37E-07 |
| HMGCR | 1.534904318 | 7.34E-12 |
| CERT1 | -0.95593537 | 1.25E-08 |
| PCDHB5 | -0.425011985 | 0.000530234 |
| PCDHB6 | -0.407498787 | 0.000265102 |
| PCDHB7 | -0.690259568 | 0.000426861 |
| CLK4 | 0.312631903 | 0.005301679 |
| PCDHB15 | -0.628263307 | 0.011972381 |
| THG1L | -0.395264256 | 0.000903182 |
| ARSB | -0.647268227 | 2.32E-06 |
| CLINT1 | -0.464381834 | 5.37E-07 |
| THBS4 | 2.646671095 | 0.000865369 |
| CNOT6 | -0.912316447 | 1.24E-08 |
| TTC1 | -0.221184177 | 0.001337121 |
| MSH3 | -0.323764178 | 0.002630598 |
| RASGRF2 | 3.036519508 | 1.12E-08 |

| | | |
|---|---|---|
| CCNG1 | 0.415477846 | 1.61E-06 |
| POLR3G | -0.894407622 | 1.56E-08 |
| DROSHA | -0.17896061 | 0.008951066 |
| LMNB1 | -2.179261549 | 2.32E-08 |
| ARRDC3 | -1.490990409 | 2.66E-09 |
| GOLPH3 | -0.919743111 | 2.57E-09 |
| SUB1 | -0.220196899 | 0.003855999 |
| NPR3 | 2.924675495 | 2.19E-05 |
| FAM172A | -1.079737835 | 3.14E-08 |
| TARS1 | -0.377663026 | 3.22E-06 |
| LNPEP | -0.597511095 | 2.61E-07 |
| PDE4D | -0.80398417 | 1.47E-07 |
| RAD1 | -1.399495036 | 1.02E-06 |
| BRIX1 | -1.117785312 | 6.78E-10 |
| RAD50 | 0.215071508 | 0.00609906 |
| IL5 | 9.044499535 | 1.83E-08 |
| ST8SIA4 | 0.585820445 | 0.000229868 |
| GNPDA1 | -0.380439754 | 3.31E-05 |
| PCDH12 | 3.102503511 | 6.61E-07 |
| NUP155 | -0.828034266 | 1.76E-08 |
| PPP2CA | -0.838979837 | 1.77E-09 |
| FGF1 | 0.855622898 | 0.007837809 |
| NR3C1 | -0.480982273 | 2.77E-06 |
| C5orf15 | -0.337906643 | 4.77E-05 |
| PPWD1 | -0.333251409 | 0.002061701 |
| TRIM23 | -0.260473967 | 0.005654971 |
| TRAPPC13 | -1.309164016 | 3.87E-07 |
| SEC24A | -1.306035276 | 2.77E-10 |
| TXNDC15 | -0.46199726 | 3.19E-05 |
| TTC33 | -0.496077354 | 0.001164427 |
| RARS1 | -0.272062138 | 0.000253171 |
| WWC1 | -1.487523034 | 1.22E-06 |
| MACROH2A1 | -0.197971801 | 0.000597424 |
| TCERG1 | -0.85094459 | 5.84E-09 |
| DPYSL3 | -0.167242161 | 0.004520299 |
| SMAD5 | -0.720299776 | 1.83E-08 |
| CSNK1A1 | -0.654429072 | 1.17E-08 |
| HMGXB3 | -1.639568914 | 1.03E-10 |
| PDGFRB | 0.544849112 | 0.012030454 |
| ATP6V0E1 | -0.396479953 | 0.000356847 |
| BNIP1 | -1.110701154 | 8.52E-06 |
| STC2 | -1.525385807 | 4.43E-08 |
| CPEB4 | 1.082109124 | 6.26E-10 |
| DBN1 | 0.603454782 | 1.86E-06 |
| ZNF346 | -0.792591467 | 1.37E-05 |
| EHHADH | 1.931758684 | 7.21E-09 |
| CNTN3 | -0.401164777 | 0.022588628 |
| SMC4 | -0.770788863 | 1.58E-09 |

| | | |
|---|---|---|
| SELENOK | 0.27392778 | 0.001581015 |
| ACTR8 | -0.783103779 | 1.12E-06 |
| TIMMDC1 | 0.157919761 | 0.013959443 |
| CRBN | -0.494602653 | 9.75E-06 |
| BCL6 | 1.72091961 | 1.24E-09 |
| ARL6 | -0.432446227 | 0.001715177 |
| NPHP3 | 0.522323905 | 3.30E-05 |
| AMOTL2 | -0.149554016 | 0.007357036 |
| NIT2 | -0.477143839 | 0.000791179 |
| KPNA1 | -0.953034172 | 5.56E-09 |
| PCCB | -0.633840746 | 6.32E-07 |
| UBE3A | -1.224068548 | 9.34E-11 |
| ARMC8 | -0.755665837 | 4.38E-07 |
| CEP70 | -0.448540754 | 0.00302107 |
| GRK7 | 6.787344502 | 7.64E-06 |
| RNF7 | -0.549399537 | 5.10E-06 |
| XRN1 | -0.325551769 | 0.000370744 |
| KAT2B | -0.408164581 | 0.000468767 |
| PDCD10 | -0.261608851 | 0.003982973 |
| WNT5A | -1.212037414 | 2.17E-08 |
| COL7A1 | 0.73265762 | 6.31E-06 |
| PRKAR2A | -0.705090001 | 3.06E-08 |
| ACAP2 | -0.77125037 | 1.11E-07 |
| ECT2 | -1.506931772 | 1.80E-10 |
| GNAI2 | -0.092474851 | 0.048458407 |
| TFG | -0.577529986 | 2.02E-07 |
| HYAL1 | 3.822407167 | 6.85E-06 |
| RPL24 | 0.383483444 | 1.00E-04 |
| CYB561D2 | -0.221842207 | 0.02588673 |
| CBLB | 0.958620065 | 3.30E-06 |
| BBX | -0.860175774 | 2.81E-09 |
| IFT57 | 0.489685495 | 0.000280771 |
| GNB4 | -1.102123368 | 2.88E-08 |
| IQCG | 1.730056513 | 3.48E-06 |
| GBE1 | -0.524011023 | 9.12E-06 |
| UMPS | -0.769898036 | 1.63E-07 |
| NCBP2 | -0.792527778 | 5.99E-09 |
| SNX4 | -1.197967276 | 1.87E-08 |
| C3orf52 | -0.368809768 | 0.008384269 |
| FRMD4B | -0.260373316 | 0.003848582 |
| SLC41A3 | -0.236179177 | 0.020048424 |
| ROPN1B | 1.144771324 | 0.027518706 |
| PLXNA1 | -0.554215064 | 3.32E-07 |
| ABTB1 | 0.593178504 | 0.000219612 |
| CSPG5 | 0.720753824 | 0.01455871 |
| KLHL18 | -0.722017465 | 5.51E-08 |
| SCAP | -0.224674648 | 0.001229967 |
| NEK11 | 1.49118517 | 0.003868815 |

| | | |
|---|---|---|
| MRPL3 | -0.462952238 | 5.93E-06 |
| HEMK1 | 1.777631648 | 1.06E-10 |
| CISH | 2.421233945 | 1.97E-08 |
| ACVR2B | -0.948419752 | 2.46E-07 |
| WDR48 | -0.605639307 | 1.21E-06 |
| COMMD2 | -1.060783448 | 4.61E-08 |
| GORASP1 | -0.784878965 | 6.82E-07 |
| ABCC5 | 0.202181451 | 0.004532074 |
| ABHD14B | 0.581048342 | 4.73E-07 |
| ABHD14A-ACY1 | 0.588762624 | 2.22E-05 |
| ARHGEF26 | -0.860982037 | 9.86E-07 |
| KLHL24 | 2.412433527 | 3.86E-12 |
| PLCH1 | 1.596823686 | 0.014162021 |
| DNAH1 | 1.779856793 | 1.27E-07 |
| SSR3 | -0.651398255 | 5.36E-08 |
| ZBTB47 | -1.263559934 | 1.39E-06 |
| TNNC1 | 2.600728793 | 0.00296751 |
| NKTR | -0.349969225 | 5.15E-06 |
| EIF4G1 | -0.611028595 | 5.00E-09 |
| SPCS1 | 0.114877366 | 0.04224979 |
| NEK4 | -0.48293827 | 4.08E-05 |
| INO80D | -0.974668819 | 3.23E-08 |
| EEF1B2 | 0.247764855 | 0.000491914 |
| DGUOK | 0.307485696 | 0.002081722 |
| MOB1A | -0.37711925 | 3.08E-06 |
| KANSL3 | -0.611260207 | 3.78E-07 |
| LMAN2L | -0.332142479 | 0.007621213 |
| RTKN | -0.350980813 | 0.003847812 |
| TTL | -0.754246488 | 1.46E-07 |
| PIKFYVE | -0.151112226 | 0.011475381 |
| FAHD2A | 0.476589171 | 0.00117388 |
| NCL | -0.262096003 | 3.10E-06 |
| ACTR1B | 0.751624349 | 1.22E-06 |
| SLC35F5 | -0.321541552 | 0.00055448 |
| ACTR3 | -0.338899658 | 1.97E-05 |
| EPB41L5 | -1.025406205 | 2.10E-06 |
| SF3B6 | 0.285876598 | 0.000364268 |
| TP53I3 | 0.810293585 | 2.39E-06 |
| STAM2 | -0.718622821 | 5.83E-06 |
| GPD2 | -0.416480551 | 7.13E-06 |
| CENPA | -1.859009081 | 2.15E-10 |
| TANC1 | 0.447752594 | 2.28E-05 |
| SLC30A3 | 1.379153807 | 3.41E-05 |
| MPV17 | 0.296380208 | 0.000569506 |
| GTF3C2 | -1.01499422 | 4.02E-09 |
| NRBP1 | -0.59474934 | 6.20E-07 |
| ITGA4 | -0.173092284 | 0.022318246 |
| PSMD14 | -0.269001173 | 0.000270797 |

| | | |
|---|---|---|
| SNX17 | 0.348063698 | 4.48E-05 |
| ASB3 | -0.372169805 | 0.002295351 |
| REEP6 | 1.606208739 | 8.69E-06 |
| PCSK4 | 3.022178428 | 0.000488123 |
| APC2 | 0.80506618 | 0.002621921 |
| IFIH1 | -0.675150554 | 0.01716744 |
| RPS15 | 0.335628272 | 1.10E-05 |
| GCA | 0.464790256 | 0.009651057 |
| INO80B | 0.806648383 | 4.41E-06 |
| MOGS | -0.180585544 | 0.005604971 |
| TTC31 | -0.733701194 | 1.11E-07 |
| CLIP4 | -0.936606865 | 3.42E-08 |
| TLX2 | 2.867035791 | 2.23E-06 |
| SPTBN1 | -0.14467813 | 0.005724722 |
| AUP1 | -0.304956871 | 7.64E-05 |
| RTN4 | -0.327291448 | 2.57E-06 |
| LOXL3 | 0.224322311 | 0.000244651 |
| DOK1 | 0.571506017 | 7.02E-05 |
| GALNT3 | 9.191631008 | 8.43E-09 |
| CCDC88A | -0.257433012 | 0.000344753 |
| MRPL19 | -1.032666599 | 2.86E-09 |
| LANCL1 | -0.614344416 | 2.38E-07 |
| WDR75 | -0.416598855 | 1.50E-05 |
| FANCL | -1.115770843 | 1.86E-08 |
| FN1 | 0.807389047 | 1.63E-08 |
| STAT1 | -0.437754084 | 4.18E-06 |
| GLS | -1.478063518 | 8.82E-11 |
| DNAH6 | 5.988650505 | 0.00064149 |
| IGFBP2 | 0.345629593 | 0.001362568 |
| IGFBP5 | 1.260228766 | 1.63E-06 |
| USP34 | -0.607126309 | 5.29E-08 |
| EFHD1 | 2.389400926 | 0.000108512 |
| KCNJ13 | 3.120470048 | 0.000330262 |
| CCT4 | 0.088538966 | 0.040575676 |
| GGCX | 0.43874121 | 1.82E-06 |
| EHBP1 | -0.846963655 | 1.94E-08 |
| TXNDC9 | -1.102575661 | 4.85E-08 |
| GNLY | 8.211395164 | 2.86E-07 |
| ST3GAL5 | -0.765560337 | 9.75E-06 |
| CHST10 | -0.788608369 | 9.34E-07 |
| MOB4 | -0.798659358 | 3.98E-07 |
| HSPE1 | -0.109410496 | 0.041317283 |
| KDM3A | 0.835092272 | 1.68E-08 |
| PLCD4 | -0.821457011 | 0.000119097 |
| ZNF142 | -1.524113728 | 5.89E-09 |
| IL1RL1 | 1.906735954 | 1.77E-08 |
| IL18R1 | 2.013003702 | 2.68E-06 |
| FHL2 | 0.342133006 | 0.000540197 |

| | | |
|---|---|---|
| UXS1 | -0.172814739 | 0.018030088 |
| ABCB6 | 0.334202832 | 0.003204549 |
| STK16 | 0.837226204 | 3.46E-06 |
| HDLBP | 0.456581074 | 1.21E-07 |
| PPP1R7 | 0.380373263 | 0.005851541 |
| ID2 | 1.504919284 | 1.16E-08 |
| TAF1B | -0.868422881 | 7.11E-06 |
| ODC1 | -0.230961463 | 0.001249138 |
| BIRC6 | -0.537205646 | 6.49E-07 |
| NOL10 | -1.124799394 | 7.65E-09 |
| PLEKHB2 | -0.720949589 | 8.85E-09 |
| GORASP2 | -0.811552123 | 4.27E-07 |
| STRN | -1.324517136 | 2.79E-10 |
| CEBPZ | -0.695122176 | 1.32E-08 |
| PRKD3 | -0.845499821 | 2.01E-08 |
| DCAF17 | -0.649748552 | 0.000142133 |
| QPCT | 0.798782475 | 0.00086719 |
| RAB3GAP1 | -0.420850012 | 0.000136714 |
| SLC25A12 | -1.549331249 | 9.57E-08 |
| RMDN2 | 0.85716947 | 0.000134541 |
| DLX2 | -2.174218951 | 9.91E-09 |
| LCT | 4.894534474 | 0.000262568 |
| DARS1 | -0.360567331 | 0.000206699 |
| SRSF7 | -0.944758132 | 1.38E-09 |
| PLCL1 | -2.381584535 | 0.000463243 |
| SLC1A4 | -0.622556307 | 2.92E-07 |
| SOS1 | -1.564749553 | 8.34E-11 |
| KYNU | 0.378094886 | 0.01379528 |
| WIPF1 | -0.315997214 | 0.000126496 |
| ORC2 | -0.782239473 | 1.78E-06 |
| COX7A2L | 0.167810607 | 0.008427016 |
| PNO1 | -0.956769732 | 9.32E-09 |
| ORC4 | -0.599185442 | 2.98E-05 |
| PLEK | 7.267575792 | 9.04E-06 |
| RND3 | 0.286981122 | 0.001022529 |
| ATF2 | -0.865169258 | 3.02E-07 |
| THADA | -0.410276427 | 0.000103987 |
| AAK1 | -1.549980518 | 3.67E-10 |
| TRAK2 | -1.469469963 | 2.94E-09 |
| C2orf42 | 1.125352219 | 2.40E-08 |
| TIA1 | -1.040247659 | 3.54E-08 |
| PCYOX1 | 0.152289145 | 0.007860293 |
| EPAS1 | -0.750756917 | 0.002117627 |
| ARID3A | 1.071957374 | 8.94E-06 |
| SUMO1 | -0.353192875 | 1.67E-05 |
| GRIN3B | 1.631848523 | 0.014735715 |
| VAX2 | 0.7918379 | 0.041875675 |
| ATP6V1B1 | 2.602119183 | 0.001094868 |

| | | |
|---|---|---|
| NFE2L2 | -1.351344402 | 1.25E-10 |
| MSH6 | -0.825173202 | 1.66E-09 |
| PLEKHA3 | -0.234382256 | 0.027790025 |
| EPHA4 | 0.560457446 | 5.84E-06 |
| PARD3B | -1.015005581 | 0.004937795 |
| FARSB | -0.220158358 | 0.011248167 |
| ALMS1 | -0.573533667 | 3.57E-07 |
| BCL9 | -1.220444599 | 1.56E-09 |
| PRRX1 | -0.211304708 | 0.01715465 |
| DHCR24 | -0.093668357 | 0.035108034 |
| DNAJC16 | 0.655214206 | 1.26E-06 |
| MARK1 | -2.507125672 | 8.32E-09 |
| RALGPS2 | -0.991801717 | 2.59E-08 |
| ANGPTL1 | 1.419310798 | 0.013779992 |
| FAM20B | -0.934635673 | 1.63E-09 |
| TCEANC2 | -0.986571456 | 9.30E-07 |
| TMEM59 | 0.390386052 | 6.63E-06 |
| LRRC42 | -0.720817404 | 1.70E-05 |
| WRAP73 | -0.936703409 | 1.37E-07 |
| ICMT | -0.585236992 | 4.46E-08 |
| RPL22 | 0.231884537 | 0.000871219 |
| CHD5 | 5.46265764 | 3.04E-10 |
| STXBP3 | -0.240872046 | 0.008395761 |
| PHF13 | -0.153974642 | 0.027919481 |
| ERRFI1 | -0.183264317 | 0.000170952 |
| PARK7 | 0.132833495 | 0.022762791 |
| KIAA1324 | 4.52067379 | 1.72E-09 |
| OPRD1 | -1.689867903 | 1.75E-05 |
| AMPD2 | -0.62147736 | 5.41E-07 |
| SRSF4 | -1.766807848 | 1.94E-11 |
| KCNC4 | -0.579094348 | 0.000514513 |
| EDEM3 | -0.613665442 | 1.25E-07 |
| WDR77 | -0.772221373 | 7.22E-08 |
| ATP5PB | 0.115284857 | 0.035222315 |
| RAP1A | -0.213841581 | 0.011095653 |
| HDAC1 | -0.207337801 | 0.00112583 |
| CAPZA1 | -0.440019147 | 3.65E-07 |
| S100PBP | -1.437232692 | 1.17E-09 |
| ASH1L | -1.503171253 | 1.13E-10 |
| SFPQ | -0.569324493 | 4.14E-09 |
| RHOU | 0.469960705 | 0.035891489 |
| ARHGEF2 | 0.19401789 | 0.001044135 |
| LAMTOR2 | 0.394247573 | 0.00020222 |
| MEF2D | 0.467155356 | 2.83E-06 |
| DLEU2L | 0.658333503 | 0.001572187 |
| C1orf21 | -0.605912987 | 0.000459137 |
| DNAJC6 | -0.374191777 | 0.000109063 |
| LEPR | 1.256704749 | 7.41E-07 |

| | | |
|---|---|---|
| IVNS1ABP | 0.278787429 | 7.56E-05 |
| KIAA2013 | -0.58077235 | 4.52E-07 |
| MFN2 | -0.614675759 | 5.19E-08 |
| SMG7 | -0.941533124 | 1.19E-09 |
| PDC | 7.306404312 | 6.21E-06 |
| SLC35D1 | -2.492218587 | 6.29E-12 |
| GADD45A | 2.660544877 | 2.03E-12 |
| WLS | -0.142121608 | 0.034073013 |
| PRDM2 | 0.497865871 | 9.27E-07 |
| RGS2 | 2.612823138 | 4.90E-13 |
| RPE65 | 4.794338938 | 0.00025874 |
| RO60 | -0.376370335 | 2.91E-05 |
| UCHL5 | -0.442025322 | 1.67E-06 |
| BCAS2 | -0.861114106 | 1.62E-06 |
| SRSF11 | -0.629914848 | 1.74E-08 |
| CTH | 1.038837487 | 4.10E-06 |
| AGMAT | 3.970057523 | 5.14E-05 |
| CFHR3 | 1.874438495 | 0.002507584 |
| CRYZ | -0.282261346 | 0.002448742 |
| PHTF1 | -0.520310754 | 2.71E-05 |
| CD58 | -0.728958972 | 0.000254032 |
| TFAP2E | 1.488281637 | 0.003977244 |
| TTF2 | -1.091344796 | 4.80E-09 |
| KIF21B | -0.740205884 | 1.33E-06 |
| MAP7D1 | -0.346910549 | 7.53E-05 |
| WARS2 | -0.494106059 | 5.39E-05 |
| OSCP1 | 0.868007543 | 0.00238567 |
| GNPAT | -0.34396263 | 0.000238698 |
| TSNAX | -0.502956695 | 1.13E-06 |
| RRAGC | 0.252524761 | 0.002806503 |
| NID1 | 0.499272707 | 1.07E-06 |
| LGALS8 | -0.247148369 | 0.000226469 |
| HPCAL4 | 0.603190146 | 0.014735715 |
| MTR | -0.683900614 | 3.56E-07 |
| RLF | 0.568165084 | 7.81E-07 |
| KMO | 0.782125839 | 0.000209627 |
| ZNF684 | 0.31770363 | 0.024214426 |
| KCNQ4 | 1.793149992 | 0.014151019 |
| RIMS3 | 3.889752506 | 8.89E-10 |
| AKT3 | -0.772727334 | 3.18E-07 |
| ETV3 | 1.144518992 | 1.47E-10 |
| ADGRL2 | -1.227055485 | 2.56E-08 |
| SDHB | 0.350640521 | 0.001599081 |
| MFAP2 | 0.226204071 | 0.041850628 |
| RPF1 | -0.310520904 | 0.000413358 |
| KDM5B | 0.352847512 | 4.16E-05 |
| UAP1 | -1.316706167 | 9.82E-11 |
| CTBS | 0.800610924 | 0.000121275 |

| | | |
|---|---|---|
| RGS4 | 1.392526234 | 0.000921108 |
| KLHL12 | 0.347273934 | 4.80E-05 |
| SSX2IP | -0.463261748 | 4.04E-05 |
| ZNHIT6 | -1.990780024 | 4.86E-10 |
| RBBP5 | -0.731102151 | 6.71E-08 |
| GBP3 | -1.949534672 | 9.16E-05 |
| GBP1 | -0.755992217 | 0.002027791 |
| GPR89A | -0.657669186 | 6.15E-05 |
| CDK18 | 1.325524444 | 2.85E-06 |
| RAB29 | -0.425171222 | 1.49E-05 |
| ECE1 | -0.252346168 | 0.000521933 |
| HMGCL | 1.415297733 | 1.33E-07 |
| GALE | 1.268664863 | 1.78E-09 |
| CD46 | -0.286006089 | 2.36E-05 |
| PRPF3 | -1.279455093 | 1.79E-09 |
| P3H1 | 0.265162723 | 0.000634333 |
| SLC2A1 | -0.630609138 | 1.57E-08 |
| EBNA1BP2 | -0.351573224 | 8.48E-06 |
| CDC20 | -0.844240711 | 1.34E-09 |
| MPL | 2.267136302 | 0.003284949 |
| ARTN | 2.479726963 | 1.21E-06 |
| B4GALT2 | -0.160598098 | 0.025241756 |
| PTCH2 | 1.598047233 | 0.000131116 |
| AKR1A1 | 0.488192739 | 5.72E-05 |
| PIK3R3 | -2.298955169 | 1.35E-10 |
| BLZF1 | -2.193582098 | 4.49E-11 |
| CCDC181 | -0.391324195 | 0.005171308 |
| SLC19A2 | -0.882329227 | 2.70E-08 |
| FAAH | 4.09257475 | 7.56E-07 |
| NSUN4 | -1.662957099 | 1.08E-08 |
| TMED5 | -0.343185781 | 6.19E-05 |
| DR1 | -1.153369679 | 1.07E-09 |
| CNN3 | -0.397454047 | 2.68E-06 |
| PRRC2C | -0.650681925 | 1.76E-09 |
| F3 | -1.185511875 | 4.07E-05 |
| ABCD3 | -0.850628835 | 3.60E-08 |
| VAMP4 | 1.156051998 | 6.34E-08 |
| DPH5 | 0.68837575 | 1.02E-05 |
| PTBP2 | 0.778589876 | 1.45E-06 |
| TNFSF4 | 2.391628135 | 1.07E-05 |
| DARS2 | -0.39500111 | 8.45E-06 |
| IRF6 | 2.266356016 | 0.003408035 |
| UTP25 | -0.263732391 | 0.000603006 |
| SERPINC1 | 5.031265592 | 0.000107775 |
| SYF2 | 0.20005598 | 0.006100199 |
| RSRP1 | -0.130557086 | 0.02928621 |
| RCOR3 | -0.464476639 | 1.06E-05 |
| MTFR1L | -0.62557685 | 6.50E-07 |

| | | |
|---|---|---|
| MAN1C1 | -0.821145673 | 0.007364924 |
| NEK2 | -0.852193601 | 9.71E-08 |
| RPS6KA1 | -0.40537115 | 3.04E-05 |
| DHDDS | -0.882339478 | 1.18E-07 |
| NSL1 | 0.240348182 | 0.002194419 |
| PROX1 | 0.822981256 | 1.68E-05 |
| ARID1A | -1.137534007 | 3.06E-10 |
| CENPF | -1.23422756 | 3.19E-11 |
| RPA2 | -0.297131119 | 0.000458722 |
| PPP1R8 | -0.703440364 | 3.98E-08 |
| SLC5A9 | -1.400398111 | 0.011428817 |
| OSBPL9 | 0.269403071 | 0.000623205 |
| ESYT2 | -0.496100535 | 9.82E-07 |
| CD3EAP | -0.763615648 | 5.16E-08 |
| MESD | -0.572950381 | 2.14E-07 |
| RCN2 | -0.447287319 | 5.55E-06 |
| MUC5B | -0.860015621 | 2.75E-08 |
| CTSD | 0.5585378 | 3.02E-07 |
| STAG1 | -1.289677357 | 4.38E-10 |
| TREH | -0.454284377 | 0.003473551 |
| IFT46 | -0.317910227 | 0.001859812 |
| MMP8 | 0.842300955 | 0.007012315 |
| ZNF541 | 8.056382608 | 6.48E-07 |
| SLC8A2 | 5.486059235 | 2.46E-05 |
| RPS25 | 0.272938993 | 2.37E-05 |
| KIF14 | -1.690075212 | 8.83E-11 |
| TNNT2 | 6.872291452 | 2.47E-05 |
| DDX59 | 0.386741508 | 6.01E-05 |
| CAMSAP2 | -0.949414301 | 7.28E-10 |
| ATF6 | -0.593430251 | 1.41E-06 |
| MREG | -2.141373326 | 3.15E-07 |
| FASTKD2 | -1.329553582 | 5.89E-10 |
| NRP2 | 0.439480878 | 3.68E-06 |
| CREB1 | -1.172337108 | 2.95E-09 |
| KLF7 | 1.040929619 | 3.93E-07 |
| B4GALT6 | -0.841841584 | 8.25E-08 |
| CA14 | 0.934829664 | 0.015883953 |
| CASC1 | 3.025943591 | 3.02E-05 |
| ATP10B | 1.462077494 | 4.72E-05 |
| SPCS2 | -0.23518566 | 0.00359044 |
| USP35 | -0.774924594 | 0.000322121 |
| ELOVL4 | 0.873474191 | 4.71E-08 |
| FILIP1 | 5.469198367 | 1.96E-05 |
| CASP8AP2 | 0.625745871 | 8.47E-08 |
| HMGN3 | -0.209058878 | 0.040820889 |
| UBE3D | 0.567504704 | 0.029562913 |
| CNR1 | -0.915671417 | 1.45E-07 |
| ANKRD13C | -2.092836196 | 5.59E-09 |

| | | |
|---|---|---|
| PHF3 | -0.336920452 | 7.35E-06 |
| FBXO30 | -0.188430064 | 0.007427394 |
| TNFAIP3 | 2.181591945 | 3.71E-10 |
| AKAP7 | -0.403452626 | 0.034224566 |
| RAB32 | 0.565203843 | 0.000109071 |
| MYB | -2.318807632 | 3.72E-05 |
| SGK1 | 0.162390087 | 0.00469403 |
| RNF146 | 1.04294213 | 9.88E-09 |
| CCN2 | 1.792941166 | 7.39E-13 |
| FBXL5 | 0.357765994 | 3.75E-05 |
| MED28 | -1.088724782 | 2.00E-09 |
| SLC16A7 | -1.554605384 | 5.58E-07 |
| RXYLT1 | 0.654321254 | 4.17E-07 |
| ZNF430 | 0.224809306 | 0.004774673 |
| DCLRE1B | -0.280570011 | 0.001449704 |
| MYL12B | 0.172835388 | 0.001922792 |
| FOXO3 | 0.367224557 | 0.000554861 |
| ARMC2 | 0.601329613 | 0.022803948 |
| RPN2 | 0.273065693 | 0.000136496 |
| TGIF2 | -1.273655789 | 4.41E-09 |
| OLFM3 | 7.861451735 | 1.30E-06 |
| ABCG2 | -0.659524057 | 0.010144691 |
| SPP1 | 1.586281185 | 1.22E-06 |
| STBD1 | -0.89180624 | 6.36E-06 |
| CCNI | 0.201620262 | 0.000786455 |
| MFSD1 | -0.576500829 | 2.76E-05 |
| RAB3GAP2 | -0.155978927 | 0.010600817 |
| PPL | 3.029449738 | 0.00391162 |
| UBN1 | -1.102327338 | 4.76E-10 |
| KLF12 | -0.171572725 | 0.029831057 |
| PCDH17 | -0.326587089 | 0.004315339 |
| HS1BP3 | 0.383548781 | 0.002788047 |
| LDAH | -0.518700807 | 2.21E-05 |
| OTUD4P1 | -1.33061846 | 4.10E-08 |
| ELL2 | 0.534773031 | 1.01E-05 |
| DNAH7 | 2.286048311 | 0.000437797 |
| CYP20A1 | 0.342862836 | 0.032330118 |
| SATB2 | -0.878584787 | 5.13E-07 |
| UBE2B | -0.294088294 | 0.000268849 |
| GDA | -1.616678053 | 6.28E-11 |
| KLF9 | -1.401290948 | 4.24E-10 |
| TJP2 | -0.907691733 | 1.82E-08 |
| ITGB1BP1 | -0.757139832 | 2.22E-08 |
| CPSF3 | 0.244760174 | 0.003619554 |
| PIGZ | 2.999702068 | 0.000997887 |
| SENP5 | -0.245123538 | 0.002431639 |
| CCDC92 | 0.741408152 | 0.001239619 |
| C1orf198 | -1.417519081 | 7.19E-10 |

| | | |
|---|---|---|
| HEATR1 | -0.398712518 | 5.12E-07 |
| PTBP3 | -0.790651786 | 4.99E-09 |
| RAD23B | -0.450279222 | 3.56E-07 |
| FKBP15 | -0.715328516 | 4.42E-07 |
| ABITRAM | -0.887743297 | 3.40E-08 |
| WDR34 | 0.168980477 | 0.028215112 |
| SET | -0.099235055 | 0.023131368 |
| PTPA | -0.321696453 | 4.01E-05 |
| RAB14 | -0.609829897 | 2.76E-08 |
| TRIM32 | -1.601869926 | 6.00E-10 |
| FBXW2 | -0.602181732 | 9.52E-08 |
| PHF19 | -0.312284227 | 0.000198768 |
| NEK6 | -0.85264314 | 6.20E-07 |
| PPP6C | -0.333731031 | 2.98E-05 |
| NDUFA8 | 0.218927046 | 0.010715499 |
| HDHD3 | 0.532849796 | 0.002606763 |
| RBM18 | -1.631081049 | 1.71E-10 |
| MAPKAP1 | -0.49819578 | 4.25E-06 |
| NR4A3 | 3.093859053 | 5.59E-11 |
| INVS | 0.272241105 | 0.047697403 |
| DENND1A | -0.565226323 | 0.000270398 |
| ALG2 | 0.369094011 | 6.42E-05 |
| VPS4B | -0.419353883 | 9.37E-06 |
| ONECUT2 | 1.13543952 | 1.72E-07 |
| ZBTB45 | 0.429422616 | 0.000957317 |
| YLPM1 | -0.769000269 | 1.00E-08 |
| FCF1 | -0.239833545 | 0.003237955 |
| IFI27L2 | 0.533510896 | 0.014868197 |
| BBOF1 | 2.061255376 | 3.63E-06 |
| NEK9 | 0.594629251 | 5.31E-06 |
| ACYP1 | 0.340624174 | 0.041900643 |
| NPC2 | 0.66456793 | 4.09E-06 |
| DNAL1 | -1.183657587 | 4.93E-06 |
| IRF2BPL | -1.913127469 | 2.99E-09 |
| LTBP2 | 2.864598214 | 2.54E-09 |
| AREL1 | -1.768555241 | 9.16E-11 |
| MLH3 | 1.08195558 | 0.039235548 |
| TTLL5 | -0.36506628 | 0.005751793 |
| DLST | -0.352712368 | 2.67E-05 |
| TGFB3 | -0.793077108 | 0.002237083 |
| ZC2HC1C | 2.719185027 | 2.39E-06 |
| SLIRP | 0.292964792 | 0.030830739 |
| RBM25 | -0.501263158 | 3.34E-07 |
| ALDH6A1 | 2.009675638 | 3.84E-08 |
| GPR68 | 2.765520718 | 5.60E-05 |
| ESRRB | 3.779136846 | 0.000120544 |
| EIF2B2 | 0.207966946 | 0.007397944 |
| NRDE2 | -0.477862313 | 1.54E-05 |

| | | |
|---|---|---|
| COQ6 | 0.314781571 | 0.004890902 |
| ZNF410 | -0.193480286 | 0.019209575 |
| GPR75 | -1.207917408 | 2.08E-05 |
| SUPT7L | -0.715772536 | 2.17E-07 |
| KLHL29 | -0.616070435 | 0.000181179 |
| TMEM214 | -0.287703461 | 0.000162876 |
| ATAD2B | -1.37846213 | 6.10E-09 |
| FKBP1B | 0.633599606 | 0.030407826 |
| ATL2 | -1.049875564 | 2.56E-09 |
| YPEL5 | 1.196425805 | 7.96E-09 |
| FAM98A | -1.12353055 | 7.72E-10 |
| YIPF4 | -0.691028428 | 2.14E-07 |
| AFTPH | 0.209051967 | 0.003826001 |
| LGALSL | -0.786587642 | 0.000143986 |
| CNRIP1 | -0.542148319 | 0.000422405 |
| BCL11A | 1.682516178 | 0.000137803 |
| CRIPT | -0.773448502 | 1.96E-07 |
| OGFRL1 | -0.765814177 | 4.59E-07 |
| SLF2 | 0.758345047 | 1.93E-08 |
| IDE | -0.62500271 | 8.20E-07 |
| IFIT3 | -1.492270805 | 2.00E-06 |
| IFIT2 | -2.039341849 | 2.97E-06 |
| GPAM | -0.628187829 | 9.32E-07 |
| PYROXD2 | 0.783472112 | 0.000283089 |
| MXI1 | 0.217967096 | 0.009651057 |
| SMNDC1 | -1.1682869 | 1.19E-09 |
| HELLS | -0.902739543 | 1.40E-08 |
| PRLHR | 8.381140975 | 2.30E-07 |
| TCTN3 | 0.451429431 | 9.58E-06 |
| AVPI1 | 2.216048058 | 3.84E-11 |
| ARMH3 | -0.568669053 | 0.000111363 |
| C10orf95 | 1.56381013 | 0.000653258 |
| GNA13 | -0.159314038 | 0.00437203 |
| KANSL1 | -0.491999106 | 0.000175231 |
| HOXB5 | -1.515849969 | 5.74E-06 |
| HOXB3 | -0.689203044 | 9.45E-07 |
| DUSP1 | 2.92234459 | 7.18E-13 |
| PANK3 | -0.727555296 | 5.31E-09 |
| MSX2 | 0.659029013 | 5.63E-05 |
| CAAP1 | -0.344205681 | 0.000583149 |
| MLANA | -1.927950217 | 0.000361319 |
| CD274 | 1.211125419 | 2.22E-06 |
| NUP43 | -0.302709259 | 0.000196111 |
| PCMT1 | 0.148348717 | 0.026402771 |
| PLEKHG1 | -1.245248256 | 0.000366084 |
| CYSTM1 | 0.667408957 | 1.84E-05 |
| ARAP3 | -0.288104023 | 0.001146557 |
| PCDHB14 | 0.447697579 | 0.038828293 |

| | | |
|---|---|---|
| MRPS14 | 0.251841555 | 0.008385578 |
| CENPL | 0.259232034 | 0.000677681 |
| SEC16B | 3.742171036 | 7.13E-06 |
| ACAT2 | 0.218603471 | 0.009438099 |
| SNX19 | -0.949891637 | 1.12E-08 |
| MSANTD2 | -1.229628532 | 6.78E-08 |
| ARR3 | 2.273185518 | 0.00016781 |
| SLC10A7 | -1.184039851 | 6.68E-06 |
| NUDCD1 | -0.725905884 | 2.36E-07 |
| ENY2 | 0.245254116 | 0.004719229 |
| MASTL | -0.174610814 | 0.0151209 |
| KIAA1217 | 0.25916407 | 0.00088676 |
| SEPTIN7P9 | 2.653066426 | 3.67E-05 |
| EPC1 | -1.506157382 | 1.68E-08 |
| CCDC77 | -0.449284292 | 4.49E-05 |
| TAF12 | 0.875552862 | 2.54E-07 |
| ENOX1 | -1.65727948 | 0.012143296 |
| MTRF1 | -1.14023434 | 9.73E-07 |
| SOHLH2 | -0.378179134 | 7.78E-05 |
| DNAJC15 | -0.596713645 | 0.002259602 |
| PROSER1 | -0.420090736 | 6.09E-05 |
| HSPH1 | -0.195833128 | 7.44E-05 |
| KBTBD7 | -1.499892509 | 2.64E-07 |
| EXOSC8 | -0.622398831 | 4.29E-06 |
| ETF1 | -0.899969743 | 3.83E-10 |
| TGFBI | 0.444586807 | 0.000160961 |
| FAM53C | 0.726130838 | 2.34E-08 |
| PAIP2 | 0.171122142 | 0.002759937 |
| KDM3B | -0.860829135 | 7.37E-09 |
| EGR1 | 4.327101506 | 6.26E-11 |
| SERP1 | 0.297553091 | 9.49E-06 |
| ZFP30 | 0.575984694 | 0.000585519 |
| NR2C1 | -0.657882062 | 5.04E-05 |
| UTP20 | -0.707986157 | 5.08E-08 |
| TMPO | -1.069390897 | 8.83E-11 |
| ARL1 | -0.672450145 | 1.20E-07 |
| MTERF2 | -0.375067905 | 0.005470999 |
| NFYB | -0.765139019 | 6.47E-07 |
| WASHC3 | 1.191805637 | 4.48E-05 |
| APAF1 | -0.482293328 | 8.10E-06 |
| DUSP4 | 0.734139149 | 2.74E-09 |
| CLU | 1.500263482 | 4.06E-09 |
| TNFRSF10B | -0.742034091 | 1.56E-08 |
| SORBS3 | -0.305824411 | 0.003575596 |
| PTK2B | 2.419020768 | 1.17E-06 |
| PPP3CC | -0.731671071 | 0.001485193 |
| RNF170 | 0.693110376 | 0.000114986 |
| UBIAD1 | -0.834951013 | 2.96E-08 |

| | | |
|---|---|---|
| TARDBP | -1.340619903 | 2.03E-11 |
| TNFRSF8 | -0.744200787 | 0.01968643 |
| LYPLA1 | -0.696298812 | 1.60E-07 |
| CRISPLD1 | -0.516352639 | 0.000214932 |
| COPS5 | -0.316772394 | 0.000108856 |
| RDH10 | -0.959880346 | 6.63E-08 |
| EPX | 3.156297303 | 0.000447662 |
| AKAP1 | -1.460555495 | 7.22E-10 |
| COIL | -1.491465714 | 5.89E-10 |
| TRIM25 | -1.12238056 | 1.84E-10 |
| SCPEP1 | 0.263040112 | 0.010578749 |
| SPOP | -1.064271746 | 5.78E-09 |
| TBX2 | -1.211188166 | 4.02E-06 |
| NACA3P | 0.387132171 | 0.000713898 |
| TEX14 | 5.517345191 | 0.009889752 |
| FAM117A | 2.533411337 | 4.34E-10 |
| NCAPH | -0.716779473 | 1.72E-06 |
| TMEM131L | -0.721453908 | 0.000352714 |
| ABCC11 | -0.496263732 | 0.038540266 |
| ADCY7 | -0.599307933 | 4.19E-06 |
| ECHDC2 | -0.593076019 | 0.000155471 |
| PSPC1 | 0.390236375 | 8.12E-05 |
| ZSCAN18 | 1.314669961 | 4.36E-06 |
| ZNF211 | 1.002254528 | 4.16E-08 |
| PDZRN3 | -0.779235017 | 8.48E-06 |
| LHX4 | -1.529234938 | 9.56E-10 |
| RNF2 | -0.786584387 | 2.04E-07 |
| TRMT1L | 0.757617283 | 6.33E-08 |
| SEC22A | -0.488382494 | 0.000365317 |
| POPDC2 | 4.027209011 | 2.90E-05 |
| B4GALT4 | -0.270248504 | 0.005566284 |
| NAA50 | -1.101751619 | 4.42E-11 |
| KIF18A | -1.859256642 | 2.53E-09 |
| DESI2 | -0.650145595 | 2.52E-08 |
| MAPK8IP1 | 0.644666232 | 9.48E-06 |
| CRY2 | 1.799171952 | 1.35E-08 |
| PEX16 | 1.338044831 | 2.66E-08 |
| DEPDC7 | -1.157819883 | 1.86E-08 |
| CAT | 0.369927406 | 8.04E-05 |
| PILRB | -0.630534231 | 8.17E-07 |
| ZMYM2 | -0.21010919 | 0.003710748 |
| GJA3 | 0.259179832 | 0.012279497 |
| TBC1D15 | 0.741786945 | 7.33E-07 |
| ADGRB2 | 0.505415806 | 0.001698703 |
| ZCCHC17 | -1.01886691 | 8.97E-09 |
| KHDRBS1 | -0.09832275 | 0.047394124 |
| TMEM39B | -0.237152306 | 0.045825417 |
| POLR3GL | 1.795026995 | 2.54E-09 |

| | | |
|---|---|---|
| ZNF639 | -1.371654509 | 2.57E-09 |
| PIK3CA | -0.22795241 | 0.004474824 |
| PDS5A | -0.691261839 | 8.96E-08 |
| TMEM156 | -1.526308903 | 0.000320945 |
| LIAS | 1.075769367 | 2.42E-06 |
| TMEM54 | 0.38351987 | 0.020482127 |
| ZSCAN20 | 1.070430198 | 7.49E-07 |
| HPCA | 0.76288963 | 0.021141184 |
| LRIF1 | 0.297716344 | 0.000198481 |
| CLCC1 | -0.547712358 | 8.54E-06 |
| GPSM2 | -1.064911335 | 2.14E-08 |
| GTDC1 | -0.720687404 | 0.000203715 |
| ZRANB3 | 0.81691516 | 0.000763099 |
| ACVR2A | 0.352597288 | 0.00666007 |
| POLK | -0.178522174 | 0.039020225 |
| SV2C | 1.408619427 | 2.65E-05 |
| FLT3 | 8.66262867 | 5.10E-08 |
| RPL21 | -0.542081281 | 1.04E-05 |
| UBL3 | -0.216627712 | 0.004361275 |
| FYTTD1 | -0.444750369 | 4.80E-06 |
| MTERF4 | -0.574246485 | 9.75E-06 |
| SASH3 | 6.913178611 | 2.40E-05 |
| OCRL | -0.833624687 | 1.70E-08 |
| PAEP | 7.4367906 | 4.07E-06 |
| MRPS2 | -0.541928107 | 9.50E-07 |
| KIAA1191 | 0.111932188 | 0.036379162 |
| COPA | 0.145711627 | 0.017300104 |
| RBBP6 | 0.540697682 | 3.76E-07 |
| ZC3H7A | -0.596088421 | 1.91E-06 |
| SERAC1 | -0.483101295 | 0.013777037 |
| LDB3 | 4.393033228 | 6.18E-09 |
| SHLD2 | -0.969952947 | 7.37E-09 |
| PRXL2A | 0.622552168 | 1.01E-05 |
| ZNF205 | -0.492931414 | 0.00404136 |
| NAA60 | -0.738741335 | 3.15E-07 |
| RPL5 | 0.271533214 | 6.47E-05 |
| ODF2L | 0.310391003 | 0.001094343 |
| SPATA1 | 2.44066778 | 0.000132932 |
| TRMT13 | -0.801156922 | 5.53E-08 |
| RWDD3 | 0.27785056 | 0.01683716 |
| ZNF644 | -0.441646458 | 3.11E-06 |
| RPAP2 | 0.355105429 | 0.000220385 |
| SLC66A2 | -0.763772135 | 1.75E-05 |
| ZMIZ2 | -0.27438531 | 0.002339824 |
| HERPUD2 | 0.37100338 | 0.000327129 |
| CBX3 | -0.759193706 | 1.51E-09 |
| HNRNPA2B1 | 0.104738378 | 0.000568036 |
| FAM126A | -0.334391052 | 7.42E-05 |

| | | |
|---|---|---|
| INHBA | 3.429278934 | 2.21E-14 |
| FKBP9 | 0.218788785 | 0.003570642 |
| NT5C3A | -0.377999803 | 0.002628693 |
| ARL4A | 0.335358709 | 0.000375559 |
| CCZ1 | -0.342141579 | 7.77E-05 |
| TWIST1 | -1.088521111 | 4.07E-05 |
| SMU1 | -0.570731761 | 1.44E-06 |
| RECK | 0.768083583 | 0.000150964 |
| TAF1L | -0.598283184 | 0.000165813 |
| ACO1 | -0.35406607 | 0.002202272 |
| PHF24 | 5.008076656 | 1.15E-07 |
| DCAF10 | -0.254782672 | 0.001226039 |
| CNTFR | 6.482415768 | 9.58E-06 |
| KIAA1549 | -0.273388246 | 0.000477766 |
| TRIM24 | -0.188108326 | 0.013146443 |
| CYREN | -0.463596696 | 2.19E-05 |
| CHST3 | -1.633806928 | 5.31E-10 |
| BICC1 | -1.390755989 | 6.27E-07 |
| CISD1 | -0.758766337 | 2.05E-06 |
| EGR2 | 3.701175093 | 1.31E-10 |
| ECD | -0.266886979 | 0.002161002 |
| P4HA1 | 0.195034894 | 0.00892117 |
| SLC25A16 | -0.470557592 | 0.004480345 |
| ZWINT | -0.575923235 | 1.43E-07 |
| VPS26A | -0.639421634 | 3.25E-07 |
| RBM19 | -0.43719538 | 0.000162912 |
| CIT | -0.48819962 | 5.27E-06 |
| ACADS | 0.68696203 | 0.006052853 |
| NME2P1 | 0.394674202 | 1.93E-05 |
| DDX54 | -0.296975117 | 0.000397759 |
| MED13L | -1.272821073 | 3.75E-10 |
| CDKN2C | -1.206642847 | 3.10E-06 |
| RNF11 | -1.004242618 | 1.19E-09 |
| RASSF8 | -1.101440801 | 3.83E-10 |
| ITPR2 | -0.368910085 | 0.000753259 |
| NECAB1 | -1.033430154 | 8.25E-06 |
| WWP1 | -1.136161753 | 9.01E-08 |
| PKN1 | 0.547266929 | 1.46E-06 |
| TRIR | -0.255959896 | 0.002000884 |
| ADGRE5 | -0.211487826 | 0.003373694 |
| WDR83 | 0.215590291 | 0.040289274 |
| GIPC1 | 0.127387944 | 0.020564897 |
| SPRYD7 | -0.366730656 | 0.003008315 |
| ATP7B | -0.834059 | 0.00067298 |
| ZC3H13 | -1.065698717 | 5.94E-10 |
| NLN | -0.816271912 | 9.95E-08 |
| CENPK | -0.543537857 | 0.00015539 |
| OPTN | 0.479390516 | 0.014882226 |

| | | |
|---|---|---|
| ITIH5 | 0.928352515 | 0.005179939 |
| ATF1 | -1.171266658 | 4.09E-08 |
| TSFM | -0.423766529 | 0.000192755 |
| ARHGAP9 | 1.68482472 | 0.001582323 |
| NCKAP1L | 7.558290931 | 3.36E-06 |
| MMP19 | 1.471189276 | 1.26E-06 |
| PFDN5 | 0.179535755 | 0.003510867 |
| SPATS2 | -0.628482622 | 3.46E-06 |
| NR4A1 | 5.027813216 | 6.97E-16 |
| HOXC13 | -0.766886744 | 6.79E-06 |
| CDK2 | -0.485063902 | 5.56E-07 |
| LRP1 | 0.554078865 | 8.62E-07 |
| HOXC11 | -1.864955106 | 6.56E-06 |
| HOXC12 | -0.880472876 | 0.000158866 |
| IKZF4 | -0.821562093 | 4.56E-05 |
| SMUG1 | -1.151818451 | 1.83E-07 |
| TUBA1B | -0.286872456 | 2.07E-06 |
| EEF1AKMT3 | 1.205744306 | 1.18E-06 |
| KBTBD4 | -0.922083712 | 4.34E-08 |
| ATPAF1 | -0.494571374 | 8.35E-06 |
| STIL | -0.984290542 | 2.80E-09 |
| HJURP | -1.92852536 | 2.58E-10 |
| IL13RA2 | 0.376181611 | 8.18E-06 |
| AMD1 | -2.004175054 | 3.14E-11 |
| NDUFAF4 | -0.684862876 | 3.55E-06 |
| USP45 | -0.337941975 | 0.009140377 |
| PLP1 | 1.261529505 | 1.34E-09 |
| MORF4L2 | 0.348777414 | 1.19E-06 |
| FAM199X | -0.647917342 | 2.34E-07 |
| METTL8 | -0.602851164 | 0.000530111 |
| SLC36A1 | -1.693465279 | 3.05E-10 |
| LPGAT1 | -1.024124677 | 1.95E-09 |
| KCNJ2 | 0.832008915 | 6.06E-05 |
| RAP2C | -0.642749545 | 2.99E-07 |
| EXOSC9 | -1.062647315 | 2.56E-07 |
| PLA2G12A | -0.35629778 | 0.001030786 |
| COQ8B | 0.721429143 | 7.75E-06 |
| PFKFB2 | 0.553177593 | 1.12E-05 |
| ZNF137P | 1.290627176 | 1.14E-05 |
| RAB38 | -0.739203593 | 2.14E-06 |
| GPR83 | 3.184218022 | 8.95E-06 |
| ago-02 | -0.340638244 | 0.000741797 |
| MXD4 | 1.291636457 | 3.05E-08 |
| CKS2 | -0.613863673 | 2.49E-07 |
| ACSL3 | 1.377065707 | 2.60E-11 |
| CHPF | 0.292918835 | 0.000123015 |
| MOGAT1 | 7.74296429 | 1.44E-06 |
| OBSL1 | 0.780254791 | 6.87E-08 |

| | | |
|---|---|---|
| ENKD1 | 0.505831873 | 0.002750054 |
| TTPAL | -0.902482624 | 4.19E-09 |
| PREX1 | 0.285034141 | 0.003263516 |
| SLC12A5 | 3.912125263 | 0.003071602 |
| ARHGAP40 | 3.988400859 | 0.002335803 |
| SDC4 | 0.462552787 | 1.66E-06 |
| NCOA3 | -0.902622459 | 1.62E-08 |
| PIGT | 0.143386205 | 0.023324641 |
| MATN4 | 2.175779763 | 2.94E-05 |
| NCOA5 | -0.350070354 | 6.96E-05 |
| VAPB | -1.049543276 | 1.41E-09 |
| PARD6B | 0.833852533 | 0.000204402 |
| CHD6 | -0.377344176 | 2.77E-05 |
| PLCG1 | -0.172719474 | 0.002864594 |
| SRSF6 | -1.622943398 | 3.86E-12 |
| ARFGEF2 | -0.329569799 | 2.72E-05 |
| ZNFX1 | -1.177746605 | 6.28E-10 |
| CSE1L | -0.462549177 | 1.66E-07 |
| TMEM189-UBE2V1 | -0.231793661 | 0.000131438 |
| RAB22A | -0.315019064 | 0.000173394 |
| STAU1 | -0.544901765 | 9.21E-08 |
| CDH26 | 4.422857659 | 0.001043831 |
| SNAI1 | 1.724893996 | 4.34E-10 |
| MOCS3 | 0.424390104 | 4.76E-05 |
| STX16 | 0.24015052 | 0.000119333 |
| PPP4R1L | 0.472338697 | 0.00186204 |
| RNF114 | -0.389560509 | 3.30E-05 |
| RBPJL | 7.640261242 | 2.09E-06 |
| PCK1 | 8.262728337 | 3.14E-13 |
| NEURL2 | 0.988949293 | 0.02396551 |
| MTRR | -0.603374174 | 1.00E-06 |
| FASTKD3 | 0.146596807 | 0.040140722 |
| PEPD | 0.273543177 | 0.012050702 |
| IQSEC2 | 0.615039946 | 0.000863112 |
| NAGK | -0.308813492 | 0.003658176 |
| MCEE | 0.763244016 | 0.000830553 |
| PAIP2B | 2.37571556 | 2.22E-07 |
| SNRNP27 | -1.057701162 | 4.42E-08 |
| MPHOSPH10 | -0.13111712 | 0.021801066 |
| IL17C | 3.893995369 | 7.38E-05 |
| USP22 | -0.466565444 | 2.42E-07 |
| ZNF576 | -1.645818272 | 1.61E-07 |
| IRGC | 7.495806202 | 3.41E-06 |
| LYPD3 | 2.986404479 | 8.54E-09 |
| USP9X | -0.262053968 | 0.000109971 |
| TRERF1 | -0.509464656 | 0.002009052 |
| PACSIN1 | 3.624231469 | 6.54E-05 |
| BTN2A2 | -0.541069308 | 0.000276252 |

| | | |
|---|---|---|
| SIRT5 | 1.282635041 | 4.57E-06 |
| MRS2 | -0.453862441 | 4.01E-05 |
| WRNIP1 | -0.277513449 | 0.002780754 |
| RRP36 | -0.762392225 | 2.37E-07 |
| BTN2A3P | -0.394436921 | 0.033271677 |
| BTN1A1 | 3.765849431 | 7.19E-06 |
| SNRPC | 0.1429378 | 0.041581586 |
| SERPINB6 | 0.239840303 | 0.003961204 |
| XPO5 | -0.618834235 | 3.77E-07 |
| ABCC10 | -0.425094955 | 0.001477539 |
| H1-3 | 4.544234758 | 9.64E-08 |
| PEX6 | 1.039427834 | 8.94E-08 |
| NQO2 | 0.490444445 | 1.34E-05 |
| OARD1 | 1.123826926 | 8.80E-08 |
| UNC5CL | 3.761563212 | 1.25E-08 |
| ZNF391 | 0.376999873 | 0.014058379 |
| RPS10 | 0.152985199 | 0.015153069 |
| MOCS1 | -0.431989505 | 0.003655009 |
| H2BC11 | 2.555862192 | 1.53E-06 |
| MED20 | -2.154565458 | 1.58E-10 |
| SPDEF | 5.331344526 | 8.47E-05 |
| TCP11 | 6.873317406 | 2.68E-05 |
| MAD2L1BP | 0.184032396 | 0.019443994 |
| APOBEC2 | 7.083879035 | 1.24E-05 |
| KLHDC3 | -0.297814127 | 0.00072668 |
| GNMT | 1.569322204 | 0.001472321 |
| MEA1 | 0.417739197 | 1.90E-05 |
| KLHL31 | 0.874464353 | 5.75E-08 |
| CDKN1A | 1.386306711 | 9.82E-11 |
| SOX4 | -0.83857023 | 5.26E-06 |
| GLO1 | -0.438968205 | 9.24E-07 |
| SSR1 | -0.78315103 | 8.68E-09 |
| RIOK1 | -0.697054368 | 4.04E-07 |
| NRN1 | 0.901574412 | 0.000244375 |
| SLC35B3 | 0.922400549 | 1.46E-05 |
| NUP153 | -1.360676359 | 1.30E-10 |
| DEK | -0.766556177 | 4.02E-09 |
| EEF1E1 | -0.574878414 | 4.06E-05 |
| LRRFIP1 | -0.382435647 | 9.06E-06 |
| EREG | 3.922295046 | 0.015823307 |
| MYRF | 0.453386101 | 0.005841769 |
| AHNAK | -0.484892874 | 7.24E-09 |
| EMC3 | -0.271734771 | 0.001068882 |
| SSUH2 | 1.499448642 | 1.05E-06 |
| SH3TC1 | -1.837648013 | 3.10E-07 |
| CNOT1 | -0.518690688 | 2.36E-07 |
| BBS2 | 0.221697403 | 0.03195515 |
| MT2A | 0.814324168 | 2.42E-09 |

| | | |
|---|---|---|
| GOT2 | -0.096572357 | 0.049048309 |
| TMTC4 | -0.353853093 | 0.026763681 |
| RAP2A | -0.219493074 | 0.000836989 |
| ABCC4 | -0.953708448 | 3.59E-08 |
| EFNB2 | 0.279385469 | 0.000130871 |
| HROB | 0.623216446 | 4.14E-05 |
| UPF3B | -0.54675662 | 1.19E-05 |
| RNF113A | -0.4004207 | 0.005970709 |
| SEPTIN6 | 0.581167096 | 4.66E-05 |
| BMP4 | 1.342136858 | 0.001064393 |
| FAM193A | -1.271027719 | 9.01E-08 |
| GRK4 | -0.444352573 | 0.015817349 |
| TEKT3 | 2.42193165 | 0.000134113 |
| MYH2 | 1.917576443 | 7.07E-05 |
| HS3ST3B1 | -0.729200051 | 0.000589232 |
| MRPS7 | -0.513714917 | 4.16E-06 |
| GGA3 | -0.364914538 | 0.000137711 |
| ARMC7 | -1.586040832 | 2.35E-08 |
| NUP85 | -0.541072679 | 7.70E-06 |
| SLC25A19 | -0.37813617 | 0.000463276 |
| MIF4GD | 0.647555433 | 7.23E-05 |
| NT5C | 0.557508073 | 1.39E-05 |
| MSTO1 | -0.18848003 | 0.024466306 |
| C1orf61 | 1.459884561 | 0.000126303 |
| TTF1 | -0.652784414 | 5.44E-07 |
| GTF3C4 | -0.526377255 | 2.10E-07 |
| DDX31 | -0.837764281 | 5.82E-07 |
| PPP1R12C | -0.235135005 | 0.020234998 |
| OPRL1 | 2.997868371 | 0.022596606 |
| PPDPF | 0.48797841 | 4.43E-05 |
| INSIG2 | -0.815865351 | 2.95E-05 |
| POLR1B | -1.182847039 | 5.34E-10 |
| CCDC93 | -1.456243363 | 2.97E-09 |
| SLC25A23 | 0.670756213 | 6.51E-05 |
| ALKBH7 | 0.555291407 | 0.005086572 |
| TNFSF9 | 1.290634334 | 8.38E-09 |
| THOC2 | -0.180984172 | 0.003280797 |
| MED1 | -0.750085919 | 5.36E-09 |
| RPL23 | 0.328355729 | 2.34E-06 |
| AC046185.1 | -0.47760695 | 0.000346671 |
| ATG4C | -0.407975412 | 0.004707502 |
| CD70 | 0.990633537 | 1.09E-08 |
| C3 | 0.991784961 | 0.000381032 |
| TRIP10 | 0.252342712 | 0.006919177 |
| GPR108 | 0.972395733 | 1.58E-07 |
| TNFSF14 | 5.418012137 | 3.55E-05 |
| FOSB | 3.093991491 | 6.27E-13 |
| OPA3 | -0.545431148 | 6.40E-06 |

| | | |
|---|---|---|
| SNRPD2 | 0.336981461 | 1.20E-05 |
| RTN2 | 1.294727797 | 5.18E-05 |
| EML2 | 0.670898506 | 0.000174942 |
| GPCPD1 | 0.933299052 | 1.28E-07 |
| SDCBP2 | 2.85714391 | 6.45E-08 |
| PANK2 | -0.151001884 | 0.017831616 |
| FAM182A | 6.835233836 | 1.10E-05 |
| GZF1 | 0.664955849 | 2.92E-07 |
| NAPB | -0.776590628 | 1.65E-05 |
| NKX2-4 | -3.307094747 | 3.18E-06 |
| CENPB | -0.627997184 | 1.28E-07 |
| PSMF1 | -0.435412373 | 4.97E-06 |
| NKX2-2 | -1.469651333 | 0.006905623 |
| DTD1 | 0.215708572 | 0.00469259 |
| TMX4 | -0.295810733 | 2.33E-05 |
| STK35 | -2.049817907 | 2.21E-11 |
| NRSN2 | 0.49366677 | 5.61E-06 |
| AP5S1 | -1.001596117 | 1.71E-06 |
| BMP2 | 0.554425462 | 0.000866551 |
| ZNF133 | -0.416297771 | 0.001879651 |
| FLRT3 | 0.661760123 | 1.45E-07 |
| PCSK2 | 2.777629968 | 0.001788632 |
| MKKS | -0.303541107 | 0.000513123 |
| BFSP1 | 3.540472993 | 1.77E-10 |
| SNRPB2 | 0.229191842 | 0.000192133 |
| TBC1D20 | -1.122374201 | 1.05E-08 |
| ITPA | -0.421823692 | 0.000465808 |
| TCF15 | 2.239413625 | 0.003122871 |
| MCM8 | -0.604436339 | 4.35E-07 |
| MRPS26 | -0.430934943 | 1.17E-05 |
| NCLN | -0.835839147 | 5.32E-09 |
| HNRNPR | -0.461736051 | 7.05E-08 |
| ZNF436 | 0.923411305 | 6.84E-08 |
| MAX | -0.914521805 | 1.24E-07 |
| ARMCX5 | -1.210080265 | 5.55E-07 |
| GDF5 | 7.054961648 | 1.25E-06 |
| MMP24 | 0.58597323 | 0.007054229 |
| NECAB3 | 0.591983343 | 0.000214222 |
| ID1 | 0.221345956 | 0.006562771 |
| ROMO1 | 0.335076238 | 0.001629655 |
| FAM83C | 3.807815528 | 5.23E-06 |
| CEP250 | -0.812462059 | 2.65E-08 |
| PLAGL2 | -0.327171236 | 0.000185159 |
| MMP24OS | 0.68617892 | 1.40E-05 |
| KDM5C | -0.112346088 | 0.039517598 |
| AMOT | -0.265436997 | 0.000482934 |
| TMEM115 | -0.747779991 | 4.74E-08 |
| PSMB2 | -0.143221497 | 0.009608122 |

| | | |
|---|---|---|
| ago-03 | -0.508382713 | 1.25E-05 |
| ST3GAL3 | 1.142783053 | 1.21E-05 |
| TMEM53 | 0.497750737 | 0.006062706 |
| HECTD3 | -0.269767396 | 0.001386991 |
| TUBGCP3 | -1.14774667 | 5.07E-09 |
| MCF2L | 1.679273539 | 5.59E-10 |
| F10 | 3.347620013 | 8.19E-05 |
| PCID2 | 1.441219939 | 5.59E-11 |
| PROZ | 2.897584839 | 3.51E-05 |
| IGFLR1 | 1.693841948 | 5.82E-07 |
| CAPNS1 | 0.331022633 | 6.35E-05 |
| PDCD2L | -0.494307457 | 0.004297379 |
| UBA2 | -0.512029601 | 3.87E-07 |
| FFAR1 | 6.062511383 | 4.89E-06 |
| COX6B1 | 0.257902051 | 0.003649624 |
| THRA | -0.481714047 | 0.00025781 |
| NR1D1 | 1.663700583 | 2.80E-08 |
| FRMD8 | -0.665263513 | 5.46E-07 |
| PRDX5 | 0.475234173 | 3.64E-05 |
| BCL2L12 | 0.255169345 | 0.008317145 |
| IRF3 | 0.49040169 | 3.22E-05 |
| PRMT1 | 0.377667828 | 3.78E-05 |
| RRAS | 0.627793102 | 0.001290292 |
| PRRG2 | 1.534357357 | 0.003301585 |
| SCAF1 | -0.552195643 | 0.000326162 |
| PRR12 | -0.539953615 | 5.07E-06 |
| SBDS | -0.259374104 | 0.000154578 |
| WNK4 | -1.077632993 | 2.31E-09 |
| BECN1 | -0.170317504 | 0.006665114 |
| PRKCG | 2.658787842 | 0.005141414 |
| TRAP1 | 0.259485564 | 0.00132667 |
| DNAJC8 | -0.388689044 | 1.86E-05 |
| AHDC1 | -0.371836162 | 5.19E-05 |
| ZNF384 | -0.618047479 | 6.69E-07 |
| EMG1 | -0.353290454 | 6.16E-05 |
| CFP | 1.770315377 | 1.09E-05 |
| TIMM17B | 0.282269603 | 0.010316201 |
| PCNX4 | -1.134335843 | 7.45E-10 |
| ATG14 | -1.658618918 | 2.35E-08 |
| SIX1 | -0.426933777 | 0.000243214 |
| RHOJ | -2.349410564 | 9.54E-08 |
| DLGAP5 | -0.947387007 | 4.94E-07 |
| L3HYPDH | -0.254519172 | 0.031277687 |
| ZBTB1 | 0.234762669 | 0.001420962 |
| TRMT5 | 0.699020881 | 5.01E-07 |
| SGPP1 | -1.513776556 | 8.30E-08 |
| PLEKHG3 | -1.973353624 | 2.96E-08 |
| RHOT1 | -0.46734491 | 4.52E-05 |

| | | |
|---|---|---|
| OMG | 1.648741478 | 0.01636512 |
| WDR60 | 0.988895872 | 1.93E-06 |
| FAM78A | -1.549926635 | 6.40E-08 |
| NUP214 | -1.327848855 | 9.33E-10 |
| CTAG2 | 0.667996171 | 0.00117581 |
| SLC10A3 | -0.543222404 | 1.58E-05 |
| HNRNPH2 | -0.482570072 | 9.34E-07 |
| TIMM8A | -0.539452901 | 7.65E-05 |
| ZC4H2 | 0.387833612 | 0.035062247 |
| CANX | 0.220981313 | 5.34E-06 |
| IPPK | -1.171386667 | 3.50E-09 |
| ZNF484 | 0.692394697 | 9.14E-05 |
| HIVEP3 | -0.130775326 | 0.035221296 |
| COX7C | 0.436664535 | 5.22E-05 |
| ABHD8 | 0.794054484 | 2.77E-05 |
| HELB | 0.59879138 | 9.84E-06 |
| RAP1B | -0.288060151 | 4.84E-05 |
| RAB3IP | -0.568239082 | 5.21E-05 |
| DYRK2 | -1.04338163 | 3.94E-09 |
| YEATS4 | -0.376325909 | 0.000211266 |
| TAS2R4 | 2.614895922 | 2.65E-06 |
| TAS2R5 | 2.206452674 | 0.000223312 |
| IDUA | 0.613098768 | 0.002014206 |
| FGFRL1 | 0.453323903 | 2.29E-06 |
| TMEM175 | 1.048971119 | 6.94E-06 |
| AUNIP | -0.804749479 | 2.61E-06 |
| FBXL12 | 0.708997526 | 5.11E-07 |
| EMC1 | -0.275599453 | 9.90E-05 |
| UBR4 | 0.108715263 | 0.024237401 |
| SIN3B | -0.471920938 | 2.94E-06 |
| SLC35E1 | -0.914703659 | 1.79E-09 |
| EPS15L1 | -0.289323809 | 0.001145623 |
| GFER | -0.764952741 | 2.66E-06 |
| PKMYT1 | -0.378409233 | 5.18E-05 |
| WFIKKN1 | 1.867402046 | 4.24E-05 |
| WDR24 | 0.473357377 | 0.000215833 |
| CHTF18 | 0.196865353 | 0.007014154 |
| MACF1 | -0.541190129 | 4.01E-08 |
| SMARCA4 | -0.261502651 | 0.000166216 |
| KDM4B | -0.374063742 | 0.002489315 |
| IL17B | 7.307167248 | 6.53E-06 |
| EMC6 | -0.210803364 | 0.022871747 |
| METTL16 | -1.621378803 | 1.54E-10 |
| TUBA4A | 1.148883955 | 0.009905242 |
| VIL1 | -0.955866162 | 7.27E-07 |
| RNF6 | -0.622170827 | 1.17E-07 |
| AKAP9 | -0.777344076 | 1.40E-08 |
| GNG11 | -0.705737357 | 1.59E-06 |

| | | |
|---|---|---|
| SEM1 | 0.477292174 | 1.15E-05 |
| HIP1 | 0.54748946 | 5.30E-07 |
| PTPN12 | -1.248822549 | 4.67E-10 |
| POR | 0.198863082 | 0.010121143 |
| STYXL1 | 0.664180823 | 0.000937791 |
| GNAI1 | -0.887500948 | 7.39E-06 |
| PEX1 | 1.346086974 | 9.14E-09 |
| MTERF1 | -0.660032707 | 1.71E-06 |
| CASD1 | -0.396453945 | 0.001599356 |
| ZNF780B | -0.419267924 | 0.0001016 |
| ZFP36 | 3.338206794 | 2.69E-13 |
| RASL11B | 3.806086894 | 1.43E-07 |
| PAICS | -0.154833457 | 0.000556745 |
| TUBGCP6 | -0.664713858 | 1.95E-05 |
| ADM2 | 1.764752034 | 1.70E-06 |
| DGCR6L | 0.441492263 | 0.001652314 |
| DGCR8 | -0.73688751 | 2.97E-07 |
| GAL3ST1 | 5.48554381 | 2.51E-05 |
| YWHAH | -1.304880174 | 1.71E-10 |
| RFPL1 | 7.305640991 | 6.09E-06 |
| GNAZ | 0.781830971 | 0.034745512 |
| MGAT3 | 1.983272062 | 8.82E-06 |
| ADORA2A | 0.640729504 | 0.03609978 |
| ATF4 | 0.450331743 | 8.27E-08 |
| RFPL3 | 3.593181612 | 7.39E-05 |
| CDC42EP1 | -1.771772429 | 5.43E-07 |
| MCHR1 | 10.17715623 | 4.70E-10 |
| TPST2 | -0.486235765 | 0.003615483 |
| BAIAP2L2 | 0.392537981 | 0.000747939 |
| MPST | 0.302140178 | 0.013089557 |
| TST | 0.798835543 | 4.81E-06 |
| APOL2 | -1.906313446 | 2.40E-08 |
| LIF | -1.70605459 | 9.98E-11 |
| APOBEC3F | 1.097977653 | 8.02E-05 |
| RIBC2 | 7.338256748 | 5.60E-06 |
| RNF112 | 4.25832729 | 1.62E-05 |
| CPA4 | -0.484259965 | 3.17E-05 |
| POT1 | 0.322091818 | 0.002532054 |
| ATP6V1F | 0.17733662 | 0.015710123 |
| LSM8 | -0.524080444 | 2.41E-05 |
| PRKRIP1 | -0.711095458 | 3.81E-06 |
| VGF | 2.079541953 | 2.46E-09 |
| PODXL | -0.441002863 | 3.68E-06 |
| STRIP2 | -0.397216807 | 0.00241705 |
| MKLN1 | 0.182281332 | 0.009773547 |
| DNAJB9 | 1.923515105 | 5.58E-11 |
| FLNC | 8.013616312 | 0.004018138 |
| CALU | -0.557672486 | 1.78E-08 |

| | | |
|---|---|---|
| CCDC136 | -0.45803936 | 0.000525454 |
| SMO | -0.570399547 | 9.75E-06 |
| IRF5 | 1.164753505 | 0.000637495 |
| LRRC17 | 1.924662757 | 7.67E-05 |
| KLHDC10 | -2.040093954 | 5.30E-11 |
| NDUFA5 | -0.322461139 | 0.000186895 |
| OPN1SW | 0.531132861 | 0.038131774 |
| MRPS12 | -0.559043791 | 2.76E-06 |
| MYO1B | -0.489339494 | 7.45E-05 |
| MTX2 | -0.423093626 | 0.000500353 |
| CHN1 | -0.588111888 | 0.00153172 |
| OSGEPL1 | 0.447307909 | 0.002631107 |
| ORMDL1 | -0.161100098 | 0.031230543 |
| HAT1 | -0.123334298 | 0.049683011 |
| HOXD13 | -2.291568954 | 1.11E-08 |
| HERC2 | -0.140735032 | 0.039534201 |
| SNRPN | 0.356410776 | 8.95E-05 |
| TWSG1 | -0.44675897 | 4.40E-05 |
| ARHGAP22 | -0.731434031 | 0.038748639 |
| EIF2AK4 | -0.304073426 | 0.000161702 |
| MYO5C | 0.783207482 | 0.035565707 |
| TTBK2 | 0.307137647 | 0.000201612 |
| CCDC32 | -0.194059353 | 0.041591183 |
| INO80 | -0.874298248 | 5.27E-08 |
| ICE2 | -0.324007547 | 0.00024425 |
| DLL4 | 3.341926019 | 1.76E-07 |
| MINDY2 | -0.456327692 | 4.44E-05 |
| IVD | -0.866858179 | 4.89E-08 |
| KNSTRN | -1.14103537 | 2.91E-09 |
| CHAC1 | 1.698786131 | 5.20E-09 |
| CLN6 | -0.687449449 | 1.52E-07 |
| ARPP19 | -0.651541816 | 8.61E-09 |
| VPS13C | 0.444589283 | 3.11E-05 |
| CALML4 | -0.503966747 | 0.000789125 |
| THAP10 | 1.042520025 | 2.95E-07 |
| ANAPC13 | -0.577156989 | 9.29E-07 |
| MBD4 | 0.442181092 | 5.86E-06 |
| COPB1 | -0.296159831 | 0.000239377 |
| SUMF2 | 0.518023433 | 1.52E-06 |
| PALLD | -0.251769996 | 0.031108017 |
| SPCS3 | -1.133840942 | 2.91E-09 |
| TPH1 | 4.212492453 | 1.81E-09 |
| E2F8 | -1.675237627 | 6.20E-07 |
| DCTD | -0.417580525 | 4.00E-06 |
| SOX15 | 0.55782441 | 0.009207748 |
| PIMREG | -0.688973494 | 5.74E-06 |
| USP6 | 2.31619013 | 5.50E-08 |
| SHBG | 4.28613597 | 0.001593255 |

| | | |
|---|---|---|
| CD68 | 0.511816278 | 9.91E-05 |
| TXNDC17 | 0.243407481 | 0.006899226 |
| FXR2 | -0.308274715 | 6.13E-05 |
| KIF1C | -0.587495313 | 4.91E-07 |
| MPDU1 | -0.193726517 | 0.005069899 |
| PHF20L1 | -0.40209404 | 1.25E-05 |
| LRRC6 | 1.480274875 | 0.005174273 |
| CCNT1 | 0.207166607 | 0.000449021 |
| PUS7L | -1.439734565 | 2.92E-09 |
| KRI1 | -0.515773502 | 9.43E-07 |
| ILF3 | -0.370973093 | 2.16E-07 |
| SLC44A2 | 0.407185708 | 0.000168266 |
| CDKN2D | 0.325495482 | 0.001644245 |
| KLK14 | 3.201047786 | 2.12E-06 |
| NGDN | -0.32274456 | 0.000393672 |
| RAB2B | -0.423185908 | 0.000370252 |
| BCL2L2 | -0.673629462 | 3.84E-07 |
| AJUBA | -2.455476966 | 1.81E-11 |
| DTD2 | -0.803396062 | 7.25E-05 |
| PARP2 | -0.959531733 | 1.55E-07 |
| FOXA1 | -0.468789452 | 0.046250276 |
| EAPP | 0.453007304 | 0.000113267 |
| MIS18BP1 | -1.101359898 | 5.13E-09 |
| NRL | 1.802799655 | 0.000151535 |
| NEDD8 | 0.182025605 | 0.028532807 |
| TEP1 | 0.228744053 | 0.007110111 |
| EPB41L4A | 0.589557717 | 0.00026728 |
| CDO1 | 1.388588013 | 3.38E-05 |
| REEP5 | -0.423230023 | 7.32E-06 |
| SEC14L1 | -0.362073682 | 2.55E-05 |
| RHBDF2 | 0.764781264 | 0.000138104 |
| AANAT | 1.986727755 | 0.00360126 |
| MAP7D3 | -1.003783754 | 1.55E-07 |
| FGF13 | -0.847353257 | 2.45E-06 |
| ASH2L | -0.375967695 | 0.000149221 |
| TTI2 | -0.366669789 | 0.000145435 |
| CDKN1C | 1.602797072 | 3.12E-05 |
| SGO1 | -1.286294619 | 3.23E-09 |
| DOHH | -0.529855339 | 0.000630146 |
| SYT5 | 1.066920873 | 0.00674616 |
| GAMT | 0.875490883 | 1.03E-05 |
| PHF10 | -0.536813415 | 0.000498595 |
| CRACR2A | -0.82456983 | 3.15E-06 |
| STARD8 | 2.338651196 | 1.10E-07 |
| SAT1 | 2.565622731 | 1.98E-10 |
| GNL3L | -1.078750847 | 1.82E-06 |
| SH3BP4 | -1.472620734 | 4.98E-10 |
| MOSPD2 | 0.494770716 | 3.12E-05 |

| | | |
|---|---|---|
| DOCK6 | 0.408549971 | 0.005809683 |
| ECSIT | 0.357315078 | 0.00049828 |
| LDLR | 2.214300681 | 2.13E-13 |
| ELOF1 | -0.173866917 | 0.003346138 |
| PRKCSH | 0.198062133 | 0.000157073 |
| THEM6 | 0.270326942 | 0.008538607 |
| NECTIN2 | 0.585647599 | 8.60E-06 |
| TOMM40 | -0.29744872 | 0.000163541 |
| APOC1 | 1.391365459 | 0.013181385 |
| XPO7 | -0.410290067 | 7.71E-05 |
| ACE2 | 3.835131525 | 0.003119111 |
| SAFB2 | -0.182606775 | 0.028559429 |
| RPL36 | 0.407611945 | 4.78E-06 |
| ATP8B3 | 1.458534995 | 0.000643717 |
| GDF1 | 0.827763066 | 0.000627162 |
| KIF1A | -0.394200874 | 0.006253419 |
| BST2 | 1.415040154 | 0.023022488 |
| SLC27A1 | 0.59044951 | 0.001343326 |
| NSUN5 | -0.153790814 | 0.031705831 |
| COLGALT1 | -0.267528043 | 3.33E-05 |
| MRPL34 | -0.315749376 | 0.000311414 |
| PGLS | 0.391234784 | 0.000414569 |
| SNX9 | -1.020162652 | 9.16E-08 |
| RTN4IP1 | -1.339738816 | 1.91E-05 |
| QRSL1 | -0.535137865 | 0.000166735 |
| MTRES1 | 0.299125609 | 0.026846009 |
| RSPH3 | 0.601676717 | 0.000498241 |
| MLLT1 | -0.470980685 | 4.88E-06 |
| AFDN | -0.702284622 | 3.02E-06 |
| ACTN4 | -0.442792185 | 1.86E-07 |
| STK33 | 6.152608084 | 9.00E-05 |
| NDUFA10 | -0.471354008 | 0.000275573 |
| ARPC1B | 0.364911762 | 1.68E-05 |
| ZSWIM6 | 0.780972003 | 8.24E-08 |
| FCHO1 | 2.92580974 | 2.45E-06 |
| UNC13A | 0.572821112 | 0.001303341 |
| MAP1S | -0.244389537 | 0.002097599 |
| GDF15 | 5.0487409 | 1.97E-07 |
| PGPEP1 | 1.423182364 | 2.05E-09 |
| IQCN | 8.951867007 | 5.85E-11 |
| LSM4 | 0.352558147 | 2.17E-05 |
| JUND | 1.240615538 | 5.17E-09 |
| HRC | 5.334705712 | 0.000169044 |
| TRPM4 | 0.857260612 | 7.10E-05 |
| ZNF557 | 0.76521195 | 5.02E-06 |
| OLFM1 | 4.756188432 | 0.000408606 |
| CAMSAP1 | -0.658869218 | 5.50E-07 |
| UBAC1 | -0.302448377 | 0.004546924 |

| | | |
|---|---|---|
| SAG | 8.823941634 | 4.07E-08 |
| ZBTB46 | 0.293272562 | 0.022942895 |
| HELZ2 | 0.390513876 | 4.34E-05 |
| CYP2E1 | 2.658524156 | 3.56E-07 |
| PNPLA7 | 0.614053009 | 0.007443944 |
| PAK4 | -0.693142032 | 9.88E-07 |
| ZNF337 | -0.466168636 | 4.82E-05 |
| CEP85 | -0.43849392 | 7.40E-06 |
| TAF4 | -1.243910186 | 1.19E-08 |
| EXOSC2 | -0.683995538 | 1.70E-07 |
| UCK1 | -0.339938634 | 0.000112809 |
| FIBCD1 | 3.158081919 | 5.43E-05 |
| PRRC2B | -0.649564034 | 1.56E-08 |
| CHMP2A | 0.427800915 | 3.23E-05 |
| TRIM28 | -0.206320724 | 1.75E-05 |
| METTL26 | 0.306004597 | 0.00055938 |
| ATG4D | -0.474936655 | 3.85E-05 |
| ZC3H4 | 1.298268326 | 3.10E-10 |
| MAP3K10 | 0.463496141 | 0.010975343 |
| LRRC47 | -0.652112019 | 1.27E-07 |
| SESN2 | 2.257860429 | 1.93E-10 |
| ATP5IF1 | 0.340898839 | 0.000107258 |
| MED18 | 0.300450949 | 0.006361993 |
| THEMIS2 | 1.541151896 | 2.12E-07 |
| CLIP1 | -0.697581287 | 9.42E-08 |
| CCDC62 | 3.537341492 | 8.07E-07 |
| HIP1R | 0.193079995 | 0.022777241 |
| ZNF317 | 0.155507387 | 0.011634437 |
| PPAN | -1.006770312 | 1.24E-08 |
| ANGPTL6 | 0.50012628 | 0.04326625 |
| SLC6A8 | -0.159062188 | 0.007567793 |
| PNCK | 1.719993968 | 0.010784523 |
| DKC1 | -0.656203753 | 2.87E-08 |
| ZNF331 | -0.556992677 | 2.03E-06 |
| ZNF236 | -0.860604058 | 1.70E-05 |
| C12orf65 | 0.694241462 | 1.43E-06 |
| NOL11 | -0.924465681 | 3.32E-09 |
| UBE4B | -0.752634176 | 7.40E-08 |
| CASZ1 | 0.485250227 | 0.025909071 |
| PKDREJ | 5.555802322 | 6.31E-10 |
| HABP4 | 0.404001322 | 0.003852836 |
| FBP2 | 0.972747996 | 0.048435277 |
| SLC35D2 | -0.519590924 | 0.00315331 |
| UBA1 | 0.162210013 | 0.003284949 |
| POLN | 2.672699341 | 8.05E-07 |
| PPIL4 | -1.077940028 | 3.16E-07 |
| ULBP2 | 0.519382803 | 0.001893935 |
| AKAP12 | 0.289075169 | 4.71E-06 |

| | | |
|---|---|---|
| SYNE1 | -0.558241845 | 2.00E-05 |
| ULBP3 | -0.494164778 | 0.016228689 |
| LATS1 | -1.138906761 | 3.18E-09 |
| EPS8L1 | 2.026646289 | 0.006152118 |
| AAR2 | -0.273204658 | 0.000624686 |
| TTLL9 | 2.103583813 | 0.00021625 |
| RBM39 | -0.766829611 | 5.80E-09 |
| ZNF341 | -1.655390444 | 4.82E-06 |
| ACSS2 | 1.217060432 | 1.27E-08 |
| EDA2R | 0.693493017 | 3.51E-05 |
| ARHGEF9 | 0.747495201 | 3.98E-06 |
| C1QL1 | -0.762851368 | 0.014151019 |
| GFAP | 1.493428986 | 0.016041115 |
| PYY | 3.73716532 | 0.000139952 |
| HIGD1B | -2.400119351 | 0.007849615 |
| ZNF227 | -0.508328434 | 5.21E-06 |
| COX4I1 | 0.144248693 | 0.038967051 |
| EMC8 | -1.231725164 | 2.24E-09 |
| GSE1 | -0.199005092 | 0.016400323 |
| GINS2 | -0.580034392 | 1.00E-06 |
| CHMP1A | -0.667729194 | 3.76E-07 |
| COX7B | 0.198897237 | 0.042161469 |
| F12 | 0.655774179 | 0.004572317 |
| NFATC1 | -0.398357999 | 0.031435112 |
| GJA9 | 2.250624347 | 0.000206998 |
| CAP1 | 0.111860593 | 0.01382147 |
| PPT1 | -0.311103461 | 1.00E-05 |
| RAB11FIP4 | 1.483347289 | 0.000140478 |
| RLIM | -0.335630435 | 1.09E-05 |
| TRAF3 | -1.612683734 | 7.72E-10 |
| HAUS8 | 0.595017061 | 0.000167503 |
| MRPS25 | -0.154450382 | 0.008438155 |
| HACL1 | -0.547904474 | 0.000371786 |
| CAPN7 | -0.825913586 | 2.26E-08 |
| RFTN1 | -0.948006551 | 2.55E-05 |
| C3orf20 | 7.467967279 | 3.75E-06 |
| RBSN | -0.453943419 | 3.56E-05 |
| GALNT15 | 6.870237481 | 2.35E-05 |
| SLC6A6 | -0.507332437 | 8.62E-07 |
| NAPSA | 1.781750552 | 9.30E-05 |
| NAPSB | 4.02170796 | 1.66E-05 |
| NR1H2 | -0.213572648 | 0.008463808 |
| LRRC4B | 0.884904329 | 3.12E-06 |
| PDLIM4 | 0.578986062 | 1.61E-06 |
| KIF3A | -0.357845219 | 8.37E-05 |
| MGAT1 | -0.79109762 | 9.58E-09 |
| GFPT2 | 0.188428857 | 0.016843601 |
| TUBG1 | 0.163849863 | 0.00509146 |

| | | |
|---|---|---|
| PSME3 | -1.207601131 | 9.08E-11 |
| RPL27 | 0.23943658 | 0.000192874 |
| PSMC3IP | -0.366927125 | 0.000101857 |
| AOC3 | 2.604934793 | 2.68E-09 |
| ACLY | 0.499894963 | 1.36E-07 |
| AOC2 | 0.918203962 | 8.37E-05 |
| NDUFA2 | 0.298042548 | 0.000289996 |
| ANKHD1 | -0.938737975 | 5.41E-09 |
| DIAPH1 | -0.628783494 | 1.25E-08 |
| NDFIP1 | -0.234906779 | 0.000630617 |
| UBE2D2 | -0.840081425 | 2.76E-08 |
| ACAP3 | 0.49205938 | 1.70E-05 |
| ANO1 | 0.277594945 | 0.007657873 |
| THOC6 | 0.359972234 | 0.000459269 |
| TRAF7 | -0.105811509 | 0.044130935 |
| BARX1 | 0.608238111 | 0.001246694 |
| NINJ1 | -0.506768796 | 0.009610857 |
| NPHP4 | 0.47179203 | 0.000643084 |
| MAP1B | -0.209872453 | 0.001279302 |
| WDR44 | -1.191403234 | 2.22E-07 |
| ZCCHC9 | 0.9865399 | 8.18E-09 |
| TOP2A | -0.985464431 | 9.30E-11 |
| RARA | 0.263497526 | 0.002336927 |
| PEX11B | -0.752675268 | 3.70E-06 |
| FMO5 | 1.257795534 | 0.000225117 |
| PIAS3 | -0.692025685 | 4.43E-07 |
| PRKAB2 | -1.03835164 | 1.50E-08 |
| CLUHP3 | -1.248841942 | 1.92E-05 |
| PDHA1 | 0.19876158 | 0.004589595 |
| MCCC2 | -0.228174961 | 0.000513689 |
| AC005261.1 | 0.946349301 | 6.40E-08 |
| ZSCAN5A | 0.29989259 | 0.009511215 |
| ZNF132 | 1.649525318 | 4.63E-07 |
| CHSY1 | -0.369054086 | 3.55E-06 |
| SNRPA1 | -0.517181482 | 0.000129409 |
| THAP1 | -0.581460423 | 0.000172792 |
| C19orf12 | -0.655588222 | 1.40E-05 |
| LRRC9 | 7.235707434 | 7.63E-06 |
| GCH1 | -1.179325669 | 1.47E-06 |
| LGALS3 | 0.647377736 | 0.000244791 |
| DNAJB1 | 1.931054299 | 1.72E-13 |
| ZSWIM4 | 0.994477779 | 2.54E-06 |
| RFX1 | 0.988458257 | 1.60E-06 |
| ZNF20 | 1.362399078 | 4.23E-09 |
| DCAF15 | -0.31078738 | 0.000410873 |
| CC2D1A | -0.630992678 | 8.03E-06 |
| LRRC41 | -0.490156434 | 1.09E-06 |
| CCT6B | 3.339529717 | 0.000321553 |

| | | |
|---|---|---|
| DHX30 | -0.515036422 | 2.21E-07 |
| RAF1 | -0.566730119 | 2.79E-07 |
| NUP210 | 0.216198096 | 0.032727194 |
| FCRLA | 0.750367016 | 0.000252634 |
| HSD17B7 | 3.269351321 | 1.95E-12 |
| ENOSF1 | -0.264528756 | 0.000613749 |
| SLX1A | -0.350233503 | 0.000132595 |
| ARFIP2 | -0.450499586 | 3.20E-05 |
| TRIM5 | 0.826016426 | 5.59E-07 |
| RRP8 | -0.236856407 | 0.010349671 |
| TIMM10B | -2.459154327 | 1.13E-10 |
| EFR3A | -0.588623677 | 7.95E-07 |
| PTCD3 | -0.741806163 | 1.30E-07 |
| MRPL35 | -0.502511213 | 3.30E-06 |
| ILKAP | -0.222167074 | 0.048485544 |
| PER2 | 1.855086486 | 2.78E-07 |
| SCLY | -0.669855125 | 9.04E-06 |
| RAN | -0.095685097 | 0.013421099 |
| CARD6 | -1.775291441 | 7.19E-07 |
| AC002316.1 | 1.476469881 | 2.52E-10 |
| CLUH | -0.362086774 | 8.42E-06 |
| INPP5K | 0.70154257 | 2.68E-07 |
| MYBBP1A | -0.636113582 | 7.87E-08 |
| RPA1 | -0.320336072 | 1.99E-05 |
| UBE2G1 | -0.806961442 | 2.73E-08 |
| TBC1D14 | -0.897877771 | 5.89E-09 |
| TMEM128 | 0.524864186 | 0.000135708 |
| COQ3 | -0.949580282 | 9.79E-05 |
| PNISR | -0.272795722 | 0.000239414 |
| POPDC3 | -0.357460984 | 0.001020556 |
| SEC61G | 0.335627853 | 0.0056125 |
| LANCL2 | -0.723610647 | 6.37E-06 |
| FIGNL1 | -1.226462465 | 8.25E-10 |
| GRSF1 | -0.437762655 | 4.66E-06 |
| ANKRD17 | -0.653426827 | 2.46E-08 |
| UTP3 | -1.262812063 | 2.57E-09 |
| ITGB4 | 1.345339553 | 1.29E-06 |
| WBP2 | 0.226450632 | 0.001690143 |
| H3-3B | -0.814276485 | 1.28E-10 |
| UNK | -0.219252656 | 0.049291747 |
| TRIM47 | -1.257435854 | 3.51E-06 |
| ZRANB2 | -0.203207819 | 0.00111929 |
| EIF5A | -0.086750274 | 0.015508545 |
| KDM6B | 1.098238034 | 3.03E-09 |
| GPS2 | 0.288286264 | 0.000100093 |
| DLG4 | 1.025105418 | 1.28E-06 |
| RIDA | 0.947794375 | 1.45E-07 |
| VPS13B | -0.74232395 | 3.04E-06 |

| | | |
|---|---|---|
| RGS22 | 8.13532235 | 4.81E-07 |
| REEP2 | 0.449162799 | 0.001128363 |
| PCBD2 | -1.44606478 | 2.90E-05 |
| SDF2 | -0.414532428 | 5.22E-05 |
| ERAL1 | -0.230878176 | 0.002516721 |
| PRMT7 | -0.54205276 | 5.25E-05 |
| NIP7 | -1.121957604 | 7.87E-10 |
| VPS4A | -0.409276377 | 1.53E-05 |
| MTSS2 | -0.601885745 | 1.52E-05 |
| HSPA12B | 2.062832512 | 7.65E-06 |
| ANKEF1 | 0.602285849 | 0.001619809 |
| SNAP25 | 1.052103155 | 0.001149492 |
| PCNA | -0.484706264 | 1.42E-07 |
| NXT1 | -0.489011118 | 0.000232023 |
| POLR3F | -1.099020631 | 1.86E-08 |
| RIN2 | -2.920917307 | 8.90E-10 |
| PTPRA | -0.485788301 | 1.33E-05 |
| DAP3 | 0.202004966 | 0.002549092 |
| KHDC4 | -0.671617036 | 1.74E-07 |
| ATP1A4 | 6.958751275 | 1.86E-05 |
| NES | -0.56813655 | 4.09E-08 |
| BCAN | -0.420232059 | 0.018011601 |
| ARHGEF11 | -1.055032891 | 5.74E-09 |
| RAB25 | 4.791870178 | 1.39E-06 |
| HAPLN2 | 4.591400514 | 0.00044473 |
| DCAF8 | -0.752381881 | 1.75E-08 |
| SYT11 | 0.239764998 | 0.000465519 |
| IGHMBP2 | -1.575577584 | 1.17E-08 |
| DPH2 | -0.645156218 | 1.57E-07 |
| TOE1 | -0.786333037 | 8.23E-08 |
| MUTYH | 1.098994471 | 2.68E-07 |
| RBM38 | -0.61193076 | 3.22E-06 |
| VSTM2L | 7.121845684 | 1.19E-05 |
| OSER1 | 0.892531522 | 1.39E-08 |
| SERINC3 | -0.285724607 | 8.91E-05 |
| PPP1R3D | 0.506672407 | 0.000309202 |
| DMGDH | 8.447647922 | 1.14E-07 |
| AP3B1 | -0.220918366 | 0.004133755 |
| ZBED3 | -0.50643552 | 8.02E-05 |
| ANGPTL3 | 2.965279956 | 0.00018654 |
| CPLANE2 | 1.580269096 | 3.71E-06 |
| DCTN4 | -0.484268628 | 9.82E-07 |
| MTUS2 | 0.784373917 | 0.049235683 |
| ZMYM5 | 1.100725793 | 1.29E-08 |
| USPL1 | -0.3875674 | 8.52E-05 |
| XPO4 | -1.335399383 | 2.18E-10 |
| CDK8 | -0.52474623 | 8.27E-06 |
| WASF3 | -1.180089895 | 3.31E-08 |

| | | |
|---|---|---|
| MYH8 | 2.451071262 | 6.10E-05 |
| MYH10 | -0.105484439 | 0.037247421 |
| SCO1 | -1.219813504 | 5.32E-10 |
| MPRIP | -0.412534297 | 7.41E-07 |
| PIK3C2B | -1.756286261 | 3.08E-07 |
| DSTYK | -0.568602898 | 3.09E-06 |
| SLC41A1 | -0.487012931 | 3.41E-06 |
| DCLK1 | 2.639287452 | 1.60E-08 |
| COG6 | -0.287404692 | 0.009567373 |
| SPART | -1.348794497 | 2.37E-10 |
| RFXAP | -0.603083118 | 0.000723148 |
| TPT1 | 0.622787929 | 5.83E-09 |
| KL | 7.081211297 | 1.58E-05 |
| RFC3 | -1.455873688 | 1.01E-10 |
| STARD13 | -0.492204362 | 5.40E-06 |
| MORC4 | -0.243572249 | 0.049119082 |
| BEX2 | 1.845506088 | 2.22E-05 |
| FAM104A | -0.769777007 | 1.26E-07 |
| SLC39A11 | -0.456223307 | 0.003710821 |
| SRRM1 | -0.903001803 | 1.32E-08 |
| KMT5C | 1.08119043 | 1.98E-05 |
| ZNF414 | -1.27823591 | 4.28E-05 |
| PDE6B | 1.626921369 | 0.018045336 |
| SLF1 | -1.017838824 | 1.37E-08 |
| WDR74 | -0.546396319 | 7.13E-06 |
| RTN3 | -0.099585939 | 0.030691063 |
| MED10 | -0.233806334 | 0.006240009 |
| PDZD2 | -0.259556614 | 0.016348962 |
| MORC2 | -0.343093487 | 0.000207842 |
| LARGE1 | -1.304290164 | 8.86E-05 |
| GSTT2B | 1.288491525 | 0.002939272 |
| SLC2A11 | 0.736661872 | 1.99E-05 |
| C1QTNF6 | 3.398867223 | 2.64E-10 |
| ADCK2 | -0.249968168 | 0.012591304 |
| MKRN1 | 0.121866264 | 0.037116633 |
| AGAP3 | -0.207214893 | 0.00459628 |
| ZNF767P | -0.738159676 | 5.17E-05 |
| BTG1 | -0.657592933 | 1.11E-07 |
| C12orf29 | -0.441850689 | 0.000128138 |
| ATP13A3 | -1.197662455 | 1.13E-10 |
| TMEM254 | 0.667364019 | 2.58E-05 |
| TMTC1 | 0.688123614 | 0.001449716 |
| KRAS | -0.703152965 | 3.84E-08 |
| IPO8 | -0.41167635 | 7.30E-05 |
| LARS1 | -0.2203957 | 0.000498821 |
| IMPA1 | -1.157072387 | 2.58E-09 |
| LRRCC1 | 0.547953167 | 0.002907143 |
| E2F5 | -0.223989858 | 0.009082716 |

| | | |
|---|---|---|
| CCDC59 | -0.581881004 | 4.07E-06 |
| SWAP70 | -0.308288275 | 0.001366845 |
| ARNTL | 1.38549597 | 4.42E-06 |
| SBF2 | -0.28350288 | 0.014411181 |
| MICAL2 | -0.384135228 | 0.028948586 |
| RRAS2 | -1.356559936 | 3.97E-08 |
| ZFC3H1 | 0.150159748 | 0.008506535 |
| TEX15 | -1.480112278 | 1.24E-06 |
| SARAF | 0.460548196 | 7.07E-07 |
| RNF122 | 1.2833399 | 0.000558259 |
| DPF2 | 0.141031831 | 0.02104407 |
| ERG28 | 0.321531127 | 0.012872318 |
| UNC79 | 4.099007836 | 1.22E-05 |
| NUMB | -0.672296631 | 5.25E-07 |
| COX16 | -0.31681722 | 0.004436337 |
| MED6 | -0.939272701 | 4.03E-07 |
| ELP3 | -0.305946239 | 0.046543496 |
| CTIF | -0.50277482 | 0.000393672 |
| MBD2 | -0.636196322 | 9.07E-06 |
| IER3IP1 | -0.754887907 | 1.21E-06 |
| CCNB1 | -1.295157565 | 1.68E-10 |
| IRAK2 | 1.125563864 | 1.64E-05 |
| THUMPD3 | -0.698742905 | 7.06E-08 |
| VHL | -0.448148685 | 6.97E-06 |
| BHLHE40 | 3.3094937 | 2.06E-13 |
| ARL8B | -0.610206394 | 1.06E-07 |
| EDEM1 | -0.156555807 | 0.011887169 |
| MEIS2 | -0.378396051 | 0.000704146 |
| DPH6 | 1.099724635 | 2.14E-06 |
| KATNBL1 | -0.41736666 | 7.86E-05 |
| PRPF38B | -1.730459805 | 2.39E-11 |
| TSPAN2 | -2.779077878 | 3.88E-06 |
| GSTM3 | 0.632203613 | 0.000340735 |
| VAV3 | -2.1673707 | 0.000523917 |
| PSRC1 | -2.162925513 | 6.26E-10 |
| SORT1 | -0.311040333 | 0.000714257 |
| WNT2B | 0.964237394 | 0.000366084 |
| PTGFRN | -0.497117486 | 9.00E-07 |
| NOTCH2 | -0.280795948 | 1.43E-05 |
| TRIM45 | 0.600393354 | 0.004452118 |
| CD101 | 2.591825454 | 2.96E-06 |
| VTCN1 | 7.905136964 | 1.96E-06 |
| NAPG | -1.137540705 | 1.83E-08 |
| SPIRE1 | -1.147468908 | 3.92E-08 |
| PPHLN1 | -0.252525055 | 0.001490689 |
| ARF3 | -0.800125788 | 6.58E-09 |
| SLC38A2 | 0.308959492 | 7.16E-07 |
| KIDINS220 | -0.612767716 | 4.68E-07 |

| | | |
|---|---|---|
| GRHL1 | 1.790038443 | 2.24E-08 |
| ROCK2 | -1.88736729 | 6.29E-12 |
| RSAD2 | 3.502462643 | 2.42E-05 |
| LPIN1 | 1.889078477 | 1.44E-10 |
| CMPK2 | -1.961089113 | 5.55E-06 |
| IL6ST | -0.40786447 | 7.91E-07 |
| FST | -1.206974962 | 1.84E-10 |
| NAV1 | -1.104743413 | 1.11E-08 |
| CDC73 | -1.639969168 | 5.79E-11 |
| TIMM17A | -0.723542574 | 1.93E-08 |
| CRB1 | 5.713915384 | 1.06E-08 |
| RPS15A | 0.13365237 | 0.041874273 |
| NARS1 | -0.270433842 | 2.71E-05 |
| RELCH | -0.807930638 | 1.13E-05 |
| FBH1 | 0.171243667 | 0.01774936 |
| RBM17 | -0.4139157 | 6.86E-05 |
| ANKRD16 | -0.630386336 | 0.007905924 |
| IL15RA | -0.983020221 | 0.005382309 |
| CCNH | -0.631768482 | 1.17E-05 |
| HRH4 | 8.078547322 | 4.37E-07 |
| TMEM241 | -0.418033494 | 0.01824425 |
| KCTD1 | 0.667426798 | 0.000347163 |
| CABLES1 | -0.701639252 | 2.00E-05 |
| SOX5 | 0.598333392 | 2.03E-07 |
| KLRD1 | 8.523804989 | 9.35E-08 |
| SPX | 3.068991416 | 0.022734321 |
| DDB2 | 0.456363795 | 3.43E-05 |
| RTL8C | -0.568134691 | 5.30E-06 |
| RAB33A | 7.469332956 | 4.32E-06 |
| RBMX2 | -0.761215724 | 4.69E-07 |
| STK26 | -0.313779021 | 0.000671797 |
| PUM1 | -0.360390745 | 1.87E-05 |
| PHC2 | -0.451102996 | 4.34E-07 |
| CDCA8 | -1.611709702 | 4.38E-11 |
| GNL2 | -0.549173569 | 6.56E-07 |
| ago-04 | -0.52860423 | 0.000473839 |
| BTF3L4 | -2.055070384 | 3.85E-11 |
| TUT4 | -0.205655258 | 0.003271474 |
| PRPF38A | -0.505299338 | 1.32E-06 |
| RNF138 | -0.672673666 | 6.50E-07 |
| ELP2 | 0.156979761 | 0.015640321 |
| DSC3 | -0.422545443 | 0.012019081 |
| FHOD3 | -0.523037348 | 0.008288527 |
| TPGS2 | 0.12244847 | 0.048458407 |
| DAGLA | -0.750258563 | 0.000116545 |
| SLC43A3 | -0.595489177 | 9.84E-08 |
| TIMM10 | -0.785334958 | 2.95E-06 |
| APLNR | 7.045797261 | 1.45E-05 |

| | | |
|---|---|---|
| FADS2 | 0.846334118 | 1.04E-07 |
| TMEM258 | 0.635135631 | 4.86E-06 |
| TCN1 | 1.501926379 | 0.00054245 |
| TMEM165 | -1.674592542 | 1.35E-10 |
| CLOCK | -1.451493032 | 2.87E-10 |
| GGACT | 1.21199781 | 0.000974649 |
| COL4A2 | 0.346386646 | 3.76E-06 |
| UBAC2 | -0.394959091 | 5.39E-05 |
| ARGLU1 | -0.460450223 | 0.000683676 |
| BIVM | -0.411131554 | 1.42E-05 |
| ERCC5 | -0.325393606 | 0.001163467 |
| TPP2 | -0.409632357 | 2.47E-05 |
| POGLUT2 | -0.485190348 | 2.45E-05 |
| CARS2 | -0.169210441 | 0.019918876 |
| ARHGAP32 | -1.584650076 | 8.60E-09 |
| ETS1 | -1.318003992 | 1.30E-10 |
| TMED7 | -0.691349376 | 4.90E-08 |
| APC | -0.256268027 | 0.000227877 |
| NREP | -0.623702662 | 2.28E-05 |
| WDR36 | -1.358375427 | 2.20E-10 |
| RFK | -1.013950832 | 1.13E-08 |
| UBQLN1 | -0.45487262 | 1.17E-06 |
| NAA35 | -0.434742678 | 0.000236926 |
| C9orf40 | -0.902313367 | 7.00E-08 |
| ANXA1 | 0.302034523 | 4.22E-05 |
| CTSL | 0.736415148 | 4.10E-08 |
| CEMIP2 | -1.53624697 | 1.39E-11 |
| AGTPBP1 | 0.299372834 | 0.022671165 |
| GOLM1 | -0.429043973 | 2.45E-06 |
| FAM189A2 | -1.325163333 | 0.00499813 |
| ISCA1 | -0.400051376 | 5.03E-05 |
| CCNJL | -1.352507881 | 2.62E-09 |
| TAOK3 | -0.225875268 | 0.013232549 |
| HNF1A | -1.423274675 | 0.000661937 |
| FBXO21 | -1.177328668 | 1.25E-08 |
| TBX3 | 0.824524473 | 3.46E-07 |
| HRK | -1.207909013 | 0.022708805 |
| RNFT2 | 0.718897214 | 0.000629208 |
| P2RX4 | 1.616788629 | 0.009345157 |
| BICDL1 | 3.234821674 | 1.16E-07 |
| DMTF1 | -0.589231712 | 3.35E-07 |
| CCDC146 | 2.033225879 | 0.000242805 |
| TMEM60 | 0.610368873 | 0.00037818 |
| HILPDA | -0.562753763 | 0.000742636 |
| RINT1 | -0.312315041 | 0.000510381 |
| SRPK2 | -0.765538784 | 5.04E-08 |
| TES | 0.193847128 | 0.006864636 |
| MDFIC | -1.199443082 | 3.27E-09 |

| | | |
|---|---|---|
| MTO1 | -0.749827237 | 8.23E-06 |
| ANKRD6 | 0.731094575 | 0.003283288 |
| CEP162 | 0.697757076 | 0.002389069 |
| SYNCRIP | -0.761859089 | 2.66E-09 |
| NT5E | 0.324595586 | 0.000117013 |
| EPHA7 | -1.054392778 | 2.47E-05 |
| ORC3 | 0.429592 | 0.002713767 |
| LCA5 | 1.027750052 | 7.82E-06 |
| MAP3K7 | -1.08330664 | 1.00E-08 |
| CGA | 7.998245314 | 6.01E-07 |
| LMO2 | 0.522749461 | 0.010882168 |
| PHF21A | 0.3633853 | 0.001973707 |
| NAT10 | -0.562936155 | 4.15E-07 |
| PRRG4 | -0.611934473 | 0.020110044 |
| CAPRIN1 | -0.452497631 | 1.08E-07 |
| DNAJC14 | -0.495072144 | 1.08E-06 |
| CD63 | 0.330158899 | 0.000186567 |
| PRPH | 3.615515853 | 1.20E-05 |
| AVIL | 2.045077539 | 1.23E-07 |
| GDF11 | -0.555304546 | 2.86E-07 |
| GLS2 | 1.267662058 | 0.039247052 |
| ITGA7 | 0.880598687 | 0.000268534 |
| FAM186B | 0.834143673 | 0.007316118 |
| RDH5 | 1.016349881 | 0.000150411 |
| AGAP2 | 1.220411368 | 0.000137319 |
| BLOC1S1 | 0.43343834 | 0.000608634 |
| CDK4 | 0.140754443 | 0.00989614 |
| TROAP | -0.724286479 | 1.21E-07 |
| TSPAN31 | 0.317653012 | 8.38E-05 |
| B4GALNT1 | 0.127181547 | 0.027857162 |
| TFCP2 | -0.873401702 | 1.46E-07 |
| COQ10A | -0.337235181 | 0.005750853 |
| PAN2 | 0.494258079 | 3.03E-06 |
| ESPL1 | -0.609242183 | 1.85E-07 |
| ZC3H10 | 1.104400281 | 1.36E-08 |
| HNRNPA1 | 0.168435399 | 0.000363683 |
| ACVR1B | -1.067069182 | 6.86E-08 |
| OS9 | 0.319639121 | 3.17E-05 |
| LTV1 | -1.020415879 | 8.38E-09 |
| MAP7 | 0.827840478 | 0.037289531 |
| CD164 | -0.802493608 | 1.98E-09 |
| NHSL1 | -0.255096134 | 0.002205454 |
| AHI1 | 0.627551963 | 0.000103423 |
| HEY2 | 2.704014828 | 3.38E-07 |
| PKIB | 1.805922814 | 0.000662984 |
| SMPD2 | -0.491387798 | 0.012647178 |
| TEC | 1.0243977 | 0.002213106 |
| PRADC1 | 0.252697757 | 0.028621562 |

| | | |
|---|---|---|
| SEMA4F | -0.715078993 | 1.34E-05 |
| CCT7 | 0.243529005 | 7.83E-05 |
| EGR4 | 4.917122012 | 6.88E-07 |
| RAB11FIP5 | -0.527764854 | 1.88E-05 |
| SMYD5 | -0.721996494 | 8.87E-07 |
| CCDC142 | 0.187976857 | 0.008534282 |
| EMX1 | 4.7918362 | 0.000236601 |
| USP15 | 0.318225675 | 6.23E-05 |
| GNS | -0.198063251 | 0.001181332 |
| CPM | 0.365100272 | 0.04139027 |
| MDM2 | 0.741758174 | 3.27E-09 |
| KLHL36 | -0.647950189 | 1.32E-06 |
| BCO1 | 3.930063251 | 7.16E-05 |
| MPHOSPH6 | -1.455811814 | 1.81E-08 |
| KIAA0513 | 1.082770593 | 1.01E-08 |
| DYNC1LI2 | -1.57168507 | 3.24E-11 |
| FBXL8 | 0.778097238 | 0.007205888 |
| FHOD1 | -0.751426339 | 5.94E-08 |
| SLC9A5 | -0.676786803 | 0.000260614 |
| ZNF670-ZNF695 | 0.55647532 | 0.018101547 |
| PCNX2 | 0.409324633 | 0.000102919 |
| KCNK1 | -0.739424449 | 0.001257538 |
| URB2 | -1.05841656 | 9.76E-10 |
| EGLN1 | -1.558614785 | 6.84E-11 |
| COG2 | -0.404407299 | 9.02E-05 |
| ABCB10 | -0.32129609 | 0.000183746 |
| NTPCR | 0.637753982 | 2.57E-06 |
| TAF5L | -0.289361802 | 9.31E-05 |
| STX6 | -0.392899158 | 2.19E-05 |
| RNASEL | 0.231312258 | 0.015596191 |
| KIAA1614 | -0.443844887 | 0.014988238 |
| CEP350 | -0.398699042 | 3.63E-06 |
| NPL | 0.531932028 | 0.001783971 |
| NIBAN1 | -0.446050451 | 5.13E-06 |
| PIGC | -0.948540096 | 4.97E-07 |
| MRPL44 | -0.778134205 | 1.73E-05 |
| PAX3 | -1.577922667 | 2.59E-07 |
| DOCK10 | -1.174060796 | 0.000527917 |
| TTLL4 | -1.391350139 | 5.21E-10 |
| USP37 | -0.622093197 | 1.17E-06 |
| ITM2C | 0.32684498 | 0.000185789 |
| SERPINE2 | 0.508084999 | 7.22E-09 |
| DNAJB2 | 0.495856476 | 6.49E-05 |
| TMBIM1 | 0.475833764 | 5.54E-06 |
| CYP27A1 | 0.475458101 | 0.014675439 |
| EIF4E2 | -1.2834249 | 1.51E-09 |
| CAB39 | -1.200860801 | 5.89E-10 |
| COX5B | 0.445221884 | 0.000354331 |

| | | |
|---|---|---|
| REV1 | -0.925341078 | 2.82E-07 |
| TSGA10 | 2.116236368 | 1.18E-07 |
| MFSD9 | -0.516872224 | 0.004532074 |
| TMEM127 | -1.280947239 | 2.51E-07 |
| TGFBRAP1 | -0.614576081 | 1.42E-06 |
| GCC2 | -0.362711138 | 5.08E-05 |
| C2orf49 | -0.660626959 | 5.89E-06 |
| ANKRD36 | -1.156510686 | 7.56E-06 |
| EPC2 | -1.451224371 | 5.59E-09 |
| ARHGEF4 | 0.293878218 | 0.02813528 |
| ALDH1L2 | -0.194780395 | 0.042658548 |
| SCYL2 | -1.084018944 | 1.51E-09 |
| APPL2 | -0.487099796 | 1.92E-05 |
| PWP1 | -0.261392912 | 0.001078045 |
| DRAM1 | -1.16537047 | 2.01E-07 |
| WASHC4 | -1.067714633 | 1.41E-09 |
| SLC41A2 | -1.168445624 | 1.25E-06 |
| NEK3 | 2.759573467 | 2.86E-06 |
| VPS36 | -1.603930374 | 3.03E-10 |
| RNASEH2B | -0.77817522 | 0.000186992 |
| CKAP2 | -1.299845441 | 5.58E-11 |
| TBC1D4 | -2.06283125 | 9.10E-12 |
| BORA | -0.933487156 | 2.64E-06 |
| LRCH1 | -1.188882319 | 4.63E-08 |
| SUCLA2 | -0.753101006 | 5.59E-06 |
| MED4 | -0.572566207 | 2.37E-05 |
| PHF11 | -0.815703016 | 0.00122839 |
| RPL13AP25 | 0.617429286 | 2.29E-07 |
| COG3 | 0.521671211 | 3.39E-06 |
| LMO7 | -0.920224732 | 1.61E-06 |
| SPRY2 | -0.710418721 | 8.02E-09 |
| NUDT15 | -0.729691454 | 4.44E-08 |
| RCBTB2 | -0.83942489 | 0.000196284 |
| LCP1 | 4.850469892 | 2.04E-07 |
| SCRN1 | -0.133112408 | 0.00334745 |
| C7orf25 | -0.774511205 | 1.71E-05 |
| TNS3 | -0.829115897 | 1.60E-07 |
| IGF2BP3 | -1.375268495 | 1.03E-10 |
| GPNMB | 0.294169338 | 0.000207668 |
| RAPGEF5 | -0.985869135 | 0.000132932 |
| RAC1 | -0.441464348 | 2.69E-07 |
| KDELR2 | -0.271615562 | 2.55E-05 |
| NUP42 | -0.962650835 | 7.80E-08 |
| ZDHHC4 | 0.467501072 | 0.000253977 |
| BZW2 | 0.182683619 | 0.009104026 |
| TBRG4 | -0.621178942 | 5.99E-07 |
| DDX56 | -0.369358874 | 3.68E-05 |
| HUS1 | -1.095610597 | 2.59E-07 |

| | | |
|---|---|---|
| CCM2 | 0.302225913 | 0.040123281 |
| CIDEB | -0.70513346 | 0.000131793 |
| TTC5 | -0.79167609 | 1.32E-05 |
| ZFHX2 | 1.555098789 | 9.61E-06 |
| ADAMTS7 | 0.924060183 | 0.009964618 |
| ABHD17C | -0.84326692 | 4.16E-08 |
| IREB2 | -0.579929193 | 1.91E-07 |
| ALPK3 | 3.487377198 | 3.96E-09 |
| CIB2 | 1.076463991 | 0.003302958 |
| CALCOCO2 | 0.274012821 | 0.000960178 |
| RSAD1 | -0.467674337 | 2.37E-06 |
| NMT1 | -0.564866586 | 1.70E-06 |
| MYCBPAP | 3.746292353 | 3.38E-06 |
| SRSF1 | -1.406826615 | 1.51E-12 |
| VEZF1 | -1.062105017 | 6.85E-10 |
| CHAD | 2.574850832 | 0.003325098 |
| TACO1 | -0.984821857 | 1.17E-06 |
| DCAF7 | -0.688148677 | 1.32E-08 |
| LIMD2 | 0.525814858 | 0.000171432 |
| BRIP1 | -0.905371997 | 9.21E-08 |
| KAT7 | -1.547180789 | 4.99E-10 |
| ACTL6A | -0.211469464 | 0.01716744 |
| NDUFB5 | 0.526739265 | 4.68E-06 |
| MRPL47 | -0.329798789 | 0.00351758 |
| TRA2B | -1.426890214 | 5.79E-11 |
| SCN2A | 1.767879546 | 4.23E-05 |
| TBR1 | 3.895871218 | 2.97E-05 |
| MARCHF7 | -0.143273672 | 0.014877801 |
| GALNT5 | 0.823259882 | 0.034781507 |
| TANK | 0.437524067 | 0.001695534 |
| GATA4 | 0.318583088 | 0.00975988 |
| EPRS1 | 0.267472128 | 5.93E-05 |
| KCTD3 | -1.119992927 | 5.89E-10 |
| RPS6KC1 | -0.952092599 | 8.79E-09 |
| CBWD2 | -0.242348262 | 0.018273116 |
| SMPD4 | -0.399521377 | 9.08E-05 |
| WDR33 | -0.847770105 | 1.40E-08 |
| IMP4 | -0.420752808 | 6.35E-06 |
| HS6ST1 | 0.180464513 | 0.032489395 |
| UGGT1 | -0.233854772 | 0.000705388 |
| GYPC | 0.446075441 | 0.00198695 |
| STAM | -0.580779678 | 1.07E-05 |
| ABI1 | -0.802660966 | 4.15E-07 |
| YME1L1 | -0.246367595 | 0.000256201 |
| DNAJC1 | -0.277720523 | 0.048225847 |
| NIPSNAP3A | 0.307781127 | 0.025803468 |
| LRRC8A | -0.706270199 | 2.93E-08 |
| CDK9 | 0.481129483 | 6.55E-06 |

| | | |
|---|---|---|
| TXN | 0.271561262 | 0.000634738 |
| ODF2 | 0.206396977 | 0.002990402 |
| ECPAS | -0.262421225 | 0.000504997 |
| TOR1B | 0.153804594 | 0.012536601 |
| SMC2 | -0.813597208 | 1.05E-08 |
| KLF4 | 1.04803281 | 3.29E-08 |
| TOR1A | -0.279363856 | 0.000132266 |
| DAB2IP | -0.863117861 | 1.26E-07 |
| STXBP1 | -0.48879536 | 8.99E-06 |
| CDK5RAP2 | -0.328693129 | 0.000277202 |
| ZFP37 | -0.760853948 | 4.71E-05 |
| SLC31A2 | -0.481818696 | 0.004361166 |
| TLR4 | -1.749862733 | 2.39E-11 |
| ZNF189 | -0.540458816 | 4.68E-06 |
| ALDOB | 5.963217892 | 2.56E-09 |
| STX17 | -0.898471049 | 1.32E-06 |
| PRPF4 | -0.60977982 | 7.28E-08 |
| USP20 | 0.486366263 | 6.26E-05 |
| BAAT | 4.034505763 | 1.11E-05 |
| ATP6V1G1 | -0.210608958 | 0.004546781 |
| TEX10 | -0.692686495 | 1.24E-07 |
| GARNL3 | 2.105544952 | 0.000714597 |
| MRPL50 | -1.355989714 | 1.71E-09 |
| DPM2 | -0.302575485 | 0.000438216 |
| WDR38 | 2.522512579 | 0.004049144 |
| HEMGN | 7.340494991 | 5.69E-06 |
| PSMB7 | 0.281636116 | 0.000134102 |
| NCBP1 | -0.599768767 | 1.98E-07 |
| ANP32B | -0.163187364 | 0.004291394 |
| RPL35 | 0.341244379 | 2.30E-05 |
| LMX1B | -1.520354747 | 0.000558098 |
| ARPC5L | -0.402065374 | 1.16E-05 |
| ENPP2 | 1.098697629 | 0.002369537 |
| DSCC1 | -1.088914585 | 1.43E-08 |
| DERL1 | -0.893507765 | 5.07E-09 |
| MYC | 1.691156722 | 1.06E-10 |
| RANBP6 | -0.690598995 | 5.11E-08 |
| POLR1E | -0.85064761 | 2.05E-07 |
| PLAA | -0.219165609 | 0.001316194 |
| UBAP2 | -1.062504014 | 6.63E-09 |
| APTX | -0.756732123 | 3.85E-07 |
| RNF38 | -0.791572081 | 6.84E-08 |
| TLN1 | 0.090349636 | 0.02112145 |
| DNAJB5 | 2.303986452 | 3.85E-08 |
| SPAG8 | 1.393092596 | 0.001063966 |
| DCTN3 | 0.246565 | 0.012510961 |
| TMEM8B | 0.796660556 | 0.000551114 |
| ALDH1B1 | -0.643627857 | 3.53E-07 |

| | | |
|---|---|---|
| HINT2 | 0.954882214 | 1.74E-07 |
| ARHGEF39 | -1.238789669 | 8.18E-08 |
| DENND4C | -0.18545617 | 0.006074563 |
| RPS6 | 0.416109729 | 4.44E-08 |
| CNPY3 | 0.236585288 | 0.005994733 |
| FOXP4 | -0.206266955 | 0.012696215 |
| PPIL1 | -0.823386192 | 7.00E-08 |
| KLC4 | 0.405517636 | 0.017485327 |
| KIF13A | -0.797711379 | 1.07E-05 |
| PIM1 | 1.911988216 | 8.44E-10 |
| CMTR1 | -0.198161954 | 0.033770328 |
| TFAP2A | -1.035474131 | 5.40E-06 |
| YIPF3 | 0.346021648 | 0.000148016 |
| FRS3 | 1.595332738 | 9.74E-06 |
| TJAP1 | -0.5584284 | 7.51E-06 |
| IRF4 | 2.332863652 | 0.001530465 |
| SLC22A23 | -0.690322255 | 0.004214349 |
| GCM1 | 8.662928676 | 5.18E-08 |
| FOXF2 | -2.209417065 | 4.90E-08 |
| BPHL | 1.868701147 | 6.61E-08 |
| HMGA1 | 0.519884847 | 8.57E-09 |
| TCF19 | -0.997358442 | 5.55E-05 |
| RANP1 | -0.244734859 | 0.048800037 |
| IER3 | 0.265688777 | 0.026437801 |
| MDC1 | -0.699384919 | 0.000141256 |
| PGBD1 | -0.42295869 | 0.005541298 |
| RNF144B | 1.282667221 | 0.000574606 |
| TAF8 | -0.257142091 | 0.012775384 |
| FAM8A1 | -0.186010796 | 0.011554083 |
| C6orf52 | 2.826560482 | 5.67E-05 |
| CPEB2 | -0.493643669 | 0.006074987 |
| FHDC1 | -1.734217339 | 4.94E-06 |
| FCHSD2 | -1.153895755 | 2.23E-07 |
| THAP12 | -0.290920667 | 5.90E-05 |
| ANKRD42 | -0.89135882 | 8.23E-08 |
| IL18BP | -0.971958904 | 6.32E-08 |
| NUMA1 | -0.689048362 | 3.07E-08 |
| CCDC90B | 0.19541997 | 0.036227323 |
| SYTL2 | 0.250651282 | 0.024191408 |
| RAB30 | 0.60696344 | 7.60E-06 |
| CREBZF | -0.789294405 | 8.96E-09 |
| PRCP | 0.23699787 | 0.009968431 |
| NARS2 | -0.259741294 | 0.015071705 |
| RNF121 | -1.027455539 | 2.66E-07 |
| MRPL15 | -0.664042179 | 6.99E-07 |
| PI15 | -0.790132358 | 8.53E-07 |
| TTPA | -1.674568437 | 0.000849837 |
| SLCO5A1 | -1.147128701 | 8.17E-06 |

| | | |
|---|---|---|
| SULF1 | 1.508210436 | 1.78E-05 |
| TGS1 | -0.40608118 | 1.35E-05 |
| SDCBP | 0.382027761 | 4.81E-06 |
| NEK1 | -0.571768429 | 7.22E-05 |
| DDX60 | -0.737245936 | 0.009248295 |
| BUD13 | -1.04856349 | 2.11E-07 |
| DCUN1D5 | -0.342250471 | 8.81E-05 |
| YAP1 | -0.681477032 | 1.24E-08 |
| SLC37A4 | 0.221869828 | 0.022655538 |
| POU2F3 | 1.372573614 | 0.036549261 |
| RDX | -0.904243097 | 2.34E-09 |
| PPP2R1B | -2.000088282 | 4.26E-11 |
| C11orf1 | 1.636234175 | 1.78E-07 |
| ALKBH8 | -0.517968868 | 6.12E-06 |
| SQOR | 2.007800504 | 0.000178106 |
| CTDSPL2 | -1.674682299 | 3.81E-11 |
| THBS1 | -1.175682592 | 7.61E-12 |
| NUSAP1 | -1.00445947 | 1.47E-09 |
| NDUFAF1 | 0.280520761 | 0.003504043 |
| KIF23 | -2.08022405 | 6.29E-12 |
| AC027088.1 | 1.180081673 | 0.003102512 |
| KNL1 | -0.897796493 | 2.46E-09 |
| HAUS2 | -0.710831087 | 2.83E-07 |
| RTF1 | -0.757407631 | 2.47E-08 |
| PARP6 | 0.660496033 | 2.66E-05 |
| RPLP1 | 0.321476047 | 5.55E-06 |
| TUBGCP4 | -0.782174955 | 1.90E-08 |
| RMDN3 | -0.399078323 | 5.71E-05 |
| UACA | 0.305987429 | 0.000163636 |
| SMAD6 | -1.449880183 | 6.00E-09 |
| ADAM10 | -0.747136717 | 1.47E-08 |
| DUOX1 | 1.663574499 | 0.00016081 |
| STRA6 | 0.361701121 | 0.023182276 |
| RSL24D1 | -0.577476801 | 1.44E-07 |
| SPTBN5 | 1.745377654 | 2.28E-09 |
| GCHFR | 0.557427774 | 0.01450294 |
| BCAR3 | -1.010579448 | 2.94E-08 |
| TTLL7 | -0.817045946 | 0.000591829 |
| FNBP1L | -0.678676792 | 3.10E-06 |
| GTF2B | 0.776750835 | 1.98E-07 |
| RABGGTB | -1.048525055 | 5.78E-10 |
| ARHGAP29 | -0.226511159 | 0.003438563 |
| IFI44 | 0.644618557 | 8.45E-05 |
| RPL7P9 | 0.290640608 | 8.11E-06 |
| CLCA2 | 1.995658156 | 0.010365687 |
| DBT | -1.055152004 | 1.56E-08 |
| RTCA | -0.338867058 | 0.000500357 |
| IFT172 | 0.244075583 | 0.007483595 |

| | | |
|---|---|---|
| SELENOI | -0.569877412 | 3.47E-08 |
| HADHB | 0.266974177 | 0.000426278 |
| ADCY3 | -0.669844638 | 1.33E-07 |
| PPM1B | -0.502312002 | 8.10E-06 |
| PNPT1 | -0.72982366 | 7.41E-07 |
| DYNC2LI1 | 1.20236069 | 6.70E-06 |
| THUMPD2 | -1.234616491 | 9.86E-07 |
| CYP1B1 | -3.063176063 | 1.89E-09 |
| ACTR2 | -0.271016555 | 1.32E-05 |
| SLC5A6 | -1.135121649 | 1.69E-09 |
| ABCG5 | 1.766015715 | 0.016159066 |
| PREPL | -0.113658349 | 0.036767185 |
| EMILIN1 | 0.558261772 | 1.95E-06 |
| FBXO11 | -0.597857551 | 2.76E-07 |
| ATRAID | 0.466522847 | 2.91E-06 |
| CENPO | -1.267238069 | 2.04E-09 |
| TRIM54 | 4.336441053 | 5.02E-06 |
| ACTR1A | -0.567862841 | 3.17E-07 |
| MFSD13A | 0.478387289 | 0.010053564 |
| MYOF | -0.66992778 | 1.34E-08 |
| LOXL4 | 1.493003334 | 2.99E-07 |
| STAMBPL1 | -0.942542229 | 2.54E-06 |
| ATAD1 | -0.804650952 | 1.69E-08 |
| KIF11 | -1.501502741 | 3.19E-11 |
| TACC2 | -1.046650569 | 4.10E-08 |
| DUSP5 | 1.810586446 | 3.06E-11 |
| CALHM2 | -0.841053396 | 5.19E-06 |
| CEP55 | -0.654621151 | 5.07E-08 |
| KIF20B | -1.372569534 | 6.21E-10 |
| ENTPD1 | -1.005461866 | 1.73E-09 |
| EXOC6 | -1.515707218 | 5.90E-07 |
| PLCE1 | -0.736073275 | 1.11E-05 |
| DBR1 | -0.695710675 | 1.01E-06 |
| DNAJC13 | -0.150931717 | 0.020059469 |
| ANXA7 | -0.19223304 | 0.010232125 |
| FAM149B1 | -0.994002539 | 6.81E-09 |
| ASCC1 | -0.577172185 | 0.00023829 |
| PLA2G12B | 7.636006656 | 2.81E-06 |
| RPS24 | 0.274189523 | 3.26E-06 |
| TET1 | -1.248174696 | 1.71E-05 |
| DNA2 | -0.917615478 | 3.01E-07 |
| MYPN | 0.810047577 | 0.003990511 |
| SMARCAL1 | 0.398595021 | 0.006375942 |
| BARD1 | -0.44856741 | 0.000194633 |
| STAT4 | 4.65030755 | 6.47E-07 |
| CARF | 1.732696263 | 9.04E-06 |
| ASNSD1 | -0.948249798 | 6.98E-09 |
| NAB1 | -0.370367535 | 6.67E-05 |

| | | |
|---|---|---|
| PPIG | -1.412798787 | 2.46E-09 |
| FASTKD1 | -0.79713723 | 0.000372716 |
| IDH1 | 0.82099872 | 1.53E-08 |
| OLA1 | 0.174983355 | 0.013228335 |
| ITPRID2 | -0.778873729 | 4.58E-09 |
| FAM117B | -1.951097617 | 3.91E-09 |
| WDR12 | -0.610599739 | 7.90E-06 |
| ABI2 | -0.577192618 | 2.38E-07 |
| ITGAV | -0.195768677 | 0.003048582 |
| SLC35A5 | -0.459857664 | 3.24E-05 |
| SLC49A4 | -0.274335295 | 0.049038784 |
| PARP9 | -1.097014829 | 1.06E-05 |
| USP8 | -0.306370055 | 0.000373477 |
| SECISBP2L | 0.238857355 | 0.000491538 |
| TMOD3 | -1.254878467 | 6.46E-10 |
| SPPL2A | -0.579052536 | 4.06E-06 |
| GLCE | -0.825957014 | 1.05E-08 |
| APH1B | -0.85836443 | 0.006496232 |
| INTS14 | -0.510909534 | 8.55E-06 |
| CILP | 2.574850832 | 0.003325098 |
| PARP16 | -1.132230904 | 1.80E-06 |
| PPCDC | -0.853145331 | 0.000113894 |
| FAM13A | -0.786510312 | 5.95E-06 |
| HERC3 | -0.354486006 | 2.88E-05 |
| HERC6 | -0.451627932 | 0.032388738 |
| PCDH10 | -0.689221583 | 1.63E-06 |
| ZGRF1 | -0.397079576 | 0.00091075 |
| AP1AR | -0.201109707 | 0.004729039 |
| COPS4 | -0.179625809 | 0.046157868 |
| HNRNPD | -0.240774057 | 0.000308997 |
| SEC31A | -0.147151202 | 0.019024988 |
| FGF5 | -1.662085506 | 0.019452738 |
| GPAT3 | -0.272922387 | 0.000322992 |
| FGF2 | -0.699043486 | 1.22E-06 |
| BMPR1B | -1.593191194 | 0.003319716 |
| RAP1GDS1 | -0.244727764 | 0.005652908 |
| LARP1B | -0.801421879 | 4.84E-06 |
| MMRN1 | 6.774837438 | 3.09E-05 |
| PDE5A | -0.477537121 | 0.021723802 |
| TRPC3 | -1.635962127 | 0.014015826 |
| NAAA | -0.452964981 | 0.013230593 |
| BMP2K | -0.235190656 | 0.006331744 |
| G3BP2 | -1.406193077 | 4.65E-11 |
| SEPTIN11 | -0.415947563 | 6.41E-07 |
| CCNG2 | 1.89655482 | 1.71E-10 |
| CNOT6L | -0.905436997 | 4.39E-07 |
| SHROOM3 | -1.482664982 | 1.00E-08 |
| CENPE | -0.837604924 | 3.26E-08 |

| | | |
|---|---|---|
| GSTCD | -0.191096413 | 0.030732157 |
| INTS12 | -0.478899634 | 0.00017045 |
| LEF1 | -1.071687922 | 5.98E-07 |
| EGF | -1.09811676 | 0.026328039 |
| PAPSS1 | 0.173243966 | 0.013157351 |
| PPP3CA | -0.578119692 | 0.001261506 |
| SLC39A8 | -0.923853006 | 9.80E-08 |
| MTTP | 6.436789506 | 3.30E-06 |
| MAPK8IP3 | -0.538035622 | 3.66E-06 |
| RGS3 | 0.403766931 | 0.003284511 |
| RNF185 | -1.37924858 | 7.97E-08 |
| ETV6 | -0.738902076 | 6.16E-06 |
| GABARAPL1 | -0.788695142 | 3.41E-06 |
| KIF21A | -0.514811707 | 4.44E-06 |
| CPNE8 | -1.283482942 | 2.40E-08 |
| YARS2 | -1.760652801 | 7.98E-10 |
| SINHCAF | -0.546992237 | 4.22E-07 |
| AEBP2 | -1.31934644 | 2.76E-09 |
| ETFBKMT | 2.029674193 | 3.60E-08 |
| ETNK1 | -1.848245451 | 9.08E-11 |
| ZCRB1 | 0.271006626 | 0.004512965 |
| TMEM117 | -1.760830379 | 9.41E-07 |
| PRICKLE1 | 0.661856711 | 0.000446435 |
| C1RL | 0.766819385 | 0.00055225 |
| NDUFA9 | -0.315892576 | 0.00143886 |
| KLRG1 | 1.738893277 | 0.000100322 |
| VAMP1 | 0.715691261 | 7.21E-06 |
| TAPBPL | 0.962098535 | 4.53E-05 |
| CD27 | 1.299636161 | 0.008651853 |
| PEX5 | -0.294424774 | 0.001825876 |
| PIANP | 2.190424677 | 2.06E-06 |
| AMIGO2 | -1.659955276 | 8.17E-11 |
| SCAF11 | -1.423830837 | 4.53E-11 |
| COL2A1 | 2.069615403 | 0.000418304 |
| LLPH | -0.268071457 | 0.001317781 |
| RPL14P1 | 0.263665377 | 2.81E-05 |
| MARCHF9 | 0.541868748 | 0.000608937 |
| INHBE | 5.069474308 | 3.13E-07 |
| GLIPR1 | -1.147805179 | 1.70E-09 |
| PHLDA1 | -1.000504647 | 2.77E-10 |
| TMEM19 | -0.921773748 | 8.70E-08 |
| LGR5 | 5.668499924 | 2.08E-09 |
| DUSP6 | -0.568281789 | 3.38E-08 |
| TMTC3 | -0.811719273 | 2.56E-08 |
| SNRPF | 0.325453195 | 0.003208635 |
| AMDHD1 | 5.483996448 | 3.02E-05 |
| NEDD1 | -1.054717396 | 8.64E-09 |
| GAS2L3 | -0.574122746 | 1.89E-06 |

| | | |
|---|---|---|
| TDG | -0.233219044 | 0.001591322 |
| RITA1 | -0.538217674 | 3.72E-05 |
| SDSL | 1.371384798 | 0.000178089 |
| MMAB | 0.43430257 | 0.003125868 |
| GLTP | -0.396454907 | 0.000290957 |
| GIT2 | -0.485729532 | 1.73E-05 |
| TCHP | -0.948106837 | 1.53E-08 |
| FAM222A | -0.82882797 | 7.83E-07 |
| NUP58 | -0.623409345 | 2.92E-08 |
| MTMR6 | -0.32832052 | 8.71E-05 |
| SLC46A3 | -0.744889407 | 0.007963303 |
| SLC7A1 | -1.264259405 | 5.42E-11 |
| PDX1 | -1.448359155 | 0.000859119 |
| SUOX | 0.822525018 | 1.25E-07 |
| CCDC65 | 0.922798229 | 0.000278084 |
| SLC39A5 | 4.766095408 | 3.68E-07 |
| TARBP2 | -0.53627782 | 7.61E-05 |
| RDH16 | 6.338135613 | 6.31E-06 |
| DHH | 3.286458023 | 0.017342392 |
| NABP2 | 0.15628042 | 0.031114143 |
| N4BP2L1 | 2.365955277 | 7.41E-06 |
| BRCA2 | 0.311051598 | 0.00015352 |
| KANSL2 | 0.239282059 | 0.00602782 |
| MAP3K12 | 0.222577026 | 0.004198384 |
| ITGB7 | 2.432076593 | 1.37E-07 |
| CSAD | 0.374427337 | 0.007699522 |
| C12orf10 | 0.239474547 | 0.00195394 |
| ANKRD52 | -1.748744018 | 1.47E-11 |
| ZNF740 | -0.597634623 | 3.52E-07 |
| WDFY2 | -1.582734576 | 1.26E-09 |
| HNRNPA1L2 | 0.518938795 | 2.22E-07 |
| LPAR6 | 2.41362138 | 1.15E-09 |
| SBNO1 | -0.112534913 | 0.031831984 |
| MORN3 | 2.596628433 | 0.000955619 |
| SETD1B | -0.48430915 | 3.60E-06 |
| VPS33A | -0.395752821 | 0.00032452 |
| VPS37B | -0.944834339 | 3.05E-08 |
| RHOF | 0.307995663 | 0.000609427 |
| DENR | -1.059077183 | 2.09E-09 |
| DIAPH3 | -0.980633855 | 1.62E-06 |
| RBM26 | -2.429696441 | 1.34E-08 |
| ZIC5 | -1.73988553 | 0.00176839 |
| ABHD13 | -0.357728694 | 4.05E-05 |
| RAB20 | -0.732498337 | 9.87E-07 |
| CUL4A | -0.375118964 | 4.51E-06 |
| CDH24 | -0.330883032 | 0.036570066 |
| REM2 | 2.473228428 | 0.000498766 |
| CBLN3 | 1.199906713 | 0.009208293 |

| | | |
|---|---|---|
| TSSK4 | 0.947620527 | 0.001114259 |
| NOVA1 | -2.612130257 | 8.09E-08 |
| FITM1 | 3.231928855 | 0.016458778 |
| TMX1 | -0.723460102 | 3.50E-07 |
| FRMD6 | -0.458198973 | 3.21E-06 |
| PELI2 | 0.436288123 | 0.004036392 |
| NAA30 | -0.706787624 | 2.38E-07 |
| ADAM21 | 3.265951641 | 0.000176906 |
| RDH12 | 6.618143764 | 4.85E-05 |
| DCAF5 | -0.388258405 | 8.13E-05 |
| RAB15 | -1.081232423 | 7.10E-07 |
| ESR2 | 1.171543495 | 0.003726002 |
| KCNH5 | 1.124844518 | 0.001415733 |
| PTGR2 | 0.478056223 | 0.004516784 |
| JDP2 | 1.45350907 | 9.39E-06 |
| AK7 | 1.536660218 | 0.013107895 |
| CLBA1 | 0.486297458 | 0.000677618 |
| WARS1 | 0.426495392 | 1.44E-06 |
| NIPA2 | -1.092966774 | 5.13E-10 |
| DUOXA1 | 7.695159179 | 2.18E-06 |
| TCF12 | -0.659315901 | 4.70E-07 |
| SERF2 | 0.500341037 | 8.95E-07 |
| ZSCAN29 | 0.185248544 | 0.00716384 |
| DUOX2 | 5.853012876 | 3.11E-05 |
| LYSMD2 | -1.411344295 | 2.51E-06 |
| FGF7 | 1.575939017 | 0.001375405 |
| BNIP2 | -1.608291334 | 7.43E-11 |
| SRP14 | 0.273280251 | 0.000469465 |
| CDAN1 | -0.885031704 | 3.37E-07 |
| TLE3 | -1.487412646 | 6.57E-09 |
| ANP32A | -0.369428496 | 9.51E-06 |
| COMMD4 | -0.142656786 | 0.034500332 |
| UBE2Q2 | -0.626284114 | 1.30E-06 |
| ETFA | 0.197036366 | 0.013701738 |
| BCL2A1 | 1.409804863 | 0.000222206 |
| SCAPER | 0.690754434 | 0.007836643 |
| WDR61 | -0.483410826 | 0.000104587 |
| NCOA2 | -0.433746936 | 0.000147193 |
| NEIL1 | -0.284521981 | 0.01384308 |
| TLNRD1 | -0.496990321 | 0.001024323 |
| TPM1 | -0.800099957 | 8.75E-09 |
| IGF1R | -0.278869066 | 4.07E-05 |
| ARRDC4 | 2.76311371 | 2.90E-12 |
| PIF1 | -2.130609155 | 1.34E-09 |
| USP3 | -0.988978272 | 5.58E-07 |
| CYP11A1 | 1.806537758 | 0.007106706 |
| BBS4 | 0.750804306 | 5.06E-05 |
| CYP1A1 | -1.226260356 | 4.81E-06 |

| | | |
|---|---|---|
| ADAMTS17 | -1.551903422 | 8.25E-06 |
| LINS1 | -0.312626979 | 0.001713365 |
| AC091230.1 | -0.185522696 | 0.030659992 |
| GOLGA6D | 7.238108345 | 8.91E-06 |
| PCSK6 | 3.680264474 | 8.00E-05 |
| CELF6 | 2.704686224 | 2.59E-05 |
| SCAMP2 | -0.31201621 | 0.000363226 |
| LMAN1L | 4.324080477 | 3.45E-05 |
| HAPLN3 | 0.953733 | 3.89E-06 |
| POLG | -1.034413613 | 2.96E-09 |
| RLBP1 | 1.452324383 | 0.000138894 |
| FANCI | -0.282616902 | 0.000110499 |
| ABHD2 | -0.274794341 | 0.000127853 |
| WDR93 | 2.121399473 | 0.00191615 |
| TICRR | -0.866908283 | 1.85E-08 |
| DET1 | 0.325372936 | 0.019685696 |
| ZNF710 | -1.680822004 | 1.94E-08 |
| UNC45A | 0.168482033 | 0.014091521 |
| FURIN | -0.149765515 | 0.013888072 |
| CRTC3 | -0.739715627 | 9.57E-07 |
| EFL1 | -0.596569241 | 1.04E-05 |
| SEC11A | 0.306162386 | 9.57E-05 |
| GLYR1 | -0.460049219 | 1.91E-06 |
| PMM2 | -1.029568823 | 3.29E-08 |
| SLC5A2 | 0.495554403 | 4.57E-05 |
| ITGAX | 2.211042238 | 1.19E-05 |
| TGFB1I1 | 0.564586986 | 4.96E-05 |
| C16orf58 | 0.343551898 | 0.00034528 |
| ARMC5 | -1.272656851 | 1.23E-08 |
| PARN | -0.278934128 | 0.000784758 |
| FTO | -0.87172825 | 3.53E-08 |
| CDR2 | 0.238984995 | 0.001146042 |
| ARHGAP17 | -0.429726009 | 2.25E-05 |
| MYLK3 | 3.062497985 | 7.69E-05 |
| NKD1 | -1.225801368 | 0.000842079 |
| DHX38 | -0.333567858 | 2.79E-05 |
| TXNL4B | -0.739427617 | 3.43E-07 |
| ZFHX3 | -0.526374504 | 3.18E-06 |
| CLEC18B | 2.446012711 | 2.02E-06 |
| CPNE2 | 0.479947673 | 0.021245822 |
| NLRC5 | -0.898410045 | 0.000152123 |
| KATNB1 | 0.413147773 | 5.69E-05 |
| KIFC3 | -0.353440876 | 4.88E-05 |
| NUDT7 | 2.075080196 | 0.000184998 |
| CMTM3 | -0.254860889 | 0.001456244 |
| CMTM2 | 7.717822129 | 1.55E-06 |
| NOL3 | 0.922241065 | 1.77E-06 |
| MAP1LC3B | 0.400584909 | 2.42E-06 |

| | | |
|---|---|---|
| CDH13 | 0.329784542 | 4.70E-05 |
| ZCCHC14 | 0.640317005 | 6.00E-07 |
| MEAK7 | -0.746053667 | 7.42E-06 |
| OSGIN1 | 0.537004672 | 0.002180126 |
| RHOT2 | -0.173047546 | 0.006066385 |
| RPL3L | 2.091625906 | 0.000428027 |
| ZSCAN32 | 0.554635362 | 3.78E-06 |
| RPS2 | 0.416644934 | 1.05E-05 |
| NDUFB10 | 0.201488599 | 0.007645985 |
| TIGD7 | 0.379789189 | 0.005374656 |
| DEF8 | 0.247357894 | 0.001625962 |
| GALNS | 0.379581743 | 0.004306426 |
| NCOR1 | -0.595167288 | 2.06E-07 |
| CDRT15P1 | -1.604341262 | 0.002093626 |
| COPS3 | -0.240899019 | 0.001455851 |
| GID4 | -0.367804252 | 0.000269091 |
| KSR1 | -0.879280304 | 7.81E-05 |
| UTP4 | 0.153558733 | 0.037521875 |
| PRPSAP2 | 0.198556003 | 0.016228689 |
| PCTP | 0.262147938 | 0.008893969 |
| C17orf80 | -0.370215305 | 2.66E-05 |
| VPS53 | -0.77472305 | 1.68E-07 |
| NPEPPS | -0.456616191 | 2.53E-06 |
| LRRC46 | 0.835841687 | 0.001076514 |
| SCRN2 | 0.561012336 | 0.000498881 |
| SSH2 | -0.947417442 | 7.90E-09 |
| RHBDL3 | -1.173204925 | 0.01495921 |
| ARSG | 1.025437238 | 0.000247783 |
| G6PC3 | 0.174295652 | 0.010466853 |
| CLTC | -0.346984209 | 2.89E-07 |
| C17orf64 | 7.766563296 | 1.51E-06 |
| PTRH2 | -0.749429054 | 6.11E-08 |
| SS18 | -1.658554535 | 5.10E-11 |
| TAF4B | -1.443112055 | 8.36E-09 |
| AFG3L2 | -0.410006677 | 9.53E-06 |
| GNAL | 1.971069081 | 7.72E-10 |
| SLC39A6 | 0.166783075 | 0.002466939 |
| RPRD1A | 0.336986132 | 6.59E-05 |
| GALNT1 | -0.405333376 | 3.38E-05 |
| ESCO1 | -0.835386336 | 1.79E-07 |
| OSBPL1A | -0.282141041 | 0.020332079 |
| GATA6 | 0.592022309 | 0.004425188 |
| GREB1L | -1.372883332 | 1.31E-06 |
| RMC1 | -0.260674227 | 0.010713536 |
| PELP1 | -0.379830754 | 6.57E-06 |
| SLC14A1 | 0.426497311 | 0.019255799 |
| WRAP53 | 0.162612711 | 0.02650567 |
| MINK1 | -1.058889212 | 4.21E-09 |

| | | |
|---|---|---|
| SAT2 | 0.704121362 | 7.20E-06 |
| ASGR1 | 1.985694855 | 0.000186511 |
| TP53 | 0.237589137 | 0.005094279 |
| CCDC40 | 0.737620224 | 0.015132282 |
| ARHGDIA | -0.681727893 | 1.05E-08 |
| TMC6 | 0.857482624 | 5.88E-06 |
| CARD14 | 2.078085401 | 2.34E-05 |
| EIF4A3 | -0.33371417 | 0.000124819 |
| CSNK1D | -0.865316717 | 5.30E-09 |
| TBCD | -0.462416909 | 0.000630146 |
| FN3KRP | -0.74567141 | 5.57E-07 |
| RPTOR | -0.9636428 | 2.93E-08 |
| FOXK2 | -0.889165729 | 6.93E-09 |
| SECTM1 | -0.38334593 | 0.030438849 |
| CEP131 | 0.527731535 | 0.00071571 |
| WDR45B | -0.934538049 | 1.32E-08 |
| CBX4 | 0.530904171 | 1.25E-05 |
| RNF165 | 1.204539021 | 0.002336463 |
| ELAC1 | 1.355875701 | 2.28E-07 |
| MBD1 | -0.737819109 | 5.56E-08 |
| SMAD4 | -1.103185415 | 1.99E-09 |
| ZCCHC2 | -1.06006898 | 1.34E-07 |
| FBXO15 | 4.491837521 | 6.28E-08 |
| PMAIP1 | 0.51365957 | 6.55E-06 |
| ERBB2 | 0.635467552 | 7.01E-07 |
| GRB7 | 3.422496922 | 2.10E-05 |
| STAC2 | 1.552421468 | 0.005304711 |
| IGFBP4 | 0.146731017 | 0.007327065 |
| FKBP10 | 0.276034656 | 3.25E-05 |
| TXNL4A | -0.221525193 | 0.003314453 |
| CACNA1A | 1.723660609 | 0.000180048 |
| BRD4 | -0.605442983 | 1.44E-06 |
| SLC39A3 | -0.327152228 | 0.008081174 |
| NFIC | -0.541366418 | 2.11E-05 |
| PLPP2 | 0.431426563 | 7.08E-05 |
| PRDM15 | -2.003127166 | 6.93E-09 |
| PFKL | 0.232443394 | 0.019430326 |
| FEM1A | -0.217492727 | 0.001842207 |
| MVB12A | 0.4498452 | 5.75E-05 |
| AC008764.1 | 0.355074721 | 0.006714174 |
| SH3GL1 | -0.44086926 | 2.55E-06 |
| DUS3L | -0.595279815 | 4.18E-06 |
| CCDC97 | -1.020858551 | 3.54E-08 |
| IFITM3 | 0.21917845 | 0.000928237 |
| PGGHG | 0.68236518 | 0.001563391 |
| COL6A1 | 0.424954545 | 1.32E-05 |
| IFNAR1 | -0.231478429 | 0.001400612 |
| SOD1 | 0.308222224 | 0.000261581 |

| | | |
|---|---|---|
| COL6A2 | 0.599254851 | 4.47E-07 |
| SIK1 | 2.233922424 | 2.58E-07 |
| TRPM2 | -0.826292991 | 0.00031446 |
| TMEM50B | -0.971713901 | 0.000196359 |
| APP | 0.243684574 | 0.000112152 |
| URB1 | -0.508225831 | 1.93E-06 |
| EMP3 | 0.803039976 | 1.37E-05 |
| SAE1 | -0.342261773 | 1.00E-05 |
| GEMIN7 | -1.540643947 | 6.45E-08 |
| WTIP | -0.658160073 | 0.004582747 |
| CAPN10 | -0.615860953 | 0.000202855 |
| MYO1F | 3.61502319 | 2.02E-06 |
| ERVK3-1 | -0.439492126 | 2.48E-06 |
| NLRP12 | 3.180573538 | 0.000671654 |
| CACNG8 | -0.329601618 | 0.007400009 |
| ZNF787 | -1.278902384 | 1.66E-06 |
| TIMM29 | -0.570993492 | 1.76E-05 |
| EVI5L | 1.688045192 | 2.82E-08 |
| SLC47A1 | -0.841273186 | 7.13E-05 |
| PSMB6 | 0.2630067 | 0.000430848 |
| SIGLEC10 | 3.925104781 | 0.02025808 |
| ZNF473 | 0.77711022 | 1.25E-07 |
| RPS11 | 0.458241626 | 3.51E-05 |
| AC020909.1 | 0.739174926 | 0.004242075 |
| RPL13A | 0.459161181 | 4.74E-08 |
| CTU1 | -0.65054844 | 0.001462393 |
| IGLON5 | 4.827862188 | 0.00031645 |
| RERE | -0.42479017 | 5.14E-06 |
| FHAD1 | 2.392576853 | 0.000627956 |
| EPHA2 | -0.685708752 | 9.78E-09 |
| EFHD2 | -0.642636577 | 1.43E-07 |
| PGD | 0.647448243 | 9.39E-08 |
| SH3BGRL3 | 0.356859083 | 1.46E-06 |
| CNKSR1 | 2.710306833 | 2.71E-05 |
| RPL11 | 0.417147948 | 3.53E-07 |
| IL22RA1 | -0.644796912 | 0.028203838 |
| C1orf216 | -0.550688301 | 5.01E-06 |
| KIAA0319L | 0.925229587 | 3.05E-08 |
| PLK4 | -1.215810025 | 9.27E-09 |
| MAP3K6 | -0.709644387 | 3.53E-05 |
| SYTL1 | 1.031822134 | 0.017002248 |
| WDTC1 | 0.173003224 | 0.033731619 |
| SERBP1 | -0.685216699 | 6.75E-09 |
| BCL10 | 1.161481937 | 7.65E-09 |
| CCN1 | 0.91144881 | 3.68E-10 |
| PRKACB | -1.78923719 | 1.04E-09 |
| PIGK | -0.308524759 | 0.000404833 |
| TINAGL1 | 0.282292873 | 4.65E-05 |

| | | |
|---|---|---|
| AZIN2 | 2.585136468 | 4.69E-09 |
| RPS8 | 0.446049434 | 1.14E-06 |
| KIF2C | -1.207690333 | 5.33E-10 |
| PTPRF | -0.202847964 | 0.001219321 |
| MOB3C | 0.566222719 | 2.56E-05 |
| SYPL2 | 1.670753497 | 1.59E-06 |
| MTF2 | -1.085880198 | 1.99E-08 |
| IGSF3 | -1.173608737 | 5.52E-10 |
| ZNF697 | -0.518793275 | 2.07E-06 |
| CTTNBP2NL | 0.550924686 | 2.98E-06 |
| STRIP1 | -0.856646167 | 1.88E-06 |
| C1orf162 | 4.395514344 | 4.30E-08 |
| PROK1 | 7.843595027 | 1.28E-06 |
| CELSR2 | 0.169446102 | 0.021925223 |
| ITGA10 | 1.789650198 | 8.48E-08 |
| GPR161 | -0.212805672 | 0.009991212 |
| ALDH9A1 | -0.530908162 | 4.14E-06 |
| ATP1B1 | -0.957534216 | 4.67E-09 |
| TIPRL | -0.236154282 | 0.000362774 |
| POGK | -1.001565337 | 4.73E-10 |
| CREG1 | -0.280397105 | 0.000133401 |
| RXRG | 1.302815088 | 0.027223415 |
| TBX19 | 0.855247161 | 0.000934903 |
| UCK2 | -1.552630761 | 8.54E-11 |
| TMCO1 | -0.296790506 | 8.45E-05 |
| POU2F1 | -2.092619884 | 7.34E-12 |
| ADCY10 | 4.19695533 | 1.18E-08 |
| COP1 | -0.818670709 | 9.08E-06 |
| NECTIN4 | 4.623235728 | 3.85E-08 |
| PPOX | 0.465425947 | 0.00094575 |
| FCGR2A | 0.513037614 | 0.008752031 |
| NUF2 | -0.899529334 | 3.29E-08 |
| RGS5 | 4.531107488 | 3.14E-12 |
| SDHC | 0.635219646 | 1.61E-08 |
| PFDN2 | 0.316810585 | 0.000300213 |
| NR1I3 | -0.841610114 | 0.001472321 |
| USP21 | -0.814362986 | 2.09E-06 |
| F13B | 8.365405821 | 2.09E-07 |
| PRCC | -0.686643193 | 3.24E-07 |
| RRNAD1 | 1.626295459 | 1.81E-09 |
| MRPL24 | 0.356632235 | 2.40E-05 |
| PIGM | -0.346532346 | 0.000282178 |
| CASQ1 | 3.164556311 | 0.000347068 |
| ISG20L2 | -1.337032923 | 1.03E-10 |
| CRABP2 | 2.256544655 | 2.71E-06 |
| ABL2 | 0.738552319 | 2.15E-08 |
| XPR1 | -0.714462395 | 6.32E-08 |
| RGS16 | 1.661591007 | 3.66E-09 |

| | | |
|---|---|---|
| TOR1AIP1 | -0.389681727 | 4.87E-06 |
| HMCN1 | 0.589791063 | 9.28E-07 |
| RGL1 | 0.299192816 | 0.006027834 |
| PRUNE1 | -0.435779267 | 0.000111577 |
| TUFT1 | 0.666039797 | 2.22E-07 |
| SF3B4 | -1.06335295 | 2.75E-10 |
| ECM1 | 0.69530601 | 6.20E-06 |
| ZNF687 | 0.783893122 | 4.07E-08 |
| SNX27 | -0.983566055 | 3.06E-09 |
| SETDB1 | -0.500179134 | 2.04E-05 |
| ADAMTSL4 | 4.232758039 | 1.62E-13 |
| MCL1 | 0.39582458 | 1.84E-08 |
| PI4KB | -0.778408387 | 1.59E-08 |
| PIP5K1A | -0.766104178 | 3.56E-09 |
| ANP32E | -0.461955355 | 1.02E-05 |
| SELENBP1 | 4.589481122 | 0.000472928 |
| CERS2 | -0.255834438 | 0.000233033 |
| LSP1P4 | -0.695934257 | 7.82E-05 |
| MRPL9 | -0.156369013 | 0.019974898 |
| ARNT | -1.08821194 | 2.07E-08 |
| POGZ | -0.528826472 | 3.93E-07 |
| C1orf56 | 0.194177593 | 0.046036498 |
| OAZ3 | 1.111017305 | 4.50E-05 |
| GOLPH3L | 0.511850397 | 5.52E-06 |
| GABPB2 | -0.506469666 | 0.000816092 |
| SYT14 | -0.975315206 | 8.20E-06 |
| KCNH1 | -2.129444746 | 8.34E-07 |
| DTL | -2.018961712 | 8.96E-12 |
| DYRK3 | 0.523855334 | 1.38E-06 |
| EIF2D | -0.253647947 | 0.00199931 |
| INTS7 | -1.193048325 | 1.88E-09 |
| VASH2 | -0.498285168 | 0.004461701 |
| SMYD2 | -0.791053331 | 1.64E-07 |
| DUSP10 | 2.220122611 | 4.00E-11 |
| HHIPL2 | 2.382935231 | 0.000392047 |
| TP53BP2 | -0.147899412 | 0.01557718 |
| ATP8B2 | 0.231525525 | 0.000695577 |
| FLG2 | 1.808924582 | 1.44E-06 |
| RAB13 | 0.53588129 | 1.10E-06 |
| TPM3 | -0.071133171 | 0.032260907 |
| NUP210L | 6.376304702 | 1.06E-08 |
| SNAPIN | -0.242214928 | 0.003537014 |
| S100A7 | 9.489785549 | 3.82E-09 |
| UBAP2L | -0.854714156 | 5.58E-09 |
| SLC39A1 | 0.432985112 | 8.06E-07 |
| HAX1 | 0.228468184 | 0.001068882 |
| CREB3L4 | 1.32386601 | 5.00E-07 |
| EFNA3 | 0.700402461 | 0.008568681 |

| | | |
|---|---|---|
| KCNN3 | 3.036275712 | 1.40E-08 |
| GATAD2B | 0.429302356 | 4.22E-06 |
| RIT1 | -0.309447075 | 0.005517105 |
| INTS3 | -0.653937049 | 3.20E-07 |
| FLG | -0.559553616 | 0.00018995 |
| C1orf131 | -0.291981968 | 0.005620642 |
| SCCPDH | 0.270350219 | 0.003942219 |
| LYST | -0.293340412 | 0.004532994 |
| MAP3K21 | -1.820987491 | 6.28E-09 |
| CEP170 | -1.050225338 | 8.71E-09 |
| ACP1 | -0.159605348 | 0.009373787 |
| SNAP47 | 0.164327802 | 0.007946701 |
| NVL | -0.383128283 | 0.000718208 |
| SDE2 | 0.631747199 | 5.53E-08 |
| DEGS1 | -0.811367459 | 2.57E-08 |
| FBXO28 | -0.533727401 | 2.14E-07 |
| GUK1 | 0.153100563 | 0.017957417 |
| CDC42BPA | -0.902337305 | 1.18E-08 |
| CNIH3 | 0.990225476 | 0.005084672 |
| C1orf35 | -0.304389288 | 0.003004763 |
| MBOAT2 | -0.528463922 | 9.35E-06 |
| PARP1 | -0.115644222 | 0.014728493 |
| PSEN2 | -0.511230172 | 0.001219829 |
| LBR | -0.845271711 | 6.58E-09 |
| WNT9A | 0.704969231 | 2.90E-05 |
| SOX13 | -0.224886871 | 0.021038071 |
| PLEKHA6 | 6.371885329 | 7.09E-05 |
| ARL8A | -0.979030074 | 1.35E-06 |
| PDIA6 | 0.172212069 | 0.00828297 |
| RHOB | 0.665555495 | 8.92E-08 |
| ATP6V1C2 | 0.12318221 | 0.038546513 |
| HNRNPLL | 0.247482164 | 0.00874334 |
| EML4 | -1.068927786 | 1.58E-09 |
| CALM2 | -0.628369006 | 2.04E-09 |
| CHAC2 | -0.772150137 | 1.98E-06 |
| RPS27A | 0.323246165 | 0.001368943 |
| VPS54 | -1.164834265 | 8.01E-08 |
| ASXL2 | -0.29168638 | 3.59E-05 |
| ETAA1 | -0.413187172 | 0.000108248 |
| MEIS1 | 0.854299831 | 0.000143274 |
| CIAO1 | -0.210236004 | 0.000781382 |
| EXOC6B | -0.906633023 | 6.45E-07 |
| SFXN5 | -1.593653243 | 3.78E-08 |
| TEX261 | -0.15802994 | 0.005109403 |
| DQX1 | 4.920319443 | 5.86E-08 |
| THNSL2 | 1.264581145 | 4.11E-08 |
| RALB | 0.203735985 | 0.003867799 |
| NT5DC4 | 1.524538724 | 0.002759651 |

| | | |
|---|---|---|
| RABL2A | 0.661795106 | 0.000175737 |
| SLC20A1 | -0.132205585 | 0.005611892 |
| LYG1 | 2.792493293 | 0.002540173 |
| AFF3 | -0.809313721 | 0.004502373 |
| UBXN4 | -0.602665676 | 2.49E-07 |
| NXPH2 | -4.879068517 | 0.000314061 |
| SPOPL | -1.634260339 | 6.78E-10 |
| POLR2D | -0.259259955 | 0.000517441 |
| AMMECR1L | -0.873971199 | 2.60E-08 |
| GALNT13 | -0.539002852 | 0.002629042 |
| PKP4 | -1.572226574 | 1.17E-09 |
| LNPK | -0.930456227 | 5.34E-08 |
| CDCA7 | -0.972063929 | 1.32E-08 |
| DLX1 | -2.835418881 | 5.58E-11 |
| UBR3 | -0.663773939 | 1.76E-05 |
| GULP1 | -0.364808545 | 0.004273615 |
| FAM171B | -0.648692034 | 1.02E-05 |
| HSPD1 | -0.149752407 | 0.000723361 |
| METTL21A | -0.503893668 | 0.002346357 |
| UNC80 | 0.972319664 | 0.026838108 |
| NBEAL1 | 0.565607162 | 9.78E-08 |
| KANSL1L | 0.237390105 | 0.017915642 |
| RHBDD1 | 0.453543421 | 0.000238698 |
| HES6 | 1.04873251 | 1.75E-05 |
| ESPNL | 0.856877907 | 0.024512254 |
| ANKMY1 | 0.296985112 | 0.030300196 |
| COPS7B | -0.546626926 | 3.45E-06 |
| DIS3L2 | -0.411904213 | 0.005507341 |
| FANCD2 | -0.437864789 | 8.02E-05 |
| TAMM41 | -1.106713855 | 0.000145304 |
| VGLL4 | -1.630788248 | 1.10E-10 |
| RAB5A | -0.594356799 | 2.55E-06 |
| RETREG2 | 0.193177906 | 0.007777273 |
| CTDSP1 | -0.842249866 | 2.41E-07 |
| CNOT9 | -1.476767925 | 4.08E-10 |
| MARCHF4 | -1.569961392 | 4.31E-07 |
| STK11IP | -0.315319272 | 0.008152583 |
| GMPPA | 0.283518526 | 0.010328866 |
| CNTN4 | 1.008104898 | 0.011862103 |
| DYNC1LI1 | -0.161358493 | 0.011028371 |
| RBMS3 | -1.481209861 | 1.42E-07 |
| OSBPL10 | -0.444573009 | 1.04E-05 |
| POMGNT2 | -0.322308344 | 0.000317863 |
| ACKR2 | 4.936778689 | 0.000172484 |
| CSRNP1 | 1.957021985 | 2.08E-11 |
| SLC25A38 | -0.24454217 | 0.001548691 |
| GOLGA4 | -0.733934168 | 3.14E-08 |
| CTDSPL | -1.746806555 | 4.93E-09 |

| | | |
|---|---|---|
| IQSEC1 | -0.348935544 | 0.000300647 |
| CAND2 | 0.582510533 | 0.028666106 |
| RPL32 | 0.309325063 | 0.000210396 |
| PTPRG | -0.223673761 | 0.001414675 |
| IL17RD | -0.439293977 | 0.024831962 |
| SHQ1 | -1.156798786 | 1.04E-08 |
| SLC25A26 | -0.368975295 | 0.016383209 |
| LRIG1 | -0.446071942 | 0.005735501 |
| AC073896.1 | 0.22613453 | 0.002697624 |
| LIMD1 | -1.924240189 | 2.60E-10 |
| NFKBIZ | 0.419640675 | 0.00015854 |
| COL8A1 | 1.554175412 | 1.13E-05 |
| NXPE3 | -0.185476755 | 0.00616179 |
| MYH15 | 1.091060568 | 0.000130086 |
| PHLDB2 | -1.227813991 | 3.48E-10 |
| RABL3 | -1.078598764 | 2.78E-06 |
| ADPRH | -1.748150388 | 1.50E-06 |
| ATG3 | -0.439241002 | 0.000192592 |
| BOC | 1.26046388 | 3.24E-05 |
| SRPRB | -0.69913115 | 7.02E-08 |
| TMEM108 | -0.935342205 | 0.002885341 |
| AGTR1 | 2.048055504 | 9.56E-07 |
| MED12L | -0.567422314 | 0.015705624 |
| EIF2A | -0.204288839 | 0.001659605 |
| ALDH1L1 | 3.741270239 | 0.023717209 |
| OSBPL11 | -1.309065624 | 1.72E-09 |
| NCEH1 | -0.231475374 | 0.004547482 |
| FAM86B2 | 2.211295499 | 0.005448034 |
| LPP | -0.545363878 | 5.69E-07 |
| RUBCN | -0.287663156 | 0.004435872 |
| AMT | 2.462838004 | 3.38E-07 |
| NICN1 | 0.350534033 | 0.028385302 |
| UCN2 | 1.988856908 | 5.55E-09 |
| DCAF1 | -1.432752496 | 3.38E-10 |
| MANF | 0.579940748 | 1.50E-05 |
| STXBP5L | 0.752752986 | 0.044880421 |
| EAF2 | 0.794030039 | 0.002153609 |
| MUC4 | 7.270706977 | 6.81E-06 |
| SLIT2 | -1.211723294 | 1.62E-07 |
| EIF2B5 | -0.376413809 | 5.39E-05 |
| VWA5B2 | 1.797368261 | 0.000822739 |
| DGKQ | 0.455061768 | 0.000494325 |
| FIP1L1 | -0.988224024 | 5.75E-08 |
| SLC26A1 | 0.908906886 | 0.000324143 |
| LYAR | -0.471563823 | 2.23E-05 |
| CORIN | 2.889764204 | 0.00012256 |
| ATP10D | -0.223382511 | 0.007456709 |
| SLC10A4 | -1.711780136 | 0.002993618 |

| | | |
|---|---|---|
| PLAC8 | 0.997774169 | 0.0139368 |
| TRMT10A | 0.978054574 | 2.98E-05 |
| KLHL8 | -1.582506787 | 5.23E-10 |
| PYURF | -0.357718115 | 3.39E-05 |
| TBCK | 0.502985716 | 0.002548249 |
| CAMK2D | -0.412616013 | 0.002139791 |
| CISD2 | -0.790926914 | 3.51E-07 |
| TIFA | -0.599341388 | 7.17E-05 |
| SPATA5 | -1.010238053 | 1.18E-06 |
| CCNA2 | -2.142925171 | 1.01E-11 |
| METTL14 | -1.348313125 | 1.37E-08 |
| USP53 | 0.23078938 | 0.004611954 |
| SETD7 | -0.821149389 | 7.11E-09 |
| NAF1 | -1.417088978 | 5.47E-08 |
| MARCHF1 | -1.524270574 | 3.84E-07 |
| RPS3A | 0.162718955 | 0.015605973 |
| PDGFC | -1.069382219 | 0.000344666 |
| CBR4 | 0.529840429 | 5.02E-05 |
| CYP4V2 | 1.020811997 | 9.74E-05 |
| MARCHF6 | -1.066443367 | 9.00E-10 |
| NKD2 | 1.178169162 | 0.028191536 |
| SRD5A1 | -0.767718684 | 1.17E-06 |
| MYO10 | -0.709270386 | 2.19E-06 |
| RPL37 | 0.279557492 | 3.35E-06 |
| SKP2 | -1.699003556 | 3.52E-11 |
| OSMR | 0.188773206 | 0.013359876 |
| PLK2 | 0.43257938 | 8.52E-06 |
| HAPLN1 | 0.915818258 | 3.62E-09 |
| LHFPL2 | -0.566902263 | 8.39E-07 |
| SSBP2 | -1.248882679 | 6.02E-05 |
| BHMT | 6.671503917 | 4.29E-05 |
| ANKRD31 | 3.566151276 | 1.66E-08 |
| IQGAP2 | -0.705022836 | 0.006878494 |
| RASA1 | -1.495328186 | 6.47E-10 |
| LIX1 | 5.721592044 | 5.33E-05 |
| PPIP5K2 | -0.324604446 | 0.000281357 |
| BDP1 | -0.433405417 | 0.000221366 |
| GTF2H2 | -0.815989253 | 0.000637711 |
| SLC30A5 | -0.228334781 | 0.000610156 |
| BTF3 | 0.279280744 | 9.20E-06 |
| FBXL17 | -0.589737238 | 6.48E-05 |
| TSLP | 2.457573773 | 0.003451162 |
| TNFAIP8 | 0.431328591 | 0.001926212 |
| FEM1C | 0.435950075 | 5.79E-06 |
| COMMD10 | -0.784288191 | 1.47E-05 |
| ATG12 | -1.209114421 | 4.63E-10 |
| YIPF5 | 0.258026283 | 0.000570537 |
| ARHGAP26 | -0.867739666 | 8.80E-06 |

| | | |
|---|---|---|
| RNF145 | -1.508219893 | 4.69E-11 |
| PCYOX1L | 0.39434989 | 0.003563296 |
| GLRA1 | 3.373058872 | 3.79E-05 |
| G3BP1 | -1.890061695 | 2.21E-12 |
| ZNF300 | -0.526387532 | 3.12E-06 |
| NHP2 | 0.162452055 | 0.010518498 |
| BOD1 | 0.263852988 | 0.001168729 |
| TENM2 | 0.693257061 | 0.000112392 |
| FAM50B | -0.931557695 | 0.002148538 |
| CDKAL1 | -1.33517019 | 1.19E-05 |
| PCDHB18P | -0.995340849 | 0.003326007 |
| PSD2 | 2.623168769 | 0.002849185 |
| LRRTM2 | 2.694558318 | 0.005955243 |
| ZMAT2 | -0.351870604 | 7.53E-05 |
| KLHL3 | 1.678322399 | 7.21E-05 |
| TRIM7 | 1.087181153 | 1.34E-05 |
| TRIM41 | -0.375569035 | 5.55E-05 |
| HIGD2A | 0.165097345 | 0.023140551 |
| PLA2G7 | 1.069415344 | 0.02099236 |
| TNFRSF21 | -0.573598091 | 6.96E-07 |
| DOK3 | -1.138336205 | 1.64E-05 |
| ABT1 | -1.001873094 | 3.23E-08 |
| PPP1R18 | -0.762858069 | 0.000185736 |
| PRIM2 | -0.692723208 | 0.000124042 |
| LGSN | 5.292839448 | 0.000200227 |
| FGD2 | 7.555076951 | 2.70E-06 |
| ANO7 | 1.703701272 | 1.77E-05 |
| TCTE1 | 4.790165376 | 0.000244691 |
| RPL7L1 | -0.519318571 | 6.81E-08 |
| NFKBIE | 1.840093502 | 2.84E-09 |
| IRAK1BP1 | -0.689385965 | 0.002061055 |
| PHIP | -0.666937638 | 4.80E-08 |
| MMS22L | -1.756038402 | 1.53E-10 |
| FAXC | -1.470964574 | 2.85E-06 |
| PNRC1 | 1.709931207 | 7.98E-10 |
| PM20D2 | -0.914757411 | 3.89E-08 |
| RARS2 | -0.296035023 | 0.046036534 |
| RNF217 | -2.157129743 | 1.70E-08 |
| ABRACL | -0.28498923 | 0.007980143 |
| SHPRH | -0.822765252 | 1.77E-07 |
| TIAM2 | -0.508383034 | 6.02E-05 |
| TMEM181 | -0.645086718 | 6.19E-08 |
| PNLDC1 | 3.954140962 | 0.004088582 |
| WTAP | -0.253576833 | 0.000566975 |
| ZMYM4 | -1.263056719 | 2.54E-09 |
| ARMT1 | -0.189929001 | 0.019646594 |
| C7orf50 | -0.210802219 | 0.029805905 |
| SDK1 | -0.781136523 | 0.020205928 |

| | | |
|---|---|---|
| EGFR | -0.762547433 | 1.97E-07 |
| CDCA5 | -0.886117927 | 4.93E-09 |
| IGFBP3 | 1.071952694 | 2.54E-09 |
| PURB | -1.080770166 | 7.98E-10 |
| AC004453.1 | 0.299137649 | 4.81E-05 |
| SSC4D | 0.739910644 | 0.01471606 |
| MDH2 | 0.133401399 | 0.016524458 |
| CCT6A | -0.252046117 | 2.20E-05 |
| PSPH | 0.285589216 | 0.010556977 |
| ATXN7L1 | -0.408560415 | 0.034852227 |
| TMEM168 | -1.403492364 | 2.42E-09 |
| MAP11 | -0.468493256 | 0.000126106 |
| SLC12A9 | -0.458213256 | 0.000214577 |
| GIGYF1 | -1.100986919 | 2.53E-08 |
| TRIM4 | 0.753337061 | 5.25E-08 |
| MEPCE | -0.400598151 | 8.76E-06 |
| TMEM209 | -0.374645571 | 6.83E-05 |
| AGBL3 | 2.121122636 | 3.66E-07 |
| ZC3HAV1L | -1.443276539 | 0.000105377 |
| TMEM140 | -0.910852374 | 1.20E-07 |
| TLK2 | -1.208039355 | 4.26E-08 |
| NOM1 | -1.487416406 | 4.57E-10 |
| NCAPG2 | -0.258652183 | 0.008715467 |
| NLGN4X | -0.716960597 | 8.16E-08 |
| SHROOM2 | -2.138460048 | 6.83E-12 |
| LUC7L2 | -1.192546607 | 4.62E-10 |
| DENND2A | 0.769721689 | 0.007933649 |
| CLTRN | 2.571503824 | 2.91E-07 |
| TMEM47 | 0.424253949 | 0.000992963 |
| LANCL3 | -1.867689924 | 0.000444907 |
| SYTL5 | 0.246008368 | 0.000308744 |
| CASK | -0.558571157 | 0.001916738 |
| KDM6A | -0.37027004 | 0.000993802 |
| SPIN2A | -1.39641724 | 0.000204167 |
| CCNB3 | 2.43497711 | 9.19E-06 |
| HDAC8 | -0.283075607 | 0.014452268 |
| ZNF182 | 0.481333855 | 0.000121591 |
| NDUFB11 | 0.293590367 | 0.000136648 |
| ZNF41 | 0.457735749 | 1.83E-05 |
| RAB41 | 1.377008313 | 0.023438921 |
| ZMYM3 | 0.320292752 | 0.000199213 |
| TAF1 | -0.698198754 | 1.18E-07 |
| NONO | -0.177416808 | 0.000288089 |
| CCDC120 | -1.190702337 | 0.000258503 |
| EBP | 0.505747754 | 4.64E-06 |
| OGT | -1.681414531 | 4.48E-12 |
| SNX12 | -0.716324196 | 1.69E-07 |
| ITGB1BP2 | 1.397137857 | 4.38E-05 |

| | | |
|---|---|---|
| IL2RG | 1.66838833 | 0.014734748 |
| GCNA | 3.045318516 | 2.71E-06 |
| DIAPH2 | -0.456744057 | 0.002134747 |
| PRPS1 | -0.687889819 | 4.54E-07 |
| RBMX | -0.13586749 | 0.005343978 |
| MCPH1 | -1.254336103 | 2.15E-08 |
| MFHAS1 | -1.35624735 | 1.35E-09 |
| FBXO25 | 0.498298067 | 0.001701426 |
| MAGEA4 | -0.106668459 | 0.001797754 |
| NSDHL | 0.91993168 | 1.51E-06 |
| CETN2 | 0.377160653 | 0.000155177 |
| RPL10 | 0.396142323 | 1.83E-05 |
| CSGALNACT1 | 0.316423085 | 0.048149418 |
| ATP6V1B2 | -0.66100112 | 2.61E-07 |
| HMBOX1 | -0.865885781 | 2.82E-05 |
| GNRH1 | -0.888004284 | 0.002947963 |
| BIN3 | -0.263993305 | 0.001631845 |
| DOK2 | 5.639294008 | 1.58E-05 |
| SLC25A37 | -1.310138837 | 6.47E-10 |
| CHMP7 | -1.5586119 | 5.54E-10 |
| DOCK5 | -1.432311021 | 2.79E-10 |
| STAR | 4.275967875 | 7.45E-07 |
| PLPBP | 0.718136105 | 1.83E-07 |
| ERLIN2 | -0.206485031 | 0.008570637 |
| TACC1 | -1.40318596 | 8.46E-11 |
| PLPP5 | 0.754069836 | 1.22E-06 |
| GINS4 | -0.516092686 | 1.41E-05 |
| NSD3 | -0.964393409 | 3.08E-09 |
| LACTB2 | 0.489931427 | 0.001985164 |
| TERF1 | -1.029584822 | 6.86E-08 |
| RPL7 | 0.361080346 | 1.18E-05 |
| SLC26A7 | 5.639757129 | 1.62E-05 |
| SYBU | -0.798769862 | 0.004687067 |
| LRP12 | -0.496826943 | 3.78E-06 |
| POLR2K | -0.258949268 | 0.000588397 |
| EIF3H | 0.616499043 | 8.85E-08 |
| UTP23 | -0.245615863 | 0.00095493 |
| NDUFB9 | 0.331681321 | 6.60E-05 |
| TATDN1 | 0.349355714 | 0.000913829 |
| ZNF7 | 0.405755588 | 0.000147682 |
| SLC39A4 | 0.580594897 | 0.016719821 |
| VLDLR | -0.843717246 | 0.000582903 |
| AK3 | -0.420805961 | 7.64E-05 |
| UHRF2 | -1.051125479 | 2.01E-09 |
| NFIB | -1.735043423 | 4.71E-09 |
| PLIN2 | 0.279009849 | 7.54E-05 |
| HAUS6 | -0.429565074 | 8.59E-07 |
| C9orf72 | 0.741912861 | 2.16E-07 |

| | | |
|---|---|---|
| ZCCHC7 | -0.260528432 | 0.003164224 |
| FBXO10 | 1.397547862 | 3.02E-09 |
| CEP78 | -1.148913985 | 6.58E-09 |
| IDNK | 1.637007244 | 0.000108204 |
| AUH | -0.764471358 | 0.00151283 |
| MFSD14B | -0.515429935 | 8.64E-07 |
| ZNF462 | -0.377746753 | 0.000110958 |
| INIP | -0.41763585 | 8.66E-06 |
| UGCG | -1.0589623 | 2.54E-09 |
| SNX30 | -0.918259799 | 2.61E-08 |
| STOM | -0.339023698 | 6.92E-06 |
| GSN | 0.702410939 | 1.70E-07 |
| MRRF | 0.702914032 | 4.02E-07 |
| WDR31 | 3.25292283 | 4.73E-08 |
| POLE3 | -0.790880742 | 1.03E-08 |
| GBGT1 | 1.351499357 | 1.02E-08 |
| SURF1 | 0.940975513 | 7.68E-07 |
| SURF2 | 0.2335223 | 0.009353563 |
| SURF6 | -0.315523388 | 0.000404746 |
| MED22 | -0.635464181 | 2.57E-06 |
| REXO4 | -0.599531593 | 1.34E-06 |
| RPL7A | 0.305030347 | 3.63E-06 |
| GTF3C5 | -0.272081529 | 0.001643211 |
| ASB6 | -0.270520252 | 0.000416096 |
| NTMT1 | 0.31463553 | 0.000528615 |
| SLC25A25 | 0.973965305 | 4.21E-09 |
| MIGA2 | -0.441120425 | 0.000414582 |
| PTGES | 0.277908918 | 0.021003283 |
| GPR107 | -0.47123217 | 1.06E-06 |
| IDI2 | -0.926068964 | 0.001338837 |
| INPP5E | -0.364603705 | 0.006444192 |
| SEC16A | -1.297511564 | 9.34E-11 |
| DPH7 | -0.870878947 | 2.84E-06 |
| NOTCH1 | 0.117533756 | 0.040283748 |
| NACC2 | -2.048822736 | 3.19E-11 |
| PROSER2 | -0.389197386 | 0.024859974 |
| COMMD3 | 0.264147589 | 0.01117253 |
| MSRB2 | 3.237936227 | 1.47E-08 |
| FAM171A1 | -0.7691137 | 1.68E-05 |
| RSU1 | -0.175310373 | 0.020502047 |
| ZEB1 | -0.249461152 | 0.000971479 |
| NRBF2 | 0.582177624 | 1.94E-05 |
| A1CF | 5.620467051 | 0.00022784 |
| CDHR1 | 1.65465824 | 0.000860821 |
| POLR3A | -1.443192258 | 7.88E-10 |
| CAMK2G | -0.22060072 | 0.012944816 |
| ADIRF | 3.193162302 | 0.001149801 |
| HTR7 | -0.946952768 | 9.51E-08 |

| | | |
|---|---|---|
| RPP30 | -0.385087212 | 0.00119816 |
| FRA10AC1 | 0.277980897 | 0.028584476 |
| ADD3 | 0.298540647 | 7.22E-05 |
| DNAJB12 | -0.81463247 | 1.20E-05 |
| EIF4EBP2 | -1.258248816 | 2.58E-10 |
| NPFFR1 | 4.870110483 | 0.0003411 |
| TCF7L2 | -0.430318976 | 3.96E-05 |
| MKI67 | -1.804172739 | 9.62E-13 |
| LRRC27 | 1.342296995 | 3.09E-05 |
| MTG1 | -0.292146535 | 0.003384608 |
| NKX6-2 | -1.069499862 | 0.034660032 |
| GSTO1 | 0.342137468 | 0.000226015 |
| TAF5 | -0.153214178 | 0.046339497 |
| PPRC1 | -0.652100598 | 3.82E-08 |
| ITPRIP | 1.77708306 | 5.37E-12 |
| CNNM2 | -1.280124443 | 9.88E-10 |
| PDCD11 | -0.517127611 | 1.68E-07 |
| ADAM12 | -0.677408402 | 0.007592354 |
| RGS10 | 0.526988108 | 0.005037725 |
| BTBD10 | -1.000954039 | 4.72E-07 |
| ADM | 0.512560542 | 0.000200502 |
| GAS2 | 1.660464711 | 0.013142363 |
| SLC5A12 | 1.035653934 | 0.014335136 |
| LIN7C | -0.64792042 | 4.75E-06 |
| PGAP2 | 0.287378461 | 0.013076931 |
| TUT1 | -0.157236422 | 0.045414211 |
| ZNF215 | -1.526262536 | 1.14E-07 |
| APIP | -0.282896377 | 0.026754214 |
| DGKZ | -0.455346998 | 2.38E-05 |
| TNKS1BP1 | 0.312532242 | 7.16E-05 |
| SSRP1 | 0.122663408 | 0.007506136 |
| PTPRJ | -0.44506494 | 9.53E-06 |
| ARFGAP2 | -0.52744168 | 1.91E-06 |
| CELF1 | -0.729741036 | 1.00E-08 |
| CCDC81 | 1.871579346 | 0.006547553 |
| SESN3 | 0.873953424 | 1.60E-07 |
| ENDOD1 | -0.849227517 | 3.12E-08 |
| KLHL35 | 4.908607973 | 1.34E-06 |
| SERPINH1 | 0.845504055 | 7.44E-09 |
| PAK1 | 0.19443634 | 0.01050163 |
| RPS3 | 0.388584937 | 1.76E-06 |
| ZC3H12C | -0.941009794 | 5.31E-08 |
| TTC12 | 1.992953999 | 1.73E-07 |
| NCAM1 | 1.311597771 | 0.007176814 |
| DRD2 | 1.854076545 | 4.28E-05 |
| NPAT | -0.164089123 | 0.021393177 |
| AASDHPPT | -0.307765171 | 3.05E-05 |
| SLX4IP | -0.22013686 | 0.04994198 |

| | | |
|---|---|---|
| LAMTOR1 | -0.320485151 | 0.000101425 |
| HYOU1 | 0.350299608 | 1.33E-06 |
| KAT14 | -0.283699379 | 0.00227756 |
| TKFC | 0.673486155 | 1.49E-06 |
| MTA2 | -0.752182395 | 5.84E-09 |
| TMEM138 | -1.438856051 | 7.77E-10 |
| FADS1 | 0.473424409 | 3.74E-07 |
| TMC2 | 8.964184317 | 2.53E-08 |
| ROM1 | 0.557973304 | 0.031211175 |
| INCENP | -1.372466041 | 2.46E-10 |
| ZP1 | 2.521557412 | 0.0061571 |
| FRG1BP | 0.417866555 | 0.013779992 |
| CPSF7 | -0.622696399 | 5.45E-08 |
| B3GAT3 | 0.419702809 | 2.07E-05 |
| EI24 | -0.446170247 | 2.33E-06 |
| CCDC15 | 0.698959209 | 3.84E-05 |
| CHEK1 | -1.374498844 | 5.31E-10 |
| ESAM | -1.180895298 | 0.000211114 |
| SIDT2 | 0.380944918 | 0.000875893 |
| TMEM25 | 2.85623075 | 1.13E-06 |
| TAGLN | 1.071781562 | 3.87E-05 |
| COMMD7 | -0.422629398 | 0.000550244 |
| C20orf144 | 3.05783036 | 0.000163161 |
| KIAA1755 | 1.766354418 | 9.28E-06 |
| OCSTAMP | 7.952441495 | 7.63E-07 |
| DSN1 | -0.47884793 | 5.48E-06 |
| SOGA1 | -1.450360532 | 5.58E-11 |
| CNBD2 | 0.579445816 | 0.012813713 |
| LSM14B | -0.823693717 | 1.49E-07 |
| CABLES2 | -0.653983767 | 9.31E-05 |
| TRPT1 | 1.271321034 | 1.08E-07 |
| NUDT22 | 0.397484137 | 0.001437859 |
| FERMT3 | 3.426589294 | 2.65E-07 |
| PLCB3 | -0.29510847 | 0.000330093 |
| MRPL49 | 0.163490393 | 0.0077252 |
| CDC42EP2 | 0.281145172 | 0.000788792 |
| FAU | 0.23139409 | 0.000294549 |
| TM7SF2 | 0.856534707 | 0.000708557 |
| VPS51 | 0.299421199 | 0.000858889 |
| ALDOA | 0.29100988 | 7.78E-07 |
| DOC2A | 1.024826288 | 9.07E-06 |
| TAOK2 | -0.532337661 | 3.26E-06 |
| TMEM219 | 0.357491045 | 0.004588559 |
| HMGA2 | -2.190667882 | 1.69E-12 |
| CNKSR2 | -1.490772237 | 3.17E-07 |
| PCDH15 | 2.564155203 | 0.000996027 |
| CWC15 | -0.541390094 | 9.07E-05 |
| FCGR1A | 5.079259066 | 2.10E-06 |

| | | |
|---|---|---|
| ARID5B | 1.436200378 | 1.95E-09 |
| DCUN1D2 | -0.544810445 | 2.56E-05 |
| TMCO3 | 0.177829572 | 0.024470525 |
| TMEM218 | 0.523818994 | 7.99E-05 |
| TIRAP | 0.523470433 | 0.000208921 |
| EEF1AKMT1 | 0.808541394 | 0.000416483 |
| LATS2 | 0.393303598 | 2.35E-05 |
| ADGRL3 | 0.578808672 | 0.029831057 |
| KIAA1328 | -0.775145504 | 0.002876099 |
| FAM124A | -0.773316246 | 0.003282813 |
| LYPD1 | -0.698292199 | 7.75E-06 |
| PDCD4 | -0.631307698 | 7.27E-07 |
| WDR17 | -0.368175526 | 0.007356942 |
| SPATA4 | 3.133374039 | 0.000708856 |
| VEGFC | -1.103961459 | 1.94E-05 |
| CNDP1 | 5.966357819 | 7.46E-06 |
| DLG2 | 1.196368672 | 0.000105644 |
| PRSS23 | -0.222279846 | 0.000657402 |
| MTMR12 | -0.703863742 | 9.68E-08 |
| DIXDC1 | -0.79349081 | 1.78E-06 |
| DLAT | -0.693929921 | 1.28E-07 |
| NKAPD1 | -0.255684276 | 0.000628622 |
| PTS | -0.509456285 | 5.74E-05 |
| PIP4K2A | -0.721412988 | 3.27E-06 |
| FREM2 | 0.276930853 | 0.000696473 |
| FOXO1 | 1.231409657 | 1.23E-08 |
| CRIM1 | -1.063640353 | 2.56E-08 |
| SEC24D | -0.671558672 | 1.50E-06 |
| DHX37 | -0.484708676 | 6.77E-06 |
| UBC | 0.251677367 | 1.55E-05 |
| ITPR1 | -0.816698617 | 4.89E-06 |
| TKTL2 | 7.00668098 | 2.14E-05 |
| PRSS53 | 0.389791372 | 0.010668823 |
| SLC7A11 | -1.664492789 | 8.04E-11 |
| NOCT | 0.302239602 | 0.003293582 |
| ENKUR | 5.824294963 | 3.98E-10 |
| GPR158 | -1.023267057 | 3.06E-06 |
| CACNA2D4 | 1.398871735 | 0.001287985 |
| DCP1B | 0.522974881 | 0.011858003 |
| CACNA1C | 4.058460792 | 0.002146554 |
| THRB | -0.948482629 | 8.19E-05 |
| NGLY1 | 0.299378379 | 0.001211911 |
| OXSM | 0.535662717 | 9.99E-06 |
| UEVLD | -0.760859671 | 2.95E-05 |
| TMEM86A | 2.63121594 | 7.01E-08 |
| C12orf45 | -0.43225267 | 0.017921735 |
| TMEM263 | 0.327926103 | 1.58E-05 |
| DLG5 | -0.882774022 | 5.25E-06 |

| | | |
|---|---|---|
| MAT1A | 2.9866604 | 0.00013434 |
| SLC2A13 | -0.235869974 | 0.00351801 |
| GXYLT1 | -2.01838299 | 1.14E-11 |
| TWF1 | -0.335547388 | 6.69E-05 |
| DIP2C | -0.464818099 | 0.007269256 |
| EIF4E | -0.615816735 | 1.35E-06 |
| TEX30 | -0.765602975 | 5.66E-07 |
| CSNK1G3 | -0.732320693 | 3.14E-08 |
| AL136982.1 | 1.830139634 | 8.28E-05 |
| SRFBP1 | -0.57541883 | 7.98E-06 |
| NPAS3 | 2.330113446 | 0.010053281 |
| FAM177A1 | -0.701395122 | 3.45E-06 |
| MBIP | -0.529048266 | 0.000635058 |
| MIPOL1 | -0.367798123 | 0.0349895 |
| TMEM18 | -0.536435085 | 1.39E-05 |
| NDUFC2 | -0.341458222 | 6.95E-05 |
| ADAMTS12 | -0.273411347 | 0.004429267 |
| NEK7 | -1.011196615 | 2.23E-09 |
| FER | -1.157993308 | 9.32E-09 |
| VIPAS39 | -0.594010513 | 0.000293119 |
| ANKRD50 | -0.362018751 | 3.65E-06 |
| UPF2 | -1.360361688 | 1.85E-09 |
| SCLT1 | -0.690691685 | 0.000180377 |
| C4orf33 | 0.658060557 | 0.000105033 |
| FRMD4A | -0.601352871 | 1.91E-06 |
| PTPRO | -1.114098456 | 0.00708933 |
| EPS8 | -0.397613449 | 1.92E-05 |
| VPS26B | -0.927740453 | 3.78E-09 |
| NCAPD3 | -0.46175068 | 1.73E-06 |
| VTI1A | -0.882425052 | 1.24E-06 |
| QDPR | -0.337246376 | 0.000530609 |
| FAM160B1 | -1.027995842 | 3.67E-08 |
| QTRT2 | -1.471426887 | 2.07E-10 |
| ZNF827 | -1.822037081 | 2.40E-09 |
| POU4F2 | 7.664990748 | 1.91E-06 |
| EDNRA | -1.907537313 | 0.000257683 |
| AKR1C2 | 0.921357461 | 0.001248327 |
| ADAM8 | 0.462656758 | 0.035778151 |
| INPP1 | -0.784343694 | 1.73E-05 |
| MFSD6 | -0.621700463 | 0.001610518 |
| RNF144A | -0.331245643 | 0.001367752 |
| ASAP2 | -1.481599849 | 1.13E-09 |
| ADAM17 | -0.19017764 | 0.005365114 |
| FLI1 | -0.475106721 | 0.031096063 |
| TMEM45B | 1.734090539 | 0.000977046 |
| WWC2 | -1.399882157 | 4.27E-10 |
| ACSL1 | -0.558786254 | 2.27E-06 |
| SLC25A4 | 0.268691528 | 0.000224012 |

| | | |
|---|---|---|
| AMN1 | 0.675004683 | 0.002106216 |
| BICD1 | -1.96304947 | 2.95E-11 |
| SAV1 | -0.991426091 | 6.07E-09 |
| CCDC122 | 1.955757434 | 2.66E-07 |
| GUF1 | -0.302833782 | 0.001051122 |
| SACS | -1.006677283 | 6.65E-11 |
| PABPC3 | 0.116904306 | 0.027508994 |
| CENPJ | -1.045881086 | 9.50E-09 |
| CCL28 | -1.31399208 | 3.53E-06 |
| GFRA1 | 3.717971379 | 2.11E-08 |
| CACUL1 | -1.7218212 | 3.52E-11 |
| TIAL1 | -1.369699891 | 8.21E-11 |
| BAG3 | -0.758834 | 1.25E-08 |
| RBM46 | 1.253376523 | 0.00291349 |
| SCHIP1 | -2.218688238 | 1.59E-09 |
| AP1S3 | -0.867539624 | 0.000702959 |
| TLCD4 | -0.289874872 | 0.030532431 |
| FAM168B | -0.667004347 | 1.18E-08 |
| PTPN14 | -1.42658619 | 5.89E-11 |
| AC073869.1 | -0.607851558 | 6.85E-06 |
| MGAT5 | -0.573643848 | 2.22E-07 |
| GPATCH11 | -0.615270342 | 3.28E-07 |
| HSPB8 | -0.349941632 | 0.02573259 |
| POU4F1 | -0.704892251 | 0.009773186 |
| SETBP1 | 0.396149263 | 0.012420714 |
| ARL14EP | -1.211636247 | 2.58E-08 |
| EPG5 | -0.585862024 | 0.000462139 |
| PSTPIP2 | 9.005375362 | 1.60E-08 |
| ATP5F1A | 0.193312937 | 0.000944054 |
| C18orf25 | -1.31905709 | 4.13E-10 |
| PDK1 | -0.61602465 | 6.94E-05 |
| PDE3B | -1.247707365 | 0.005649022 |
| TCF7L1 | 0.949028671 | 0.000438316 |
| TGOLN2 | -0.623090541 | 5.36E-09 |
| SH2D6 | 1.544280252 | 0.045896937 |
| UHMK1 | -1.041326095 | 4.38E-11 |
| POC5 | -0.662548023 | 0.000122138 |
| CWF19L2 | -0.996606178 | 3.86E-07 |
| JMY | 0.304558185 | 0.00013827 |
| HOMER1 | 0.31102187 | 0.000104546 |
| XRCC4 | -0.529306918 | 0.00218203 |
| ZNF773 | 0.568124556 | 0.000127724 |
| AC010326.1 | -1.778806518 | 4.60E-07 |
| SUV39H2 | -0.763144249 | 1.69E-07 |
| DCLRE1C | -0.797494424 | 0.000134995 |
| OLAH | 3.448605711 | 0.011814774 |
| RPP38 | -0.961967553 | 7.10E-06 |
| NMT2 | -0.267153405 | 0.02588673 |

| | | |
|---|---|---|
| ZNF837 | 2.509520332 | 3.69E-06 |
| USP12 | -0.795930313 | 1.03E-07 |
| CAMK4 | -0.474203366 | 4.71E-06 |
| TRIM36 | 0.650052683 | 3.96E-06 |
| ZFP36L2 | -2.339626416 | 2.21E-12 |
| PAN3 | -0.766664278 | 8.86E-07 |
| PLEKHH2 | 0.800179337 | 0.000100193 |
| PFKM | 0.160151544 | 0.010732639 |
| TMEM123 | -0.4076374 | 2.20E-07 |
| GRIA4 | 1.852148499 | 0.001209609 |
| IGSF10 | -2.105817137 | 0.000833992 |
| SPEF2 | 2.430287799 | 2.52E-08 |
| MBNL1 | -1.115820204 | 2.58E-10 |
| NADK2 | -0.776913136 | 2.24E-06 |
| GPD1L | -0.909930397 | 1.20E-08 |
| CCNO | -0.455204455 | 0.00509146 |
| DDX4 | 7.717822129 | 1.55E-06 |
| SLC30A6 | -0.289607125 | 0.000417542 |
| RASGRP3 | 1.983939349 | 0.000571727 |
| SAR1B | -0.490836526 | 1.74E-06 |
| GPR180 | 0.254069891 | 0.000516295 |
| WDR78 | 4.683422119 | 1.52E-06 |
| FARP1 | -0.221160228 | 0.000741399 |
| IFIT5 | -0.767924284 | 3.03E-08 |
| HHEX | -1.056044282 | 1.50E-07 |
| UTRN | -0.181501861 | 0.015871179 |
| PTPRK | -1.044673901 | 5.89E-07 |
| GGPS1 | 0.12375917 | 0.045731793 |
| ZNF117 | 1.776465641 | 4.34E-10 |
| MARVELD2 | 2.133407375 | 0.006676585 |
| MED21 | -2.296657472 | 1.12E-10 |
| PLOD2 | -0.654231703 | 2.74E-07 |
| ADGRA3 | -1.64129651 | 1.20E-10 |
| SREK1IP1 | -1.502488481 | 2.96E-09 |
| LGI2 | 1.404508686 | 0.009651057 |
| CWC27 | -0.337865689 | 0.003221344 |
| MR1 | 2.073955417 | 0.00018479 |
| SRP19 | -0.558292577 | 1.98E-06 |
| CENPH | 0.319298166 | 0.005037276 |
| CDYL | -1.312534704 | 2.38E-08 |
| CARHSP1 | -0.548588528 | 8.22E-07 |
| TXNDC11 | 0.556737738 | 2.23E-06 |
| DAB2 | -1.221947854 | 7.65E-09 |
| ANAPC1 | -0.350774254 | 0.00054195 |
| CAST | -0.237095853 | 0.001470305 |
| SCOC | 0.401423867 | 6.02E-06 |
| CLGN | 0.757531586 | 0.039611847 |
| CETN3 | 0.896358583 | 1.37E-06 |

| | | |
|---|---|---|
| SMARCA5 | -0.25497925 | 0.000499016 |
| SYCP2L | -0.88029503 | 0.028864678 |
| BMP6 | 0.83459356 | 4.40E-05 |
| RASSF3 | -1.059936463 | 5.58E-09 |
| HNRNPU | -0.294334579 | 9.53E-06 |
| RANBP2 | -0.476184293 | 2.60E-07 |
| AHCTF1 | -0.74112097 | 4.04E-09 |
| MERTK | 3.866346169 | 6.46E-08 |
| TMEM87B | -1.397327166 | 2.50E-08 |
| NR4A2 | 2.275192147 | 2.51E-10 |
| RBMS1 | -1.574931757 | 9.68E-10 |
| CD96 | 0.119546889 | 0.01869939 |
| ADGRF1 | -0.455913765 | 0.037256004 |
| FAM49B | -0.316791019 | 0.000280779 |
| ASAP1 | -1.100808304 | 2.12E-09 |
| TRAPPC8 | -0.324004568 | 0.002528204 |
| LINC00467 | 2.255066792 | 1.26E-05 |
| INO80C | 0.71016843 | 0.000277 |
| LPCAT1 | -1.323087821 | 3.01E-07 |
| PLEKHG4B | 4.101597933 | 0.00322149 |
| ADPRHL1 | 1.974818953 | 3.01E-06 |
| CMTM7 | -0.580203411 | 5.03E-06 |
| FBXL2 | 0.443935952 | 0.017977299 |
| UBP1 | -1.674055895 | 2.42E-11 |
| RPIA | -1.237887207 | 1.55E-08 |
| PTPRD | -1.166104938 | 0.028500808 |
| LURAP1L | 1.698879171 | 0.003716051 |
| CNKSR3 | 0.382653614 | 0.001021299 |
| GTF2E1 | 0.781701033 | 4.01E-07 |
| ZDHHC7 | -0.758246219 | 4.19E-08 |
| CMIP | -1.333702672 | 2.08E-08 |
| PID1 | -1.362547477 | 0.006731585 |
| TRIP12 | -0.194826895 | 0.001023704 |
| FBXO36 | 2.876896813 | 3.05E-07 |
| KCTD15 | -0.891863692 | 1.76E-08 |
| ZNF599 | 1.231541084 | 2.55E-06 |
| MCOLN2 | -0.609974383 | 0.001935971 |
| DDAH1 | -1.894980166 | 1.26E-10 |
| SREK1 | -0.793660542 | 5.41E-09 |
| CHD1 | -0.581727584 | 2.05E-07 |
| DGKE | 0.834123522 | 0.000985982 |
| HS2ST1 | -0.365821544 | 2.04E-05 |
| CACNA2D1 | -0.779667336 | 9.14E-06 |
| NUS1 | -0.646994597 | 1.22E-07 |
| PPP2R5E | -1.382372943 | 6.32E-10 |
| GRAP | 4.253046625 | 0.001212281 |
| SLC5A10 | 2.402644501 | 0.000139463 |
| IMPACT | -0.471582032 | 5.48E-05 |

| | | |
|---|---|---|
| SDHAF4 | 3.068339608 | 3.03E-07 |
| THY1 | -2.69007094 | 0.003913372 |
| TBCEL | 0.465322421 | 2.66E-05 |
| JPH3 | 3.340018928 | 1.05E-06 |
| OTULIN | -1.670125914 | 8.38E-11 |
| UBASH3B | -1.583102817 | 3.58E-11 |
| ROBO4 | 1.294792229 | 9.97E-07 |
| ROBO3 | 0.706851699 | 0.000457655 |
| PANX3 | 1.683005642 | 0.001685527 |
| TBRG1 | 0.274807101 | 0.000284985 |
| NRGN | -0.174596547 | 0.038499292 |
| RETREG1 | -0.640154473 | 0.000140004 |
| CDH12 | -0.760221757 | 7.20E-05 |
| TOMM70 | -1.004976559 | 4.25E-10 |
| ABI3BP | 1.783004579 | 2.21E-05 |
| ANGPT1 | -0.981014948 | 1.42E-07 |
| PITPNC1 | 0.277789479 | 0.002148538 |
| CC2D1B | -0.237290844 | 0.003108788 |
| PRKCA | -0.972274469 | 3.63E-08 |
| LRRK1 | -0.637097097 | 0.000160383 |
| ABCA10 | 1.068166975 | 5.40E-05 |
| MIA3 | 0.170094188 | 0.005024497 |
| DISP1 | -1.260661786 | 1.76E-06 |
| NEIL2 | 0.514482873 | 0.000307379 |
| OBSCN | 1.607522827 | 1.77E-06 |
| LONRF1 | -1.052372863 | 1.59E-06 |
| TRIM11 | -0.389971805 | 3.42E-05 |
| ENAH | -0.756905204 | 4.98E-09 |
| SH3RF1 | -1.412518745 | 9.42E-08 |
| BUB3 | -1.030555284 | 2.82E-10 |
| MMP21 | 5.551502956 | 2.32E-05 |
| CNTNAP3B | -0.286042836 | 0.033010177 |
| SRSF12 | -1.426165973 | 0.025939978 |
| PDLIM3 | 0.359034788 | 0.00379741 |
| SORBS2 | 0.496566661 | 0.002849013 |
| CEP170P1 | -0.694147883 | 0.000444554 |
| BTG3 | -0.855626856 | 8.64E-07 |
| PDE1C | -1.329562883 | 1.13E-10 |
| RABGEF1 | -0.37244906 | 4.59E-05 |
| JAM2 | 1.816043199 | 0.005433697 |
| GABPA | -0.836125921 | 7.92E-08 |
| ADAMTS1 | -0.679290037 | 0.002905834 |
| TSEN2 | -0.951500203 | 2.54E-06 |
| SLFN13 | 1.128228212 | 0.000862226 |
| XPC | 0.69443783 | 9.52E-08 |
| FLCN | 1.039300639 | 8.98E-08 |
| DPH3 | -0.382507772 | 4.44E-05 |
| OXNAD1 | -0.539277371 | 0.000309809 |

| | | |
|---|---|---|
| PLCL2 | -2.599815713 | 1.10E-05 |
| CXXC1 | -0.502442531 | 7.82E-06 |
| SKA1 | -1.597646814 | 7.61E-07 |
| PPP4R1 | -0.958547252 | 3.50E-08 |
| CCDC144B | 0.626784935 | 1.59E-06 |
| MPPE1 | 0.823142907 | 3.52E-05 |
| USP43 | -0.71173238 | 0.001349252 |
| RAB6B | 0.446916847 | 0.005256567 |
| EME1 | 0.469220739 | 1.21E-05 |
| EPHB1 | -1.724004203 | 1.27E-08 |
| ACSS1 | -0.37303202 | 0.024295713 |
| ANKRD40 | -1.083837605 | 7.10E-10 |
| ZNF18 | 1.963334854 | 7.22E-10 |
| VOPP1 | -0.872462105 | 4.52E-07 |
| SEPTIN14 | -0.96518599 | 0.031404364 |
| APOOL | -0.758163901 | 8.64E-07 |
| CYP2U1 | 2.072230223 | 1.89E-08 |
| RSPH10B | -0.232496922 | 0.029825999 |
| FBXL18 | -1.001286972 | 6.23E-08 |
| AK9 | 2.022380005 | 6.16E-05 |
| KLF10 | 0.867035731 | 3.14E-09 |
| AZIN1 | -0.455907621 | 1.43E-07 |
| ATP6V1C1 | -0.796237662 | 1.25E-07 |
| OTUD6B | -1.06650268 | 1.37E-09 |
| GTF3C6 | -0.33427148 | 0.000237655 |
| AGPAT5 | -0.837732281 | 1.77E-08 |
| MMS19 | 0.396811943 | 4.71E-05 |
| ZFYVE27 | -0.886065626 | 2.87E-08 |
| GOLGA7B | 2.084670146 | 4.25E-06 |
| TRMT44 | -0.994928386 | 4.36E-05 |
| SLC25A28 | -0.405975878 | 0.001320528 |
| HSPA13 | 0.470890917 | 1.30E-06 |
| USP25 | -0.948816048 | 1.59E-05 |
| GRAMD2B | -0.61905389 | 0.006067929 |
| ZCCHC10 | -0.457075026 | 2.62E-05 |
| C16orf87 | 0.374050182 | 0.001063572 |
| RHOC | 0.40652069 | 1.13E-05 |
| PPM1J | 0.938068398 | 0.031924593 |
| DBI | 0.392448897 | 0.001065598 |
| SLC16A1 | -1.11767705 | 1.99E-10 |
| HEATR3 | -0.6454881 | 4.97E-06 |
| TRIM74 | 2.911969432 | 3.84E-05 |
| NIFK | -0.452897321 | 7.22E-06 |
| OXA1L | 0.221180193 | 0.006802289 |
| SLC7A7 | 1.21145782 | 0.01657269 |
| LARP1 | -1.009881883 | 1.31E-10 |
| CNOT8 | -0.472749784 | 9.45E-06 |
| SETD9 | 1.411095396 | 9.12E-07 |

| | | |
|---|---|---|
| MIER3 | -0.902495001 | 1.32E-08 |
| NUP205 | -0.549491278 | 4.68E-07 |
| ZKSCAN2 | -0.575174625 | 4.20E-06 |
| RBM45 | 0.536882165 | 0.001262101 |
| TTN | 1.892630881 | 2.56E-06 |
| PDIA4 | 0.434126314 | 5.73E-06 |
| KCTD18 | 1.408465474 | 5.17E-08 |
| FAM126B | -0.263058076 | 0.001920049 |
| FLACC1 | 1.539058262 | 0.002202394 |
| TMEM237 | -0.580492434 | 0.000409042 |
| FZD7 | 0.715333579 | 9.47E-07 |
| FMN2 | -1.231924119 | 3.13E-08 |
| RNF20 | -0.218628649 | 0.000718813 |
| PPARGC1B | -2.401481039 | 1.97E-09 |
| SLC26A2 | -0.29002655 | 3.12E-05 |
| LSM11 | -2.358604576 | 3.13E-11 |
| SAXO1 | 3.800931207 | 1.93E-06 |
| RRAGA | -0.174959206 | 0.002158732 |
| SLC24A2 | 2.169146842 | 2.57E-06 |
| PXYLP1 | -0.357600441 | 0.000609427 |
| RASA2 | -0.480058975 | 5.19E-05 |
| TMBIM4 | 0.460242904 | 0.000264334 |
| VBP1 | -0.149885574 | 0.013145466 |
| RAB39B | 2.086010386 | 5.94E-08 |
| MICU3 | -1.202023443 | 0.000174671 |
| VPS37A | -0.37514729 | 0.000311414 |
| KIF5A | 0.66059793 | 0.001088629 |
| MAGEA8 | 7.304113185 | 6.39E-06 |
| PSD3 | -0.815104027 | 1.39E-06 |
| CARNMT1 | -1.420526715 | 1.09E-09 |
| MCU | -0.753677156 | 1.52E-06 |
| ELMSAN1 | 0.341853535 | 4.43E-05 |
| CFAP70 | 3.575896318 | 7.11E-08 |
| GNAQ | -0.784740505 | 5.25E-08 |
| GPR61 | 3.57317602 | 2.53E-07 |
| MMP16 | -0.781150797 | 5.41E-09 |
| ADK | -0.307865356 | 0.009599845 |
| KCNMA1 | 0.565181297 | 0.001451881 |
| DCK | -1.004185366 | 6.28E-08 |
| ALX3 | -0.551293063 | 0.039315154 |
| DPY19L4 | -0.438321035 | 3.67E-05 |
| NDUFAF6 | 0.923356979 | 1.50E-05 |
| ART3 | 0.812893711 | 8.36E-05 |
| WHAMM | -0.349124819 | 0.020531478 |
| N6AMT1 | -2.306587777 | 6.58E-09 |
| RWDD2B | 1.378266755 | 1.40E-08 |
| USP16 | -0.574578996 | 1.30E-06 |
| CCT8 | 0.166619324 | 0.003959051 |

| | | |
|---|---|---|
| NAA11 | -0.59048746 | 8.41E-07 |
| BACH1 | 1.064331799 | 1.73E-09 |
| TIAM1 | -0.847021506 | 2.16E-07 |
| RPGR | 0.37209719 | 0.00397584 |
| PCGF6 | -0.380373453 | 0.001198117 |
| ANKRD9 | 0.460065823 | 0.001688571 |
| SFR1 | 0.704624008 | 0.000192684 |
| SFXN2 | -1.204068221 | 1.52E-06 |
| ATP5MPL | 0.372972909 | 0.00025295 |
| FGF18 | 1.840695647 | 0.0038134 |
| PCDH1 | 0.382299437 | 6.44E-05 |
| UQCRB | 0.688519531 | 2.18E-07 |
| PTDSS1 | -0.387464147 | 3.85E-06 |
| RPL30 | 0.304236205 | 5.79E-05 |
| FAM122C | 0.964750899 | 0.002123339 |
| SUPV3L1 | -0.805654978 | 1.19E-06 |
| FAM122B | -0.294029547 | 0.000218297 |
| EEF1A1 | 0.418759535 | 5.08E-06 |
| FBXO43 | 0.528916818 | 0.000599023 |
| HK1 | 0.222199636 | 0.000948244 |
| TYSND1 | -0.660016216 | 4.18E-05 |
| PHF6 | -0.276749043 | 0.000168694 |
| CD109 | 0.144747669 | 0.004175701 |
| UBE2L6 | 0.203202422 | 0.013093218 |
| ZDHHC5 | -0.995410334 | 8.82E-10 |
| ZFAND3 | -0.168484866 | 0.029894406 |
| NPTN | -0.525242786 | 2.25E-06 |
| KAT6B | -0.271212796 | 0.000968177 |
| SAMD8 | -0.178243316 | 0.007487758 |
| RAB11FIP1 | -0.233582642 | 0.020444318 |
| UTP14A | -0.631187717 | 1.72E-07 |
| AIFM1 | -0.270123232 | 0.002156639 |
| MAPK13 | 1.450270349 | 1.91E-05 |
| BAG4 | -0.149201982 | 0.04837589 |
| TBC1D31 | -0.475900707 | 2.93E-05 |
| ATAD2 | -0.894942005 | 2.78E-09 |
| FBXO32 | 1.96524811 | 1.47E-09 |
| ZNF689 | -2.236001001 | 8.65E-10 |
| PRR14 | 0.713965085 | 8.27E-07 |
| FBRS | 0.142119638 | 0.036331667 |
| FRRS1 | -0.363523772 | 0.045871044 |
| PHKG2 | 0.565163942 | 2.17E-06 |
| MFSD14A | -0.63971247 | 3.37E-07 |
| SASS6 | -2.103189278 | 1.93E-09 |
| ITGAD | 2.815872801 | 0.001879651 |
| MALSU1 | -1.027759213 | 1.96E-09 |
| VPS8 | 0.642462884 | 0.000591243 |
| GALK2 | 0.25041021 | 0.025696401 |

| | | |
|---|---|---|
| MPV17L | -1.198393489 | 4.42E-06 |
| BUB1B | -1.604752415 | 9.35E-10 |
| EIF4A2 | 0.716970631 | 1.23E-09 |
| BRPF1 | -0.250315164 | 0.00213674 |
| TATDN2 | -1.038971748 | 1.12E-09 |
| GHRL | 1.299676686 | 0.036581463 |
| SEC13 | 0.246946319 | 0.000544595 |
| FAM92A1P1 | -0.694176386 | 0.001058833 |
| EXOG | -1.926019495 | 3.24E-10 |
| ZFYVE9 | -1.244657627 | 9.10E-09 |
| ATP2B2 | 7.341240332 | 6.06E-06 |
| SMG1 | -0.178355668 | 0.000539611 |
| FCHO2 | -1.013195393 | 1.90E-07 |
| RBPMS | -0.681937454 | 0.000124072 |
| ODR4 | -0.752000896 | 5.95E-06 |
| CPT2 | 0.315890155 | 0.000235463 |
| NECAP2 | -0.847265212 | 1.32E-07 |
| LRP8 | -1.018689873 | 2.07E-09 |
| CDCP2 | 5.088905965 | 0.000136352 |
| PAXIP1 | -0.906472473 | 1.73E-06 |
| STEAP2 | -0.596532168 | 0.049946939 |
| CLDN12 | -0.538892978 | 4.47E-07 |
| MMP14 | 0.453859895 | 2.10E-06 |
| FZD1 | -0.446265883 | 2.08E-05 |
| GATAD1 | -0.960910095 | 2.96E-08 |
| ZFHX2-AS1 | 0.540707901 | 0.001204067 |
| CLEC18A | 2.397498206 | 0.000118453 |
| DHRS4 | 0.873995899 | 0.002906886 |
| CLEC18C | 4.210911454 | 0.003693928 |
| ST3GAL2 | -0.368325109 | 0.000861481 |
| DHRS1 | 0.74585807 | 0.000465097 |
| CACNA1D | 0.699614633 | 0.041425182 |
| ARSL | 0.814081876 | 0.000783228 |
| AASDH | 1.561070947 | 8.34E-10 |
| ZNF19 | 2.124603118 | 2.84E-07 |
| RNF111 | -0.297066895 | 0.000338911 |
| CCNB2 | -0.877410258 | 1.49E-08 |
| MYO1E | -0.217796814 | 0.000957514 |
| APPL1 | -1.259659466 | 5.78E-10 |
| VPS26C | -1.255115981 | 3.53E-09 |
| DYRK1A | -0.67524918 | 3.84E-07 |
| KCNJ15 | 2.567831374 | 0.00143414 |
| ETS2 | -0.203922104 | 0.048472337 |
| LCA5L | 2.308664905 | 2.54E-07 |
| TMEM164 | -0.620055987 | 5.82E-07 |
| TAB3 | -0.520798523 | 4.66E-07 |
| C9orf43 | 1.875179327 | 1.26E-05 |
| PALM2AKAP2 | -0.423767287 | 2.12E-06 |

| | | |
|---|---|---|
| ZNF618 | -0.817981782 | 1.59E-07 |
| TMEM268 | -1.849323777 | 3.38E-09 |
| SNX22 | 0.268927541 | 0.000631366 |
| UBN2 | -0.346205586 | 6.35E-05 |
| BRAF | -0.792437859 | 1.63E-07 |
| ACAN | 0.803714897 | 0.000117688 |
| PSMG3 | -0.15433577 | 0.043355319 |
| WDR19 | 1.010073817 | 2.64E-08 |
| SLC37A3 | -0.83956583 | 2.36E-08 |
| FMNL2 | -1.049236757 | 1.01E-09 |
| GAREM2 | -0.656972833 | 0.002950693 |
| SPPL3 | -0.511921087 | 1.91E-05 |
| TNFRSF14 | 1.06547353 | 0.007453216 |
| PANK4 | -0.321942768 | 0.02785207 |
| C12orf43 | -0.839132675 | 5.92E-07 |
| RER1 | -0.322750279 | 0.000138221 |
| RADIL | 1.611104302 | 0.009798576 |
| SKI | -0.239981558 | 0.000188049 |
| WIPI2 | -0.247918991 | 0.000727438 |
| LDLRAP1 | -1.085864189 | 6.68E-08 |
| AGAP1 | -0.585469226 | 6.59E-06 |
| KRTCAP3 | 1.982640914 | 6.59E-05 |
| ANKRD61 | 1.082431642 | 0.00626801 |
| PAFAH2 | -1.104110144 | 1.21E-06 |
| EXTL1 | -0.418967232 | 0.000706082 |
| BABAM2 | 0.542518725 | 0.000645037 |
| WDR66 | 1.066654948 | 0.001417058 |
| DUSP2 | 4.042508956 | 2.57E-13 |
| UBXN11 | 0.521155065 | 8.52E-06 |
| NLRP14 | 2.412985424 | 0.006035353 |
| PTPDC1 | -0.901488245 | 4.64E-06 |
| GALNT14 | -0.636220787 | 0.000920846 |
| HPD | 3.625651194 | 6.42E-05 |
| RHPN1 | 0.684974998 | 1.35E-05 |
| TPRG1L | 0.386827629 | 0.000242401 |
| XDH | -0.932057095 | 0.007901745 |
| CNNM4 | 0.491800742 | 6.28E-06 |
| EYA3 | -0.595006468 | 8.38E-07 |
| DZIP1L | -1.341462899 | 4.73E-05 |
| FANCC | -0.949295251 | 1.60E-06 |
| MRAS | 0.547289783 | 0.00134604 |
| WASF2 | -0.283867761 | 0.00010125 |
| ABHD3 | -0.598603605 | 7.77E-05 |
| FAIM | -1.125937452 | 2.62E-05 |
| COLEC12 | 1.694785862 | 4.72E-05 |
| RNF207 | -0.977368976 | 2.55E-05 |
| CUL4B | -0.609196892 | 2.52E-07 |
| GPR153 | 1.023377874 | 1.78E-08 |

| | | |
|---|---|---|
| SLC13A3 | 4.374534172 | 3.17E-10 |
| RHBDL2 | 1.830567287 | 0.000266436 |
| SHROOM4 | -0.632471568 | 4.04E-05 |
| H2BC5 | 3.231022003 | 3.45E-09 |
| H4C8 | 1.968732668 | 0.00150422 |
| MITD1 | -0.269440309 | 0.045670238 |
| EIF5B | -0.269435674 | 0.000103987 |
| RIBC1 | 2.215400414 | 1.20E-05 |
| CNOT11 | -1.019670304 | 2.56E-08 |
| KCNB1 | 6.901779028 | 4.80E-09 |
| NRG2 | -1.28756681 | 7.57E-05 |
| AHCYL2 | -0.659879985 | 5.39E-06 |
| B4GALT5 | -1.44945233 | 3.12E-11 |
| SPATA2 | 0.200800985 | 0.005870963 |
| FAM86C1 | 1.385126843 | 3.17E-08 |
| DNAH3 | 1.940167828 | 0.000122277 |
| PPP1R9A | -0.976578849 | 1.92E-06 |
| ZC3H18 | -1.10577795 | 5.93E-08 |
| GDPD5 | 1.47120382 | 2.32E-08 |
| ALAS2 | 4.851543887 | 2.03E-07 |
| PPP1R15B | -0.168060855 | 0.001475453 |
| COPG2 | 0.296832234 | 0.002733991 |
| EMSY | -1.202908966 | 2.87E-08 |
| PAGE5 | 0.361938885 | 4.73E-05 |
| GPAT4 | -0.466113937 | 3.87E-06 |
| PKD1L1 | 3.812561085 | 2.24E-05 |
| ELK4 | -1.339647471 | 1.18E-10 |
| SLAMF8 | 5.07355056 | 1.33E-08 |
| SLC45A3 | -0.932552111 | 4.02E-07 |
| NBL1 | -0.157784234 | 0.0187413 |
| F11R | 0.344800257 | 0.005899728 |
| USF1 | -0.676575569 | 5.12E-07 |
| SPATA2L | 0.8142331 | 5.09E-05 |
| NIT1 | 0.998711076 | 8.58E-08 |
| DEDD | -0.910600267 | 3.51E-08 |
| ZNF276 | -1.382925551 | 1.19E-09 |
| B4GALT3 | -0.14860874 | 0.021380462 |
| ADAMTS4 | 0.999040355 | 0.001691134 |
| NDUFS2 | 0.176962963 | 0.010769908 |
| TOMM40L | -0.815864181 | 6.76E-07 |
| MPZ | 0.763566142 | 0.000130409 |
| CCAR2 | -0.33597226 | 3.22E-05 |
| CACHD1 | -1.155119489 | 1.03E-06 |
| CDC42SE2 | -1.275860529 | 1.89E-09 |
| RAPGEF6 | -1.229104032 | 4.13E-09 |
| EPB41 | -0.896089492 | 5.88E-08 |
| MIS18A | -0.549469552 | 2.87E-05 |
| ALG8 | -0.164897837 | 0.039611847 |

| | | |
|---|---|---|
| CFAP298 | -0.490356006 | 1.30E-05 |
| SYNJ1 | -0.398922388 | 7.39E-05 |
| PAXBP1 | -1.008131241 | 5.82E-07 |
| MRPL10 | 0.862875723 | 1.59E-08 |
| GART | -0.215315807 | 0.001571233 |
| SON | -0.695732111 | 1.15E-09 |
| DONSON | -0.39670914 | 0.000247029 |
| SV2A | -0.481376098 | 6.58E-06 |
| LAD1 | 0.830954691 | 0.005842197 |
| STC1 | 2.291007472 | 3.38E-11 |
| TNNI1 | 5.197956058 | 1.71E-07 |
| CSRP1 | 0.600915487 | 2.48E-07 |
| HOXB13 | -0.701469211 | 0.000100861 |
| KCNE2 | 3.761942163 | 8.16E-06 |
| RCAN1 | 0.503211023 | 3.02E-07 |
| UBE2Z | -0.120404048 | 0.018976605 |
| SNF8 | 0.773049506 | 3.69E-07 |
| CCDC24 | 0.610591346 | 0.017102444 |
| RUNX1 | -1.003147895 | 1.71E-09 |
| IGF2BP1 | -2.207819649 | 3.86E-12 |
| AC005041.1 | 1.623450883 | 2.50E-07 |
| MORC3 | 0.672231993 | 7.02E-07 |
| CHAF1B | -0.640250219 | 0.000133888 |
| HLCS | -0.492723442 | 7.67E-05 |
| GOLGA6A | 4.664896943 | 0.001285229 |
| ARHGAP27 | 0.877621345 | 0.001058833 |
| ADPGK | -0.159111161 | 0.005537655 |
| CYB5R1 | 0.547272419 | 5.38E-06 |
| PSMD4 | 0.424325052 | 2.99E-05 |
| PSMB4 | 0.270905651 | 1.46E-05 |
| BTG2 | 3.671593281 | 7.83E-13 |
| HK2 | -1.209715971 | 2.43E-10 |
| C1R | 0.434495339 | 0.025309607 |
| CELF3 | 6.10861209 | 4.12E-06 |
| ALDH4A1 | 0.447763745 | 0.002177492 |
| STARD9 | -0.233021323 | 0.024688483 |
| THEM4 | -0.166052987 | 0.014536568 |
| UBR1 | -0.371384074 | 2.86E-05 |
| MED8 | -0.899212275 | 6.43E-08 |
| ISL2 | 0.573748391 | 0.024214426 |
| RSPRY1 | -0.673127266 | 4.26E-07 |
| CCDC17 | 3.638393099 | 3.46E-07 |
| NAE1 | -0.231842551 | 0.002378919 |
| TMEM69 | 0.69211785 | 2.08E-06 |
| DRC7 | 7.976729215 | 6.36E-07 |
| EFCAB14 | -1.023577685 | 2.21E-09 |
| SPON2 | 1.195559235 | 2.68E-06 |
| CTBP1 | -0.62576352 | 2.22E-06 |

| | | |
|---|---|---|
| TPPP3 | 0.87721978 | 0.00994635 |
| ZFYVE28 | 1.445629251 | 2.27E-05 |
| CARMIL2 | 0.5433171 | 0.006464974 |
| C16orf86 | 2.043618324 | 7.46E-06 |
| FAM131B | 0.851046731 | 0.014195079 |
| RGS12 | -0.678341499 | 0.000192193 |
| ZYX | -0.418527483 | 2.68E-06 |
| ABR | -0.416667387 | 0.009913582 |
| LYPD5 | 2.042178652 | 4.45E-05 |
| CCDC107 | -1.217968968 | 4.71E-06 |
| ZNF222 | 1.495750505 | 1.67E-08 |
| NPR2 | 1.011051914 | 7.19E-05 |
| ZNF233 | 3.658983535 | 2.53E-09 |
| GNE | -0.602142 | 2.22E-07 |
| CALM3 | -0.504576863 | 1.20E-07 |
| DFFA | -0.227928096 | 0.00045631 |
| CCDC28B | 1.402911365 | 2.80E-07 |
| IQCC | 0.622121002 | 7.78E-05 |
| BSDC1 | -0.467529606 | 1.11E-05 |
| ZBTB8A | -0.912827602 | 0.000122463 |
| ATAD3B | -0.645359834 | 1.63E-06 |
| SSU72 | -0.298655125 | 0.00030471 |
| UBE2J2 | -0.761358483 | 1.57E-07 |
| ZNF362 | 0.89700035 | 2.50E-05 |
| NR2F6 | -0.768412534 | 2.45E-07 |
| CCDC58 | 0.310852022 | 0.043441442 |
| VMA21 | -0.184569809 | 0.008303248 |
| CILP2 | 2.288829717 | 8.32E-09 |
| FAM86C2P | 2.467440385 | 8.31E-07 |
| SLC37A1 | -1.08495064 | 0.027173478 |
| WDR4 | -0.966105281 | 8.76E-06 |
| NDUFV3 | 0.344137317 | 0.001934523 |
| PKNOX1 | -1.484275859 | 3.13E-08 |
| CBS | 0.451114087 | 0.007106726 |
| U2AF1 | -0.434774465 | 3.27E-05 |
| RRP1B | -1.162741251 | 7.00E-10 |
| PDXK | -1.118185808 | 9.59E-09 |
| G6PD | 0.169779007 | 0.004885223 |
| CSTB | 0.351567757 | 5.11E-05 |
| RRP1 | -1.256927085 | 1.41E-07 |
| AGPAT3 | -0.556182503 | 1.05E-06 |
| TRAPPC10 | -0.771097625 | 2.61E-07 |
| ICOSLG | 1.720228961 | 0.001850575 |
| CFAP410 | -0.563199393 | 0.000593882 |
| ZNF66 | 0.716711946 | 0.004383391 |
| FAM207A | -0.969905898 | 0.000186449 |
| RALGDS | -0.186752696 | 0.028099518 |
| LSS | 0.355189484 | 0.014286785 |

| | | |
|---|---|---|
| VAV2 | -1.075116395 | 1.67E-08 |
| MCM3AP | -0.387387862 | 1.34E-05 |
| C21orf58 | -0.900653903 | 4.21E-05 |
| PCNT | -0.671658772 | 2.30E-07 |
| DIP2A | -0.557466395 | 6.70E-05 |
| S100B | 0.89746689 | 2.81E-06 |
| PRMT2 | 0.184539489 | 0.021765289 |
| ADAMTS13 | 2.65257976 | 0.000968676 |
| CACFD1 | -0.450264671 | 0.049418918 |
| SLC2A6 | 0.971563188 | 0.000187242 |
| ZNF761 | 0.551788742 | 1.70E-06 |
| ZNF714 | -0.608387514 | 1.26E-06 |
| GPSM1 | -0.188370453 | 0.025224279 |
| C19orf47 | -1.120188529 | 1.22E-06 |
| CFAP157 | -0.379238161 | 0.013119475 |
| ST6GALNAC6 | -0.319827405 | 0.013503461 |
| SHKBP1 | 0.17364844 | 0.027648912 |
| ZDHHC12 | 0.238641517 | 0.021711342 |
| PKN3 | -0.450702831 | 0.002488771 |
| AC020929.1 | 2.913531967 | 0.00093684 |
| COX6B2 | 5.698108633 | 7.88E-09 |
| TAOK1 | -1.281448048 | 8.46E-11 |
| MED27 | -0.269603969 | 0.029629676 |
| DEDD2 | 0.854179008 | 1.46E-07 |
| SIK3 | -0.216474451 | 0.009676064 |
| NEK8 | 1.890415895 | 3.16E-07 |
| TLCD1 | 0.946631076 | 4.53E-05 |
| PCSK7 | -1.226661311 | 2.11E-09 |
| SAFB | -0.27685112 | 0.000158593 |
| S100A1 | 0.883978987 | 1.93E-05 |
| CHTOP | -0.570044414 | 1.29E-07 |
| CXCR5 | -1.528239694 | 2.28E-08 |
| ZBTB7B | 0.424062168 | 0.000788792 |
| FLAD1 | -0.139150909 | 0.041038458 |
| SHC1 | -0.277494422 | 1.88E-05 |
| VPS11 | 0.430045716 | 2.98E-05 |
| ADAR | -0.611509676 | 5.16E-09 |
| IL6R | -0.55250785 | 0.000194049 |
| UBE2Q1 | -0.546996851 | 3.34E-07 |
| CHRNB2 | 2.553070202 | 1.34E-07 |
| ANO10 | 0.281885041 | 0.013786906 |
| FDPS | 0.29233949 | 0.000291752 |
| RUSC1 | 0.357925757 | 0.000277034 |
| GBAP1 | 1.789394177 | 3.76E-08 |
| PAQR6 | 0.631884233 | 6.32E-05 |
| PMF1 | 0.181416344 | 0.027588342 |
| LMNA | 0.134863357 | 0.031989299 |
| NBEAL2 | -0.327697213 | 0.000969602 |

| | | |
|---|---|---|
| CCDC12 | -0.622900403 | 0.000958209 |
| PPP1R35 | 0.339278895 | 0.023516833 |
| GPATCH4 | -0.834033316 | 7.11E-09 |
| FGFR4 | 0.534341649 | 0.000244791 |
| CYP3A4 | 4.718709069 | 0.000356685 |
| NACC1 | -0.837146859 | 1.91E-08 |
| LY6K | -0.542272554 | 7.88E-06 |
| IER2 | 1.5291962 | 5.95E-12 |
| ZNF394 | -0.230093917 | 0.004540222 |
| CPSF4 | -0.541250275 | 2.86E-05 |
| TONSL | -0.27046807 | 0.000790233 |
| RECQL4 | 0.155448331 | 0.014161474 |
| LRRC14 | -0.517418095 | 5.77E-06 |
| ZNF333 | 1.20411515 | 3.12E-09 |
| FOXH1 | 0.934974396 | 0.00134604 |
| ORAI2 | -0.239762572 | 0.000696007 |
| ALKBH4 | -1.551088165 | 4.14E-06 |
| MRNIP | 1.120303868 | 1.13E-10 |
| SQSTM1 | 1.203811018 | 1.25E-11 |
| MGAT4B | 0.125151373 | 0.023438921 |
| RPL8 | 0.431677077 | 9.27E-06 |
| MAML1 | -0.534142473 | 4.48E-07 |
| MFSD12 | -0.582776593 | 7.28E-08 |
| AC007663.1 | 3.330473407 | 0.0187413 |
| CCDC116 | 3.514779298 | 1.08E-07 |
| DVL3 | -0.321660765 | 0.000140403 |
| AP2M1 | 0.25856378 | 0.000135689 |
| ABCF3 | -0.276826001 | 0.001887834 |
| PCYT1A | -1.001759436 | 4.56E-09 |
| FBXO27 | 0.520342293 | 8.57E-05 |
| U2AF1L4 | -0.497780432 | 0.023724665 |
| THAP8 | 1.283011955 | 3.52E-05 |
| ZNF382 | 5.640093108 | 5.94E-07 |
| LRRC56 | 0.827619205 | 0.037840007 |
| PLXDC1 | 1.94891194 | 0.000249441 |
| GRIN2C | 2.18414314 | 8.04E-07 |
| FDXR | 0.951221457 | 2.98E-06 |
| SAP30BP | -0.802285428 | 1.07E-07 |
| ACOX1 | 0.156574442 | 0.043005293 |
| PRPSAP1 | -0.850640438 | 6.42E-07 |
| CYGB | 2.825281853 | 0.001822047 |
| SRSF2 | -1.987165855 | 9.00E-13 |
| ZNF577 | 0.372916675 | 0.019629773 |
| TMEM143 | 1.063481749 | 0.000343253 |
| ALDH16A1 | 0.322314052 | 0.000535351 |
| ITGA5 | 0.28577471 | 8.91E-05 |
| SIGLEC16 | 3.65966389 | 3.73E-05 |
| LSM12 | -1.282710688 | 3.05E-09 |

| | | |
|---|---|---|
| JOSD2 | 0.509641015 | 0.006735481 |
| FAM171A2 | 0.881075403 | 0.000328653 |
| DBF4B | -1.432525293 | 8.94E-09 |
| PLCD3 | 0.319398268 | 0.000259539 |
| FMNL3 | -0.960479538 | 5.03E-09 |
| RACGAP1 | -1.092197865 | 4.90E-10 |
| LARP4 | -0.590168782 | 5.61E-08 |
| GRASP | 2.376830396 | 1.44E-07 |
| RAVER1 | -0.631334465 | 2.38E-07 |
| SPC24 | -1.561486192 | 3.36E-08 |
| ALOX15 | 5.699998423 | 1.36E-05 |
| TREML1 | 8.155890512 | 3.43E-07 |
| ADCY10P1 | 1.341240168 | 2.98E-05 |
| ZNF653 | -0.803591467 | 0.00102381 |
| MED11 | -0.385510401 | 0.019179614 |
| RNASEK-C17orf49 | 0.386091571 | 2.50E-05 |
| TNFSF13 | 1.611716411 | 2.00E-06 |
| SENP3 | -0.413766397 | 4.25E-06 |
| FGF11 | 1.27413967 | 1.60E-05 |
| RPL26 | 0.2839965 | 3.32E-05 |
| POLR3K | -0.492913805 | 1.56E-05 |
| SNRNP25 | -0.569704357 | 1.57E-05 |
| WDR90 | -0.281456733 | 0.00167478 |
| JMJD8 | 0.213408366 | 0.001082646 |
| SPSB3 | 0.701229102 | 1.91E-06 |
| CCNF | -2.236604156 | 1.37E-11 |
| TBC1D24 | -0.850930163 | 4.73E-08 |
| AMDHD2 | 0.280679921 | 0.015388694 |
| BICDL2 | 2.166681999 | 0.000173394 |
| ZNF75A | 0.937779689 | 1.31E-07 |
| ADCY9 | -0.869995599 | 1.88E-07 |
| CLPB | -0.489065147 | 2.19E-05 |
| NEU3 | -1.295909797 | 2.88E-09 |
| PPP1R32 | 4.619274192 | 8.03E-08 |
| GNG3 | 1.514271933 | 0.001959 |
| UBXN1 | 0.318961009 | 0.0002127 |
| LBHD1 | 0.992357749 | 1.95E-09 |
| TTC9C | -0.393247425 | 0.001057745 |
| TAF6L | -0.580761943 | 0.0002957 |
| STX5 | 0.201950215 | 0.012632773 |
| RPL29 | 0.397176739 | 6.22E-07 |
| ITIH3 | 5.024664947 | 0.00018995 |
| SYVN1 | 0.621422768 | 1.69E-07 |
| ZFPL1 | 0.571936308 | 0.000414267 |
| RPS6KA4 | -0.453300749 | 0.000104488 |
| TPCN2 | -0.655298128 | 0.000362844 |
| CMPK1 | -0.213483139 | 0.000993802 |
| COA7 | -1.620463793 | 4.12E-11 |

| | | |
|---|---|---|
| ZYG11B | -0.378563276 | 3.36E-05 |
| CZIB | -0.194004705 | 0.032721229 |
| MAGOH | -0.461920966 | 0.000301755 |
| PARS2 | 1.751359598 | 6.47E-10 |
| LEXM | 5.66576828 | 4.35E-05 |
| USP24 | -0.781275156 | 8.45E-08 |
| NOL9 | -0.328056192 | 0.000196759 |
| PRKAA2 | -0.448262643 | 0.000164505 |
| KLHL21 | 0.215529439 | 0.000267279 |
| ZSWIM5 | -1.280192466 | 0.001377896 |
| GMEB1 | -1.668236977 | 1.77E-08 |
| SLC45A1 | 1.443799811 | 0.028995029 |
| SELENON | 0.12170646 | 0.041037823 |
| AK4 | -1.443804612 | 5.47E-10 |
| JAK1 | -0.617853974 | 1.36E-07 |
| RAVER2 | -0.71564177 | 1.01E-06 |
| LZIC | -0.992245785 | 7.02E-08 |
| KNCN | 9.135129598 | 1.02E-08 |
| SLC25A34 | 3.327447241 | 1.37E-08 |
| AKR7A3 | 1.646769091 | 0.018850448 |
| DHRS3 | 0.910331264 | 5.74E-06 |
| MATN1 | 1.838062306 | 4.77E-05 |
| LAPTM5 | 0.940057752 | 0.002077037 |
| PEF1 | -0.193430854 | 0.013069214 |
| SYNC | -0.249896554 | 0.008452158 |
| RBBP4 | -0.240130617 | 0.000123015 |
| KIAA1522 | 0.543562341 | 2.37E-05 |
| TSSK3 | 1.171089771 | 8.55E-06 |
| CAMK2N1 | -0.405021303 | 0.010715499 |
| SCNN1D | 1.044494326 | 0.022858501 |
| MXRA8 | 0.722683374 | 0.001999497 |
| FAAP20 | 0.252798868 | 0.003027341 |
| MEGF6 | 7.367957806 | 7.88E-06 |
| NFIA | -0.69385093 | 0.00268594 |
| MYSM1 | -0.834579283 | 2.40E-08 |
| USP1 | -1.023217633 | 7.34E-10 |
| FUBP1 | -0.96137611 | 9.80E-10 |
| NEXN | -2.192018617 | 1.58E-07 |
| DNAJB4 | 0.775737339 | 6.34E-06 |
| ADGRL4 | -0.474807672 | 0.014801164 |
| LRRIQ3 | 2.048005069 | 4.66E-07 |
| TYW3 | -1.164934777 | 5.61E-10 |
| LHX8 | -2.732390642 | 6.92E-05 |
| FAM102B | -0.840547021 | 1.06E-06 |
| HENMT1 | 0.844703403 | 0.000623617 |
| WDR63 | 3.762904063 | 0.007289158 |
| GBP2 | 0.690390551 | 0.002154044 |
| ATXN7L2 | -1.362647018 | 1.47E-05 |

| | | |
|---|---|---|
| ZNF326 | 0.168628923 | 0.031926382 |
| GFI1 | 0.798168166 | 0.007687642 |
| KCNT2 | 0.7816487 | 0.013717091 |
| AGL | -0.345025569 | 0.000154883 |
| EXTL2 | -0.788694411 | 5.34E-08 |
| SLC30A7 | -0.927950279 | 5.92E-09 |
| ZNF281 | -0.577654665 | 5.55E-07 |
| ARPC5 | -0.346678404 | 3.84E-06 |
| NLRP3 | 0.675947481 | 0.01302302 |
| ZNF496 | -0.736713265 | 4.76E-08 |
| TRIM58 | -1.648283982 | 3.29E-10 |
| KCNJ9 | 7.083771194 | 5.76E-06 |
| IGSF8 | 0.303734276 | 0.001031782 |
| DDR2 | 0.855269974 | 2.91E-09 |
| PEA15 | 0.119812096 | 0.008388158 |
| PEX19 | -0.244549075 | 0.002398569 |
| NCSTN | 0.160157275 | 0.024704923 |
| VANGL2 | 0.608981798 | 0.004128128 |
| SLC9C2 | 8.70279065 | 4.71E-08 |
| C1orf74 | 0.948530979 | 2.14E-07 |
| FLVCR1 | -1.570017404 | 1.80E-10 |
| ATF3 | 3.283549515 | 1.36E-12 |
| AXDND1 | 4.510573251 | 0.000718725 |
| IER5 | 1.48719629 | 3.72E-10 |
| BPNT1 | -0.455350712 | 1.96E-05 |
| SPATA17 | 1.205897153 | 0.018285349 |
| BROX | -0.325521879 | 4.11E-05 |
| NBPF20 | 0.282040261 | 0.002485663 |
| TFB2M | -0.784272769 | 5.53E-08 |
| KLHDC8A | 4.837574587 | 0.000505718 |
| PKDCC | -0.632678768 | 0.0349895 |
| B3GALNT2 | -0.236645034 | 0.017449553 |
| MAPKAPK2 | -0.303928393 | 0.000122116 |
| IL24 | -0.803623324 | 1.29E-06 |
| FCMR | -0.984075859 | 0.015123047 |
| CAPN2 | -0.219268528 | 0.000229868 |
| WDR26 | -0.596445986 | 6.92E-08 |
| REL | 0.241038244 | 0.00495469 |
| PUS10 | 0.679812166 | 0.001089993 |
| DISC1 | 0.425705544 | 0.018581107 |
| MEMO1 | -0.472962958 | 0.001951165 |
| DPY30 | -0.296536059 | 0.001120527 |
| MAIP1 | -0.735934538 | 8.96E-05 |
| ARL5A | -1.178028167 | 1.37E-09 |
| CLHC1 | 1.187591272 | 1.73E-05 |
| CFAP36 | 0.44893524 | 8.50E-05 |
| NUP35 | -0.547808043 | 0.000781691 |
| CCDC138 | -1.235519661 | 4.00E-06 |

| | | |
|---|---|---|
| SMC6 | -0.428458729 | 5.71E-05 |
| VSNL1 | -0.507104127 | 0.018681456 |
| CCDC74A | -0.454193128 | 0.041483184 |
| H3-3A | -0.274842726 | 0.000120231 |
| COQ8A | 0.372377857 | 4.27E-05 |
| EN1 | -0.979298114 | 0.003698983 |
| SGCB | -0.504845444 | 2.54E-06 |
| CCDC140 | 2.283984372 | 0.00058635 |
| SGPP2 | 1.978232725 | 2.69E-05 |
| SMARCAD1 | -1.456189159 | 4.13E-10 |
| PDLIM5 | -1.093422744 | 1.80E-09 |
| RPRD2 | -2.106451556 | 5.37E-12 |
| CTSS | 1.107654521 | 0.029968935 |
| PACRGL | -0.603256461 | 0.000112142 |
| BNIPL | 0.969754845 | 0.042376966 |
| TNFAIP8L2 | 2.19773752 | 0.003871737 |
| LYSMD1 | -1.888982518 | 1.15E-07 |
| SCNM1 | -0.340645671 | 0.000515849 |
| TMOD4 | 1.234064352 | 0.001169346 |
| ERCC3 | -0.487256636 | 7.26E-06 |
| RNF149 | -0.578128352 | 2.90E-05 |
| IWS1 | -0.802565763 | 1.37E-07 |
| BOLA3 | 0.300288073 | 0.024672149 |
| CDC42EP3 | -1.000168345 | 1.75E-09 |
| S100A11 | 0.177238418 | 0.008146685 |
| DHX57 | -0.887672787 | 3.32E-08 |
| SPRR2D | 0.669433685 | 0.011858003 |
| TGFA | -0.57932005 | 6.77E-09 |
| CCNYL1 | -0.383817601 | 7.43E-05 |
| FZD5 | -0.456289629 | 0.009308225 |
| DCAF16 | -0.895269408 | 5.20E-09 |
| C1orf189 | 7.95586464 | 7.42E-07 |
| GNPDA2 | -0.955325514 | 2.37E-08 |
| PAQR3 | -1.133451712 | 1.27E-09 |
| NIPAL1 | 0.770694068 | 9.65E-08 |
| ANTXR2 | -0.730503778 | 5.01E-05 |
| MRPS18C | 0.473643247 | 0.000335233 |
| CGGBP1 | 0.870889527 | 1.59E-09 |
| ABRAXAS1 | 0.682478956 | 1.47E-06 |
| GPR155 | 0.738424015 | 0.00957053 |
| PBXIP1 | 1.08054099 | 2.73E-08 |
| PYGO2 | -0.447781705 | 2.29E-06 |
| DCST2 | 3.183242043 | 2.66E-05 |
| DCST1 | 1.285205667 | 0.008735387 |
| COL6A3 | 0.585428905 | 1.72E-05 |
| INAVA | -1.353169 | 6.70E-07 |
| LINC01116 | -0.528363076 | 0.008518955 |
| YY1AP1 | -0.534767884 | 2.18E-06 |

| | | |
|---|---|---|
| KBTBD8 | 0.801381794 | 2.02E-06 |
| EOGT | -1.648419642 | 1.19E-09 |
| NAXE | 0.194646681 | 0.006598313 |
| POGLUT1 | -0.629854388 | 1.93E-06 |
| SLC22A15 | -1.665771468 | 1.39E-06 |
| ATP1A1 | 0.453765335 | 3.53E-07 |
| LRRC58 | -1.09338025 | 1.40E-10 |
| FSTL1 | -0.261157499 | 0.015877665 |
| LMOD1 | 4.309270923 | 1.27E-09 |
| ELF3 | 7.001050954 | 1.68E-05 |
| TMEM183A | -0.351680052 | 3.93E-05 |
| TMEM169 | 2.873121832 | 3.28E-09 |
| ARPC2 | 0.217405787 | 0.002686769 |
| TSACC | -2.072755165 | 9.47E-06 |
| TMEM79 | 0.319523956 | 0.003644757 |
| SSR2 | -0.276188081 | 3.02E-05 |
| ADORA1 | -0.796693316 | 0.000662328 |
| CIP2A | -0.557664879 | 4.68E-06 |
| EOMES | -2.358592564 | 0.001106308 |
| CWC22 | -0.334035019 | 0.000230292 |
| AZI2 | -0.738308519 | 6.64E-07 |
| TGFBR2 | -1.193442509 | 2.11E-09 |
| ANKZF1 | 0.791849381 | 1.61E-07 |
| HDAC11 | 1.443511447 | 1.32E-05 |
| GLB1L | 1.827507428 | 2.59E-07 |
| STT3B | -0.212574652 | 0.002220748 |
| CHCHD4 | -0.516078578 | 1.31E-05 |
| NFASC | 1.731048009 | 2.38E-06 |
| SGO2 | -1.807064247 | 3.76E-11 |
| SERPINI1 | 1.416033031 | 2.81E-05 |
| CLASP2 | -0.913577684 | 3.37E-08 |
| SUCLG1 | 0.20069989 | 0.010196759 |
| NUAK2 | 1.429913533 | 6.09E-07 |
| PRKCI | -0.564215805 | 1.39E-06 |
| AIM2 | 0.639491612 | 0.020677939 |
| EFHB | 3.7853013 | 5.81E-06 |
| RPL22L1 | 0.194456218 | 0.002455778 |
| PPM1L | -1.294806637 | 6.94E-07 |
| ICA1L | -1.135895866 | 5.46E-05 |
| SNHG16 | -1.212750078 | 5.17E-10 |
| RYBP | 0.377950673 | 4.56E-06 |
| PPP4R2 | -0.464454094 | 9.24E-07 |
| CD200R1 | 6.033618678 | 0.001737043 |
| GTPBP8 | -0.78724937 | 2.80E-06 |
| NEPRO | -1.088499508 | 2.73E-08 |
| SPICE1 | -0.601577401 | 9.57E-06 |
| CCDC191 | 2.013452178 | 0.000246854 |
| NKX6-1 | 0.682115528 | 0.000270737 |

| | | |
|---|---|---|
| CDS1 | -1.963171358 | 1.51E-06 |
| WDFY3 | -0.916283219 | 2.05E-07 |
| COX18 | 0.792087857 | 2.49E-05 |
| ALB | 8.330018574 | 6.01E-07 |
| ATXN7 | -0.395047899 | 9.70E-06 |
| PSMD6 | -0.103812207 | 0.04096838 |
| ADAMTS9 | -0.5419363 | 0.041791046 |
| PPM1K | -0.428900685 | 0.000294156 |
| GMPS | -0.481474733 | 2.54E-06 |
| TIPARP | -0.423289864 | 4.17E-06 |
| CCNL1 | 0.171371594 | 0.004163663 |
| PTX3 | 0.373811925 | 0.002251742 |
| HESX1 | 4.122887577 | 6.00E-08 |
| DCLK3 | 8.492473893 | 9.66E-08 |
| SLMAP | -1.552576349 | 1.30E-10 |
| RPL9 | -0.63074858 | 2.75E-07 |
| SMIM14 | 1.217256866 | 1.83E-07 |
| RPP14 | -1.21876336 | 2.34E-09 |
| ABHD6 | 0.324577734 | 0.004249556 |
| C3orf67 | 0.832928674 | 0.003221682 |
| RBM47 | -0.890119795 | 2.16E-05 |
| APBB2 | -0.974200135 | 1.19E-07 |
| IL17RE | 2.2006421 | 3.30E-05 |
| CRELD1 | 0.885797874 | 1.35E-06 |
| PRRT3 | 0.60111432 | 0.005448305 |
| PCOLCE2 | -0.420647547 | 0.032267515 |
| U2SURP | -0.592935753 | 1.44E-07 |
| MTMR14 | 0.210906456 | 0.010227461 |
| TTC14 | -1.135900255 | 3.13E-09 |
| CXCL3 | 1.289072606 | 0.00023047 |
| MTHFD2L | 0.637118621 | 2.18E-05 |
| CXCL1 | 0.812849419 | 4.54E-07 |
| RCHY1 | -0.73813914 | 8.68E-06 |
| GYG1 | 0.143608235 | 0.030775494 |
| TOPBP1 | -0.738894675 | 2.84E-08 |
| RYK | -0.96210478 | 3.53E-08 |
| UCN | 3.317365037 | 1.44E-06 |
| ZNF513 | 0.449718047 | 0.000687087 |
| SLC4A1AP | -1.236134427 | 5.80E-08 |
| PLB1 | 1.770980645 | 0.021839619 |
| SPDYA | -0.669831505 | 0.00013109 |
| KIAA1143 | 0.458751888 | 5.29E-06 |
| KIF15 | -1.199390793 | 1.17E-08 |
| TGM4 | 6.871264811 | 2.36E-05 |
| WDR43 | -1.660378316 | 3.55E-11 |
| ZDHHC3 | -0.582271045 | 6.31E-07 |
| SLC6A20 | 7.972389542 | 1.02E-06 |
| LZTFL1 | 0.365169642 | 0.000467275 |

| | | |
|---|---|---|
| ELP6 | -0.340121338 | 0.002759651 |
| DTX3L | -0.893370767 | 2.09E-08 |
| ZNF148 | -0.974375497 | 4.24E-09 |
| NMNAT3 | 3.188653301 | 0.000329676 |
| SMIM12 | -1.180859686 | 1.65E-07 |
| ZMYM6 | -0.36163147 | 0.001459174 |
| TPRA1 | -0.154856064 | 0.037032413 |
| YEATS2 | -0.435947909 | 3.76E-05 |
| MEAF6 | -0.90616385 | 2.77E-08 |
| SNIP1 | -0.199140172 | 0.006297133 |
| POLR2H | -0.186989135 | 0.009433504 |
| KLF15 | 2.38208906 | 2.25E-08 |
| TMEM41A | -0.921431671 | 3.66E-07 |
| SENP2 | -0.618463262 | 2.81E-06 |
| RHO | 7.526402291 | 2.96E-06 |
| RFC4 | -0.783692295 | 1.23E-06 |
| BAP1 | -0.53054923 | 6.15E-07 |
| TKT | 0.389270616 | 8.88E-07 |
| PRKCD | -0.29905607 | 0.001656956 |
| SFMBT1 | -1.45254652 | 1.47E-07 |
| GNL3 | -0.463299246 | 4.93E-07 |
| PBRM1 | -1.055452108 | 1.64E-09 |
| UVSSA | 0.511682982 | 8.62E-05 |
| TASOR | -0.582629121 | 9.31E-08 |
| SLBP | -0.4569009 | 1.69E-06 |
| SLC51A | 1.106488952 | 0.002054909 |
| UBXN7 | 0.617633174 | 2.18E-07 |
| RNF168 | -0.602760974 | 6.72E-06 |
| PIGX | -0.574653801 | 8.87E-05 |
| MELTF | 0.347260626 | 4.44E-05 |
| S100P | 2.147593434 | 0.00138689 |
| EXO5 | -0.746088476 | 1.97E-05 |
| CLDN19 | 5.119413226 | 2.50E-07 |
| ERMAP | 2.151241423 | 4.09E-09 |
| ZNF691 | 0.736684489 | 3.88E-06 |
| SGMS2 | -1.050880227 | 8.33E-07 |
| METAP1 | -0.592827124 | 2.99E-07 |
| DNAJB14 | -0.694148722 | 1.19E-06 |
| H2AZ1 | -0.32224172 | 7.18E-05 |
| SLC9B2 | -0.841775003 | 4.35E-05 |
| BDH2 | 0.73336674 | 0.006558653 |
| PGRMC2 | -0.140088131 | 0.044629903 |
| CDC25A | -1.90275018 | 3.14E-11 |
| ZNF589 | 0.187987765 | 0.011636203 |
| FBXW12 | 2.861354464 | 0.000355312 |
| CCDC51 | 0.377479814 | 0.000196414 |
| ATRIP | -0.534821198 | 9.67E-06 |
| SHISA5 | 0.30910396 | 0.000177044 |

| | | |
|---|---|---|
| SPRY1 | -0.6750294 | 0.000533154 |
| BSN | 1.208655791 | 4.60E-06 |
| APEH | 0.147966944 | 0.042386177 |
| HSPA4L | -0.95238801 | 1.35E-07 |
| MFSD8 | -0.778476323 | 9.19E-05 |
| MON1A | -1.008993251 | 3.57E-07 |
| MST1R | 1.432743235 | 0.001688958 |
| RAD54L2 | -0.814920471 | 2.36E-08 |
| GRM2 | 7.283897296 | 1.34E-11 |
| DUSP7 | -0.813630587 | 2.35E-08 |
| POC1A | -0.793559001 | 2.37E-05 |
| PPM1M | 1.056379248 | 1.07E-07 |
| WDR82 | -1.090720054 | 1.99E-10 |
| PITX2 | 0.600782878 | 0.000455664 |
| C4orf3 | -0.55987182 | 5.24E-05 |
| PRSS12 | 0.469953569 | 0.007157044 |
| HMGB2 | -1.460439738 | 7.33E-11 |
| SAP30 | -0.363558518 | 0.009047657 |
| HAND2 | -2.330499754 | 9.52E-05 |
| MAD2L1 | -0.756988741 | 2.21E-08 |
| MAP9 | -0.360020704 | 0.000560614 |
| CEP44 | -1.356219232 | 1.15E-07 |
| ASB5 | 7.886793085 | 8.59E-07 |
| TMEM144 | -0.561527029 | 0.010108751 |
| GASK1B | 0.553681922 | 0.035160848 |
| NAA15 | -1.316966497 | 1.26E-10 |
| ARFIP1 | -1.047097576 | 1.57E-08 |
| ICE1 | -0.789289689 | 4.27E-09 |
| ABCE1 | -0.686475066 | 1.71E-08 |
| OTUD4 | -1.145404569 | 5.89E-10 |
| LSM6 | -0.416121135 | 0.013782914 |
| PRMT9 | -0.626181626 | 0.000266436 |
| ITGA2 | 0.242238497 | 0.000473783 |
| MOCS2 | -0.179768665 | 0.007185034 |
| SLC45A2 | 5.443133498 | 1.77E-08 |
| TMEM161B | -0.768587432 | 1.14E-07 |
| LMBRD2 | 0.347979607 | 0.00330167 |
| NIPBL | -0.825667809 | 5.80E-08 |
| ADGRV1 | 0.257486842 | 0.030311314 |
| SLC25A46 | -0.609236109 | 2.14E-07 |
| STARD4 | 1.826225018 | 8.12E-09 |
| PGGT1B | -1.453870137 | 9.91E-09 |
| F2RL2 | -0.594375116 | 0.001143454 |
| CCDC112 | 0.489328817 | 0.001209687 |
| CMBL | 0.271420133 | 0.001918745 |
| C5orf63 | -0.566519284 | 0.040788358 |
| F2RL1 | -1.775913928 | 7.50E-07 |
| AGGF1 | -0.375613179 | 4.38E-05 |

| | | |
|---|---|---|
| WDR41 | -0.543783811 | 8.31E-06 |
| PRDM9 | 4.322375457 | 0.00127783 |
| NDUFS4 | 0.397353057 | 0.006759745 |
| ESM1 | -1.448002872 | 3.71E-09 |
| GRPEL2 | -0.851108845 | 8.89E-09 |
| CDC20B | 8.475184557 | 1.59E-07 |
| ARSK | 0.319110992 | 0.010822714 |
| RHOBTB3 | -0.611724531 | 1.02E-06 |
| GPX8 | -0.317212541 | 0.000117075 |
| TIGD6 | 0.985520106 | 5.07E-08 |
| SPZ1 | 8.416733866 | 1.29E-07 |
| SERINC5 | -1.266860042 | 1.71E-08 |
| CAGE1 | 5.010199165 | 2.76E-09 |
| CASP3 | -0.611227282 | 2.38E-07 |
| PRIMPOL | 0.653945308 | 2.61E-05 |
| EGFLAM | -1.448128805 | 2.10E-05 |
| CFAP97 | -0.543849513 | 1.91E-07 |
| RICTOR | -0.700937265 | 1.15E-07 |
| TENT2 | -0.766895769 | 3.77E-07 |
| ANKRA2 | 1.424036893 | 1.88E-06 |
| UTP15 | -0.916661736 | 1.96E-08 |
| TLR3 | -0.905736379 | 0.000255573 |
| KLKB1 | 4.758269838 | 0.000737859 |
| CCDC127 | 0.338031661 | 0.001335267 |
| SEPTIN8 | -0.320725032 | 0.000107583 |
| SHROOM1 | -1.496938957 | 9.88E-08 |
| GDF9 | 1.613332553 | 1.36E-05 |
| UQCRQ | 0.150792224 | 0.014680997 |
| LEAP2 | 2.632989662 | 4.17E-06 |
| SLC35A1 | -0.740240529 | 1.85E-06 |
| GRIK2 | 0.288513987 | 0.028649749 |
| CGAS | 4.751026888 | 0.00029854 |
| FABP7 | 1.166994238 | 0.000657402 |
| CITED2 | 2.92670137 | 9.00E-13 |
| CREBRF | 3.170811096 | 4.63E-12 |
| DCBLD1 | -0.43522148 | 0.001208714 |
| SFXN1 | -0.996408989 | 1.78E-09 |
| TMEM200A | -1.348924534 | 0.000122129 |
| PDSS2 | -0.798838493 | 2.35E-05 |
| STXBP5 | -1.153169878 | 1.62E-06 |
| IL31RA | -0.861377254 | 0.003170258 |
| ANKRD55 | -1.924631448 | 0.038398208 |
| DAGLB | 0.192212881 | 0.023020626 |
| STK17A | -0.168653653 | 0.00428527 |
| TRA2A | -1.043806106 | 1.10E-08 |
| FAM183BP | 7.791424046 | 1.33E-06 |
| GALNT10 | -0.822607566 | 2.27E-08 |
| SAP30L | -1.023402305 | 1.20E-07 |

| | | |
|---|---|---|
| RPS14 | 0.286726859 | 1.39E-05 |
| MYOZ3 | 9.306275351 | 6.29E-09 |
| COG5 | 0.826352091 | 1.17E-07 |
| BMT2 | 0.827539161 | 3.90E-07 |
| GPR85 | 1.305226238 | 9.08E-05 |
| SLU7 | 0.503966913 | 1.78E-06 |
| PTTG1 | -0.179029391 | 0.0116822 |
| CAMLG | 1.036836117 | 2.97E-09 |
| KCNK5 | -0.550698078 | 0.001653313 |
| KIF6 | 1.83023853 | 3.61E-06 |
| STEAP1 | -0.973677051 | 0.000627956 |
| CDCA7L | -0.674182873 | 1.28E-07 |
| MIOS | -1.489629735 | 4.55E-07 |
| USP49 | 1.771177488 | 3.62E-10 |
| SYTL3 | 3.604428735 | 0.000238394 |
| HEY1 | -0.941288915 | 0.000215551 |
| ZNF704 | -2.275233396 | 7.95E-07 |
| FABP5 | -0.310149533 | 0.019856298 |
| TAGAP | 7.501158202 | 5.33E-06 |
| CHMP4C | -0.711117421 | 0.046580284 |
| SLC13A4 | 1.322675611 | 0.000171218 |
| BRI3 | 0.533412413 | 2.77E-05 |
| LMTK2 | -1.123139285 | 2.34E-09 |
| CTSB | 0.192977662 | 0.002065775 |
| DLC1 | -0.573690863 | 8.96E-07 |
| ADCY1 | -0.963653333 | 0.000412405 |
| C8orf48 | 1.067305556 | 0.000272381 |
| HNF4G | -2.320702508 | 0.001090663 |
| PEX2 | 0.326428478 | 0.000199334 |
| RAD21 | -0.592878934 | 1.83E-08 |
| TNFRSF11B | -0.761517293 | 0.004211995 |
| PHKG1 | 0.73507623 | 0.006496945 |
| EN2 | 0.788083725 | 1.86E-05 |
| KCNV1 | -0.931983212 | 0.041114817 |
| CSMD3 | 1.577041838 | 0.002879967 |
| DNAAF5 | -0.43533341 | 2.83E-05 |
| OSGIN2 | -0.639171 | 5.59E-07 |
| SUN1 | -0.355503147 | 1.39E-05 |
| OXR1 | -0.664425676 | 8.72E-07 |
| TMEM74 | 0.615598306 | 0.004652617 |
| FAM86FP | 1.684122843 | 5.18E-05 |
| GPR146 | 2.410598426 | 1.16E-07 |
| TMEM184A | 1.28098862 | 4.61E-08 |
| MICALL2 | 0.651264993 | 2.85E-05 |
| CDK5 | 0.468955229 | 0.000818924 |
| SLC4A2 | 0.133448886 | 0.024075724 |
| FASTK | -0.308341446 | 0.000697026 |
| PHAX | -1.215262044 | 1.58E-09 |

| | | |
|---|---|---|
| FZD6 | -0.571073811 | 1.39E-06 |
| SLC25A32 | -0.843998974 | 1.68E-08 |
| DCAF13 | -0.120563285 | 0.035860978 |
| TP53INP1 | 2.135715514 | 1.71E-10 |
| INTS8 | -0.71704701 | 5.42E-08 |
| VIRMA | -0.204668779 | 0.001697812 |
| FREM1 | 0.563304239 | 0.006399059 |
| GEM | 4.99422532 | 1.53E-13 |
| WASHC5 | -0.222697291 | 0.014316967 |
| FAM219A | -0.390064569 | 0.000308842 |
| C9orf24 | 4.863964229 | 0.000332019 |
| SNAPC3 | -0.467472904 | 3.53E-06 |
| MYORG | 1.648287522 | 1.76E-08 |
| TMEM65 | -0.751139771 | 3.78E-05 |
| PSIP1 | -0.358955845 | 2.09E-05 |
| CCDC171 | 0.64128463 | 0.004730096 |
| UBAP1 | -0.483551412 | 0.000181199 |
| NIPSNAP3B | -1.267153137 | 0.011791961 |
| ABCA1 | 0.942461751 | 4.78E-05 |
| NFIL3 | 2.385669218 | 6.46E-12 |
| METTL2B | -0.875672277 | 1.39E-07 |
| FXN | -0.860975355 | 2.58E-05 |
| KDM1B | 0.195032252 | 0.020971899 |
| HGSNAT | -0.385398711 | 0.00135455 |
| KIF27 | 2.705268263 | 9.93E-10 |
| C9orf64 | -1.738216602 | 4.14E-07 |
| HNRNPK | 0.301677176 | 1.21E-06 |
| SVEP1 | 0.654516644 | 0.013130211 |
| ANKS6 | -0.507504365 | 2.20E-05 |
| PGAP4 | -1.472433512 | 1.82E-05 |
| ZHX1 | -0.285235154 | 0.000145088 |
| DYNLT3 | 0.338991431 | 7.90E-05 |
| METTL27 | 1.323471424 | 1.31E-05 |
| MID1IP1 | -0.499760327 | 0.000417561 |
| KIAA1958 | -0.380040031 | 0.000171762 |
| RNF183 | 9.453044727 | 3.58E-09 |
| PCDH19 | -1.549551623 | 0.003306665 |
| PIGA | 0.368196825 | 6.06E-05 |
| VEGFD | 2.36949852 | 0.001616009 |
| STRBP | -0.784434112 | 1.26E-06 |
| GAPVD1 | -1.066420766 | 1.06E-09 |
| ATP7A | -0.520816913 | 0.000258614 |
| ZNF367 | -0.768912833 | 2.02E-07 |
| NLGN4Y | -0.373576014 | 0.029690644 |
| HDX | 1.020256731 | 9.20E-05 |
| NDUFB6 | 0.293703031 | 0.000791239 |
| AQP7 | 9.301499215 | 1.45E-08 |
| NOL6 | -0.312012498 | 2.98E-05 |

| | | |
|---|---|---|
| AQP3 | 1.686208535 | 1.71E-08 |
| VCP | -0.353821663 | 6.13E-07 |
| STOML2 | 0.451177103 | 4.88E-07 |
| BRWD3 | -0.569511726 | 2.10E-06 |
| SLITRK5 | -1.538513081 | 2.03E-08 |
| MELK | -0.766137627 | 9.12E-07 |
| OTUD1 | 2.236700402 | 1.12E-08 |
| ARHGAP12 | -0.683674515 | 9.34E-08 |
| FAT3 | -0.339784012 | 0.002400648 |
| HECTD2 | -0.627652053 | 0.000160319 |
| FBXO33 | 0.411055588 | 6.13E-05 |
| INTS6L | -1.007137656 | 0.000252181 |
| CLDN2 | 3.541910086 | 0.009949222 |
| WRN | -0.736608404 | 4.25E-05 |
| MARCHF8 | -0.988787232 | 0.000731302 |
| CFL2 | -0.698177156 | 9.61E-08 |
| SUGT1 | -0.394128762 | 5.98E-05 |
| GTF2A1 | -1.253908406 | 5.40E-09 |
| ZCCHC24 | -0.293861626 | 0.000187249 |
| PGM2L1 | 0.56085693 | 0.000977594 |
| SLC16A9 | 1.323070841 | 0.018752341 |
| CRYL1 | 1.559110553 | 0.0001288 |
| REEP3 | -0.992930791 | 2.46E-08 |
| HEPACAM | 5.722881983 | 4.07E-05 |
| SKA3 | -1.361386092 | 1.31E-09 |
| DDIAS | -1.09538716 | 1.20E-07 |
| PCF11 | 0.915218819 | 2.99E-09 |
| PKNOX2 | 1.867970545 | 4.57E-07 |
| LRR1 | -0.615356947 | 5.20E-06 |
| RPL36AL | -0.299831756 | 0.000132122 |
| DNAAF2 | -0.661039624 | 5.97E-06 |
| C10orf25 | -1.086047951 | 0.045064391 |
| ZNF22 | -0.722995429 | 2.39E-07 |
| KLHDC2 | -0.484083139 | 1.03E-05 |
| EML5 | -1.332891508 | 1.13E-07 |
| NEMF | -0.683231952 | 5.67E-07 |
| RPUSD4 | -1.274862801 | 1.71E-09 |
| ARF6 | -0.170482364 | 0.004959532 |
| TTC8 | 0.47613318 | 0.001735288 |
| NOXRED1 | 2.628772305 | 3.99E-06 |
| KBTBD6 | -1.303901365 | 2.32E-09 |
| NUDT5 | -0.17758019 | 0.017545375 |
| OXGR1 | -1.535476935 | 0.001027312 |
| BEND7 | -0.666972948 | 0.026419902 |
| ATP5F1C | 0.230038195 | 0.00217455 |
| PRPF18 | -0.512618696 | 0.001287782 |
| TAF3 | -0.455997235 | 2.21E-05 |
| VDAC2 | 0.296607672 | 0.000813119 |

| | | |
|---|---|---|
| SLC18A2 | -0.984845437 | 1.12E-06 |
| PDZD8 | -0.484659096 | 4.69E-07 |
| ABRAXAS2 | -0.292865436 | 0.000278912 |
| QSOX2 | -0.197407412 | 0.001922792 |
| FAM204A | -0.564157569 | 1.37E-05 |
| NSD1 | -1.322483248 | 8.33E-11 |
| ENOX2 | -0.677491098 | 4.43E-05 |
| GHITM | 0.147284569 | 0.008770364 |
| SNAPC4 | -0.297500221 | 0.004693921 |
| PMPCA | -0.639642516 | 1.78E-07 |
| SPACA9 | 1.174046365 | 0.0001442 |
| TSC1 | -0.794638939 | 2.48E-07 |
| BORCS5 | -1.075581448 | 2.35E-05 |
| DIPK1B | 2.383951245 | 1.16E-09 |
| ZMYND19 | -0.255821688 | 0.009716717 |
| RET | 5.201186656 | 0.000127761 |
| DDX21 | -0.389302853 | 1.18E-07 |
| BMS1 | -0.433172127 | 1.43E-05 |
| STK32C | 0.514205641 | 0.005146444 |
| PIP4P1 | 0.758303668 | 1.46E-05 |
| METTL17 | -0.243705589 | 0.001608376 |
| ARHGEF40 | 0.180008113 | 0.043780243 |
| ZNF219 | 0.632534355 | 0.000822925 |
| CASP7 | -1.277794356 | 4.19E-09 |
| METTL3 | -0.540230605 | 1.58E-06 |
| SALL2 | 0.598294099 | 0.002484244 |
| TRUB1 | -0.159083802 | 0.044325918 |
| ERICH6B | 2.402487192 | 4.28E-05 |
| ZFYVE1 | 0.760759799 | 4.75E-06 |
| HSPA12A | 0.173763877 | 0.030830739 |
| SHLD2P1 | -0.983979403 | 1.63E-06 |
| FRAT1 | 1.573363713 | 1.22E-06 |
| ANKRD2 | 2.430472027 | 0.00471942 |
| E2F7 | -1.073307896 | 1.57E-09 |
| ARHGAP42 | -2.256282766 | 6.37E-06 |
| PACSIN3 | 0.260104971 | 0.001453795 |
| SLC39A13 | -0.771443816 | 3.85E-07 |
| AGBL2 | 1.681321287 | 0.00114266 |
| CPSF2 | -0.653067239 | 4.79E-07 |
| SMCO2 | 6.772642866 | 3.47E-05 |
| MOAP1 | 0.701268342 | 9.89E-07 |
| IFI27L1 | 0.990003036 | 0.004611954 |
| IFI27 | 3.651328756 | 1.74E-06 |
| CLMN | -0.485487338 | 7.83E-05 |
| CCDC38 | 4.440734827 | 2.37E-06 |
| PTER | 1.281544463 | 9.01E-08 |
| CACNB2 | 0.987661369 | 0.002450346 |
| ARL5B | 0.395380214 | 1.28E-05 |

| | | |
|---|---|---|
| SMCO4 | -0.789844496 | 0.005683797 |
| CEP295 | -0.64258031 | 1.24E-06 |
| TAF1D | -1.111411715 | 3.83E-10 |
| AMOTL1 | -1.330152797 | 1.89E-11 |
| HTRA1 | 0.307855469 | 0.020418878 |
| CEP57 | 0.218729014 | 0.001798991 |
| TCP11L2 | 2.966316051 | 4.61E-08 |
| SPRED1 | -1.932307231 | 8.77E-12 |
| GPR176 | -0.85989759 | 4.31E-08 |
| AC126323.1 | 0.514083603 | 2.82E-05 |
| GPT2 | -0.493064962 | 1.01E-05 |
| RPUSD2 | 0.668243317 | 1.60E-06 |
| HIF1AN | -0.180889271 | 0.007050292 |
| ZFYVE19 | 0.214502721 | 0.025631943 |
| FBN1 | -0.240627449 | 0.005881597 |
| BRD7 | -0.593737501 | 6.35E-07 |
| CKB | 0.606690104 | 2.17E-07 |
| TRMT61A | -0.923275578 | 4.33E-07 |
| BTRC | -1.510948775 | 1.19E-09 |
| BAG5 | -0.207360546 | 0.001759563 |
| LARP6 | 1.432188308 | 1.32E-08 |
| API5 | -0.765837007 | 2.28E-08 |
| ZNF319 | 0.515853352 | 0.000341359 |
| HPS6 | -0.504777185 | 5.66E-05 |
| SENP8 | 2.323311421 | 2.97E-09 |
| NOLC1 | -0.465272409 | 6.54E-08 |
| ALKBH3 | -0.714494028 | 0.000107821 |
| COPS2 | -0.438236179 | 1.15E-06 |
| GABRB3 | 0.714741596 | 0.000202823 |
| SGPL1 | -1.230159356 | 2.03E-08 |
| PCBD1 | 0.316791113 | 0.019160949 |
| ARIH1 | -0.782238898 | 1.75E-08 |
| C16orf71 | 2.635633409 | 1.26E-07 |
| SCN3B | 4.278133095 | 2.58E-10 |
| COX11 | -0.611497286 | 6.32E-07 |
| ZNF202 | -0.360203304 | 0.0004223 |
| STXBP4 | -1.989460292 | 4.29E-06 |
| CUL5 | -1.607056236 | 8.37E-10 |
| WBP1L | 0.485415749 | 1.11E-05 |
| BORCS7 | -0.329691866 | 0.004334829 |
| PLEKHF1 | 1.138226813 | 7.72E-06 |
| ANAPC16 | 0.260854258 | 0.001605354 |
| SMPD1 | 0.372526621 | 0.001740422 |
| APBB1 | 0.303329315 | 0.019540335 |
| SYNPO2L | 5.280315199 | 9.65E-05 |
| C11orf65 | 2.38140421 | 3.41E-05 |
| TRIM44 | -0.237850719 | 4.84E-05 |
| ILK | 0.159239883 | 0.027405843 |

| | | |
|---|---|---|
| TAF10 | 0.230354552 | 0.000742829 |
| TPP1 | 0.337011728 | 0.000366084 |
| DCHS1 | 0.519277253 | 0.013006193 |
| USP54 | -0.525400922 | 8.12E-05 |
| C11orf74 | 1.151627471 | 9.14E-05 |
| WDR88 | 2.441561362 | 2.95E-06 |
| CYB5R2 | 0.394154966 | 0.038757589 |
| SERPINB8 | -0.705104889 | 1.12E-06 |
| TUB | 0.262640735 | 0.008312855 |
| IDH3A | -0.402293527 | 7.31E-06 |
| WDR72 | 0.607310575 | 5.52E-06 |
| PLD4 | 1.622335537 | 6.42E-05 |
| ZMAT1 | 4.421359505 | 4.63E-06 |
| XRRA1 | -0.987028083 | 5.45E-08 |
| RNF169 | -0.977534859 | 1.10E-08 |
| RPL27A | 0.333165116 | 0.001096683 |
| DENND2B | -0.346668072 | 0.010878436 |
| CDYL2 | -1.592356292 | 7.49E-10 |
| CENPN | -0.256353521 | 0.000589646 |
| ATMIN | -0.403688897 | 5.19E-06 |
| C16orf46 | 1.934680186 | 4.04E-07 |
| PKD1L2 | 6.213589514 | 5.18E-10 |
| LEO1 | -0.804997372 | 1.43E-07 |
| ZNF143 | -1.070639471 | 2.31E-05 |
| TMX3 | -0.168292681 | 0.006919595 |
| WEE1 | -1.454505002 | 6.21E-10 |
| HDGFL3 | -0.705769078 | 1.55E-07 |
| NDST2 | 0.289228015 | 0.026403162 |
| MCM7 | -0.102915453 | 0.012542209 |
| ZNF3 | -0.269494682 | 0.000715287 |
| ZSCAN21 | 0.541831143 | 0.003346738 |
| RIMKLB | 1.801258894 | 1.13E-12 |
| TK2 | -1.52442789 | 4.32E-07 |
| SEC11C | 0.292904273 | 0.03113672 |
| CPLX4 | 6.912180263 | 2.73E-05 |
| TMEM135 | -0.343765241 | 0.001748413 |
| IQCD | 1.579720962 | 0.000279725 |
| NDEL1 | -0.172038563 | 0.013615579 |
| CENPV | 0.505549239 | 0.045150451 |
| RRAD | 3.618770579 | 8.56E-05 |
| CIAO2B | 0.313055707 | 0.000461424 |
| CFAP52 | 2.576877468 | 0.000327654 |
| HSP90B1 | 0.575226437 | 1.02E-08 |
| ATF7IP2 | 0.791139619 | 1.67E-06 |
| MMP10 | 2.826466505 | 0.000365317 |
| TVP23A | -0.381099289 | 0.026732456 |
| BEX3 | 0.374559241 | 3.72E-05 |
| TMPRSS5 | 0.5411834 | 0.00547806 |

| | | |
|---|---|---|
| COG1 | -0.563688048 | 2.39E-06 |
| PLEKHA7 | -1.029897763 | 0.000862438 |
| ZNF606 | 1.814409998 | 3.07E-09 |
| B2M | 0.280248967 | 4.20E-05 |
| ZNF592 | -0.563328489 | 2.22E-06 |
| CASC4 | -0.353217676 | 3.45E-05 |
| NNMT | -1.257812263 | 1.01E-08 |
| AP1G1 | -1.017691298 | 1.25E-09 |
| SLFN5 | 3.236830088 | 7.50E-14 |
| STRCP1 | 7.12616806 | 1.22E-05 |
| YPEL4 | 2.808558889 | 1.60E-07 |
| PPIB | 0.337769241 | 0.000979139 |
| LDHC | 0.956266141 | 0.008057156 |
| CIAO2A | -0.216185225 | 0.004546924 |
| FAM111A | -0.988877286 | 1.69E-09 |
| KIF7 | -0.45308218 | 0.000494141 |
| TMEM170A | 0.320429477 | 0.000232218 |
| ANPEP | 0.213792464 | 0.00111042 |
| RBPMS2 | -1.208165129 | 4.70E-05 |
| NAV2 | -0.567951139 | 0.017746138 |
| ANKDD1A | 0.865269691 | 5.81E-05 |
| C18orf54 | -1.312122715 | 8.71E-09 |
| DCTN5 | -0.665952394 | 2.04E-08 |
| PLK1 | -1.285514866 | 4.93E-10 |
| CLPX | -0.632299332 | 8.62E-07 |
| ZBTB39 | -0.932493481 | 3.13E-08 |
| TAC3 | 7.421210236 | 4.54E-06 |
| MYO1A | 10.87972534 | 7.50E-11 |
| NEMP1 | 0.327155421 | 1.75E-05 |
| STAT6 | 0.445976045 | 3.61E-05 |
| PATL1 | -1.07761531 | 1.25E-09 |
| STX3 | -1.169104135 | 4.85E-09 |
| MRPL16 | -0.494331403 | 4.24E-06 |
| PIP4K2C | -0.432108276 | 1.02E-05 |
| YWHAB | -0.645520256 | 4.14E-09 |
| TSC22D4 | -0.441518522 | 8.34E-05 |
| DIS3L | -1.143186971 | 9.95E-08 |
| CCNDBP1 | 0.593322012 | 6.34E-06 |
| EPB42 | 4.030965848 | 9.08E-05 |
| SMAD3 | -0.971208156 | 1.78E-09 |
| MAP1A | 0.797302373 | 1.46E-07 |
| MAPRE2 | -0.25755092 | 0.000558234 |
| EVA1C | -0.696296593 | 0.02840827 |
| MARS1 | -0.18429478 | 0.000791867 |
| MBD6 | -0.448243205 | 5.96E-06 |
| CNPY4 | 0.435993468 | 0.003966314 |
| PDIA3 | 0.359981905 | 7.01E-06 |
| NUDT21 | -0.416023208 | 5.10E-07 |

| | | |
|---|---|---|
| NAT16 | 2.042516241 | 0.00147289 |
| NKX3-1 | 0.628342157 | 6.05E-06 |
| AL357033.1 | 2.433612711 | 0.001890066 |
| TEF | 1.272163922 | 1.55E-06 |
| MEI1 | 4.825812912 | 0.000233442 |
| PBX3 | -1.418464636 | 2.35E-08 |
| PHB | 0.181390087 | 0.003166217 |
| SNRPD1 | -0.423977313 | 1.79E-05 |
| TTC16 | 4.33393062 | 0.002637243 |
| SAMD14 | 1.087464713 | 7.60E-06 |
| PIP5KL1 | 2.223716934 | 6.12E-06 |
| FAM102A | -0.879755723 | 1.63E-06 |
| ACSF2 | 1.345529892 | 1.95E-05 |
| GOLGA2 | 0.725766587 | 3.77E-08 |
| TRUB2 | -0.488272111 | 1.30E-05 |
| SLC27A4 | -0.401613713 | 3.27E-05 |
| CERCAM | 0.265098544 | 0.008609758 |
| DOLPP1 | -0.436944481 | 4.97E-05 |
| CCDC103 | 1.594654443 | 5.82E-06 |
| ENDOG | -0.730424262 | 0.000249722 |
| ISLR2 | 3.435428924 | 0.015871179 |
| SP2 | -0.51907838 | 1.43E-05 |
| PRR15L | 2.194775566 | 0.003656052 |
| COQ7 | -0.47696958 | 0.000213957 |
| GOLGA6C | 4.23933302 | 0.003293582 |
| FBXO22 | -0.437339238 | 4.52E-06 |
| TBC1D2B | -0.978785632 | 1.00E-08 |
| KATNAL2 | 3.352132531 | 1.31E-08 |
| HDHD2 | 0.541689215 | 0.000102253 |
| ZNF91 | 0.556970218 | 3.74E-07 |
| RNF214 | 0.417866447 | 5.31E-05 |
| CDK12 | -2.306426705 | 1.36E-12 |
| DUS2 | -0.763933564 | 0.000325168 |
| ENGASE | 0.624390206 | 1.44E-05 |
| RBFOX3 | 2.013070853 | 0.000376888 |
| ATP5MG | 0.256917543 | 0.005103626 |
| TBC1D16 | -0.511167548 | 5.84E-07 |
| MYO5B | 1.94134394 | 4.37E-06 |
| ACAA2 | 0.444816695 | 0.003820703 |
| RRM1 | -0.295674812 | 0.000117136 |
| OR51B5 | 0.661071371 | 0.000663275 |
| FN3K | 2.569416961 | 3.64E-05 |
| PRRT2 | 1.057996812 | 0.000312732 |
| ZNF23 | 1.079900185 | 2.81E-08 |
| IRGQ | -0.756657115 | 3.61E-07 |
| ZNF226 | -0.788455684 | 1.79E-06 |
| ZNF180 | -0.37892063 | 0.000221919 |
| PPP2R3B | 1.069773709 | 0.003754214 |

| | | |
|---|---|---|
| ZNF668 | -1.334078487 | 2.70E-07 |
| ZNF646 | 0.961976538 | 4.58E-09 |
| SMG8 | -0.479244417 | 3.43E-06 |
| RAB8A | -0.613545243 | 2.14E-07 |
| GPX4 | 0.48310761 | 4.37E-06 |
| MIDN | 2.030895932 | 3.86E-12 |
| JSRP1 | 0.432929032 | 0.008833229 |
| KLHL26 | 1.104320848 | 1.44E-08 |
| GATAD2A | -0.481741909 | 3.91E-06 |
| MVD | 1.00436483 | 1.31E-06 |
| CDT1 | -0.388799013 | 6.41E-05 |
| TRAPPC2L | -0.361503081 | 0.003570894 |
| ANKRD11 | -0.678419219 | 2.26E-07 |
| SPATA33 | 0.723732475 | 8.28E-05 |
| RSKR | 0.834995033 | 1.40E-06 |
| PROCA1 | 2.235261603 | 1.88E-06 |
| RPL13 | 0.514448039 | 0.006589034 |
| ZNF641 | 0.605672823 | 0.00031354 |
| DHRS13 | -0.308480701 | 0.019350421 |
| KMT2D | -0.990383503 | 5.81E-10 |
| CORO6 | 2.54221973 | 0.004991281 |
| RHEBL1 | 0.673466683 | 0.001292813 |
| TUBA1A | -0.256757467 | 0.000244236 |
| TUBA1C | -0.893410328 | 1.05E-09 |
| ZNF610 | 5.464342163 | 1.93E-09 |
| ZNF528 | 0.458328153 | 0.001747724 |
| ZNF701 | 1.820688448 | 9.73E-09 |
| SERTAD3 | 0.930342764 | 4.37E-06 |
| NCKAP5L | 0.723631677 | 6.20E-08 |
| RAB4B | -0.518876131 | 0.000849513 |
| GPD1 | 2.122897146 | 7.46E-06 |
| PROSER3 | -0.456720418 | 0.000681396 |
| CYP2S1 | 1.408834005 | 0.000186076 |
| AXL | -0.121889149 | 0.009004333 |
| ZNF526 | 0.703093575 | 0.000528267 |
| TRAPPC9 | 0.35994525 | 0.027587342 |
| ZNF146 | -0.179565039 | 0.000631272 |
| ZNF283 | -0.747896119 | 1.73E-06 |
| C19orf33 | 0.982245011 | 0.000835708 |
| YIF1B | 0.417709791 | 0.000120907 |
| DNAAF3 | 1.327553209 | 2.21E-05 |
| PSCA | 1.178171491 | 0.015017051 |
| DAPK3 | -0.67864763 | 9.86E-06 |
| EEF2 | 0.524855359 | 8.34E-10 |
| CHAF1A | -1.126594201 | 2.20E-09 |
| UBXN6 | 0.18335852 | 0.034451786 |
| HDGFL2 | 0.255591397 | 0.003206194 |
| PLIN4 | 1.167751671 | 0.007991107 |

| | | |
|---|---|---|
| SEMA6B | 0.411431435 | 0.002264369 |
| NXN | -0.819957116 | 0.001137613 |
| TLCD3A | -1.959291904 | 1.66E-10 |
| GLOD4 | -0.425125676 | 3.54E-05 |
| MFSD3 | 0.43141399 | 0.00443728 |
| KIFC2 | 0.916248118 | 3.18E-05 |
| SRR | -0.441282127 | 3.31E-06 |
| TSR1 | -0.535840414 | 1.01E-07 |
| TRPV3 | 5.658598145 | 1.44E-05 |
| HSD11B1L | 1.174182686 | 9.21E-06 |
| CYB5D2 | 1.110667766 | 2.92E-06 |
| C19orf48 | -0.192656165 | 0.002080848 |
| ZNF83 | 0.286972299 | 0.000850714 |
| KRT80 | -0.362811481 | 3.11E-05 |
| OTUB1 | -0.207647247 | 0.002733991 |
| AC010323.1 | 0.377141949 | 0.031250861 |
| CD320 | 0.198897158 | 0.014890982 |
| SPRYD3 | -1.283252539 | 6.68E-09 |
| IGFBP6 | 0.630812787 | 3.01E-06 |
| ZNF558 | 1.156208428 | 2.17E-08 |
| NDUFV1 | 0.222258772 | 0.005752183 |
| CDK2AP2 | 0.887984609 | 3.05E-07 |
| NUDT8 | -0.950404266 | 0.000110499 |
| TBX10 | -1.395643563 | 4.86E-06 |
| AC011511.1 | 0.262713328 | 0.028167626 |
| PRDX2 | 0.415432926 | 1.37E-05 |
| MIS12 | 0.734026408 | 1.76E-08 |
| MRPL58 | 0.180274631 | 0.027322514 |
| ATP5PD | 0.508994397 | 2.30E-05 |
| TMEM88 | 4.602089282 | 1.59E-08 |
| SRP68 | -0.328117224 | 0.000482934 |
| MGAT5B | -0.610634753 | 0.000495048 |
| TMEM68 | -1.011706001 | 1.40E-07 |
| GHDC | 0.692805175 | 3.03E-05 |
| FAM234A | 0.265235864 | 0.000568858 |
| SOST | 8.098607371 | 4.02E-07 |
| ZNF598 | -0.302038361 | 0.000156232 |
| MLST8 | 0.169654692 | 0.022750724 |
| E4F1 | 0.197519095 | 0.013309467 |
| DNASE1L2 | 1.217175843 | 0.000423284 |
| ABCA3 | 2.36678877 | 1.17E-05 |
| KCTD5 | -1.099283833 | 2.44E-09 |
| SRRM2 | -0.759755787 | 3.48E-10 |
| ZNF597 | 0.85376178 | 8.93E-05 |
| NLRC3 | 3.33065731 | 8.46E-07 |
| SDHAF2 | -1.064815967 | 1.47E-08 |
| DDB1 | -0.132415531 | 0.008782528 |
| VPS37C | -1.27809687 | 1.03E-08 |

| | | |
|---|---|---|
| VWCE | 3.006869198 | 1.76E-08 |
| BEST1 | 0.286990321 | 2.94E-05 |
| FTH1 | 0.224986518 | 1.17E-05 |
| BSCL2 | 0.837528552 | 1.44E-07 |
| POLR2G | 0.576314638 | 7.00E-06 |
| SLC3A2 | -0.637803071 | 7.63E-09 |
| SPINDOC | -0.486871853 | 2.39E-06 |
| ATG16L2 | 0.375483645 | 0.009874825 |
| C2CD3 | -0.424613949 | 0.000113815 |
| TRANK1 | 1.886415744 | 1.53E-10 |
| TTC21A | 2.160783135 | 4.29E-07 |
| RPSA | 0.119358263 | 0.008262486 |
| CTNNB1 | 0.217085936 | 3.53E-05 |
| ULK4 | 0.639021326 | 0.001421046 |
| FADD | -0.685526436 | 5.13E-07 |
| NAALADL1 | 4.247355514 | 4.81E-07 |
| SAC3D1 | -0.684986781 | 5.37E-05 |
| BATF2 | 3.127274098 | 5.14E-09 |
| SF1 | -0.397509847 | 5.16E-07 |
| MAP4K2 | 0.312540559 | 0.004367563 |
| MAJIN | 7.526402291 | 2.96E-06 |
| CCDC88B | 1.517337705 | 0.004436337 |
| PBK | -0.467311817 | 5.11E-06 |
| SCARA5 | 1.148242135 | 1.31E-05 |
| COPS6 | 0.259482112 | 0.000268534 |
| PAFAH1B2 | -0.914802444 | 1.41E-09 |
| NUDT16L1 | 0.353576552 | 0.000684905 |
| KIAA1586 | -2.700107453 | 4.41E-11 |
| RAB4A | -0.373109801 | 0.00111042 |
| OR1F1 | 2.759505775 | 0.000712046 |
| SETD5 | -0.723220128 | 2.26E-08 |
| VASN | 0.764548089 | 0.000303733 |
| FAM83B | -1.125075794 | 0.004573165 |
| THAP9 | 2.178479437 | 1.83E-09 |
| RNF187 | -0.133484809 | 0.016169206 |
| HOOK3 | -1.527639111 | 1.90E-09 |
| MAPK1IP1L | -0.322213486 | 2.17E-05 |
| DDIT4 | 0.651427082 | 2.30E-08 |
| RBPJ | -1.066132659 | 2.60E-10 |
| ZCCHC4 | -1.34075137 | 1.16E-06 |
| TTC39C | -0.670478206 | 0.000201294 |
| GLYCTK | -0.480153873 | 0.015091384 |
| GNG4 | 0.246265113 | 0.001199675 |
| UBTD2 | 0.140385945 | 0.022466327 |
| IRF2BP2 | -1.601871333 | 1.28E-10 |
| NT5DC2 | -0.626781717 | 3.49E-07 |
| SMIM4 | 0.285892539 | 0.046969743 |
| COA6 | -0.703152616 | 3.03E-06 |

| | | |
|---|---|---|
| KIF5C | -1.200077552 | 4.98E-07 |
| MGAT2 | -0.565125986 | 8.27E-07 |
| BMI1 | -0.854701707 | 7.27E-09 |
| THAP11 | -0.541094711 | 1.96E-06 |
| MMADHC | -0.138814293 | 0.029199791 |
| PDHB | 0.215763313 | 0.001041165 |
| PXK | -0.441314372 | 0.00022896 |
| H1-4 | 3.816342575 | 1.07E-07 |
| PCMTD1 | -0.193441509 | 0.039681122 |
| MPLKIP | 0.26665706 | 0.00171221 |
| IRF2 | 0.384512141 | 0.002932877 |
| MOBP | 9.645928915 | 4.23E-05 |
| CX3CR1 | 11.32932163 | 2.42E-11 |
| XIRP1 | 4.34688236 | 1.63E-06 |
| SCN11A | 6.493084349 | 9.38E-06 |
| ARF4 | -0.845367294 | 1.41E-08 |
| HLA-DPA1 | 0.184597862 | 0.017200551 |
| SEPTIN2 | -0.249376577 | 4.89E-05 |
| DTYMK | -0.182184517 | 0.0272995 |
| ING5 | -0.467622831 | 4.43E-05 |
| MLKL | -1.674547459 | 3.33E-09 |
| RFWD3 | -0.873376973 | 2.91E-09 |
| COG7 | -1.051256295 | 1.78E-05 |
| CDC40 | -1.158914081 | 4.24E-08 |
| STIP1 | -0.103077267 | 0.015141594 |
| HR | -1.555315579 | 5.17E-11 |
| RAB31 | -0.947406517 | 1.77E-06 |
| REEP4 | -0.210794603 | 0.049291847 |
| TNXB | 2.906251018 | 1.76E-05 |
| ATXN2L | -0.19781512 | 0.001374117 |
| PHYHIP | 0.904556246 | 0.008918442 |
| CCDC110 | 3.697326228 | 1.34E-06 |
| FEN1 | -1.453460247 | 2.39E-11 |
| MTCL1 | -1.3504149 | 1.60E-07 |
| HEXIM2 | 1.273331218 | 8.33E-06 |
| FNTA | -0.471776243 | 9.33E-06 |
| SERINC2 | 0.579565302 | 0.000481399 |
| TRAPPC11 | -0.160759964 | 0.020155859 |
| ING2 | 0.397780978 | 0.00016385 |
| CDKN2AIP | 0.450978266 | 9.83E-06 |
| SNRNP48 | -0.944624102 | 8.80E-08 |
| SLC20A2 | -0.574747216 | 3.02E-06 |
| DYNLRB2 | 4.166221116 | 0.001859812 |
| TMUB2 | -0.477452575 | 9.90E-05 |
| ADAM9 | -0.16388086 | 0.021590907 |
| GDNF | -2.437797295 | 5.26E-07 |
| AXIN2 | 1.675442751 | 0.010279623 |
| ZNF30 | 0.693116611 | 1.58E-05 |

| | | |
|---|---|---|
| LRATD2 | -1.69192081 | 3.57E-12 |
| LDLRAD4 | 1.619683093 | 0.004788331 |
| KCTD19 | 1.574063861 | 0.041856484 |
| SLC16A4 | 0.403758639 | 0.007920786 |
| AHCYL1 | 0.186298345 | 0.000829303 |
| DNAJC21 | -0.189637118 | 0.004324831 |
| NPNT | 7.792517616 | 1.21E-06 |
| SEMA4C | -0.910865316 | 1.47E-08 |
| CNNM3 | -0.330705288 | 0.005953396 |
| TET2 | -1.242901892 | 6.85E-09 |
| TCTN2 | -0.519125373 | 0.007098802 |
| SHOX2 | -0.573030517 | 0.000250348 |
| PPIP5K1 | -0.490520163 | 0.000721387 |
| TSPAN5 | -0.400919252 | 0.001123918 |
| ZBTB5 | 0.43742195 | 2.88E-05 |
| CHTF8 | -0.588762411 | 7.37E-07 |
| ADAL | 0.288322755 | 0.011614884 |
| LCMT2 | 0.248293673 | 0.000848252 |
| SNTB2 | -0.877059849 | 1.19E-08 |
| ZNF507 | -1.875850178 | 9.37E-11 |
| STX18 | -0.359434561 | 0.006295571 |
| ZBTB49 | 1.238604046 | 1.25E-07 |
| GFM1 | -0.494028605 | 1.61E-06 |
| DDX19A | -0.563385699 | 6.68E-07 |
| ATOH8 | -1.069398417 | 0.002647338 |
| TNIP2 | -0.848008671 | 1.97E-05 |
| C2orf68 | 0.452811694 | 1.58E-05 |
| RNF181 | 0.53902155 | 2.83E-05 |
| VAMP5 | 0.356509827 | 0.016580873 |
| MAT2A | -1.008757905 | 6.24E-11 |
| ZNF608 | -0.59143853 | 0.00077462 |
| LETM1 | -0.970037806 | 5.23E-09 |
| TMEM129 | 0.58424798 | 1.27E-05 |
| PPIC | 0.345537088 | 0.00531558 |
| SPRY3 | -0.723556622 | 0.010347452 |
| CEP120 | 0.21558928 | 0.008425862 |
| STXBP6 | -1.514421879 | 0.007451157 |
| MFF | -0.552673005 | 9.20E-06 |
| LGALS9 | 2.9602268 | 9.46E-05 |
| PXDC1 | -0.68704717 | 1.47E-05 |
| E2F6 | -0.949335848 | 5.20E-07 |
| COMMD8 | -0.314420865 | 0.004122546 |
| UQCRFS1 | -0.447337452 | 4.36E-06 |
| COL4A3 | 1.46572763 | 0.030675807 |
| MAP2K1 | -0.49716965 | 1.22E-06 |
| IRS1 | -0.433796973 | 8.33E-05 |
| MECP2 | -0.13494775 | 0.016681298 |
| AR | -0.838456666 | 0.000172792 |

| | | |
|---|---|---|
| VXN | 2.314005762 | 3.88E-06 |
| SLC25A6 | 0.553599601 | 4.90E-07 |
| CHST14 | -0.732194677 | 1.72E-07 |
| PARM1 | -0.609412192 | 0.01618917 |
| CSNK1G1 | -0.495831116 | 3.06E-06 |
| ZNF354A | 0.778619417 | 4.12E-07 |
| ATF5 | -0.814061179 | 1.88E-07 |
| UBE2V2 | -0.57768758 | 1.75E-07 |
| ZBTB43 | 0.758087427 | 3.81E-08 |
| CPT1C | 3.837603185 | 3.18E-10 |
| XPO6 | -0.894586327 | 1.22E-08 |
| MN1 | 5.183240116 | 8.89E-09 |
| APEX2 | -0.924998941 | 1.42E-08 |
| NSMCE1 | 0.832507747 | 1.78E-05 |
| IL13 | 7.404211953 | 4.43E-06 |
| RAB3B | -0.579752088 | 0.006010083 |
| CD2BP2 | -1.129384133 | 3.24E-10 |
| TBC1D10B | -0.52383111 | 1.67E-06 |
| RAB24 | 0.582680642 | 7.75E-06 |
| THBS3 | 0.835475766 | 2.59E-06 |
| SLC50A1 | 1.802070014 | 2.32E-10 |
| EFNA1 | 1.585113759 | 1.72E-08 |
| SH3TC2 | -1.449621409 | 1.76E-09 |
| CXCL11 | 6.052387482 | 0.000112231 |
| NMD3 | -1.06708111 | 6.87E-09 |
| ADRB2 | 0.417517085 | 0.012696068 |
| GPRIN1 | -0.86957939 | 2.08E-07 |
| KCNAB1 | 0.691376433 | 0.01967796 |
| PGM2 | -0.549424481 | 6.68E-07 |
| STK32A | 3.226835293 | 0.000401769 |
| IL1RAPL1 | -0.522674758 | 0.000852004 |
| MINAR1 | 1.484384293 | 3.15E-08 |
| SLC33A1 | 0.263791895 | 0.001000364 |
| SNUPN | -0.866091288 | 3.37E-06 |
| SIN3A | 0.419162669 | 3.50E-06 |
| ARL13B | -0.344571386 | 0.000448086 |
| PTK2 | -0.609118539 | 9.34E-07 |
| PTAFR | 5.758100382 | 8.79E-05 |
| PTPN9 | -0.616432798 | 1.10E-06 |
| CXCL8 | 3.894083035 | 7.50E-14 |
| SDC2 | -0.358656384 | 0.004569413 |
| MMGT1 | -0.671542537 | 6.96E-08 |
| PLEKHA2 | -0.647057651 | 0.001170685 |
| CLIC4 | -0.1009507 | 0.010461081 |
| METTL15 | -1.145497755 | 5.91E-05 |
| ZNF280A | 3.428466166 | 9.92E-05 |
| ZEB2 | -0.590621063 | 3.51E-05 |
| PCBP1 | -0.711109482 | 1.68E-09 |

| | | |
|---|---|---|
| HINT1 | 0.261213723 | 0.000181064 |
| DTWD2 | -1.488168488 | 2.54E-08 |
| BNC1 | 1.405462206 | 8.55E-07 |
| DFFB | -1.644491253 | 1.41E-06 |
| ANTXR1 | -0.238369572 | 0.000819308 |
| CKAP2L | -1.214280282 | 8.84E-10 |
| C15orf40 | 0.570689354 | 1.10E-05 |
| RAMAC | -0.510624074 | 1.39E-05 |
| APLF | 2.877273791 | 0.017115502 |
| RGPD8 | -0.258630032 | 0.017755705 |
| LUZP1 | -1.008210727 | 6.21E-10 |
| BUB1 | -0.888458676 | 1.66E-08 |
| SPNS1 | -0.207163797 | 0.010035748 |
| LRRC45 | -0.322937559 | 0.038792322 |
| CHRNA5 | -1.017643392 | 0.00010989 |
| AGPAT2 | 0.349606888 | 0.001343362 |
| ASPSCR1 | -0.392403279 | 0.018306328 |
| FASN | 0.921755224 | 7.79E-11 |
| DUS1L | -0.392792158 | 6.05E-05 |
| GPS1 | 0.212890739 | 0.000699398 |
| RFNG | 0.684140427 | 2.56E-05 |
| DCXR | 1.029914964 | 2.70E-07 |
| ZNF32 | -0.777208726 | 9.06E-06 |
| LDB2 | 2.549938195 | 0.000583428 |
| LIMS1 | -1.015225312 | 7.56E-09 |
| NLGN1 | -0.287619693 | 0.033462559 |
| TAPT1 | -0.773280108 | 3.59E-07 |
| HNRNPF | -0.524950214 | 1.66E-07 |
| BTD | -0.686499327 | 3.79E-06 |
| CSGALNACT2 | -0.973750479 | 1.65E-08 |
| ROBO1 | -0.533167302 | 5.51E-07 |
| AVEN | -0.312798778 | 0.038595881 |
| P2RY1 | -0.695625246 | 0.000275371 |
| TRIM56 | 0.351295787 | 0.001879651 |
| AHSP | 7.638439334 | 2.06E-06 |
| WNT10B | -0.779650752 | 0.001320457 |
| CALML6 | 2.212277903 | 0.001467903 |
| SYAP1 | -0.171879707 | 0.008696445 |
| ITGAM | 6.588077852 | 2.32E-05 |
| TOR1AIP2 | 0.114419894 | 0.014692982 |
| TM4SF1 | -0.361015175 | 4.82E-05 |
| OTUD3 | -0.630066552 | 1.70E-06 |
| GUSB | 0.238460341 | 0.003315042 |
| BRD3 | 0.458796249 | 9.04E-06 |
| KLF13 | -1.115816562 | 1.49E-06 |
| ZFPM2 | -1.101546198 | 2.25E-06 |
| ZNF768 | 0.151629907 | 0.031684566 |
| TMEM42 | 1.137508813 | 5.42E-07 |

| | | |
|---|---|---|
| MAP3K2 | -1.432113747 | 1.71E-07 |
| SF3B5 | -0.261723919 | 0.001854614 |
| ZNF35 | -0.782808262 | 1.82E-05 |
| TIGD4 | 2.172598624 | 4.97E-07 |
| IFFO2 | 0.161833788 | 0.018195023 |
| NLGN2 | 0.337925186 | 0.000228667 |
| CHD3 | 0.745247987 | 1.50E-08 |
| ALCAM | -0.689226092 | 5.96E-06 |
| YWHAG | -0.686815158 | 1.08E-09 |
| UBE2E3 | -0.569751652 | 2.86E-06 |
| CNTROB | -0.721056647 | 2.26E-06 |
| KCNAB3 | 2.340290544 | 7.13E-10 |
| FAM153A | 3.344772539 | 0.00021621 |
| SIMC1 | -0.714241588 | 8.52E-06 |
| AC106795.1 | -0.229367803 | 0.004866551 |
| NSG2 | 3.221228797 | 0.00048472 |
| NIPA1 | -0.988004046 | 5.31E-09 |
| FOXD4 | 2.055425314 | 0.000775522 |
| HNRNPA3 | 0.198206991 | 0.000415528 |
| SIK2 | -0.590723905 | 1.03E-05 |
| RNF150 | -0.652048719 | 0.00059371 |
| CCDC144A | 0.471017374 | 0.000279019 |
| AL512625.1 | 0.616735489 | 0.001566682 |
| HOXD4 | -1.599296268 | 0.007115966 |
| CHRNB1 | 0.685493607 | 0.00093684 |
| USP38 | -0.205446643 | 0.003560336 |
| SLC16A5 | 1.113374586 | 2.75E-07 |
| ANKK1 | 7.323813629 | 0.001054934 |
| ADPRM | 1.561109848 | 6.83E-08 |
| PWWP2A | -1.233488916 | 9.32E-09 |
| USP47 | -0.357386104 | 0.000319371 |
| PDCD6IP | 0.12154581 | 0.020444318 |
| ZNF212 | -0.365723742 | 0.001686493 |
| MRAP | -0.835412572 | 0.047179625 |
| FAM161A | -0.601988498 | 0.000223965 |
| ZNF282 | -0.530580038 | 5.05E-06 |
| GLB1 | -0.235680429 | 0.007837019 |
| FAXDC2 | 0.520790354 | 0.003450227 |
| CRTAP | 0.481946791 | 1.39E-07 |
| ELP5 | -0.349255199 | 6.35E-05 |
| GABARAP | 0.364449512 | 0.000988535 |
| CDK1 | -1.601344941 | 4.96E-11 |
| UBB | 0.371654909 | 6.50E-07 |
| NFRKB | -1.306574077 | 9.93E-10 |
| PRDM10 | -0.200237002 | 0.025390525 |
| B3GNT2 | 1.106392594 | 2.08E-08 |
| FOS | 5.587437879 | 5.35E-15 |
| SMAD1 | -1.481125942 | 0.000589232 |

| | | |
|---|---|---|
| SP7 | 6.774837438 | 3.09E-05 |
| TCAF2 | -0.546120779 | 0.000520768 |
| SEMA3E | -1.021679837 | 0.027446916 |
| LRRN2 | 5.487348874 | 3.26E-08 |
| SLC30A1 | 1.795059121 | 7.61E-12 |
| DCLK2 | -1.681013204 | 0.000153765 |
| ZNF804A | -2.627420917 | 1.73E-09 |
| TMEM182 | 0.744987639 | 0.000870155 |
| KRT8 | 0.549261067 | 1.97E-06 |
| ADORA2B | -1.037347127 | 2.11E-06 |
| SDR9C7 | 7.044885087 | 1.40E-05 |
| METTL7B | -0.456145338 | 0.003884434 |
| AC021066.1 | 1.635640506 | 0.000140234 |
| HARS1 | -0.466929283 | 8.06E-07 |
| NFXL1 | 0.304885313 | 0.001133695 |
| DENND5B | -0.41425428 | 0.000140161 |
| CD14 | 2.217112183 | 4.01E-05 |
| RIOX1 | -1.805374481 | 2.20E-09 |
| SPATA24 | 0.708028688 | 0.044786007 |
| RALGAPB | -0.794576023 | 1.13E-08 |
| MZB1 | 3.520798553 | 0.005878378 |
| SLC23A1 | 0.827530309 | 0.048884114 |
| NPAS2 | -1.029798997 | 3.53E-09 |
| LONRF2 | -0.733054062 | 2.51E-06 |
| NUDT9 | -0.498145299 | 0.002019992 |
| PA2G4 | -0.122307489 | 0.008719621 |
| COX7B2 | -0.716778873 | 0.026166302 |
| ELOVL6 | -0.499909354 | 8.65E-06 |
| PFKFB3 | -0.742397363 | 4.43E-08 |
| ARL6IP1 | -0.78188457 | 1.88E-09 |
| SMAGP | -0.473492993 | 3.43E-05 |
| CDH2 | 0.195625899 | 0.002355311 |
| EMB | -0.762602312 | 1.03E-07 |
| DLGAP1 | 2.921213659 | 1.24E-08 |
| STAT2 | 0.46960251 | 2.84E-05 |
| IRF2BP1 | 0.379101962 | 0.001515615 |
| FOXA3 | 4.897641166 | 0.000175557 |
| SLC26A5 | 2.967892264 | 0.000104488 |
| COMMD5 | -1.029773464 | 1.20E-07 |
| GTSF1 | 0.35110901 | 0.005109804 |
| DPY19L2P2 | -0.415497669 | 0.009974132 |
| ZNF16 | 0.468988851 | 0.000998683 |
| ARMC10 | -0.247805609 | 0.006231308 |
| RNF34 | -0.736908569 | 1.02E-07 |
| ACYP2 | 0.94686424 | 0.000114729 |
| ATF7 | -0.146296617 | 0.027383006 |
| RASA4B | 1.537747125 | 6.65E-05 |
| TTLL6 | 2.397135403 | 0.000854902 |

| | | |
|---|---|---|
| POLH | -0.595563514 | 2.69E-05 |
| KCNS3 | 0.810087702 | 9.21E-05 |
| KIF5B | -0.779269926 | 3.23E-09 |
| GPR37 | 0.452227203 | 0.042280257 |
| AKAP13 | -0.662567893 | 9.65E-08 |
| CDCA4 | -1.807444467 | 1.03E-10 |
| CHCHD7 | 0.178705036 | 0.011846337 |
| USP32 | -0.176141017 | 0.029126349 |
| CEL | 1.930279068 | 0.001964978 |
| PPM1D | 0.907788404 | 1.45E-08 |
| AC093323.1 | 0.842730128 | 1.14E-08 |
| KBTBD2 | -0.248269019 | 0.000277426 |
| RIOX2 | -0.77270914 | 9.10E-08 |
| TRIAP1 | -0.624620322 | 1.39E-06 |
| KIAA0232 | 0.139699069 | 0.016314361 |
| MTSS1 | 0.798066536 | 4.99E-05 |
| TMEM43 | -0.189514559 | 0.001524688 |
| RNF139 | -0.3630051 | 0.000852526 |
| RPS9 | 0.292319486 | 0.000127882 |
| GSTA4 | 1.193448511 | 1.83E-06 |
| MSANTD4 | 0.157883586 | 0.02106628 |
| PAQR8 | 1.401179731 | 2.18E-07 |
| NUDT6 | 0.799007189 | 0.000135744 |
| TPT1-AS1 | 1.066689856 | 5.84E-07 |
| TANC2 | -0.552374659 | 5.03E-07 |
| DNAJC24 | -1.30333764 | 3.50E-09 |
| ZNF160 | 0.660308538 | 1.47E-07 |
| ZNF415 | 2.473420373 | 7.10E-08 |
| PDGFD | -1.421391328 | 2.30E-06 |
| HS6ST2 | -0.422799901 | 3.04E-05 |
| PYGO1 | -1.980598693 | 5.43E-07 |
| PKIA | -0.671360142 | 0.000471928 |
| TSNARE1 | -0.490990753 | 0.017386832 |
| PATE1 | 4.325686904 | 0.001200606 |
| SOX7 | 1.099925411 | 0.036516899 |
| C11orf24 | -0.557835994 | 2.82E-06 |
| FAM86JP | 2.452423306 | 1.72E-09 |
| TRMT61B | -1.155834641 | 5.52E-07 |
| INSR | -0.257417076 | 0.008349063 |
| MFN1 | -0.657055175 | 6.56E-07 |
| GIMAP8 | 8.985888098 | 2.55E-08 |
| KCNMB3 | -1.162756209 | 2.01E-05 |
| KCNG3 | -2.303505488 | 0.000842079 |
| PRKCE | -1.419238594 | 5.88E-07 |
| JAGN1 | -0.172950051 | 0.007873948 |
| TADA3 | 0.588226023 | 1.91E-06 |
| SOCS5 | -0.241511768 | 0.001309045 |
| C9orf16 | 0.614718814 | 6.96E-06 |

| | | |
|---|---|---|
| MORN4 | 0.695315528 | 6.38E-05 |
| ZNF672 | -1.03356087 | 7.83E-09 |
| ZNF692 | -1.394982577 | 1.37E-08 |
| NAIF1 | -0.524295799 | 0.000211906 |
| RBKS | 5.28250242 | 2.54E-09 |
| TMEM126B | -0.584837745 | 6.99E-06 |
| TRIM8 | -0.911423445 | 2.35E-07 |
| NETO2 | -0.96952927 | 3.85E-09 |
| CLDN20 | 1.466679792 | 0.005507391 |
| CDC42BPG | 1.986727755 | 0.00360126 |
| JUNB | 3.034325701 | 1.43E-11 |
| FAM241B | -0.934348387 | 0.00808288 |
| UGT2B7 | 1.217950803 | 0.043143128 |
| SHCBP1 | -1.503272378 | 1.03E-10 |
| SOSTDC1 | 7.583192658 | 2.48E-06 |
| FAM98B | -1.054690631 | 1.67E-09 |
| ZNF439 | 0.759746045 | 0.000462653 |
| ZNF440 | 0.80757098 | 4.72E-08 |
| GAA | 0.792435926 | 9.51E-07 |
| CANT1 | -0.632485071 | 6.39E-07 |
| PGAM1 | -0.170875278 | 0.001508288 |
| CHD7 | -1.963527036 | 5.27E-11 |
| ESCO2 | -0.954335092 | 2.72E-08 |
| LURAP1 | 1.560656146 | 1.35E-05 |
| CLCN5 | -1.750345426 | 6.32E-10 |
| TPPP | 1.626106787 | 3.00E-06 |
| APLN | -1.078084049 | 0.003676528 |
| MRPL36 | -0.65383447 | 7.88E-06 |
| ZNF581 | 0.431314005 | 0.000379018 |
| NAT1 | 1.078905634 | 1.67E-05 |
| MCC | -0.483057092 | 0.000149186 |
| CDK5R2 | 1.730075315 | 0.004779089 |
| DSEL | -1.4352873 | 4.08E-07 |
| POLR1C | -0.41556369 | 1.84E-05 |
| ASXL1 | -1.284273986 | 4.44E-10 |
| ZNF562 | -0.437978378 | 5.71E-06 |
| ZNF318 | -1.083425975 | 6.26E-09 |
| ZNF561 | -0.322478617 | 8.10E-05 |
| WIPF2 | -0.191596533 | 0.001955064 |
| NLRP5 | 3.259113771 | 0.000420955 |
| LRRC8C | -1.998039018 | 4.88E-10 |
| LRRC8D | -0.900022997 | 6.10E-09 |
| PPID | -0.259229755 | 0.00312918 |
| LPAR3 | 0.506201456 | 7.13E-06 |
| PTGER4 | 0.98640609 | 0.004232698 |
| TBCA | 0.350481894 | 0.000535574 |
| BCL2L1 | -1.498506696 | 6.06E-10 |
| PLRG1 | -0.16823058 | 0.025326553 |

| | | |
|---|---|---|
| RAB4B-EGLN2 | -0.397364216 | 0.000197192 |
| CLSTN1 | -0.12203858 | 0.018446872 |
| CXXC5 | -0.210380507 | 0.043499258 |
| ZNF274 | 0.218308644 | 0.004019229 |
| PIK3CD | 0.6565128 | 5.71E-05 |
| SLC25A33 | -0.164437697 | 0.045690739 |
| ENC1 | -1.58005365 | 1.58E-10 |
| SPSB1 | 1.05274658 | 2.19E-07 |
| BPTF | -1.488382283 | 5.10E-11 |
| NMRAL2P | -1.489107737 | 0.000656812 |
| PLEKHG5 | 0.377560615 | 0.001261278 |
| ATF7IP | -0.344776132 | 1.06E-05 |
| RGS19 | -0.267181598 | 0.029593119 |
| TCEA2 | 0.527467 | 0.00020038 |
| ANO5 | -1.122342018 | 0.01650403 |
| HDAC3 | 0.1597187 | 0.043816393 |
| GPHN | -1.129577832 | 2.30E-06 |
| VAT1L | 0.633499455 | 0.000218255 |
| TMEM51 | 0.388911459 | 0.024018217 |
| CAMTA1 | 0.782930259 | 1.96E-05 |
| SPATA5L1 | -1.56789333 | 2.30E-08 |
| GATM | 3.610343947 | 6.36E-07 |
| NHLH1 | 2.654266924 | 1.68E-07 |
| SLFNL1 | -0.892012542 | 0.000799359 |
| BCL2 | -1.447494983 | 8.90E-08 |
| RHNO1 | 0.438224873 | 5.42E-06 |
| CTPS1 | -1.332245638 | 2.72E-10 |
| METTL18 | -1.0168426 | 3.10E-06 |
| COL8A2 | 2.366971717 | 1.11E-05 |
| PWWP2B | -0.596428703 | 0.005114368 |
| PCDHB1 | 3.392168575 | 2.15E-06 |
| EXOSC10 | -0.545917568 | 1.08E-06 |
| ZNF570 | 0.461306137 | 0.000690379 |
| MLLT3 | -0.686824178 | 1.31E-05 |
| FAM90A1 | 2.24469883 | 1.51E-06 |
| RRM2 | -1.647494342 | 1.80E-12 |
| TRAPPC12 | -0.508815943 | 0.000206209 |
| RPS21 | 0.452369801 | 3.76E-06 |
| C3AR1 | 1.673103886 | 0.000221051 |
| MRM3 | -0.249951481 | 0.005714433 |
| PTEN | -0.855101701 | 1.20E-08 |
| RNASEH1 | -0.318871584 | 0.00257577 |
| PRNP | 0.149503294 | 0.000588773 |
| FRMD5 | -0.502674578 | 3.99E-05 |
| TLN2 | -0.444498268 | 0.005408247 |
| FBXW10 | 5.5410052 | 7.16E-08 |
| SRGAP2C | 0.571310883 | 1.20E-06 |
| SCG2 | 3.435725636 | 4.05E-13 |

| | | |
|---|---|---|
| ZNF57 | 1.329793891 | 8.66E-07 |
| SHLD1 | 1.09190212 | 4.79E-05 |
| LDHAL6B | 2.907890405 | 2.05E-05 |
| SYNPO | -0.384937518 | 0.000185431 |
| ZNF556 | 1.228236529 | 7.57E-05 |
| ZNF554 | 2.26201099 | 9.47E-07 |
| THOP1 | -0.257231422 | 0.002077101 |
| EPHX4 | -0.913631087 | 0.001725666 |
| LAMB2 | 0.453186002 | 4.49E-06 |
| USP19 | -0.238386072 | 0.000374735 |
| QARS1 | 0.233542768 | 0.001108663 |
| ORMDL3 | -1.160140275 | 9.23E-08 |
| KLF11 | 0.992103971 | 8.65E-06 |
| LRRC15 | -0.482836116 | 0.048802378 |
| SMN1 | -0.554231012 | 1.46E-06 |
| EIF2AK3 | 0.944372751 | 1.68E-07 |
| KRCC1 | 3.520899088 | 3.86E-12 |
| CYCS | -0.766620194 | 1.59E-09 |
| SLFN12 | -0.807163123 | 4.71E-07 |
| SLC9C1 | 7.666183637 | 1.84E-06 |
| FRMD3 | -1.164265447 | 3.56E-08 |
| MTBP | -0.702582662 | 2.82E-05 |
| MALT1 | -2.276271564 | 2.09E-11 |
| ISG20 | 0.892254927 | 4.99E-08 |
| MBOAT1 | -1.749178556 | 3.62E-05 |
| ID4 | 2.989914226 | 1.31E-07 |
| CXCR6 | -0.955239932 | 0.043678303 |
| CEBPB | 1.548042309 | 2.30E-09 |
| PAIP1 | -0.42249273 | 0.000547256 |
| C5orf34 | -0.281350605 | 0.017622308 |
| SERHL | -1.36259783 | 6.74E-06 |
| NEGR1 | -0.745317875 | 3.03E-06 |
| ZNF131 | -0.155596191 | 0.014815768 |
| MACROD2 | 1.180894724 | 0.000420481 |
| DPAGT1 | 0.281631607 | 0.000343543 |
| BSG | 0.229630544 | 7.27E-05 |
| HINFP | 1.093538698 | 3.05E-08 |
| CERS6 | -0.680402804 | 3.18E-07 |
| TP53RK | -0.809325609 | 1.04E-07 |
| B3GALT1 | -0.722529367 | 0.016941835 |
| BPGM | -1.039353605 | 7.42E-07 |
| POP7 | -1.241496181 | 1.56E-08 |
| ALG14 | -0.959839664 | 0.000334008 |
| STARD5 | 0.859915282 | 0.000523254 |
| IL16 | 0.633999736 | 0.003437514 |
| ABCG4 | 3.139963335 | 2.22E-08 |
| GNB2 | 0.474957951 | 6.43E-07 |
| CFAP53 | 1.209652667 | 0.001395608 |

| | | |
|---|---|---|
| OR5B2 | 6.871264811 | 2.36E-05 |
| MCRIP2 | 0.379616009 | 0.015754778 |
| ARNT2 | 0.176815409 | 0.008390044 |
| GNG12 | -0.177002889 | 0.000186895 |
| PRSS27 | 1.770741438 | 0.002601173 |
| MYOZ2 | 5.574841492 | 0.000491611 |
| SYNPO2 | 3.510195884 | 1.32E-05 |
| TTC36 | 4.237383384 | 0.003125868 |
| GTPBP2 | -0.578835662 | 1.01E-05 |
| FGGY | 2.153973116 | 0.003641738 |
| FUT9 | -0.765469859 | 0.002743635 |
| TCEAL1 | 0.397395414 | 0.014383101 |
| ZNF24 | -0.679007949 | 3.26E-08 |
| MANEA | -1.51756809 | 6.40E-09 |
| RAB40A | 1.745755404 | 0.003891304 |
| AFF1 | -0.924217035 | 8.39E-08 |
| FIBP | 0.277849787 | 0.000446808 |
| CARNS1 | 1.401636246 | 0.007720305 |
| PPP1CA | 0.13445709 | 0.014168814 |
| HCFC1 | -1.006370129 | 5.70E-10 |
| RASGRP1 | 0.924453252 | 0.019518915 |
| CHCHD1 | -0.57645771 | 5.50E-05 |
| SMPDL3A | 0.503620413 | 0.03896086 |
| RND1 | 2.410721986 | 1.58E-06 |
| RAD9A | 0.203923387 | 0.02634934 |
| AGAP5 | 0.993552812 | 5.16E-05 |
| TMEM134 | 0.456166479 | 0.02342169 |
| FAM71D | -0.826927163 | 0.00056293 |
| CORO1B | 0.672583298 | 1.26E-07 |
| FUT10 | -0.993809836 | 2.19E-06 |
| LRRC20 | 1.205632257 | 2.48E-05 |
| MUS81 | -0.791176704 | 7.42E-07 |
| TMEM217 | 1.662831803 | 0.000904139 |
| ZNF596 | 2.907651171 | 3.29E-10 |
| TMCC1 | 0.670320345 | 7.01E-05 |
| NAA16 | -1.032969746 | 1.33E-07 |
| EFCAB12 | 2.431169257 | 0.008233864 |
| PSME3IP1 | -0.382419358 | 2.69E-05 |
| RAB43 | -0.301715933 | 0.004340145 |
| CBWD1 | -0.234460571 | 0.020480624 |
| HOXC5 | -1.312119166 | 0.000118536 |
| DCP2 | -1.638735617 | 3.13E-11 |
| RPL38 | 0.335197955 | 2.74E-05 |
| OVOL1 | 3.82889598 | 3.29E-05 |
| CES4A | 4.916093134 | 0.00044506 |
| CES2 | 0.682675791 | 1.32E-06 |
| PDP2 | -0.610220687 | 5.36E-07 |
| SP3 | -0.334591501 | 1.86E-05 |

| | | |
|---|---|---|
| LSM2 | -0.271002404 | 0.048230301 |
| DMXL1 | -0.654283821 | 7.82E-07 |
| METAP1D | -0.477924285 | 0.010965126 |
| EGFL7 | 0.734042218 | 0.002476569 |
| NADSYN1 | 0.766944394 | 3.18E-06 |
| DHCR7 | 0.91751083 | 2.85E-08 |
| LVRN | 1.825960802 | 3.72E-05 |
| RNASEH2C | -0.238205539 | 0.001224748 |
| MYEOV | 0.511145688 | 0.008626407 |
| ANKRD13D | -0.22558592 | 0.009885661 |
| MYD88 | -0.996464045 | 2.27E-08 |
| MRGPRD | 4.6784278 | 0.000390666 |
| OXSR1 | -1.062341637 | 7.24E-10 |
| PHF8 | -0.969644979 | 1.18E-08 |
| LCLAT1 | -1.767142334 | 2.53E-10 |
| ADH6 | 3.387496395 | 0.000207057 |
| MIR4435-2HG | -0.659343609 | 6.65E-05 |
| FRG2C | 2.89920139 | 0.039611847 |
| AC007318.1 | 0.239896577 | 0.000131828 |
| KAT5 | -0.602769418 | 3.21E-06 |
| SH3RF3 | -0.605355809 | 0.002027319 |
| GXYLT2 | -0.403055132 | 0.009737686 |
| DCAKD | 0.358731925 | 0.001683186 |
| ARPP21 | 0.524818924 | 0.025951126 |
| TADA2B | -0.582751043 | 5.24E-06 |
| CCDC96 | 1.61122829 | 1.07E-05 |
| GRK2 | -0.33233334 | 7.86E-05 |
| RELA | -0.371628751 | 1.69E-05 |
| HECTD4 | -0.630098755 | 4.62E-07 |
| FAM222B | -0.730335107 | 2.02E-07 |
| BNC2 | -0.980273356 | 3.40E-08 |
| RXFP4 | 2.640575031 | 0.000131437 |
| COQ2 | -0.701529001 | 0.000556184 |
| HSPA6 | -0.95494348 | 0.001719647 |
| TRMT112 | 0.134796029 | 0.014649229 |
| KDM2A | -0.778361347 | 1.97E-08 |
| NOC3L | -0.889989158 | 6.04E-09 |
| COMMD1 | 0.214289888 | 0.001529425 |
| RAPH1 | -0.957592017 | 2.36E-08 |
| MTX1 | 0.25885396 | 0.002086277 |
| ADCY5 | 1.793892595 | 0.000312235 |
| PARP14 | -0.76670876 | 8.82E-08 |
| CKS1B | 0.498966719 | 5.02E-06 |
| AHSA2P | -1.241536014 | 4.07E-10 |
| TUBB8B | 2.018018873 | 0.002510378 |
| MFSD4B | 0.505862431 | 0.000707486 |
| VANGL1 | -0.761419302 | 3.10E-08 |
| GLRX | 0.47504675 | 0.000205219 |

| | | |
|---|---|---|
| IQCB1 | 0.512444142 | 6.56E-05 |
| GOLGB1 | -0.461094685 | 2.23E-06 |
| DMRT2 | -1.75869132 | 4.46E-07 |
| PLAC8L1 | 1.881412051 | 0.000264518 |
| SLC2A14 | 3.153402761 | 8.11E-09 |
| GPR137 | 0.153850645 | 0.030612437 |
| MMRN2 | 3.441260024 | 2.74E-08 |
| ZNF449 | -0.578544703 | 0.001592553 |
| ZBTB21 | 1.315731631 | 2.04E-10 |
| PPP1R3B | 1.562094269 | 2.72E-08 |
| FAM86B3P | 1.514723431 | 1.02E-06 |
| STOX2 | -0.434983726 | 0.044050661 |
| MAP3K11 | -0.342741781 | 7.20E-05 |
| TRIB1 | 1.527293334 | 1.80E-09 |
| SFT2D3 | -0.714070509 | 0.000186822 |
| AC097637.1 | 1.086414132 | 2.59E-07 |
| DAG1 | 0.208562587 | 4.79E-05 |
| NAA20 | -0.337762365 | 4.56E-05 |
| EHBP1L1 | 0.260715904 | 0.000171287 |
| THAP2 | 1.52476404 | 1.45E-09 |
| RNF26 | -0.580772517 | 8.55E-07 |
| ZNRD2 | -0.210347462 | 0.003710803 |
| SMARCC1 | -0.336166993 | 5.65E-06 |
| ZNF417 | -0.212259998 | 0.017413569 |
| FKBP2 | 0.551617756 | 0.000156137 |
| LTA | 5.152774786 | 7.18E-05 |
| VEGFB | 0.563017807 | 4.21E-05 |
| PEAK1 | -0.9881418 | 2.54E-09 |
| TNFRSF10C | -0.946485078 | 0.008698564 |
| GMPPB | -0.327129739 | 6.36E-05 |
| ZNF622 | 0.202284516 | 0.004929151 |
| CSPG4 | 0.810234303 | 3.50E-10 |
| NABP1 | -1.501994948 | 2.42E-08 |
| ADGRF3 | 3.200215777 | 2.34E-08 |
| CHD2 | 0.38752889 | 8.38E-06 |
| CCDC106 | 0.699906438 | 0.000165521 |
| CEP83 | -0.938805209 | 6.43E-07 |
| SULT1B1 | 1.665725232 | 3.21E-05 |
| NUDT4 | -0.213065448 | 0.004207216 |
| PC | 0.258360976 | 0.002950784 |
| SCAI | -0.662961862 | 2.64E-05 |
| NMNAT1 | 1.573283575 | 1.60E-08 |
| LRFN4 | -0.364624589 | 0.000943641 |
| TRAPPC3L | 7.00668098 | 2.14E-05 |
| SLC19A1 | -0.56656688 | 7.02E-06 |
| RCE1 | -0.548332161 | 9.57E-05 |
| UQCRH | 0.288462389 | 0.000250123 |
| EIF1AX | -0.988032421 | 6.74E-09 |

| | | |
|---|---|---|
| BCLAF3 | -0.626588392 | 0.000114588 |
| SUSD5 | 0.273961338 | 0.000718677 |
| HEG1 | -1.226652393 | 2.31E-10 |
| WFIKKN2 | 8.175750101 | 3.01E-07 |
| C11orf80 | 0.526440593 | 0.008672943 |
| TOMM20 | -0.440249957 | 2.96E-07 |
| AGFG1 | -0.983687439 | 4.89E-09 |
| STAT5B | -0.304232044 | 0.001795407 |
| CNP | -0.135333405 | 0.010745129 |
| JUP | 0.576598436 | 0.005079684 |
| TDRD12 | 3.459387584 | 0.000437072 |
| EIF1 | -0.183506934 | 0.000166674 |
| ENDOV | 1.030087214 | 1.44E-06 |
| RNF213 | 0.565822209 | 4.25E-07 |
| TIGD3 | 2.284169818 | 1.89E-07 |
| MARCHF10 | 2.334632601 | 8.37E-07 |
| PLK3 | 1.65618011 | 5.31E-10 |
| DPY19L1 | -0.740046019 | 5.36E-08 |
| AC013489.1 | -1.101983312 | 3.53E-05 |
| ZNF791 | -0.12727296 | 0.03163107 |
| PHC3 | -0.784488876 | 6.85E-08 |
| CBX2 | -0.590610942 | 7.42E-07 |
| SPTBN2 | 0.405014256 | 2.91E-05 |
| GOLIM4 | -0.403638259 | 2.07E-05 |
| RBM4B | -1.638623619 | 9.57E-10 |
| ATP5MD | 0.247322936 | 0.001061465 |
| HOXB2 | -1.546799068 | 1.42E-09 |
| MARCHF3 | -0.866875587 | 0.006520272 |
| RBM4 | -0.7979391 | 2.09E-08 |
| XXYLT1 | -0.373169967 | 0.00030471 |
| UBXN2A | -0.855785029 | 3.68E-07 |
| TCAP | 1.593372194 | 1.32E-05 |
| CCS | 0.700052281 | 1.97E-05 |
| NRROS | -0.980477792 | 0.003917506 |
| CEP19 | 2.117241002 | 1.12E-08 |
| FBXO45 | -0.530025157 | 5.96E-07 |
| GNG5 | 0.235957034 | 0.010799585 |
| FAM3C2 | -0.986180882 | 2.06E-09 |
| C9orf131 | 2.431230112 | 0.000213368 |
| CTSF | 0.588706003 | 0.000192229 |
| MSRB3 | -0.985496639 | 7.27E-07 |
| LEMD3 | -0.345046799 | 3.32E-05 |
| C16orf91 | -0.409129956 | 0.000271062 |
| TLR10 | 4.307090806 | 0.001033285 |
| TLR1 | 1.504418157 | 0.000180736 |
| TLR6 | -1.053614469 | 1.17E-05 |
| RGMB | -0.924140522 | 1.17E-08 |
| ZDHHC24 | 0.588054042 | 0.001994192 |

| | | |
|---|---|---|
| AC020659.1 | 0.979281346 | 0.000767256 |
| TRMT10C | -0.87411766 | 3.72E-09 |
| CTU2 | -1.012711975 | 5.57E-07 |
| MGA | -0.782223063 | 2.04E-08 |
| C12orf66 | 0.622681557 | 6.33E-05 |
| SNX31 | 9.621347754 | 2.71E-09 |
| ADCY6 | 0.140212636 | 0.021400142 |
| PITPNA | -0.866855402 | 5.00E-09 |
| DDX23 | -0.493539173 | 6.39E-07 |
| ZNHIT2 | -0.571551794 | 0.001346956 |
| ZBTB4 | -0.517772307 | 5.00E-06 |
| TNK1 | 1.921043959 | 2.25E-07 |
| ZHX3 | -0.950646363 | 3.32E-09 |
| PHLDA3 | 0.510841695 | 3.47E-06 |
| SLC16A13 | 2.576649977 | 4.14E-08 |
| STAG3L3 | -0.827285223 | 8.35E-06 |
| EXO1 | -1.584640527 | 3.12E-10 |
| RALGAPA1 | -0.455867886 | 0.000424047 |
| LIG4 | -0.345803037 | 9.05E-05 |
| GTF2IRD2B | 1.721301172 | 1.06E-07 |
| ATP2A2 | -1.057823824 | 4.96E-11 |
| ZWILCH | -0.412168723 | 1.56E-05 |
| RPL4 | 0.236141393 | 3.69E-05 |
| SNAPC5 | 0.247006567 | 0.006051256 |
| C12orf76 | 1.323391902 | 1.45E-07 |
| GALNTL6 | -1.026228467 | 0.008416002 |
| BBS1 | 0.36966301 | 0.000999376 |
| DENND4A | -0.514426941 | 8.10E-06 |
| MFSD4A | 1.854899965 | 0.000101524 |
| TTC9B | 2.444103892 | 0.008913149 |
| MYO1H | 2.41672284 | 2.00E-05 |
| MRPL11 | 0.176588037 | 0.027265849 |
| IL20RB | -1.161165304 | 0.023181223 |
| GOLT1A | 1.984333976 | 0.005811006 |
| AKIRIN1 | -1.118580189 | 5.96E-10 |
| MSL2 | 0.147101617 | 0.015349642 |
| ZNF497 | 0.765084003 | 7.44E-05 |
| TRAM1L1 | -1.480962542 | 6.50E-06 |
| ANGEL2 | 0.420781948 | 6.16E-06 |
| UGT8 | -1.513899749 | 8.82E-10 |
| IQCK | 0.826327706 | 0.000316217 |
| SLC29A2 | 0.60894199 | 0.001500938 |
| B4GAT1 | -0.362244584 | 0.000366084 |
| TMEM167A | -1.439609324 | 5.03E-11 |
| RESF1 | 0.207748277 | 0.005224898 |
| FGFBP3 | 1.238852491 | 8.26E-05 |
| NR1D2 | 1.182219695 | 3.48E-10 |
| PABPC5 | 1.981812922 | 2.43E-06 |

| | | |
|---|---|---|
| BRMS1 | -0.280575821 | 0.000515031 |
| RPL15 | 0.262736376 | 6.42E-06 |
| FAM241A | -0.987923174 | 4.81E-08 |
| SRP72 | -0.350834438 | 2.01E-05 |
| RIN1 | -0.314972199 | 0.0029684 |
| ODAPH | 9.29529347 | 2.07E-08 |
| THAP6 | -0.440869294 | 0.000144529 |
| CEP135 | -0.397366873 | 0.000661725 |
| FZD4 | -0.378331369 | 0.001205744 |
| DENND6A | -1.431494541 | 2.31E-09 |
| PDE12 | -1.181792956 | 5.53E-10 |
| GLMN | -1.214501112 | 6.36E-06 |
| DNAH12 | 6.91682617 | 0.012813945 |
| YIF1A | 0.403987876 | 0.000161139 |
| RSRC1 | -0.537075637 | 0.000529886 |
| RAB1B | -0.126967499 | 0.025341241 |
| METTL15P1 | -0.60080753 | 0.009691762 |
| MICOS13 | 0.416775956 | 0.014433412 |
| C3orf33 | -1.308703297 | 6.48E-05 |
| SEZ6L2 | 0.424931992 | 0.000543786 |
| ASPHD1 | 0.54306219 | 0.007679207 |
| AMZ1 | 2.279119326 | 0.000216042 |
| DHX36 | -0.292031744 | 0.000534535 |
| ZIC4 | 1.344236412 | 0.000408775 |
| AC026271.1 | -0.35918168 | 1.50E-05 |
| FBXW8 | 0.594450539 | 0.000114003 |
| KLC2 | -0.368830765 | 7.50E-05 |
| CTBP2 | -0.75464489 | 2.96E-08 |
| ZDHHC14 | -0.449426481 | 0.005054928 |
| SNHG29 | 0.487126601 | 7.82E-08 |
| UBE2C | -1.041849438 | 1.99E-09 |
| GK5 | -0.61851434 | 5.55E-05 |
| VCPIP1 | 0.305267709 | 9.05E-05 |
| PDIK1L | -0.334461099 | 0.003372873 |
| SPSB4 | -1.453613289 | 1.13E-05 |
| TRAF6 | -0.609302424 | 3.74E-05 |
| TVP23C | 0.430629313 | 0.001381904 |
| PACS1 | -0.380159895 | 0.000218357 |
| MARCKSL1 | -0.284424154 | 4.74E-05 |
| SH3BP5L | -0.591692875 | 1.12E-06 |
| TMEM51-AS1 | 3.598079773 | 0.000273812 |
| YPEL2 | 1.365023715 | 1.35E-07 |
| ABO | 5.746375351 | 7.02E-05 |
| FAM182B | 3.869981737 | 1.56E-06 |
| PPM1E | -0.664540311 | 8.53E-05 |
| FAM131A | 1.054991803 | 2.83E-06 |
| CSRP2 | 0.808465056 | 2.42E-06 |
| INHBC | 4.729383627 | 1.07E-06 |

| | | |
|---|---|---|
| DDIT3 | 5.015933857 | 6.91E-14 |
| DCTN2 | 0.321669904 | 8.22E-05 |
| ZNF408 | 0.760425769 | 2.19E-06 |
| CTDSP2 | -0.45864767 | 2.86E-06 |
| CKAP5 | -0.409996963 | 5.75E-07 |
| ARHGAP1 | 0.239360838 | 0.001600626 |
| ATG13 | -0.287724342 | 0.000312024 |
| CHST1 | -0.676380196 | 4.68E-05 |
| GOLGA8A | 0.578401687 | 2.16E-07 |
| VWA3A | 3.63511165 | 8.09E-07 |
| CENPS | 0.386517448 | 0.002601917 |
| DOLK | -0.458814397 | 0.000473743 |
| PHYHD1 | 7.76823162 | 1.30E-06 |
| CATSPER1 | 1.000448834 | 0.000588919 |
| CCNE2 | -1.158345967 | 2.46E-09 |
| PHYKPL | -0.442090552 | 0.000306524 |
| ZNF519 | 0.303368237 | 0.016538498 |
| LSM1 | -0.558096841 | 0.000122315 |
| TMEM9B | -0.846672535 | 3.92E-07 |
| NRIP3 | -0.625570723 | 0.002565229 |
| EIF1AD | -0.809393902 | 4.17E-08 |
| SMAD2 | -0.830586899 | 4.24E-08 |
| EIF3F | 0.407952994 | 8.33E-05 |
| ZNF25 | 2.273979625 | 3.99E-09 |
| ARL10 | -0.330129048 | 0.033208184 |
| PCSK1 | 1.20887755 | 2.95E-06 |
| RFESD | 1.178293202 | 1.89E-05 |
| CCDC14 | -0.472819173 | 9.23E-07 |
| SART1 | -0.165272974 | 0.010812673 |
| PPP2R2D | -0.526046606 | 3.72E-05 |
| MCTP1 | -0.465367348 | 0.0023299 |
| POLD4 | 1.401122862 | 5.79E-09 |
| CLCF1 | 1.111285681 | 9.79E-08 |
| TSGA10IP | 5.366300968 | 3.43E-05 |
| UBQLNL | 7.43539413 | 4.03E-06 |
| LIPT2 | 1.361720154 | 2.72E-05 |
| CABP4 | 8.22929726 | 2.75E-07 |
| ALG10B | 0.336825696 | 0.000486178 |
| LONRF3 | -1.435711517 | 2.54E-06 |
| UCP3 | 1.474902093 | 2.90E-05 |
| UCP2 | 0.859565218 | 0.000139789 |
| C11orf68 | -0.639136756 | 3.87E-06 |
| PAAF1 | 1.012838661 | 1.31E-05 |
| RAB6A | -0.489273154 | 1.96E-06 |
| FOSL1 | -1.889413541 | 1.36E-12 |
| ERCC4 | -0.281450366 | 0.001135584 |
| CCDC85B | -1.322600866 | 3.01E-06 |
| TMEM70 | -0.608107734 | 4.98E-07 |

| | | |
|---|---|---|
| LINC00476 | 1.410395866 | 4.64E-06 |
| RPS6KB2 | -0.281691208 | 0.00306898 |
| TOM1L2 | 0.661406623 | 6.15E-07 |
| ZNF77 | 2.19356524 | 2.23E-09 |
| GPR156 | 1.320964153 | 3.73E-05 |
| B3GNTL1 | -0.777755725 | 0.00138568 |
| MLXIP | -1.37013002 | 3.37E-10 |
| BAK1P1 | -0.699419519 | 0.007527548 |
| NR2F1 | -1.251562257 | 2.48E-05 |
| C15orf54 | -2.793082695 | 1.17E-07 |
| TTLL11 | 0.706339561 | 0.001599709 |
| TOMM5 | -0.793264383 | 8.36E-09 |
| RUVBL1 | -0.117039831 | 0.048931606 |
| SFN | 2.055386243 | 0.003548453 |
| CCDC168 | -0.679060047 | 0.015360706 |
| CTDNEP1 | -0.305439622 | 5.97E-05 |
| ETV4 | 0.164453551 | 0.011859344 |
| FAM172BP | 3.798709858 | 5.05E-05 |
| TMEM270 | 3.507911551 | 0.000349663 |
| RPL7AP66 | 0.349151394 | 3.15E-05 |
| ZDHHC21 | -0.866522484 | 1.31E-07 |
| TSPEAR | 1.707383782 | 0.000832529 |
| ARL4D | 1.002292247 | 1.37E-06 |
| UBE2O | -0.201884157 | 0.00280602 |
| ORAI3 | 0.85576917 | 0.000189078 |
| DENND2C | 0.623036976 | 0.013888072 |
| PLEKHD1 | 8.348752232 | 2.34E-07 |
| TUBB6 | -0.449942645 | 8.60E-07 |
| LYSMD3 | 0.422966985 | 4.07E-06 |
| AMIGO3 | -0.946217542 | 9.47E-06 |
| ZNF613 | 0.512009402 | 0.001746887 |
| AC007160.1 | 1.174956312 | 0.019839821 |
| RPL23P2 | 1.197266239 | 0.017605539 |
| MBLAC2 | 0.292999732 | 0.010546815 |
| SLC35A4 | -0.390710255 | 6.25E-06 |
| CRYBG2 | 3.559960505 | 8.57E-05 |
| IP6K1 | -0.529263316 | 6.14E-07 |
| SSNA1 | 0.3271232 | 0.001339038 |
| CSTF3 | 0.863623478 | 9.86E-09 |
| YES1 | -0.228923149 | 0.000775271 |
| CHMP6 | -0.570794378 | 0.000771048 |
| DLEU1 | -0.262201217 | 0.003941814 |
| TMEM39A | -0.878914737 | 6.11E-08 |
| TCP11L1 | -0.724826973 | 4.44E-06 |
| CCDC57 | 0.53110455 | 2.53E-05 |
| SPHK1 | -0.536209214 | 0.000871219 |
| ATAD5 | -0.894843012 | 1.77E-08 |
| SMIM19 | 1.494503517 | 6.64E-07 |

| | | |
|---|---|---|
| ZNF404 | 1.563004577 | 4.53E-06 |
| ANAPC2 | -0.885081175 | 1.81E-07 |
| SLC35G1 | -1.292628774 | 1.66E-09 |
| ZNF135 | 1.15588611 | 6.85E-05 |
| COX8A | 0.377393821 | 0.000124729 |
| ZSCAN2 | 1.050742222 | 5.89E-05 |
| B3GNT4 | -0.263757051 | 0.001247411 |
| HSD11B2 | -2.150857308 | 0.002274606 |
| CRLF3 | -0.932228393 | 3.02E-08 |
| EID2 | -0.581534697 | 5.82E-06 |
| DMRTA1 | -1.866285992 | 2.07E-07 |
| GJC3 | 3.08453284 | 0.000307379 |
| RIMS2 | 1.044738641 | 0.006535936 |
| KCMF1 | -0.591090959 | 4.96E-07 |
| SPRYD4 | -0.251171691 | 0.028228921 |
| SYNE3 | 0.420341624 | 7.55E-05 |
| CLK2 | -0.485629789 | 0.000617621 |
| LPCAT4 | -0.428733354 | 0.000925631 |
| SLCO3A1 | -0.692504822 | 2.49E-05 |
| WDR25 | 2.011845218 | 2.15E-08 |
| PLAAT3 | 0.266661742 | 0.014162021 |
| PHLDB3 | 0.780602192 | 0.000172403 |
| GNG7 | 0.808738757 | 0.001037241 |
| USF3 | -0.301802043 | 0.000485519 |
| CNTD1 | -0.26379668 | 0.00150491 |
| DCAF4L2 | 1.717841759 | 0.027250193 |
| AC008969.1 | 1.544977898 | 4.89E-07 |
| B3GNT5 | -0.630246707 | 9.61E-06 |
| LMNB2 | -0.458446282 | 9.26E-07 |
| RMDN1 | 0.374547401 | 0.000201252 |
| MEX3C | -0.890869685 | 2.08E-08 |
| RNF152 | -0.979828141 | 0.001456475 |
| MYO1D | -0.615393728 | 0.002699591 |
| C20orf197 | 1.671268994 | 0.002116816 |
| FOXL1 | 2.187967949 | 3.91E-06 |
| LRRC37A | 0.733643051 | 0.000534016 |
| FOXC2 | -1.626949279 | 2.02E-08 |
| BDNF | 0.435083666 | 0.003950153 |
| SCAND2P | -0.606607376 | 4.55E-07 |
| CCDC121 | 2.260059838 | 4.90E-08 |
| ACSF3 | -0.247649221 | 0.036813686 |
| BOK | -0.440897091 | 0.008626131 |
| ZNF843 | 8.854670339 | 2.48E-06 |
| RBIS | 0.269628832 | 0.005786038 |
| PFN4 | 2.998185742 | 1.11E-07 |
| CDK5R1 | 1.282702528 | 9.27E-09 |
| RUFY1 | -0.501870251 | 9.56E-05 |
| LRRC37A3 | 3.500622074 | 2.60E-10 |

| | | |
|---|---|---|
| VSIG10 | -0.198771431 | 0.0384344 |
| MIR7-3HG | 4.713429488 | 0.000312623 |
| IRX5 | -1.145438629 | 5.21E-05 |
| METRNL | 1.387704157 | 3.18E-05 |
| FAM91A1 | -0.883085925 | 8.92E-09 |
| AL358781.1 | -0.772086222 | 5.24E-08 |
| WSB2 | -0.743277542 | 3.88E-08 |
| TCEANC | 2.634412943 | 6.70E-07 |
| PNMA1 | 0.147062921 | 0.044996865 |
| MAMSTR | 1.966310235 | 0.000878198 |
| TYMSOS | 1.192022164 | 0.001906288 |
| ANKLE2 | -1.209489999 | 5.53E-10 |
| EFCAB5 | 2.220515103 | 0.000393749 |
| TOB2P1 | 1.268200314 | 0.00348105 |
| MUC20 | 3.379515473 | 2.49E-05 |
| THAP4 | -0.342681029 | 0.000214127 |
| NFATC2IP | -0.237394581 | 0.000803046 |
| FAM89B | -0.329657724 | 0.002185672 |
| SHMT1 | -0.655812647 | 5.08E-06 |
| DPP7 | 0.563968605 | 1.51E-05 |
| SMCR8 | -0.175411487 | 0.022186423 |
| MTHFR | 2.011727912 | 1.99E-10 |
| DEAF1 | -0.451104003 | 0.011925197 |
| MTX3 | -0.72922955 | 1.92E-08 |
| TMEM80 | -0.405997303 | 0.008889922 |
| ZDHHC13 | -0.43161908 | 0.007348047 |
| SLC38A9 | -0.813513882 | 0.00011576 |
| ACER2 | 2.342683416 | 4.62E-10 |
| WDR73 | -0.603847957 | 5.40E-07 |
| POLE | -0.356357674 | 1.95E-05 |
| PHETA2 | 7.669756584 | 2.89E-06 |
| DSCAML1 | 4.325686904 | 0.001200606 |
| RHOG | -0.77146636 | 3.97E-07 |
| EPS8L2 | 0.934978544 | 0.000427058 |
| ANO6 | -0.656928771 | 4.86E-08 |
| NUDT4B | 0.353340373 | 0.000760159 |
| FAM210A | -0.457731873 | 3.45E-05 |
| TALDO1 | 0.221287944 | 0.00350252 |
| NAP1L4P1 | -0.966840962 | 0.000352595 |
| RIMKLA | 1.307081961 | 0.00232535 |
| RPS6KA3 | -0.657406135 | 1.90E-08 |
| B3GNT8 | 4.730539222 | 0.001138936 |
| PUS1 | -1.310347516 | 1.93E-08 |
| PCNPP5 | 0.288124546 | 0.022621959 |
| CHD9 | -0.942569318 | 2.86E-09 |
| GATD1 | -0.650214008 | 4.79E-06 |
| FZD8 | 0.881079283 | 2.79E-08 |
| GJD4 | 3.28890859 | 1.08E-05 |

| | | |
|---|---|---|
| CLDN22 | -1.046521447 | 0.010262294 |
| TOP3A | -0.934879612 | 1.58E-08 |
| ZBTB38 | 1.695841183 | 7.39E-13 |
| CCDC71 | 0.769678217 | 3.72E-07 |
| AC024940.2 | 1.11003763 | 4.77E-05 |
| LRRN4CL | 1.631337865 | 0.002719736 |
| HIC1 | -0.267512027 | 0.010141245 |
| PPFIA3 | 0.724119417 | 0.000103992 |
| MAGEF1 | -0.953331455 | 3.66E-08 |
| SAMD9L | -1.553022168 | 6.61E-08 |
| ZFAS1 | 0.224175478 | 0.001442442 |
| PAWR | -0.706950251 | 5.24E-08 |
| TGIF1 | -0.41222309 | 5.49E-05 |
| MIEF2 | 1.438876871 | 9.26E-07 |
| NIM1K | 2.896497049 | 2.88E-06 |
| ERICH5 | 6.097757194 | 3.32E-10 |
| OR2T8 | 1.198698502 | 0.035149312 |
| NR2C2 | -0.962209226 | 1.32E-09 |
| GPR4 | 5.756072612 | 5.07E-09 |
| CAVIN1 | -0.633739503 | 1.70E-07 |
| ARIH2 | -0.630987293 | 1.06E-07 |
| RBM44 | 1.847327117 | 0.002185848 |
| ZBTB33 | -0.461668355 | 5.10E-06 |
| IRX3 | 0.301933082 | 0.027382006 |
| SLC25A22 | -0.642556876 | 1.91E-07 |
| RABEP2 | -0.787349165 | 8.50E-05 |
| TBL1XR1 | -1.882725852 | 7.40E-11 |
| SAMD12 | -0.911346823 | 0.002355361 |
| C18orf32 | 0.292621966 | 0.000187607 |
| PIDD1 | 0.987457921 | 1.58E-07 |
| ZNF491 | 1.991401432 | 4.30E-07 |
| RPLP2 | 0.472172598 | 5.18E-07 |
| HASPIN | -0.783700436 | 7.37E-07 |
| JUN | 3.613874356 | 2.18E-07 |
| CSTF2T | -0.694870485 | 5.11E-07 |
| PGBD5 | 3.233616392 | 1.46E-07 |
| GBA | 0.865331083 | 4.90E-07 |
| IL17RA | -0.213113098 | 0.016114181 |
| PNPLA2 | 0.27059851 | 0.000537807 |
| MBOAT4 | 4.677524333 | 0.00036691 |
| AGTRAP | 0.864441797 | 4.31E-06 |
| THAP5 | 0.823281137 | 9.76E-09 |
| CD151 | 0.201747065 | 0.006387984 |
| POLR2L | -0.237614822 | 0.004213348 |
| NECTIN3 | -0.129496428 | 0.029825999 |
| ANXA2R | 0.728040179 | 0.00757091 |
| TMEM94 | -0.224928254 | 0.010100544 |
| FLII | 0.382212166 | 3.22E-06 |

| | | |
|---|---|---|
| SOX12 | -0.874824083 | 4.42E-06 |
| HNRNPA0 | -0.386093632 | 3.16E-06 |
| AC025171.1 | 0.530796629 | 0.003651532 |
| YIPF7 | 5.064184086 | 0.00010041 |
| MYOZ1 | 6.619362672 | 4.77E-05 |
| KCNJ10 | 1.734253501 | 0.008373174 |
| TENM3-AS1 | 0.999976298 | 0.000509481 |
| ZNF620 | 1.053546754 | 8.13E-05 |
| TMEM187 | 3.495750643 | 1.37E-08 |
| GRB2 | -0.617886263 | 1.19E-07 |
| UBE2N | -0.769903674 | 5.45E-09 |
| SPATA31C2 | -3.212232648 | 5.21E-09 |
| ZNF354C | -0.315838742 | 0.000785384 |
| MAMDC4 | 1.149402462 | 3.61E-06 |
| CENPBD1 | -0.26822118 | 0.020874011 |
| BET1L | -0.406013376 | 0.0001817 |
| RIC8A | -0.12056078 | 0.041798642 |
| IMP3 | -1.00541726 | 2.14E-09 |
| ASB8 | -0.225586379 | 0.011268485 |
| DMAP1 | -0.815128386 | 2.32E-07 |
| ADAMTSL1 | 0.696872632 | 0.028567991 |
| IMPDH2 | 0.550525599 | 2.66E-08 |
| MLF1 | 0.460935805 | 0.000899261 |
| NDUFAF3 | 0.698883867 | 5.45E-07 |
| C2orf69 | -0.915081075 | 1.03E-07 |
| TWF1P1 | -0.149684599 | 0.008554441 |
| TSSK6 | 1.160994698 | 0.004767011 |
| BOLA1 | 2.844870728 | 1.77E-06 |
| PDE4DIP | -0.209458573 | 0.003159729 |
| DDX10 | -0.556906393 | 1.66E-05 |
| NDUFV2 | 0.426344127 | 0.000751618 |
| DALRD3 | 0.144356395 | 0.047819455 |
| ZNF114 | 2.76849457 | 3.77E-06 |
| ZNF518B | 0.236455272 | 0.000500188 |
| LCORL | -0.338677382 | 0.000247123 |
| ZNF454 | 1.532606891 | 4.15E-05 |
| SH2B1 | -0.200770174 | 0.004647696 |
| VN1R1 | 1.352049485 | 0.00876374 |
| PLEC | -1.103436497 | 7.46E-05 |
| ZNF543 | 0.355030025 | 0.014957307 |
| GALNT11 | -0.59607055 | 1.23E-05 |
| SLITRK1 | 0.582754078 | 0.001384254 |
| WDR6 | -0.888132902 | 2.08E-09 |
| GEN1 | -1.435214492 | 1.50E-10 |
| TMEM11 | -0.648572455 | 2.08E-05 |
| ZNF354B | -0.184554196 | 0.044559981 |
| SHISA3 | -2.310787528 | 0.000675344 |
| PLEKHM3 | 0.32310586 | 0.03657717 |

| | | |
|---|---|---|
| FAM220A | 0.968561882 | 2.93E-08 |
| DNAJC22 | -0.375404504 | 0.003493768 |
| BEND3 | -2.490279115 | 2.45E-09 |
| RPS3AP5 | 0.368494054 | 1.38E-06 |
| TIMM23B-AGAP6 | -0.877548057 | 0.038403608 |
| GLDC | 0.51693335 | 0.019260731 |
| H3P16 | -0.274332898 | 0.000884749 |
| MCMDC2 | 1.727535056 | 0.000394538 |
| RPL10P16 | 0.407498291 | 2.14E-07 |
| P4HTM | 4.098993451 | 0.002920113 |
| KLHL11 | 0.274072766 | 0.013131779 |
| CTXN1 | 0.423845488 | 0.008147195 |
| CA8 | -2.905230667 | 4.12E-06 |
| EPM2AIP1 | -1.106098557 | 2.97E-09 |
| MAF | -0.994147779 | 2.27E-06 |
| CTNNBIP1 | 0.210064104 | 0.024130898 |
| ERN1 | 2.028818822 | 1.86E-08 |
| GPR35 | 3.292475649 | 7.61E-06 |
| ACTG1P1 | 5.053397577 | 3.82E-09 |
| AC092656.1 | 5.885734367 | 3.01E-05 |
| CSRNP3 | 1.904535092 | 1.87E-05 |
| PARP10 | -0.308009756 | 0.005787561 |
| SUZ12 | -0.683934087 | 4.82E-08 |
| NSUN3 | -0.371664171 | 0.002567404 |
| KCTD12 | -1.150668123 | 6.60E-07 |
| DHFR2 | 0.182751707 | 0.0216393 |
| AL450998.1 | 0.297961778 | 0.000277112 |
| RPP25 | -1.447233003 | 9.33E-10 |
| GRINA | 0.316889583 | 3.54E-05 |
| STX19 | 5.963794442 | 6.26E-09 |
| FAM219B | -1.490950215 | 2.54E-09 |
| RIIAD1 | 7.712634457 | 4.41E-06 |
| MPI | -0.324029163 | 0.000149567 |
| TRIM73 | 1.307336989 | 1.44E-05 |
| TMEM52 | 1.271671423 | 0.026298305 |
| TMEM139 | 1.36799238 | 0.000129603 |
| EFCAB13 | 0.991013693 | 4.44E-05 |
| MSC | -0.608121155 | 0.00459628 |
| APOLD1 | -0.122619047 | 0.037038667 |
| EXOSC4 | -0.643578345 | 2.57E-06 |
| TAF7 | -0.582853457 | 2.36E-08 |
| ZNF852 | -1.297691923 | 1.50E-05 |
| PFAS | -0.607650535 | 7.56E-08 |
| HYI | -0.382607605 | 0.00132708 |
| CYBC1 | -0.259000945 | 0.008260716 |
| ZNF552 | -0.736547893 | 0.000304835 |
| GAK | -0.295323936 | 0.000704146 |
| TUFM | 0.257294276 | 2.07E-05 |

| | | |
|---|---|---|
| ERICH3 | 9.423140444 | 4.99E-09 |
| RMI1 | -0.155062061 | 0.017227221 |
| CTC1 | 0.61703646 | 3.35E-05 |
| FBXO34 | 0.202643798 | 0.008737561 |
| LINC00324 | 3.191732359 | 1.06E-07 |
| MRFAP1L1 | -0.811943838 | 1.42E-08 |
| SNX18 | -0.616659265 | 1.21E-06 |
| EXD1 | 5.619256018 | 1.77E-05 |
| AURKB | -0.906543886 | 1.24E-08 |
| MRFAP1 | -0.345305873 | 9.83E-07 |
| C3orf38 | -0.405155028 | 4.43E-05 |
| TMEM107 | 1.444174529 | 9.32E-07 |
| RRS1 | -2.092649486 | 5.88E-11 |
| EXOC3L1 | 3.060631032 | 1.17E-05 |
| TRIML2 | 1.119115243 | 0.01507284 |
| RCC2 | -0.649853178 | 1.07E-08 |
| IGSF22 | 3.253367615 | 0.000248072 |
| C9orf50 | 6.77593355 | 3.11E-05 |
| ZFP42 | 1.585074312 | 3.10E-10 |
| AC020907.1 | 0.413927077 | 0.01896659 |
| FAM133A | -0.49272273 | 0.005599864 |
| DPM3 | 0.472389472 | 0.001727757 |
| PER1 | 3.070727693 | 7.39E-13 |
| H3P47 | -0.256280577 | 0.000839375 |
| TMTC2 | -0.416656802 | 1.06E-05 |
| HES7 | 2.509189705 | 2.87E-06 |
| FARSA | -0.174306337 | 0.004198384 |
| SPTY2D1 | 0.274990924 | 0.000176524 |
| SAMD4B | -0.316625364 | 3.13E-05 |
| MTUS2-AS1 | 2.753561453 | 2.98E-06 |
| ALOXE3 | 3.608297793 | 0.027786367 |
| EDC3 | -1.377347158 | 9.65E-10 |
| PXT1 | 5.355378757 | 7.78E-05 |
| GGN | 0.950796818 | 0.008672975 |
| ZNF664 | -0.385200681 | 8.01E-07 |
| CALR | 0.169300384 | 7.51E-05 |
| MAGED1 | 0.357400837 | 3.04E-05 |
| GVQW3 | 1.034124933 | 2.11E-05 |
| LDLRAD3 | -0.86037286 | 1.81E-08 |
| SMCO3 | 2.687908019 | 7.86E-05 |
| MEIS3P1 | 1.122634188 | 4.66E-05 |
| PTPN11 | -0.593729053 | 8.39E-09 |
| FAM156B | -0.89353107 | 1.67E-08 |
| WSCD1 | 1.058462196 | 0.000372451 |
| CLK3 | 0.646126693 | 6.50E-07 |
| ELMOD2 | -0.50040066 | 0.000168078 |
| EGR3 | 2.456822372 | 4.10E-12 |
| CATSPERE | 2.890453056 | 0.000208767 |

| | | |
|---|---|---|
| LINC00174 | 0.441922596 | 0.045250499 |
| GEMIN4 | -0.969939764 | 6.73E-09 |
| FJX1 | -1.078095866 | 4.97E-07 |
| KLHL28 | 1.36100132 | 9.59E-09 |
| ZBTB18 | -0.270985959 | 0.000984057 |
| EEF1A1P27 | 2.945831408 | 0.024591232 |
| C14orf28 | 3.631733358 | 2.00E-09 |
| ALOX12B | 2.368011037 | 0.008223233 |
| EIF3J-DT | 0.938868293 | 9.89E-05 |
| LBX2 | 0.991093527 | 0.00020083 |
| DNHD1 | 3.088652495 | 6.67E-11 |
| SLITRK4 | -1.220413421 | 1.79E-05 |
| HTR1D | 2.665466809 | 0.000379471 |
| RNF151 | 4.471404808 | 1.29E-06 |
| CIITA | -1.916233146 | 4.52E-11 |
| ZFPM1 | 0.62795583 | 0.000583194 |
| PLD6 | -0.335824995 | 0.008802873 |
| CDC42EP4 | -0.429102249 | 3.16E-05 |
| DGKZP1 | -0.320081148 | 0.000632018 |
| ZBTB42 | -0.46191323 | 0.041442901 |
| LACC1 | -1.83215263 | 3.39E-06 |
| MAF1 | 0.395596256 | 6.59E-06 |
| WDR97 | 2.241952362 | 3.76E-07 |
| PCED1B | 1.405798248 | 0.000346634 |
| PIPOX | 5.037890019 | 4.74E-07 |
| ATP8B5P | 1.155225162 | 8.36E-05 |
| OR7E22P | 6.867151347 | 2.94E-05 |
| MRGPRX4 | -0.800342607 | 0.009588011 |
| PCBP1-AS1 | -0.714310547 | 2.12E-07 |
| MYADM | -0.512668765 | 3.22E-06 |
| MROH1 | -1.571290887 | 1.26E-08 |
| SERTAD2 | -0.232213529 | 0.000258077 |
| AKAP5 | -0.770703448 | 0.003619991 |
| NKPD1 | 3.822475837 | 8.10E-07 |
| GIPC3 | 1.670279996 | 0.002199559 |
| RNF227 | 1.37971102 | 3.07E-07 |
| NLRP11 | -2.398822889 | 6.75E-07 |
| TIGD5 | -0.560883464 | 0.031474204 |
| PDXDC1 | -0.413057184 | 2.03E-05 |
| C1orf194 | 2.466185847 | 8.74E-05 |
| ZNF154 | 1.476656412 | 0.000168739 |
| R3HDM2 | 0.503836354 | 3.41E-05 |
| NRXN1 | -1.301436904 | 0.014824709 |
| SEPHS2 | -0.131937928 | 0.010227769 |
| BBS10 | 0.771919278 | 7.34E-08 |
| FIZ1 | -0.55063722 | 7.61E-05 |
| PUF60 | -0.154733794 | 0.010001416 |
| SSC5D | 2.216273907 | 3.43E-07 |

| | | |
|---|---|---|
| DCTPP1 | -0.526821511 | 1.06E-05 |
| NAIPP2 | -0.510847336 | 0.046875857 |
| TSHZ1 | -0.655954088 | 2.04E-06 |
| PSTK | -0.819524004 | 0.000498821 |
| SOCS4 | 0.361792523 | 1.51E-05 |
| ZADH2 | -1.181235517 | 7.66E-09 |
| AC093909.1 | -0.284513012 | 0.006845481 |
| ZNF48 | 1.330810409 | 5.86E-08 |
| FAM71E2 | 4.374942667 | 3.68E-06 |
| C3orf80 | -1.606969058 | 0.000364202 |
| ANKRD18A | -1.257402478 | 1.24E-06 |
| TMEM86B | 2.000395916 | 6.28E-07 |
| SEPTIN1 | 2.730704917 | 3.53E-06 |
| TRNAU1AP | 1.764552184 | 5.16E-09 |
| EXOC3 | -0.368215108 | 8.81E-05 |
| AC017081.1 | 2.121638873 | 6.48E-05 |
| TDRD6 | 0.896652428 | 0.045567948 |
| ACTA2-AS1 | 1.869073549 | 1.49E-05 |
| MED14 | -0.657118033 | 3.51E-07 |
| FAHD1 | -1.122849186 | 2.92E-07 |
| TDRP | 2.068821468 | 0.000357899 |
| RCC1 | 0.416680626 | 1.03E-06 |
| F2 | 3.842949957 | 3.78E-05 |
| FO393411.1 | 0.291414814 | 0.00058866 |
| HERC2P3 | -0.547993406 | 0.00015712 |
| ZNRF2 | -1.335701733 | 5.37E-06 |
| RRH | 2.426715847 | 0.003429314 |
| ZNF816 | 1.597139413 | 4.24E-10 |
| FGD6 | -0.613610826 | 1.39E-05 |
| LINC01869 | 5.028110894 | 0.000241478 |
| AL049610.1 | 1.017403107 | 0.016985239 |
| OAZ2 | -0.373981311 | 1.64E-05 |
| CCDC43 | -0.753601483 | 2.05E-07 |
| MEIOC | 1.545980219 | 6.62E-06 |
| FZD2 | -0.859368871 | 3.17E-05 |
| TIGD2 | 0.693252144 | 1.85E-05 |
| HCLS1 | 5.65795885 | 0.001205122 |
| ZNF609 | -1.283797565 | 1.64E-10 |
| PAK2 | -0.624479382 | 6.21E-08 |
| CCDC66 | -0.584020851 | 1.42E-05 |
| EMC3-AS1 | 0.899910181 | 6.00E-06 |
| MCFD2 | -0.265753209 | 3.09E-05 |
| HARBI1 | 0.591454884 | 5.02E-05 |
| C11orf71 | 1.988055389 | 1.20E-07 |
| TPRXL | 5.033385529 | 0.000113742 |
| SERTM1 | 0.883894405 | 0.007065079 |
| GAS1 | -0.956642814 | 0.000188092 |
| ZNF571 | 1.304770653 | 1.99E-06 |

| | | |
|---|---|---|
| GLIPR1L2 | 2.946993894 | 0.024591232 |
| MIGA1 | -0.325252524 | 9.56E-05 |
| NRIP1 | -0.296104216 | 8.81E-05 |
| ZSCAN4 | 4.422077965 | 0.000917626 |
| RNF182 | -0.835359026 | 6.35E-07 |
| C9orf139 | 1.230153539 | 0.023748102 |
| H2AC6 | 2.147136523 | 4.56E-09 |
| SRP9P1 | -0.181472486 | 0.007752162 |
| SKIDA1 | -0.607724094 | 0.013544073 |
| H2BC4 | 1.998593358 | 2.94E-06 |
| MB21D2 | -1.014637365 | 2.79E-06 |
| SSTR2 | 1.104454463 | 0.001968318 |
| ZNF594 | 0.461340471 | 4.79E-05 |
| PCGF5 | -0.76952084 | 2.92E-07 |
| SLC47A2 | 2.771648046 | 6.94E-06 |
| MAB21L1 | -1.136518011 | 7.32E-05 |
| YOD1 | -0.569270262 | 5.22E-07 |
| TMEM64 | -0.599867363 | 1.29E-06 |
| LINC02363 | 4.167446888 | 0.001839068 |
| CHRM4 | 2.191217737 | 0.00017846 |
| S1PR5 | 0.834120594 | 0.001548691 |
| SMG1P3 | -0.271531234 | 0.001162828 |
| GPR157 | -0.599114962 | 0.00021006 |
| WDFY3-AS2 | 3.552996809 | 3.41E-08 |
| SLC36A4 | -0.55333771 | 3.43E-05 |
| ZDHHC20 | -0.580620492 | 1.69E-07 |
| ZFP3 | 4.315826903 | 0.007289158 |
| ARSJ | -1.978156537 | 2.59E-10 |
| HOXC9 | -1.20921829 | 3.09E-05 |
| HOXC10 | -0.980640526 | 4.74E-08 |
| PSMG4 | 0.722632679 | 6.12E-05 |
| ZNF443 | 0.495070817 | 0.000978865 |
| PDIA3P1 | 0.312224575 | 5.57E-06 |
| GREM2 | 4.008139577 | 3.60E-06 |
| SSR4 | 0.523564231 | 5.16E-05 |
| CAPS2 | 3.073124327 | 4.23E-06 |
| CUEDC1 | -1.258996884 | 8.12E-09 |
| D2HGDH | 0.342923952 | 0.013337493 |
| OXTR | -1.456770159 | 4.86E-06 |
| CMTR2 | -0.29803542 | 0.000121377 |
| FAM83H | 1.45886953 | 0.011286272 |
| GPR62 | 1.764938611 | 0.002189164 |
| ZNF572 | 2.531177096 | 1.49E-08 |
| ST20 | 0.384637847 | 0.028104123 |
| PITPNB | -1.795410016 | 1.50E-11 |
| TCEAL8 | 0.387937742 | 0.0002957 |
| LRRC57 | -0.471969123 | 3.35E-05 |
| MRPL14 | -1.026637946 | 1.24E-07 |

| | | |
|---|---|---|
| GPR137C | 1.45244148 | 0.000463668 |
| BBS12 | 2.197481873 | 8.90E-10 |
| ZFP82 | 0.373585886 | 0.000303377 |
| LSMEM1 | 1.042447198 | 0.011361904 |
| NQO1 | -0.762807955 | 8.63E-10 |
| AEN | -0.364901072 | 7.73E-06 |
| FKRP | -0.644663856 | 1.47E-06 |
| SLC25A42 | 1.703531126 | 7.53E-08 |
| FCRL6 | 7.00011059 | 1.83E-05 |
| METTL23 | 0.325623987 | 0.002913612 |
| HIGD1A | -0.152035647 | 0.046648692 |
| MAPK15 | 4.243016506 | 7.97E-06 |
| EHMT1 | -1.056683041 | 1.97E-08 |
| AC105219.1 | 0.598190599 | 0.013123422 |
| ENTR1P2 | 0.824068902 | 0.000916726 |
| F2R | -0.576835507 | 3.85E-07 |
| ZNF707 | -0.466956076 | 0.004039172 |
| NPM1 | 0.51327656 | 7.37E-09 |
| DHTKD1 | -0.170693963 | 0.021345135 |
| H2AW | 0.820031791 | 0.000415452 |
| ZNF746 | -0.391739527 | 0.000117077 |
| TLCD5 | 1.352924488 | 3.08E-09 |
| ZNF322 | -0.438019067 | 0.000108831 |
| LRRC75A | 0.491635737 | 2.26E-06 |
| CFAP65 | 4.575486402 | 0.001581267 |
| DDX60L | -0.71916623 | 0.002184131 |
| SYNE4 | 4.922969203 | 1.06E-06 |
| OGFOD3 | -0.677108933 | 4.33E-05 |
| AATK | 1.320210762 | 0.002615819 |
| DDN | 3.796094298 | 1.41E-10 |
| SAGE1 | -0.364953304 | 0.003226253 |
| SOX2 | -1.764261164 | 3.54E-06 |
| ZNF678 | -0.405310338 | 0.000172273 |
| RAP2B | -0.184912479 | 0.01615815 |
| ZBTB2 | 0.255715642 | 0.014146973 |
| ACBD4 | 1.472199693 | 2.40E-05 |
| RPL24P4 | 0.257127699 | 0.002199131 |
| FANCB | -1.693447906 | 2.86E-09 |
| SETD2 | -0.526080285 | 2.44E-07 |
| MRPS23 | -0.151583747 | 0.046368619 |
| GPR135 | -0.609931168 | 0.007901745 |
| SLX1B | -0.459268787 | 0.017084028 |
| ZFP41 | 1.148459982 | 5.48E-08 |
| PHLDA2 | -0.440683276 | 0.000954424 |
| PLAG1 | 1.056068862 | 2.05E-08 |
| YIPF6 | -0.756262545 | 3.34E-08 |
| ZBTB20 | 1.388207601 | 0.000416303 |
| DIPK2A | -0.44725968 | 7.08E-05 |

| | | |
|---|---|---|
| C5orf30 | -0.434455497 | 3.55E-05 |
| GPR3 | 0.716531083 | 0.000170001 |
| SIAH2 | -0.96913526 | 9.53E-08 |
| LINC00471 | 2.236024994 | 6.87E-05 |
| LSM10 | -0.944585554 | 3.24E-07 |
| KCTD9P2 | -0.725763518 | 0.000181449 |
| RELL1 | -0.369717633 | 0.041850628 |
| RFX7 | -0.595559058 | 9.99E-07 |
| SLC35C1 | -0.908502926 | 4.23E-08 |
| RNF41 | 0.638439868 | 4.94E-08 |
| SLC2A4 | 3.105542182 | 5.57E-06 |
| IBA57 | -1.691067752 | 2.10E-08 |
| C5orf24 | -0.782433316 | 3.16E-08 |
| COA4 | -0.572795962 | 7.37E-07 |
| PRKAG1 | 0.399889628 | 7.40E-06 |
| GINS3 | -1.374983101 | 6.38E-09 |
| SNRPE | -0.305053733 | 0.003254866 |
| RTKN2 | -0.946587668 | 4.65E-05 |
| PNMA8A | -0.932940823 | 6.85E-09 |
| MGAT4C | -2.525175909 | 0.00132732 |
| IDH2 | 0.382419354 | 0.008956541 |
| TMEM259 | -0.212987124 | 0.002973858 |
| GET1 | 0.224220208 | 0.02386408 |
| TNRC18 | -0.625253224 | 9.29E-07 |
| DEXI | -0.664481084 | 0.000173292 |
| NOP10 | -0.292180027 | 0.004801321 |
| TDRKH | 1.067857055 | 3.17E-08 |
| ZNF708 | -1.33457184 | 2.44E-08 |
| IST1 | -0.145963549 | 0.020039746 |
| ERCC6L2 | 0.309839987 | 0.00029815 |
| CREB3L2 | 0.10808574 | 0.036991681 |
| TP53TG1 | 1.574912121 | 1.48E-05 |
| TSEN54 | -0.603321956 | 3.56E-05 |
| RGMA | 1.868201595 | 1.23E-05 |
| UBA7 | 1.928147618 | 3.65E-06 |
| MRPS16 | -0.475615428 | 1.10E-05 |
| RAD51B | 1.209293585 | 0.010152732 |
| ARL6IP4 | 0.299025296 | 0.000946751 |
| EXT1 | -1.65266463 | 1.41E-10 |
| SHMT2 | 0.27890785 | 2.25E-05 |
| MOB2 | -1.161023066 | 1.79E-05 |
| ATP6AP2 | 0.216364028 | 0.000931117 |
| CYB5D1 | -0.94787175 | 6.10E-08 |
| FAM153B | 1.860120782 | 0.000617452 |
| BACE2 | 0.178793298 | 0.006303297 |
| SYNM | -1.665500625 | 2.42E-11 |
| FIGN | -2.237814932 | 1.71E-09 |
| B4GALNT4 | 1.072384762 | 8.40E-06 |

| | | |
|---|---|---|
| C8orf33 | -0.716112891 | 3.87E-08 |
| DCAF4L1 | 1.006919665 | 0.000314269 |
| SPACA6 | -1.060442138 | 0.001088629 |
| ZSCAN22 | 0.757420153 | 2.71E-06 |
| KCNJ14 | -1.644155841 | 8.67E-06 |
| C1S | 4.38827953 | 8.58E-09 |
| GLTPD2 | 3.026421951 | 3.43E-05 |
| KBTBD3 | 0.96300429 | 0.000203548 |
| FAM87A | 1.660454422 | 0.016313932 |
| PLCXD1 | -1.166182208 | 3.27E-06 |
| RPL27AP5 | 0.74084503 | 0.013539816 |
| DNM1P46 | 5.615012096 | 2.58E-10 |
| TRAPPC6B | -1.041223704 | 1.33E-08 |
| PGBD4 | 0.607227267 | 1.41E-05 |
| NPLOC4 | -0.507561049 | 7.69E-07 |
| KCNK4 | 5.061416767 | 9.96E-05 |
| TEX19 | 2.363609596 | 6.26E-07 |
| TSHZ2 | 1.119801296 | 0.00379741 |
| CAPN12 | 0.34918074 | 0.000403061 |
| EXOC7 | -0.321292841 | 3.00E-05 |
| KPNA2 | -1.107221683 | 2.26E-10 |
| BGN | 0.555246278 | 0.000368656 |
| CEP97 | -0.460622039 | 5.76E-05 |
| FES | 3.672351655 | 4.01E-07 |
| GLRX5 | -0.496792862 | 2.90E-05 |
| FAM104B | 0.381558322 | 0.01732533 |
| MXRA7 | 0.208174198 | 0.006595018 |
| LIMK2 | -0.272005631 | 0.013262563 |
| MFSD5 | -0.710741355 | 1.16E-05 |
| RWDD4 | -0.33227862 | 0.003355284 |
| SATB1 | 0.453582496 | 0.004634033 |
| CSF1R | 2.684491754 | 1.34E-05 |
| EPHB3 | -1.070830565 | 3.92E-05 |
| ACTL10 | 0.953588594 | 0.001648806 |
| LINC00334 | 4.673905018 | 0.000349487 |
| TRAK1 | -0.996290669 | 3.80E-07 |
| TSPAN10 | 0.99230546 | 0.039971656 |
| PLCB1 | -0.674465059 | 0.000129675 |
| SKA2 | -0.565705995 | 1.27E-06 |
| CCNYL2 | -0.58026684 | 0.049516951 |
| LINC01006 | 0.785778728 | 0.043053287 |
| TTC3 | -0.255419294 | 0.001173125 |
| PPP1R27 | 2.525980178 | 0.00159531 |
| BRICD5 | -0.823437036 | 0.001379106 |
| IGIP | 1.607004677 | 5.94E-08 |
| TSKU | -0.50580957 | 8.92E-05 |
| ANXA2 | 0.15743763 | 0.006598313 |
| SEPHS1P1 | -1.920307636 | 0.00262908 |

| | | |
|---|---|---|
| HOXB4 | -2.485201897 | 7.98E-10 |
| PAQR7 | -0.451921524 | 0.001981284 |
| MAFA | 0.941654394 | 0.038236617 |
| RPS17 | 0.286082759 | 0.003045256 |
| HCAR2 | 3.55921803 | 8.80E-05 |
| CCDC87 | 3.155374954 | 0.021881779 |
| C1orf116 | 3.252046349 | 4.26E-06 |
| TMEM198B | 1.519524568 | 2.76E-09 |
| CRIP2 | 0.463022128 | 1.58E-05 |
| DDX28 | 0.332701117 | 0.001536686 |
| ACBD3 | 0.409024133 | 6.37E-06 |
| C16orf72 | -0.312721756 | 1.79E-05 |
| RRP7BP | -0.622462807 | 0.002030146 |
| VMO1 | 2.693681174 | 0.000147873 |
| PRKCZ-AS1 | 1.169901823 | 0.003458792 |
| RPL35A | 0.320673227 | 2.51E-05 |
| ZNF721 | 0.438084761 | 8.48E-06 |
| TSPEAR-AS2 | 2.189428644 | 1.43E-07 |
| C11orf54 | -0.293259631 | 0.013076931 |
| CEP63 | 0.213004105 | 0.022802945 |
| SRPRA | -0.619804282 | 1.05E-08 |
| EWSR1 | -0.44062813 | 6.30E-07 |
| HMGN4 | -0.77287649 | 9.78E-08 |
| SPATA13 | -0.586641133 | 2.38E-07 |
| GJC1 | -2.121296484 | 7.82E-12 |
| CNOT10 | -0.30076295 | 0.00174336 |
| MTA1 | 0.181967633 | 0.032082273 |
| CADM1 | -0.378133293 | 3.84E-05 |
| ZNF320 | -0.320024883 | 0.000252144 |
| C12orf60 | 2.611312671 | 1.42E-09 |
| PYCR1 | 0.449756121 | 7.77E-06 |
| NAA38 | 0.379944117 | 9.04E-05 |
| SLC8A1 | -1.11235638 | 2.94E-06 |
| SLC25A21 | -1.171260319 | 0.00031092 |
| ABAT | 1.825195803 | 4.16E-06 |
| SLC25A10 | 0.296762426 | 0.002209854 |
| RGPD6 | -0.236322804 | 0.007875631 |
| FAM133CP | 0.616203628 | 0.008417712 |
| WBP2NL | 1.784206504 | 0.001642827 |
| NKX2-5 | -0.696899892 | 4.05E-05 |
| AFMID | 0.401591412 | 0.002906361 |
| GAS6 | 0.427138305 | 0.012176748 |
| NEB | 2.752164783 | 1.61E-07 |
| BEGAIN | 0.890091514 | 0.017928909 |
| ARHGEF37 | 3.774675284 | 5.27E-11 |
| PTGDR2 | 3.642421431 | 2.45E-09 |
| CEP57L1 | -0.785019924 | 0.000155042 |
| GPR19 | 1.891284607 | 2.57E-07 |

| | | |
|---|---|---|
| RABIF | -0.526948813 | 5.40E-05 |
| TMEM119 | 6.494837556 | 2.71E-06 |
| BX322650.1 | -0.682932233 | 2.48E-05 |
| HSP90AB3P | 0.362361147 | 1.07E-06 |
| RUVBL2 | 0.253942608 | 0.000325495 |
| GDPGP1 | 0.344031913 | 0.040422569 |
| WT1-AS | 0.927713748 | 0.000661998 |
| PRR36 | 1.98225237 | 0.000835147 |
| LINC01547 | 0.839620224 | 0.000835091 |
| PTTG1IP | -0.267765788 | 0.000106993 |
| DDX41 | -0.121607288 | 0.041850628 |
| ABHD16B | 1.538472258 | 2.61E-05 |
| PLGLB1 | 0.720225085 | 0.004636204 |
| DAZAP2 | -0.640890291 | 2.44E-06 |
| CCBE1 | -0.645116082 | 0.002650333 |
| RPSAP19 | 0.358139881 | 6.06E-06 |
| MAGEA2B | -0.608116671 | 5.68E-08 |
| TMEM121B | 2.282321683 | 0.002466941 |
| ZNF623 | 0.362924935 | 0.000138803 |
| TUBB | -0.276558298 | 0.003437514 |
| CCDC125 | -0.712260349 | 0.003339742 |
| BCOR | -1.762501807 | 4.43E-11 |
| JRKL | 1.091764823 | 5.78E-09 |
| SYNDIG1L | 8.015700405 | 6.83E-07 |
| FHL3 | 0.754445174 | 5.28E-06 |
| PMCH | -1.052201936 | 0.003994064 |
| C19orf71 | -1.003721611 | 6.92E-06 |
| RIPK4 | 0.672345491 | 3.92E-06 |
| LRIT3 | 2.676733014 | 6.60E-05 |
| TRIM61 | 3.753856898 | 2.35E-05 |
| OR7E38P | 0.522560224 | 0.005840833 |
| AC138932.1 | -0.243679241 | 0.0022345 |
| SH2D7 | 3.184768726 | 0.000502054 |
| MX2 | -0.207409752 | 0.004894978 |
| EP400 | -0.70506712 | 1.08E-07 |
| MEX3B | 1.492273821 | 1.70E-07 |
| PI4KAP2 | -0.562989764 | 0.006344506 |
| TENT5C | 2.354614593 | 3.05E-08 |
| UTP11 | -0.964484438 | 1.32E-08 |
| PSMG1 | -0.302407946 | 0.002447437 |
| PRR14L | -0.4087011 | 6.85E-06 |
| SERHL2 | -1.488795223 | 4.24E-05 |
| RNF5 | 1.178445987 | 0.010546815 |
| SETD3 | -0.303513838 | 0.003204169 |
| ZNRF3 | -0.517736505 | 0.000122551 |
| H3C13 | 3.526453632 | 1.25E-06 |
| SMG1P5 | 0.335488799 | 0.001690438 |
| FAM167B | 1.559208176 | 0.015059031 |

| | | |
|---|---|---|
| MRPL54 | -0.414323704 | 0.015871179 |
| ZNF438 | 0.815706567 | 0.00074783 |
| HMCES | -0.466188743 | 1.35E-05 |
| DGCR6 | 1.981601576 | 0.001120527 |
| ZNF530 | -0.306614881 | 0.010448121 |
| KLHL25 | -0.41201496 | 0.004395693 |
| TRMT12 | -1.401790344 | 9.36E-09 |
| GUSBP1 | 0.244022337 | 0.037655944 |
| PSG9 | 4.642535126 | 0.000785426 |
| ALYREF | -0.384312805 | 0.000215246 |
| UPP1 | 0.455770989 | 0.001256262 |
| TRIM52 | 0.188371538 | 0.005702542 |
| LHFPL6 | -1.230320018 | 1.74E-06 |
| CMTM4 | -0.87471615 | 4.20E-08 |
| TMEM50A | -0.376553745 | 4.56E-05 |
| TBK1 | -1.609597777 | 1.33E-09 |
| CBX6 | -1.579526625 | 7.05E-11 |
| TBL3 | -0.451569386 | 1.23E-05 |
| KREMEN1 | -0.719228792 | 0.001884669 |
| TRAIP | -0.77390153 | 9.36E-06 |
| AIFM3 | 2.105755196 | 0.005737674 |
| KCTD16 | 6.223347593 | 1.89E-05 |
| B3GALT5 | -1.570581778 | 2.74E-06 |
| ZNF703 | 0.531647711 | 0.002868478 |
| TUBA8 | 1.097361452 | 0.0113936 |
| LIN9 | -1.352506265 | 2.14E-07 |
| BTBD9 | -0.716452202 | 3.07E-05 |
| ANKRD45 | 1.920138612 | 2.09E-05 |
| GPR39 | -0.80959307 | 2.22E-06 |
| ZNF730 | 1.311314885 | 2.99E-07 |
| KIRREL1 | -0.284806316 | 7.91E-06 |
| IQGAP3 | -0.331730306 | 8.99E-05 |
| TOB2 | -1.925255062 | 4.66E-11 |
| SCN5A | 1.750057796 | 0.001906327 |
| TTC32 | 0.53069651 | 0.004603654 |
| SDR42E2 | 3.756575169 | 0.004493327 |
| DUSP5P1 | 1.727709879 | 0.004078619 |
| HTR7P1 | 1.900344911 | 2.36E-08 |
| PRKX | -2.833154949 | 1.88E-10 |
| KMT5A | -0.334028407 | 8.00E-05 |
| SMTN | -0.799247554 | 2.24E-08 |
| NPW | 2.871180921 | 2.22E-06 |
| PP2D1 | 1.5352845 | 0.002767689 |
| COA3 | -0.525390121 | 5.16E-06 |
| ST6GALNAC3 | -0.953681156 | 2.71E-06 |
| PTP4A2 | -0.638669326 | 4.47E-08 |
| ACTG1 | 0.090609867 | 0.000945284 |
| DENND5A | -0.281788419 | 0.005865479 |

| | | |
|---|---|---|
| CTAG1B | 0.467089656 | 0.00049553 |
| DIABLO | -0.196899141 | 0.008023984 |
| VPS33B | 0.336761565 | 0.006659848 |
| ADAP2 | 6.170416465 | 0.000524881 |
| FAM120C | 0.766308398 | 9.02E-07 |
| BRD7P2 | -0.606241513 | 7.11E-06 |
| TRIML1 | 4.714749058 | 0.000438271 |
| EIF3C | 0.179541092 | 0.000119687 |
| NIPSNAP1 | 0.726809132 | 1.60E-07 |
| LRTOMT | 0.199173279 | 0.022339148 |
| NR2C2AP | -0.436864349 | 3.51E-05 |
| SCFD2 | -0.684670924 | 3.45E-06 |
| KCNJ12 | 0.571392978 | 0.025771176 |
| AC026410.1 | 1.504561987 | 0.014027685 |
| PGP | 0.156148596 | 0.042339897 |
| C22orf46 | -0.279576678 | 0.003340193 |
| SNRNP35 | -0.791268989 | 3.22E-06 |
| IRAK1 | -0.286824824 | 2.56E-05 |
| C11orf72 | 3.611849522 | 9.88E-05 |
| PCDH9 | 0.709928952 | 4.22E-07 |
| ACOT1 | -0.679207161 | 0.009348298 |
| OAF | 0.231712013 | 0.000180089 |
| ALDH1A3 | -0.822424407 | 1.55E-10 |
| CDR1 | 3.654008537 | 1.25E-08 |
| POU6F1 | 1.23339856 | 0.000455257 |
| TM2D3 | 0.24954224 | 0.003342817 |
| TSSC4 | -0.921034704 | 1.84E-07 |
| PRKD1 | -0.740860632 | 0.000608634 |
| ZDHHC23 | -1.680360154 | 2.21E-08 |
| RPL23AP82 | 0.787776657 | 0.001983724 |
| EFNA5 | -1.345106675 | 5.30E-07 |
| H1-5 | 2.99572887 | 0.001249467 |
| SPATA32 | 5.080021382 | 3.05E-09 |
| PKP3 | 0.504430425 | 0.004787032 |
| CSF1 | -0.714991344 | 0.000333116 |
| PLA2G6 | 1.196291431 | 0.000133044 |
| A3GALT2 | 3.69883314 | 5.17E-05 |
| SS18L1 | -0.724456045 | 2.59E-07 |
| TOP1MT | 0.390569318 | 0.001032585 |
| COPB2 | 0.127803213 | 0.029035356 |
| LRRC19 | 3.775436848 | 3.92E-06 |
| THAP7 | -0.308297531 | 0.005698297 |
| AP001062.1 | -0.936892729 | 0.000759363 |
| KNTC1 | -0.358269543 | 0.000115437 |
| CCR10 | 0.710625803 | 0.012279647 |
| NCMAP | 7.843595027 | 1.28E-06 |
| WDR27 | 0.510475833 | 6.18E-05 |
| TXNRD2 | -0.609576716 | 0.000291347 |

| | | |
|---|---|---|
| POU3F2 | -0.974773417 | 2.01E-07 |
| PTP4A3 | 3.857003195 | 8.21E-05 |
| NEU1 | 0.935857425 | 3.96E-05 |
| TMEM255B | 1.811094966 | 4.35E-06 |
| HDDC3 | 0.988353892 | 3.29E-06 |
| ZFP1 | 0.554217461 | 2.12E-05 |
| SOCS3 | 0.24835386 | 0.021645828 |
| SLITRK6 | -1.142784337 | 2.86E-09 |
| LPAR5 | 4.678789143 | 8.56E-10 |
| XPOT | -0.368272606 | 2.73E-06 |
| KRBA2 | 5.744577122 | 6.32E-05 |
| ZNF93 | 0.379496064 | 0.000316915 |
| SEPTIN9 | -0.118486109 | 0.021994252 |
| ODF4 | 9.233944088 | 2.48E-08 |
| FOXD4L4 | 3.184212815 | 0.00030057 |
| CDCA2 | -0.66025373 | 8.19E-07 |
| AMER1 | -1.076121451 | 1.85E-08 |
| ZBTB40 | -1.492585777 | 3.09E-10 |
| H2BC21 | 1.821096971 | 1.12E-07 |
| CLDN6 | 2.385106424 | 0.000271062 |
| EIF4ENIF1 | -0.553254236 | 0.00033867 |
| SERINC4 | 1.792353504 | 9.45E-08 |
| ATL3 | -1.452329194 | 5.61E-10 |
| MGAT4EP | 7.086542092 | 1.33E-05 |
| UBE2G2 | -0.713118244 | 1.65E-08 |
| OSBP2 | -0.369803953 | 0.042772196 |
| ZBTB7C | 4.836747197 | 3.18E-08 |
| PRR16 | -2.010622696 | 3.53E-05 |
| TMED9 | 0.113776162 | 0.029327039 |
| TMEM186 | -0.198260873 | 0.046183533 |
| RBM33 | -0.970528042 | 7.13E-09 |
| BTBD6 | -0.303656545 | 0.003482777 |
| RBM43 | 0.966648035 | 2.14E-05 |
| SUMO3 | -0.302282416 | 0.000255736 |
| JAG2 | 1.573949343 | 0.003226779 |
| FMNL1 | 2.716610974 | 2.87E-10 |
| NUTM2A | 3.477463039 | 9.77E-07 |
| WT1 | -2.356571219 | 8.96E-09 |
| ZFP90 | 0.80866029 | 2.64E-07 |
| FAM227A | 1.755633243 | 1.52E-06 |
| CHRM5 | 4.279320966 | 2.67E-07 |
| SORCS2 | 0.942281314 | 0.004668781 |
| TMEM106A | 1.409030588 | 8.64E-06 |
| SIVA1 | -0.231675561 | 0.023406425 |
| BRI3BP | -1.196885929 | 3.72E-10 |
| AP3M1 | -1.034728281 | 5.45E-09 |
| F8 | 3.175714482 | 9.32E-08 |
| CA13 | -1.545075121 | 2.46E-06 |

| | | |
|---|---|---|
| UBOX5 | 0.935971438 | 0.000143857 |
| MAFF | 0.411906556 | 0.000462624 |
| BRF1 | -0.534631129 | 0.000466193 |
| ANKS1B | -0.720077434 | 0.001156515 |
| NELFA | -1.312604787 | 1.60E-08 |
| C5orf47 | 8.232933498 | 3.14E-07 |
| AC000068.1 | 1.441230349 | 0.001429365 |
| FLRT2 | -2.06099339 | 3.43E-07 |
| INTS5 | -0.471065455 | 3.33E-05 |
| RPS27L | 0.417743265 | 8.45E-05 |
| MANEAL | -1.063032771 | 2.52E-08 |
| FAM43A | 1.814725736 | 1.63E-08 |
| HSF1 | -0.214333725 | 0.00198039 |
| H2BC13 | 3.394592779 | 4.27E-05 |
| INPP5J | 1.760599331 | 2.87E-06 |
| LRRC37B | -0.882950539 | 0.000488393 |
| DDX51 | -1.244059702 | 6.84E-09 |
| NOMO2 | 0.17935031 | 0.014842858 |
| LINC00482 | 5.917280698 | 8.15E-06 |
| SIGIRR | 0.439230427 | 0.013924026 |
| NRBP2 | 0.656779833 | 0.000162115 |
| ZNF445 | -0.538869078 | 2.03E-06 |
| PGBD2 | 0.420545323 | 3.30E-05 |
| TCEAL9 | 0.425465539 | 0.00053657 |
| GP1BA | -1.013823166 | 0.007361541 |
| PRPF39 | -0.666066534 | 8.89E-07 |
| MAGEA11 | 0.253548673 | 0.013905022 |
| PPIL6 | 1.660456809 | 0.000875596 |
| ZNF74 | 0.46589897 | 0.000122465 |
| KIAA0825 | 2.689919167 | 1.66E-06 |
| UBALD2 | -0.531617324 | 7.93E-06 |
| CDNF | 4.299155089 | 4.92E-06 |
| ZBTB37 | -0.254898288 | 0.00119403 |
| IL3RA | 1.102276725 | 0.000950825 |
| CCDC137 | -0.8508304 | 4.73E-08 |
| RGPD2 | 0.695199723 | 0.000112809 |
| ARL15 | -1.475118822 | 2.64E-09 |
| C12orf56 | 3.538882353 | 6.35E-07 |
| AC027601.1 | 7.12530457 | 1.13E-05 |
| SOCS1 | 1.726204311 | 0.000377096 |
| GAS2L1 | -0.747835662 | 1.55E-05 |
| HS6ST3 | -0.792308439 | 0.004825603 |
| TNFAIP8L1 | -0.722828431 | 0.000393414 |
| RAD51D | -1.362902237 | 1.98E-08 |
| MAPK11 | 0.552078058 | 0.011634437 |
| SP140L | -0.597306076 | 0.000272636 |
| MRPL30 | -0.877452866 | 8.47E-08 |
| SMYD3 | 0.888245498 | 0.000489139 |

| | | |
|---|---|---|
| SH3BGR | 3.030040588 | 1.32E-05 |
| FAM174B | 1.070330179 | 0.017544518 |
| ZSWIM9 | 0.931332493 | 1.81E-05 |
| TMEM179B | 0.55598109 | 1.92E-06 |
| PARPBP | -1.285199431 | 7.11E-09 |
| STAC3 | 1.718115477 | 0.000553627 |
| ROR1 | -0.734659053 | 0.000298673 |
| SDHAP1 | -0.938054177 | 0.003848057 |
| AC138393.1 | 1.56608412 | 7.34E-08 |
| IRF7 | 1.911512939 | 0.000423931 |
| L3MBTL1 | 3.355278801 | 8.38E-11 |
| BRCC3 | -0.711915624 | 4.40E-07 |
| PDE6G | 3.862533315 | 3.23E-05 |
| PRKG1 | -0.930119388 | 0.021728894 |
| NR2F2 | -1.887540025 | 8.63E-11 |
| LSAMP | 0.648301297 | 0.001000009 |
| AHNAK2 | 1.025200724 | 6.84E-09 |
| OLFML2A | 0.219754902 | 0.032157192 |
| SP1 | -0.361644253 | 3.19E-05 |
| ACTBP7 | 3.125010544 | 1.69E-06 |
| MRPL40 | 0.242322499 | 0.008832588 |
| PDIA2 | 3.513306338 | 8.10E-05 |
| PCGF3 | -0.804876225 | 2.63E-08 |
| LMLN | 1.659913282 | 6.88E-09 |
| P4HB | 0.365642317 | 3.87E-07 |
| PSMD13 | 0.329143437 | 0.000837826 |
| PBX1 | 0.527516434 | 0.000326162 |
| NDUFA4L2 | 1.082511756 | 0.00675093 |
| SHC4 | 2.086972614 | 1.74E-09 |
| AC034236.1 | 0.423764017 | 4.53E-07 |
| ZFP36L1 | -0.576427689 | 3.87E-07 |
| UBE2L3 | -0.575092076 | 5.14E-07 |
| BRWD1 | -1.375999397 | 1.97E-09 |
| PMEL | 2.675297146 | 7.78E-07 |
| SYN3 | 1.207062183 | 0.035860978 |
| POU3F1 | 2.444241543 | 3.86E-06 |
| ZBTB3 | -0.978329748 | 0.000791282 |
| MYBL1 | -0.49892808 | 1.11E-05 |
| MOSMO | -0.874521473 | 1.95E-09 |
| DRG1 | -0.528072436 | 1.82E-05 |
| ANKFY1 | -1.29720858 | 1.81E-10 |
| YTHDF3 | -0.537529247 | 2.57E-07 |
| SRL | 9.898561551 | 1.26E-09 |
| IFIT1 | -0.798077987 | 0.004247929 |
| KCNQ5 | -1.09962671 | 6.36E-07 |
| MORF4L1 | -0.907748946 | 4.87E-09 |
| NLRP9 | 3.238825612 | 1.22E-07 |
| WDR53 | 0.488606455 | 0.00033704 |

| | | |
|---|---|---|
| SLC52A2 | -0.531892473 | 8.06E-07 |
| PIGP | 0.87585256 | 0.000176779 |
| IKZF1 | 6.86818075 | 2.62E-05 |
| PCYT2 | 0.446864948 | 0.000148632 |
| NAT8L | -0.37693227 | 0.001842692 |
| ARL17A | -1.085113103 | 0.003966669 |
| RPL12P4 | 0.598242828 | 4.55E-08 |
| AL035411.1 | -0.263265363 | 0.000265219 |
| DNAH14 | 0.509961378 | 0.000132018 |
| ZNF829 | -1.023730579 | 5.53E-07 |
| THNSL1 | 2.208523132 | 1.67E-10 |
| IFITM1 | 0.738470217 | 0.026631282 |
| FFAR3 | 7.202297888 | 1.06E-05 |
| LINC00839 | -0.564776256 | 0.000283711 |
| KLHDC8B | 0.324953841 | 0.047228217 |
| SETD4 | -1.139624229 | 4.62E-05 |
| RNPC3 | 0.944775906 | 5.95E-07 |
| IRS2 | 0.978696464 | 5.91E-09 |
| C7orf61 | 4.598377539 | 2.94E-06 |
| FAM186A | 4.900741389 | 0.000171026 |
| BICD2 | -0.5438389 | 6.00E-07 |
| SLITRK2 | -0.566342933 | 0.036903226 |
| RASA3 | -0.403973484 | 3.09E-05 |
| LRCH3 | -0.531752971 | 0.001602542 |
| LEMD1 | 3.695424116 | 0.01106092 |
| NDUFA13 | 0.186441299 | 0.017505367 |
| AC021092.1 | 1.38533701 | 6.51E-10 |
| ZNF529 | -0.336133601 | 0.000917626 |
| ZNF284 | 0.490249346 | 0.001382692 |
| TAL2 | 3.721119611 | 8.49E-06 |
| MATN1-AS1 | 1.761793614 | 3.43E-06 |
| AIDA | -0.370962814 | 1.81E-05 |
| C15orf41 | -1.23357949 | 3.46E-07 |
| AC012085.1 | -0.181258082 | 0.002119087 |
| NBPF6 | 1.793060158 | 1.43E-05 |
| CYP2R1 | 1.284150029 | 4.99E-05 |
| LRRC70 | 2.946494222 | 1.64E-06 |
| PIP5K1C | -0.168244806 | 0.011803949 |
| OR9M1P | 6.724095173 | 3.61E-05 |
| C2orf76 | 1.573377554 | 6.07E-06 |
| POLR3C | -0.267855243 | 0.001597762 |
| UBL4B | 8.522814142 | 8.58E-08 |
| CCDC84 | -0.737001167 | 4.18E-05 |
| BCL9L | -1.636834287 | 3.08E-10 |
| KIF18B | -1.822571492 | 1.20E-10 |
| ZNRF1 | -0.666844384 | 0.000166659 |
| SAPCD2 | -0.72159974 | 3.65E-07 |
| EDARADD | 0.131151923 | 0.019419242 |

| | | |
|---|---|---|
| SOWAHB | 8.120138048 | 4.10E-07 |
| KLHL32 | 6.956814904 | 2.22E-05 |
| MRTFB | -1.517646215 | 9.44E-09 |
| BTLA | 7.86665498 | 1.12E-06 |
| ZNF17 | 0.982453263 | 8.58E-08 |
| KDM4D | 2.310921549 | 1.28E-08 |
| GPAT2 | 4.576583276 | 0.012695794 |
| TOR3A | -0.704461848 | 6.92E-08 |
| PPP1CC | -0.810395302 | 6.32E-10 |
| ZNF555 | 2.496953278 | 5.87E-07 |
| NAP1L3 | -0.262377553 | 0.020680415 |
| CA5BP1 | -0.389738851 | 0.000611716 |
| BACE1 | 0.893174395 | 1.24E-09 |
| RGS9BP | 1.119585098 | 0.048433635 |
| RXRA | -1.606874357 | 5.51E-09 |
| ANKRD37 | 0.898885047 | 5.17E-06 |
| ZNF75D | 1.481274262 | 9.58E-08 |
| KRT10 | -0.450364295 | 1.73E-05 |
| CCDC30 | 3.317256678 | 6.14E-07 |
| NKRF | -1.22891074 | 1.25E-08 |
| KPNA4 | -1.549095597 | 3.13E-11 |
| ZNF501 | 2.077062981 | 4.81E-07 |
| ZNF197 | 1.488693176 | 2.45E-10 |
| RPS23 | 0.519982549 | 3.15E-08 |
| BTN3A2 | -0.907817871 | 9.64E-07 |
| PCLO | 0.785487714 | 3.59E-08 |
| INSIG1 | 4.304307369 | 6.91E-14 |
| ZNF396 | 3.183145994 | 2.64E-07 |
| TMEM222 | -0.298825337 | 0.002006872 |
| ARHGAP30 | 3.424600196 | 3.87E-08 |
| SEPTIN10 | 0.525531407 | 6.55E-06 |
| FAM86B1 | 2.416729565 | 1.42E-07 |
| FOXD2 | -0.905614451 | 0.001064563 |
| GPATCH8 | -1.056282618 | 7.43E-10 |
| CEACAM19 | 1.978543982 | 9.04E-06 |
| NF2 | -1.620703837 | 4.16E-10 |
| SMIM29 | 0.959189993 | 1.32E-06 |
| UBE2H | -0.635258438 | 9.78E-08 |
| HPDL | -1.445889316 | 5.27E-06 |
| KATNA1 | -0.562105868 | 0.001827491 |
| ARAP1 | -0.391053704 | 0.000175017 |
| KIF24 | -0.520297125 | 9.29E-05 |
| PDE2A | 0.882905832 | 0.016301644 |
| PRG2 | 3.694386922 | 0.000104485 |
| PRR5 | -1.184117139 | 3.99E-05 |
| ZFP91 | -1.220902453 | 6.93E-11 |
| C17orf58 | -0.690751735 | 1.83E-05 |
| BCDIN3D | 1.298850381 | 3.00E-08 |

| | | |
|---|---|---|
| LYRM7 | -0.559346314 | 6.45E-06 |
| MST1L | 1.172331828 | 0.009603328 |
| BCR | -0.293410123 | 0.006098768 |
| SPIN4 | -1.445344499 | 9.63E-10 |
| SPIN2B | -1.251542811 | 5.34E-06 |
| HYAL3 | 0.463334656 | 0.001679729 |
| VSIG10L | 1.953041841 | 1.50E-08 |
| ZNF397 | 1.325467721 | 1.35E-09 |
| ZSCAN30 | 2.568882934 | 1.30E-10 |
| TPCN1 | 0.396874453 | 0.00055182 |
| HEXIM1 | -0.518692844 | 5.02E-07 |
| KRT14 | 0.406782549 | 0.020186239 |
| TRABD2A | 2.331129685 | 4.36E-06 |
| PDZD7 | 1.31716642 | 1.44E-05 |
| POFUT2 | 0.1823048 | 0.044944385 |
| QRFPR | -0.742975472 | 0.002251868 |
| ERCC6L | -0.272076282 | 0.000627113 |
| TMEM17 | -0.892962192 | 0.000402799 |
| ZDHHC17 | -0.525192887 | 1.80E-06 |
| P2RY4 | 4.418973152 | 0.00097228 |
| ZNF395 | -0.317073535 | 0.000449034 |
| ZACN | -0.470854044 | 3.02E-06 |
| CHCHD2P9 | 0.392275044 | 0.004671918 |
| PPARA | 0.499917507 | 0.000341261 |
| TMEM232 | 4.11872115 | 6.51E-05 |
| KANK3 | 1.219376972 | 0.005374143 |
| TMPRSS6 | 1.686297508 | 0.00265461 |
| TMEM216 | -0.389719702 | 0.009486937 |
| RPS19BP1 | 1.200954498 | 2.38E-09 |
| TMEM262 | 0.418359103 | 0.002595519 |
| C3orf70 | -1.697359584 | 1.85E-06 |
| TEAD1 | -1.536619695 | 1.50E-11 |
| PLCD1 | 1.30924204 | 1.17E-05 |
| MITF | 0.413415097 | 0.000213005 |
| HEATR4 | 2.309628554 | 0.000859769 |
| NAP1L1 | 0.245064601 | 1.59E-05 |
| CMC1 | -0.700880256 | 5.73E-07 |
| SLIT1 | 4.748423072 | 8.49E-07 |
| LYPD6 | -0.457584096 | 0.048497273 |
| AKR1C1 | 0.987750151 | 0.011986396 |
| SPATA21 | -0.908351434 | 3.86E-08 |
| RNF220 | -0.885633477 | 3.62E-08 |
| SHTN1 | -1.128598238 | 2.61E-06 |
| BAGE2 | 0.265656324 | 0.004850438 |
| AL162231.1 | 1.257746478 | 0.00046669 |
| ZNF546 | 1.44690536 | 1.44E-06 |
| MT1X | 1.974186634 | 1.30E-06 |
| FNBP1 | -0.845746203 | 7.89E-08 |

| | | |
|---|---|---|
| DYNC2H1 | 0.415035845 | 6.09E-05 |
| BCAM | 0.274721778 | 0.02949604 |
| RSBN1L | -1.565590222 | 5.45E-10 |
| EPOR | 1.877250556 | 4.67E-09 |
| TAF9B | -0.523833006 | 2.14E-06 |
| MAGI2 | 1.586687325 | 8.70E-06 |
| CHP1 | 0.361009084 | 6.23E-05 |
| C11orf96 | 3.927363058 | 5.36E-10 |
| COL4A1 | 0.154869705 | 0.003984522 |
| HSPA14 | -0.70557814 | 4.35E-07 |
| SIRT7 | -0.219972839 | 0.026117234 |
| PRR13P5 | 0.197322497 | 0.034045192 |
| IFT140 | 0.91030829 | 1.15E-05 |
| POTEG | 1.058108793 | 0.000296509 |
| AGMO | 1.311767895 | 2.68E-06 |
| TLR5 | 4.038111675 | 2.19E-09 |
| USP7 | -0.894858342 | 8.20E-09 |
| COX8C | 7.370150861 | 5.50E-06 |
| PLEKHN1 | 0.954748576 | 0.023836573 |
| TERF1P2 | -0.813377495 | 0.016051036 |
| ZNF385C | 2.872658161 | 4.14E-05 |
| TET3 | -1.192077714 | 6.01E-10 |
| ZNF286A | -0.654834302 | 5.36E-07 |
| ISG15 | 0.928820848 | 0.000484053 |
| TCL6 | 4.590441124 | 0.000453797 |
| C17orf97 | 0.930782987 | 0.000803556 |
| RGPD1 | 0.473457752 | 5.22E-05 |
| DHRS4L2 | 0.576889724 | 0.002052549 |
| VMAC | 0.671579133 | 0.003280797 |
| HAPLN4 | 1.415448473 | 0.015032697 |
| B3GLCT | -0.587279263 | 5.02E-05 |
| SPRY4 | -1.017489272 | 1.72E-09 |
| KRT18P59 | 1.465677432 | 0.000231666 |
| TMEM203 | -0.648458458 | 1.00E-06 |
| DNAJB13 | 2.552129076 | 5.27E-05 |
| TCEA1 | -0.181031243 | 0.003320811 |
| FANCA | -0.993667251 | 5.75E-08 |
| SECISBP2 | -0.532106429 | 7.01E-06 |
| C9orf153 | 1.140821639 | 0.025899414 |
| SEMA4D | 0.522515073 | 4.13E-05 |
| LIN28B | -1.355718257 | 3.64E-08 |
| DNAH17 | 1.827204996 | 4.42E-07 |
| MCRS1 | -0.378461557 | 0.000176584 |
| FANCM | -0.682523359 | 3.60E-06 |
| CARD9 | 0.653197212 | 5.46E-05 |
| PEAR1 | 1.09555196 | 0.004785926 |
| TMEM202 | 6.673854288 | 4.15E-05 |
| SOWAHD | 4.669486231 | 2.80E-08 |

| | | |
|---|---|---|
| ZFP69 | -0.459430399 | 0.001085981 |
| TMEM220 | 2.339136078 | 5.79E-08 |
| H1-2 | 0.833522592 | 0.000235861 |
| PLSCR3 | 0.796556724 | 3.52E-07 |
| EIF4EBP1 | 0.158303801 | 0.010371732 |
| CCDC157 | 0.610650343 | 0.000985572 |
| TTC24 | 3.275588651 | 9.33E-06 |
| PALM3 | 0.569976075 | 0.007702361 |
| SHISA7 | 3.467081654 | 0.000177528 |
| LRRC74B | 2.948044185 | 0.001158543 |
| LCN10 | 6.912180263 | 2.73E-05 |
| C2orf66 | 3.69277468 | 0.011839452 |
| AC091057.1 | -1.545510559 | 7.57E-08 |
| PMS2CL | -0.401674633 | 0.003217301 |
| DNER | 1.011552902 | 0.006001805 |
| CPSF4L | 4.322375457 | 0.00127783 |
| KLHL17 | -0.511494932 | 0.001857458 |
| KCTD9P3 | -0.944934672 | 0.00375947 |
| RINL | 1.272969061 | 2.30E-05 |
| HNRNPA1P61 | 2.207830732 | 0.000177585 |
| OR7D2 | 9.076099095 | 1.25E-08 |
| AC026412.1 | -0.399254169 | 0.018222739 |
| SNHG28 | 4.675529089 | 3.37E-09 |
| MEIS3P2 | 1.636419204 | 1.74E-05 |
| S100A3 | 1.117988189 | 4.02E-05 |
| UBQLN2 | -0.443785978 | 2.52E-06 |
| C19orf67 | 7.401354604 | 5.71E-06 |
| ZNF490 | 1.954943674 | 2.52E-10 |
| CLCN1 | 5.391840379 | 3.12E-05 |
| NWD1 | 2.438612607 | 0.000263259 |
| RNF133 | 3.670380199 | 0.007641846 |
| RAB42 | 3.429912032 | 2.40E-06 |
| C11orf95 | -1.532125346 | 8.99E-10 |
| GPR89B | -0.881954145 | 4.33E-07 |
| AGRN | 0.103409437 | 0.046817158 |
| TMPPE | 0.371396592 | 0.012529655 |
| ZNF626 | 0.806287627 | 1.27E-05 |
| HEPACAM2 | 9.736745175 | 2.46E-09 |
| ZC3H6 | 5.172455891 | 0.006026744 |
| LINC00265 | -0.549098385 | 0.008681267 |
| PRKAR1B | 0.349069016 | 0.00402642 |
| NUTM2B | 4.129915412 | 1.05E-10 |
| NCR3LG1 | -0.88610972 | 3.75E-07 |
| DCUN1D3 | 0.539633131 | 1.70E-05 |
| POTEE | 1.093057294 | 5.21E-06 |
| AD000671.1 | 0.630016379 | 0.000148593 |
| ZNF793 | 1.025653982 | 2.45E-05 |
| TUBB4B | -0.736878181 | 2.89E-10 |

| | | |
|---|---|---|
| AGAP4 | -0.627661135 | 0.00329089 |
| AC010442.1 | -0.702273428 | 0.002544951 |
| COMMD6 | 0.78780238 | 1.16E-05 |
| HYKK | 1.365872987 | 6.81E-05 |
| ZNF383 | 0.395700073 | 0.003833009 |
| HES4 | 2.729381209 | 6.91E-06 |
| ZNF669 | 0.921833251 | 3.24E-07 |
| CENPP | -1.065146918 | 6.78E-09 |
| ZNF559 | -1.618820757 | 1.70E-09 |
| SBK1 | 2.08927014 | 0.007315606 |
| SLC38A3 | 1.143727114 | 0.001047246 |
| GTF2F2 | -0.879971134 | 4.38E-07 |
| FAM92A | -0.418355911 | 2.80E-05 |
| FOCAD | 0.193961177 | 0.017349036 |
| H3-5 | -0.525676547 | 2.71E-06 |
| GPAT2P2 | 3.49093796 | 9.91E-06 |
| PPP3R2 | 7.693404731 | 1.74E-06 |
| SELL | 2.047636483 | 0.026191746 |
| NANOS2 | 4.232341059 | 1.70E-05 |
| CERKL | -0.290117077 | 0.008832588 |
| SLC24A5 | -1.28555258 | 0.004429267 |
| H2AX | -0.853282748 | 3.70E-07 |
| SERPINA5 | 0.727383777 | 0.000199745 |
| C19orf54 | 0.703541867 | 7.93E-06 |
| LCTL | -0.487092595 | 0.007316118 |
| NCCRP1 | 7.527714283 | 3.07E-06 |
| C22orf34 | -0.69817732 | 0.022119219 |
| AF279873.1 | -0.506544195 | 0.007922503 |
| SRSF10 | -0.831581889 | 1.02E-09 |
| DUSP28 | 1.917895194 | 9.42E-08 |
| CCDC9B | -1.31804408 | 5.23E-10 |
| NBR1 | 0.309969886 | 0.000128214 |
| RALGAPA2 | -0.506502051 | 0.000407893 |
| NDOR1 | -0.192162794 | 0.015344569 |
| FBLL1 | 1.968047215 | 4.09E-05 |
| PAQR9 | -1.181843933 | 0.022213268 |
| CFAP54 | 3.076503056 | 0.000377459 |
| CLN3 | -0.454505335 | 0.001344975 |
| FAM72B | -0.56914992 | 3.45E-06 |
| ASAH2 | -1.246875415 | 4.92E-05 |
| NANOS1 | -1.114848821 | 5.94E-05 |
| ZNF177 | 1.225591337 | 0.000159403 |
| RTL6 | -1.453839451 | 3.73E-10 |
| DPYD | -0.215257225 | 0.023454765 |
| S100A16 | 0.29758236 | 0.000594865 |
| PTAR1 | -0.945276825 | 2.92E-09 |
| SAXO2 | 4.792974234 | 2.53E-10 |
| H1-9 | -1.358117889 | 5.05E-05 |

| | | |
|---|---|---|
| RHCE | 4.316310006 | 2.83E-05 |
| SLC4A5 | 0.373902165 | 0.011630738 |
| CYP51A1-AS1 | 2.559615937 | 0.00024613 |
| ZDHHC9 | -0.473736677 | 8.92E-05 |
| QRFP | 1.508436892 | 0.000504416 |
| SMIM15 | -0.886305604 | 6.59E-09 |
| TMEM120B | -0.173036535 | 0.011082331 |
| RBM34 | -0.202950039 | 0.007213294 |
| TMEM198 | 1.328313111 | 5.57E-06 |
| SPRED3 | 0.479589571 | 0.046340286 |
| ZNF548 | 0.418555645 | 0.000201747 |
| MTF1 | -0.300400718 | 0.000188964 |
| TMEM201 | -0.915370339 | 1.81E-08 |
| NHLRC3 | 0.307464245 | 0.017277668 |
| ZDHHC11 | -1.201099828 | 1.29E-07 |
| CNR2 | 8.567321168 | 8.18E-08 |
| LINC00910 | 2.516366174 | 6.54E-06 |
| RPL14 | 0.339238008 | 9.53E-06 |
| BEND4 | -1.102642201 | 4.12E-06 |
| RPSAP47 | 0.162041153 | 0.008844055 |
| ZNF563 | 2.028278719 | 1.92E-05 |
| RPL10AP2 | 0.463472015 | 5.88E-05 |
| FBF1 | 1.046424069 | 6.55E-07 |
| ASTL | 6.237743196 | 3.66E-06 |
| MSL1 | -1.624793593 | 4.86E-11 |
| AC099489.1 | 5.843699552 | 1.32E-08 |
| LRRK2 | -1.092941988 | 0.002637967 |
| GJB3 | -1.607232378 | 6.37E-06 |
| TRMT2B | -0.442080098 | 0.020293987 |
| HACD4 | 0.855403726 | 0.006998743 |
| CFAP126 | 2.419962324 | 1.47E-06 |
| USP32P1 | 0.265688039 | 0.016560323 |
| FAM120AOS | 0.234583786 | 0.001789783 |
| MSANTD1 | 1.825661746 | 0.001435791 |
| NELFB | -0.239375808 | 0.001114267 |
| LRRC66 | 4.433916304 | 3.40E-06 |
| ZNF292 | 0.510303036 | 1.69E-06 |
| HUS1B | 1.445572607 | 0.000552616 |
| KCTD21 | 1.846648314 | 2.54E-10 |
| SBSN | 5.487932478 | 3.36E-08 |
| ADAT2 | -0.998456283 | 1.88E-06 |
| SHLD2P3 | -0.619019603 | 0.030741138 |
| NDUFA4 | 0.255955976 | 0.002346208 |
| ANKDD1B | 1.86726511 | 0.001961655 |
| ALKBH2 | -0.6754427 | 5.21E-06 |
| RNFT1 | -0.676353034 | 0.000685909 |
| FAM111B | -1.319081836 | 1.19E-09 |
| H1-0 | 0.143098057 | 0.003982268 |

| | | |
|---|---|---|
| LITAF | -0.175143806 | 0.005615065 |
| TMEM120A | 0.561382102 | 0.000123014 |
| ARID2 | -1.540227666 | 1.72E-10 |
| RIMKLBP1 | 1.706108034 | 3.52E-09 |
| SF3B3 | -0.382710766 | 2.10E-06 |
| SP6 | 6.086045344 | 7.20E-08 |
| CLDN4 | 1.356636738 | 3.30E-06 |
| FAM47E | 1.761417191 | 0.000632014 |
| JPT1 | -0.334462565 | 4.28E-05 |
| ZNF527 | -0.524581147 | 0.00207761 |
| S100A13 | 0.388972436 | 4.18E-05 |
| ZNF33A | -1.740224962 | 2.86E-09 |
| OR14I1 | 3.6493795 | 0.000196414 |
| PCDH18 | 1.323854213 | 1.27E-06 |
| ZNF600 | -0.242513037 | 0.035729373 |
| DPY19L2P1 | -1.697727073 | 1.21E-07 |
| PAX8-AS1 | 2.08550076 | 8.10E-09 |
| AC069277.1 | 0.669932312 | 0.036732415 |
| LINC00943 | 3.267956576 | 1.23E-05 |
| TSPYL1 | -0.289945758 | 0.000109063 |
| ALKAL2 | 1.375303127 | 0.03001381 |
| ZKSCAN3 | 1.361049997 | 0.000190862 |
| FOXR2 | -1.451367176 | 1.27E-08 |
| RRP7A | -0.642201508 | 1.99E-06 |
| LIN54 | -0.171512515 | 0.032311612 |
| FAM53B | -0.946387254 | 1.31E-07 |
| BNIP5 | 8.638477057 | 1.02E-07 |
| S100A14 | 5.451057374 | 4.23E-06 |
| KAZN | -1.131812608 | 6.43E-06 |
| SLC35E2B | 0.793487858 | 1.04E-07 |
| RPS2P46 | 0.478778478 | 1.35E-05 |
| NEMP2 | -0.762367154 | 8.75E-05 |
| ALG1L | 3.112144246 | 8.06E-05 |
| HMGB1 | -0.299231709 | 1.67E-06 |
| GJB4 | 2.354520967 | 0.003038856 |
| BLOC1S2 | -0.261409736 | 0.000758531 |
| SYCP2 | 3.153577418 | 3.56E-06 |
| ZNF724 | 1.355389951 | 2.12E-09 |
| ZNF699 | 1.020982736 | 5.08E-06 |
| TDRD7 | -0.613624245 | 0.000689273 |
| SERPINA3 | 0.699269782 | 9.77E-07 |
| AKR1C3 | 1.094614173 | 0.000607686 |
| SPATS2L | 0.547590151 | 1.43E-06 |
| ZNF250 | 1.399970452 | 3.25E-07 |
| ZNF79 | 1.869595644 | 2.65E-09 |
| S100A4 | 0.585488496 | 1.54E-06 |
| FAT4 | -0.745246757 | 4.67E-05 |
| ACADSB | 0.627837608 | 2.52E-06 |

| | | |
|---|---|---|
| STK40 | -0.321271228 | 9.16E-05 |
| RPS2P4 | 0.741989344 | 2.72E-05 |
| SEMA4A | 4.726241086 | 6.26E-10 |
| HRCT1 | -1.410050753 | 0.004766239 |
| MPHOSPH8 | -0.440699365 | 1.06E-05 |
| RNF216P1 | -0.866004111 | 1.93E-07 |
| EEF1A1P5 | 0.417755342 | 9.87E-09 |
| GREB1 | 0.691578528 | 0.01304509 |
| ZNF766 | 0.985427949 | 5.74E-08 |
| RYR1 | 1.784933611 | 0.005664574 |
| SRGAP3 | 2.532869059 | 6.00E-10 |
| FAM217B | -0.81152502 | 9.98E-09 |
| LCOR | 0.11471247 | 0.034959054 |
| SUPT5H | -0.165748503 | 0.004530786 |
| XPNPEP3 | 0.301809283 | 0.002842025 |
| ZNF107 | -0.773658105 | 7.86E-08 |
| PPIA | 0.144878913 | 0.03516562 |
| ZNF836 | 0.842614227 | 0.000855568 |
| ZNF493 | 1.235114539 | 1.76E-09 |
| GTF2IRD2 | 1.879358255 | 2.90E-05 |
| SUPT3H | 0.921285259 | 0.007240763 |
| NIF3L1 | -0.83373567 | 2.62E-06 |
| GARS-DT | 0.210253041 | 0.017544518 |
| ATP2A1 | 2.374133173 | 3.60E-08 |
| ZNRD1ASP | 1.346449456 | 4.43E-05 |
| POM121 | -0.661334891 | 2.97E-08 |
| ZBTB44 | -1.944964913 | 2.03E-11 |
| STK31 | 2.645329146 | 0.000765735 |
| ZNF729 | 1.125026598 | 0.001762187 |
| CD55 | 0.581691891 | 4.61E-06 |
| ZNF565 | 0.763777745 | 0.000781992 |
| NTNG2 | 1.599889597 | 0.00429794 |
| WDR5 | -0.87083271 | 8.19E-09 |
| TRRAP | -0.460659481 | 5.46E-06 |
| NUDT11 | -0.67799137 | 9.18E-05 |
| SRGAP2B | 0.831350001 | 1.29E-08 |
| ASB13 | -0.311609942 | 0.032095811 |
| SLC35F1 | -0.317979841 | 0.027601059 |
| ZNF34 | 0.894578575 | 0.000132839 |
| ZNF781 | 0.523575993 | 0.010369395 |
| INCA1 | 1.708456292 | 4.69E-05 |
| PTPN1 | -0.586372107 | 1.57E-07 |
| EVL | -1.151150048 | 1.86E-05 |
| THEM5 | 2.998612794 | 0.000956222 |
| NOXO1 | -1.163274126 | 0.000221182 |
| EPHB4 | -0.573257625 | 6.25E-06 |
| ZNF765 | 0.747400822 | 4.41E-07 |
| ZNF124 | -0.331538597 | 0.006505888 |

| | | |
|---|---|---|
| XRCC6 | 0.176679871 | 0.00122962 |
| S100A5 | 1.133379822 | 0.049224937 |
| C20orf204 | 2.343640288 | 6.07E-09 |
| PPP1R26 | 0.335071932 | 0.000295403 |
| NBPF4 | 1.773761389 | 4.46E-05 |
| TSC22D2 | -0.370590933 | 0.000249681 |
| NPIPB15 | 1.421094068 | 2.12E-05 |
| ZNF569 | -1.620194798 | 2.46E-09 |
| ARMCX4 | -0.604525104 | 0.000749904 |
| YRDC | -0.93241484 | 4.71E-09 |
| ZNF777 | 0.207109529 | 0.024502033 |
| PIK3R4 | -0.947647664 | 7.56E-09 |
| ZNF775 | 0.748695276 | 0.000267178 |
| ZNF605 | -0.270288797 | 0.006681075 |
| TRAPPC2 | -0.789405321 | 7.04E-05 |
| MYL6B | 0.552676485 | 2.57E-06 |
| ZNF799 | 1.170654527 | 1.46E-07 |
| SIAH1 | -0.545045173 | 1.59E-05 |
| C20orf96 | 0.609887292 | 0.022222646 |
| ESRRG | 1.260701713 | 0.005054652 |
| SULT1A1 | -0.907161935 | 0.003902863 |
| ARL9 | 2.058195484 | 0.000276565 |
| PRPF40A | -0.315303044 | 4.30E-05 |
| GDAP2 | -0.627228123 | 2.35E-06 |
| TCEAL3 | 0.320118511 | 0.029345485 |
| ANAPC7 | -0.419868958 | 2.13E-05 |
| TPK1 | -0.722945386 | 0.014168814 |
| AFAP1 | -1.610810159 | 7.20E-11 |
| NACA | 0.423565408 | 1.96E-07 |
| MYO18A | -0.285070106 | 0.000206781 |
| BORCS6 | 1.971151954 | 1.83E-08 |
| FAM72A | -0.525218662 | 6.64E-06 |
| SULF2 | 0.584967891 | 1.05E-06 |
| HBG2 | 1.749644655 | 8.26E-05 |
| LAMA2 | 0.782503434 | 0.004812978 |
| PLXNB2 | 0.666150231 | 1.14E-07 |
| XRCC2 | -0.837955027 | 4.42E-08 |
| MYO6 | -0.303383999 | 0.004933456 |
| MRTFA | -0.494848375 | 0.00038017 |
| ZNF782 | 0.435588767 | 0.003054721 |
| POTEF | 1.133118603 | 3.39E-05 |
| ZNF846 | 2.456539 | 9.41E-08 |
| MMP1 | 1.763562064 | 3.24E-08 |
| TCF4 | -0.715298657 | 8.48E-05 |
| WNK3 | 1.91365203 | 8.91E-05 |
| HRH1 | 1.637380604 | 1.52E-09 |
| RABL6 | -0.269152096 | 0.003399295 |
| ZNF136 | 0.237065544 | 0.003301954 |

| | | |
|---|---|---|
| ZKSCAN5 | 0.29097066 | 0.000205219 |
| ZNF502 | 1.055099237 | 3.55E-05 |
| TRAPPC4 | -0.252779655 | 0.004779089 |
| TECPR2 | 0.784979743 | 5.30E-06 |
| TLR7 | 8.55123424 | 7.80E-08 |
| LINC00173 | 1.88727283 | 0.007269256 |
| ZFP62 | 0.26167739 | 0.001380046 |
| TOMM7 | 0.259554907 | 0.001764845 |
| TRPV1 | 1.193019338 | 3.95E-06 |
| ZNF33B | -1.169713123 | 2.52E-08 |
| AC009022.1 | 0.690848338 | 0.002708553 |
| ZNF512B | -0.464173201 | 5.70E-06 |
| AMZ2 | 0.987534526 | 3.04E-09 |
| NF1 | -0.605642888 | 1.16E-07 |
| VKORC1L1 | -0.82285183 | 1.23E-08 |
| ZNF418 | 0.519972592 | 0.014773438 |
| DAPK1 | 2.198142886 | 2.59E-09 |
| COL27A1 | 0.236097639 | 0.002370273 |
| GM2A | 0.389279877 | 8.40E-05 |
| H2AC13 | 4.167776524 | 5.49E-08 |
| S100A2 | 0.810201211 | 2.32E-06 |
| SNHG17 | -0.578553631 | 7.27E-07 |
| ZNF700 | 0.254155041 | 0.002779896 |
| CD47 | -0.77082754 | 9.25E-08 |
| TLE1 | -0.525094736 | 0.000417786 |
| MAML3 | 1.246735993 | 1.95E-09 |
| H2AC11 | 3.911694972 | 4.41E-06 |
| STRN3 | -0.787941841 | 1.97E-05 |
| NPIPB10P | 1.366946787 | 0.000112809 |
| CTBP1-DT | 0.303122029 | 0.01383133 |
| ZSCAN16 | 1.138347663 | 3.41E-05 |
| MVB12B | -1.147164701 | 1.39E-06 |
| ILRUN | -0.982740029 | 3.81E-09 |
| AC008758.1 | 1.873174067 | 5.56E-09 |
| POTEI | 0.923431349 | 3.23E-05 |
| ARID5A | 0.648566599 | 0.003813822 |
| PPTC7 | 0.126993398 | 0.018419264 |
| NHLRC2 | -1.578947339 | 2.58E-10 |
| H2AC7 | 4.40165903 | 0.021881779 |
| ZFP28 | 0.605915241 | 1.98E-05 |
| CBWD3 | -0.224467694 | 0.041580203 |
| SCN8A | -1.084241467 | 5.55E-06 |
| LAMB3 | 0.997370728 | 1.25E-05 |
| H2BU1 | 3.030649004 | 1.36E-07 |
| ANKRD36B | -1.215111302 | 3.73E-05 |
| ARHGEF12 | -0.652349412 | 2.67E-08 |
| ZNF252P | 0.234762854 | 0.001103145 |
| FLNA | -0.067860904 | 0.018620012 |

| | | |
|---|---|---|
| SRGAP1 | -0.755537097 | 6.97E-07 |
| FAM3C | -1.063623363 | 1.82E-09 |
| NOP9 | -0.883803253 | 1.63E-07 |
| SLC39A10 | 0.981572409 | 5.99E-09 |
| SCOC-AS1 | 2.781146298 | 9.48E-05 |
| CASP4 | -0.629089369 | 4.23E-05 |
| AP2A1 | 0.238668542 | 0.004510539 |
| ZNF585A | 1.264143345 | 2.03E-09 |
| FUT11 | -0.656898295 | 4.86E-06 |
| SMIM10L2B | 4.072672752 | 3.22E-05 |
| ANXA4 | 0.842252368 | 7.75E-06 |
| WDR5B | 0.755481825 | 2.65E-05 |
| NPIPB9 | 0.708062545 | 0.003251189 |
| WDR45 | 0.334283417 | 0.000201152 |
| ZNF138 | -1.08636516 | 7.17E-07 |
| ZNF429 | 2.887847648 | 2.30E-10 |
| ZNF470 | 0.401038353 | 0.000230647 |
| SERTAD1 | 1.812112637 | 7.60E-11 |
| ZNF100 | -1.17573882 | 3.82E-09 |
| CXorf40B | -1.151424413 | 6.38E-07 |
| ZNF398 | -0.61582737 | 7.84E-06 |
| ZSCAN25 | -0.622449497 | 1.76E-05 |
| ANXA6 | 0.309788264 | 0.007066169 |
| ZNF441 | 0.828824152 | 4.23E-07 |
| GMFB | -0.662603179 | 3.91E-08 |
| ZNF420 | -0.199297096 | 0.029506594 |
| ZNF763 | 3.233946029 | 5.89E-08 |
| ZMYM1 | -0.29462512 | 0.000823024 |
| ZSCAN26 | -0.637750282 | 0.001028731 |
| MAFG | -0.489293898 | 1.38E-06 |
| ARRDC1 | 0.210913718 | 0.027226923 |
| KIAA1671 | -0.218016782 | 0.023602527 |
| ZNF300P1 | 0.495492519 | 0.006220246 |
| DYNC1H1 | 0.181717766 | 0.000184998 |
| SLC6A17 | 2.921002239 | 2.21E-07 |
| ZGPAT | -0.867448711 | 2.20E-07 |
| SLC25A29 | -0.214906071 | 0.018215438 |
| PGAP1 | -0.340361662 | 0.000722801 |
| SRC | -0.828716547 | 1.03E-07 |
| ZNF682 | -1.688255989 | 3.30E-08 |
| ZNF772 | 0.321228314 | 0.000441467 |
| ZNF257 | -0.321843956 | 0.001097736 |
| PCNX3 | -0.229543705 | 0.000534205 |
| ACSL5 | 2.276635246 | 0.000400769 |
| LRRC8B | -1.100137321 | 1.68E-07 |
| AC107956.1 | 0.622810469 | 3.99E-05 |
| H3C12 | 3.460873732 | 2.19E-06 |
| ZNF785 | 2.386279337 | 6.80E-05 |

| | | |
|---|---|---|
| PSMD12 | -0.495615978 | 1.52E-06 |
| MAGEA6 | -0.230041281 | 0.001286523 |
| PIWIL2 | 4.010608004 | 1.59E-07 |
| MIRLET7BHG | -0.69987136 | 0.00748685 |
| NOL4L | 0.316381738 | 0.000872051 |
| CYSRT1 | 1.819973913 | 0.000123666 |
| ZSCAN5B | 2.070176995 | 0.002249974 |
| ENTPD4 | -1.573476002 | 2.58E-10 |
| C1D | -0.79438618 | 8.97E-06 |
| TBC1D9B | -0.485246683 | 1.69E-06 |
| H4C11 | 1.843376637 | 0.000223059 |
| KANK2 | -0.391610119 | 5.55E-06 |
| EIF4BP6 | 0.140962748 | 0.007307027 |
| C6orf141 | 0.702675192 | 0.000222151 |
| GTF2E2 | -1.074608329 | 2.07E-07 |
| RAD54B | -0.865173472 | 4.24E-07 |
| RAMP2-AS1 | 3.108802116 | 1.12E-07 |
| FITM2 | -0.426881065 | 0.00010077 |
| BLM | -0.896438264 | 1.07E-07 |
| DDI2 | -1.461664826 | 4.70E-11 |
| TRIM33 | -1.045816783 | 1.19E-09 |
| LRP10 | 0.259024316 | 0.000538931 |
| PELI1 | 0.806840067 | 8.69E-07 |
| AC008543.1 | 1.041313635 | 0.000299079 |
| ZNF655 | -0.119636979 | 0.022037299 |
| UAP1L1 | 1.618568122 | 7.81E-09 |
| BNIP3P1 | 0.305490438 | 0.000719397 |
| ZNF98 | -0.44707388 | 0.001174119 |
| ZNF786 | 0.475925226 | 0.000521853 |
| ZNF517 | 0.423843412 | 0.020530072 |
| ZNF675 | -0.230743225 | 0.014411181 |
| SLC22A5 | -1.434675112 | 5.69E-07 |
| DACT3 | 0.773046311 | 0.002117627 |
| ADARB1 | -1.356517448 | 2.87E-09 |
| ZNF860 | -0.483494582 | 0.002597525 |
| HTT | -0.284399165 | 0.00038014 |
| C5AR1 | 2.992238316 | 1.02E-07 |
| H3C4 | 4.216091186 | 3.67E-06 |
| SHPK | -1.277433535 | 6.79E-09 |
| MAP3K5 | -1.930439422 | 2.27E-06 |
| OGDHL | 0.751347296 | 0.030282004 |
| GSTK1 | 0.188053205 | 0.032983262 |
| HNRNPAB | -0.78244942 | 3.19E-09 |
| STMN3 | 0.644302188 | 0.000101771 |
| COL13A1 | 0.494208264 | 0.025211789 |
| ZNF695 | -1.468200122 | 1.95E-08 |
| SLC2A10 | 2.060923288 | 1.91E-06 |
| ZNF665 | 2.901001261 | 3.62E-09 |

| | | |
|---|---|---|
| RPF2 | -1.034618995 | 3.10E-08 |
| FAM177B | 4.233029226 | 2.26E-07 |
| MIB2 | 0.686438019 | 1.51E-05 |
| MYO5A | -0.43533658 | 3.27E-06 |
| IRF1-AS1 | -1.502522711 | 0.003544068 |
| ATG7 | -1.007995571 | 2.57E-08 |
| SIPA1L1 | -0.452934596 | 0.000651441 |
| TTC30A | 0.726260842 | 1.58E-05 |
| PIGN | -0.400595336 | 0.016678239 |
| ZNF624 | 1.916514237 | 6.68E-07 |
| HHLA3 | 0.872072158 | 0.001143865 |
| HOXA4 | -0.804155605 | 0.001330197 |
| TOPORS | 0.462314355 | 1.47E-06 |
| BCO2 | 2.888100679 | 3.02E-05 |
| ENTPD6 | -0.612798479 | 7.69E-07 |
| CCDC154 | 4.023716421 | 0.000143055 |
| FAR1 | -0.721966196 | 7.81E-08 |
| CPLANE1 | -0.234282157 | 0.039412255 |
| VN1R5 | 3.857866598 | 8.25E-05 |
| ZNF615 | 0.354236846 | 0.001428798 |
| CXorf40A | -0.732639319 | 8.99E-06 |
| CDC42SE1 | -0.417728932 | 2.68E-06 |
| SERPINB2 | 3.997041099 | 7.62E-05 |
| DNAH10 | 4.392530863 | 5.53E-09 |
| AL157838.1 | -1.10223828 | 0.034754975 |
| SPTAN1 | 0.341285408 | 1.90E-06 |
| PARVA | -0.473499562 | 3.59E-06 |
| FAM114A1 | 0.422361074 | 0.006162759 |
| RPE | -1.446915392 | 2.61E-09 |
| ZNF460 | 2.859223581 | 3.87E-10 |
| CR1L | 5.253229929 | 0.000103553 |
| PHF2 | -1.034274311 | 1.92E-07 |
| C14orf178 | 7.341240332 | 6.06E-06 |
| PSAP | 0.440422185 | 6.99E-07 |
| S100A10 | 0.128740877 | 0.008057569 |
| CFAP43 | 4.293629471 | 3.27E-09 |
| RPL37A | 0.383556512 | 2.11E-05 |
| TXNRD3 | 1.234368474 | 2.73E-05 |
| MCMBP | -1.497812421 | 6.58E-11 |
| EME2 | 0.360075357 | 0.000141875 |
| KLHDC1 | 4.204062284 | 9.34E-08 |
| ZNF81 | 0.560452846 | 7.82E-06 |
| TAF13 | -0.549537425 | 1.73E-06 |
| ZNF780A | -1.087500102 | 7.69E-08 |
| ATAD3A | -0.296220673 | 0.000361831 |
| ZNF461 | 0.43640604 | 0.005332626 |
| AC122129.1 | 1.757907245 | 1.42E-05 |
| CCDC180 | 3.334334476 | 3.22E-08 |

| | | |
|---|---|---|
| SLC9A8 | 0.655625855 | 1.75E-06 |
| H4-16 | 1.172072252 | 0.002248832 |
| SLC22A20P | 3.601039324 | 1.23E-05 |
| OR8G1 | 4.235465542 | 3.88E-06 |
| INKA2 | 1.197875951 | 5.16E-09 |
| GPAA1 | 0.411426266 | 1.04E-05 |
| SGTB | -1.540940264 | 1.40E-08 |
| ZNF790 | 0.959497208 | 1.46E-05 |
| NKIRAS1 | -0.614747135 | 1.16E-05 |
| MEIG1 | 6.666862319 | 1.55E-06 |
| KIF13B | 0.333051701 | 0.007477832 |
| ADH5 | 0.191074229 | 0.002100927 |
| H2BC12 | 0.803895888 | 0.000195107 |
| TEAD4 | -1.261429057 | 2.09E-08 |
| SPG7 | -0.287759354 | 0.000690418 |
| HRNR | 0.422926368 | 0.022319628 |
| ZNF677 | -0.769764812 | 4.14E-05 |
| ERO1A | -0.79715274 | 2.92E-08 |
| ZNF823 | 1.582295372 | 3.81E-09 |
| ZNF347 | 1.273892403 | 2.53E-09 |
| PLCG2 | 1.228687647 | 0.002528739 |
| ZNF71 | -0.712867021 | 9.79E-05 |
| S100A6 | 0.50245311 | 1.59E-08 |
| RPL12 | 0.425466309 | 1.67E-06 |
| ZNF121 | -0.16225394 | 0.010163469 |
| MPZL1 | -1.071574224 | 1.21E-09 |
| VPS13A | -0.424377674 | 7.57E-05 |
| AKAP17A | -0.608580627 | 0.000312124 |
| GOLGA6L9 | -0.543162733 | 0.002080471 |
| LEKR1 | 2.286245265 | 9.92E-06 |
| C1orf122 | 0.203541379 | 0.011932799 |
| SNHG12 | -0.910857614 | 4.36E-08 |
| AL592490.1 | -0.376348299 | 0.016490258 |
| NOL8 | -0.52866878 | 3.81E-07 |
| CCDC151 | 1.564805086 | 0.000164459 |
| MRPL42 | -0.572737341 | 1.34E-06 |
| ENTPD7 | -0.567455197 | 4.50E-05 |
| FCGR1B | 6.258836292 | 1.39E-09 |
| ZNF335 | -0.873712459 | 3.63E-07 |
| ZNF560 | 0.44641959 | 0.004430116 |
| RPS4X | 0.406791417 | 3.23E-07 |
| ZNF273 | -0.292665151 | 0.032900516 |
| ZNF84 | 0.351732692 | 8.33E-05 |
| MAK16 | -0.591254743 | 2.09E-07 |
| ZNF667 | 5.302911489 | 0.000110854 |
| SIRPA | -0.436032272 | 0.001640605 |
| GRK6 | 0.32667909 | 0.001682985 |
| MARCHF5 | -0.774167113 | 1.28E-07 |

| | | |
|---|---|---|
| ZBTB14 | -1.223457859 | 2.14E-07 |
| CD2AP | -1.62785608 | 1.62E-09 |
| SFI1 | -0.647591318 | 0.00018972 |
| ZNF248 | -1.436080124 | 4.89E-07 |
| CHSY3 | -1.627185415 | 2.31E-06 |
| LPAR1 | -1.798352068 | 4.62E-05 |
| ZNF544 | -0.79512036 | 2.68E-07 |
| TMEM229B | 0.417854283 | 0.011602556 |
| SOWAHC | 1.417804871 | 1.70E-09 |
| ZNF770 | -0.132276195 | 0.020849028 |
| ZNF849P | 1.399188602 | 1.37E-05 |
| NPIPB6 | 2.753352691 | 3.14E-06 |
| HMGN5 | 0.878572456 | 0.000523447 |
| MIER1 | 0.190090633 | 0.013299908 |
| MAN1A2 | -1.010036143 | 1.01E-08 |
| SVIP | -0.19974036 | 0.011284607 |
| ZNF251 | 1.420356657 | 6.83E-08 |
| DDRGK1 | 0.258658958 | 0.012435228 |
| TFDP1 | -0.621056235 | 2.05E-08 |
| ZNF334 | 0.41489989 | 0.008440017 |
| SZT2 | 0.574758524 | 2.00E-06 |
| ZXDA | -0.622042795 | 0.03343989 |
| RPS6KL1 | 0.450563431 | 0.027869435 |
| AC092143.1 | -0.182433094 | 0.00077769 |
| CACNA1E | 6.724095173 | 3.61E-05 |
| QRICH1 | -0.569945353 | 2.55E-07 |
| DDX42 | -0.34936046 | 1.07E-05 |
| RPL23A | 0.445824339 | 6.57E-07 |
| STYX | -1.485277923 | 1.17E-08 |
| UBL5 | 0.213141173 | 0.004617281 |
| HELZ | -0.890390306 | 1.58E-08 |
| TMEM116 | 1.409064787 | 7.20E-07 |
| UCKL1 | -0.233591742 | 0.007971828 |
| ZNF485 | 1.046327835 | 1.11E-05 |
| PEG3 | 2.535800993 | 1.20E-08 |
| SDAD1 | -0.52101522 | 4.57E-06 |
| ZKSCAN8 | 0.392421116 | 1.35E-05 |
| PHETA1 | 0.480354012 | 0.000242885 |
| HYLS1 | -0.867760174 | 1.25E-07 |
| ZNF442 | 0.650898611 | 0.006121513 |
| ZNF813 | 1.048984659 | 6.35E-07 |
| HOXC4 | -1.233757232 | 2.31E-07 |
| PIM3 | 1.168704482 | 1.25E-09 |
| ASPH | -0.210233292 | 0.001562225 |
| SPRED2 | -0.635955348 | 3.31E-07 |
| WWP2 | -0.405204511 | 0.000104488 |
| GFPT1 | -0.806975503 | 1.67E-08 |
| UVRAG | -1.32383899 | 1.04E-08 |

| | | |
|---|---|---|
| ZNF26 | -0.323433764 | 0.000429379 |
| ITSN2 | -0.418811879 | 1.92E-05 |
| NTRK1 | 1.23017257 | 0.010872114 |
| OGA | -0.998832536 | 4.24E-10 |
| ZNF658B | 2.832208656 | 2.58E-10 |
| TCAF1 | 0.693976362 | 5.56E-08 |
| ZNF69 | 0.777692164 | 4.54E-05 |
| TXNRD1 | -2.006601335 | 3.14E-13 |
| ZNF583 | -0.887134438 | 0.000107877 |
| ZNF568 | -0.646171437 | 6.51E-05 |
| ZXDB | -0.259326342 | 0.009964779 |
| ZNF480 | -1.50677016 | 6.21E-10 |
| ZNF587 | -0.174513792 | 0.007294274 |
| TPM2 | 0.355109664 | 1.81E-05 |
| FLVCR1-DT | 1.267644001 | 7.43E-05 |
| SH3BGRL2 | -0.633181146 | 0.000269164 |
| ZNF808 | 0.560499808 | 3.83E-05 |
| ANKRD35 | 9.135129598 | 1.02E-08 |
| YTHDF2 | -0.503819774 | 3.01E-07 |
| NBR2 | 1.018072076 | 0.000191179 |
| TMA16 | -1.696269987 | 3.27E-09 |
| ATL1 | 0.785588658 | 0.044571992 |
| CNGA1 | 2.455995194 | 6.85E-05 |
| MAFK | 0.587054726 | 9.67E-07 |
| ARMH1 | 5.289778199 | 4.45E-05 |
| GPN1 | -0.360265995 | 0.000187668 |
| ZNF28 | 0.927540201 | 9.15E-09 |
| ZNF511 | -0.392939023 | 0.000898643 |
| ZNF627 | 2.623861052 | 6.25E-12 |
| WDHD1 | -1.15766997 | 4.81E-08 |
| ZNF789 | 0.684517561 | 7.01E-06 |
| CTNND1 | -1.243090797 | 8.63E-11 |
| DDX39B | 0.170309084 | 0.034781507 |
| SLC34A3 | 1.620372896 | 0.000321545 |
| SH2D1B | -1.146082731 | 0.008744386 |
| ARC | 4.965428403 | 7.50E-14 |
| AC073343.1 | 2.605094285 | 0.000293753 |
| NUDT16 | -0.315379674 | 0.00016401 |
| TLK1 | -1.353554889 | 1.32E-07 |
| LRBA | 0.788604147 | 0.000287996 |
| BAZ1A | -1.167287734 | 9.53E-10 |
| COPS8 | -0.585843214 | 3.00E-06 |
| PPIAP22 | 0.227527768 | 0.000222419 |
| CCDC69 | -1.026676021 | 0.000363126 |
| MDM4 | -0.535520385 | 9.34E-06 |
| RYR2 | 0.628978677 | 0.000817545 |
| ZNF534 | 1.677301582 | 0.011595453 |
| KLHL9 | -1.865242342 | 2.76E-11 |

| | | |
|---|---|---|
| NCOA6 | -1.677652274 | 7.31E-11 |
| TAT | 8.33190418 | 2.64E-07 |
| C6orf89 | 0.17077772 | 0.008127746 |
| CALM1 | -0.550058788 | 2.97E-08 |
| TAFA2 | -0.889966839 | 0.001271366 |
| TTC37 | -0.125984236 | 0.031425735 |
| TUSC1 | -0.977761186 | 8.43E-07 |
| SLC9A6 | -1.10566942 | 2.56E-08 |
| MT-ND6 | 0.530222085 | 9.52E-09 |
| IPO9 | -0.230134378 | 0.000224384 |
| CEP290 | 1.090709332 | 1.05E-08 |
| SSBP3-AS1 | 1.443469954 | 0.000224464 |
| MT-CO2 | 0.67927105 | 1.16E-09 |
| TOGARAM1 | 0.379477986 | 0.000386056 |
| DLL1 | 4.756556575 | 0.001440212 |
| ANKRD13B | -0.281726891 | 0.021792527 |
| UNC13B | -0.308381708 | 0.000577588 |
| MT-CYB | 0.398937093 | 3.85E-05 |
| LDB1 | -1.116341707 | 2.77E-09 |
| PPP1R14C | -0.753171768 | 0.000212707 |
| CTR9 | -0.226380917 | 0.0008996 |
| SMOC1 | 1.564037454 | 3.65E-06 |
| F5 | 4.82744295 | 0.000213797 |
| ZNF652 | -0.280782251 | 0.01394602 |
| SMURF1 | -0.616544147 | 1.16E-06 |
| SLC5A3 | -0.919615384 | 3.29E-10 |
| MTCO3P12 | 0.136017054 | 0.028320213 |
| GPATCH3 | -0.678356231 | 8.87E-05 |
| PLXNB3 | 0.530545561 | 2.23E-05 |
| OXCT2 | -1.619487713 | 0.012036465 |
| RPL10A | 0.467852541 | 1.61E-07 |
| COLGALT2 | -0.700427782 | 0.00191103 |
| MT-ND2 | 0.213636446 | 1.31E-06 |
| FAM169A | -1.101573544 | 3.60E-08 |
| ZNF830 | -0.951221844 | 3.59E-07 |
| GRIN3A | 1.032874106 | 0.007601543 |
| MT-ND5 | 0.532268844 | 8.78E-07 |
| TMEM184B | -0.359873883 | 9.21E-05 |
| MTOR | -0.437942968 | 4.91E-06 |
| SCAMP5 | -0.446298067 | 0.000519052 |
| ZNF521 | 0.540530529 | 0.014924048 |
| ALPK2 | -2.052632766 | 2.09E-07 |
| LRIG2 | -0.226723408 | 0.008154245 |
| MT-CO1 | 0.264014231 | 4.61E-08 |
| PNP | -0.491927265 | 2.27E-06 |
| PAX9 | -0.333038514 | 0.005438496 |
| FOXJ3 | -0.366701103 | 7.90E-06 |
| ZNF358 | 0.423269753 | 0.000515834 |

| | | |
|---|---|---|
| SFT2D1 | -0.583362825 | 4.16E-05 |
| CD247 | 1.371434509 | 0.000490416 |
| CHAMP1 | 0.151297998 | 0.005364946 |
| ARHGAP11A | -1.901713918 | 7.82E-12 |
| HMGN2 | -0.156450596 | 0.001680588 |
| UBE2J1 | -1.153939734 | 2.27E-08 |
| GJC2 | 1.795970463 | 0.000393406 |
| OPA1 | -0.419713157 | 2.22E-05 |
| DENND4B | -0.459812262 | 2.27E-06 |
| RYR3 | -0.476122966 | 0.043018229 |
| ZNF277 | 1.021315253 | 1.94E-05 |
| MT-ND3 | 0.079795037 | 0.041968983 |
| KTI12 | -0.557617751 | 1.91E-05 |
| SELENOT | -1.000862974 | 1.96E-09 |
| ARHGEF15 | 7.00668098 | 2.14E-05 |
| RUSC2 | -0.92781889 | 2.07E-08 |
| FICD | 1.688176872 | 2.33E-08 |
| OSTC | -0.29575752 | 0.003942731 |
| HSD3BP5 | 0.480905404 | 0.009588979 |
| TSEN15 | -0.753427024 | 4.79E-08 |
| LTN1 | -0.173431463 | 0.005802737 |
| RUNDC1 | -0.502770328 | 4.24E-06 |
| CCDC152 | 0.937599893 | 0.028244676 |
| STKLD1 | 1.345921295 | 0.002963659 |
| TYW1 | -0.681886318 | 3.36E-06 |
| DCAF12 | -0.298431707 | 0.000119428 |
| SFMBT2 | -0.266392677 | 0.003162006 |
| ITPRIPL1 | -2.063766048 | 6.57E-09 |
| MT-ND4 | 0.652100434 | 6.12E-08 |
| SMC5 | -1.366440066 | 1.90E-10 |
| MT-ND1 | 0.130521855 | 0.000120971 |
| PRMT6 | -1.511835734 | 1.54E-10 |
| SHISA4 | 0.737659063 | 4.66E-05 |
| CIPC | -0.615543192 | 7.41E-06 |
| CAPZA2 | -0.498380726 | 8.16E-07 |
| MT-ATP6 | 0.303575312 | 1.98E-08 |
| TOP1 | -0.519083369 | 3.65E-07 |
| PRC1 | -0.997488787 | 1.80E-09 |
| MAP3K3 | -0.444552256 | 1.52E-05 |
| L1CAM | 0.359892347 | 2.06E-05 |
| SREBF2 | 0.361019839 | 4.58E-05 |
| C1orf174 | -0.42057517 | 4.67E-05 |
| SPOUT1 | -0.863802491 | 3.06E-08 |
| RPL39 | 0.386538577 | 6.37E-05 |
| DZIP3 | 1.558461939 | 7.22E-09 |
| KIAA0753 | -1.035093851 | 1.02E-07 |
| DCLRE1A | -1.052225044 | 7.31E-09 |
| ATG9A | 0.129597125 | 0.035351596 |

| | | |
|---|---|---|
| NOS1AP | -0.544769166 | 0.035215598 |
| APRT | 0.413420722 | 9.33E-05 |
| GPRASP1 | 1.831647264 | 9.60E-05 |
| MAGEE1 | -0.573090227 | 0.009274319 |
| MT-CO3 | 0.139443918 | 1.61E-05 |
| ZFP2 | 3.890815477 | 4.76E-05 |
| L3MBTL3 | -1.527299848 | 3.12E-05 |
| DMD | -0.450834971 | 0.046642318 |
| MFAP3L | 0.355505197 | 0.047446914 |
| SMG5 | -0.759657364 | 1.20E-08 |
| KIFBP | -1.289097599 | 7.21E-09 |
| TGM2 | 0.535037913 | 5.47E-07 |
| PJA2 | 0.653220692 | 4.59E-08 |
| RORB | -1.150016942 | 5.50E-07 |
| SGMS1 | -0.660247261 | 1.58E-05 |
| SNORA31 | 2.224506443 | 0.000225778 |
| RNU5A-1 | 2.598019414 | 0.002564468 |
| SNORA63 | 4.017099271 | 0.00020245 |
| RNA5S17 | 1.640916818 | 0.018436202 |
| RNA5S6 | 1.249399362 | 0.04356547 |
| SNORD14E | -0.828180346 | 0.024454192 |
| SNORA65 | 1.514361636 | 0.001452277 |
| RN7SKP275 | -5.169195626 | 0.000131941 |
| SNORA71C | -1.336827974 | 0.024591288 |
| RNU6-915P | 6.774837438 | 3.09E-05 |
| SNORA23 | 1.490264834 | 0.002637967 |
| AL162581.1 | 0.592639845 | 0.025039738 |
| SNORD6 | -1.974760704 | 0.002966795 |
| RNU6-652P | 2.21878436 | 0.005737006 |
| TDRKH-AS1 | 1.544144291 | 0.000443577 |
| ZNF525 | 0.750211653 | 1.36E-05 |
| POLH-AS1 | -2.546118747 | 0.000394116 |
| AC012454.1 | 6.962616657 | 2.04E-05 |
| LINC00449 | 1.396829463 | 0.00265958 |
| C6orf47 | -1.69902826 | 0.006965932 |
| AC083799.1 | -0.526074205 | 0.004974796 |
| COX20 | -1.101737257 | 1.39E-09 |
| CHML | -0.820200851 | 6.71E-09 |
| IBA57-DT | 1.507678473 | 0.003144567 |
| TATDN3 | 0.471207824 | 0.007424377 |
| SERTAD4-AS1 | -1.013933742 | 8.08E-05 |
| MIR29B2CHG | -1.746708207 | 3.06E-05 |
| RAET1G | 1.734569535 | 0.012191082 |
| C1orf53 | 0.62646226 | 0.024334189 |
| SAMD5 | -1.501555809 | 3.38E-06 |
| ECT2L | 2.308245485 | 0.000724799 |
| AL645568.1 | 1.285748967 | 0.004183287 |
| CENPW | -0.772050971 | 5.43E-05 |

| | | |
|---|---|---|
| MSTO2P | -0.169807637 | 0.019589245 |
| SPRN | -1.248777754 | 2.24E-05 |
| FAM229B | 0.828179301 | 0.007181166 |
| EEF1AKMT2 | -0.363854337 | 0.010789994 |
| CCDC162P | 3.943765005 | 0.019683275 |
| ADAMTSL4-AS1 | 4.918766992 | 3.96E-09 |
| H3C14 | 4.380107768 | 9.73E-12 |
| H2AC18 | 3.276321362 | 1.42E-09 |
| H2BC18 | 2.975494181 | 3.88E-10 |
| H3C15 | 3.194003112 | 7.35E-08 |
| HSD3BP4 | 2.907341738 | 1.44E-05 |
| ATP1A1-AS1 | 0.658939892 | 1.36E-05 |
| RBM20 | -1.706787787 | 3.31E-06 |
| C6orf163 | 2.360737641 | 0.01761627 |
| GDI1 | -0.116692785 | 0.030653909 |
| PCMTD2 | 0.73444301 | 2.21E-08 |
| SOX18 | -1.148857934 | 0.002101557 |
| LIME1 | -1.574341706 | 5.97E-07 |
| HSP90B3P | 0.519891353 | 3.09E-05 |
| LINC00632 | 2.390092427 | 2.67E-07 |
| SAMD13 | 2.668383458 | 0.00010125 |
| LINC01270 | 0.800699787 | 0.002178772 |
| LRRC73 | 1.221899488 | 0.001445361 |
| LINC00963 | 0.351358689 | 0.001589269 |
| SYS1 | -0.978329273 | 4.51E-09 |
| INPP5B | -0.46478373 | 0.001336394 |
| MAFB | 2.056758156 | 3.05E-05 |
| TRAF3IP1 | -0.27056789 | 0.019168778 |
| CHIC1 | -2.121959635 | 9.56E-10 |
| NAP1L6P | 2.007205247 | 4.16E-06 |
| GIGYF2 | -1.01344256 | 6.40E-09 |
| RUFY2 | -0.849423461 | 5.32E-05 |
| GGTA1P | 1.688069616 | 0.001582126 |
| PHACTR4 | -1.639096694 | 4.15E-10 |
| ASAH2B | -1.466794508 | 2.06E-07 |
| AGAP6 | -1.348781941 | 2.64E-07 |
| TIMM23B | -0.780943536 | 3.02E-05 |
| ZDHHC18 | -0.907182598 | 4.61E-07 |
| AGAP9 | -0.681888848 | 0.000579712 |
| BMS1P1 | -0.612233254 | 0.009442063 |
| MACO1 | 0.51757317 | 1.95E-05 |
| GDF5OS | 5.552487361 | 2.13E-05 |
| ZDBF2 | 1.263037872 | 6.33E-10 |
| RPL12P1 | 0.965369792 | 0.009357892 |
| RPL12P16 | 0.575487401 | 6.95E-08 |
| KIFC1 | -1.581329287 | 2.64E-08 |
| DAXX | -0.780457912 | 2.14E-07 |
| BMPR2 | -0.95740463 | 4.76E-08 |

| | | |
|---|---|---|
| PFDN6 | 0.371552306 | 0.006231308 |
| RXRB | -0.295148114 | 0.020147727 |
| OXLD1 | 0.736674802 | 2.74E-05 |
| LINC02731 | 2.946189097 | 9.98E-05 |
| COL11A2 | 0.963329142 | 0.004781982 |
| HLA-DOA | 0.659158406 | 0.002039615 |
| HNRNPCP2 | 0.16769434 | 0.000623205 |
| BRD2 | -1.549728948 | 2.54E-07 |
| COL5A2 | 0.421279952 | 1.81E-06 |
| SPIN3 | -0.583380824 | 0.000356479 |
| NBDY | 0.285643936 | 0.021275849 |
| TNRC6C-AS1 | 2.412000093 | 5.54E-07 |
| HLA-DRA | 0.313062527 | 0.016376031 |
| PJVK | 3.694503737 | 4.32E-07 |
| MRPL38 | -0.135038163 | 0.041630507 |
| STK19 | -0.580306694 | 0.047592269 |
| DXO | -0.941791185 | 0.003809532 |
| SDHD | -0.338640834 | 3.51E-05 |
| PKP4-AS1 | -1.778260367 | 0.000760159 |
| XAGE1B | 0.260141997 | 0.015246568 |
| HSPA1B | -0.480442811 | 5.26E-06 |
| MBD5 | -0.328929855 | 0.040289274 |
| GPANK1 | 0.498007669 | 0.01140397 |
| FAM155A | -1.37765246 | 2.87E-05 |
| BAG6 | 0.296055935 | 0.00200374 |
| PRRC2A | -0.326328726 | 0.000594638 |
| ZNF814 | -0.34048214 | 6.41E-05 |
| ZNF551 | -0.297738985 | 0.000193151 |
| MICA | 0.447249211 | 0.046177919 |
| ZNF805 | 1.118805159 | 6.73E-06 |
| HLA-C | 1.013230278 | 0.001809775 |
| AL450124.1 | 1.306148316 | 0.031684566 |
| PPP1R10 | 2.246798447 | 3.14E-12 |
| ABCF1 | -0.519165386 | 0.004773171 |
| DDR1 | 0.399749366 | 0.003200726 |
| GNL1 | 0.417439995 | 0.00089058 |
| HLA-E | 0.394841263 | 0.010048324 |
| ZNF468 | 0.518499405 | 1.11E-05 |
| ZNF616 | 1.62395049 | 1.46E-09 |
| DISP3 | 2.592966662 | 0.002717503 |
| RACK1 | 0.23391536 | 2.25E-05 |
| TBC1D8 | -1.07648611 | 7.11E-08 |
| LINC02210 | -0.867013573 | 0.009581054 |
| CBY3 | 3.095496065 | 0.019217144 |
| C5orf60 | 3.696830633 | 0.000209964 |
| AC010624.1 | 6.917165493 | 2.09E-05 |
| FAM153CP | 4.289649572 | 1.22E-05 |
| GABBR1 | -0.61844279 | 0.016665893 |

| | | |
|---|---|---|
| MIR1915HG | -0.96500654 | 0.001133415 |
| STARD7-AS1 | -0.654393487 | 0.00035982 |
| TRIM27 | -1.168833054 | 8.92E-08 |
| AC008429.1 | 2.113983246 | 0.000171044 |
| FOXD4L5 | 1.185017374 | 0.047516585 |
| ZNF204P | 0.832315111 | 0.003040385 |
| AL590399.1 | 0.623268356 | 0.013100774 |
| TTC25 | 0.866939637 | 0.001284049 |
| ADGRF5P2 | 3.414959704 | 1.05E-05 |
| MRPL53 | 0.246683579 | 0.001134564 |
| ATXN2 | -0.919703824 | 1.06E-07 |
| TCTN1 | 0.699934938 | 5.02E-06 |
| ZBTB48 | 0.791117896 | 1.96E-06 |
| FAM201A | -0.90785945 | 0.001418669 |
| IGFL4 | 6.774837438 | 3.09E-05 |
| MZT1 | -0.59829533 | 1.06E-05 |
| ZNF155 | 0.543931858 | 0.000214065 |
| UQCC3 | 0.742694585 | 0.00010041 |
| FBXO48 | 2.37794274 | 1.77E-09 |
| ATP6V0E2-AS1 | 1.052847573 | 1.97E-05 |
| ZNF783 | -0.653639708 | 2.37E-05 |
| ZNF425 | 5.091572929 | 6.44E-13 |
| BLACE | 8.232933498 | 3.14E-07 |
| PCDHA9 | -1.326072498 | 5.05E-09 |
| PCDHA8 | -1.146567047 | 3.32E-08 |
| PCDHA7 | -1.012419717 | 1.91E-08 |
| PCDHA5 | -1.376935603 | 1.99E-09 |
| PCDHA4 | -1.268806475 | 5.65E-09 |
| PCDHA2 | -1.06761222 | 2.15E-08 |
| PCDHA1 | -1.424884733 | 2.92E-09 |
| MS4A13 | 3.095162029 | 4.68E-07 |
| SPIRE2 | 0.677475539 | 0.00073737 |
| AARD | 0.654759512 | 0.002604225 |
| PKHD1L1 | 4.548281308 | 0.000506695 |
| SLFN12L | -1.19215011 | 0.018595197 |
| LINC02397 | 4.675601922 | 0.000500779 |
| SLC35B4 | -1.475279365 | 2.39E-10 |
| TMEM231 | 0.389600509 | 0.001566999 |
| FAM205A | 7.582561345 | 2.60E-06 |
| C4orf47 | 3.253067621 | 0.000373262 |
| SDHAF1 | -0.389412838 | 0.010616775 |
| ARID3C | 4.591400514 | 0.00044473 |
| PSENEN | 0.378360054 | 0.000121479 |
| C11orf91 | 1.764985188 | 4.09E-05 |
| LINC00654 | 4.510573251 | 0.000718725 |
| ZBTB10 | 0.201509244 | 0.003458792 |
| C4orf46 | -0.544702196 | 2.61E-05 |
| CCDC144NL | -1.881901485 | 3.60E-08 |

| | | |
|---|---|---|
| LGR4 | 0.249050912 | 0.013888072 |
| KRT17P7 | 4.589481122 | 0.000472928 |
| AC105052.1 | 0.373964594 | 0.035729373 |
| SPDYE2 | -0.954918804 | 0.000321904 |
| E2F4 | -0.990824532 | 3.26E-09 |
| KRT17P5 | 7.845177365 | 1.87E-06 |
| PDE7A | -1.194599508 | 1.07E-08 |
| SNX2 | 0.202589308 | 0.015868317 |
| SAP25 | 1.197462777 | 0.005468208 |
| NT5M | 0.965385155 | 0.007509636 |
| ADGRG1 | 0.741219658 | 8.77E-08 |
| IPO7 | -0.41858924 | 5.39E-07 |
| PRR13 | 0.398152959 | 0.000132106 |
| TECPR1 | -0.192026458 | 0.046343664 |
| CFI | 0.580558422 | 0.005746853 |
| SAMD9 | -1.311077016 | 8.38E-10 |
| KRT81 | 0.298901883 | 0.019209575 |
| TP53TG3D | 1.382224339 | 0.000776283 |
| TP53TG3C | 1.019921382 | 0.006283995 |
| ATP6AP1L | 3.978077063 | 4.94E-08 |
| CCDC85C | -0.433534541 | 4.70E-06 |
| SPDYE18 | -1.044706579 | 0.028577485 |
| AC004980.1 | 0.435251795 | 0.023459996 |
| NAP1L4 | -1.042160574 | 1.83E-06 |
| SMG1P2 | 0.414013461 | 0.000401406 |
| TMEM256 | 0.820720746 | 4.01E-05 |
| CHKB-DT | 3.577019414 | 6.76E-07 |
| CPT1B | -0.460338342 | 0.007805177 |
| SMN2 | -0.472004507 | 0.001912434 |
| POM121B | -0.782228825 | 2.37E-05 |
| HMGN1 | 0.28093386 | 3.51E-05 |
| EIF3CL | 0.150745273 | 0.006178524 |
| LINC01446 | -0.867236429 | 8.47E-06 |
| LCMT1 | -0.377639924 | 0.019603777 |
| LINC00898 | 5.199348126 | 0.000122465 |
| MFSD2B | 1.210811448 | 1.42E-05 |
| CDPF1 | 0.689160113 | 0.001283764 |
| SMIM11A | 0.893399945 | 0.000205294 |
| DPF3 | 1.177695394 | 0.000614512 |
| ETFRF1 | 0.31782652 | 0.002013288 |
| C17orf107 | 2.568405745 | 5.70E-06 |
| ITSN1 | -0.215757147 | 0.024270557 |
| ITPRIPL2 | -0.49414712 | 1.06E-06 |
| AL359878.1 | 1.532308579 | 0.002547964 |
| C5orf51 | -0.426171375 | 5.55E-06 |
| CATSPER2P1 | 0.99100384 | 0.000220333 |
| GAGE1 | 1.046536352 | 0.005054464 |
| LOH12CR2 | 1.358180416 | 0.032095811 |

| | | |
|---|---|---|
| CYS1 | 6.708599568 | 4.88E-10 |
| PLPP6 | -0.313931696 | 0.006159127 |
| GMNC | 2.256413852 | 0.001533207 |
| TTC23L | 4.945625199 | 0.005398171 |
| RFPL3S | 3.471313827 | 1.11E-05 |
| RPS3AP47 | 0.422693029 | 0.000462139 |
| ZNF316 | -0.485874737 | 7.08E-05 |
| SRRM2-AS1 | 3.758184751 | 3.42E-08 |
| OLIG2 | 0.756488642 | 0.000878647 |
| RNPS1 | -0.395166233 | 5.39E-06 |
| HSP90AB2P | 0.422165838 | 2.86E-05 |
| AC105345.1 | 5.594161384 | 2.41E-05 |
| DNAJC19 | -0.338815415 | 0.042808876 |
| JPT2 | -0.59309711 | 5.33E-08 |
| SERPINB5 | 2.621009368 | 0.000369161 |
| TMEM191C | 1.575272563 | 0.047843698 |
| HERC2P9 | -0.251731569 | 0.011867124 |
| Z69890.1 | 3.024876857 | 6.25E-06 |
| ATP10A | -0.350855866 | 0.027097732 |
| DUXAP8 | -0.189113536 | 0.009968431 |
| TSSK2 | -1.570541933 | 0.000291365 |
| HNRNPA1P4 | 0.648558519 | 5.16E-08 |
| NELFE | 0.538982222 | 0.000771077 |
| RGL2 | 0.670866762 | 0.002757416 |
| TAP2 | -0.476616049 | 0.007987281 |
| HCP5 | 0.935853254 | 0.024569333 |
| HCG27 | 2.687362682 | 0.002994183 |
| ZBTB12 | -0.718154695 | 0.040922506 |
| EHMT2 | -0.399711048 | 0.007558784 |
| HSPA1L | 0.979638308 | 0.00192136 |
| CLIC1 | 0.311771753 | 0.026000875 |
| H1FX-AS1 | 0.407152557 | 0.044466919 |
| RAB12 | -0.799132903 | 1.50E-07 |
| HLA-B | 0.397759993 | 0.025167631 |
| NRM | 0.654995426 | 0.025234997 |
| DHX16 | -0.412146646 | 0.012563517 |
| HLA-A | 0.391667813 | 0.003300215 |
| HLA-F-AS1 | 1.755361346 | 0.009337352 |
| HACD2 | -1.26274982 | 3.82E-10 |
| CFAP44 | 0.893498629 | 6.65E-06 |
| LNP1 | 2.055710389 | 1.13E-08 |
| VGLL3 | -1.465889566 | 3.02E-05 |
| ZCWPW2 | 5.600727652 | 9.12E-05 |
| ANKRD28 | -1.044689283 | 1.16E-09 |
| COLQ | 3.432069133 | 3.87E-07 |
| THUMPD3-AS1 | 0.776869801 | 6.84E-08 |
| RNU1-27P | 1.574753032 | 0.015732015 |
| SNORA22B | 1.013122573 | 0.036657712 |

| | | |
|---|---|---|
| AC106802.1 | 6.8249165 | 2.75E-05 |
| SNORD101 | -1.873151119 | 0.000644839 |
| RNU6-36P | 1.225099525 | 0.044467445 |
| RNU1-2 | 1.700008315 | 0.000296962 |
| SNORA15B-1 | 1.72079423 | 0.014577426 |
| RNVU1-15 | 5.961886984 | 1.21E-09 |
| SNORA2B | 2.823785539 | 7.94E-05 |
| SNORD15B | -3.54123046 | 1.76E-05 |
| RNVU1-14 | 5.516485674 | 1.03E-07 |
| SNORA66 | 1.257031177 | 0.005021612 |
| MIR621 | -1.061391584 | 0.00053255 |
| MIR616 | 4.291576734 | 1.40E-05 |
| SNORD12C | -3.973289712 | 0.003894643 |
| SNORD83A | -1.289781486 | 0.005691193 |
| SNORA48 | 4.551737182 | 3.64E-06 |
| MT-TF | -0.589859789 | 0.001364687 |
| MT-RNR2 | 0.174978797 | 1.02E-05 |
| MT-TQ | 0.53167094 | 0.000763467 |
| MT-TW | -0.402812283 | 0.046341191 |
| MT-TY | -0.264579899 | 0.00102029 |
| MT-TS1 | 0.530509597 | 4.71E-06 |
| MT-TD | -0.677034962 | 0.022876998 |
| MT-TG | -0.380448864 | 0.007643131 |
| MT-TR | -0.795384133 | 0.0009708 |
| MT-TH | -0.235875629 | 0.036347063 |
| MT-TS2 | -0.363582393 | 0.012862642 |
| MT-TL2 | -0.33191542 | 0.013773109 |
| MT-TT | -0.570419865 | 0.000166852 |
| MT-TP | 0.45361369 | 3.41E-05 |
| GPX3 | 0.393178465 | 0.014373795 |
| SELENOH | 0.15527593 | 0.014593447 |
| DIO1 | 3.937956649 | 1.52E-05 |
| STK38L | -0.265227866 | 0.000486601 |
| SACM1L | -1.211929325 | 9.88E-10 |
| MT-RNR1 | -0.371614554 | 1.54E-07 |
| TSN | -0.812674935 | 6.12E-08 |
| SLC48A1 | -0.306761275 | 0.0111707 |
| PRR22 | -0.732260789 | 0.004766703 |
| SNORD67 | -1.104310099 | 0.040917683 |
| SNORD17 | -1.009599659 | 1.97E-06 |
| SNORA53 | 1.875759936 | 0.000184127 |
| ZRSR2P1 | 0.413180375 | 0.00686245 |
| AC064799.1 | 0.256439819 | 5.10E-05 |
| LINC01089 | 0.332039572 | 0.028552914 |
| LINC02693 | -1.324310996 | 5.81E-10 |
| KRTAP2-3 | -5.479573732 | 9.31E-08 |
| LINC02656 | 7.640868051 | 2.20E-06 |
| RTL8B | -0.876032704 | 0.001129192 |

| | | |
|---|---|---|
| ST13P5 | 0.235378756 | 0.008607645 |
| RPL15P3 | 0.233039563 | 0.00216259 |
| MT-ND4L | 0.55194108 | 3.06E-10 |
| RPL23AP4 | 2.529283459 | 4.55E-07 |
| BX088651.1 | 1.331799547 | 0.036903538 |
| AC016747.1 | 0.658985769 | 0.000355102 |
| PTTG3P | -0.284797398 | 0.028229138 |
| ZNF580 | 0.697906231 | 1.13E-05 |
| ZNF611 | 0.300851708 | 0.00110049 |
| SYT3 | 0.902825035 | 0.025781938 |
| NUP62 | -1.73026097 | 4.00E-11 |
| COX20P1 | -0.651326689 | 0.000550244 |
| AL117350.1 | -1.609565206 | 0.001750288 |
| DENND1B | -2.536822355 | 1.13E-09 |
| C1orf220 | -1.449362092 | 0.000397302 |
| Z99572.1 | 1.307847937 | 5.08E-05 |
| SFT2D2 | -1.004589918 | 9.37E-10 |
| FGFR1OP | -0.574102023 | 3.66E-05 |
| AC092447.1 | -1.075547425 | 0.012510747 |
| HMGB3P6 | -0.684888568 | 0.001257679 |
| RPL31P11 | 0.756711482 | 0.018808697 |
| SCAF8 | -0.999550953 | 4.95E-09 |
| CFAP45 | 3.487500722 | 4.30E-07 |
| AL356535.1 | 0.558292149 | 0.001044261 |
| AC007256.1 | -0.838954499 | 0.001384649 |
| AC092279.1 | -0.326172533 | 0.000771077 |
| NPM1P46 | 0.435736569 | 4.60E-06 |
| BTF3L4P3 | -0.864277579 | 0.045102661 |
| AL021939.1 | -0.622357376 | 0.032120583 |
| TCTEX1D2 | 0.780714668 | 0.04478433 |
| CRYGS | 1.030286678 | 0.019381018 |
| RPL7AP60 | 0.915847778 | 0.048849961 |
| KLHL23 | -2.137033417 | 1.43E-09 |
| RPL22P1 | 0.404841078 | 4.07E-06 |
| AP003040.1 | 0.483169392 | 0.044906686 |
| TRIM59 | -1.001583885 | 7.68E-10 |
| MLLT11 | 0.176647846 | 0.004083773 |
| ASIC3 | 0.918640566 | 0.000734481 |
| AL049697.1 | -0.563971882 | 0.001254799 |
| CCDC183 | 0.962103727 | 0.006742829 |
| RPL12P38 | 0.621108853 | 1.10E-05 |
| PPP1R2P10 | -1.02608782 | 0.001244078 |
| EEF1A1P16 | 0.353680844 | 0.000422415 |
| NPM1P32 | 0.245172854 | 0.025476054 |
| H3P4 | 1.930977592 | 0.000663275 |
| SUPT4H1 | -0.58620119 | 2.65E-05 |
| RPL12P42 | 0.601287916 | 0.023924627 |
| EEF1B2P6 | 0.417948353 | 8.87E-05 |

| | | |
|---|---|---|
| AC090114.1 | 0.367397075 | 0.000886009 |
| NRAS | -0.614478529 | 1.28E-07 |
| ZNF625-ZNF20 | 1.247278808 | 1.78E-08 |
| HNRNPA3P6 | 0.257975011 | 0.00023762 |
| AC008481.1 | 0.463628262 | 0.02313824 |
| AC008481.2 | 3.33446931 | 0.000374508 |
| RPL7AP34 | 0.40778257 | 0.036463939 |
| AL122020.1 | 0.396173792 | 0.000493723 |
| RPS7P11 | 0.31420816 | 4.45E-05 |
| NPM1P50 | 1.558216793 | 0.030377165 |
| QTRT1 | -0.249291778 | 0.008691183 |
| CHUK | -1.036616299 | 2.52E-09 |
| PCNPP3 | 1.430835627 | 3.38E-05 |
| MXD3 | 0.195727193 | 0.001529129 |
| RPS3P6 | 0.555577488 | 0.001775859 |
| GSTM2 | 1.123759337 | 1.77E-06 |
| NAP1L1P3 | 0.169235634 | 0.041926768 |
| COG8 | -0.614943 | 3.17E-07 |
| AC105052.2 | 0.567447221 | 0.014084029 |
| ARHGAP19 | -1.68428742 | 1.03E-10 |
| LCAT | 0.663618879 | 0.000753259 |
| MAGEA12 | -0.382191918 | 4.08E-05 |
| PTPRCAP | 1.90262043 | 0.000681838 |
| ANXA2P1 | 0.402546655 | 0.011625255 |
| GPC2 | 0.992098612 | 0.02170594 |
| HSPD1P1 | -0.203345722 | 0.001038371 |
| RPLP1P6 | 3.715269047 | 2.81E-07 |
| VTI1BP2 | 2.784795404 | 1.83E-05 |
| RPL18AP3 | 0.434527836 | 2.73E-07 |
| AC007068.1 | 0.606126401 | 0.004489516 |
| SIPA1 | 0.491456335 | 2.07E-05 |
| FTH1P3 | 0.207278696 | 0.030910513 |
| ERV3-1 | 1.520813592 | 1.89E-08 |
| SYNJ2BP | -0.899777655 | 4.10E-08 |
| ARL2 | 0.357798073 | 0.000101729 |
| TTLL13P | 3.425134233 | 3.05E-07 |
| AL731556.1 | 0.622169207 | 0.014737892 |
| AC132008.1 | 7.08299031 | 1.30E-05 |
| SRA1 | -0.190109496 | 0.005558729 |
| STIMATE | -1.178811716 | 5.61E-09 |
| RARRES2P2 | 3.346964536 | 0.000230784 |
| DNAJC9 | -1.558096893 | 1.20E-10 |
| RPLP0P6 | 0.515757301 | 5.78E-09 |
| C8orf82 | 0.714017665 | 0.000220077 |
| AL390039.1 | -0.341269288 | 0.023526004 |
| CPLX3 | 4.054513976 | 6.10E-06 |
| ZBTB9 | 0.359239553 | 0.010678301 |
| TMX2 | -0.169663445 | 0.005496984 |

| | | |
|---|---|---|
| AL049873.1 | 0.475237163 | 7.25E-05 |
| SLX1A-SULT1A3 | -0.686568657 | 1.66E-07 |
| RPL7AP50 | 0.385008705 | 3.34E-05 |
| RPL11P3 | 0.974319131 | 0.002013516 |
| HEXA | 0.196394086 | 0.023075061 |
| RPSAP54 | 0.332635866 | 0.000713654 |
| LEPROT | -0.36273333 | 6.27E-05 |
| LBH | 2.283168358 | 1.93E-07 |
| ADAT3 | 0.594542948 | 0.013796284 |
| PPP1CB | -0.677219554 | 2.06E-08 |
| SULT1A4 | -0.752300187 | 0.008081852 |
| LAT | 1.427489198 | 3.14E-05 |
| PGGT1BP1 | 1.00959777 | 0.014657492 |
| NCKIPSD | -0.51880257 | 1.86E-05 |
| SLC25A5P3 | 0.610678312 | 0.003220017 |
| LDHBP2 | 0.180197884 | 0.042795937 |
| SEC14L1P1 | -0.563793758 | 0.00703055 |
| S1PR3 | 2.406731352 | 1.93E-09 |
| SLC35F6 | -0.257448345 | 0.001291529 |
| RPL17P50 | 0.958026185 | 0.000778599 |
| AL590762.2 | 4.927726177 | 0.000247783 |
| PIGCP1 | -2.188665483 | 6.64E-08 |
| FAM209B | 3.68778687 | 4.83E-05 |
| DDAH2 | 0.822707137 | 0.031471111 |
| RPS29 | 0.40485036 | 1.32E-05 |
| ZNF337-AS1 | -0.262235911 | 0.003376228 |
| CENPBD1P1 | 0.411550638 | 3.47E-05 |
| AC020898.1 | 0.401868201 | 0.000126982 |
| ZNF134 | -0.822081017 | 5.43E-08 |
| ACTBP2 | -0.343703303 | 0.002320147 |
| DDX47 | -0.773668423 | 3.13E-08 |
| ZNF845 | 1.352693437 | 6.06E-10 |
| TMX2P1 | -0.176444051 | 0.01487652 |
| EMP2 | -0.462215735 | 3.68E-06 |
| KCTD11 | 1.377168663 | 4.35E-08 |
| RPL21P75 | 0.286968722 | 7.11E-05 |
| AC044787.1 | 0.426983192 | 1.42E-05 |
| EEF1B2P2 | 0.481607649 | 0.0077252 |
| YBX1P10 | 0.20041981 | 0.000599332 |
| AC092798.1 | 3.882899249 | 1.63E-07 |
| NPM1P6 | 0.482947799 | 5.50E-07 |
| RPL13AP7 | 0.496433101 | 1.14E-05 |
| LINC01521 | 0.731300326 | 0.009442063 |
| PPM1N | 1.422314554 | 0.000580254 |
| CEACAM16 | 7.586974944 | 3.21E-06 |
| SLC23A3 | 0.335575063 | 0.017646104 |
| DNASE1 | 0.861713857 | 2.49E-08 |
| MDP1 | 1.222831973 | 2.33E-05 |

| | | |
|---|---|---|
| CSNK1E | -0.495746356 | 2.24E-06 |
| GALT | 0.346165686 | 0.004186529 |
| CLDN9 | 4.83236174 | 0.00119298 |
| AC091153.1 | 0.498511173 | 7.75E-06 |
| AC005840.1 | 0.437305616 | 0.001698703 |
| KRT18P17 | 1.212206611 | 0.000128869 |
| DNAJB1P1 | 2.118001397 | 4.39E-07 |
| ITGA1 | 0.128161627 | 0.022990534 |
| AC019080.1 | 2.975692052 | 2.46E-07 |
| NUDT19 | -0.945104511 | 2.18E-07 |
| ZNF726 | 0.939915143 | 3.71E-06 |
| ZNF99 | 3.784819935 | 6.03E-06 |
| AC010615.1 | -1.138325099 | 0.014545341 |
| TAX1BP3 | -0.468302568 | 1.71E-05 |
| RPL7AP14 | 2.850405829 | 0.001144659 |
| AP1G2 | 0.858842286 | 2.64E-06 |
| ZNF90 | 0.350597086 | 4.19E-05 |
| NAXD | -0.845817742 | 3.94E-08 |
| PGAM1P7 | -1.290356069 | 0.027148538 |
| MEF2B | 1.793797766 | 5.50E-06 |
| GANC | 0.597956964 | 0.00011449 |
| RPSAP61 | 0.389270882 | 0.000196104 |
| RRM2P3 | -1.405601081 | 5.06E-07 |
| AL034370.1 | -1.57732294 | 6.57E-08 |
| TTLL3 | 1.39536618 | 6.34E-08 |
| REPIN1 | -0.157361427 | 0.026045628 |
| ZNF891 | -0.807147885 | 5.98E-06 |
| SMIM7 | -0.748996241 | 8.97E-08 |
| UCA1 | 0.524240147 | 0.002412507 |
| AC011999.1 | 4.18755448 | 0.002446956 |
| CPSF1P1 | 2.044590053 | 8.14E-07 |
| GNAQP1 | -0.576142529 | 0.006394915 |
| CPNE1 | 0.331787581 | 9.62E-05 |
| PAXIP1-AS2 | -0.779642282 | 0.000641058 |
| TPT1P5 | 0.7141839 | 0.000395428 |
| LYRM4 | -0.528940663 | 0.000873961 |
| MYCBP | -0.695948358 | 4.96E-07 |
| AC094103.1 | -2.680400917 | 2.58E-05 |
| AC132008.2 | -0.714364074 | 0.000307695 |
| AMZ2P1 | 0.928865867 | 4.25E-05 |
| XPOTP1 | -0.342400333 | 0.000120775 |
| ZNF788P | -1.326480867 | 9.80E-10 |
| TTC41P | 1.089675421 | 0.004313143 |
| EEF1A1P12 | 0.406174825 | 8.77E-06 |
| RPS4XP1 | 0.458091922 | 0.000271549 |
| KRT18P10 | 0.532317706 | 0.006119031 |
| C19orf38 | 4.220005007 | 4.38E-07 |
| HNRNPA1P10 | 0.628929817 | 2.73E-09 |

| | | |
|---|---|---|
| C17orf67 | 1.642256998 | 0.000330926 |
| SETP21 | 1.168623293 | 0.049929394 |
| AC010336.1 | 0.200208114 | 0.048179764 |
| FIS1 | 0.357458903 | 0.000597943 |
| ANKRD36BP1 | -1.480575755 | 5.97E-09 |
| KCTD9P4 | -0.914634356 | 0.016567203 |
| AC124312.1 | 0.266142194 | 0.009636526 |
| AGGF1P1 | 2.92723251 | 1.66E-07 |
| SCART1 | 2.871854458 | 4.78E-10 |
| KRT8P14 | 2.680031568 | 0.00117581 |
| APTR | 2.057671507 | 1.05E-07 |
| SPDYE3 | -0.64089601 | 0.004680197 |
| SOGA3 | 2.663915371 | 6.83E-06 |
| AC133644.1 | -1.19296189 | 0.014502282 |
| HAUS3 | -0.797944986 | 2.64E-07 |
| VSTM5 | 4.410478121 | 0.000790727 |
| RPS3AP26 | 0.340989284 | 1.14E-06 |
| TUBAP2 | -0.371071267 | 2.33E-05 |
| LRRC37A4P | -0.62589473 | 0.011367858 |
| VPS33B-DT | 4.217152062 | 4.55E-06 |
| FAM187A | 1.661770292 | 2.55E-06 |
| RCN1P2 | 0.11531944 | 0.045743318 |
| TPT1P6 | 0.378118306 | 0.000735229 |
| RPL7P1 | 0.253694176 | 5.29E-06 |
| KRT42P | 8.838335935 | 2.75E-08 |
| PPME1 | -0.56757677 | 3.17E-07 |
| STARD10 | 1.059869202 | 0.000139635 |
| ZNF705E | 3.948559011 | 8.96E-07 |
| RPS15AP1 | 0.384435405 | 2.01E-05 |
| GTF2IRD2P1 | 2.428198831 | 4.92E-07 |
| NUTM2D | 4.28524062 | 8.85E-11 |
| RPS19P1 | 0.443574426 | 0.002960962 |
| RPSAP6 | 1.84287058 | 0.044501766 |
| B3GNT10 | 2.318164693 | 5.54E-08 |
| IFRD2 | -0.530963663 | 9.70E-07 |
| CAPN14 | -1.892920449 | 0.005571274 |
| AC005562.1 | -0.983001275 | 2.22E-06 |
| HNRNPUL2 | -0.634923835 | 1.04E-08 |
| CSKMT | 3.738638447 | 2.69E-13 |
| SEPTIN7P2 | 1.155655439 | 2.40E-06 |
| AL161756.1 | 1.049009702 | 0.044887019 |
| POLR2J4 | 1.030122467 | 6.64E-07 |
| AC010468.1 | 0.442742236 | 7.43E-06 |
| NPM1P31 | 0.569967908 | 0.017545375 |
| CYCSP55 | -0.9747935 | 0.000465808 |
| MPRIPP1 | -0.285600579 | 0.018481759 |
| DDX12P | -0.443329311 | 0.002297355 |
| UPF3AP2 | 0.588119958 | 0.000407018 |

| | | |
|---|---|---|
| RAD51AP2 | 4.813598642 | 1.06E-06 |
| EVPLL | 9.535149972 | 2.81E-09 |
| SMTNL1 | 6.667999476 | 1.63E-06 |
| RPS9P1 | 0.939244895 | 0.000442854 |
| LINC01588 | 1.777805471 | 2.68E-05 |
| RPL23AP3 | 0.592250649 | 0.001446268 |
| ZSWIM7 | 0.769797711 | 0.000334889 |
| ARHGEF28 | -0.383764822 | 0.001028066 |
| TBC1D26 | 7.465232185 | 3.87E-06 |
| PPIAP29 | 0.315461326 | 5.67E-05 |
| AKAP17BP | 3.342662898 | 0.000266688 |
| HSPD1P7 | 1.343088686 | 0.001342162 |
| CHCHD2P2 | 0.250873836 | 0.047395073 |
| DNAJA1P3 | -0.460406984 | 0.001409878 |
| RTL10 | -0.678713621 | 8.66E-06 |
| PHB2 | 0.199786834 | 0.00184441 |
| AL008729.1 | 1.022389705 | 0.016822629 |
| RPL13P12 | 0.390655298 | 8.25E-06 |
| GNL3LP1 | -0.596096491 | 0.015689054 |
| FDPSP5 | 0.751000423 | 0.018786774 |
| CD27-AS1 | 1.431834601 | 1.54E-07 |
| NEURL4 | -0.486183457 | 4.42E-06 |
| ALOX12-AS1 | 1.328440164 | 7.45E-05 |
| NPM1P24 | 0.45000846 | 3.30E-05 |
| KRT18P11 | 0.326902165 | 0.020961686 |
| TERF1P4 | -0.805254523 | 0.000799939 |
| UBXN2B | -0.731273646 | 9.68E-08 |
| BX322639.1 | 0.719476008 | 3.50E-06 |
| PRSS41 | 7.001990739 | 1.60E-05 |
| AC141586.1 | 0.388122677 | 0.002935432 |
| AC138409.2 | -2.246325377 | 2.55E-07 |
| NLRP2B | 7.579400771 | 4.62E-06 |
| MUC5AC | 1.43443915 | 0.009758997 |
| MSMP | -0.527751104 | 0.0341161 |
| GOLGA6B | 4.894534474 | 0.000262568 |
| FAM166B | 4.402632665 | 9.17E-06 |
| LINC00680 | 0.401444728 | 0.004370481 |
| PEX26 | -0.622100485 | 4.31E-07 |
| AL353795.1 | 0.795023979 | 0.040283748 |
| RBMXP2 | 0.369199149 | 0.00021896 |
| UBE2QL1 | 2.038465532 | 8.58E-07 |
| LINC02449 | 1.092751081 | 0.00130131 |
| LINC02649 | 2.121977899 | 0.004048457 |
| FASTKD5 | -0.388210741 | 2.26E-05 |
| GOLGA8B | 0.848336625 | 2.81E-08 |
| DHRS4-AS1 | 1.425278708 | 5.08E-08 |
| AP000533.2 | -1.645321019 | 3.58E-05 |
| AP000523.1 | 0.520671326 | 0.001590218 |

| | | |
|---|---|---|
| HOMEZ | 2.110676986 | 2.34E-08 |
| DDX3X | -0.853310482 | 1.61E-10 |
| AC127502.1 | -0.966383785 | 0.022061528 |
| NPM1P12 | 0.580136483 | 0.000930346 |
| EIF5P1 | -0.736132854 | 0.001139702 |
| HSPA1A | 0.270200552 | 0.002448841 |
| KRT18P23 | 1.576460286 | 0.025424456 |
| SLC25A5P1 | 0.652579013 | 0.005581518 |
| MRPL3P1 | -0.425723449 | 0.000185159 |
| MYL5 | 0.423899093 | 0.001955064 |
| ACTG1P3 | 1.571739659 | 0.006072142 |
| MIR17HG | -1.847127745 | 1.26E-08 |
| ZNF407 | -0.237321814 | 0.016120398 |
| MCM3AP-AS1 | -0.560243566 | 0.000384649 |
| NPEPL1 | 0.439347925 | 0.003436798 |
| ZNF663P | 6.018801353 | 7.90E-06 |
| RPL17-C18orf32 | 0.278320497 | 0.000277426 |
| BCRP3 | 2.026157887 | 0.000866946 |
| CALM2P3 | -0.640507126 | 1.54E-05 |
| HNRNPA1P7 | 0.610836642 | 2.45E-09 |
| AC007731.2 | 1.069884009 | 9.38E-06 |
| IFIT1P1 | 1.854452578 | 0.007309542 |
| RPS4XP4 | 7.306404312 | 6.21E-06 |
| EFCAB8 | 4.352334949 | 3.78E-06 |
| LINC00189 | 0.984614005 | 9.20E-05 |
| C20orf202 | 7.004806617 | 1.72E-05 |
| RSC1A1 | -0.630164973 | 4.00E-07 |
| TMEM242 | 1.384887812 | 1.46E-06 |
| TMEM167B | 0.38642643 | 0.000186407 |
| ARHGAP27P1-BPTFP1-KPNA2P3 | -0.237917614 | 0.038053367 |
| FAM72D | -0.310202377 | 0.000475886 |
| TNFRSF25 | -0.480344741 | 0.005776998 |
| SLC35E2A | 1.166722781 | 0.000137456 |
| KRT18P12 | 0.87788459 | 0.049184149 |
| AL451074.2 | 6.820671122 | 3.18E-05 |
| TSTD1 | 4.172141617 | 5.13E-06 |
| AL354702.1 | 0.789385471 | 1.54E-07 |
| CROCCP2 | -0.665372549 | 2.70E-05 |
| C1orf167 | 1.021359179 | 0.006187061 |
| TTC34 | 1.973698187 | 0.007424431 |
| MMP23A | 1.719629772 | 0.015084904 |
| AC078819.1 | -0.13116804 | 0.031792506 |
| GAPDHP72 | 0.236539727 | 0.028960151 |
| VN1R13P | 6.822795279 | 2.73E-05 |
| AL109918.1 | 0.472644395 | 0.00174939 |
| RPS2P55 | 0.55542689 | 9.38E-08 |
| AC009403.1 | 0.714833896 | 0.0208399 |
| RPL10P1 | 1.357196417 | 0.004704494 |

| | | |
|---|---|---|
| TPT1P4 | 0.46216916 | 4.58E-05 |
| AL135903.1 | -0.767470541 | 9.35E-05 |
| PPIAP31 | 0.212534367 | 0.023942428 |
| FNIP1 | 0.965692194 | 1.20E-07 |
| AL139100.2 | 0.33068889 | 9.92E-05 |
| AL031777.1 | 8.608629213 | 6.30E-08 |
| IQCB2P | 1.602913283 | 0.000138121 |
| AL355864.1 | 5.517331315 | 5.59E-05 |
| RPS10P3 | 0.403307776 | 0.008524616 |
| AL390719.1 | 0.790508464 | 0.001800986 |
| ZNF192P2 | 9.106358593 | 1.11E-08 |
| AL109955.1 | 3.128634222 | 6.91E-05 |
| AC016739.1 | 0.427749466 | 3.08E-06 |
| RPS27AP11 | 0.426130698 | 0.003457722 |
| TATDN2P2 | 0.557388488 | 0.004745801 |
| AC136632.1 | 0.370303089 | 3.21E-06 |
| NEPNP | 1.833135554 | 0.007250089 |
| MORF4L1P1 | -0.21056163 | 0.000304381 |
| TENM3 | -0.150985805 | 0.021174144 |
| LYPLA1P3 | -0.46372563 | 4.89E-05 |
| RAET1K | 2.047083672 | 1.56E-05 |
| SLC25A20P1 | 8.464924604 | 1.44E-07 |
| AL591135.1 | -0.730168563 | 6.16E-05 |
| AL590867.2 | 0.408552094 | 6.82E-07 |
| H2AZP3 | -0.47410524 | 0.000490001 |
| LINC00339 | 1.253862479 | 1.77E-07 |
| MIF-AS1 | 0.473376646 | 3.97E-06 |
| GAPDHP63 | 0.18738953 | 0.024894718 |
| AC006971.1 | -0.593642016 | 0.041967417 |
| AL117344.1 | 4.999739211 | 0.000119692 |
| BRD7P4 | -0.557474164 | 0.008294742 |
| CEBPZOS | -0.161030234 | 0.021347942 |
| NUFIP1P1 | -0.644249308 | 0.000265502 |
| AL033519.3 | 0.383819724 | 0.000225612 |
| NPM1P37 | 0.409847283 | 0.018459734 |
| AL592114.1 | 0.372206262 | 0.004676996 |
| HMGB1P20 | 5.010022036 | 0.001181211 |
| RNASEK | 0.372789218 | 6.10E-05 |
| AC138392.1 | -0.155810261 | 0.026435253 |
| RPS20P2 | 0.898725256 | 0.01160645 |
| PIP5K1P1 | -0.772194483 | 0.0256585 |
| ZNF602P | 2.921584907 | 0.000191096 |
| AC125494.1 | -0.604229877 | 1.69E-06 |
| CATSPERZ | 3.777178468 | 3.42E-05 |
| RPL23P8 | 0.190801368 | 0.013913442 |
| AL355802.1 | 0.497776552 | 0.00114266 |
| NBPF1 | 0.155953076 | 0.016035995 |
| AL603766.1 | 0.191660395 | 0.010993531 |

| | | |
|---|---|---|
| FTH1P8 | 0.248596757 | 0.001744657 |
| AP000580.1 | 1.240325314 | 3.02E-05 |
| UMAD1 | -0.236128815 | 0.024519191 |
| FAM228B | 2.76467789 | 9.24E-08 |
| BMPR1AP1 | -0.611535914 | 0.000928454 |
| ZNF433-AS1 | 0.665674355 | 0.002192923 |
| AL133268.1 | -1.93924681 | 1.80E-09 |
| AL133260.1 | 0.362050799 | 0.000158896 |
| AL137784.1 | 0.850870513 | 0.011462409 |
| RPSAP45 | 0.245580191 | 0.025819222 |
| GAPDHP42 | 7.306404312 | 6.21E-06 |
| ZSCAN12P1 | 2.797203867 | 1.70E-07 |
| AL161787.1 | 0.207615872 | 0.010466541 |
| RPL12P8 | 0.503177837 | 0.016553531 |
| BTF3P7 | 1.266712354 | 0.024735429 |
| AL391422.1 | 7.76823162 | 1.30E-06 |
| AL353151.1 | 1.537588861 | 0.013332426 |
| HNRNPA1P12 | 0.549202469 | 1.11E-05 |
| LINC02076 | 1.869643832 | 5.64E-05 |
| ZGLP1 | 1.882765757 | 4.85E-07 |
| VAMP2 | 0.552452124 | 4.28E-05 |
| H2BC19P | 2.178612265 | 4.76E-08 |
| AL356234.1 | -0.840418079 | 0.016665893 |
| AL139095.2 | 0.493656135 | 2.64E-05 |
| HMGB1P13 | 7.433296986 | 4.86E-06 |
| PKMP3 | 7.06186844 | 1.10E-06 |
| RPL35P2 | 0.360485369 | 0.000163814 |
| Z86062.1 | 0.454350218 | 0.038280751 |
| RPL21P28 | 0.284311207 | 0.000281342 |
| MTMR9LP | 1.654801258 | 0.000107595 |
| RPL21P119 | 0.317008429 | 7.79E-05 |
| RPL21P16 | 0.322445083 | 3.13E-06 |
| RPS18P9 | 0.393300529 | 0.047882425 |
| SNORA81 | 3.283135625 | 4.42E-05 |
| SNORD100 | -1.475880947 | 0.002123151 |
| PPP3CB-AS1 | 1.76305638 | 1.45E-08 |
| C6orf226 | 0.851670029 | 0.035480278 |
| PPP3R1 | -0.962876167 | 2.53E-09 |
| FANCG | -0.290557326 | 0.000692209 |
| AP4M1 | 0.253351754 | 0.011533459 |
| PLXNA4 | 2.186861453 | 0.002384994 |
| MAGEA3 | -0.248401103 | 0.000113227 |
| ZNF816-ZNF321P | 0.810176402 | 0.009588979 |
| ZBED8 | 1.043143574 | 4.62E-08 |
| HMSD | -0.636862038 | 0.045743318 |
| FAM200A | 0.510604233 | 3.92E-05 |
| PPP2R2A | -1.526649167 | 8.16E-10 |
| C19orf73 | 2.803463019 | 0.000166573 |

| | | |
|---|---|---|
| ZNF880 | -1.195285082 | 1.25E-05 |
| TRIM16 | -0.937830264 | 7.27E-07 |
| DENND10P1 | 0.258107173 | 0.005923937 |
| HEPN1 | 7.165575523 | 1.28E-05 |
| LINC01465 | -1.947777781 | 0.005764725 |
| APOL6 | -1.117243755 | 1.29E-07 |
| FADS3 | 0.330394409 | 0.005237924 |
| CCNL2 | -1.047183954 | 9.33E-10 |
| UBA52 | 0.322139185 | 1.26E-06 |
| EXOC3-AS1 | 1.867863865 | 2.03E-06 |
| TIAF1 | -0.497303169 | 0.000188613 |
| BTBD19 | 1.026417996 | 7.11E-05 |
| RAB6C | -0.481919655 | 0.00538728 |
| HDAC4-AS1 | 4.219107393 | 0.002147409 |
| KIAA2012-AS1 | 4.680827859 | 0.000540354 |
| POTEJ | 0.902413582 | 0.013936993 |
| CYTOR | -0.292529477 | 0.005618268 |
| AC079305.2 | -0.908650266 | 0.029228618 |
| DCDC2B | 2.342538192 | 0.000218937 |
| RN7SKP16 | -3.974924419 | 3.71E-05 |
| SNORD12B | -2.690065506 | 0.00022676 |
| AL353769.1 | 4.359159322 | 0.000887458 |
| RNU6-77P | 5.124208656 | 0.000299524 |
| SNORA79B | 1.32084561 | 0.022791744 |
| H2BP1 | 2.651106606 | 7.20E-07 |
| AL590666.1 | 1.345897715 | 0.025539905 |
| FTH1P10 | 0.198344431 | 0.002918241 |
| AL445685.1 | -0.74499699 | 0.006193932 |
| RPS10P7 | 1.77454325 | 7.73E-05 |
| LINC00397 | 1.326256411 | 0.038122287 |
| AL590632.1 | 2.950808851 | 0.001474169 |
| TCAF1P1 | 0.870464858 | 5.35E-08 |
| AL441992.2 | 0.62883668 | 0.045731793 |
| NUTM2A-AS1 | -1.021691774 | 1.69E-07 |
| EXOSC6 | -0.16221811 | 0.037041686 |
| AC135983.3 | -1.442277428 | 4.57E-05 |
| CDRT15 | -1.473457242 | 0.025753661 |
| CSNK1A1P1 | -0.477804615 | 0.042084813 |
| EEF1A1P8 | 3.208538713 | 5.43E-07 |
| RPS15AP5 | 0.680551127 | 0.033747358 |
| ZNF844 | 1.492791307 | 3.83E-10 |
| TMSB4XP4 | 1.191405748 | 0.004119055 |
| AC073136.1 | -0.380602602 | 0.000338833 |
| TINCR | 0.904163325 | 0.001397836 |
| FNDC3CP | 7.199837387 | 8.61E-06 |
| AL513523.3 | 0.798079843 | 0.028666106 |
| AC241585.1 | -0.514502358 | 0.005143815 |
| LINC00370 | 5.488595819 | 4.51E-08 |

| | | |
|---|---|---|
| UHRF2P1 | -1.191655246 | 0.019785283 |
| ADH5P3 | 1.185017374 | 0.047516585 |
| AC027612.1 | -1.827920441 | 0.006489323 |
| LINC01422 | 8.55123424 | 7.80E-08 |
| NSUN5P1 | 0.170604525 | 0.014755882 |
| BX842568.2 | -0.652206069 | 0.000134102 |
| CCDC18-AS1 | 2.410303469 | 1.02E-07 |
| MIR503HG | 1.817898987 | 2.44E-05 |
| TSSC2 | 2.10310508 | 0.013797782 |
| PRR3 | -0.998006727 | 5.34E-05 |
| LINC00205 | 0.99752766 | 1.17E-05 |
| CD99P1 | 1.445373196 | 0.010788005 |
| AL513217.1 | 3.441581945 | 0.018737502 |
| ENTPD3-AS1 | 1.323814101 | 0.000217434 |
| CERS1 | 0.496424751 | 0.001893455 |
| RPS20P14 | 0.410140493 | 4.85E-05 |
| CFL1P1 | 2.19642849 | 0.000669114 |
| AL603910.1 | 3.622099358 | 8.29E-05 |
| FAM95B1 | -1.822378862 | 0.010051909 |
| NPM1P19 | 0.724605277 | 0.047890371 |
| NBEAP3 | 0.973843136 | 0.001340778 |
| OSER1-DT | 3.842620779 | 6.82E-12 |
| AC016738.1 | -0.636895327 | 0.001172988 |
| AFG3L1P | -0.331687422 | 0.003123889 |
| AC068491.1 | -0.981096894 | 0.000187607 |
| HNRNPRP1 | -0.598954601 | 0.001249371 |
| EDRF1-DT | 4.376243828 | 2.93E-07 |
| EPB41L4A-AS1 | 0.411717503 | 9.75E-05 |
| CCNT2-AS1 | -1.360484118 | 0.007782359 |
| AC005076.2 | 3.092910693 | 3.89E-08 |
| CPTP | 0.237919497 | 0.019271673 |
| AC092106.1 | -0.822029282 | 0.018553128 |
| AL049795.1 | 1.130163946 | 0.001820619 |
| AL139811.1 | 1.377535312 | 0.009618166 |
| SNHG14 | 0.452380999 | 9.41E-07 |
| UBE2FP1 | 1.059521669 | 0.00083414 |
| AC245452.1 | -0.623723494 | 5.01E-05 |
| RPS24P8 | 0.360086643 | 0.002640075 |
| AC021148.1 | 2.48011123 | 2.21E-08 |
| AL591846.2 | 0.639034904 | 0.024045465 |
| INHBA-AS1 | 3.759390338 | 8.41E-07 |
| PTPN2P2 | -0.983793236 | 0.047831219 |
| ATG12P2 | 4.836676438 | 0.000513949 |
| UBE2SP2 | -0.692050854 | 2.21E-07 |
| AC009506.2 | 0.841114366 | 0.037863298 |
| DNAJC27-AS1 | 3.557381762 | 1.88E-06 |
| AL137802.1 | 4.869319321 | 0.000285145 |
| C5orf66 | -0.598900917 | 3.26E-06 |

| | | |
|---|---|---|
| BX119904.1 | -1.017035946 | 0.048228494 |
| MRRFP1 | 0.48760764 | 0.037953135 |
| RPSAP18 | 0.650445587 | 0.007284623 |
| ARHGEF2-AS1 | 9.074291685 | 2.01E-08 |
| SLC25A5-AS1 | 0.436425787 | 0.001468539 |
| AF196972.1 | 1.561864308 | 0.033347513 |
| TMCO1-AS1 | 1.968426359 | 1.39E-05 |
| AC017104.1 | 1.458003798 | 0.000946579 |
| HSP90AA2P | 0.140442356 | 0.000952388 |
| AC007683.1 | 0.484439928 | 0.001023482 |
| STK24-AS1 | 2.36514681 | 0.000602854 |
| KRT18P27 | 0.929252816 | 0.044012267 |
| ADM5 | 0.922916323 | 2.73E-05 |
| PRKAR2A-AS1 | 2.135659738 | 9.35E-07 |
| PIGUP1 | -0.87263461 | 0.019530154 |
| ATXN1L | -0.203599318 | 0.001351024 |
| AC104162.1 | 2.973406831 | 0.001172753 |
| AC245100.1 | 7.526402291 | 2.96E-06 |
| AC138150.1 | -0.453523374 | 0.011963526 |
| KRT8P45 | 2.013481226 | 8.84E-05 |
| SMIM13 | -1.026656624 | 4.35E-09 |
| AC092810.1 | -0.705843954 | 0.01951243 |
| EIF4BP3 | 0.166844979 | 0.009821449 |
| HNRNPA1P48 | 0.601997684 | 7.25E-09 |
| SVIL-AS1 | -1.350798185 | 1.31E-07 |
| TNK2-AS1 | -1.300999869 | 0.034852227 |
| RTCA-AS1 | 3.101634671 | 8.17E-06 |
| HCG18 | 0.656926963 | 0.009263754 |
| AL121723.1 | 0.961643544 | 0.007933649 |
| RPS27AP16 | 0.386610525 | 2.56E-06 |
| AL391095.2 | 0.943552391 | 0.000372109 |
| PDSS1P2 | 2.66941194 | 4.45E-05 |
| AC026954.1 | 5.263530902 | 4.85E-05 |
| SH3BP5-AS1 | 0.736843448 | 8.25E-05 |
| IPO8P1 | -0.297693908 | 0.002816342 |
| ZNF812P | -0.439547278 | 0.000670414 |
| LAMTOR5-AS1 | 1.910795849 | 2.98E-06 |
| AC020688.1 | -1.071304941 | 3.22E-05 |
| CEP57L1P1 | -1.057499684 | 0.004667315 |
| PCOLCE-AS1 | 0.931342735 | 0.000190331 |
| AC099850.2 | 1.628317055 | 4.49E-05 |
| MUC20P1 | 7.236508162 | 7.79E-06 |
| CSNK2B | 0.501787964 | 0.044655968 |
| EIF4A2P4 | 1.330431595 | 1.46E-07 |
| CETN4P | 4.389268903 | 4.19E-06 |
| TMEM72-AS1 | 7.203936 | 1.41E-05 |
| AC096677.2 | 1.38097358 | 0.000103335 |
| TMEM183B | -0.158310665 | 0.020398707 |

| | | |
|---|---|---|
| BTF3P9 | 0.294677479 | 0.005901774 |
| RPL12P17 | 1.020468988 | 0.016462462 |
| LINC00240 | 2.54329239 | 9.92E-06 |
| OPA1-AS1 | 4.717830429 | 0.000331803 |
| RPL29P11 | 0.347709599 | 7.77E-05 |
| YBX1P1 | 0.200177647 | 0.001103297 |
| AC011447.1 | 0.445516175 | 0.000539592 |
| MRPL20-AS1 | 0.962092076 | 4.09E-07 |
| AC138028.2 | -0.50632889 | 0.006872945 |
| AC007899.1 | -0.670007478 | 0.031416986 |
| RPS4XP16 | 2.066995838 | 1.25E-08 |
| AP001347.1 | 2.557002356 | 0.000183981 |
| LINC00863 | 2.086208169 | 6.87E-09 |
| AL391684.1 | 2.326089329 | 7.47E-08 |
| LINC00184 | 4.236082246 | 0.003368726 |
| AL137014.1 | -1.295724061 | 0.012083021 |
| AL353150.1 | 3.897563401 | 6.21E-06 |
| AL390066.2 | -1.160897518 | 0.017894739 |
| UBE2D3P1 | -0.322716799 | 0.002769215 |
| LINC01058 | 7.582561345 | 2.60E-06 |
| HMGB3P22 | 6.774837438 | 3.09E-05 |
| AC012485.1 | -1.32660799 | 0.000402369 |
| RPL23AP2 | 0.425568182 | 2.15E-06 |
| PFN1P4 | 1.337458548 | 0.003962316 |
| RPL3P7 | 0.5759989 | 1.21E-05 |
| CR388220.1 | 0.37911245 | 0.008039505 |
| RANP4 | 5.337768563 | 0.00016815 |
| DYNC1I2P1 | -0.212235125 | 0.021651451 |
| SLC9A3-AS1 | -0.849757091 | 0.017551417 |
| NPM1P39 | 0.492678514 | 4.52E-07 |
| DUTP6 | 3.452137745 | 1.98E-06 |
| AL590617.2 | -0.979095751 | 3.20E-05 |
| PLEKHM1 | -1.559857339 | 1.24E-06 |
| RPS12P26 | 0.414550692 | 0.04895828 |
| AC246787.1 | 0.624866856 | 0.000584328 |
| AC078883.1 | -0.649685028 | 0.044646588 |
| DUXAP9 | -0.404529916 | 5.30E-05 |
| RPS27AP12 | 0.598155918 | 0.00155817 |
| ST13P6 | 0.2684815 | 0.003165486 |
| TAF1A-AS1 | 3.475277138 | 1.46E-07 |
| AC074281.1 | -1.184156696 | 0.048824979 |
| AL121987.1 | 7.16221043 | 9.60E-06 |
| PPIAP9 | 0.614760709 | 0.046815536 |
| LHFPL3-AS2 | 7.642686996 | 2.98E-06 |
| RPL23AP18 | 0.521331915 | 0.002953776 |
| SNX18P3 | 0.738193534 | 0.002668306 |
| ARL6IP1P2 | -0.523384933 | 0.018595049 |
| PPP1R26-AS1 | 0.710679909 | 0.010346746 |

| | | |
|---|---|---|
| AC009506.3 | -0.518980562 | 0.002774417 |
| NRSN2-AS1 | 2.533935713 | 5.48E-08 |
| AC016700.1 | -0.957124676 | 0.028902542 |
| CR786580.1 | 2.262256121 | 2.14E-06 |
| AC104134.1 | 0.774782604 | 0.00019956 |
| RBMS1P1 | -1.366166031 | 4.55E-07 |
| BOLA3-AS1 | -1.997292138 | 6.85E-09 |
| RPS15AP10 | 1.61150811 | 0.000193873 |
| RAB6C-AS1 | 0.833052117 | 0.00151025 |
| FDPSP1 | 1.168524789 | 0.001492589 |
| RFPL1S | 9.37478777 | 5.23E-09 |
| JPX | 0.921716738 | 1.52E-07 |
| BX119917.1 | 0.60191716 | 0.00145709 |
| NUTM2B-AS1 | -0.895942012 | 3.05E-07 |
| AL354707.1 | 2.378492248 | 9.01E-05 |
| LINC00475 | 2.987084854 | 6.17E-06 |
| LINC01703 | 1.451011026 | 0.005362134 |
| AC245452.2 | 0.889142915 | 0.046737475 |
| LINC02476 | 1.069095245 | 0.00538277 |
| UBE2D3P2 | -0.544168997 | 0.004121459 |
| AC093155.1 | -0.65615729 | 0.003909451 |
| RPL35P5 | 0.250183653 | 0.010096862 |
| BX248409.1 | 0.682322171 | 0.023724665 |
| LINC02158 | 1.812229977 | 0.014316041 |
| ZNF469 | -1.512338052 | 3.94E-08 |
| MTND2P28 | 0.187658628 | 0.000131385 |
| IPO7P2 | -0.23699004 | 0.003078946 |
| LAGE3P1 | 2.280484857 | 0.015631303 |
| HNRNPA1P35 | 0.631201247 | 1.24E-06 |
| SLC26A6 | 0.328076678 | 0.000324251 |
| ASH2LP3 | 7.12530457 | 1.13E-05 |
| FGD5-AS1 | -0.136048808 | 0.007969521 |
| NPM1P18 | 0.412292842 | 0.005191011 |
| LINC00431 | 0.999996472 | 0.015000721 |
| LINC01389 | 1.613860987 | 0.037361437 |
| TRAM2-AS1 | 1.031761916 | 9.05E-05 |
| AL080250.1 | 2.475976213 | 8.47E-06 |
| FAM229A | 1.031522707 | 2.38E-06 |
| ERCC6 | -0.230180428 | 0.014115493 |
| MKRN7P | 7.694574593 | 1.98E-06 |
| AL355490.2 | 1.228863025 | 0.046261631 |
| RUSC1-AS1 | 0.196080696 | 0.014719569 |
| AC245100.2 | -0.533172536 | 0.01203324 |
| C3orf86 | 7.719547343 | 1.63E-06 |
| LINC00115 | 0.904211231 | 0.032545537 |
| BMS1P11 | 0.648120283 | 0.036092104 |
| HLA-DQA1 | 0.515598798 | 0.00987683 |
| FRG2B | 3.430968748 | 0.000105019 |

| | | |
|---|---|---|
| RIMKLBP2 | 2.820708399 | 4.31E-08 |
| NOL7 | -0.237562618 | 0.003608945 |
| ELFN1 | 1.983672133 | 5.65E-06 |
| MTND1P23 | 0.19337377 | 0.018843228 |
| PIGBOS1 | -1.025966256 | 7.88E-05 |
| LINC01534 | 1.704268275 | 0.000170888 |
| CKS1BP2 | 0.399577188 | 7.35E-05 |
| PBX2 | 0.439910582 | 0.003938432 |
| BX005266.2 | 0.644958203 | 0.021928805 |
| CCT8P1 | 0.211394102 | 0.002593583 |
| AC092802.2 | 4.106908019 | 3.76E-07 |
| TLK2P1 | -1.008288264 | 4.05E-08 |
| MEMO1P1 | -0.550777978 | 0.000131216 |
| PAICSP1 | 1.145473214 | 0.020440332 |
| PHF2P2 | -1.479640119 | 1.26E-05 |
| LINC00623 | -0.516490671 | 0.007970185 |
| USP12PX | -0.89789237 | 0.001325566 |
| AC113935.1 | 0.334789906 | 5.44E-06 |
| UQCRFS1P1 | -0.306035161 | 0.000701256 |
| MRPS18B | 0.492122969 | 0.006921341 |
| AC009487.2 | -0.702213722 | 7.09E-09 |
| LINC01907 | 7.585715246 | 2.67E-06 |
| BAIAP2-DT | 0.80274522 | 0.000455135 |
| Z98752.1 | 4.131851248 | 0.002579904 |
| EIF4A1P5 | -0.466320179 | 0.026090604 |
| AP4B1-AS1 | -0.481391307 | 0.016113225 |
| TEX22 | 1.280188087 | 0.004350313 |
| SGMS1-AS1 | 2.215696108 | 9.73E-09 |
| RPL26P19 | 0.410357453 | 6.01E-07 |
| PRELID3BP1 | 0.719374137 | 0.047857779 |
| RPLP0P1 | 9.17388688 | 9.27E-09 |
| NPIPB14P | 1.100581234 | 0.002197409 |
| RPL37AP1 | 0.435510343 | 6.53E-05 |
| HLA-DMB | -0.394202497 | 0.031374418 |
| ARPC3P1 | 0.251197164 | 0.036258034 |
| TMEM191A | 0.972241739 | 0.034911856 |
| ZNF192P1 | 2.711578927 | 2.29E-07 |
| RPS3AP38 | 4.566810451 | 7.69E-07 |
| NUP50-DT | 2.633866213 | 7.35E-06 |
| AC005682.1 | 6.618143764 | 4.85E-05 |
| NMD3P1 | -0.935167667 | 0.000948112 |
| PSPC1-AS2 | -5.212568068 | 0.000489655 |
| ACTG1P24 | 2.995622511 | 0.023721134 |
| RPL10AP6 | 0.538325771 | 2.78E-07 |
| TERF1P5 | -0.886568592 | 0.000878583 |
| AC016831.1 | 0.449884691 | 0.03657717 |
| IFNA20P | 3.872155173 | 0.00039493 |
| AL031727.1 | 0.367408957 | 0.000157197 |

| | | |
|---|---|---|
| SLC16A1-AS1 | -0.805818033 | 7.72E-05 |
| AC108161.1 | 0.390659943 | 0.04075716 |
| GAPDHP32 | 2.871573481 | 3.29E-06 |
| RPL22P3 | 0.329024464 | 0.03602308 |
| AL390198.1 | 3.215681437 | 0.023612983 |
| UPF3AP1 | 0.436868797 | 0.012062961 |
| TMEM185B | 0.552045293 | 3.88E-06 |
| RPS7P10 | 0.332869213 | 8.87E-05 |
| AL445487.1 | -0.673365568 | 0.020847594 |
| AC012512.1 | 0.822421649 | 8.10E-05 |
| RPL7P22 | -0.56170606 | 0.000393928 |
| SCDP1 | 1.663946783 | 1.87E-08 |
| CYP4F26P | -1.887916438 | 0.00273221 |
| FTH1P20 | 0.198637602 | 0.003189053 |
| FTLP3 | 0.271742146 | 0.000282902 |
| AL365356.1 | -1.647842206 | 0.001880698 |
| KIF4B | -0.824611404 | 1.07E-05 |
| MTND4P10 | 7.12530457 | 1.13E-05 |
| IGBP1P1 | 0.589022411 | 0.024780166 |
| LINC01535 | 1.379018871 | 0.000617452 |
| ENTPD1-AS1 | -0.65642298 | 0.000116106 |
| AL034380.1 | 2.156253505 | 0.003957202 |
| AL161452.1 | -0.59659852 | 0.031266485 |
| AL513175.1 | -0.595870755 | 1.93E-06 |
| HSBP1L1 | 0.999652578 | 0.00772595 |
| CUTALP | 0.506979307 | 4.13E-05 |
| AL606760.1 | 1.072446474 | 0.000204889 |
| TBCAP1 | 0.281183852 | 0.000446918 |
| TLK1P1 | -0.966233241 | 0.041781363 |
| CACTIN-AS1 | -0.333167291 | 0.035878664 |
| MEIS1-AS3 | 3.801299656 | 0.006432935 |
| AC006001.2 | 1.456965054 | 0.015267397 |
| LINC02774 | 3.752356718 | 2.37E-05 |
| AC092164.1 | 8.632056389 | 1.30E-07 |
| AC010894.2 | 3.396684984 | 0.000180345 |
| RPSAP17 | 0.323466555 | 0.042708691 |
| ATE1-AS1 | 4.352334949 | 3.78E-06 |
| AC005154.1 | 2.536842686 | 0.00474523 |
| AL354733.1 | 1.77098875 | 0.007165582 |
| E2F6P3 | -0.818439735 | 0.000138221 |
| ERVMER34-1 | 2.297043387 | 1.16E-07 |
| AL451069.1 | 7.4367906 | 4.07E-06 |
| BSN-DT | 5.112838671 | 2.03E-08 |
| PDZPH1P | 6.796391694 | 7.55E-06 |
| CEP164P1 | -0.65601857 | 5.92E-05 |
| IL9RP3 | 1.587956505 | 0.010993531 |
| RPS4XP2 | 0.531825251 | 0.000341953 |
| DANCR | 0.275609472 | 0.000321545 |

| | | |
|---|---|---|
| RPL12P19 | 1.245309907 | 0.00679695 |
| COX6A1P2 | 0.310636368 | 6.40E-05 |
| LINC01203 | 0.745311705 | 0.002124224 |
| AL009174.1 | 0.327483588 | 0.003371506 |
| AC007285.1 | 1.611337134 | 0.002328511 |
| SLC8A1-AS1 | -1.376130035 | 0.007387241 |
| LINC00511 | -0.331246544 | 0.001961655 |
| ITGB2-AS1 | -1.35860167 | 0.004265703 |
| AC022483.1 | -0.881663959 | 0.000182051 |
| C14orf132 | 0.999517691 | 0.049117699 |
| WDR46 | -0.155353234 | 0.025816188 |
| RPL41P1 | 0.452896559 | 3.42E-05 |
| AC107983.1 | 0.670004754 | 0.002033285 |
| AC005912.1 | 0.29103919 | 0.000562036 |
| LINC02798 | 2.19953561 | 0.000748489 |
| RBMX2P5 | -0.668380623 | 0.02684005 |
| RPS28P7 | 0.481439576 | 1.12E-06 |
| LMCD1-AS1 | 2.249175781 | 0.000477766 |
| ZNF717 | 0.567617387 | 0.000172483 |
| PGBP | 5.40762703 | 0.000129594 |
| SUB1P1 | 0.371310139 | 0.006331088 |
| AC079145.1 | 5.342656068 | 9.21E-08 |
| PFN1P6 | 1.541267834 | 2.99E-06 |
| AL157935.2 | 1.171343163 | 0.00104917 |
| AL133346.1 | 0.802948667 | 0.042772196 |
| AL672291.1 | 7.199837387 | 8.61E-06 |
| AC012063.1 | 1.076780394 | 0.013379313 |
| MIS18A-AS1 | 1.748566073 | 8.48E-05 |
| SMIM2-AS1 | 1.821812386 | 0.005249554 |
| KLLN | 3.30585334 | 9.60E-08 |
| ELL2P1 | 0.876771504 | 1.68E-05 |
| AL035541.1 | 0.541217682 | 0.011625255 |
| AC067942.1 | 0.581784672 | 0.006282147 |
| AC140076.1 | 1.031493201 | 0.025477926 |
| AL139396.1 | -1.696198487 | 0.017723521 |
| AC005042.1 | 1.571735778 | 0.019842774 |
| HAUS6P1 | -0.586624818 | 1.32E-06 |
| PARG | -1.161105747 | 2.57E-08 |
| HNRNPKP2 | 0.245309144 | 0.00053657 |
| RBM26-AS1 | 4.80198526 | 1.36E-12 |
| HLA-DRB4 | 0.25347997 | 0.010299393 |
| TP73-AS1 | -0.406168467 | 0.019315483 |
| FTH1P16 | 0.32731749 | 0.000599483 |
| EIF4A2P2 | 1.882952459 | 0.006714846 |
| AL133410.1 | 7.497145996 | 3.27E-06 |
| AC007386.1 | 4.220143675 | 0.001626747 |
| KIF9-AS1 | 2.031617573 | 3.46E-06 |
| ZMYM4-AS1 | -0.936350717 | 0.031435381 |

| | | |
|---|---|---|
| AKR1B15 | 4.965279701 | 0.000139789 |
| STK4-AS1 | 3.917346007 | 5.19E-08 |
| AC099066.2 | -0.97900658 | 0.009060647 |
| SCAMP4 | 0.247074555 | 0.003048582 |
| RPL7P6 | 0.226381182 | 0.049884189 |
| AC096540.1 | 1.812229977 | 0.014316041 |
| AC016394.2 | 2.67137719 | 8.83E-08 |
| ADGRF5P1 | 1.517973233 | 3.51E-05 |
| AP001324.1 | 0.393511684 | 3.99E-06 |
| CERS6-AS1 | -0.690012 | 5.46E-06 |
| RPL3P2 | 0.600233406 | 1.47E-05 |
| AL390728.4 | -0.141139324 | 0.028869365 |
| LINC00355 | 3.705942685 | 6.42E-13 |
| RPL23AP74 | 0.486528481 | 3.67E-06 |
| AC091607.1 | -0.495305156 | 0.029229568 |
| AC006042.1 | 2.858443541 | 0.000313321 |
| C4A | 4.50724666 | 0.002089429 |
| RCC2-AS1 | -0.604037869 | 0.002255984 |
| SUCLA2P1 | -0.905899874 | 0.044138554 |
| AL031282.1 | 0.891503409 | 4.38E-05 |
| AC012358.3 | 1.416165155 | 0.005503647 |
| INKA2-AS1 | 2.618343379 | 0.000448859 |
| SLC9A7P1 | -0.781529279 | 0.007685881 |
| AC138969.3 | -0.258915472 | 0.021305846 |
| DPY19L2P3 | -0.399592051 | 0.034680681 |
| PSPC1P1 | 1.442537597 | 4.41E-05 |
| AL590652.1 | 5.292729189 | 5.44E-05 |
| AC016596.1 | 5.42306759 | 1.07E-09 |
| AC092634.3 | 2.034648203 | 0.002410792 |
| FRMD8P1 | -0.905324822 | 0.006010083 |
| THAP12P3 | 1.401901741 | 0.004644759 |
| AC011933.1 | 0.585363497 | 0.00132989 |
| AC073343.2 | 2.532488486 | 0.001539933 |
| HNRNPA1P46 | 3.486128767 | 9.18E-07 |
| AL158835.1 | 5.036735973 | 3.86E-07 |
| AL390729.1 | 4.465487986 | 0.000832529 |
| AC114763.1 | 1.578471289 | 0.009393464 |
| AL158823.1 | 1.109813071 | 0.004090714 |
| RPL7P47 | 0.360700628 | 0.002006991 |
| HLA-DRB1 | 0.336582954 | 0.000634637 |
| MELTF-AS1 | -0.853858702 | 0.005628341 |
| ST13P19 | 0.50481575 | 0.007166866 |
| AL096678.1 | 1.561932221 | 0.016423386 |
| AC105339.1 | 5.112336634 | 0.000174929 |
| AC078927.1 | -0.258564631 | 0.045648903 |
| CASP16P | 4.0027346 | 4.72E-05 |
| TLE1P1 | -0.753189289 | 0.037116633 |
| AL031963.1 | 1.05313424 | 0.008409549 |

| | | |
|---|---|---|
| AL020996.1 | 0.753483734 | 0.027196846 |
| GEMIN8P4 | 1.772896774 | 2.14E-06 |
| AL512353.1 | 3.234627521 | 2.85E-08 |
| PTPN2P1 | -0.580539726 | 0.005436867 |
| AC131235.2 | 0.481817502 | 1.71E-06 |
| ATF4P3 | 0.394212549 | 0.021269658 |
| HCG11 | -1.124824627 | 4.17E-07 |
| NACA4P | 0.329508514 | 0.001385821 |
| GAPDHP1 | 0.245945964 | 0.000133137 |
| AC093495.1 | 0.74264011 | 0.000710228 |
| COL6A4P2 | 8.368349056 | 1.52E-07 |
| MT-ATP8 | 0.25871198 | 0.005163054 |
| C4B | 2.932793973 | 0.030756372 |
| AL021707.2 | 1.273534158 | 4.44E-05 |
| KATNBL1P6 | -0.319879002 | 0.021734273 |
| LYPLA2P1 | -0.810628107 | 0.002377801 |
| PCAT6 | 3.736228033 | 0.000136779 |
| BMS1P17 | -1.678580868 | 4.29E-06 |
| AL512770.1 | -0.47649018 | 0.045707829 |
| AL669831.1 | 1.225723106 | 0.029765307 |
| AC015971.1 | 0.55198708 | 0.003284766 |
| LINC01004 | 2.465336036 | 2.13E-07 |
| CCT6P1 | 0.442258404 | 0.005688172 |
| AL139260.1 | 2.344548251 | 0.000206067 |
| SDAD1P1 | 1.46295572 | 7.83E-07 |
| AC098484.1 | -1.17565091 | 0.016708785 |
| RPS19P7 | 0.490820233 | 0.002020711 |
| AP006222.1 | 1.525637066 | 0.000392206 |
| OST4 | 0.2226787 | 0.003449308 |
| C2orf92 | 0.735993016 | 0.008284348 |
| MICE | 2.294712604 | 0.001753235 |
| RAB11FIP1P1 | -1.470846825 | 9.80E-05 |
| EEF1A1P11 | 0.467195563 | 7.44E-07 |
| CCDC183-AS1 | 0.499377889 | 0.018950229 |
| NUTM2E | 3.243197293 | 8.30E-09 |
| SPCS2P4 | -0.246459009 | 0.005794406 |
| AL139011.1 | -1.478160986 | 0.007026884 |
| HK2P1 | -1.041776656 | 3.85E-06 |
| ZNF883 | -1.153542629 | 4.85E-07 |
| HDAC2-AS2 | 2.2928973 | 0.001803181 |
| HOTAIR | 1.205963678 | 0.002274483 |
| HNRNPCP7 | 1.623246958 | 0.000637758 |
| PSMD10P1 | 1.931975056 | 0.005159652 |
| TNKS2-AS1 | 3.016835255 | 6.59E-06 |
| LINC02575 | -0.926789303 | 0.012322633 |
| AL450306.1 | -1.249079915 | 5.59E-06 |
| FAF2P1 | 3.467618375 | 0.006816744 |
| LINC01128 | -1.348256951 | 1.52E-09 |

| | | |
|---|---|---|
| FAM207BP | -0.741614624 | 0.00264651 |
| AC064807.1 | 3.321367916 | 8.74E-05 |
| AC072022.1 | 1.927564143 | 0.00227756 |
| MIR4500HG | -2.695685444 | 0.003834737 |
| AL160408.2 | -1.024329141 | 1.25E-06 |
| CT45A5 | -0.593760273 | 2.98E-05 |
| AL133410.2 | -0.565042098 | 0.038612485 |
| AC104653.1 | 5.254412009 | 0.00010512 |
| SEPTIN7-AS1 | 3.062712871 | 6.38E-07 |
| UBAC2-AS1 | 4.23272192 | 5.10E-06 |
| SKIV2L | 0.35829312 | 0.00083818 |
| RASA4CP | 1.962114189 | 0.002468336 |
| KPNA2P2 | -0.983323631 | 0.008938424 |
| RPS13P2 | 0.404250652 | 2.27E-06 |
| UBE3AP2 | -1.2590574 | 9.56E-05 |
| HOMER2P1 | 1.919875291 | 0.0064705 |
| UBTFL6 | 8.444164071 | 2.02E-07 |
| AC097658.2 | 0.461155947 | 9.16E-05 |
| AKR7A2P1 | 2.406723574 | 1.39E-05 |
| AL450992.2 | 4.641642706 | 0.000650225 |
| AL365366.1 | 0.667064022 | 0.006664355 |
| AC091729.3 | 3.097469902 | 3.14E-11 |
| AF127577.1 | -0.507637056 | 0.02939359 |
| AL449283.1 | 4.790849248 | 0.000344666 |
| PRRT1 | 2.059709027 | 0.000236956 |
| LINC01165 | 8.648484178 | 5.48E-08 |
| HSPA8P8 | -0.286030782 | 0.000359029 |
| CALM2P2 | -0.651322318 | 7.95E-07 |
| LINC02587 | -1.570992653 | 0.046277597 |
| AC026403.1 | 0.58277587 | 9.86E-08 |
| VIM-AS1 | 0.538806385 | 0.004730096 |
| KANSL1L-AS1 | 3.284727364 | 2.39E-06 |
| RPS7P4 | 0.336992091 | 0.008452197 |
| ANKRD10-IT1 | -0.270617023 | 0.02313824 |
| AC114488.1 | 1.981454808 | 0.005262262 |
| ADAM1A | -1.232899337 | 0.000199534 |
| AL358473.1 | 2.243917653 | 0.000117409 |
| PTGES3P3 | -0.22695287 | 0.0261187 |
| PNPT1P1 | -0.648514652 | 0.001377065 |
| HNRNPA1P8 | 0.475484242 | 4.17E-05 |
| ST13P13 | 0.410428081 | 0.043820339 |
| TUBB8P1 | 1.118549919 | 0.00678634 |
| KRT8P12 | 1.316305619 | 2.51E-08 |
| ACAP2-IT1 | 1.353224265 | 0.00088699 |
| MTCO2P12 | 0.641645673 | 4.62E-05 |
| HYI-AS1 | 2.032003037 | 0.000821607 |
| HMGN2P19 | 1.322145396 | 0.008400836 |
| SZT2-AS1 | 0.721366716 | 0.045522742 |

| | | |
|---|---|---|
| LINC00452 | 2.425389546 | 1.93E-07 |
| USP24P1 | -1.598778284 | 0.000180883 |
| LINC01715 | 4.447941996 | 6.77E-09 |
| RALGAPA1P1 | -0.340738307 | 0.000126208 |
| AL512625.2 | 1.195955374 | 0.007645985 |
| RGS17P1 | 2.239413625 | 0.003122871 |
| PATL2 | 5.570238616 | 1.03E-08 |
| AC092155.2 | 0.389912171 | 0.048860272 |
| AC053503.1 | 3.315908945 | 0.013515947 |
| ZFYVE9P1 | -1.499996327 | 0.002288175 |
| LINC01204 | 2.528062935 | 1.48E-06 |
| TNPO1P3 | -0.439537984 | 0.000711236 |
| ACVR2B-AS1 | 1.008803377 | 0.014464679 |
| RPL21P93 | 0.278099576 | 0.016909764 |
| RPL4P4 | 0.191669885 | 0.000157865 |
| ITGB1-DT | 3.292346366 | 0.020096485 |
| RPL26P6 | 0.559653528 | 0.018270562 |
| ZNF492 | -0.189752681 | 0.035556241 |
| AC009237.3 | 0.903481773 | 0.00017486 |
| SOS1-IT1 | -3.189871353 | 4.16E-07 |
| AL136531.1 | 2.509324495 | 5.17E-05 |
| AL096803.2 | 6.823856256 | 2.69E-05 |
| MRPS18AP1 | 2.571663782 | 4.67E-05 |
| AL021396.1 | 8.701330674 | 4.52E-08 |
| AC013444.1 | 5.225633821 | 0.000114663 |
| XIST | -0.823001841 | 1.84E-10 |
| AL391825.1 | 0.544940883 | 0.045126335 |
| ZNF688 | 0.489123355 | 0.044629903 |
| DDX50P1 | 0.656066732 | 0.005834848 |
| ARSD-AS1 | 0.984449845 | 0.004627185 |
| PGA3 | 7.039400522 | 2.29E-05 |
| RPSAP20 | 7.769342796 | 1.32E-06 |
| RHEBP1 | -0.250834374 | 0.010561433 |
| EIF4EP2 | -0.955114655 | 1.32E-08 |
| TOB1-AS1 | 5.69831003 | 0.000504416 |
| RPL5P4 | 0.463252961 | 0.000832678 |
| ALMS1-IT1 | -1.460550287 | 1.18E-06 |
| LRRC37A15P | 1.645682159 | 3.14E-05 |
| RPL4P6 | 3.770632411 | 1.38E-07 |
| AL162231.2 | 4.723025631 | 5.10E-07 |
| RPL10P6 | 0.422659847 | 0.000917626 |
| MTAPP2 | -0.695264437 | 0.029644124 |
| TMEM254-AS1 | 5.518578676 | 1.02E-09 |
| ACBD6 | -1.036119023 | 8.96E-09 |
| AL021707.3 | 0.559750391 | 0.020221549 |
| CT45A11P | 2.717278548 | 0.000493856 |
| AL122010.1 | -2.481252098 | 0.001009157 |
| AL158156.1 | 1.4200493 | 0.017759401 |

| | | |
|---|---|---|
| AC008267.2 | 2.912074108 | 5.79E-06 |
| AL450405.1 | 0.305358743 | 5.80E-05 |
| FTH1P5 | 0.289892486 | 0.003431101 |
| RPL4P5 | 0.204593103 | 0.010499101 |
| PHBP9 | 0.842428748 | 0.046111489 |
| SPATA31C1 | -2.067064726 | 0.004107774 |
| LINC02603 | 0.438594514 | 0.049929394 |
| HNRNPA1P59 | 0.975431026 | 0.001581859 |
| AC078817.1 | 0.385195195 | 0.002640675 |
| AL353743.2 | 3.04535235 | 0.000792664 |
| AC010422.1 | 3.457123756 | 0.000433795 |
| AL022476.1 | 2.849473306 | 0.001193526 |
| AL359921.1 | 1.149513414 | 0.012354533 |
| TPI1P2 | 2.3368394 | 4.20E-08 |
| DDX10P2 | -0.699330109 | 0.013570297 |
| KARS1P2 | 6.959718539 | 1.80E-05 |
| AC009245.1 | 0.217833877 | 0.003057679 |
| EMC1-AS1 | 0.886220327 | 0.00780885 |
| TFAMP1 | 3.081635655 | 1.55E-07 |
| ANKRD18B | -0.588636872 | 0.000215833 |
| U73166.1 | 3.212954683 | 1.45E-07 |
| PA2G4P4 | -0.137394075 | 0.028024096 |
| PSMG3-AS1 | 3.613765054 | 1.06E-09 |
| AL139383.1 | 1.685920416 | 0.029305449 |
| RPL19P21 | 0.384368938 | 0.001787292 |
| THAP7-AS1 | 2.065687973 | 3.63E-07 |
| COL6A4P1 | 3.488094986 | 6.68E-06 |
| LIMD1-AS1 | -2.035909652 | 8.98E-08 |
| AL392046.1 | 5.163987785 | 1.22E-09 |
| AC021078.1 | -0.52987706 | 1.79E-07 |
| FAM133DP | 0.628876652 | 0.000528002 |
| ZNF32-AS2 | -1.036976129 | 0.004710569 |
| AL139095.3 | 3.341892784 | 0.014031093 |
| AC021016.1 | 0.555252136 | 0.000676703 |
| GTF2IRD1P1 | -1.905533835 | 2.73E-06 |
| PHBP12 | 4.225321339 | 0.001623669 |
| AC092683.1 | -1.243340364 | 8.63E-06 |
| RPS23P8 | 0.352242457 | 7.59E-06 |
| KLHL7-DT | 2.495519531 | 3.52E-05 |
| TNPO1P2 | -0.56659745 | 0.000123531 |
| THAP12P1 | -0.968481748 | 0.024768226 |
| CEACAM22P | 2.995622511 | 0.023721134 |
| ENO1-AS1 | 3.364872299 | 0.000115814 |
| LINC01001 | 1.13596267 | 0.003665503 |
| AC092171.2 | -0.984955395 | 3.88E-08 |
| AL021937.1 | 1.654929498 | 0.004307195 |
| VN1R108P | 2.57484816 | 0.000558632 |
| AC005532.1 | 4.113037473 | 0.004485764 |

| | | |
|---|---|---|
| AC241584.1 | 0.553773434 | 0.022514386 |
| LINC01293 | -0.416619231 | 0.001195002 |
| RPL31P2 | 1.189118856 | 0.045900048 |
| RPL23AP15 | 7.694574593 | 1.98E-06 |
| TSEN15P1 | -1.076574602 | 0.005540724 |
| AGAP10P | -0.503724628 | 0.043945029 |
| AC005077.4 | 3.356712258 | 0.001002054 |
| RPS18P12 | 0.453321784 | 2.46E-05 |
| AL391807.1 | 2.888578321 | 0.00023211 |
| AL139241.1 | 1.208850486 | 0.001675378 |
| AC079250.1 | 0.556839941 | 2.64E-05 |
| HSBP1 | 0.258379216 | 0.001712497 |
| RPL7P32 | 0.36333263 | 0.000416062 |
| CDC20P1 | -0.734725641 | 9.09E-06 |
| CR788250.1 | 0.3537159 | 0.002249198 |
| LINC00326 | 0.734193655 | 0.001492589 |
| LINC00271 | 2.715643891 | 0.000574951 |
| NPM1P9 | -0.544232172 | 0.034281352 |
| RPL22P24 | 2.488401281 | 1.73E-06 |
| AL031666.1 | 2.688853452 | 0.000423112 |
| MORF4L2-AS1 | 0.967481966 | 0.037746615 |
| PTP4A1P1 | 1.318781814 | 0.000231321 |
| COX11P1 | -0.96786746 | 0.012566007 |
| YBX1P2 | 0.246589506 | 0.002586255 |
| LINC00852 | 1.025044049 | 0.009917785 |
| SPRY4-AS1 | 1.763701558 | 0.005828944 |
| AL157884.2 | 7.080321009 | 1.83E-05 |
| ZNF826P | -0.883635694 | 4.28E-06 |
| CFAP58-DT | 4.124608493 | 1.32E-06 |
| C1DP1 | -0.37922289 | 0.008391257 |
| ITPR1-DT | 4.064234855 | 0.008610423 |
| CFAP97D1 | 7.401354604 | 5.71E-06 |
| AGER | 1.982319967 | 0.000541588 |
| AC112484.1 | 3.60374901 | 2.16E-08 |
| MAP4K3-DT | 2.92276774 | 1.50E-06 |
| BEND3P1 | -0.778156473 | 9.14E-05 |
| AC111200.2 | 1.056855277 | 0.002405095 |
| WARS2-AS1 | 0.592995331 | 0.000171745 |
| Z97353.1 | 0.383451364 | 0.027042202 |
| RNF2P1 | 1.648030024 | 0.000127436 |
| GORAB-AS1 | 2.534641623 | 4.32E-07 |
| AC018868.1 | 0.347058307 | 0.014478903 |
| AL358472.1 | 0.91439234 | 0.003370674 |
| LINC01516 | 2.038296259 | 0.008506535 |
| WASIR2 | 3.503921623 | 0.014279681 |
| RB1-DT | 4.644567237 | 2.35E-06 |
| FAM225A | -1.361347682 | 1.27E-07 |
| AL157932.1 | 3.046407491 | 0.000780672 |

| | | |
|---|---|---|
| HINT1P1 | 0.77830904 | 0.040996651 |
| LINC01249 | 1.146121624 | 0.049959198 |
| AC245100.4 | 0.959447141 | 0.00818972 |
| NEK2P2 | -0.704247947 | 0.02247737 |
| RPL7P21 | -0.639187178 | 0.000221919 |
| FAHD2CP | 1.961174457 | 0.000310443 |
| SHQ1P1 | 1.704905033 | 2.85E-06 |
| LINC01363 | 2.166456998 | 1.95E-07 |
| DLEU2 | -1.214817867 | 1.14E-07 |
| AC007098.1 | -1.16941015 | 0.013712952 |
| AL354733.2 | 1.997730594 | 0.001337729 |
| LUARIS | 1.312381039 | 0.044126471 |
| AL590428.1 | 1.364620384 | 0.000428027 |
| AL033397.1 | 1.130461136 | 0.011318889 |
| Z97180.1 | -0.949882486 | 3.19E-05 |
| AL356276.2 | -0.971250033 | 0.011879774 |
| AL354993.1 | 7.718972494 | 1.57E-06 |
| AC004895.1 | 4.323480052 | 0.001224117 |
| LINC00899 | 1.858675048 | 3.10E-05 |
| LINC-PINT | 1.475340357 | 2.74E-08 |
| VN1R110P | 9.315067239 | 5.81E-09 |
| AC079922.1 | 0.457620884 | 0.02787299 |
| AL136454.1 | 0.395915518 | 5.90E-06 |
| TMEM44-AS1 | 2.234224006 | 1.85E-08 |
| DBIL5P | 1.264277337 | 0.002156488 |
| PIK3CD-AS2 | 1.283510877 | 0.003382699 |
| DOC2GP | 2.889730447 | 0.000286948 |
| AC009502.1 | 1.573408721 | 0.018608555 |
| AC019097.1 | -0.457566404 | 0.000143455 |
| LINC01819 | 0.598152717 | 0.002887406 |
| AL513523.4 | 3.860120422 | 2.28E-05 |
| RPS7P2 | 7.12530457 | 1.13E-05 |
| AL162377.1 | 1.936031993 | 0.000119025 |
| AL807752.3 | -0.796557555 | 3.66E-05 |
| KF459542.1 | -1.236048004 | 5.17E-05 |
| GAPDHP37 | 7.43539413 | 4.03E-06 |
| TAPBP | -0.239430391 | 0.019147941 |
| RPS7P3 | 0.341197879 | 0.01517452 |
| AC004837.3 | 8.038493763 | 4.94E-07 |
| DPY19L1P2 | -0.62812456 | 0.016438893 |
| AC007383.3 | -0.935092824 | 0.002704584 |
| LINC02818 | 2.837791416 | 0.000441922 |
| AL355490.3 | -0.341353436 | 0.034974038 |
| ANXA2P2 | 0.187502703 | 0.000105377 |
| AC074183.1 | 1.521643148 | 1.42E-06 |
| LEF1-AS1 | -0.779459115 | 0.045299612 |
| FAAHP1 | 4.12890654 | 0.001821164 |
| LSM12P1 | -0.711655214 | 5.75E-07 |

| | | |
|---|---|---|
| ZBED9 | 1.632232296 | 4.56E-09 |
| SLC4A1APP1 | -1.197536418 | 1.27E-06 |
| TMEM253 | 2.343073185 | 7.33E-05 |
| RPL31P7 | 1.006084572 | 0.033544628 |
| AC012313.1 | 0.946409452 | 2.71E-06 |
| RFX3-AS1 | 2.808375052 | 6.06E-06 |
| TMA7 | 0.168328637 | 0.032026368 |
| RPS15AP12 | 0.359204612 | 0.000410366 |
| ST13P4 | 0.245171321 | 0.008824593 |
| RAP2C-AS1 | 2.973115665 | 0.028618386 |
| AL161909.1 | 0.339045922 | 1.02E-05 |
| AL137013.1 | -0.974398052 | 0.007441964 |
| FTH1P7 | 0.2027471 | 0.005643722 |
| AC098824.1 | -1.430406287 | 8.36E-05 |
| AC008440.1 | -0.661007172 | 0.018524485 |
| LINC02043 | 1.759367822 | 0.003582361 |
| RPS27AP2 | 1.180734521 | 0.013440094 |
| Z74021.1 | 0.290768516 | 0.00037818 |
| SEMA3B-AS1 | 2.616690647 | 0.000104311 |
| AL050404.1 | 1.075178686 | 4.78E-05 |
| TAP1 | -0.450519608 | 0.024542501 |
| FTLP2 | 0.41501031 | 0.004885099 |
| RPS3AP25 | 0.578109776 | 0.013026114 |
| AC015712.1 | 1.205318272 | 0.006078608 |
| SMIM26 | -0.217220693 | 0.042078168 |
| AJM1 | 0.641421483 | 0.002014206 |
| MHENCR | 1.831822331 | 1.18E-06 |
| EMSLR | -1.413910765 | 7.41E-07 |
| AL133517.1 | -2.513449866 | 0.002206614 |
| SLC16A6P1 | -1.096191884 | 0.013284138 |
| BMPR1AP2 | -0.920565701 | 9.90E-05 |
| TMA16P2 | -1.893041737 | 0.003715003 |
| EEF1B2P3 | 0.290437229 | 9.79E-05 |
| AC010900.1 | 2.243730971 | 0.004114067 |
| AC098820.2 | 1.841752955 | 0.005378582 |
| AL592437.2 | -1.473115073 | 1.41E-05 |
| AL391058.1 | 0.226005402 | 0.026553898 |
| ETF1P1 | -0.865273085 | 0.048491384 |
| AC027612.3 | -0.342982144 | 0.02871602 |
| AC093673.1 | 0.668713796 | 0.017448908 |
| RPL3P4 | 0.586772842 | 1.83E-08 |
| KIAA1614-AS1 | -0.721585782 | 0.035195078 |
| KANTR | 1.004196929 | 0.001426527 |
| AL683813.2 | 1.264141904 | 0.018369164 |
| AC026412.2 | -0.44670389 | 0.009349012 |
| PRPS1P2 | 0.782330735 | 0.000795686 |
| LINC01431 | 1.459639877 | 0.012180672 |
| GOLGA8N | 4.502458093 | 1.20E-06 |

| | | |
|---|---|---|
| IDI2-AS1 | 2.404245797 | 2.92E-09 |
| PHBP10 | -0.559848457 | 0.040558993 |
| ZNF687-AS1 | 2.32856074 | 1.69E-06 |
| LINC00665 | -0.597471327 | 9.64E-07 |
| BDH2P1 | 0.469843003 | 0.047973443 |
| NUTM2HP | 2.690570672 | 0.002813074 |
| AC135050.1 | -1.647260907 | 4.37E-07 |
| BMS1P22 | -1.458010595 | 0.000117395 |
| SDCBPP3 | 0.455323606 | 0.029502257 |
| SLCO4A1-AS1 | 0.524750626 | 0.036191074 |
| DUX4L50 | 1.102003447 | 0.007775663 |
| AC231533.1 | -1.305674888 | 0.000372084 |
| PET117 | -0.847966033 | 5.22E-07 |
| AL137845.2 | -0.882446852 | 0.000260818 |
| AL590708.1 | 3.525211349 | 1.85E-05 |
| LYRM9 | 4.360384135 | 0.000149043 |
| AL451081.2 | 0.412759377 | 0.046342434 |
| RPS11P5 | 0.376034532 | 0.001119783 |
| USP12PY | -1.395125939 | 0.001860168 |
| LINC00342 | -1.39360923 | 1.97E-05 |
| RPL23AP87 | 1.66477542 | 0.000795889 |
| AL512844.1 | 0.250564755 | 0.007434379 |
| SNHG15 | -0.69705806 | 6.87E-06 |
| LINC00327 | -1.295170708 | 0.003380426 |
| AL592435.1 | 0.712357527 | 0.014517925 |
| FAM157B | 0.861327897 | 0.013359876 |
| SNHG7 | 0.896666092 | 7.46E-09 |
| AL354872.1 | -1.187960513 | 0.010509757 |
| AC006970.2 | 3.337844771 | 0.000301736 |
| AC244453.2 | 5.224523569 | 2.93E-07 |
| AC243772.2 | 5.712748783 | 5.43E-10 |
| EEF1A1P14 | 0.785179545 | 0.003091679 |
| LINC01271 | 1.197397276 | 0.002270176 |
| RPL23AP25 | 1.766763932 | 5.54E-06 |
| CCDC144NL-AS1 | -0.710628482 | 1.70E-07 |
| HOXB-AS3 | -1.334067512 | 0.000180097 |
| VN1R20P | 3.414937608 | 0.000279501 |
| CTAGE7P | 1.084732714 | 0.000850776 |
| HMGA1P8 | 0.94198262 | 0.000776696 |
| AC138356.2 | -0.638083933 | 0.0012915 |
| AC008105.1 | 3.946697859 | 7.72E-10 |
| AL161457.2 | 1.008712264 | 0.01366138 |
| AC093157.1 | 3.854368311 | 1.35E-10 |
| RNF224 | 1.309490331 | 0.003649624 |
| AC108479.1 | 0.841291128 | 0.000320724 |
| AC004987.2 | -0.372122288 | 0.01891652 |
| NPIPB7 | 3.995258284 | 1.19E-05 |
| AC002524.1 | -0.865019104 | 6.10E-06 |

| | | |
|---|---|---|
| RPL21P134 | 0.179483649 | 0.016120848 |
| AC006042.2 | -1.35222337 | 0.030740727 |
| GPX1 | 0.242045956 | 0.000255561 |
| AC009362.1 | 0.338366031 | 0.01031899 |
| AC092447.5 | 1.507587246 | 0.000444876 |
| TMEM18-DT | 0.970070997 | 0.006861283 |
| RASA4DP | 1.493947111 | 0.004713997 |
| LINC01346 | 6.998228118 | 2.41E-05 |
| MIPEPP3 | 1.485054936 | 0.000171631 |
| PFN1P1 | 0.31610601 | 0.002579904 |
| Z83844.2 | 1.470952978 | 0.003341588 |
| AL139002.1 | -1.251118687 | 0.005003544 |
| LINC01719 | 3.740016637 | 3.86E-12 |
| AL162430.2 | 0.436722958 | 0.000291161 |
| EIF3FP3 | 0.24812239 | 0.000393672 |
| AL009181.2 | -0.864779116 | 0.024749027 |
| HOTAIRM1 | 1.719600451 | 4.19E-07 |
| AGGF1P2 | 3.29124691 | 7.61E-06 |
| AC125238.2 | -2.055127714 | 3.67E-05 |
| AL445524.1 | 0.597605077 | 0.007894588 |
| EEF1A1P6 | 0.423773668 | 1.04E-08 |
| AC008105.2 | 3.027665699 | 9.99E-06 |
| OR5B1P | 6.825976011 | 2.94E-05 |
| AL356653.1 | -0.304770549 | 0.003403306 |
| RANP8 | 2.8903706 | 0.000807162 |
| ZNF529-AS1 | 0.571769684 | 0.041107925 |
| B4GALT1-AS1 | 1.874044863 | 0.001682042 |
| AL050331.1 | 0.491683879 | 2.29E-05 |
| LINC00513 | 4.915605156 | 9.11E-10 |
| AC231533.2 | 0.42146725 | 0.02614067 |
| AC246785.2 | 6.588863005 | 0.013698421 |
| CYP51A1P2 | 1.234492516 | 2.22E-07 |
| AL138789.1 | -1.07791435 | 0.000110911 |
| LINC02609 | 2.060067444 | 3.42E-05 |
| BTF3P5 | 0.24696607 | 0.043116398 |
| TWIST2 | -1.168055186 | 0.000381276 |
| LINC01137 | 0.746734582 | 0.008907259 |
| GAS6-AS1 | 2.943577526 | 5.23E-09 |
| SLC26A4-AS1 | 6.606980738 | 2.22E-06 |
| AL512353.2 | 3.228944968 | 0.01512263 |
| AL929472.3 | -1.079322153 | 4.40E-05 |
| AC007969.1 | 0.299309289 | 0.000630146 |
| CDC27P1 | -1.323212591 | 0.000162876 |
| AC139887.1 | 1.669440844 | 0.013610034 |
| H2BC15 | 2.46988604 | 2.18E-06 |
| EIF4HP1 | -0.134617676 | 0.019381451 |
| AC093732.1 | 2.670961093 | 0.002428459 |
| SELENOTP1 | -0.780135638 | 1.67E-05 |

| | | |
|---|---|---|
| BX927235.2 | 1.467836187 | 0.023181223 |
| MGAT2P1 | -1.33495361 | 0.020841318 |
| AC009302.1 | -0.925219851 | 1.72E-05 |
| YEATS2-AS1 | -0.594260775 | 0.00892899 |
| GTF2F2P2 | 6.673854288 | 4.15E-05 |
| AC026202.2 | 2.258808647 | 0.004522113 |
| RPL10P9 | 0.497066693 | 4.69E-07 |
| RPS28 | 0.401195824 | 3.16E-05 |
| MT1XP1 | 1.629731804 | 0.000335565 |
| AC008443.1 | 1.041057181 | 3.65E-08 |
| UQCRHL | 0.223523549 | 0.002345276 |
| UBE2SP1 | -0.726221607 | 3.50E-08 |
| RPSAP14 | 0.491041687 | 0.002579904 |
| GDI2P2 | -0.336272901 | 0.011929666 |
| AC092017.2 | 0.675214386 | 0.001553226 |
| GAPDHP22 | 10.13274416 | 6.88E-10 |
| RPL5P34 | 0.321864967 | 0.002842135 |
| EIF2AK3-DT | 2.974170594 | 3.37E-07 |
| TXNP6 | 0.32896489 | 0.000462675 |
| PAGE2 | 0.334005897 | 0.001855114 |
| AC074117.1 | 1.703900738 | 1.24E-08 |
| AL049552.1 | 2.976462019 | 0.000146587 |
| RPS15AP11 | 0.554995213 | 0.000388117 |
| SRP14P2 | 0.31420697 | 0.020712398 |
| TPT1P1 | 0.757301506 | 0.019880034 |
| AL138878.2 | 1.022662394 | 0.010646959 |
| RPL13AP6 | 0.615861866 | 0.001562225 |
| AC073529.1 | 1.523490714 | 0.005103626 |
| AL158835.2 | 2.68217253 | 0.000582657 |
| NAP1L4P3 | 0.168423757 | 0.037033029 |
| AL035446.1 | 0.92628867 | 0.003181177 |
| AL513165.1 | -1.097914849 | 2.07E-05 |
| RNASEH1-AS1 | 1.144438588 | 5.66E-07 |
| HSPA8P1 | -0.12931453 | 0.019136817 |
| AC244394.2 | 7.371611122 | 4.97E-06 |
| ZFX-AS1 | 2.478729464 | 0.001017269 |
| AL109618.1 | -0.835174024 | 1.99E-08 |
| AP000936.3 | 0.705312551 | 0.000972771 |
| AC099560.2 | 0.358011594 | 3.04E-06 |
| AL592293.2 | -0.504490217 | 5.05E-05 |
| ST13P18 | 0.25709543 | 0.030112304 |
| AL604028.2 | 1.932686783 | 2.06E-05 |
| RPS4XP11 | 0.402748601 | 0.003215443 |
| AC007405.2 | 7.404211953 | 4.43E-06 |
| PFN1P3 | 3.67642185 | 2.68E-08 |
| TATDN1P1 | 0.667697847 | 0.000100502 |
| USP27X-AS1 | 1.347401925 | 0.000351887 |
| AL592546.2 | 2.490814438 | 0.000483479 |

| | | |
|---|---|---|
| CCDC188 | 1.870803868 | 0.001059769 |
| ZNF37BP | 0.191179178 | 0.003629508 |
| AL353743.4 | 2.996870474 | 4.21E-06 |
| BX005266.3 | 7.375255581 | 6.27E-06 |
| MAGI2-AS3 | 0.78640377 | 6.64E-08 |
| PINLYP | -2.04065486 | 0.01547903 |
| ACBD3-AS1 | 0.743626971 | 0.002909159 |
| RPL34-AS1 | 3.352293861 | 0.000642488 |
| SP2-AS1 | 1.221509407 | 0.000192874 |
| RPL13AP20 | 0.501802061 | 8.96E-06 |
| AC073072.1 | -0.628823241 | 0.046900179 |
| PPP1R2P1 | 2.111213614 | 0.001580233 |
| PTGES3P1 | -0.26575135 | 0.000177057 |
| AC005041.2 | 7.467967279 | 3.75E-06 |
| AL161719.1 | 1.462285668 | 0.002491168 |
| AL354751.1 | 3.891820017 | 2.11E-06 |
| FAM133B | 0.316864356 | 0.000437797 |
| LNCTAM34A | 1.611900937 | 0.00416061 |
| API5P1 | -0.44552442 | 0.000116821 |
| H2BP2 | 6.549033739 | 7.43E-10 |
| AL355483.3 | 2.932217771 | 0.001177748 |
| CCT6P3 | 0.306743989 | 0.041392147 |
| AL161449.1 | 0.988408365 | 0.046934722 |
| AC125807.1 | 0.704970513 | 0.000732835 |
| MCIDAS | -1.304672969 | 1.79E-05 |
| MAPKAPK5-AS1 | -0.205284136 | 0.034680951 |
| JRK | -0.69847779 | 1.89E-06 |
| RPSAP9 | 0.42568384 | 2.16E-05 |
| MED14OS | -0.993142906 | 0.024780498 |
| AL162151.2 | 0.585323079 | 9.09E-06 |
| LERFS | 0.75885155 | 0.000182452 |
| SDCBP2-AS1 | 2.563954639 | 3.13E-10 |
| AL139289.1 | 2.09280926 | 8.45E-05 |
| AC074351.1 | 8.539219162 | 1.04E-07 |
| UPF3AP3 | 1.6108926 | 0.022801914 |
| NPIPB2 | 1.661020266 | 0.000110795 |
| GAS5 | -0.975034283 | 1.69E-09 |
| AC144530.1 | 0.208109251 | 0.017661458 |
| MGAT4FP | 6.722959852 | 3.84E-05 |
| E2F6P1 | -0.587906066 | 0.001031375 |
| WASH4P | 0.654368985 | 0.001758836 |
| AC012618.3 | 2.441272227 | 0.000123111 |
| TPT1P9 | 0.496155947 | 2.39E-06 |
| RPS3AP6 | 0.375582742 | 1.71E-07 |
| ECI2-DT | 3.333192787 | 1.99E-05 |
| AC007314.1 | -1.303329277 | 0.000954424 |
| AL353705.4 | -0.599494007 | 0.000373943 |
| CR759913.1 | 0.455076186 | 0.020406516 |

| | | |
|---|---|---|
| RPL23AP42 | 0.320705005 | 9.33E-07 |
| HNRNPUL2-BSCL2 | -0.255491612 | 0.000238517 |
| TEX21P | 2.841830586 | 7.76E-05 |
| SNHG20 | -0.68946466 | 0.000279483 |
| AC098484.2 | 3.252660716 | 0.000230657 |
| ATP5PDP4 | 0.465049867 | 0.00613091 |
| AC010883.1 | 2.09557665 | 0.000378208 |
| GTF3C2-AS1 | -0.902556652 | 6.28E-05 |
| AL133415.1 | 0.499921254 | 2.12E-06 |
| FABP5P7 | -0.334210821 | 0.000645808 |
| FTH1P2 | 0.165131049 | 0.007974172 |
| AC244034.1 | 0.476890031 | 0.001668236 |
| AC098934.2 | 3.477751173 | 6.86E-10 |
| AC244453.3 | 2.154961144 | 0.000803556 |
| AL023775.1 | -1.292228355 | 2.19E-06 |
| DAAM2-AS1 | 1.205163845 | 0.004531798 |
| C19orf81 | 3.966019384 | 2.98E-08 |
| AL035456.1 | -1.529747236 | 4.04E-07 |
| TECRP1 | 0.208286181 | 0.004102407 |
| PPIAP3 | 2.985489162 | 1.46E-05 |
| AL021154.1 | 3.488069192 | 3.12E-07 |
| AC245052.3 | 1.745882926 | 0.000505432 |
| SUMO1P3 | -0.252419991 | 0.009311364 |
| CHCHD2P6 | 0.173535901 | 0.018876236 |
| AC099654.2 | 1.458864693 | 3.23E-05 |
| ANKRD65 | 5.499252182 | 5.17E-08 |
| AL157400.3 | 2.95883934 | 0.000849837 |
| BRD3OS | -0.179277302 | 0.017795522 |
| ZSCAN31 | 2.362677033 | 1.44E-06 |
| AL445931.1 | 1.224494129 | 0.040283748 |
| LINC02532 | 4.274434841 | 0.001376888 |
| RPSAP16 | 7.793610381 | 1.20E-06 |
| B3GALT4 | 7.997296219 | 5.72E-07 |
| AL121672.2 | 3.111783577 | 0.000201198 |
| C12orf75 | 0.189187877 | 0.011506751 |
| LINC01366 | 1.677371109 | 0.000923505 |
| HGH1 | -0.379906109 | 0.000288089 |
| RPL39P3 | 0.364368957 | 7.19E-07 |
| PPP1R3E | 0.64668252 | 0.000981112 |
| TATDN2P3 | 1.291445592 | 0.013484614 |
| TSPY26P | 2.775568716 | 8.24E-06 |
| HLA-F | 1.097747703 | 0.015257157 |
| MSH5 | 0.424499062 | 0.047962206 |
| AC016712.1 | 0.396746701 | 0.002904273 |
| Z82188.2 | 2.157496328 | 0.006596021 |
| BX284668.4 | 1.228042245 | 0.046131408 |
| AL360181.2 | 1.631017033 | 0.000282674 |
| AC093155.2 | -0.6631264 | 0.006008763 |

| | | |
|---|---|---|
| AC074117.2 | 3.689106542 | 0.000237159 |
| RAMACL | -0.561292298 | 3.55E-05 |
| MCF2L-AS1 | 1.776815676 | 0.000965634 |
| AP000925.1 | 3.707719848 | 0.004043307 |
| FAUP1 | 0.312401525 | 0.003407273 |
| AL354733.3 | -0.609145687 | 0.024287061 |
| HM13-IT1 | -1.217305456 | 0.00774457 |
| LINC00957 | 1.924248154 | 0.024839488 |
| DUSP8P5 | -0.849327727 | 0.011534803 |
| PVRIG2P | -0.449908619 | 0.026643782 |
| RPS29P16 | 2.788768474 | 9.34E-05 |
| SSR4P1 | 2.72940169 | 9.31E-08 |
| RPEL1 | -1.010459875 | 0.000110499 |
| AL596202.1 | 1.034437611 | 4.79E-05 |
| AC113174.1 | 0.935140135 | 0.001469578 |
| SPAAR | 2.714773258 | 0.00025295 |
| NOTCH4 | 2.019560572 | 0.001198043 |
| AC104837.1 | -0.645766843 | 0.010079814 |
| CYMP-AS1 | 8.634505662 | 6.09E-08 |
| AC007041.1 | -0.76413732 | 0.000413494 |
| KRT18P63 | 2.730552005 | 0.00030471 |
| LINC01278 | -0.593350453 | 0.000208929 |
| AC098934.3 | 0.370294677 | 0.001003316 |
| HAUS6P3 | -0.494129001 | 5.93E-05 |
| DBF4P1 | -0.842235328 | 7.41E-07 |
| AC073046.1 | -2.087840826 | 1.33E-05 |
| RPS2P7 | 0.475002656 | 0.000293914 |
| HIPK1-AS1 | 5.654020931 | 5.62E-05 |
| MSC-AS1 | -1.091883499 | 6.03E-05 |
| AC087499.4 | 0.475687838 | 0.014468199 |
| RPL6P27 | 0.294465867 | 7.62E-07 |
| AC005822.1 | -1.937615986 | 3.35E-09 |
| SUMO1P4 | -0.390904367 | 0.026402366 |
| GAPDHP65 | 0.223264212 | 0.001497299 |
| AC004674.2 | 0.627458154 | 0.010517344 |
| AL355472.1 | 0.301847222 | 0.000950631 |
| FAM21EP | 1.427986476 | 0.012684608 |
| RNF148 | 3.21869896 | 1.84E-05 |
| NUS1P1 | -0.826960453 | 1.63E-06 |
| LINC01647 | 3.152097254 | 5.43E-07 |
| AC064850.1 | -0.729281255 | 0.000794543 |
| AL356599.1 | -0.324407857 | 0.019888843 |
| H3P6 | -0.302834119 | 1.21E-05 |
| AC090286.3 | 1.137521755 | 0.02974211 |
| ABHD16A | 0.888906077 | 0.007922231 |
| NPM1P26 | 0.507614661 | 0.0043341 |
| LINC00894 | 0.732967463 | 0.005812227 |
| DICER1-AS1 | 4.366410257 | 0.002718645 |

| | | |
|---|---|---|
| ANKRD34C | 10.42833976 | 2.33E-10 |
| GABPAP | -0.791993606 | 2.18E-06 |
| AC013268.3 | 2.997522749 | 0.000953525 |
| AC009299.3 | 8.231318499 | 2.50E-07 |
| AL390860.1 | 1.88153918 | 0.000117865 |
| KIAA0040 | -0.524068731 | 6.21E-06 |
| AC000089.1 | 0.496750415 | 3.65E-06 |
| OLMALINC | 3.107287084 | 6.29E-12 |
| AC005540.1 | -1.179572421 | 0.000650225 |
| GSN-AS1 | 1.199318361 | 5.45E-05 |
| AC078777.1 | -1.012500368 | 0.000124312 |
| TSPEAR-AS1 | 1.708015969 | 4.02E-05 |
| CPB2-AS1 | 4.054694069 | 5.37E-07 |
| ASH1L-AS1 | 0.524353035 | 0.031316523 |
| NEXN-AS1 | 1.364763357 | 0.001347622 |
| COX7CP1 | 0.483163359 | 0.005729398 |
| MORC2-AS1 | -1.650010009 | 0.000247847 |
| AC004898.1 | -0.553522195 | 0.04716567 |
| AL591043.2 | 4.15098949 | 0.004335883 |
| MYCBP2-AS1 | -0.998593808 | 0.000441922 |
| GAPDHP14 | 0.502266202 | 0.025498525 |
| RPL17P36 | 0.208729576 | 0.007231379 |
| HSPB1P1 | 0.836755719 | 0.000300921 |
| AL020995.1 | 1.677301582 | 0.011595453 |
| ACTG1P4 | 3.152314628 | 8.32E-05 |
| COX10-AS1 | 0.21370358 | 0.024192887 |
| ZBTB22 | 0.466703447 | 0.024417479 |
| PRELID3BP10 | 0.523244255 | 0.027012061 |
| AL445231.1 | 4.886746308 | 3.83E-08 |
| AC245014.1 | 5.973468487 | 4.51E-05 |
| TMEM147-AS1 | 0.967037424 | 7.45E-08 |
| PRADC1P1 | 2.473419894 | 4.59E-05 |
| KDM4A-AS1 | -0.9549244 | 0.000245515 |
| LINC01376 | 3.752313134 | 1.78E-06 |
| VEZF1P1 | -0.961561355 | 0.000564286 |
| AC009404.1 | 1.366393145 | 0.000323576 |
| EI24P2 | -0.49373614 | 0.045522742 |
| ZBED5 | -0.87892901 | 1.69E-08 |
| EEF1E1P1 | 1.131955141 | 0.04629047 |
| RPL34P34 | 0.481963423 | 0.025918585 |
| NCR3 | 6.961651229 | 1.88E-05 |
| TRHDE-AS1 | -0.959950968 | 4.33E-06 |
| SCAT8 | 0.510157548 | 0.004305857 |
| AL445183.3 | -0.805542963 | 0.000504232 |
| CCDC200 | 4.039290245 | 2.31E-09 |
| DDX11L2 | 1.976051752 | 0.000284945 |
| AC009237.8 | 0.911621784 | 0.033060895 |
| MFF-DT | 3.360808687 | 4.45E-07 |

| | | |
|---|---|---|
| AC099336.2 | 0.416531097 | 3.36E-05 |
| UBE2L5 | -0.561576479 | 0.002001061 |
| AL035458.1 | -0.862233651 | 4.54E-05 |
| GCNT1P1 | 8.689919787 | 5.04E-08 |
| PKMP1 | 0.229169727 | 0.005484666 |
| AC107081.2 | 0.240294376 | 0.003932984 |
| LINC00896 | 3.422076364 | 0.000166381 |
| AL135791.1 | 1.980544878 | 0.002123339 |
| LINC01424 | 1.237658269 | 0.038222509 |
| NPM1P40 | 0.474953339 | 0.01948139 |
| KPNA2P1 | -1.496008749 | 0.00538728 |
| H3P36 | -0.322538833 | 0.006614396 |
| MED28P7 | 2.890362793 | 0.033317752 |
| RPL13AP5 | 0.51136141 | 1.12E-06 |
| RAD23BP1 | -0.578220873 | 0.001312149 |
| ZNF853 | 0.62744847 | 0.000931502 |
| AC127070.1 | 1.983906211 | 0.002909795 |
| PPT2 | 0.34607864 | 0.01141318 |
| PRKG1-AS1 | 1.912876789 | 0.033884287 |
| MTX1P1 | 0.348897969 | 0.029288623 |
| MAP3K2-DT | 0.916843277 | 0.02416155 |
| HLA-DPB1 | 0.476347163 | 0.040720082 |
| EIF1AXP1 | -0.295714104 | 4.37E-05 |
| ARHGEF38 | -0.722114006 | 0.049909469 |
| LINC01010 | 7.264438095 | 1.92E-05 |
| LINC01844 | 2.31974216 | 0.010192864 |
| RPL31P63 | 0.543602004 | 0.002137738 |
| COX7A2P2 | 0.677449239 | 0.012230019 |
| AC092809.4 | -0.500594912 | 0.000365335 |
| INTS6-AS1 | 3.924921861 | 3.92E-11 |
| AL391650.1 | -0.23587251 | 0.040575676 |
| AL513175.2 | -0.385423674 | 0.015088836 |
| AL157884.3 | 4.681488325 | 0.002738036 |
| RPL24P8 | 0.408038198 | 0.000149177 |
| SNX25P1 | -0.734763821 | 0.002630598 |
| ELOA-AS1 | -1.819339291 | 2.35E-08 |
| BCYRN1 | -0.692790463 | 0.012326929 |
| Z97634.1 | 1.361323531 | 0.004983238 |
| AC024560.1 | 7.612627638 | 2.25E-06 |
| BET1-AS1 | 3.010561638 | 7.53E-05 |
| LINC01554 | 1.746830677 | 0.019494687 |
| MIR600HG | 0.762491737 | 0.000875596 |
| AC109635.1 | 2.895630887 | 0.001427027 |
| AL157937.1 | 0.71834656 | 0.030170074 |
| AL162411.1 | 3.004629614 | 1.71E-05 |
| AL157938.2 | -0.804746488 | 0.011630738 |
| EDRF1-AS1 | -1.387590033 | 0.000245238 |
| RPL12P12 | 0.415895269 | 0.012481086 |

| | | |
|---|---|---|
| ZEB1-AS1 | 0.944059283 | 0.000202543 |
| AC018738.1 | 0.413755514 | 2.39E-05 |
| AC005550.2 | -1.500670168 | 0.00510747 |
| AL512353.3 | 2.968731358 | 0.000361857 |
| AL732372.2 | 1.083274908 | 0.00092801 |
| AC073254.1 | -1.493697538 | 2.75E-07 |
| AC242426.2 | 1.057504668 | 0.000275248 |
| AL449403.2 | 3.333222908 | 7.07E-05 |
| AL080243.4 | 0.495073974 | 8.90E-07 |
| Z99289.1 | 2.140070841 | 4.60E-06 |
| FTH1P11 | 0.2351472 | 0.004344864 |
| AC092447.7 | 1.385446463 | 0.000213206 |
| ANO7L1 | 2.827921933 | 0.001726132 |
| RLIMP2 | -0.608428122 | 0.031926382 |
| SMG1P1 | 0.321792427 | 0.003653956 |
| TTN-AS1 | 2.320069666 | 0.000942199 |
| AL121992.1 | 4.482748101 | 2.16E-06 |
| SRRM1P3 | -1.40201154 | 0.001543545 |
| AC008267.5 | 1.671715419 | 0.000103706 |
| AC246785.3 | 8.717282443 | 6.69E-08 |
| CDC42P6 | -0.374944527 | 1.70E-05 |
| BX088651.4 | 0.635839647 | 0.04986208 |
| NRBF2P3 | 0.978208007 | 0.019769628 |
| XRCC6P1 | 0.285380559 | 0.010917642 |
| FOXD2-AS1 | 1.032126893 | 0.000163884 |
| RPSAP11 | 1.368782874 | 0.007518986 |
| CAMTA1-DT | 2.95240052 | 0.000212912 |
| ASS1P12 | 1.62430381 | 4.51E-06 |
| CECR7 | -0.643176083 | 0.016314361 |
| CDC27P2 | -1.559267294 | 0.001825017 |
| CDK2AP2P2 | 3.371365718 | 0.000127657 |
| C10orf143 | 3.328217163 | 7.56E-07 |
| LINC01409 | 1.526225817 | 1.24E-06 |
| AC034102.1 | 0.511902879 | 1.82E-06 |
| WAKMAR2 | 3.209826207 | 5.37E-07 |
| PKN2-AS1 | 6.957783397 | 1.99E-05 |
| RPSAP15 | 0.319015254 | 7.61E-05 |
| LINC00857 | -1.127471944 | 1.31E-05 |
| AC243919.1 | 0.288347765 | 5.85E-06 |
| PYY2 | 1.70800004 | 0.004939888 |
| OXCT2P1 | -0.875031236 | 0.015621404 |
| AC004522.4 | 7.164734951 | 1.13E-05 |
| C2orf74 | 1.126590405 | 5.00E-06 |
| RPS15AP38 | 0.51913079 | 0.030725258 |
| KRR1P1 | 4.222339093 | 1.68E-07 |
| Z95152.1 | 2.331910422 | 8.92E-05 |
| AC002075.2 | 0.474496948 | 0.018925776 |
| AC078991.1 | 0.420908429 | 0.001591322 |

| | | |
|---|---|---|
| DCTN1-AS1 | 2.603983262 | 0.00459628 |
| AL034379.1 | 1.01891209 | 0.003248451 |
| AC079922.2 | 1.012349736 | 6.90E-05 |
| FAM200B | -0.424681348 | 0.000214222 |
| GGTA2P | 2.367386572 | 0.000360257 |
| AC073332.1 | 1.077256585 | 0.003048851 |
| ADAMTSL4-AS2 | 5.263937201 | 1.22E-07 |
| LINC00211 | 4.672982303 | 0.000718677 |
| AL157713.1 | -0.267676045 | 0.001877882 |
| AL773545.3 | 0.913327634 | 0.017858861 |
| NFYAP1 | 3.226833082 | 2.67E-09 |
| AC073130.1 | -3.306008594 | 6.04E-05 |
| NALT1 | 2.73415762 | 4.39E-05 |
| AC097523.2 | 1.600761999 | 0.000542732 |
| PRKCQ-AS1 | -1.648565221 | 8.72E-06 |
| MTCO1P12 | 0.219065331 | 9.53E-06 |
| FLG-AS1 | -0.189891644 | 0.047461343 |
| PTENP1 | -1.27080868 | 4.43E-09 |
| RPL35P1 | 0.285688711 | 0.021961029 |
| AC116347.1 | 0.317709955 | 0.044444554 |
| AC093110.1 | 0.635920165 | 0.011876643 |
| ARMC4P1 | 4.454753312 | 0.000696468 |
| AC138035.1 | 1.092201184 | 7.05E-06 |
| AC009133.1 | -0.523650112 | 1.97E-07 |
| AL355574.1 | 1.185400278 | 0.007643478 |
| AC009244.1 | -0.306435486 | 0.001497444 |
| LRRC37A2 | 0.645962137 | 0.025226094 |
| AL031284.1 | -0.281783824 | 0.020419455 |
| RPL9P7 | 0.446107248 | 1.50E-05 |
| GOLGA2P5 | 0.958362957 | 0.000160595 |
| LINC02806 | 7.637223491 | 2.29E-06 |
| LINC01410 | 1.505144527 | 3.36E-06 |
| Z95327.2 | -0.327040674 | 0.034246346 |
| ARPC3P2 | 1.40614493 | 0.005271851 |
| AC008753.1 | 0.558605796 | 0.049319549 |
| RPS2P35 | 0.639460932 | 0.000344666 |
| AHCYP2 | 3.404805028 | 1.55E-05 |
| PAXBP1-AS1 | 2.43079259 | 5.36E-08 |
| LRIG2-DT | 1.638967949 | 0.003225445 |
| TARDBPP2 | -0.884623814 | 0.030065181 |
| TMEM250 | -0.988328229 | 9.75E-09 |
| OR2W3 | 0.763935714 | 0.009500215 |
| MYO5BP2 | 1.913872049 | 5.65E-05 |
| AC006210.2 | -1.194501624 | 4.43E-05 |
| LINC00707 | -1.116078423 | 5.26E-08 |
| PAGE2B | 1.31044324 | 0.006663833 |
| AL603839.3 | 0.98081829 | 0.030604129 |
| SCARNA12 | -1.209051711 | 0.000666265 |

| | | |
|---|---|---|
| SNORD125 | -2.258906849 | 0.012374869 |
| NCK1-DT | 2.083152257 | 9.47E-06 |
| AC008026.1 | 0.341304239 | 0.000429182 |
| CLRN1-AS1 | 7.33974928 | 5.51E-06 |
| CASTOR1 | 1.125860309 | 0.012393025 |
| RNF103 | -0.276378211 | 0.000821544 |
| RBM14 | -2.579223527 | 2.81E-12 |
| ATP6V1B1-AS1 | -2.417309753 | 1.56E-06 |
| LLPH-DT | 1.002624013 | 0.027601409 |
| HNRNPA1P26 | 0.640493496 | 0.024900325 |
| ALKBH6 | 0.340397205 | 0.03896086 |
| PCDHA13 | -1.411036608 | 5.21E-09 |
| TMED10P2 | 3.546043753 | 0.010514531 |
| Z68871.1 | -1.454931536 | 0.002344871 |
| KCNMB3P1 | -1.763727224 | 0.013223907 |
| AC091212.1 | 2.285495641 | 0.000422229 |
| AC011979.1 | 0.210311367 | 0.019787443 |
| RPL7P38 | 8.349497096 | 1.95E-07 |
| CASTOR3 | 0.681029185 | 0.001285919 |
| MYLK-AS1 | 1.579982737 | 0.002869855 |
| RPL37P2 | 0.199450904 | 0.014841759 |
| HMGN1P7 | 6.8249165 | 2.75E-05 |
| AC073610.1 | 2.547857361 | 4.55E-05 |
| AC004865.2 | 4.026240568 | 0.002181966 |
| AL157392.3 | -0.505401045 | 2.25E-05 |
| AL355864.2 | 8.283657457 | 2.03E-07 |
| NME1 | 0.211057 | 0.001224507 |
| APOBEC3G | 0.950889072 | 4.08E-05 |
| AC004890.1 | 1.280617762 | 0.000115974 |
| TLR9 | 5.368996229 | 4.22E-10 |
| WBP1 | 0.494838619 | 1.38E-05 |
| MRPS17 | -0.502108843 | 0.000127595 |
| AC002310.2 | 1.879748716 | 0.005954204 |
| RPL21P39 | 0.523128582 | 0.002740689 |
| SUGT1P3 | 1.527416501 | 0.00686245 |
| AC063944.1 | 7.693404731 | 1.74E-06 |
| GET4 | -0.46818021 | 7.01E-05 |
| GAPDHP27 | 3.443518984 | 5.57E-06 |
| C1orf226 | -0.617469435 | 1.33E-05 |
| ADSL | -0.223021628 | 0.002912208 |
| STIM2-AS1 | 2.431629731 | 0.002332723 |
| AL133163.1 | 4.45138939 | 3.22E-06 |
| LINC00888 | 1.443056081 | 1.91E-06 |
| AC104563.1 | 0.401689497 | 4.07E-05 |
| AMY2B | 2.013043457 | 2.65E-08 |
| AC069431.1 | 7.887705527 | 7.69E-07 |
| RPSAP12 | 0.248428707 | 0.001653337 |
| ETF1P2 | -0.593717262 | 0.005750853 |

| | | |
|---|---|---|
| PSMD12P | -0.634031611 | 0.004610615 |
| AC087385.1 | -2.286688001 | 0.001498209 |
| PCDHGC3 | -0.109769305 | 0.035160042 |
| CYMP | 7.303348702 | 6.85E-06 |
| AL512306.2 | 6.182186865 | 2.13E-05 |
| ZNF542P | -0.374071339 | 0.000158657 |
| COX19 | -0.964252467 | 1.31E-05 |
| RPS27P29 | 0.377135423 | 0.002980781 |
| GAPDHP33 | 2.363449621 | 1.04E-05 |
| GHRLOS | 2.427153811 | 4.37E-07 |
| AL450384.2 | 2.517501699 | 2.86E-05 |
| RPS2P5 | 0.546040074 | 3.30E-09 |
| PPIL3 | -0.362654428 | 0.014416107 |
| RPL23AP7 | 0.973994109 | 0.000335065 |
| RPS4XP13 | 0.284341629 | 0.026450124 |
| AC010343.1 | 0.408395167 | 0.000108512 |
| EGFL8 | 1.966143621 | 0.005394984 |
| AC004801.2 | -2.170669724 | 2.71E-05 |
| SAMMSON | 1.928976797 | 4.09E-05 |
| LRRFIP1P1 | -0.911010585 | 0.000135134 |
| LINC00973 | 0.795739195 | 0.001019917 |
| AC022494.1 | -0.695626702 | 0.021089199 |
| RPL7AP10 | 1.951314308 | 0.002325102 |
| AL365273.1 | -1.146575493 | 0.001470034 |
| RPL3P9 | 0.560077031 | 0.008956541 |
| LINC02067 | 1.489978008 | 6.42E-05 |
| AQP1 | 1.20258149 | 0.000743748 |
| AL096701.3 | 1.416731159 | 0.003064978 |
| RPS6P25 | 0.393137169 | 7.72E-05 |
| AP001024.1 | 0.56131838 | 0.000515693 |
| AC106872.2 | 1.124894863 | 9.99E-05 |
| ISY1 | -0.566249958 | 7.20E-06 |
| PNMA2 | 0.641164102 | 0.002084013 |
| AC117382.1 | -0.363797893 | 0.0071409 |
| AL512306.3 | 4.374140863 | 0.001045189 |
| LRRD1 | 0.702229084 | 1.32E-07 |
| AC010655.2 | 1.416748666 | 0.000108855 |
| RPS27P21 | 0.242370186 | 0.019248068 |
| AC026877.1 | 0.252097292 | 0.015571838 |
| RPL9P25 | 1.968765963 | 9.27E-05 |
| TMEM189 | -0.181757635 | 0.006963275 |
| RDH14 | 0.247350666 | 0.002675147 |
| AC063952.2 | 2.217338865 | 0.025818147 |
| AC020633.1 | -2.642757804 | 1.29E-05 |
| RPL15P2 | 0.501340943 | 0.02556936 |
| AC021074.1 | 0.354376581 | 0.00030125 |
| RPL23AP64 | -1.693099851 | 0.001682698 |
| MIF | 0.422678061 | 0.000193125 |

| | | |
|---|---|---|
| AC021723.1 | 1.180257661 | 0.000100052 |
| AL157400.4 | 2.746725269 | 0.000186511 |
| AC114490.1 | 1.418258296 | 0.000285273 |
| TPM3P9 | 0.897474843 | 4.16E-07 |
| AP000873.1 | -0.620481202 | 0.017753288 |
| AC004951.3 | 1.510199659 | 0.009229746 |
| AC093019.2 | -2.185347196 | 0.000374518 |
| CYP51A1P1 | 1.143132188 | 5.35E-06 |
| CEACAMP10 | 3.884060809 | 0.010732639 |
| YAE1 | 0.529951526 | 3.55E-05 |
| AC115220.1 | 1.444666437 | 0.011751605 |
| RN7SL809P | -1.33495361 | 0.020841318 |
| AC063944.2 | 4.314632398 | 1.55E-05 |
| AL136126.1 | 0.427601305 | 0.000255877 |
| CRCP | -0.897585154 | 2.62E-08 |
| ENPP7P4 | 2.459561522 | 0.000553322 |
| AC106712.2 | 0.844016348 | 0.001864505 |
| LINC02614 | 2.103077397 | 1.01E-06 |
| PPATP1 | 0.323441623 | 0.005559264 |
| WWTR1-AS1 | 0.210766419 | 0.019354982 |
| CDRT1 | 2.75443107 | 0.000185159 |
| AL603962.1 | 1.856207573 | 0.004082769 |
| RPL36A | -0.28321171 | 0.000783088 |
| PMS2P11 | 0.790452216 | 0.02321302 |
| AC007688.2 | 0.271223568 | 0.011797052 |
| PDXP | -0.171685634 | 0.019093468 |
| HLA-DMA | 0.472064618 | 0.002210923 |
| AC104619.1 | 1.342156051 | 0.002909593 |
| SSX2 | -0.47117387 | 0.047893726 |
| HSPA8P9 | -0.274763658 | 0.026904048 |
| AL355032.1 | 0.300112561 | 0.00302107 |
| PSMC1P1 | 0.178203262 | 0.001020389 |
| AGGF1P3 | 4.012873544 | 1.07E-05 |
| CBX5P1 | 3.570975441 | 0.005294299 |
| ACTG1P20 | 0.330510876 | 0.047369125 |
| ARPC4 | -0.189620239 | 0.002808402 |
| AC018475.1 | 0.439948645 | 0.000742622 |
| CORT | 1.360133207 | 0.00092801 |
| PAQR9-AS1 | 2.310448876 | 7.52E-05 |
| AC245096.1 | 2.496742656 | 0.001363227 |
| AC114728.1 | 0.422292017 | 0.000419406 |
| AC069499.2 | -1.28791765 | 0.004716122 |
| INMT | 3.042387111 | 0.00107921 |
| RPL31P49 | 0.516057264 | 0.035058674 |
| ARPC1A | -0.177605994 | 0.009822772 |
| AP001062.3 | -0.90975843 | 0.00608648 |
| FABP5P3 | 4.373072962 | 0.001054824 |
| RPL7AP30 | 0.360350377 | 1.37E-05 |

| | | |
|---|---|---|
| RPSAP52 | -1.1600077 | 0.000699398 |
| AC002467.1 | 0.912016505 | 0.000472032 |
| C8orf58 | -0.381407749 | 0.001170454 |
| AL627309.5 | 1.058113897 | 0.004585025 |
| AC079944.2 | -0.329135054 | 0.005769639 |
| HOGA1 | 2.924423855 | 0.000183772 |
| PWP2 | -0.466616842 | 3.87E-05 |
| AC079447.1 | -0.460128761 | 0.005443673 |
| PI4KA | 0.197154929 | 0.009140377 |
| HYPK | 0.463536607 | 3.89E-06 |
| RPL9P28 | 1.252391862 | 0.005708935 |
| RPL7AP6 | 0.368933466 | 6.00E-07 |
| RPL7AP31 | 0.479643616 | 0.001255246 |
| MUC20-OT1 | -0.80356999 | 0.000379471 |
| RPL9P32 | 0.446591556 | 0.023824407 |
| MTFP1 | -0.905226203 | 4.05E-05 |
| SNHG3 | -0.439681258 | 4.51E-06 |
| SERBP1P3 | -0.707759332 | 0.040822255 |
| LINC02657 | 0.670876233 | 0.024759735 |
| ABCF2P1 | -0.392195847 | 0.043800811 |
| AC093789.1 | 1.370489952 | 0.000235593 |
| ARHGDIG | 1.362940182 | 0.004153969 |
| NECTIN3-AS1 | 5.542889111 | 7.47E-05 |
| ARFGAP3 | -0.211002517 | 0.049418918 |
| BGLAP | 2.057485788 | 7.41E-06 |
| LINC00996 | 5.23740422 | 5.38E-05 |
| AC018635.1 | -2.725730559 | 7.60E-07 |
| AC092597.1 | 0.232972364 | 0.031711773 |
| RPL5P3 | 0.491318414 | 0.049479707 |
| AP000893.1 | 7.340494991 | 5.69E-06 |
| AC108488.1 | 0.730466096 | 0.020332079 |
| AC022400.3 | 1.251319259 | 1.43E-06 |
| STAG3L5P | -0.309228135 | 0.021517598 |
| AC073861.1 | 0.364021133 | 1.22E-06 |
| RPL12P32 | 0.714067984 | 0.000769378 |
| AC023906.1 | 0.674974644 | 0.017429235 |
| INHCAP | 5.128592756 | 0.000834622 |
| BMS1P4 | 1.714219858 | 9.66E-07 |
| FAM91A3P | -0.534739396 | 0.005857298 |
| RPL12P30 | 7.612008947 | 2.21E-06 |
| RPL7AP11 | 0.443061246 | 2.69E-05 |
| AC091429.1 | 0.657323285 | 5.24E-06 |
| AC109992.1 | 0.757415002 | 0.002686769 |
| MRPL20 | -0.408953204 | 1.61E-05 |
| LINC00960 | -0.536792818 | 0.000102716 |
| MAGEA5 | 2.472242801 | 0.004048488 |
| AC007620.2 | -1.362751568 | 3.01E-08 |
| SERPINB10 | 3.625651194 | 6.42E-05 |

| | | |
|---|---|---|
| RPL21P11 | 0.279481138 | 0.005288044 |
| AL645608.5 | 1.267019715 | 0.007079137 |
| MBL1P | -1.360319258 | 0.00052775 |
| DECR2 | 0.499820802 | 0.001984464 |
| GNG10 | -0.171230952 | 0.025561671 |
| CNTF | -0.797991263 | 0.000383788 |
| RPS27AP1 | 0.406035299 | 0.036516899 |
| RTL5 | 2.068176116 | 3.47E-08 |
| RPSAP26 | 0.250470111 | 0.000886184 |
| RPL23AP81 | 1.730310502 | 0.000686077 |
| ZNF702P | 0.612995748 | 2.39E-05 |
| AC117395.1 | 1.856794287 | 0.004038846 |
| GLYCTK-AS1 | -1.403370543 | 0.035790782 |
| AP5Z1 | -0.217417474 | 0.021081613 |
| AC113398.1 | 0.326956762 | 0.002266061 |
| AC068756.1 | 1.299526164 | 0.009766411 |
| ZNF709 | 1.878813931 | 4.41E-09 |
| STRC | 5.750165725 | 0.000115072 |
| ERVW-1 | 4.436388445 | 4.15E-08 |
| AC007182.2 | 0.255892227 | 0.007316118 |
| RPSAP3 | 0.930765738 | 7.77E-05 |
| AC096992.1 | 7.046708888 | 1.57E-05 |
| AC005062.1 | 0.242721776 | 0.001354354 |
| RPL12P35 | 1.418154682 | 0.00052248 |
| AC012158.1 | 3.609552805 | 8.65E-06 |
| ARHGEF26-AS1 | 3.178697569 | 0.000589724 |
| AC107032.1 | 0.280378649 | 0.004745801 |
| MRPL33 | -0.376503676 | 0.000304537 |
| MICAL3 | -1.144575195 | 4.32E-10 |
| AC093270.3 | 1.466751773 | 0.023919126 |
| AC092944.1 | 5.360569367 | 2.80E-08 |
| AC087343.1 | 0.400846835 | 0.000171398 |
| AC115223.1 | 0.31869252 | 1.59E-05 |
| PPAN-P2RY11 | -1.10435478 | 2.62E-09 |
| AC006252.1 | 2.562382713 | 3.24E-06 |
| PCDHAC2 | -1.403117719 | 3.08E-09 |
| STON1 | 1.108753293 | 0.000528002 |
| RPL30P14 | 0.672450116 | 0.010172642 |
| PRAF2 | 0.337538377 | 0.000212143 |
| VSIG8 | 7.909682001 | 7.83E-07 |
| RPL31P61 | 1.082493074 | 0.011785977 |
| AC008494.1 | 1.576828647 | 0.01109182 |
| STMP1 | -0.150749786 | 0.033445509 |
| KCTD7 | -0.181884337 | 0.00697511 |
| EFNA4 | 1.593854913 | 8.13E-09 |
| AC090543.3 | -0.54785914 | 0.007254849 |
| TICAM2 | 0.305990866 | 0.00234328 |
| RPL23AP49 | 3.58755091 | 8.58E-08 |

| | | |
|---|---|---|
| RPL5P30 | 1.285429282 | 0.000430692 |
| AC106820.1 | 0.650057734 | 0.008390668 |
| NAA80 | 1.134782787 | 1.16E-06 |
| TNFRSF6B | -0.874310367 | 0.022034773 |
| AC024940.3 | 0.676880814 | 0.000224696 |
| HNRNPKP4 | 0.261631373 | 0.00010003 |
| AC004967.1 | 0.820544056 | 0.003422137 |
| LINC00880 | 1.693919372 | 0.017819239 |
| IL10RB | 1.220849795 | 1.68E-06 |
| ZNF487 | 1.467240695 | 0.00038478 |
| RPS4XP14 | 0.733449781 | 0.002722504 |
| WDR92 | -0.514861038 | 0.004476729 |
| NME2 | 0.320536023 | 9.60E-06 |
| AC018638.5 | -0.614292866 | 0.012556347 |
| AC006254.1 | 2.212423131 | 1.71E-07 |
| AC009108.1 | 2.295818814 | 0.004531344 |
| DUBR | -0.583753579 | 0.001488501 |
| LEFTY1 | 7.318773652 | 3.43E-06 |
| CFAP57 | 0.669373933 | 0.017392673 |
| NPIPB5 | -0.118396054 | 0.041541371 |
| RPLP0P2 | 0.723185582 | 0.011879279 |
| EEF1A1P10 | 1.678858084 | 0.000122737 |
| TMEM35B | 0.490828988 | 0.008772986 |
| ST13P15 | 0.27653919 | 0.034570319 |
| AC006547.2 | -0.846014582 | 0.001266942 |
| AP000942.1 | 0.509992362 | 0.016228048 |
| RN7SL832P | 2.126281279 | 1.92E-05 |
| RPL12P6 | 0.553653826 | 0.000125364 |
| AC011495.1 | 0.482827035 | 1.77E-05 |
| TIPARP-AS1 | 1.84879826 | 0.001480148 |
| MRPS6 | -0.508345282 | 0.001428581 |
| ZNF512 | -0.802727523 | 1.81E-05 |
| AL390195.2 | -0.809119109 | 2.77E-07 |
| RPL23AP65 | 0.267445806 | 0.000238894 |
| RTL9 | -0.979025677 | 0.007438947 |
| ACY1 | 0.633301074 | 1.67E-06 |
| NFS1 | -0.736935949 | 2.50E-07 |
| AC093591.1 | 0.239023709 | 0.02729509 |
| FAM86DP | 0.255508826 | 0.003393208 |
| DDOST | 0.273950532 | 1.61E-05 |
| LINC01011 | 1.356100907 | 0.003028885 |
| TMEM199 | -1.418488317 | 3.16E-09 |
| AC007566.1 | 2.446633564 | 3.58E-06 |
| AC093278.1 | 4.853483043 | 8.72E-07 |
| RPS4XP6 | 0.425093332 | 4.69E-05 |
| AC110749.1 | 0.343361676 | 0.042046849 |
| DNAJC25-GNG10 | -0.150862046 | 0.043617943 |
| PDCL3P4 | 1.192776545 | 2.57E-07 |

| | | |
|---|---|---|
| ATP1B3-AS1 | 1.434055057 | 0.000427486 |
| AC010973.2 | 0.238359001 | 0.032988699 |
| WWP1P1 | -0.86198637 | 0.000360464 |
| P2RY11 | -2.260425416 | 4.65E-11 |
| PBX2P1 | 0.496007569 | 0.008096449 |
| AC091153.3 | 2.691440962 | 2.34E-05 |
| TMEM141 | 0.980967432 | 3.05E-06 |
| ZFAND6P1 | 0.642718863 | 0.014593447 |
| TMEM225B | 3.074551726 | 0.000421873 |
| GMCL2 | -0.534651602 | 0.025876178 |
| AC133134.1 | 0.347261837 | 0.019705799 |
| PKD1P1 | -0.236098099 | 0.006212535 |
| RPL32P29 | 3.156612246 | 0.000126157 |
| DBNDD2 | 0.767052689 | 0.000378505 |
| GATA2-AS1 | 0.472038772 | 0.031831984 |
| AL589743.1 | -0.38501976 | 4.78E-06 |
| AC024293.1 | 0.317320543 | 2.96E-05 |
| AL138759.1 | -1.169290919 | 0.013901023 |
| RPL7P23 | 0.32587811 | 9.26E-05 |
| AC116533.1 | 0.404160962 | 9.38E-08 |
| ENO1P1 | 0.13160155 | 0.021174144 |
| RBM12 | -2.022897522 | 1.36E-12 |
| AC005154.2 | 1.11473137 | 0.037522352 |
| LILRA6 | 2.392458813 | 0.001270303 |
| KRT18P34 | 3.406468064 | 1.30E-07 |
| AC098583.1 | 0.41798293 | 0.000863112 |
| ODCP | 2.425812334 | 0.001041398 |
| CCDC13 | 4.180016306 | 9.16E-07 |
| MIATNB | -1.15884828 | 0.004306135 |
| TPTEP2 | 1.09078158 | 0.000151944 |
| LINC02086 | 3.094446552 | 0.019350553 |
| UBE2V1 | -0.192956391 | 0.000628799 |
| PTCHD4 | 0.931934418 | 0.000116224 |
| AC020658.1 | 0.845661243 | 0.049846186 |
| BX679664.3 | 0.290662965 | 5.31E-05 |
| AL132656.2 | -0.637504087 | 0.02017648 |
| N4BP2L2 | -1.073904077 | 1.22E-09 |
| GABPB1-AS1 | 2.834574214 | 1.36E-12 |
| AC136604.2 | 4.208871398 | 2.16E-08 |
| LIFR-AS1 | 1.752433542 | 5.28E-05 |
| AC107959.1 | -0.566428895 | 0.000525214 |
| LINC00847 | 1.487149256 | 1.50E-07 |
| SMARCA5-AS1 | -0.284950067 | 0.047833263 |
| MALINC1 | 1.795994268 | 3.92E-07 |
| RNF139-AS1 | 1.835698787 | 5.21E-07 |
| LINC00861 | 7.693404731 | 1.74E-06 |
| EEF1A1P4 | 0.629086764 | 1.77E-05 |
| AC124242.1 | 2.707601013 | 0.000283192 |

| | | |
|---|---|---|
| AC008393.1 | 2.89431689 | 7.86E-08 |
| AP003396.1 | -0.504519111 | 0.014245831 |
| LINC02447 | 4.30975812 | 6.06E-08 |
| AP000866.1 | 3.276956445 | 2.21E-08 |
| LINC02709 | 1.496179614 | 7.24E-05 |
| LINC00461 | -1.048489622 | 4.53E-05 |
| NEAT1 | 2.031418265 | 1.67E-13 |
| RORA-AS1 | 2.076991576 | 0.0006542 |
| AP000787.1 | 1.260271527 | 0.003351039 |
| SCAMP1-AS1 | 0.468559217 | 0.02250981 |
| BDNF-AS | 2.266487186 | 6.57E-06 |
| AC022075.1 | 1.270979448 | 0.006599348 |
| ZNF585B | 0.798205055 | 8.35E-06 |
| NADK2-AS1 | 2.339074761 | 3.65E-05 |
| AC097382.2 | -1.096198305 | 1.80E-09 |
| LINC02202 | 6.961651229 | 1.88E-05 |
| CEBPA | 1.147767284 | 0.000547573 |
| RAD51-AS1 | 1.583869823 | 1.87E-05 |
| NSMCE1-DT | 2.52627152 | 0.001364092 |
| AC025164.2 | 1.921746667 | 8.57E-09 |
| SNHG6 | 0.451079199 | 2.67E-05 |
| LINC01184 | 0.293389863 | 0.011787097 |
| AC093752.1 | 1.400969748 | 8.72E-07 |
| AC090515.3 | 2.847845959 | 4.77E-05 |
| RAB30-DT | 1.136284244 | 1.01E-05 |
| NUDT16P1 | 4.554314221 | 0.00062713 |
| AC016065.1 | 1.364483531 | 0.0001037 |
| AP002026.1 | 3.151236421 | 2.89E-07 |
| AC107959.2 | 3.256444596 | 0.00052076 |
| RRS1-AS1 | -1.548439897 | 0.008516989 |
| KCTD21-AS1 | 1.809407965 | 1.06E-08 |
| UBR5-AS1 | 0.451995153 | 0.020166939 |
| SBF2-AS1 | -0.904319827 | 0.00070077 |
| AC116535.2 | 3.612340813 | 4.34E-05 |
| AC113382.1 | 1.488643968 | 0.024304736 |
| PRR7-AS1 | -0.639430829 | 0.000552638 |
| EXTL3-AS1 | 3.55005895 | 2.36E-08 |
| AL049543.1 | 8.648484178 | 5.48E-08 |
| LACTB2-AS1 | 1.00029549 | 0.000983669 |
| LINC002481 | 4.715191416 | 0.000301696 |
| LINC02288 | 3.554856465 | 0.000309026 |
| UBE2D3-AS1 | 2.048240295 | 1.45E-05 |
| AC093162.3 | 1.445742677 | 0.014889853 |
| CACNA1C-AS1 | 7.976247657 | 6.20E-07 |
| PICART1 | 4.263286907 | 1.47E-08 |
| RASSF8-AS1 | 1.014557688 | 5.52E-06 |
| H2AJ | 1.702204595 | 7.73E-07 |
| RGMB-AS1 | 1.773779987 | 2.66E-05 |

| | | |
|---|---|---|
| AL157938.3 | 1.826326805 | 3.84E-05 |
| GPR176-DT | 3.196403813 | 1.51E-06 |
| DNM1P35 | 3.00816642 | 0.000621217 |
| UBAP1L | 2.253201537 | 1.27E-07 |
| Z84485.1 | 3.009838735 | 0.000591439 |
| PGAM5 | -0.547584583 | 4.82E-07 |
| AC009126.1 | 1.430598464 | 0.000283159 |
| AC090204.1 | 0.452340522 | 0.030778981 |
| AP000873.2 | 0.347336582 | 0.036086042 |
| AC011346.1 | 4.872410541 | 4.69E-07 |
| ZBED5-AS1 | 2.156287619 | 1.33E-06 |
| ZCCHC3 | -0.640919937 | 2.75E-07 |
| LY6E-DT | 1.42095555 | 0.006453768 |
| AC010547.1 | 2.482458718 | 0.002719736 |
| AC090061.1 | 1.341474049 | 0.017200551 |
| TMED2-DT | 0.332473166 | 0.00155789 |
| DNAJC3-DT | 1.893401944 | 1.25E-07 |
| GPRC5D-AS1 | 4.771064323 | 0.013850133 |
| OIP5-AS1 | 0.47179363 | 1.75E-06 |
| CKMT2-AS1 | 0.355188397 | 0.02568076 |
| SPTY2D1OS | 0.554989216 | 0.001705989 |
| TWF2 | 0.570263469 | 4.13E-07 |
| CPEB2-DT | 5.277991258 | 0.00010309 |
| MARS2 | -0.643310301 | 3.34E-06 |
| MTND4P12 | 0.723779642 | 5.02E-07 |
| LRP4-AS1 | 4.538426209 | 2.07E-11 |
| AC120114.1 | 1.76628492 | 6.29E-05 |
| AC068446.2 | 2.274282151 | 4.86E-07 |
| AC008966.1 | 0.95975907 | 0.000100428 |
| AC006064.2 | 0.331548889 | 0.000206281 |
| AC024896.1 | -0.337573959 | 0.015731977 |
| AC022364.1 | 0.920361081 | 0.002117627 |
| SEC24B-AS1 | 2.776955492 | 0.001629655 |
| FOXD1-AS1 | -1.655591721 | 1.50E-06 |
| AC005329.1 | 0.465774468 | 0.007139524 |
| UBA6-AS1 | -0.894926927 | 7.68E-06 |
| BCKDHA | 0.368268896 | 0.000901315 |
| AC067942.2 | -0.528166462 | 0.004731117 |
| SMURF2P1 | -1.133234279 | 0.010528472 |
| LINC01194 | 1.199606007 | 0.002199417 |
| LINC02150 | 8.337546556 | 2.59E-07 |
| AC098487.1 | 4.31354768 | 7.22E-09 |
| GAPDHP60 | 0.246753715 | 0.004571123 |
| AC026785.2 | 0.303045946 | 0.028600005 |
| AC106872.4 | 7.199016329 | 8.57E-06 |
| LINC02438 | 1.440279939 | 0.009999297 |
| AC023794.2 | 3.72688345 | 1.51E-07 |
| SUMO2P17 | 0.402771091 | 0.021631003 |

| | | |
|---|---|---|
| LUCAT1 | -0.844480104 | 0.006085525 |
| WHAMMP2 | -2.282998635 | 3.56E-05 |
| TNXA | 1.930986197 | 0.003005091 |
| LINC02434 | 3.332504615 | 1.31E-12 |
| AC104066.3 | 3.965386074 | 1.69E-11 |
| PCDHAC1 | -1.276139404 | 4.99E-09 |
| FOSL1P1 | -1.443140701 | 0.000176633 |
| LINC02119 | 1.907237319 | 1.24E-08 |
| ABHD14A | 0.259336774 | 0.042339897 |
| LINC02062 | 2.691044298 | 2.48E-05 |
| ASNSP1 | 3.170383304 | 0.043504434 |
| SRP14-AS1 | 2.660204058 | 2.72E-08 |
| MTATP6P1 | 0.307876401 | 1.24E-06 |
| AC024558.1 | 2.259692133 | 7.96E-06 |
| TECRP2 | 0.729475595 | 0.042879092 |
| AC026410.3 | -0.450982936 | 0.000755597 |
| NPM1P21 | 0.43855768 | 1.52E-05 |
| GLDCP1 | 0.777762681 | 0.003903463 |
| STIMATE-MUSTN1 | -1.061468174 | 1.16E-08 |
| AC139491.2 | 3.239703592 | 6.01E-09 |
| HMGA1P2 | 0.533936717 | 0.004961138 |
| RBM14-RBM4 | -0.961987641 | 5.31E-09 |
| AC106872.5 | 6.961651229 | 1.88E-05 |
| AC010273.2 | 0.467888024 | 0.043910797 |
| AP003498.1 | 2.408876168 | 2.35E-05 |
| LINC02102 | 1.411998892 | 0.018507865 |
| LINC02484 | 1.55222778 | 6.04E-06 |
| AC097534.1 | -1.026770837 | 0.004062695 |
| AC010627.1 | 5.282635579 | 0.000128214 |
| AC024587.1 | -0.863501201 | 0.001645541 |
| AC097512.1 | 0.402892378 | 0.04057285 |
| HNRNPH1P3 | 0.35699572 | 0.025175061 |
| USP46-AS1 | 0.741065613 | 0.003023726 |
| TNFSF12-TNFSF13 | 1.752488581 | 7.41E-06 |
| ICE2P1 | 2.690013736 | 6.97E-05 |
| AC022126.1 | 2.520734835 | 0.000583348 |
| FMN1 | -0.914382128 | 1.77E-07 |
| AC046134.2 | 3.074870309 | 5.06E-07 |
| AC131254.1 | 2.130927743 | 0.002249978 |
| BCLAF1P1 | -1.140295 | 3.28E-06 |
| KRT8P46 | 1.674681646 | 0.000518863 |
| AL136360.1 | 3.230857192 | 0.014974552 |
| AC008771.1 | -0.825333517 | 0.004480129 |
| ZNF436-AS1 | 2.162793299 | 1.34E-08 |
| HAUS5 | -0.186532471 | 0.018078692 |
| SUDS3P1 | -1.167924935 | 0.000555557 |
| PCDHA11 | -1.444222889 | 2.76E-09 |
| AC106744.1 | 5.403360346 | 7.29E-05 |

| | | |
|---|---|---|
| HSPD1P5 | -0.189343239 | 0.018217216 |
| TMEM132D-AS1 | 1.894985821 | 0.001887362 |
| ATP1B1P1 | 1.942129857 | 0.001430379 |
| CDH12P2 | -1.688575654 | 0.017638883 |
| ATP5MGP3 | 0.83582117 | 0.041073286 |
| EEF1A1P9 | 0.453807448 | 1.51E-06 |
| LINC00939 | 9.436317945 | 4.49E-09 |
| AMD1P3 | -0.760711448 | 0.00620424 |
| UGDH-AS1 | 3.431370521 | 0.000122129 |
| NPM1P27 | 0.43731035 | 1.17E-08 |
| LINC01060 | 0.452755163 | 0.011274686 |
| HMGB3P17 | 8.470068226 | 1.69E-06 |
| AC092834.1 | -2.972592032 | 6.87E-05 |
| RBMXP4 | 0.748213472 | 0.031335509 |
| ZNF324B | 1.016588073 | 2.15E-07 |
| RBBP4P1 | -0.254212026 | 0.01117253 |
| AC006160.1 | 3.715541551 | 4.66E-05 |
| LINC01234 | 0.862222076 | 1.18E-07 |
| SERBP1P5 | -0.14466411 | 0.012176027 |
| AC034231.1 | 1.500401346 | 0.000161661 |
| AC004832.3 | -0.61006407 | 0.001357838 |
| AP000295.1 | 0.781097662 | 0.002733991 |
| HOXC13-AS | 0.886547423 | 0.01715805 |
| NOP14-AS1 | 0.685379956 | 1.42E-06 |
| THEGL | 5.211413645 | 6.46E-05 |
| SRD5A3-AS1 | 1.421945098 | 0.003978618 |
| ZNF564 | 1.188026063 | 1.53E-08 |
| FER1L5 | 4.058460792 | 0.002146554 |
| AC093890.1 | -0.939505681 | 0.000200426 |
| AC083829.1 | 2.483308746 | 0.000790715 |
| AC026427.1 | 8.906170458 | 2.30E-08 |
| AC092647.5 | -0.499551133 | 0.000436055 |
| EAF1-AS1 | 0.366319903 | 0.000258498 |
| HSD3BP3 | 3.98980602 | 1.35E-05 |
| AC012636.1 | 3.520798553 | 0.005878378 |
| AC011330.1 | -0.493551277 | 0.005998029 |
| LINC02021 | 2.1315557 | 0.000135459 |
| AC138819.1 | 1.757952178 | 0.003237666 |
| EEF1A1P19 | 0.433017048 | 6.36E-06 |
| PVT1 | -0.329821891 | 0.044820269 |
| AC021106.1 | -1.153158921 | 0.002164023 |
| MCPH1-AS1 | 2.251315148 | 2.02E-05 |
| PDCD6 | -0.330702595 | 9.65E-05 |
| AC034207.1 | 0.4513795 | 0.021677962 |
| UQCRBP3 | 4.929207864 | 0.00021046 |
| AL157400.5 | -1.732135666 | 0.008754798 |
| AL355315.1 | 0.30588597 | 0.018229817 |
| AC104116.1 | 1.012009974 | 6.97E-05 |

| | | |
|---|---|---|
| TMEM158 | -0.45307372 | 0.013884298 |
| AC079848.1 | 3.246786124 | 0.025648977 |
| SNX18P23 | 2.735706201 | 0.000131409 |
| ARPIN-AP3S2 | -0.288508702 | 0.003547769 |
| SLC7A11-AS1 | -1.717762154 | 4.79E-05 |
| LINC01018 | 6.618143764 | 4.85E-05 |
| YJEFN3 | 0.609278778 | 4.52E-06 |
| SH3TC2-DT | -1.555330985 | 7.86E-06 |
| AP000866.2 | -2.631317519 | 0.00010077 |
| AC017013.1 | -0.543397307 | 0.038630387 |
| DNAH10OS | 2.305267644 | 2.13E-05 |
| AC107982.1 | 1.410499227 | 4.95E-05 |
| PCDHA10 | -1.178376979 | 8.90E-09 |
| AC078881.1 | 1.09222338 | 0.001621908 |
| AC004803.1 | 2.553100367 | 1.50E-07 |
| AC026801.1 | 6.961651229 | 1.88E-05 |
| MYLK-AS2 | 6.871264811 | 2.36E-05 |
| EEF1A1P13 | 0.443298358 | 2.82E-07 |
| AC091180.4 | 0.22616109 | 0.002031311 |
| LINC01618 | 6.469091211 | 1.18E-05 |
| LINC02762 | 2.844855306 | 0.002158886 |
| ZNF718 | 0.830309772 | 4.27E-05 |
| SMIM20 | 0.581636223 | 6.88E-06 |
| AC079140.2 | 0.617911752 | 1.06E-05 |
| AC104596.1 | 2.949858969 | 0.036649176 |
| PURPL | 3.335027063 | 6.28E-10 |
| AC020659.2 | -0.657508932 | 0.036222691 |
| AP006623.1 | -0.842159465 | 0.013584402 |
| AC112722.1 | 3.067964654 | 0.000723043 |
| AACSP1 | -1.052061333 | 0.000229908 |
| AC122718.1 | 0.550167381 | 0.000131664 |
| LRRC37BP1 | -1.286914797 | 6.74E-09 |
| GMPSP1 | -0.493686426 | 0.000151813 |
| CHCHD10 | 0.648945705 | 0.004648383 |
| FAM218A | 1.120812914 | 0.000641058 |
| LINC02145 | 6.77593355 | 3.11E-05 |
| INTS6P1 | -1.742826502 | 6.21E-06 |
| GPR162 | 1.767734604 | 0.000588174 |
| KRT8P33 | 2.391568313 | 1.23E-07 |
| LINC02059 | 7.437488347 | 4.26E-06 |
| ATP6V1E2 | 1.362189821 | 7.72E-07 |
| AC010406.1 | 7.122711146 | 1.12E-05 |
| GLI4 | 1.087559183 | 1.97E-07 |
| AL732372.3 | 1.476019701 | 0.000595138 |
| AC010210.1 | 6.152693361 | 1.72E-05 |
| AC008581.1 | -0.955179499 | 0.016624161 |
| AC012645.1 | 2.568908962 | 5.94E-05 |
| AL031073.1 | 4.644162743 | 0.001370035 |

| | | |
|---|---|---|
| AC068580.4 | 0.652012575 | 1.31E-05 |
| AC023794.5 | 9.346522375 | 1.08E-08 |
| ST3GAL1P1 | 2.719682734 | 0.000660993 |
| LINC00491 | -1.890632763 | 9.08E-05 |
| AC106872.8 | 6.665724289 | 2.22E-06 |
| SELENOP | 1.904157727 | 0.000183345 |
| AC016687.2 | 7.003868569 | 1.61E-05 |
| TPM3P6 | 1.58731629 | 2.95E-07 |
| LINC02381 | -0.577322045 | 0.00239022 |
| AC105250.1 | 0.736412851 | 1.95E-06 |
| AC127070.2 | 3.434138524 | 7.66E-06 |
| AC112484.2 | 2.389733681 | 0.000777647 |
| ZBED3-AS1 | 1.220657654 | 0.002069452 |
| AC021087.2 | -0.3328855 | 0.011904014 |
| HNRNPKP1 | 0.402465586 | 0.001607502 |
| AC106872.9 | 3.6493795 | 0.000196414 |
| RNPS1P1 | -0.313010219 | 0.000387017 |
| AC125807.2 | -1.609536586 | 2.63E-08 |
| AC008443.6 | 4.256237464 | 1.18E-05 |
| SMAD1-AS1 | -1.822646401 | 0.000267118 |
| GMDS-DT | 3.629543348 | 3.45E-07 |
| AC138956.1 | 1.344839607 | 0.021674467 |
| AC097467.3 | 0.75467849 | 0.005129725 |
| AL035458.2 | 1.825655735 | 2.73E-05 |
| AC008629.3 | 7.08299031 | 1.30E-05 |
| AC016924.1 | 4.37200433 | 0.001087522 |
| AC093752.2 | 1.385098549 | 0.007244222 |
| AC016687.3 | 2.603983328 | 0.000208946 |
| AC012055.1 | 0.791607246 | 0.005142325 |
| AC025470.2 | 1.251449691 | 3.92E-07 |
| GAPDHP35 | 0.422455014 | 0.006819126 |
| AC022296.3 | 1.738691954 | 0.002678681 |
| SNHG21 | 0.918816314 | 4.42E-05 |
| AC136604.3 | 1.418380729 | 0.020375385 |
| AC244502.1 | 7.00011059 | 1.83E-05 |
| ZFPM2-AS1 | -0.293354355 | 0.046571054 |
| CUL1P1 | -1.329240774 | 0.022755513 |
| AC087854.1 | -2.168334203 | 0.003911574 |
| AC112184.1 | 2.330864251 | 7.73E-06 |
| BMS1P2 | 0.862141841 | 0.00055566 |
| CTBP2P4 | -0.414652626 | 0.015230011 |
| YBX1P5 | 6.774837438 | 3.09E-05 |
| AF117829.1 | 1.190881988 | 5.51E-06 |
| MRPS30-DT | 1.4095488 | 1.83E-05 |
| AP002490.1 | -0.706048336 | 1.80E-07 |
| HOXC-AS3 | 1.853819492 | 0.00085211 |
| AC020661.1 | 0.901489093 | 0.025568453 |
| AC010261.2 | 7.201478166 | 9.56E-06 |

| | | |
|---|---|---|
| ZNF674 | 0.902490099 | 1.39E-05 |
| AC015631.1 | 1.372875712 | 0.00013298 |
| LINC00923 | 1.292655782 | 6.95E-05 |
| LINC01337 | 2.593493005 | 0.003218113 |
| AL691442.2 | 0.984673256 | 0.001162941 |
| AC004069.1 | 0.95548575 | 0.031220466 |
| LINC02228 | 0.959668033 | 0.039798818 |
| SMIM15-AS1 | -0.520294743 | 0.013358437 |
| ALG1L2 | 1.832540571 | 0.000186486 |
| LINC02384 | 8.213032461 | 2.64E-07 |
| AC107391.1 | 6.963581473 | 2.29E-05 |
| RTN3P1 | -0.251242505 | 0.006392464 |
| IRF5P1 | 7.040315991 | 1.94E-05 |
| AP000350.4 | 0.486091425 | 3.27E-05 |
| WWC2-AS2 | 1.857553243 | 0.002388637 |
| ZNF550 | 0.396101245 | 2.57E-05 |
| LINC01301 | 1.182899745 | 0.002733991 |
| AC145285.2 | 3.419248407 | 2.48E-05 |
| FKBP4P1 | 1.61847674 | 0.014020387 |
| AC068134.3 | -1.16963076 | 0.003786044 |
| AC106744.2 | 5.178549865 | 6.21E-08 |
| FOXD1 | -1.759418572 | 1.46E-08 |
| PPIAP11 | 0.267064283 | 0.005771803 |
| CENPS-CORT | 0.618078075 | 0.000786032 |
| AC005324.3 | 0.380684801 | 0.028901284 |
| SNX18P24 | 1.803272874 | 0.005753675 |
| AC091887.1 | 1.85729933 | 0.00210662 |
| MALAT1 | -0.432089762 | 2.07E-05 |
| DDX3P3 | -0.781441615 | 0.00248194 |
| AC099520.1 | 1.9147259 | 0.00019445 |
| AC008427.2 | -0.688436925 | 0.005428697 |
| ABCA11P | 0.529646426 | 0.001728879 |
| HADHAP1 | 0.469048061 | 0.024403381 |
| AC104825.1 | 3.893021549 | 8.55E-09 |
| NAIPP4 | -0.610256919 | 0.021150209 |
| PCDHA12 | -1.454365827 | 3.14E-09 |
| FAM86EP | 2.730976345 | 8.65E-10 |
| AC007684.1 | 3.798685693 | 0.00642187 |
| RNA5SP268 | 3.817264453 | 9.82E-05 |
| AC004707.2 | -0.278127151 | 0.045091372 |
| AC055854.1 | 1.491949206 | 0.027068322 |
| AC009630.2 | -0.551860261 | 0.001044135 |
| AC104986.1 | 1.359548983 | 0.016787368 |
| HOXA10-AS | -1.6901529 | 6.49E-06 |
| AC037459.2 | -0.251596665 | 0.002074021 |
| AC110998.1 | -1.231587376 | 0.00016781 |
| SHLD3 | -0.75360066 | 5.40E-05 |
| CCDC71L | 1.034902419 | 7.18E-09 |

| | | |
|---|---|---|
| HOXA10 | -1.901296189 | 2.52E-09 |
| AC022217.3 | 2.522252755 | 0.001195002 |
| LINC01606 | 2.328397297 | 0.001834269 |
| TMEM200B | -0.901662798 | 2.75E-05 |
| SERPINE3 | -4.40788255 | 3.89E-06 |
| C1orf210 | 8.060024042 | 4.72E-07 |
| AZIN1-AS1 | 4.29943932 | 1.62E-07 |
| AL606534.4 | 2.330113446 | 0.010053281 |
| AC115837.1 | 0.173626842 | 0.003596631 |
| TUG1 | 0.365395153 | 2.07E-06 |
| AC084024.3 | -0.604457135 | 0.000647691 |
| GUSBP16 | -1.141948648 | 0.0270481 |
| TRNP1 | -0.177404936 | 0.016055754 |
| AC124242.2 | 1.302038787 | 0.03000181 |
| AC104561.1 | 1.881857788 | 0.002994183 |
| AC083843.1 | 8.016637511 | 5.84E-07 |
| PCAT1 | 2.398616765 | 0.002788374 |
| AC091185.1 | -1.077198535 | 0.041125697 |
| AF121898.1 | 1.625199925 | 9.82E-05 |
| NACA2 | 0.406597295 | 3.31E-05 |
| FAM86HP | 2.488908535 | 4.99E-07 |
| SEPTIN10P1 | 0.560255532 | 0.003678874 |
| HOXA-AS2 | -0.944680264 | 0.002114961 |
| RNF5P1 | 0.528900205 | 0.006002057 |
| AFG3L2P1 | -0.475009658 | 0.038748639 |
| AC107959.3 | 2.059979638 | 0.02165788 |
| AC084026.2 | 4.103860645 | 3.14E-05 |
| AP002852.1 | 2.302262702 | 0.002814576 |
| AC022893.1 | 1.085454119 | 0.004235006 |
| AF279873.3 | -0.919554643 | 0.00514674 |
| AC023632.2 | 3.030581866 | 1.93E-06 |
| ALG11 | 0.59416426 | 3.68E-06 |
| MINCR | 1.855158641 | 3.33E-07 |
| ATXN7L3B | -1.423764465 | 1.08E-10 |
| AC010834.1 | 0.363634481 | 0.009651057 |
| AC015909.2 | 1.825079251 | 0.00981723 |
| OTUD6B-AS1 | -0.648271733 | 1.81E-06 |
| LNCOC1 | -0.435118623 | 6.58E-05 |
| AC025442.1 | -1.246659303 | 0.005679818 |
| UTP14C | -0.493491192 | 4.71E-06 |
| PSAT1P1 | -1.054951583 | 0.022815313 |
| AC022868.1 | 0.453497498 | 0.044005247 |
| AC010834.3 | 3.078606314 | 3.81E-08 |
| SLC2A3P1 | -0.972261548 | 3.50E-08 |
| AC131025.1 | 7.497145996 | 3.27E-06 |
| LINC01608 | 4.18114522 | 0.000149466 |
| AC087752.3 | 3.325182353 | 3.02E-05 |
| LINC01419 | 0.896954681 | 7.30E-09 |

| | | |
|---|---|---|
| AC067904.2 | -0.998265872 | 0.043313547 |
| AC104115.2 | 2.120262695 | 0.001562529 |
| MAPRE1P1 | -0.696858188 | 0.01590598 |
| THAP12P7 | -0.312553724 | 0.001979137 |
| VPS13B-DT | 2.049197928 | 0.00018387 |
| AC008663.2 | 4.594026732 | 0.000677113 |
| ALG1L13P | 1.383250329 | 0.003886775 |
| AC100810.1 | 1.138872723 | 0.039547836 |
| ZNF260 | -0.808508138 | 1.18E-07 |
| RPL10P8 | 0.779454998 | 0.018704303 |
| MAP2K1P1 | -0.628190419 | 0.004144113 |
| BRIX1P1 | -0.975617382 | 0.003662054 |
| AC083843.2 | 7.815325753 | 1.27E-06 |
| AC105206.2 | 1.262970177 | 0.001340686 |
| AC024451.3 | -0.660381533 | 0.029825999 |
| LYN | -1.247973095 | 5.30E-08 |
| SLC2A3P4 | 2.281974579 | 1.50E-05 |
| PINX1 | -0.484918958 | 0.004739982 |
| AC108449.1 | 1.219575478 | 0.030760212 |
| CRYZL2P-SEC16B | 2.444902824 | 8.38E-05 |
| MAPK6P5 | -0.60262901 | 0.026643782 |
| AC113191.1 | -0.896581022 | 0.001664074 |
| PRXL2AP2 | 6.399768465 | 1.22E-05 |
| LINC01933 | 4.225321339 | 0.001623669 |
| PKIA-AS1 | 1.327550543 | 0.026559053 |
| KRT8P3 | 1.286092917 | 7.22E-05 |
| AP003692.1 | -0.614284499 | 0.014316041 |
| CKS1BP7 | 0.386907251 | 0.002163951 |
| HOXA-AS3 | -1.500585812 | 0.000158958 |
| AL355432.1 | 3.228699546 | 0.000497093 |
| LRRC24 | -0.518937641 | 0.01241105 |
| MSH5-SAPCD1 | 0.842536144 | 0.007292612 |
| CHKB-CPT1B | -0.444544887 | 0.002665201 |
| LINC02584 | -1.021866318 | 0.030620462 |
| AP000942.2 | -0.645567394 | 0.000162115 |
| AP003392.1 | -1.237567308 | 0.048458407 |
| RCC2P6 | -0.408547817 | 0.044122109 |
| AP002765.1 | 7.556363372 | 2.73E-06 |
| AP5B1 | -1.248677272 | 1.35E-08 |
| AL354920.1 | 3.648152528 | 2.25E-08 |
| AC026191.1 | 1.886811091 | 8.32E-07 |
| CHMP4A | 0.434159063 | 0.000751618 |
| PHBP2 | 1.467903286 | 0.005543695 |
| LINC02750 | 1.43995101 | 0.000749626 |
| AP001816.1 | 2.753772976 | 1.24E-06 |
| AC239804.1 | 3.554723617 | 1.13E-06 |
| AC036111.1 | 2.130827055 | 0.000446391 |
| AP000873.3 | 2.733355978 | 1.37E-06 |

| | | |
|---|---|---|
| AC069287.2 | -0.556257239 | 0.024901004 |
| OR7E126P | -1.705712497 | 0.03001381 |
| AC084337.1 | -1.191911225 | 0.014334824 |
| CSNK2A3 | -0.900974167 | 2.37E-09 |
| AP000662.1 | 3.293734998 | 7.59E-05 |
| AP002336.2 | 2.142655408 | 0.000850714 |
| AP001783.1 | 3.897360224 | 7.52E-05 |
| AP001000.1 | -1.060163657 | 2.39E-09 |
| AC027031.2 | -0.420917328 | 0.002009052 |
| WAC-AS1 | 0.494666332 | 3.42E-05 |
| RTL1 | 1.068607915 | 0.000789159 |
| AC091053.1 | -1.529135896 | 0.000485411 |
| AP002387.2 | 2.071784998 | 2.17E-05 |
| FPGT | -0.909967348 | 1.18E-07 |
| AP001970.1 | 5.052897688 | 0.000210167 |
| GLYATL1P2 | -1.649064427 | 0.000231368 |
| AL157756.1 | 4.423405156 | 9.16E-06 |
| AP000879.1 | -0.763225796 | 1.17E-06 |
| MEX3A | -1.176764473 | 1.03E-08 |
| AP001931.1 | -0.788430281 | 0.002398727 |
| NAP1L1P1 | 0.246715273 | 0.003156055 |
| EEF1G | 0.47085184 | 9.58E-08 |
| AL662828.1 | 0.28008397 | 0.036820406 |
| EGLN1P1 | -1.042583676 | 0.021186634 |
| SYS1-DBNDD2 | 0.59822672 | 0.001116935 |
| AP003032.2 | 8.871613028 | 2.75E-08 |
| RNF185-AS1 | -1.291578927 | 0.000669391 |
| AP001372.2 | 1.074855254 | 2.75E-05 |
| AC024405.2 | 2.099106829 | 2.15E-06 |
| LINC02551 | 2.230462423 | 2.00E-05 |
| AP005018.2 | -0.811788233 | 0.000685583 |
| NPIPA2 | 0.448010502 | 0.02445131 |
| MPV17L2 | 0.370673883 | 0.000143756 |
| TMEM9B-AS1 | 1.838348199 | 0.002611723 |
| AP006287.1 | 2.847606543 | 0.001413783 |
| ATP6V1G2-DDX39B | 0.369794196 | 0.001850819 |
| AP001267.1 | -0.464498756 | 5.53E-06 |
| SUGT1P4-STRA6LP | -1.182857863 | 0.001134842 |
| BX248516.1 | 0.378676018 | 0.001932389 |
| BORCS8 | -0.376706374 | 0.014836489 |
| AP000866.4 | 4.139639862 | 5.02E-05 |
| ALG1L9P | 1.428131777 | 0.003158982 |
| AP000781.2 | -0.507044937 | 4.31E-06 |
| DPP3 | 0.260897446 | 0.000904026 |
| STX16-NPEPL1 | 0.289476901 | 0.004700539 |
| ANKHD1-EIF4EBP3 | -0.952712309 | 9.60E-09 |
| AP001767.2 | 7.85035855 | 0.003705094 |
| ARL6IP1P3 | -0.990777671 | 0.012219174 |

| | | |
|---|---|---|
| STRA6LP | 2.190763683 | 2.36E-05 |
| AC109635.3 | -0.510812357 | 0.045825417 |
| AC069185.1 | 3.183975932 | 4.75E-05 |
| AL020996.2 | 0.471950398 | 0.027746491 |
| AP001646.2 | 0.55038698 | 0.047400615 |
| ZFP91-CNTF | -1.124288871 | 1.22E-08 |
| AF186192.2 | 6.962616657 | 2.04E-05 |
| AC005324.4 | -0.364022421 | 0.001922744 |
| CHMP1B | 1.636680406 | 3.68E-11 |
| AP003392.3 | 0.238524084 | 0.028989233 |
| CCDC84-DT | 1.023944737 | 2.03E-05 |
| AP002360.2 | 1.583549776 | 2.83E-06 |
| TOLLIP-AS1 | 2.508960489 | 0.000722646 |
| HTD2 | -0.479084361 | 0.00014308 |
| AC026894.3 | 7.043059096 | 1.43E-05 |
| MUC5B-AS1 | -1.892538359 | 0.003605067 |
| PGAM1P8 | 0.775414085 | 0.006919177 |
| NPM1P35 | 0.470515377 | 0.036077856 |
| MIR100HG | -1.114260591 | 1.56E-09 |
| AP006621.3 | 1.16971691 | 0.000431305 |
| OR8G5 | 2.130954647 | 0.003685657 |
| AP003557.1 | 5.062109082 | 9.75E-05 |
| EID1 | -0.605448026 | 1.86E-08 |
| OR5BA1P | 8.492811232 | 9.53E-08 |
| AC004923.4 | 2.168556038 | 0.003191082 |
| AP001922.5 | 7.199837387 | 8.61E-06 |
| AL096711.2 | 1.953285588 | 3.23E-05 |
| AP001830.1 | 4.073706814 | 0.007659198 |
| AL133352.1 | 0.18214859 | 0.028987785 |
| NOX5 | 1.280082284 | 0.001530465 |
| AC023946.1 | -0.82592609 | 2.03E-06 |
| AP000640.2 | 3.79824021 | 1.98E-05 |
| AC015689.1 | -1.280965821 | 0.014460065 |
| Z97989.1 | -1.442137019 | 0.014921513 |
| AC022182.3 | -0.212837503 | 0.034690809 |
| PCDHA3 | -1.030458551 | 2.64E-08 |
| OR8G3P | 2.391474716 | 5.89E-05 |
| AC023538.1 | -1.165024273 | 0.010231623 |
| AL354813.1 | 0.899456631 | 0.046412036 |
| AC008750.2 | 6.725229658 | 3.55E-05 |
| CD44-AS1 | 1.235993097 | 0.000427019 |
| AP002812.5 | 1.47000276 | 0.023997378 |
| AP003486.1 | 1.121816555 | 5.48E-06 |
| AC108471.3 | 3.962991118 | 1.35E-05 |
| AC109635.5 | 0.632033187 | 0.004499262 |
| AP002990.1 | 0.471924662 | 9.04E-08 |
| AC005363.1 | -0.848343083 | 0.011721521 |
| AP002748.4 | 1.606333156 | 3.13E-05 |

| | | |
|---|---|---|
| AL356215.1 | 1.348689801 | 0.003307674 |
| NPIPB8 | 1.284097618 | 4.34E-05 |
| AP000873.5 | 2.516314573 | 0.000208921 |
| ZNF252P-AS1 | 1.903716182 | 0.00238567 |
| FDXACB1 | -1.384814837 | 8.94E-06 |
| AC073651.1 | 2.342327529 | 0.002699396 |
| BRWD1-AS2 | 2.618827577 | 0.000495983 |
| AC006511.3 | 1.787653868 | 0.003672786 |
| RAB44 | 5.867255335 | 9.06E-06 |
| AC023790.2 | 8.686972337 | 6.06E-08 |
| PCDHB17P | -0.759543066 | 0.00013431 |
| AC073585.1 | 1.463432437 | 2.58E-05 |
| AP002373.1 | 0.782258209 | 0.029946675 |
| TRIL | -3.289513917 | 3.11E-06 |
| SNHG1 | 0.112825737 | 0.019355882 |
| AC011462.1 | 0.379411606 | 0.001153986 |
| AC009533.3 | 0.801130694 | 0.014263892 |
| KRT8P11 | 6.656861847 | 1.56E-06 |
| AL117348.2 | 0.321212774 | 0.008416978 |
| AP003717.1 | 2.998910186 | 0.001195982 |
| PXN-AS1 | -0.300137672 | 0.000275212 |
| DENND5B-AS1 | -1.008123684 | 3.21E-05 |
| AL138752.2 | -0.784131415 | 2.11E-08 |
| LINC00346 | 0.609359968 | 0.010626484 |
| AC008813.1 | 0.471643608 | 0.040137558 |
| AP000777.2 | -0.389258897 | 0.035450851 |
| AP000439.2 | -1.753713447 | 1.54E-07 |
| AC027277.1 | -0.713801886 | 0.003308549 |
| AC026362.1 | 0.272475366 | 0.000791059 |
| CTSO | 1.22320459 | 0.039467072 |
| DNAAF4 | 0.389271163 | 0.037173911 |
| URB1-AS1 | 0.867606005 | 0.00557365 |
| ZNF432 | 0.875068311 | 1.90E-07 |
| AP000721.1 | 0.383722152 | 0.002620936 |
| AP001453.2 | 0.722793941 | 0.00062277 |
| LINC01152 | 2.184796453 | 0.001009567 |
| AC027290.2 | -1.141737268 | 0.000100805 |
| ATF4P4 | 0.730868951 | 0.000891548 |
| AP002518.1 | 6.276957371 | 1.85E-05 |
| ZNF10 | 0.575314883 | 3.14E-05 |
| ZNF486 | 0.143943781 | 0.039235548 |
| SUPT16HP1 | -0.374753886 | 0.000135901 |
| HMBS | 0.474385961 | 7.60E-05 |
| AC021092.2 | 0.320475916 | 0.004204414 |
| RPL41P2 | 0.335022364 | 4.47E-05 |
| ABCC6P1 | 3.989253888 | 1.27E-05 |
| AP002748.5 | 0.371043962 | 0.001947461 |
| HSPA8P5 | -0.379351001 | 0.013261874 |

| | | |
|---|---|---|
| AC069234.2 | -0.729090943 | 7.37E-07 |
| UBE2NP1 | -0.725613812 | 0.023173266 |
| AL357673.1 | 1.492595326 | 0.003186222 |
| AP002383.2 | -2.563304854 | 2.99E-09 |
| AP003419.1 | 1.372306738 | 1.51E-07 |
| POLG2 | -0.477884221 | 4.06E-05 |
| SMIM10L1 | -1.107006684 | 5.41E-05 |
| AC156455.1 | 1.873441131 | 0.005803188 |
| AP003108.2 | -0.791738945 | 1.74E-06 |
| AC010186.2 | 0.481519432 | 0.040790824 |
| LINC02393 | -1.035103613 | 0.010134232 |
| AP002414.2 | -1.247508 | 9.53E-06 |
| AC092747.2 | 1.18419514 | 0.009217481 |
| ZBTB11-AS1 | -0.434028506 | 0.003138986 |
| AC112777.1 | -1.208145272 | 1.91E-09 |
| AC025423.2 | 1.558880287 | 5.65E-09 |
| ZNF350 | 1.064290299 | 8.43E-07 |
| AP001160.1 | -0.483641016 | 0.006320489 |
| AP002784.2 | 0.372732786 | 5.53E-05 |
| AC009511.2 | -1.991940262 | 2.09E-06 |
| ZNF253 | 0.176129998 | 0.025354202 |
| AC005832.3 | -0.771648048 | 0.024237401 |
| AP000777.3 | -0.564063399 | 0.000580158 |
| AC048338.1 | -0.229131052 | 0.005571274 |
| AP002770.2 | 0.852127552 | 0.026879234 |
| AC005840.3 | 0.948969199 | 0.035735361 |
| AC090023.2 | 2.438612607 | 0.000263259 |
| AP000763.4 | -1.284951556 | 4.20E-05 |
| KRT17P8 | 4.90460761 | 0.000276252 |
| AC131009.2 | -2.727009284 | 0.000917761 |
| AL513165.2 | 0.365570898 | 0.003185479 |
| SNRPEP2 | -0.175751282 | 0.025738342 |
| AC137590.1 | -1.38951436 | 0.000321553 |
| AP006296.1 | -0.701431016 | 0.000312695 |
| AP003721.4 | 4.21602812 | 3.49E-08 |
| AP001453.3 | 4.404306461 | 1.27E-06 |
| DENND11 | -0.985519197 | 4.86E-06 |
| LSM14A | -1.016206659 | 3.96E-08 |
| RRN3P3 | -0.391173247 | 0.046697588 |
| TMPO-AS1 | -1.035392273 | 8.47E-07 |
| CR383656.1 | -1.552603853 | 4.70E-06 |
| AC009318.1 | 2.431943735 | 3.74E-06 |
| HNRNPA1P50 | 0.444551987 | 0.045936635 |
| GATC | -0.538042721 | 2.09E-07 |
| AC007569.1 | 7.237308466 | 8.20E-06 |
| AC079601.2 | 1.035412822 | 5.75E-05 |
| LINC01619 | -1.468632227 | 0.00471731 |
| AC008014.1 | 2.252726941 | 0.002117627 |

| | | |
|---|---|---|
| ZNF271P | -1.828938592 | 5.12E-10 |
| AC127164.1 | 2.327732735 | 1.93E-05 |
| AL132780.1 | -2.397420385 | 0.017205107 |
| ZBED6 | -2.774279032 | 6.29E-12 |
| AC012555.1 | 4.253046625 | 0.001212281 |
| RACGAP1P | -0.685138315 | 0.049018924 |
| MGAM | 5.441916811 | 1.61E-08 |
| AC068888.1 | 0.341783752 | 0.000832529 |
| AC025165.2 | 0.192686676 | 0.037100102 |
| AC048341.1 | 1.180983337 | 3.43E-06 |
| AC008770.1 | 1.786313445 | 0.002757416 |
| VENTXP3 | 0.557633599 | 0.012862499 |
| FNTB | -0.52237953 | 6.75E-05 |
| AC023509.1 | 0.351958241 | 0.000398641 |
| AC025257.1 | -1.212905184 | 3.58E-05 |
| AC023055.1 | -0.26406246 | 0.001625057 |
| AC089984.1 | 1.73793797 | 0.001454713 |
| AC004241.1 | 2.785515458 | 1.72E-07 |
| AL157935.3 | 0.350227058 | 0.007030704 |
| AC063948.1 | 2.069362894 | 0.002312775 |
| PPP1R12A-AS1 | 2.102694844 | 1.00E-06 |
| HSPD1P4 | -0.183779684 | 0.001661193 |
| ZNF625 | 0.739895104 | 0.000276077 |
| GALNT4 | 0.213536732 | 0.006610951 |
| SCAT2 | -1.020959492 | 0.007794955 |
| AC073957.2 | 2.793552171 | 5.14E-07 |
| AC084032.1 | 7.694574593 | 1.98E-06 |
| AC055716.2 | -0.596897426 | 0.031989299 |
| PSMA3-AS1 | -0.270991205 | 0.004290736 |
| AC004024.1 | 5.377393926 | 9.20E-05 |
| BTBD10P1 | -1.437363126 | 0.00196825 |
| AC025265.1 | 1.782335676 | 0.001537966 |
| GIHCG | 1.99174505 | 3.71E-06 |
| LBX2-AS1 | 1.290422162 | 6.68E-07 |
| CPNE8-AS1 | 1.882342795 | 0.004779565 |
| CNPY2 | 0.178485075 | 0.004546924 |
| AC078860.2 | 8.352101158 | 1.63E-07 |
| AC002996.1 | 0.969882766 | 0.008176401 |
| AC010200.1 | 1.717139168 | 0.006238949 |
| CNOT2-DT | 2.26654435 | 1.32E-07 |
| AC011611.4 | 1.344853191 | 0.001222785 |
| AC007298.2 | 2.867121037 | 2.02E-07 |
| AL512310.8 | -1.442617604 | 7.31E-05 |
| DDN-AS1 | 0.559143846 | 0.046316942 |
| AC025165.3 | -0.62146529 | 0.014641271 |
| CUX1 | -0.456172426 | 2.25E-05 |
| TEN1 | -0.425506667 | 0.040512074 |
| P2RX5-TAX1BP3 | -0.38468048 | 4.40E-05 |

| | | |
|---|---|---|
| QRSL1P3 | 3.939711772 | 6.38E-06 |
| AC126614.1 | 5.088905965 | 0.000136352 |
| HMGA1P3 | 0.422867255 | 0.009649624 |
| AC026111.1 | -0.528464768 | 0.012775384 |
| AC011603.3 | -0.292265185 | 8.11E-06 |
| AC073569.1 | 4.788492683 | 0.000272843 |
| AC009779.3 | 1.149359734 | 7.78E-06 |
| BCDIN3D-AS1 | 1.42443426 | 2.13E-07 |
| AC093014.1 | -0.608401532 | 0.00814075 |
| AC008149.1 | -0.540735863 | 0.020059128 |
| SLC7A5P2 | -0.944099864 | 1.06E-06 |
| AC004801.5 | 2.084475218 | 0.001833012 |
| AC125611.3 | -1.167535517 | 2.79E-10 |
| AC002351.1 | 3.341801103 | 0.000301869 |
| RPL10P13 | 1.446117479 | 0.027432716 |
| CHURC1 | -0.325348892 | 0.020721983 |
| AC009779.4 | 0.769445278 | 0.000225415 |
| C17orf49 | 0.426766373 | 8.23E-05 |
| AC034102.6 | 0.626078906 | 0.010530406 |
| ITFG2-AS1 | 1.028246344 | 0.018961261 |
| AC125611.4 | 1.659328599 | 9.95E-05 |
| PCNPP1 | 0.623519212 | 0.029651294 |
| RTEL1 | -0.427215978 | 0.000495983 |
| AL139099.2 | -0.584809593 | 6.42E-07 |
| RBM8B | -0.329570101 | 4.63E-05 |
| PDF | -0.547789495 | 3.92E-06 |
| AC048337.1 | 4.167446888 | 0.001839068 |
| LINC00641 | 2.701444238 | 1.48E-10 |
| AL132777.1 | -0.60889943 | 3.42E-05 |
| AL132780.2 | 2.792916695 | 5.08E-05 |
| AC012651.1 | 5.043254153 | 7.72E-07 |
| AL139011.2 | -0.584839889 | 3.00E-06 |
| AL049779.1 | 0.554389397 | 0.013065242 |
| CHMP4BP1 | 0.524495166 | 0.014656948 |
| LINC02207 | 1.706816636 | 0.001235312 |
| PPIAP6 | 0.434275548 | 5.08E-05 |
| SPESP1 | 1.195118968 | 2.48E-07 |
| AL157871.1 | -0.871200458 | 0.012162189 |
| LINC00239 | 1.684428842 | 0.011095653 |
| AL157871.2 | 3.84859692 | 4.18E-06 |
| AP001781.2 | -0.723234115 | 0.000701256 |
| AC068896.3 | -0.396796655 | 0.021009322 |
| AC025162.1 | -0.475287163 | 0.005399664 |
| SPECC1L-ADORA2A | -0.219727678 | 0.002769113 |
| AC005519.1 | -1.086477159 | 0.018659215 |
| NBEAP1 | 2.925132758 | 0.00019552 |
| PPIAP4 | 3.876866647 | 0.005385809 |
| SERPINE4P | 1.006097466 | 0.004487915 |

| | | |
|---|---|---|
| AL160006.1 | -0.626481891 | 0.000170853 |
| BCL2L2-PABPN1 | -0.262319703 | 0.008431103 |
| SYNJ2BP-COX16 | -0.221469285 | 0.037134518 |
| HSPE1P2 | 0.315068762 | 0.006591081 |
| AC005520.3 | -0.173328041 | 0.040045386 |
| ARHGAP5-AS1 | 0.621990971 | 0.00218196 |
| LINC01397 | 4.774426715 | 1.36E-06 |
| LINC00638 | 1.77570416 | 0.000208929 |
| AC023310.2 | -1.356568924 | 0.003546297 |
| SLC25A21-AS1 | 1.422390519 | 3.62E-05 |
| AL132711.1 | 7.639046884 | 2.02E-06 |
| PRC1-AS1 | -0.538671936 | 0.000758169 |
| AL135999.1 | 0.566547859 | 0.000536499 |
| AL162231.3 | 0.657606004 | 0.033956112 |
| AC025884.1 | 1.537977096 | 0.000270584 |
| AL121603.2 | -0.73492031 | 0.005515938 |
| AL132800.1 | 5.061416767 | 9.96E-05 |
| LINC01579 | -0.469876408 | 0.007224452 |
| AL133453.1 | -1.500718596 | 8.99E-06 |
| AL049779.2 | -2.065385508 | 1.23E-06 |
| AL356019.2 | 1.84390369 | 0.000138495 |
| AL161670.2 | -0.86261096 | 0.002789438 |
| LINC01580 | 2.589421728 | 0.004586439 |
| AL121594.1 | -0.402390436 | 7.29E-05 |
| LINC00520 | 8.368716547 | 1.57E-07 |
| OR11H7 | 4.788492683 | 0.000272843 |
| RNASE4 | 1.434951338 | 0.005700275 |
| AC137834.1 | 0.479880563 | 0.003390697 |
| MC1R | 0.249758947 | 0.010168378 |
| AL135818.1 | 7.433296986 | 4.86E-06 |
| AC007040.2 | -0.785922344 | 0.000332859 |
| NPM1P5 | 0.423704695 | 1.09E-05 |
| AL132639.2 | 3.620964981 | 0.001530465 |
| TUBB3 | -0.155069028 | 0.001955087 |
| AL118558.1 | 0.642548605 | 0.016719821 |
| AL137129.1 | -0.843210822 | 2.16E-06 |
| HMGN1P3 | 2.268695685 | 0.003232559 |
| AL133523.1 | 8.873430697 | 2.48E-08 |
| UBE2F-SCLY | -0.397950426 | 0.001812258 |
| AL135978.1 | -1.090181254 | 1.77E-05 |
| AL355075.4 | 0.783259292 | 0.041574479 |
| AC005479.1 | 1.659588876 | 0.012350134 |
| AL049869.1 | -1.694517031 | 4.06E-05 |
| AL139317.3 | -1.43264297 | 1.92E-08 |
| POC1B-GALNT4 | 0.217714457 | 0.006127802 |
| SHLD2P2 | -1.296671734 | 0.00069069 |
| AC025884.2 | 1.79601455 | 0.003605067 |
| AL157791.1 | -1.184156696 | 0.048824979 |

| | | |
|---|---|---|
| NDUFC2-KCTD14 | -0.319623889 | 0.001142939 |
| AC103996.3 | 3.207285148 | 3.08E-05 |
| AL132780.3 | 1.092507502 | 3.89E-06 |
| AL445363.2 | 2.572598814 | 0.003544068 |
| AL109766.1 | -0.430992975 | 0.002832796 |
| RANBP20P | -1.371122274 | 2.45E-05 |
| AC004816.1 | 1.739807346 | 0.000217235 |
| ADAM20P1 | 3.501751735 | 0.000203432 |
| GNRHR2P1 | -0.474341794 | 0.000198296 |
| AL163636.3 | 1.862624263 | 0.002550789 |
| AC023024.1 | 4.026126735 | 3.59E-07 |
| AC090971.2 | -1.372590548 | 2.76E-05 |
| AC020658.2 | 0.946727764 | 0.039705288 |
| PRKXP1 | 1.050344448 | 0.009901799 |
| ITGB3 | 0.794877281 | 2.50E-08 |
| AC004943.1 | 1.26452367 | 0.034822021 |
| AC103739.1 | 1.834774314 | 0.012214253 |
| LINC02323 | 1.44503206 | 0.028244676 |
| USP3-AS1 | -0.580442996 | 0.005901774 |
| AC018904.1 | -1.080695117 | 0.020767592 |
| AC104590.1 | 2.686448842 | 0.002973858 |
| AC087477.2 | 1.115270762 | 0.006913562 |
| AC087639.1 | 8.633281535 | 8.19E-08 |
| BUB1B-PAK6 | -1.025713012 | 5.56E-05 |
| ZNF710-AS1 | -1.275007252 | 9.92E-05 |
| AC103739.2 | 3.683047325 | 6.35E-06 |
| ACTG1P17 | 6.238575253 | 7.78E-05 |
| AC087632.2 | 0.240025917 | 0.040720082 |
| AC105129.2 | 7.529680116 | 3.92E-06 |
| AC011921.1 | 4.673905018 | 0.000349487 |
| AL590133.2 | -0.29393744 | 0.000606585 |
| AC026951.1 | -1.907211012 | 5.38E-05 |
| AC090825.1 | 0.71282877 | 0.044179399 |
| AC108449.2 | -1.641784992 | 0.000547334 |
| DCAF13P3 | -0.540232813 | 0.035403749 |
| AC090695.2 | 1.496242036 | 0.022676717 |
| TGIF2-RAB5IF | -0.486536905 | 1.16E-05 |
| ISCA1P4 | 4.626179802 | 0.001031723 |
| HNRNPA1P45 | 1.636606813 | 0.005081272 |
| CTXND1 | 2.6254468 | 4.02E-05 |
| AC027237.5 | -1.106711397 | 0.000880942 |
| THTPA | 0.295917739 | 0.02653956 |
| AC105339.3 | 4.178371727 | 2.30E-05 |
| AC023355.2 | 0.369007937 | 0.013781012 |
| AC023968.2 | -0.528113806 | 0.004436972 |
| MRPL46 | -0.858200714 | 6.05E-07 |
| AL132640.2 | 5.261724743 | 5.22E-05 |
| AC051619.4 | 3.016464766 | 0.000113218 |

| | | |
|---|---|---|
| AC022613.2 | 4.306311505 | 4.76E-10 |
| LINC00052 | 1.147644704 | 0.00615255 |
| AL136295.5 | -0.371796855 | 0.005288044 |
| AC091045.2 | -0.631888164 | 0.003081287 |
| HMGN1P26 | 8.578151407 | 9.29E-08 |
| AC090607.2 | -1.031084426 | 0.00224467 |
| CERNA1 | 2.791276047 | 0.002584345 |
| AC015712.2 | -0.84846495 | 8.53E-10 |
| RBM17P4 | -0.79662082 | 0.005854443 |
| AC015712.4 | 3.451942923 | 1.00E-06 |
| EEF1A1P22 | 0.718078136 | 0.002119087 |
| AC079328.2 | -0.706983584 | 4.99E-05 |
| AC104046.1 | 1.570187777 | 0.005581518 |
| ST20-AS1 | 1.212995137 | 1.86E-06 |
| AL132640.3 | 8.211395164 | 2.86E-07 |
| AC090825.2 | 7.305640991 | 6.09E-06 |
| AC009996.1 | -1.043249584 | 0.045028334 |
| DNM1P47 | 3.976757527 | 6.12E-06 |
| AC010478.1 | 3.58409956 | 7.90E-11 |
| AC126323.2 | 9.737608171 | 1.77E-09 |
| AC084757.3 | 3.477071051 | 2.04E-06 |
| HSP90B2P | 0.583718054 | 2.36E-07 |
| AC022087.1 | 3.668267051 | 0.007626055 |
| CRTC3-AS1 | 2.021127232 | 0.004030422 |
| NCAPGP2 | -0.874104228 | 0.042427666 |
| AC068234.1 | 0.869254509 | 4.24E-08 |
| AL138976.2 | 2.398904392 | 0.001110211 |
| AC009065.2 | 1.466911966 | 0.003162058 |
| AC012640.4 | 3.350991875 | 4.16E-09 |
| AC009093.1 | -1.570626812 | 0.006312137 |
| AC002519.1 | 7.814787595 | 1.39E-06 |
| AC018845.1 | 6.671503917 | 4.29E-05 |
| LYPD8 | -0.54850955 | 0.047455774 |
| AC072061.1 | 1.314869176 | 0.001321166 |
| ELOCP2 | -0.199946289 | 0.014243779 |
| AL390728.6 | -1.434163833 | 5.98E-05 |
| AC009113.1 | 2.079845325 | 2.20E-08 |
| AC126323.3 | 0.6936326 | 0.003121398 |
| NR4A1AS | 5.049396135 | 5.35E-15 |
| AC106820.2 | -1.613424325 | 0.019407493 |
| ACTG1P15 | 3.499470902 | 0.010900413 |
| HNRNPLP2 | 0.265513818 | 0.000681711 |
| AC011939.1 | -0.556835404 | 0.044841888 |
| AC022167.1 | -0.712396894 | 0.004186529 |
| AL050341.2 | 0.384452239 | 0.034443161 |
| AC124068.1 | -0.353940199 | 0.004944737 |
| IL21R-AS1 | 3.288680311 | 6.52E-05 |
| RBM15B | -0.115333575 | 0.023946463 |

| | | |
|---|---|---|
| AC107068.1 | -1.002516899 | 4.02E-05 |
| AC009120.2 | -0.229912217 | 0.006723987 |
| AC093010.3 | -0.934043564 | 0.022912692 |
| AL928711.1 | 3.183808526 | 0.000466049 |
| AL133338.1 | 1.576191199 | 2.29E-05 |
| HOXB7 | -1.124628729 | 7.62E-07 |
| NORAD | 0.393754227 | 7.80E-08 |
| AL512408.1 | 2.321781596 | 1.47E-06 |
| AC023824.1 | -1.423794344 | 0.00746313 |
| AF274858.1 | 2.335960285 | 0.000625198 |
| AC106886.1 | 3.857003195 | 8.21E-05 |
| MIR762HG | 2.375676632 | 8.27E-08 |
| DDX59-AS1 | 8.755690437 | 6.81E-08 |
| AC093752.3 | 2.008526835 | 1.55E-05 |
| AC009163.3 | 2.025558589 | 0.005913659 |
| AC034111.1 | 3.647994915 | 0.01205643 |
| DNM1P33 | 6.67267955 | 4.13E-05 |
| AC008074.2 | 3.112382843 | 4.13E-06 |
| AC012435.2 | 2.491433805 | 1.76E-06 |
| AOC4P | 2.458791885 | 0.001641769 |
| AC026464.2 | 0.509170111 | 0.010981067 |
| AC138028.4 | -0.469493047 | 0.001762187 |
| AC068987.1 | -2.026731671 | 0.011427876 |
| AC008915.3 | -1.079089393 | 0.000211493 |
| CSPG4P13 | 0.809268973 | 5.51E-05 |
| AC145350.2 | 0.926839199 | 0.02246938 |
| AC126696.1 | 8.264448429 | 2.58E-07 |
| LINC02846 | 2.330777774 | 7.34E-05 |
| AC124798.1 | -1.049915858 | 0.001178335 |
| CD2BP2-DT | 1.589995249 | 3.83E-06 |
| AC009119.1 | 0.960615251 | 0.023494442 |
| KDM7A-DT | 0.620579255 | 0.011285011 |
| ZNRD2-AS1 | 0.729583355 | 0.004259315 |
| AC099778.1 | 1.38773365 | 0.000149392 |
| AC084262.1 | 2.38677195 | 0.000638173 |
| AL035071.1 | 1.698281626 | 1.61E-05 |
| LINC02166 | 2.020914752 | 0.001242137 |
| SNHG19 | 0.815717792 | 2.88E-06 |
| AC124944.2 | 0.892524663 | 0.017742455 |
| AC026471.1 | 1.623821766 | 8.44E-09 |
| AC105036.3 | -0.703580725 | 0.000935443 |
| AC093525.2 | -0.163195413 | 0.011716757 |
| AC068338.2 | 1.043629567 | 0.000183492 |
| AC022167.2 | -0.865541856 | 1.25E-07 |
| AC009019.1 | -0.694416026 | 0.001119897 |
| AC137932.1 | 1.280445965 | 0.005396253 |
| SLX1B-SULT1A4 | -0.987122676 | 3.52E-05 |
| AL133367.1 | 2.285907594 | 3.17E-05 |

| | | |
|---|---|---|
| AC019294.2 | 1.519265387 | 0.029227986 |
| AC106820.4 | -1.456611722 | 0.011305194 |
| AC009088.2 | -0.338290483 | 0.001246465 |
| AL008727.1 | 4.910059009 | 6.69E-07 |
| AC104024.2 | 0.775362306 | 0.00069644 |
| HEXA-AS1 | 3.069662306 | 1.54E-06 |
| AL353708.1 | 2.272762726 | 0.002133105 |
| AC106028.3 | 1.165227437 | 0.027266573 |
| AC026464.3 | -0.542361851 | 0.017164016 |
| LDC1P | 7.044885087 | 1.40E-05 |
| WBP11P1 | -0.477081172 | 0.029971028 |
| AC022336.2 | 2.685453469 | 0.002931654 |
| AC110079.1 | 0.50579877 | 0.001155826 |
| AC012414.4 | 2.746100107 | 1.95E-05 |
| AC040169.2 | 4.057006337 | 0.006174279 |
| LINC02367 | 0.422224365 | 0.000421924 |
| AL031709.1 | -1.610845885 | 0.008146537 |
| ATP2A1-AS1 | 0.664069935 | 0.026383493 |
| C16orf95 | 1.142592835 | 0.000495402 |
| AC018557.1 | -1.735933747 | 1.15E-09 |
| AC126696.2 | -1.17246246 | 0.001324431 |
| AC023794.7 | 2.559614559 | 0.001375892 |
| AL353719.1 | 2.488597465 | 0.001662062 |
| AC008870.2 | 0.36874208 | 0.003023726 |
| MYL12BP1 | 0.282173476 | 0.049762589 |
| AC109347.1 | 2.645265235 | 1.30E-05 |
| FAM157C | 0.976403359 | 3.71E-05 |
| AC023043.1 | 0.536153267 | 0.041420971 |
| AC132872.1 | 1.165360178 | 0.000296552 |
| ERVK13-1 | -1.172296666 | 2.46E-07 |
| AC127459.1 | -1.749857927 | 1.69E-05 |
| BNIP3P5 | 1.693298704 | 0.015829431 |
| AC069224.2 | 2.400346315 | 0.002547817 |
| AC011374.1 | 7.198194824 | 8.80E-06 |
| STAM-AS1 | 1.328289735 | 0.000470515 |
| AC012531.1 | -1.11157292 | 0.000247813 |
| AL590004.3 | 1.562425416 | 0.00033422 |
| Z98885.2 | 4.996579377 | 0.000258185 |
| AC025917.1 | 1.142730778 | 0.012176027 |
| AC068870.1 | 5.11298681 | 0.000167785 |
| AC026471.2 | 5.003116967 | 1.40E-07 |
| SNAI3-AS1 | 3.161602599 | 5.98E-07 |
| AC114811.2 | 3.80142865 | 0.000122315 |
| AC092718.3 | 2.898364343 | 1.30E-05 |
| HERC2P5 | -0.566045229 | 0.026061453 |
| AC237221.1 | 3.498677799 | 0.011961573 |
| AL096870.2 | 0.671874902 | 1.32E-05 |
| AL591848.3 | -1.454105467 | 0.00210763 |

| | | |
|---|---|---|
| AC133552.1 | 6.958751275 | 1.86E-05 |
| SLC7A5P1 | 1.047728681 | 0.00554881 |
| AC009690.1 | 0.766722068 | 0.00052011 |
| AC139256.2 | 2.535957137 | 0.000865369 |
| AC007823.1 | 4.198128721 | 0.00140273 |
| AC093520.1 | 1.824719944 | 6.26E-05 |
| AC012321.1 | -1.162286203 | 3.32E-08 |
| AC061975.1 | 5.530954086 | 2.86E-09 |
| KRT8P50 | 1.128144408 | 0.04849653 |
| AC092803.1 | 2.005350826 | 0.000871274 |
| CEROX1 | 1.48308234 | 0.002226194 |
| AC109460.2 | -1.505700865 | 1.62E-06 |
| TAT-AS1 | 3.704098688 | 6.40E-05 |
| AC105020.4 | 0.88132902 | 6.29E-07 |
| AC106886.2 | 0.638039644 | 0.011779518 |
| LINC00565 | 1.335196098 | 3.73E-07 |
| AL158206.1 | 2.172103652 | 2.00E-09 |
| CCPG1 | -0.149188843 | 0.014370021 |
| AL158212.3 | -1.358408728 | 7.87E-06 |
| LINC01311 | 1.839991542 | 1.92E-05 |
| AC016134.1 | 7.815325753 | 1.27E-06 |
| LINC00622 | 0.718400232 | 5.37E-05 |
| CAPN10-DT | 0.473556568 | 0.030962596 |
| AL356489.2 | -1.038016143 | 0.004071586 |
| AL390195.3 | 0.978187626 | 0.003855999 |
| AC119674.1 | 3.434299772 | 1.56E-06 |
| AL136982.4 | 1.713262448 | 0.000292322 |
| AC132825.3 | 1.751057596 | 0.024913991 |
| AC105046.1 | 1.855116373 | 3.33E-06 |
| SULT1A3 | -0.917989187 | 1.89E-07 |
| AC036108.2 | -1.155601831 | 1.19E-08 |
| AC092718.4 | -0.893992838 | 6.91E-05 |
| AC009139.2 | -0.742866296 | 0.016598312 |
| AL441883.1 | 1.607664648 | 0.011371446 |
| AC009053.3 | 1.903494502 | 4.26E-06 |
| AC073476.3 | 3.409412327 | 1.17E-07 |
| AC234775.3 | 3.136168156 | 7.56E-05 |
| AC093904.4 | 8.366142181 | 1.75E-07 |
| TMEM178B | -0.944371379 | 0.000153953 |
| AL049555.1 | -1.866842439 | 0.003482249 |
| RBFADN | 1.988678344 | 1.09E-05 |
| AC133548.2 | 2.666057639 | 5.84E-05 |
| AL137802.2 | 3.627517329 | 5.82E-08 |
| AC023908.3 | 1.327152482 | 5.19E-05 |
| ADAMTS7P3 | 3.28700168 | 2.81E-05 |
| AC091167.2 | 0.199241585 | 0.002338429 |
| EPPK1 | 4.730682325 | 1.66E-12 |
| AC012182.2 | -1.245445605 | 0.046140582 |

| | | |
|---|---|---|
| AC112484.3 | 3.710720267 | 2.56E-07 |
| AC107027.3 | -0.735521673 | 0.019846937 |
| AC018845.3 | 2.543315334 | 0.004858052 |
| Z95115.1 | 1.919129811 | 3.89E-05 |
| AL365475.1 | -0.921009014 | 0.010393708 |
| AL162231.4 | 1.333115044 | 0.001551542 |
| AC103706.1 | 1.444200462 | 0.003553727 |
| ZNF865 | 0.767525253 | 3.64E-06 |
| AC021483.2 | -0.227844472 | 0.043678303 |
| BOP1 | 0.162496472 | 0.004266986 |
| ANKRD26P1 | 3.76720728 | 0.017912263 |
| AL136038.5 | 3.677306034 | 0.00780885 |
| Z97055.2 | 2.337306195 | 0.002435381 |
| AC008870.4 | -0.822604441 | 0.014335136 |
| ATP2C2-AS1 | 3.259679432 | 6.40E-08 |
| AC126323.6 | -1.168233029 | 5.98E-06 |
| AC002550.1 | -0.928766295 | 1.56E-05 |
| AC105275.2 | 6.725229658 | 3.55E-05 |
| LARP4P | -0.496657315 | 0.008530756 |
| AC010168.2 | 1.210891599 | 0.000582914 |
| LINC01355 | 1.565711285 | 2.33E-06 |
| AC006538.1 | 0.680299208 | 0.015849644 |
| PYCARD-AS1 | 3.224590119 | 2.68E-05 |
| AC010491.1 | 3.18617231 | 6.05E-07 |
| VPS9D1-AS1 | 0.301060696 | 0.02961913 |
| AC138627.1 | 7.199837387 | 8.61E-06 |
| AC100756.2 | 3.013132419 | 0.000136239 |
| AL022069.1 | 5.16002604 | 0.000841937 |
| TMEM202-AS1 | 0.894792169 | 0.002308898 |
| LINC01003 | 1.923193254 | 1.67E-06 |
| AC135048.1 | 0.696119564 | 0.004620775 |
| TBILA | 1.320049553 | 0.005540724 |
| AL031714.1 | -0.231554721 | 0.010679883 |
| TP53TG3B | 1.622494827 | 0.00190183 |
| AC012615.1 | 0.793705262 | 0.006129974 |
| AL596244.1 | -0.578561286 | 0.032964365 |
| FSCN1P1 | -0.50164189 | 0.005509194 |
| AP003119.3 | -0.968811463 | 0.002344873 |
| TMEM249 | 1.360148907 | 0.000453985 |
| AC005632.3 | 2.523185462 | 0.001219829 |
| AC233266.2 | 1.208935422 | 0.005315161 |
| AC092115.3 | -1.108533905 | 7.52E-07 |
| AC114947.2 | 0.620252343 | 0.000829271 |
| AC091230.2 | 3.867294399 | 6.16E-08 |
| AC010547.4 | 1.067621279 | 3.88E-08 |
| SUB1P3 | 0.30904379 | 0.008597695 |
| AC093525.6 | 1.119982546 | 0.008544145 |
| AC036108.3 | 0.769173174 | 0.005930708 |

| | | |
|---|---|---|
| LINC02177 | 3.569889966 | 0.005291917 |
| LINC02605 | 0.81415984 | 0.027575306 |
| AL031600.3 | -0.327401529 | 0.030675807 |
| C15orf65 | 0.892462411 | 0.03422542 |
| Z92544.2 | 0.377247074 | 0.019426238 |
| AC009065.9 | -1.438712912 | 0.015849644 |
| AC093591.2 | -1.37497901 | 0.018667973 |
| AC012213.2 | 8.676338734 | 5.00E-08 |
| AC018362.1 | -0.437564351 | 0.00094587 |
| Z97986.1 | -0.837412964 | 0.021332539 |
| AC134682.1 | 4.635113222 | 0.000394149 |
| AC092134.1 | 1.955989328 | 0.00362851 |
| H2BC20P | 2.990705706 | 8.58E-13 |
| AC009163.5 | 0.739474643 | 0.007014154 |
| AL031708.1 | 0.446963237 | 5.62E-05 |
| AC117382.2 | 3.238223668 | 0.00024935 |
| AC099518.5 | -0.498193347 | 0.000878414 |
| LINC02616 | 3.378676093 | 2.92E-08 |
| AC027228.2 | -1.123936527 | 1.21E-05 |
| AC012184.3 | 2.645397917 | 6.28E-07 |
| LINC02582 | 1.23853168 | 0.000872258 |
| AC006058.1 | -0.540527421 | 3.99E-05 |
| TCF24 | 2.002983535 | 0.008593321 |
| AC005606.2 | -0.967020756 | 0.000163114 |
| ISY1-RAB43 | -0.266389689 | 0.004077441 |
| AC007406.5 | 0.718742877 | 0.000588805 |
| TUBB8P7 | 4.722363397 | 7.37E-07 |
| AC138932.4 | -0.802045637 | 0.000673041 |
| AC018362.2 | -0.893577709 | 0.001841964 |
| LINC00662 | 0.69049343 | 1.48E-05 |
| AL358933.1 | 3.738102034 | 0.000149058 |
| AC093249.6 | -1.066484978 | 0.000289015 |
| AC124283.1 | -2.119833668 | 5.55E-08 |
| MIA | 0.393891091 | 1.31E-05 |
| AC130462.3 | -0.803147581 | 0.003439001 |
| AC005670.1 | 7.610770794 | 2.30E-06 |
| AC144831.1 | 2.455069957 | 0.000171774 |
| AC108134.2 | 2.185936949 | 0.000374608 |
| RPL23AP84 | 1.719993968 | 0.010784523 |
| AC026954.2 | 0.249521472 | 0.002508939 |
| AC127496.1 | -2.744139459 | 4.97E-05 |
| AC007336.1 | -1.330062581 | 0.000765969 |
| AC005920.4 | 6.867151347 | 2.94E-05 |
| TCF20 | -0.333426131 | 7.90E-05 |
| LINC01974 | 8.525454919 | 1.41E-07 |
| SNORD3B-2 | 1.324337407 | 0.006829302 |
| AC015912.1 | -1.410066701 | 3.49E-06 |
| AC010614.1 | 1.434007074 | 0.017684618 |

| | | |
|---|---|---|
| AC133065.1 | 2.442874138 | 0.001281806 |
| LINC02175 | 4.997714224 | 8.33E-07 |
| AC020978.5 | -1.601280429 | 4.98E-10 |
| KRT39 | 7.003868569 | 1.61E-05 |
| AC007952.4 | -1.204908904 | 0.013203795 |
| AC008914.1 | -0.411036564 | 0.000529755 |
| CORO7 | -0.512726966 | 0.010976359 |
| AC027796.3 | -0.472217708 | 3.21E-05 |
| AGK | -0.315451599 | 0.002893201 |
| AL160291.1 | 1.606442528 | 0.003961204 |
| AC145207.2 | -0.721240676 | 3.88E-08 |
| AC007613.1 | 1.02949763 | 0.001026229 |
| AC004771.3 | -1.025379122 | 0.009272759 |
| AC006435.1 | 1.26021502 | 8.77E-06 |
| LINC01569 | 3.148844215 | 6.93E-09 |
| TMEM256-PLSCR3 | 0.830492055 | 7.85E-07 |
| AC037482.1 | 7.952441495 | 7.63E-07 |
| AC133065.2 | 5.998768899 | 6.92E-06 |
| AP000705.5 | 1.838055346 | 0.003037185 |
| TXNP4 | 0.359402435 | 0.021385571 |
| AC120057.2 | 0.377440174 | 0.002821111 |
| AL022341.2 | 0.709254308 | 0.026857532 |
| AC090617.3 | -0.385882728 | 0.008121722 |
| AC018512.1 | 0.660472921 | 0.0301013 |
| AC087741.1 | 0.762586416 | 0.00012043 |
| TMEM231P1 | 1.907236991 | 6.13E-06 |
| LINC01979 | 3.058252596 | 4.14E-07 |
| AC133552.2 | 1.044955562 | 0.030241991 |
| TDO2 | 4.829884676 | 0.000214932 |
| AC099489.2 | 6.777028881 | 3.25E-05 |
| AC139530.2 | 0.312238142 | 0.000931256 |
| OVCA2 | -0.574526525 | 2.23E-06 |
| AC005722.3 | 9.046574667 | 1.36E-08 |
| AC133552.3 | 7.525745863 | 3.02E-06 |
| AC009121.3 | -1.565704762 | 0.026567943 |
| AC010401.1 | -0.507486472 | 0.014581216 |
| AC217785.2 | 1.980312157 | 1.18E-06 |
| SSBP1 | -0.282224367 | 0.006701788 |
| MRPL12 | 0.230460605 | 0.001818441 |
| AC145207.3 | 0.395979049 | 4.36E-05 |
| AC113189.2 | 1.162259066 | 0.001912734 |
| TMPOP2 | -1.728433944 | 5.98E-05 |
| CCNQ | -0.413910372 | 0.002099902 |
| ALOX12P2 | 1.035219362 | 0.001504159 |
| EIF4A1P9 | 1.012125595 | 0.03654551 |
| AC005921.3 | -1.130831774 | 1.35E-05 |
| GTF2I | -0.320663256 | 0.000296448 |
| ZNF234 | 0.186465379 | 0.011856687 |

| | | |
|---|---|---|
| AC007114.1 | 4.010154594 | 1.31E-05 |
| ROCK1P1 | -0.839857288 | 0.000509123 |
| AC108134.4 | 3.970057523 | 5.14E-05 |
| AC090617.5 | 1.51862536 | 9.45E-05 |
| ZNF213-AS1 | 2.083772928 | 1.00E-08 |
| AC009121.4 | 3.914693528 | 0.002726178 |
| AC040162.3 | -0.480712046 | 4.12E-05 |
| SLC27A3 | 0.648939331 | 0.025836271 |
| AC087500.2 | 3.898765983 | 0.004728704 |
| AC087292.2 | -1.898088657 | 4.50E-07 |
| RYKP1 | -0.589779114 | 0.009850051 |
| AC217785.3 | 2.976504957 | 1.72E-08 |
| ATP5F1AP3 | 0.422495903 | 0.0093257 |
| AC006111.2 | 0.693939096 | 0.006431118 |
| KRT12 | 4.862974852 | 0.000193125 |
| AC087190.3 | -0.217554829 | 0.001094457 |
| RPS7P1 | 0.218670101 | 3.62E-05 |
| AC020978.7 | -1.547686261 | 4.07E-08 |
| AC003965.1 | 3.930249735 | 0.004667858 |
| SCAMP3 | 0.178372228 | 0.006462657 |
| AC068234.2 | 1.994898841 | 8.87E-05 |
| KRT23 | 2.272154133 | 0.008481457 |
| AC133552.4 | 7.613246072 | 2.35E-06 |
| TAPT1-AS1 | 0.559750884 | 0.012883466 |
| AF001548.2 | 1.696356533 | 0.015877665 |
| PPIAL4A | 0.575018005 | 0.042311164 |
| AC004477.1 | 1.969969396 | 1.39E-06 |
| AC110597.3 | -1.572601198 | 4.68E-06 |
| SRSF8 | -1.377871774 | 7.13E-10 |
| FAM72C | -0.474498919 | 1.81E-05 |
| AK4P1 | -0.895683951 | 3.34E-06 |
| UBBP4 | 0.148942493 | 0.040455283 |
| AC145207.4 | 0.690156455 | 0.001082663 |
| AP000919.1 | -1.138593973 | 3.32E-08 |
| BPTFP1 | 0.612507178 | 0.021976865 |
| LINC02864 | -0.452805714 | 0.000395149 |
| LINC00667 | 0.836893333 | 5.31E-08 |
| AC018521.1 | 0.681100024 | 0.012675338 |
| AC135178.3 | 1.078959348 | 3.85E-09 |
| RDM1P5 | 4.812994803 | 1.19E-06 |
| SINHCAFP1 | -0.889472707 | 0.003436329 |
| AC022211.3 | -0.265959112 | 0.016232469 |
| LINC00672 | 2.511825916 | 0.006107361 |
| AP000845.1 | 2.985670401 | 1.25E-05 |
| SNORD3A | 1.557416721 | 8.20E-05 |
| AC015818.3 | 7.766563296 | 1.51E-06 |
| NBPF11 | 0.22982157 | 0.005152006 |
| AC087393.2 | -0.53230966 | 0.000454678 |

| | | |
|---|---|---|
| AC073508.2 | -0.189838294 | 0.034439958 |
| AC015813.1 | 3.656886048 | 5.09E-12 |
| AC104984.2 | 4.827862188 | 0.00031645 |
| AL353997.2 | 9.783241211 | 1.64E-09 |
| AC055811.2 | 0.390998167 | 0.033671317 |
| AC239868.1 | 1.89738541 | 5.03E-05 |
| LINC00909 | 1.092408187 | 3.27E-07 |
| AC016596.3 | 0.443126515 | 4.27E-06 |
| CCDC58P1 | -1.140791914 | 0.000185484 |
| AC134978.1 | -0.579921988 | 9.01E-05 |
| NOTCH2NLA | 0.714981717 | 2.19E-06 |
| SNRPGP2 | -0.187848011 | 0.04956569 |
| DYNLL2 | 0.146437037 | 0.010272184 |
| MYH4 | 3.5249801 | 2.03E-08 |
| MIR4653 | 2.308277043 | 0.003352979 |
| NCMAP-DT | 0.568930179 | 0.012126894 |
| AC061975.4 | 4.598856694 | 0.000696172 |
| SUZ12P1 | -0.629566676 | 0.00872251 |
| AL359922.1 | -0.856734606 | 3.67E-05 |
| MIR4257 | 5.659968521 | 1.48E-05 |
| AC015674.1 | -0.210717045 | 0.037106957 |
| SNX19P3 | -1.030336403 | 0.013993604 |
| LINC00526 | 0.699298868 | 0.001416964 |
| AC009630.3 | -0.596137344 | 0.002714009 |
| AC138696.1 | 1.294638548 | 2.76E-07 |
| SEPTIN4-AS1 | 4.373072962 | 0.001054824 |
| AC018521.5 | 1.524327 | 0.001433445 |
| SH3GL1P2 | -1.068149764 | 0.041147356 |
| AGAP12P | -0.644435248 | 0.019557824 |
| GJA5 | -1.328291634 | 0.001023966 |
| TSPOAP1-AS1 | 0.708629763 | 0.022552807 |
| LRRC37A7P | -1.422622631 | 0.030659519 |
| AC005726.3 | 1.080477156 | 0.00132937 |
| SRP72P1 | -0.462572622 | 0.025482513 |
| AL359922.2 | -0.322574423 | 0.020965372 |
| AC010761.3 | 1.080164798 | 0.003901495 |
| AC103810.2 | 3.896519708 | 7.11E-05 |
| RBM8A | -0.337861746 | 1.06E-05 |
| AP005263.1 | 0.344190734 | 0.018008426 |
| PGDP1 | 0.727669473 | 7.00E-05 |
| AC005726.4 | 0.475203025 | 0.035108034 |
| AC099850.3 | -0.425655904 | 0.022418847 |
| AC130324.2 | -1.761595622 | 0.007349794 |
| TIMM23 | -0.947221433 | 3.85E-08 |
| RN7SL219P | 3.473260502 | 1.14E-05 |
| AC093484.3 | 0.332546811 | 1.19E-05 |
| AC099850.4 | -0.466330922 | 0.001343461 |
| AP005119.2 | -0.857714229 | 0.001168911 |

| | | |
|---|---|---|
| AC012447.1 | 2.283537301 | 0.00064173 |
| AC132938.4 | 3.911970849 | 3.77E-05 |
| KRT18P55 | 4.394830802 | 2.62E-08 |
| RNF115 | -0.364143889 | 3.72E-05 |
| AC015922.2 | 0.843824998 | 0.004198177 |
| FCGR1CP | 5.357584164 | 9.21E-05 |
| WDR45BP1 | -0.655647726 | 0.001798486 |
| C21orf59-TCP10L | -0.414836949 | 0.014752201 |
| AC002094.2 | -1.741913539 | 2.26E-05 |
| TUFMP1 | -1.116951315 | 1.12E-06 |
| RARA-AS1 | 2.158485829 | 4.32E-06 |
| RPL17 | 0.219275592 | 0.000790233 |
| MAFG-DT | -0.468470104 | 0.025566309 |
| AC074143.1 | 1.176201556 | 0.001245295 |
| LINC01970 | 2.469997796 | 0.004123382 |
| AC005821.1 | 3.754120782 | 1.87E-08 |
| AC103808.3 | 0.842587862 | 0.014284671 |
| AP001099.1 | 6.276252481 | 2.83E-06 |
| ZNF488 | -0.71264748 | 0.030586208 |
| AC018413.1 | 1.161875564 | 0.002528739 |
| AC005899.5 | -1.509140287 | 2.12E-06 |
| AC138207.6 | 6.671503917 | 4.29E-05 |
| SEC22B | -0.736145998 | 9.31E-08 |
| FSBP | -0.872787632 | 4.41E-06 |
| EEF1E1-BLOC1S5 | -0.601393682 | 0.000578977 |
| LINC01029 | -0.315076079 | 1.83E-05 |
| AP000919.2 | 1.403898182 | 0.024059892 |
| TXNIP | 1.959770508 | 4.10E-09 |
| SRGAP2 | -0.198581864 | 0.001796054 |
| POLRMTP1 | -0.259536811 | 0.042857362 |
| BAHCC1 | 0.440271288 | 0.001271366 |
| AC015813.2 | -1.344562227 | 4.24E-10 |
| AC004585.1 | -1.389474447 | 0.007500107 |
| RASSF5 | -1.488203217 | 2.01E-06 |
| STRADA | -0.606894318 | 5.02E-06 |
| KLF14 | 1.491265636 | 3.34E-05 |
| ARIH2P1 | -0.817819036 | 0.009057915 |
| AC098850.3 | 0.488414111 | 0.000365527 |
| AC061975.5 | 9.097529559 | 1.28E-08 |
| AC005410.2 | 3.110361013 | 0.000598453 |
| AC090774.2 | -2.086285466 | 6.39E-06 |
| AP002478.1 | 4.802454787 | 2.10E-08 |
| SNHG25 | 1.988619682 | 3.05E-08 |
| AP005061.2 | 6.140886537 | 0.000870061 |
| NCOA4 | -0.26603883 | 0.000204016 |
| AC005288.1 | -0.901146286 | 9.37E-08 |
| AC127024.5 | -1.56248024 | 3.59E-05 |
| AC005697.2 | 4.21337317 | 5.08E-06 |

| | | |
|---|---|---|
| MAGEA10-MAGEA5 | 0.787554473 | 0.011689435 |
| AC093484.4 | -1.996285424 | 3.94E-05 |
| AC087164.1 | -0.729380227 | 0.003378725 |
| AL135905.1 | 3.085854095 | 5.44E-07 |
| MYO15B | 0.721353828 | 0.033422443 |
| AC015688.4 | -1.795592085 | 0.012123475 |
| AP005136.3 | -1.650328165 | 0.015732015 |
| KPNA2P3 | -0.692495467 | 0.000252181 |
| AC132825.4 | 3.020928108 | 1.55E-07 |
| LINC00663 | 4.727819251 | 7.33E-05 |
| AC006116.2 | 1.201393205 | 0.001064055 |
| ZNF793-AS1 | 2.654860045 | 5.84E-05 |
| AC006213.1 | 1.282815449 | 6.33E-07 |
| AC067852.1 | 0.928120586 | 0.027174276 |
| AC006116.3 | 4.431031669 | 3.32E-07 |
| AC022916.2 | 3.534743534 | 5.04E-06 |
| AC067852.2 | 0.227505168 | 0.048700234 |
| AC087289.1 | 0.941274436 | 0.02028496 |
| AL050343.2 | -0.343486531 | 0.035450851 |
| AC090227.2 | 0.876062177 | 0.01762754 |
| AC060780.1 | 1.532842722 | 7.11E-08 |
| AC012615.2 | -0.978265053 | 0.009901799 |
| AC007780.1 | 0.83755898 | 0.000162849 |
| AC067968.1 | 0.344419961 | 0.002986235 |
| LRRC37A16P | -0.425451391 | 0.012175927 |
| ZNF850 | 0.556480565 | 5.39E-05 |
| AC006213.2 | 1.071621948 | 0.000210586 |
| PTGES3L | 1.107229864 | 0.03496705 |
| AC018761.2 | 0.616493198 | 0.003722594 |
| UXT-AS1 | 0.465906625 | 0.032165535 |
| CCDC58P3 | -0.673904641 | 0.021924599 |
| CTBP2P7 | 6.722959852 | 3.84E-05 |
| ILF3-DT | 1.162008993 | 9.17E-07 |
| AC060766.1 | -1.002688451 | 5.14E-05 |
| ZNF561-AS1 | 0.870382915 | 0.014033785 |
| AD000671.2 | 1.449820041 | 8.77E-06 |
| AC008105.3 | 2.661437405 | 0.000174256 |
| AC016588.1 | -1.089807013 | 0.005426469 |
| AC090360.1 | 0.920996627 | 0.002380007 |
| AD000091.1 | 0.905309094 | 0.000426765 |
| LINC01842 | 2.348759644 | 0.001044261 |
| AC093227.1 | -2.323675349 | 1.15E-05 |
| AC091152.2 | 1.203788251 | 0.005840087 |
| CHMP1B-AS1 | 1.733534859 | 1.65E-09 |
| AC005837.2 | 0.692737457 | 0.024037223 |
| AC022098.1 | 0.52983275 | 0.006014917 |
| PHF5CP | -0.900943699 | 0.049929394 |
| AC008770.2 | 1.052470089 | 7.78E-05 |

| | | |
|---|---|---|
| AP001029.2 | -0.800297804 | 0.000312732 |
| AC007773.1 | 3.41797061 | 1.07E-07 |
| AC020915.1 | -0.264921834 | 0.000543828 |
| AC005336.2 | 4.240182734 | 2.27E-08 |
| AC010504.1 | 1.512784377 | 0.014314226 |
| WDR7-OT1 | -1.359859416 | 0.035303904 |
| AC012615.6 | -0.973443265 | 0.005227907 |
| AP005264.3 | 7.044885087 | 1.40E-05 |
| AC105105.1 | -2.364894577 | 9.00E-06 |
| AC020928.1 | -1.049809554 | 0.047357968 |
| AC006504.3 | 1.381088986 | 0.021216008 |
| PARD6G-AS1 | 1.341873993 | 9.38E-05 |
| LINC01140 | -1.570973025 | 0.002978876 |
| MAP3K14-AS1 | 0.905015725 | 2.07E-06 |
| AC090409.1 | -2.171463886 | 8.86E-06 |
| ATF7-NPFF | 0.53250299 | 0.011285011 |
| AC138150.2 | 0.99995296 | 0.000258498 |
| AC012569.1 | -0.869870433 | 0.000637664 |
| RNFT1-DT | 2.39856271 | 1.63E-06 |
| AC011511.4 | -0.681777204 | 1.38E-07 |
| AC015911.4 | -1.337500062 | 8.35E-06 |
| AC104532.1 | 0.747605123 | 0.001207149 |
| SELENOKP1 | 2.372614426 | 0.001232824 |
| SNHG30 | 0.502671358 | 0.0135618 |
| SNHG22 | 3.176154362 | 0.002569304 |
| EIF4A2P1 | 1.308448311 | 0.036382139 |
| AC060780.2 | 1.332767172 | 0.021520461 |
| ZNF833P | 1.548029137 | 0.00052011 |
| GEMIN7-AS1 | -0.79096273 | 0.006152118 |
| RPL9P29 | 1.417155747 | 0.014692982 |
| MIR924HG | -0.767562717 | 0.004127161 |
| AC008649.2 | 3.609522825 | 3.65E-06 |
| AC011447.3 | 0.38162002 | 0.046076466 |
| AC090236.1 | -2.195654292 | 0.000474975 |
| AC090229.1 | 1.323331451 | 0.027539932 |
| LINC01901 | -2.120164642 | 0.004374887 |
| AC016582.2 | 0.35129619 | 0.002779739 |
| AC087289.3 | 0.158888506 | 0.022605017 |
| AC006116.7 | 3.081364805 | 0.000456693 |
| AC005786.3 | -0.672758633 | 4.03E-05 |
| AD000671.3 | 1.2818704 | 1.39E-05 |
| AL353997.3 | 8.0389551 | 4.93E-07 |
| AC005944.1 | 0.771654763 | 0.000143552 |
| ZNF571-AS1 | 0.997190204 | 0.002140149 |
| AC008736.1 | 0.9780285 | 0.000995576 |
| CIRBP-AS1 | -0.863913877 | 0.045649238 |
| NFE2L3P1 | -1.164217587 | 0.024913991 |
| ZNF887P | -1.27792286 | 4.85E-05 |

| | | |
|---|---|---|
| ZNF285 | 1.464023274 | 5.35E-07 |
| AC011446.1 | -0.875019285 | 1.08E-05 |
| AC020916.1 | -0.429984818 | 0.003166361 |
| LINC01864 | 1.599149144 | 0.010802036 |
| AC008735.2 | -0.661611448 | 0.015340653 |
| MIR497HG | 0.365692535 | 0.001044436 |
| S1PR2 | -0.549685634 | 0.045084938 |
| AC060766.4 | -1.112207432 | 0.001551543 |
| AC006116.9 | 5.747162723 | 1.40E-09 |
| AC011477.2 | -1.293847998 | 0.008652844 |
| AC006504.5 | 0.406656264 | 0.001532918 |
| AC011477.3 | 3.707535737 | 2.85E-05 |
| AC060780.3 | 1.462717397 | 0.020074054 |
| AC011446.2 | 1.553327779 | 6.09E-10 |
| AC016590.1 | 1.958689453 | 0.028785378 |
| AC011511.5 | 0.378828822 | 0.009445445 |
| AC138430.1 | 1.389834445 | 0.01409563 |
| AC067852.3 | 2.5617971 | 7.87E-07 |
| RPL7L1P5 | 4.407117364 | 0.003311982 |
| AC060766.5 | -0.924334958 | 0.000181545 |
| AC010327.5 | 2.447622039 | 0.001092709 |
| AC125437.1 | -1.726481511 | 0.004580288 |
| AP001357.2 | -1.146146948 | 0.003908744 |
| AC004156.1 | 3.742711542 | 0.023466386 |
| ZNF224 | 0.844714259 | 2.44E-08 |
| AC091551.1 | -1.075471166 | 9.17E-06 |
| AC104365.4 | -2.513020218 | 0.011899307 |
| AC020922.1 | 4.750691692 | 1.70E-06 |
| AC012254.3 | 1.553383518 | 2.08E-06 |
| SINHCAFP2 | -0.591677945 | 6.37E-05 |
| AC060766.7 | -1.012379695 | 6.28E-06 |
| AC009005.1 | 1.33176119 | 0.002713767 |
| EML2-AS1 | 1.026525787 | 0.00046401 |
| AC022098.3 | 3.371555073 | 0.000337559 |
| SMIM22 | 3.806406452 | 3.35E-05 |
| LIN37 | 1.175739872 | 3.27E-06 |
| AC087289.5 | -1.438309817 | 0.000735347 |
| NDUFA7 | 0.392523066 | 0.010811981 |
| MZF1-AS1 | 1.495364173 | 5.80E-08 |
| AL356740.1 | 4.051123957 | 0.004061222 |
| AC022144.1 | 3.123986323 | 3.58E-06 |
| AC024075.1 | -1.11043895 | 0.026086236 |
| AC008750.3 | 6.773740543 | 3.21E-05 |
| AC139769.2 | 2.512349221 | 0.000133064 |
| AC010328.1 | 1.558187188 | 0.01350549 |
| PTOV1-AS1 | 0.244690381 | 0.011452686 |
| AC005253.1 | 0.321326158 | 0.035354697 |
| NBPF12 | 0.385308047 | 0.000249616 |

| | | |
|---|---|---|
| NAPA-AS1 | 1.273880637 | 0.026335953 |
| AC006539.2 | -2.570312808 | 0.000534444 |
| AC123912.1 | 3.967656648 | 5.33E-05 |
| AC008982.1 | 0.410751766 | 2.76E-05 |
| AC008764.2 | -1.148652314 | 0.028549673 |
| AC010615.2 | -0.370002718 | 0.003591618 |
| AC026304.1 | 2.373671213 | 8.89E-05 |
| AC010336.4 | -0.566867765 | 0.044138554 |
| AC004076.1 | 0.856331301 | 0.012037701 |
| SMIM17 | 4.527188441 | 0.006069312 |
| AC002985.1 | 0.569992717 | 0.018140326 |
| AC005261.2 | -0.240953859 | 0.049536167 |
| AC137932.3 | -0.41624743 | 0.012070906 |
| EEF1A1P7 | 0.532586901 | 0.00426603 |
| ARL14EPL | 0.899428338 | 0.027195012 |
| AC090004.1 | 0.390000639 | 0.019920012 |
| AL589765.5 | 1.624671613 | 0.016406414 |
| LINC02560 | 2.974828637 | 0.000826597 |
| AC008764.3 | -1.266066697 | 1.14E-05 |
| LRRC2-AS1 | 4.322375457 | 0.00127783 |
| FAM156A | -0.57632776 | 0.003925745 |
| AC092279.2 | -0.520685943 | 0.005188919 |
| FENDRR | 7.635397867 | 3.22E-06 |
| AC132192.2 | 1.550998273 | 3.38E-05 |
| SSX2B | -0.461745001 | 0.00195459 |
| AC008403.1 | 0.991513244 | 0.000657877 |
| AC026202.3 | 0.763145637 | 0.01247256 |
| AC020915.3 | -0.603275869 | 0.004780578 |
| VN1R83P | -2.812348802 | 0.000157197 |
| AC008403.2 | 0.538164464 | 0.027184939 |
| AC003002.2 | -1.509655546 | 0.001101753 |
| AC012313.4 | 1.350894161 | 0.001435929 |
| AC011815.1 | 1.092162082 | 0.001016988 |
| AL031282.2 | 0.42017673 | 0.000299643 |
| AC011466.1 | 2.344989257 | 2.53E-05 |
| MAGEA2 | -0.56786765 | 0.000747499 |
| AP003680.1 | 1.539604002 | 0.019943705 |
| AC006486.1 | -0.541530153 | 0.001485836 |
| AC005759.1 | 6.725229658 | 3.55E-05 |
| CTAG1A | 0.612742345 | 8.11E-06 |
| LINC00664 | 1.741158655 | 5.94E-08 |
| AC011495.3 | 1.353783583 | 0.023505492 |
| AC010643.1 | 3.434453845 | 1.98E-07 |
| MTCO3P39 | 3.105895915 | 1.34E-05 |
| RPL7P50 | 2.579939226 | 2.08E-06 |
| AC008764.4 | -0.471396128 | 0.029623709 |
| AC027307.3 | 1.278718946 | 0.012524993 |
| AC004264.1 | -1.584498782 | 0.003160889 |

| | | |
|---|---|---|
| AL118506.1 | 1.071719565 | 3.09E-05 |
| AC008878.3 | -0.777006277 | 3.73E-07 |
| ESPNP | -1.365770707 | 0.011607947 |
| AC008758.5 | 1.588345802 | 0.000513629 |
| A1BG-AS1 | 0.387802076 | 0.001967321 |
| AC012313.5 | 1.735716777 | 2.62E-06 |
| CSAG3 | 0.447496307 | 0.000504622 |
| CT45A1 | -0.136669988 | 0.040122487 |
| CKS1BP3 | 0.48136534 | 2.13E-05 |
| MIA-RAB4B | 0.384719492 | 2.67E-05 |
| MAN1B1-DT | 2.536260468 | 0.000417904 |
| AC092070.2 | 1.486442378 | 0.010600817 |
| AC003006.1 | 0.530551586 | 0.036487082 |
| MTRNR2L12 | 0.485182763 | 0.018997972 |
| AC016629.1 | 2.844581112 | 3.14E-05 |
| AC012313.6 | 3.394592779 | 4.27E-05 |
| CALR3 | 7.998245314 | 6.01E-07 |
| AC009955.4 | 1.072665703 | 0.029165744 |
| AC007842.1 | 0.934148374 | 7.67E-06 |
| AC010328.3 | 2.053556986 | 0.00203767 |
| AC010636.2 | 1.352839142 | 0.004032809 |
| TRABD2B | -0.901298625 | 0.016534338 |
| AL137002.1 | 2.679187923 | 6.27E-05 |
| AC011452.2 | -1.686832971 | 5.08E-08 |
| AC010422.3 | 0.540709695 | 0.000903444 |
| AC008894.2 | -0.637342993 | 2.50E-07 |
| AC011445.2 | 3.704952891 | 3.77E-05 |
| AC011503.1 | 1.355285906 | 0.000272115 |
| IKBKG | -0.284273998 | 0.01379528 |
| ZNF587B | -1.013093358 | 9.31E-08 |
| PTOV1-AS2 | -1.146669399 | 0.002010073 |
| AC008764.6 | 0.790502852 | 0.006331088 |
| AC008750.8 | 0.674471025 | 0.004123649 |
| SPIB | 4.015170276 | 1.11E-05 |
| LINC01224 | -0.7666921 | 9.00E-05 |
| AC011503.3 | 1.349236724 | 0.001710435 |
| AC024075.3 | 0.590213544 | 0.034809113 |
| AC010618.3 | 0.445380623 | 0.013054399 |
| AC012313.8 | 1.01540284 | 0.014329621 |
| AC010319.4 | -1.315027337 | 0.02614067 |
| ERVK9-11 | 3.132753911 | 5.41E-09 |
| AC003973.1 | 5.600837551 | 4.07E-05 |
| AC024257.3 | 1.026832107 | 0.039390926 |
| AC003002.3 | 1.300388281 | 0.00016182 |
| AC093677.2 | 0.820278411 | 0.029785802 |
| AP001350.1 | -1.329385238 | 0.000142182 |
| CT45A10 | -0.419240285 | 0.000310562 |
| AC010422.5 | 0.273596582 | 0.032167701 |

| | | |
|---|---|---|
| AC005523.2 | -0.420707777 | 7.75E-06 |
| RPARP-AS1 | 1.321378073 | 1.71E-06 |
| NOP53-AS1 | 1.733687278 | 0.01259502 |
| AC008982.2 | -1.529699025 | 0.003203166 |
| AC010422.6 | 1.587760673 | 1.65E-10 |
| NBPF9 | -0.142498745 | 0.03568108 |
| AL691432.1 | 0.801634522 | 0.010208631 |
| AC008758.6 | 1.356437371 | 0.029870996 |
| ZIM2-AS1 | 5.620351932 | 2.54E-08 |
| AC010336.6 | 1.267856548 | 0.018031621 |
| KCNQ1OT1 | -1.123104664 | 7.75E-07 |
| AL513523.13 | -0.830970158 | 0.000685255 |
| AC022150.4 | -1.904483518 | 1.20E-08 |
| IPO5P1 | -1.773943983 | 5.34E-07 |
| EGLN2 | -0.409114465 | 3.08E-05 |
| AC112491.1 | 0.314346449 | 0.004305857 |
| ARHGAP19-SLIT1 | -1.060109064 | 5.43E-05 |
| SNHG8 | 0.815634702 | 2.03E-08 |
| COMMD3-BMI1 | -0.559714728 | 3.24E-05 |
| RMRP | 1.257335533 | 0.00072886 |
| AC234772.2 | -1.706940466 | 0.003590809 |
| AL606834.1 | -1.321434237 | 0.000792973 |
| AL049840.4 | -1.318139398 | 0.000630617 |
| Z98884.2 | -1.229247117 | 0.00531149 |
| DDIT4-AS1 | 0.712792814 | 2.20E-07 |
| AC004817.3 | 3.560873415 | 4.54E-06 |
| PCF11-AS1 | 0.657839932 | 0.004578714 |
| AL049840.5 | -1.462009455 | 0.000572678 |
| AC090181.2 | -1.202465652 | 0.016224409 |
| AL117336.1 | 3.482360806 | 1.27E-07 |
| FMC1-LUC7L2 | -1.220496068 | 6.63E-09 |
| AL049840.6 | -1.279236827 | 1.76E-05 |
| SPACA6P-AS | 3.233569214 | 6.01E-07 |
| AC010359.2 | 2.88943426 | 0.033729484 |
| AC006064.4 | 0.194687713 | 7.01E-05 |
| AC010969.2 | -1.785489082 | 1.80E-05 |
| AC018809.2 | -0.895780658 | 0.001165049 |
| AC146944.3 | -0.703322803 | 0.033729484 |
| AC010531.6 | 2.923844522 | 0.00023382 |
| AL137247.2 | 1.9019051 | 0.001109616 |
| ZNF559-ZNF177 | 0.624984711 | 0.002386435 |
| AC232271.1 | -0.968052991 | 0.002853264 |
| AC009108.3 | 3.10254148 | 1.67E-07 |
| AC026691.1 | 3.111256429 | 1.69E-06 |
| BMS1P7 | 0.721479774 | 0.043013589 |
| AL020997.3 | 2.324688123 | 0.000351511 |
| AC025165.5 | 1.685661268 | 0.000976687 |
| AC127502.2 | -1.870536671 | 5.22E-07 |

| | | |
|---|---|---|
| AC090589.3 | -3.6683954 | 0.016798292 |
| MIR222HG | -0.430928456 | 2.06E-05 |
| GAS5-AS1 | 1.294865263 | 0.000491914 |
| AC012065.4 | 3.820809559 | 0.000115191 |
| AC027020.2 | 1.513655331 | 0.000344659 |
| MICOS10-NBL1 | -0.186156824 | 0.014572414 |
| AC068620.2 | 0.766348906 | 0.02076386 |
| AL591806.4 | 0.359299899 | 0.033223757 |
| NCBP2AS2 | -0.698554608 | 0.000230919 |
| AC104109.2 | 3.305136334 | 3.60E-07 |
| AP002840.2 | 3.133374039 | 0.000708856 |
| BIVM-ERCC5 | -0.261332802 | 0.00199931 |
| AC104695.3 | 7.269142197 | 7.40E-06 |
| AC079880.2 | -1.106065412 | 0.000200954 |
| NBPF8 | -0.353040654 | 5.70E-05 |
| AC015849.1 | 6.619362672 | 4.77E-05 |
| NDUFB8P2 | 1.131962671 | 0.010793576 |
| H4C15 | 4.039082842 | 4.51E-09 |
| AL121758.1 | -0.80008989 | 4.70E-09 |
| BORCS7-ASMT | 0.954527258 | 0.008183946 |
| SMC2-AS1 | 3.512764074 | 7.20E-05 |
| POC1B-AS1 | 1.177899679 | 0.000858507 |
| HMGN3-AS1 | 0.863231917 | 0.001143865 |
| PFN1P2 | 0.912746078 | 0.001753804 |
| CAHM | 1.770183063 | 0.00102283 |
| AC099343.3 | 3.527471183 | 0.000101435 |
| AC026367.1 | 3.04535235 | 0.000792664 |
| YTHDF2P1 | -0.386977152 | 0.011255207 |
| AL391422.4 | 1.843487991 | 5.24E-07 |
| PEBP1P2 | 0.351489779 | 0.029474073 |
| AL158163.1 | 0.811673401 | 0.011940795 |
| AL353622.1 | 2.514334786 | 5.72E-06 |
| TSIX | -1.067460498 | 6.49E-11 |
| TAF15 | -0.166802455 | 0.004998759 |
| HNRNPA1P67 | 1.942415656 | 0.003343407 |
| AC095055.1 | 7.320318287 | 6.93E-11 |
| BUD13P1 | 2.330438597 | 0.002631107 |
| AC005034.4 | -1.488075522 | 1.64E-07 |
| PRMT1P1 | 0.758564894 | 0.008913353 |
| AL035413.1 | 0.6332793 | 0.021182028 |
| AP000436.1 | 4.275575408 | 0.001393239 |
| RPS10-NUDT3 | 0.26220401 | 0.000682686 |
| LINC00221 | -0.326785945 | 0.007549075 |
| AC016727.1 | -0.554032846 | 0.000384234 |
| SRGAP2D | 0.783144373 | 5.31E-06 |
| H4C14 | 4.046427286 | 1.53E-13 |
| RASL10B | 1.507982513 | 0.006719743 |
| AL049844.2 | -1.77309714 | 0.00016401 |

| | | |
|---|---|---|
| TAS2R6P | 1.63622834 | 0.024994135 |
| AC008013.4 | 3.066579538 | 0.000102459 |
| CT45A9 | -0.493257307 | 0.024722817 |
| LPP-AS2 | 1.247085851 | 7.42E-05 |
| AC016355.1 | 2.910791245 | 3.22E-06 |
| AC015849.5 | 2.967678802 | 5.45E-06 |
| AC091982.2 | 7.4367906 | 4.07E-06 |
| AL589674.1 | -0.692490749 | 0.000259332 |
| HMGB1P51 | 4.376274474 | 0.001092216 |
| LRRC37A9P | 3.71934745 | 5.40E-07 |
| AC013244.1 | 2.853199971 | 0.001205012 |
| TLCD4-RWDD3 | 0.282740723 | 0.0236855 |
| NUDT4P2 | 0.379377687 | 0.000391504 |
| AC018647.2 | 1.688063391 | 3.62E-10 |
| RPL23AP88 | 1.849885064 | 0.012758741 |
| AC016831.4 | 5.403936572 | 0.000623383 |
| AC240274.1 | 1.011450897 | 5.30E-08 |
| TMCC1-AS1 | 2.27325313 | 2.90E-08 |
| SRXN1 | -0.803493958 | 5.05E-09 |
| AC010201.2 | 1.454491999 | 0.020578383 |
| AL035417.2 | 4.30825678 | 0.00103475 |
| AL117336.2 | 1.750447918 | 3.08E-06 |
| HTATSF1P2 | 0.761015693 | 0.001691306 |
| NBPF19 | -0.205660594 | 0.000737394 |
| AL445228.3 | 2.582436299 | 0.001835307 |
| NBPF10 | -0.154663795 | 0.033804958 |
| CT45A2 | -0.380793714 | 0.017587614 |
| AC011477.5 | 5.60036169 | 3.29E-05 |
| BNIP3P17 | -1.750410226 | 6.21E-05 |
| ARF4P1 | -0.608945551 | 0.000924784 |
| Z83843.1 | -1.283838826 | 1.43E-05 |
| LINC02427 | 10.64709035 | 1.43E-10 |
| BNIP3P11 | 0.658055953 | 0.009942764 |
| AC108463.3 | -1.345261263 | 0.016094848 |
| LIX1L | -0.879431655 | 1.13E-08 |
| ATP2B1-AS1 | 2.610184612 | 4.94E-07 |
| AC112220.2 | -2.200189246 | 0.000188697 |
| AC099343.4 | 2.252734834 | 4.01E-08 |
| AC097359.3 | 4.960036354 | 0.000239407 |
| AC010998.2 | 1.260422039 | 0.009297424 |
| AC233992.2 | 1.365502737 | 0.000467215 |
| AL390208.1 | 1.434766838 | 0.000144738 |
| AC008608.2 | 3.521921361 | 0.005942988 |
| AL355802.3 | -0.984688203 | 0.001200607 |
| AC080013.3 | 1.191947394 | 0.046900179 |
| AL118558.3 | 2.729662323 | 1.91E-05 |
| AC026740.1 | -2.688153972 | 6.54E-05 |
| AC008875.1 | 3.050469725 | 0.024986121 |

| | | |
|---|---|---|
| AL589666.1 | -0.631879664 | 0.000568219 |
| AC011337.1 | 1.912472899 | 4.33E-05 |
| AL603832.3 | 0.65000763 | 8.77E-05 |
| AC022400.5 | 1.492051642 | 2.40E-05 |
| AC008937.3 | 1.027908072 | 0.000341938 |
| AC087501.4 | -2.135290026 | 0.00551693 |
| AL162258.1 | 0.491595126 | 0.00016182 |
| AC073195.2 | 2.107642646 | 0.006185029 |
| LINC01215 | 8.194097283 | 2.82E-07 |
| AC104118.1 | -0.941688565 | 0.007581358 |
| AGAP11 | 1.970178803 | 3.76E-05 |
| AL109811.2 | 2.720351181 | 1.18E-07 |
| AL035530.2 | 4.936023339 | 0.000162383 |
| AC120053.1 | 3.110192802 | 1.40E-07 |
| AC012467.2 | -0.47108124 | 0.006562771 |
| AL139274.2 | 2.506917517 | 0.000117482 |
| AC064807.4 | 6.917165493 | 2.09E-05 |
| AL162258.2 | 1.128748399 | 2.04E-05 |
| ANKRD34A | 0.913297048 | 0.002934564 |
| AL136984.1 | 4.222507834 | 0.001592965 |
| AP002907.1 | 1.329915406 | 0.001017784 |
| AL445647.1 | 1.209326509 | 1.38E-05 |
| GTF2H5 | 0.598878396 | 4.05E-05 |
| AC007390.2 | -0.456561128 | 0.000173642 |
| AL365181.2 | 1.30327656 | 2.82E-07 |
| AC005618.1 | 3.347666115 | 4.16E-09 |
| AL139423.1 | 0.705429249 | 0.022236376 |
| AC004233.2 | 2.756041384 | 0.00021345 |
| AC080013.4 | -1.236464304 | 0.013102292 |
| AC087623.2 | 0.951489318 | 0.005558729 |
| AL024498.1 | 2.050124345 | 0.03269266 |
| AC244517.1 | 2.078244915 | 0.005087354 |
| AC008906.2 | 2.345486763 | 0.002878478 |
| AC008966.2 | 4.288635655 | 4.40E-07 |
| AP006545.2 | 5.29036887 | 4.43E-05 |
| AC022400.6 | 1.278897231 | 4.20E-05 |
| AL359643.3 | 1.003538714 | 0.001431494 |
| ARF4-AS1 | 1.189522546 | 0.019362661 |
| AL365330.1 | -1.394632128 | 0.000579974 |
| AC138696.2 | 2.038908913 | 0.020453664 |
| U47924.2 | 0.3056104 | 0.001558932 |
| AC005041.3 | 0.929581341 | 0.007010229 |
| AP003392.5 | 1.653638485 | 0.0101212 |
| AL356512.1 | 1.376980791 | 0.001049849 |
| H2AC19 | 0.559762933 | 0.011648221 |
| AL136304.1 | 1.422836627 | 0.015820642 |
| AC004908.1 | 0.999088067 | 0.016637333 |
| AL138831.2 | 4.220143675 | 0.001626747 |

| | | |
|---|---|---|
| AC044849.1 | 0.624142732 | 0.003609116 |
| AC021242.3 | 0.583197076 | 0.02697956 |
| AL138724.2 | -0.902748656 | 5.36E-05 |
| AC092687.3 | 3.335391505 | 9.88E-05 |
| AL031963.3 | 1.616842403 | 0.002841638 |
| AL445309.1 | 4.004297774 | 4.33E-05 |
| NUDT3 | -0.127737861 | 0.031189689 |
| KMT2B | -0.572817704 | 3.81E-07 |
| AC011816.2 | 7.499153443 | 3.73E-06 |
| AC093297.2 | -1.155287375 | 0.000108163 |
| AL137003.1 | -1.083346962 | 0.044618449 |
| AC107952.2 | 3.25528354 | 0.00020059 |
| AC074032.1 | 1.471665346 | 0.003789597 |
| POM121C | -0.861635596 | 1.70E-09 |
| CD24 | 2.209973999 | 4.01E-06 |
| AL365181.3 | -0.2863352 | 0.000397899 |
| AC022384.1 | -0.989655306 | 7.02E-09 |
| FAM47E-STBD1 | -0.968207005 | 6.64E-06 |
| U91328.1 | 1.919456442 | 3.76E-05 |
| AL024507.3 | 0.612887724 | 0.007244318 |
| AL020996.3 | -0.532983816 | 0.043744491 |
| AC104958.2 | 1.409546281 | 0.000463463 |
| AL645608.7 | 3.212109924 | 0.000490229 |
| LINC01023 | 2.005577798 | 0.001388744 |
| AC099522.2 | 2.641720764 | 2.65E-08 |
| AC017048.3 | 6.56273294 | 5.59E-05 |
| GTF2IP13 | 1.87646733 | 0.000334002 |
| AL596325.2 | -1.775008375 | 0.002322205 |
| AC139887.4 | 2.965678381 | 0.000221588 |
| AC016394.3 | -0.849944532 | 0.001096683 |
| AC007308.1 | -1.804516815 | 0.010546815 |
| AC010655.3 | 1.64169014 | 0.000190999 |
| ZNF595 | 0.269988491 | 0.001737043 |
| AC073073.2 | 0.694497159 | 0.00556105 |
| AC015982.2 | 0.906984785 | 0.008804809 |
| MAGI1-IT1 | 3.972442636 | 1.99E-05 |
| AC026464.6 | -0.522294319 | 1.14E-05 |
| AC067750.1 | 1.797799554 | 8.21E-06 |
| AC006027.1 | -2.204767079 | 5.88E-06 |
| GTF2IP20 | 1.627736323 | 8.33E-08 |
| AC005020.2 | 0.907476995 | 0.020950864 |
| AL358472.2 | 0.449264358 | 0.04573052 |
| AP000317.2 | -0.669770453 | 0.008752031 |
| AL590560.3 | 3.442220443 | 0.000325624 |
| AL021707.6 | 1.394051114 | 0.000997375 |
| PCDHB16 | -0.514432412 | 0.000111854 |
| AC124016.2 | 2.022773319 | 0.000675479 |
| AC006333.1 | 1.145522327 | 0.000276715 |

| | | |
|---|---|---|
| AC010997.4 | 1.833135554 | 0.007250089 |
| GAS6-DT | 2.6024856 | 5.14E-05 |
| AL359091.3 | 1.826741863 | 0.002483026 |
| AC010913.1 | 1.042611353 | 0.021498843 |
| AL121658.1 | 1.342196287 | 0.009955122 |
| AL022322.1 | 1.564604415 | 0.001061887 |
| ADIRF-AS1 | 3.456566559 | 6.29E-12 |
| AC007881.3 | 3.947956258 | 0.004741847 |
| AC005323.2 | 2.088572628 | 0.00028515 |
| STAG3L5P-PVRIG2P-PILRB | -0.452851883 | 1.77E-05 |
| AC083798.2 | -0.922323785 | 2.36E-08 |
| AC093726.1 | -1.225443726 | 0.010652342 |
| AC115284.1 | 0.743519993 | 0.000382344 |
| AL596094.1 | 2.707068346 | 0.000299254 |
| AC004854.2 | -0.869181417 | 0.002294438 |
| AC005696.2 | -1.157597011 | 3.87E-05 |
| AC104109.3 | -0.54283359 | 0.000488439 |
| AC019131.2 | 2.81116684 | 0.000278937 |
| AC245060.4 | -0.655583219 | 0.00017486 |
| AC015712.6 | 2.154673089 | 2.47E-07 |
| AC098850.4 | 1.036148799 | 0.00047568 |
| U62317.2 | -0.637424525 | 0.001773345 |
| AC073610.2 | 0.213953663 | 0.018195161 |
| AL022328.1 | 1.035979759 | 0.049891106 |
| AL391834.1 | 3.061814952 | 1.98E-06 |
| AC069544.1 | 1.238466489 | 0.019430326 |
| AL159169.2 | -2.567928159 | 0.001276444 |
| AP000525.1 | -1.122853884 | 0.004845259 |
| AC113189.4 | 1.24364587 | 2.23E-05 |
| LINC01578 | 0.374177486 | 0.000291074 |
| AL355987.3 | 1.452903893 | 0.000482934 |
| AL109827.1 | 0.35646585 | 0.009448424 |
| ATP6V1FNB | 1.480425872 | 2.04E-06 |
| AL353708.3 | 0.993742937 | 0.008440017 |
| AL122035.2 | -1.347347438 | 0.000272411 |
| AC009237.14 | 0.289805672 | 0.03234497 |
| AC022400.7 | 0.270265719 | 0.012540553 |
| AC005832.4 | -0.284159299 | 0.049174824 |
| AL391121.1 | -0.688445165 | 0.017062071 |
| AC083862.2 | -0.61743972 | 0.002494991 |
| AC079834.2 | 5.058539369 | 0.000341354 |
| AC093799.1 | 4.273293444 | 0.001390271 |
| AP000553.2 | 1.807924936 | 0.022016838 |
| RBAK-RBAKDN | 0.72735082 | 0.034878853 |
| AC024243.1 | 2.008454297 | 0.002754403 |
| AL365181.4 | 7.740700415 | 1.91E-06 |
| AC012360.3 | 0.575613114 | 0.000374735 |
| AL355388.1 | 2.32240727 | 1.18E-05 |

| | | |
|---|---|---|
| ARL2-SNX15 | 0.365314475 | 0.000343625 |
| AC010864.1 | 3.381532041 | 6.53E-08 |
| AL360270.3 | 0.731407711 | 0.006534318 |
| AC008124.1 | -0.591772909 | 0.000150259 |
| AC009690.3 | 0.901633187 | 0.02162473 |
| AL358472.3 | 2.276783623 | 0.000174238 |
| DGCR9 | 1.736404746 | 0.000524741 |
| LINC02035 | 0.949346679 | 0.000124633 |
| AC007684.2 | 4.072672752 | 3.22E-05 |
| AL390726.4 | -2.791349835 | 5.97E-05 |
| AL365203.2 | 1.704425152 | 4.09E-06 |
| ATF4P2 | 3.079100855 | 0.000429684 |
| AC012531.2 | -1.117455909 | 0.000268746 |
| AL121845.2 | -1.502940744 | 0.000164589 |
| AC012531.3 | -1.271846206 | 0.002580972 |
| AC005046.2 | -1.554393116 | 7.02E-06 |
| AL354694.1 | 3.827250718 | 0.012418709 |
| AL359921.2 | 1.63672963 | 0.000109476 |
| CDC37L1-DT | 2.875998618 | 7.00E-07 |
| AL449106.1 | 1.815247487 | 0.00091595 |
| AC017083.1 | 1.670914516 | 0.023167244 |
| GRIN2B | 2.127783375 | 0.00399611 |
| AC009309.2 | 2.480742002 | 4.42E-06 |
| AC007378.1 | -0.806471811 | 0.036331667 |
| AC019129.2 | 3.448490824 | 0.000198502 |
| AC144652.1 | 4.895709844 | 2.08E-11 |
| NBPF26 | -0.310666936 | 0.000239459 |
| LINC02604 | 1.758165741 | 5.92E-08 |
| AL355512.1 | 4.371865846 | 8.86E-06 |
| BX537318.1 | 1.3387192 | 0.043874414 |
| LINC00653 | 1.423898725 | 8.97E-06 |
| AL138963.4 | 0.53176684 | 2.37E-06 |
| AC073957.3 | 1.937610085 | 1.92E-06 |
| AL121845.3 | -0.971253051 | 1.28E-07 |
| AL359736.1 | -0.431715041 | 0.001039818 |
| AC010655.4 | -0.302349877 | 0.00573029 |
| AC006946.2 | 0.732218271 | 0.015248514 |
| H3-2 | 1.132992325 | 0.002520503 |
| AC002059.1 | 2.685515779 | 0.000867068 |
| AC008695.1 | -0.936379965 | 2.80E-07 |
| AL355816.2 | -0.851805188 | 0.00040978 |
| AL391834.2 | 3.510445261 | 1.01E-09 |
| AC102953.2 | -0.815277821 | 0.001186894 |
| AC097376.3 | 2.310269166 | 1.82E-07 |
| AL022328.4 | 0.8079459 | 0.048592111 |
| AL049839.2 | 0.703099695 | 1.06E-06 |
| ZBTB8B | 0.655018956 | 0.001887228 |
| AC017083.2 | -1.014746799 | 6.00E-07 |

| | | |
|---|---|---|
| AP000350.6 | 1.246525947 | 0.001310198 |
| AC009237.15 | 2.423532348 | 0.000232596 |
| AC024560.3 | 4.896944721 | 0.000266684 |
| AL121749.1 | 2.972586116 | 0.001090565 |
| AC058791.1 | 5.640584968 | 2.27E-09 |
| AC099329.2 | 8.210165997 | 3.61E-07 |
| AC078846.1 | -1.945709059 | 4.59E-08 |
| TM4SF19-TCTEX1D2 | 0.4989843 | 0.03281378 |
| PAXIP1-AS1 | 2.439548188 | 1.08E-08 |
| AL451074.6 | 2.479094599 | 0.00145353 |
| AC096586.2 | -1.338287024 | 0.001792204 |
| AC069222.1 | 2.581996749 | 1.72E-06 |
| AL356488.3 | 1.344858147 | 0.001370035 |
| AC083880.1 | 1.793165326 | 0.002799512 |
| AC017083.3 | -0.750816277 | 6.14E-06 |
| AL136982.7 | 2.680031568 | 0.00117581 |
| AC004877.1 | 4.923809112 | 4.67E-10 |
| AC004951.4 | -0.408457416 | 0.034652346 |
| AC004080.6 | -1.011629323 | 0.001171924 |
| AC006946.3 | 2.693771989 | 0.00345512 |
| AL645608.8 | 1.804416231 | 8.15E-05 |
| AC006480.3 | 2.515893377 | 1.57E-05 |
| AC093788.1 | -2.06613498 | 0.002596784 |
| AL139124.1 | 2.522290198 | 0.000607262 |
| AL133215.3 | 3.986774737 | 0.002678297 |
| AC096992.2 | 3.237507644 | 3.08E-06 |
| AC011841.1 | 1.231537103 | 0.000279725 |
| H4C12 | 2.617243634 | 0.000103618 |
| CWC25 | -0.437298769 | 0.000199585 |
| AC131009.3 | -1.039631587 | 0.00609161 |
| CDHR5 | 1.639305323 | 0.02233451 |
| LENG9 | 0.908421604 | 0.018158249 |
| AC009283.1 | 1.528423417 | 0.00061736 |
| SMIM11B | 0.662575635 | 0.034099618 |
| EEF1D | 0.202611119 | 0.001296306 |
| AL355377.1 | 7.043972366 | 1.39E-05 |
| AL731571.1 | 0.376222502 | 0.036558835 |
| EPOP | 0.564229191 | 3.67E-05 |
| SRCIN1 | 0.845199201 | 0.000984892 |
| ZNHIT3 | 0.322910194 | 0.018041561 |
| AL139260.2 | 0.346367065 | 0.003565106 |
| AC011295.1 | 0.339797513 | 3.11E-07 |
| AC100757.1 | -1.382492078 | 7.84E-05 |
| CT45A7 | -0.638243199 | 0.001107239 |
| AC091271.1 | 1.769828424 | 6.58E-05 |
| LHX1 | -1.573261789 | 1.77E-08 |
| AC092670.1 | 0.322274994 | 0.001744657 |
| AL139089.1 | 0.83474204 | 0.048688521 |

| | | |
|---|---|---|
| AC106782.4 | 4.265533673 | 3.04E-05 |
| AC007686.3 | -1.325993786 | 5.88E-05 |
| AL592188.1 | 1.520229018 | 6.32E-09 |
| AC005258.1 | 0.488043826 | 0.012317376 |
| CYFIP1 | -0.601809 | 2.36E-07 |
| RNVU1-29 | 1.630482595 | 0.00013793 |
| ARHGAP23 | -0.973591309 | 1.52E-08 |
| AL136040.1 | 2.165003209 | 1.38E-05 |
| H2BC8 | 3.931258074 | 4.05E-08 |
| USP27X | 0.571399525 | 0.002071491 |
| AL096828.3 | 4.119385182 | 0.004352664 |
| AL133227.1 | 1.654289505 | 0.017120014 |
| AF254983.1 | -0.790251585 | 0.023182276 |
| TAF9 | 0.28777002 | 0.000374426 |
| AC139494.2 | -1.743118075 | 0.005425309 |
| AL731566.2 | -1.517792064 | 0.000669345 |
| AC211476.5 | 1.431390843 | 0.020790567 |
| NOL12 | -0.856598754 | 6.25E-05 |
| AC012313.9 | 2.34733458 | 0.000528002 |
| AL031667.3 | 1.434091991 | 0.019280907 |
| H3C8 | 1.299508138 | 0.027405843 |
| AL049794.1 | 3.047719468 | 4.74E-06 |
| RN7SL2 | 0.484321026 | 0.049418918 |
| AC239804.2 | 6.873317406 | 2.68E-05 |
| AC024909.1 | -0.530791593 | 3.64E-06 |
| FAM27E3 | 1.140264621 | 0.007937454 |
| INO80B-WBP1 | 0.789304378 | 7.45E-08 |
| AC133551.1 | 2.971400376 | 0.000693724 |
| NATD1 | 0.663010817 | 8.63E-06 |
| AC011815.2 | 8.297991996 | 3.02E-07 |
| RNVU1-27 | 8.495507157 | 1.41E-07 |
| SOCS7 | -1.711427174 | 6.24E-11 |
| AC009163.7 | 0.793520674 | 0.032634675 |
| SNORA73A | 1.930542824 | 0.000298803 |
| ADRA2B | 6.790311635 | 0.000156226 |
| LINC02334 | 2.353072334 | 7.41E-05 |
| AL355974.1 | 2.715265715 | 0.000598605 |
| CU633967.1 | -0.640283926 | 5.05E-05 |
| AC032011.1 | 3.314854405 | 2.09E-05 |
| ZNF658 | 2.289213419 | 4.96E-11 |
| AC103691.1 | 0.583937749 | 0.016378891 |
| TPTE | -0.965371191 | 0.003129645 |
| AL121772.1 | 1.85786019 | 6.35E-07 |
| AL162390.2 | 3.91383793 | 0.002758676 |
| AC245060.5 | -0.698773285 | 0.015089804 |
| SEC22B2P | -1.05015173 | 0.009946272 |
| AC114271.1 | 0.461580787 | 0.000618755 |
| RNVU1-25 | 7.121845684 | 1.19E-05 |

| | | |
|---|---|---|
| AC142525.1 | -0.708247108 | 9.61E-05 |
| AC092119.2 | 1.497140429 | 0.004316393 |
| YWHAE | -0.143646682 | 0.022680744 |
| RCC1L | -0.437552105 | 2.86E-05 |
| CU639417.1 | 0.834507104 | 0.018442225 |
| AC080038.2 | 1.345740272 | 0.018620873 |
| LILRB3 | 2.86646453 | 3.38E-05 |
| PCCA-DT | -0.833133535 | 0.000491679 |
| AL669942.1 | -1.784555209 | 0.006345236 |
| AC073052.2 | -0.993107983 | 5.60E-06 |
| AC125257.2 | 1.424147952 | 0.001650673 |
| AC106782.5 | 2.311495338 | 0.000103834 |
| AC106886.3 | 2.745399625 | 5.58E-05 |
| MYO5BP1 | 2.590425509 | 1.89E-05 |
| AC005332.4 | -1.502098303 | 0.011846792 |
| ARHGEF10 | -0.68975362 | 1.74E-05 |
| ARHGAP11B | -1.326071541 | 9.04E-06 |
| H3C6 | 3.326037926 | 1.40E-06 |
| AC120498.9 | -1.4067678 | 0.020254832 |
| AC016747.4 | 1.095656518 | 1.55E-05 |
| Z82195.2 | 8.13661659 | 3.76E-07 |
| NPHP3-ACAD11 | 0.764047415 | 1.63E-06 |
| MAFIP | -0.579849509 | 0.014801164 |
| MRM1 | -1.528631886 | 1.31E-06 |
| AC009804.1 | 1.940204577 | 0.000518102 |
| AL139384.1 | 2.05996215 | 0.000440683 |
| ZKSCAN2-DT | -0.631956868 | 0.015830719 |
| RN7SL600P | 4.711667608 | 1.70E-09 |
| AC026356.1 | -1.941512341 | 6.18E-05 |
| PTCHD3P1 | -2.214847192 | 3.21E-05 |
| AC092794.1 | 3.089588562 | 0.00082975 |
| ZNF280B | 7.612008947 | 2.21E-06 |
| PRAME | 0.190732315 | 0.011184981 |
| AL121581.1 | 0.846443899 | 1.65E-05 |
| MLLT6 | -0.200454383 | 0.00323364 |
| TP53TG3E | 1.5494566 | 0.003327247 |
| PPP4R3B | -0.755235869 | 1.44E-08 |
| NUDT18 | 1.693741422 | 3.66E-05 |
| TFPT | 1.741856909 | 0.010459027 |
| GGNBP2 | 0.331817584 | 0.000477063 |
| AC025594.2 | 2.773716927 | 3.37E-05 |
| ZNF2 | 0.723758039 | 0.001134546 |
| AC241952.2 | 0.468447029 | 0.009413075 |
| SEC22B3P | -1.144600188 | 0.02949604 |
| AC048341.2 | -0.856482813 | 0.004588247 |
| AL161431.1 | 1.166492647 | 7.66E-10 |
| AL121906.2 | -1.154725337 | 0.038974587 |
| TLCD2 | 0.75027326 | 0.00635049 |

| | | |
|---|---|---|
| CFL1P5 | 1.158313822 | 0.021042349 |
| RNVU1-26 | 3.272473785 | 0.000346634 |
| LINC02340 | 6.837997479 | 1.25E-06 |
| AC004542.3 | 0.704148105 | 0.019537356 |
| AL354950.2 | -1.021753743 | 0.004076758 |
| AL138781.1 | 9.047035413 | 1.43E-08 |
| AL355493.3 | 6.725229658 | 3.55E-05 |
| FGD5P1 | 4.062131196 | 9.92E-06 |
| AC092111.1 | 2.014689928 | 1.68E-05 |
| FCGBP | 4.540805423 | 8.77E-12 |
| DHRS11 | 0.650842161 | 0.00121928 |
| AL391095.4 | 1.230716418 | 0.003539392 |
| AL845472.1 | 2.431146668 | 9.07E-06 |
| AC105020.6 | -1.020995005 | 5.57E-06 |
| FP565260.1 | -0.451647126 | 0.002993618 |
| AC025031.4 | 0.940597022 | 0.045088337 |
| LINC01730 | 1.069105864 | 0.014941671 |
| AC133552.5 | 2.418709736 | 0.00195658 |
| AC100835.2 | 3.458871361 | 8.67E-05 |
| RNVU1-19 | 5.643507869 | 7.70E-09 |
| AC008115.3 | -1.914815818 | 6.75E-06 |
| AC241377.3 | -0.848655857 | 0.004076346 |
| XKR5 | 4.022300912 | 3.89E-05 |
| AC024051.7 | 1.255391605 | 0.034718516 |
| AC139677.2 | 0.787733666 | 0.000631178 |
| AL035461.3 | -1.023081971 | 0.014061646 |
| AATF | -0.448001808 | 7.93E-06 |
| AC090527.4 | 1.446921955 | 0.015018983 |
| H2BC9 | 1.374173587 | 0.031224987 |
| AC008622.2 | -1.312472234 | 2.69E-08 |
| AL049794.2 | 4.220143675 | 0.001626747 |
| AC092747.4 | -2.120991736 | 1.91E-06 |
| AC091982.3 | 2.570477468 | 3.68E-06 |
| RIMBP3 | -0.946773455 | 0.006122042 |
| TUBGCP5 | -0.635438909 | 0.008948321 |
| SMARCB1 | -0.291542006 | 0.038184109 |
| AC009133.4 | -1.032851099 | 3.77E-05 |
| AL139385.1 | -0.832816254 | 0.00372723 |
| U2AF1L5 | -0.560205896 | 0.000256402 |
| DUSP14 | 0.487351529 | 0.00013181 |
| EIF3FP1 | 2.064527554 | 6.77E-06 |
| AL355001.2 | 1.222278238 | 0.003913708 |
| SIK1B | 2.228770255 | 3.42E-11 |
| WDR81 | -0.390973614 | 0.016556914 |
| UHRF1 | -1.292007797 | 2.55E-10 |
| ORAI1 | 0.21305936 | 0.009178299 |
| NAIP | -1.437640479 | 3.35E-05 |
| CU633904.2 | -0.79945976 | 1.38E-07 |

| | | |
|---|---|---|
| AC008073.3 | 5.835575536 | 3.30E-05 |
| AC037198.1 | -1.173771925 | 7.79E-05 |
| AC026356.2 | -1.83708667 | 0.000741399 |
| FUT8-AS1 | 1.21537172 | 0.014014062 |
| SLC22A18 | 0.857275259 | 0.028659142 |
| AC009118.2 | -1.318315991 | 8.27E-05 |
| WHAMMP3 | -1.668768647 | 5.19E-05 |
| RN7SL1 | 1.098503336 | 0.000180419 |
| H4C9 | 1.251637706 | 7.03E-05 |
| ERMARD | 0.835666795 | 0.003233395 |
| AC245014.3 | 5.564355385 | 2.82E-06 |
| RNH1 | 0.331888206 | 0.028826204 |
| TADA2A | -1.423035712 | 1.98E-06 |
| AL442125.1 | -0.762222926 | 0.012160743 |
| AC007786.2 | 5.222618811 | 0.000193151 |
| AC011043.1 | -0.637863174 | 0.002825807 |
| AC024884.2 | 2.629413448 | 0.001693024 |
| ARL17B | -0.756456905 | 0.000999353 |
| FRG1HP | 0.716470264 | 0.000114019 |
| PIP4K2B | -0.54289185 | 3.92E-07 |
| AL157712.1 | 5.819536842 | 5.30E-05 |
| UBE2NL | -1.493098844 | 0.000219113 |
| AC016876.4 | 2.068591941 | 0.003971776 |
| AC004241.3 | 1.554505748 | 1.33E-05 |
| AC025287.2 | 2.126500806 | 0.00039017 |
| AC005393.1 | 3.208844839 | 6.97E-08 |
| AC004076.2 | -1.040107617 | 0.036581463 |
| GPR179 | 6.160817949 | 1.73E-10 |
| AL136962.1 | -1.110143947 | 0.014877012 |
| MYH11 | 1.014464099 | 0.000625198 |
| AC239799.1 | 5.55604631 | 1.97E-09 |
| AL133243.3 | -0.768252576 | 0.009071803 |
| AP001505.1 | -2.203658057 | 5.71E-06 |
| AL353726.1 | -1.154386008 | 0.01702369 |
| HERC2P2 | -0.561301014 | 7.27E-07 |
| AL442067.1 | 2.583528383 | 0.001766569 |
| RAB7B | -0.755812069 | 0.000709975 |
| AP000351.1 | 7.200657999 | 8.95E-06 |
| AC007950.2 | 4.323288774 | 0.002788047 |
| AL161891.1 | -1.079679654 | 7.19E-06 |
| AL136295.6 | 2.793871602 | 0.001924333 |
| RNA5-8SN4 | 1.275425532 | 0.007599786 |
| AC243773.1 | -1.871681608 | 0.004798239 |
| CSPG4P10 | -0.387970591 | 0.006669894 |
| AC142472.1 | 2.082529241 | 7.00E-06 |
| AC105137.2 | 3.147613794 | 0.000615207 |
| AC004079.4 | 1.638928943 | 0.010518426 |
| FAN1 | -0.293991191 | 0.000522653 |

| | | |
|---|---|---|
| AC068234.3 | 7.160525075 | 1.02E-05 |
| AC092117.1 | 2.674928983 | 1.55E-06 |
| AL133216.2 | 1.639871617 | 0.007991891 |
| SERPINF2 | 3.02558933 | 0.000580254 |
| AC016590.3 | 2.002643329 | 0.000494936 |
| AC015922.3 | 0.701563007 | 0.00119565 |
| AC023157.3 | 1.250338659 | 0.006742829 |
| H2AC16 | 3.602376766 | 0.00024635 |
| GSTT4 | 4.373072962 | 0.001054824 |
| AL121772.3 | -1.397516764 | 0.001424711 |
| AL158063.1 | 7.273051107 | 7.59E-06 |
| LENG1 | 1.862302345 | 0.038068701 |
| H4C5 | 2.052818338 | 0.002030846 |
| AL513314.2 | 1.355696054 | 0.016624161 |
| AL590096.1 | 4.20042138 | 1.82E-05 |
| AL031663.3 | 2.665489323 | 0.048308893 |
| GTF2IP1 | -0.426176057 | 1.67E-06 |
| CU634019.2 | -0.748149611 | 4.27E-05 |
| STAG3L2 | -0.907645987 | 3.67E-06 |
| H2AC8 | 3.637553342 | 3.89E-05 |
| FP565260.3 | 1.25679064 | 4.36E-05 |
| AC211476.10 | 1.678931796 | 0.000652016 |
| AL589743.5 | 1.512089472 | 0.013194149 |
| AC073569.3 | 1.353296563 | 0.007910657 |
| LINC00235 | 1.386403788 | 0.000621428 |
| LINC00869 | -0.439922996 | 0.005139671 |
| F8A3 | -0.459977932 | 0.000299175 |
| AL589743.6 | 0.62842422 | 0.000611108 |
| AL139384.2 | 1.874371178 | 0.00572833 |
| PIGW | 0.24450751 | 0.037113671 |
| AC006449.5 | 1.580747599 | 1.05E-05 |
| AC007325.2 | 2.475491399 | 0.000819981 |
| RPPH1 | 0.786677157 | 0.048742284 |
| H2BC7 | 4.411721603 | 1.61E-06 |
| PCGF2 | 0.177507489 | 0.021861069 |
| AC004812.2 | 0.673653857 | 0.015542121 |
| AL109976.1 | 3.937956649 | 1.52E-05 |
| AC084876.1 | -0.816078549 | 0.032374129 |
| AC090246.1 | 3.366833351 | 0.017458214 |
| AC048344.4 | -2.311686818 | 0.0001288 |
| AC010654.1 | 2.480538894 | 0.000233868 |
| AC145212.1 | -0.810086315 | 0.000870892 |
| SEC22B4P | -0.819291325 | 0.039259545 |
| AC069234.5 | 2.24304579 | 0.003648759 |
| MARCKS | -0.639593891 | 4.44E-06 |
| CPEB1 | 0.674391476 | 0.009535114 |
| CEBPB-AS1 | 2.791161341 | 7.53E-08 |
| RN7SL473P | 7.32405282 | 0.001060416 |

| | | |
|---|---|---|
| AP001527.2 | -1.042069366 | 0.005459755 |
| PKD1L3 | 5.233948462 | 0.000227877 |
| AC087392.5 | 1.713808852 | 0.009084653 |
| AC010536.3 | 4.254256585 | 0.001196504 |
| AP000351.2 | 1.541767805 | 0.00248021 |
| AC007996.1 | -3.080397191 | 3.08E-08 |
| DDX52 | -1.008630992 | 1.46E-08 |
| AP000944.2 | 3.100221219 | 2.86E-06 |
| AC005363.2 | 4.480524893 | 0.001582454 |
| WNT3 | -1.598176232 | 0.001964978 |
| AC007906.2 | 3.245324213 | 1.77E-06 |
| TIMM22 | -0.382998263 | 0.003266744 |
| AL139407.1 | 2.053624953 | 7.83E-05 |
| AL121583.1 | 2.758756993 | 0.000517893 |
| AP003900.1 | 1.851119936 | 2.72E-07 |
| AC159540.2 | -1.214947026 | 6.44E-06 |
| AL109811.3 | -0.690374653 | 4.32E-07 |
| GOLGA6L17P | 1.165474693 | 0.037329277 |
| AL442128.2 | 0.985660429 | 0.013390395 |
| PGM5P2 | -0.557749399 | 0.019496738 |
| MROH6 | 0.378069403 | 0.004366143 |
| PSMB3 | 0.128128082 | 0.033544628 |
| AL138478.1 | -0.500220616 | 0.023912297 |
| RN7SKP175 | 3.352293861 | 0.000642488 |
| AL391988.1 | -1.596823462 | 0.002389341 |
| SNORA50C | 3.130245369 | 2.04E-08 |
| SPC25 | -1.401078572 | 3.00E-05 |
| DDX6P2 | -1.018762216 | 0.00169347 |
| AL357500.2 | -2.118166925 | 0.003237955 |
| MRPL45 | 0.269791511 | 0.001652867 |
| AC107308.1 | -1.260344559 | 0.036681849 |
| AC006449.7 | 1.734514101 | 1.11E-06 |
| AC244489.3 | 0.353122116 | 0.039469662 |
| SNORA67 | 2.647901108 | 0.001133602 |
| AC107075.1 | 0.245204356 | 0.01401279 |
| AC139100.2 | 1.82712866 | 7.42E-05 |
| AL627171.3 | 7.079430199 | 2.19E-05 |
| AC245690.1 | -0.773961447 | 0.00141155 |
| RDM1 | 1.507942238 | 0.012205197 |
| LY6E | 0.139295922 | 0.024749027 |
| MT2P1 | 0.849231219 | 0.000304598 |
| GOLGA8J | 2.210445859 | 0.002598642 |
| AC135050.6 | -2.112081542 | 6.50E-07 |
| AL358075.4 | -1.168637275 | 0.000267667 |
| AL354811.1 | 0.833749051 | 0.001269724 |
| RNA5-8SN1 | 0.698623518 | 0.002652408 |
| AL118505.1 | 1.379226538 | 0.016425906 |
| AC011043.3 | 1.48568998 | 0.02913491 |

| | | |
|---|---|---|
| TRIM69 | 8.25097071 | 3.08E-07 |
| CLN8 | -0.397289987 | 0.00290178 |
| PRICKLE4 | 1.41002729 | 0.000222534 |
| RNA5-8SN2 | 0.524488756 | 0.016534338 |
| TSTA3 | 0.296942298 | 0.005032038 |
| MYO19 | 0.315038158 | 0.000626402 |
| ZNF229 | -0.584167657 | 3.23E-05 |
| AL031673.1 | 2.334730642 | 2.49E-05 |
| AC121338.2 | 3.012413031 | 0.000871775 |
| AL354696.2 | 2.843424563 | 0.000158437 |
| AC243791.2 | -0.945002837 | 0.002040717 |
| AC138393.3 | 1.743468847 | 0.000416303 |
| NAPRT | 0.271432474 | 0.035696306 |
| CHMP1B2P | -0.525225932 | 0.00525504 |
| SLC43A2 | -0.523436727 | 0.036732858 |
| AC007325.3 | -0.652169472 | 0.005172725 |
| AL589182.2 | 0.417172766 | 0.032095811 |
| TBC1D3E | 1.229368798 | 0.019205985 |
| TSEN34 | -0.498944459 | 0.049641867 |
| ZNF426-DT | 1.662024508 | 0.01761627 |
| C11orf98 | -0.163598035 | 0.011630464 |
| AC037198.2 | -1.555839127 | 9.15E-07 |
| AZU1 | 4.168480143 | 2.20E-05 |
| AC141557.2 | 0.851562481 | 0.045670238 |
| H2AC17 | 3.43015969 | 1.66E-05 |
| AC100847.1 | 1.313195979 | 0.008607645 |
| NKILA | 2.533278864 | 2.40E-08 |
| AC005332.6 | -0.611866717 | 0.000176746 |
| AC090510.3 | -1.005913475 | 0.000613851 |
| AL136295.7 | 0.745653491 | 0.014880231 |
| LINC00624 | -0.74130306 | 0.002453399 |
| AC007325.4 | 1.018660987 | 0.030401309 |
| H3C10 | 3.791738526 | 1.12E-09 |
| AC073508.3 | -0.891642332 | 0.000875596 |
| TP53TG3F | 2.607875931 | 0.001062675 |
| AC117503.2 | 0.542518579 | 0.031462926 |
| AC090616.6 | -1.349744502 | 0.005408423 |
| CU638689.1 | 1.716629911 | 0.021497106 |
| CEP83-DT | 2.705510696 | 4.31E-07 |
| AC006213.6 | 1.518027583 | 4.98E-05 |
| AC002310.5 | -0.673030852 | 0.019880034 |
| CR381653.1 | 0.91304687 | 0.000996439 |
| AC244517.4 | -0.50350337 | 0.041580203 |
| AL031587.5 | -0.301879724 | 0.008775875 |
| AC127070.4 | -1.301270214 | 0.024913077 |
| AC138907.8 | -1.10166499 | 0.010060022 |
| AC006378.1 | 3.716341364 | 0.001540191 |
| AC132938.5 | 2.412309312 | 0.000200323 |

| | | |
|---|---|---|
| AL031602.1 | 6.822795279 | 2.73E-05 |
| AL138847.2 | 3.716125957 | 0.008089109 |
| AC093909.6 | -0.54109535 | 0.00038118 |
| AL031009.1 | 0.630493091 | 0.027162076 |
| AC002350.2 | -0.714756059 | 0.041566269 |
| FP671120.5 | -1.324892152 | 1.65E-06 |
| AL662907.1 | 3.212720772 | 2.72E-08 |
| OR51B2 | 2.502208411 | 4.13E-06 |
| AL356863.1 | 7.407776011 | 5.97E-06 |
| AC005225.4 | 3.382079433 | 0.000198324 |
| AC022211.5 | 0.486407112 | 0.011125918 |
| AC011447.6 | 1.601585203 | 0.006494562 |
| AC102945.2 | -1.524144719 | 0.005700275 |
| AC244517.5 | 0.443081024 | 0.015830719 |
| AC080080.1 | 2.797530062 | 4.20E-06 |
| AC015813.5 | -0.876554994 | 2.55E-08 |
| AL807752.6 | 1.143962061 | 0.012654199 |
| AL022322.2 | 1.417934929 | 0.003853912 |
| AC022400.8 | -0.399256686 | 0.030578764 |
| AC243964.4 | 0.910100662 | 2.14E-05 |
| AC138470.1 | 1.341667309 | 0.029381833 |
| AC093535.2 | 3.584041387 | 0.000127821 |
| AL356585.2 | -1.614259036 | 0.001886885 |
| AL590326.1 | 2.182022322 | 0.005132133 |
| AC011912.1 | 1.452856042 | 0.004926628 |
| AC126474.1 | -1.242482099 | 0.001415733 |
| AC129492.7 | 2.2074201 | 1.80E-06 |
| AL353726.2 | -0.586999914 | 0.015871179 |
| AC233300.1 | 1.7377485 | 0.007713012 |
| AC003681.1 | -0.727376095 | 1.28E-06 |
| AC141586.5 | 2.477171887 | 0.001608907 |
| CTBP2P9 | 2.800257577 | 0.002326669 |
| AL662907.2 | 0.725103455 | 0.040308431 |
| AC135048.4 | 1.410529143 | 0.000158657 |
| AC068669.1 | 1.937227914 | 0.000243627 |
| AC134043.3 | 4.502794723 | 0.000581412 |
| AC015813.6 | 1.935654747 | 3.76E-11 |
| AC008522.1 | -1.073131714 | 1.76E-08 |
| AC122688.3 | -0.57747437 | 0.036212656 |
| AP003108.3 | 0.350319221 | 0.036668107 |
| AC007614.1 | 6.772642866 | 3.47E-05 |
| AC020604.1 | 3.357205783 | 0.000226053 |
| AC087741.3 | 2.361335271 | 0.000317439 |
| AL078621.3 | 1.391273928 | 0.003445848 |
| AC012020.1 | 4.513493877 | 0.000721523 |
| AL136164.3 | 1.279203052 | 0.004167973 |
| AL139288.2 | 3.815488804 | 0.000112392 |
| AC020910.6 | 1.077337923 | 0.000560986 |

| | | |
|---|---|---|
| AJ003147.3 | 3.83597549 | 3.70E-05 |
| AC096636.1 | -2.670568004 | 0.000104417 |
| AC021945.1 | -0.939875672 | 0.015112451 |
| AC092692.1 | 4.374140863 | 0.001045189 |
| AL359195.2 | 6.352855785 | 2.32E-06 |
| AC079331.1 | 6.8249165 | 2.75E-05 |
| AL031123.3 | 3.043417275 | 2.76E-05 |
| AC137723.1 | 4.997650788 | 0.001238826 |
| AC003973.2 | -0.300130153 | 0.045337485 |
| AL158801.6 | -0.432429389 | 8.85E-05 |
| AL359183.1 | 1.376998383 | 0.004823356 |
| AC112497.2 | -0.457867456 | 0.011476686 |
| AL445363.3 | -1.128623584 | 5.96E-06 |
| AC092279.3 | -1.494765303 | 0.01774747 |
| AC087164.2 | -0.400266475 | 0.011041222 |
| AL513497.1 | 2.460646774 | 9.88E-10 |
| AC006277.1 | 3.767629021 | 2.07E-06 |
| Z99129.4 | 1.429711522 | 4.29E-05 |
| AL353763.1 | 3.033416904 | 5.88E-07 |
| AC090498.1 | 0.372510262 | 5.78E-06 |
| AC004623.1 | 1.37819134 | 0.007902018 |
| AL355377.2 | 6.773740543 | 3.21E-05 |
| AC083843.4 | 1.606736168 | 0.005099914 |
| AC093525.8 | -0.982379183 | 0.016900495 |
| AC115618.3 | 0.577906928 | 0.02984094 |
| AC008764.8 | 0.970176826 | 3.84E-05 |
| AC006486.2 | 1.606389326 | 2.24E-05 |
| AC005261.6 | 1.288371861 | 0.001597762 |
| AC010435.1 | -0.762011969 | 0.030080379 |
| AL845472.2 | 0.87781821 | 0.000126106 |
| AC020763.4 | -0.57340916 | 1.97E-05 |
| AC109326.1 | 2.294679918 | 3.04E-10 |
| AC067930.5 | 0.16746598 | 0.020337556 |
| AL353795.3 | 3.812312361 | 1.18E-05 |
| AC131280.1 | 2.791276047 | 0.002584345 |
| AC020978.9 | -0.647668832 | 0.001006415 |
| AL451064.2 | 1.212005264 | 0.032425439 |
| AC005703.6 | -0.97793448 | 0.039705288 |
| AC131649.2 | 2.049247698 | 1.27E-05 |
| AL603839.4 | 3.605974585 | 6.12E-07 |
| AL359922.3 | -1.265129718 | 0.003766374 |
| AC006128.1 | 0.942958685 | 0.001837055 |
| AC005336.3 | 6.818359605 | 1.10E-06 |
| SNX18P12 | 1.90879271 | 0.014383101 |
| AC007342.7 | 1.290714621 | 0.006306182 |
| LINC02033 | 7.608911638 | 2.93E-06 |
| KMT5AP1 | -0.323493345 | 0.00345147 |
| AL022311.1 | 0.199176567 | 0.002155704 |

| | | |
|---|---|---|
| AC132938.6 | -1.091144184 | 0.045968127 |
| AC008764.9 | 3.275017277 | 2.22E-05 |
| AC105339.5 | 4.317017729 | 5.72E-06 |
| AC118344.4 | 0.912564689 | 0.001767977 |
| AC005899.9 | 1.115313013 | 8.30E-05 |
| AC013394.1 | 0.734579777 | 0.000214706 |
| LINC00552 | 3.2315415 | 5.71E-09 |
| AC138907.9 | -1.261731636 | 0.035499033 |
| AC015909.5 | 1.144222858 | 0.032421924 |
| BCLAF1P2 | -1.355811005 | 1.13E-10 |
| AC005538.2 | 0.249560869 | 0.031472449 |
| AL031846.2 | -0.643686921 | 0.032446515 |
| AL356273.6 | 2.314480809 | 8.79E-05 |
| AC083964.2 | 7.305640991 | 6.09E-06 |
| AC004453.2 | 4.429035688 | 1.12E-08 |
| AC124864.3 | -3.506218853 | 0.013425574 |
| AC006511.4 | 1.659558273 | 5.47E-05 |
| AC091152.4 | 3.791764887 | 3.38E-06 |
| AC134407.3 | -0.950702534 | 0.028523318 |
| Z73965.1 | 3.183975932 | 4.75E-05 |
| AL356585.4 | -2.181107934 | 2.31E-05 |
| AL138831.3 | 4.388958994 | 0.003218602 |
| AL032819.2 | 7.585084998 | 2.53E-06 |
| AC008895.1 | -1.162823811 | 0.046964438 |
| AL137783.1 | 8.478254098 | 1.01E-07 |
| GVQW2 | 3.412154472 | 5.55E-08 |
| AC023024.2 | 3.263552116 | 0.00053255 |
| AC011815.3 | 3.642578039 | 8.05E-05 |
| AC008079.1 | 0.53914765 | 0.047528361 |
| AP001350.2 | -1.368097678 | 0.001042188 |
| AC116407.4 | 1.097023415 | 0.023890547 |
| AC022336.4 | 2.664170215 | 0.000210439 |
| AC118282.4 | 2.654413929 | 1.47E-06 |
| AC104581.4 | -0.863571504 | 3.33E-07 |
| AC004241.5 | 2.082113914 | 0.000162628 |
| AL022069.2 | 4.099290543 | 6.36E-07 |
| AC130304.1 | -0.569034117 | 0.001047036 |
| AC023818.1 | -2.507798136 | 1.34E-05 |
| AL353763.2 | -0.978544423 | 1.85E-05 |
| AC011447.7 | -1.250991307 | 6.55E-06 |
| AC126474.2 | -2.4336365 | 3.48E-06 |
| AC008555.7 | 2.933454793 | 2.65E-06 |
| AL022393.1 | 2.246869152 | 0.001295566 |
| PLAC4 | -4.331541453 | 1.13E-05 |
| AL021920.2 | -2.646297742 | 7.84E-06 |
| AC073857.1 | -2.356785375 | 1.12E-07 |
| AL662795.2 | 0.828458659 | 0.017943392 |
| AL132780.5 | 1.68160741 | 0.004061222 |

| | | |
|---|---|---|
| AC027290.3 | 0.581948676 | 0.000826368 |
| AP000892.4 | -1.136471059 | 2.68E-08 |
| AL049844.3 | -0.997778371 | 2.58E-06 |
| AC009078.3 | -2.986500265 | 0.000263306 |
| AC133065.3 | -2.457621367 | 4.14E-07 |
| AC022413.2 | 1.556895359 | 0.000507784 |
| AL136531.3 | 4.602014333 | 4.52E-06 |
| PCDH20 | -0.384426463 | 0.013779992 |
| AL359258.3 | 1.870429941 | 3.08E-05 |
| AC022107.1 | 1.745880844 | 2.09E-08 |
| AD000864.1 | 1.320376803 | 0.033864824 |
| AC005831.1 | -1.291767629 | 1.27E-07 |
| AC026401.3 | -1.56259957 | 1.62E-05 |
| GGT4P | -2.967286188 | 5.52E-05 |
| AC106886.5 | -0.37905782 | 2.71E-05 |
| AC012254.5 | 7.401354604 | 5.71E-06 |
| MTCO1P1 | 3.693166073 | 0.000109449 |
| AC233723.2 | 1.844428453 | 5.26E-05 |
| AC131212.4 | -2.179747489 | 0.006057589 |
| AC025262.3 | -0.855760124 | 0.006802289 |
| AC074183.2 | 1.40440766 | 8.15E-07 |
| CTBP2P10 | 3.251197768 | 2.34E-05 |
| AC020917.4 | -1.670106511 | 0.019877156 |
| AC009084.2 | 7.234906282 | 7.71E-06 |
| AC127496.7 | -2.100520051 | 2.33E-05 |
| AC011466.4 | 2.261515293 | 0.008083418 |
| AC092159.3 | 7.553789421 | 2.96E-06 |
| AP002364.1 | 2.566378666 | 7.51E-06 |
| AC019080.5 | 1.874975479 | 0.005309156 |
| AL353898.3 | 2.310095263 | 0.003638274 |
| AP000648.4 | -0.601563439 | 0.000513016 |
| AL109806.1 | 3.810183232 | 7.80E-06 |
| AC022532.1 | 3.178697569 | 0.000589724 |
| AC093525.10 | 0.861087308 | 0.035565707 |
| AC132872.4 | -0.357252752 | 0.024684967 |
| AC115284.2 | 1.770509236 | 2.82E-05 |
| AL357673.2 | 7.793064099 | 1.19E-06 |
| AC084876.2 | 2.049064016 | 8.41E-05 |
| FP565260.6 | -1.12863929 | 0.000113815 |
| AC006058.4 | -0.833925402 | 0.000440314 |
| FP236383.3 | -1.371547696 | 3.48E-07 |
