## Supplementary file S2 for "Resveratrol targets G-quadruplexes to exert its pharmacological effects"

**Supplementary file 2. Functional enrichment analysis of upregulated DEGs**

| term_name | term_id | source | p_value |
|---|---|---|---|
| organonitrogen compound metabolic process | GO:1901564 | GO:BP | 1.50E-30 |
| cellular nitrogen compound biosynthetic process | GO:0044271 | GO:BP | 9.56E-30 |
| biological regulation | GO:0065007 | GO:BP | 1.32E-28 |
| organic cyclic compound biosynthetic process | GO:1901362 | GO:BP | 2.20E-28 |
| regulation of primary metabolic process | GO:0080090 | GO:BP | 7.46E-27 |
| regulation of nitrogen compound metabolic process | GO:0051171 | GO:BP | 2.10E-25 |
| regulation of biological process | GO:0050789 | GO:BP | 2.31E-24 |
| heterocycle biosynthetic process | GO:0018130 | GO:BP | 7.40E-23 |
| aromatic compound biosynthetic process | GO:0019438 | GO:BP | 8.23E-23 |
| developmental process | GO:0032502 | GO:BP | 2.86E-22 |
| nucleobase-containing compound biosynthetic process | GO:0034654 | GO:BP | 3.24E-22 |
| multicellular organismal process | GO:0032501 | GO:BP | 3.42E-22 |
| regulation of response to stimulus | GO:0048583 | GO:BP | 6.05E-22 |
| cytoplasmic translation | GO:0002181 | GO:BP | 7.00E-22 |
| anatomical structure development | GO:0048856 | GO:BP | 1.54E-21 |
| regulation of cellular process | GO:0050794 | GO:BP | 3.41E-21 |
| positive regulation of biological process | GO:0048518 | GO:BP | 1.25E-20 |
| small molecule metabolic process | GO:0044281 | GO:BP | 1.65E-20 |
| response to stress | GO:0006950 | GO:BP | 2.59E-20 |
| cell differentiation | GO:0030154 | GO:BP | 8.92E-20 |
| cellular developmental process | GO:0048869 | GO:BP | 9.29E-20 |
| positive regulation of cellular process | GO:0048522 | GO:BP | 1.08E-18 |
| protein metabolic process | GO:0019538 | GO:BP | 1.37E-18 |
| regulation of DNA-templated transcription | GO:0006355 | GO:BP | 7.19E-18 |
| regulation of RNA biosynthetic process | GO:2001141 | GO:BP | 7.30E-18 |
| generation of precursor metabolites and energy | GO:0006091 | GO:BP | 9.25E-18 |
| regulation of transcription by RNA polymerase II | GO:0006357 | GO:BP | 1.16E-17 |
| RNA biosynthetic process | GO:0032774 | GO:BP | 2.50E-17 |
| regulation of nucleobase-containing compound metabolic p | GO:0019219 | GO:BP | 2.50E-17 |
| multicellular organism development | GO:0007275 | GO:BP | 2.79E-17 |
| DNA-templated transcription | GO:0006351 | GO:BP | 3.02E-17 |
| regulation of cell communication | GO:0010646 | GO:BP | 1.02E-16 |
| regulation of signaling | GO:0023051 | GO:BP | 1.27E-16 |
| transcription by RNA polymerase II | GO:0006366 | GO:BP | 3.31E-16 |
| regulation of RNA metabolic process | GO:0051252 | GO:BP | 6.27E-16 |
| cellular response to stimulus | GO:0051716 | GO:BP | 8.10E-16 |
| response to stimulus | GO:0050896 | GO:BP | 8.87E-16 |
| response to abiotic stimulus | GO:0009628 | GO:BP | 9.95E-16 |
| response to organic substance | GO:0010033 | GO:BP | 1.21E-15 |
| phosphorus metabolic process | GO:0006793 | GO:BP | 3.28E-15 |
| tissue development | GO:0009888 | GO:BP | 4.04E-15 |
| anatomical structure morphogenesis | GO:0009653 | GO:BP | 5.90E-15 |
| regulation of signal transduction | GO:0009966 | GO:BP | 7.07E-15 |
| phosphate-containing compound metabolic process | GO:0006796 | GO:BP | 1.12E-14 |
| regulation of multicellular organismal process | GO:0051239 | GO:BP | 1.13E-14 |
| cell development | GO:0048468 | GO:BP | 6.76E-14 |

| | | | |
|---|---|---|---|
| system development | GO:0048731 | GO:BP | 1.15E-13 |
| cellular response to chemical stimulus | GO:0070887 | GO:BP | 1.18E-13 |
| regulation of metabolic process | GO:0019222 | GO:BP | 2.57E-13 |
| transport | GO:0006810 | GO:BP | 4.46E-13 |
| carboxylic acid metabolic process | GO:0019752 | GO:BP | 1.44E-12 |
| oxoacid metabolic process | GO:0043436 | GO:BP | 3.66E-12 |
| small molecule biosynthetic process | GO:0044283 | GO:BP | 4.69E-12 |
| organic acid metabolic process | GO:0006082 | GO:BP | 4.79E-12 |
| response to external stimulus | GO:0009605 | GO:BP | 5.18E-12 |
| lipid metabolic process | GO:0006629 | GO:BP | 5.76E-12 |
| positive regulation of response to stimulus | GO:0048584 | GO:BP | 7.37E-12 |
| cell death | GO:0008219 | GO:BP | 7.37E-12 |
| positive regulation of metabolic process | GO:0009893 | GO:BP | 7.55E-12 |
| programmed cell death | GO:0012501 | GO:BP | 1.07E-11 |
| animal organ development | GO:0048513 | GO:BP | 1.07E-11 |
| monocarboxylic acid metabolic process | GO:0032787 | GO:BP | 1.43E-11 |
| regulation of apoptotic process | GO:0042981 | GO:BP | 1.46E-11 |
| regulation of programmed cell death | GO:0043067 | GO:BP | 1.72E-11 |
| organelle organization | GO:0006996 | GO:BP | 1.82E-11 |
| localization | GO:0051179 | GO:BP | 1.95E-11 |
| positive regulation of multicellular organismal process | GO:0051240 | GO:BP | 2.00E-11 |
| cellular response to stress | GO:0033554 | GO:BP | 4.06E-11 |
| regulation of biological quality | GO:0065008 | GO:BP | 5.43E-11 |
| cell communication | GO:0007154 | GO:BP | 5.44E-11 |
| regulation of cellular metabolic process | GO:0031323 | GO:BP | 5.66E-11 |
| positive regulation of cellular metabolic process | GO:0031325 | GO:BP | 5.91E-11 |
| steroid biosynthetic process | GO:0006694 | GO:BP | 6.23E-11 |
| response to oxygen-containing compound | GO:1901700 | GO:BP | 7.35E-11 |
| organic hydroxy compound metabolic process | GO:1901615 | GO:BP | 8.22E-11 |
| intracellular signal transduction | GO:0035556 | GO:BP | 1.11E-10 |
| apoptotic process | GO:0006915 | GO:BP | 1.11E-10 |
| catabolic process | GO:0009056 | GO:BP | 1.21E-10 |
| regulation of intracellular signal transduction | GO:1902531 | GO:BP | 1.26E-10 |
| signaling | GO:0023052 | GO:BP | 1.61E-10 |
| aerobic respiration | GO:0009060 | GO:BP | 2.42E-10 |
| organonitrogen compound biosynthetic process | GO:1901566 | GO:BP | 2.57E-10 |
| amide metabolic process | GO:0043603 | GO:BP | 2.65E-10 |
| regulation of macromolecule metabolic process | GO:0060255 | GO:BP | 3.77E-10 |
| regulation of localization | GO:0032879 | GO:BP | 4.12E-10 |
| cell activation | GO:0001775 | GO:BP | 4.18E-10 |
| organic hydroxy compound biosynthetic process | GO:1901617 | GO:BP | 4.18E-10 |
| inflammatory response | GO:0006954 | GO:BP | 5.61E-10 |
| organophosphate metabolic process | GO:0019637 | GO:BP | 5.83E-10 |
| lipid biosynthetic process | GO:0008610 | GO:BP | 5.83E-10 |
| oxidative phosphorylation | GO:0006119 | GO:BP | 6.15E-10 |
| regulation of transport | GO:0051049 | GO:BP | 6.40E-10 |
| energy derivation by oxidation of organic compounds | GO:0015980 | GO:BP | 1.00E-09 |
| signal transduction | GO:0007165 | GO:BP | 1.04E-09 |

| | | | |
|---|---|---|---|
| electron transport chain | GO:0022900 | GO:BP | 1.06E-09 |
| positive regulation of macromolecule metabolic process | GO:0010604 | GO:BP | 1.27E-09 |
| negative regulation of nitrogen compound metabolic proces | GO:0051172 | GO:BP | 1.43E-09 |
| carbohydrate derivative metabolic process | GO:1901135 | GO:BP | 2.33E-09 |
| regulation of protein metabolic process | GO:0051246 | GO:BP | 2.77E-09 |
| regulation of biosynthetic process | GO:0009889 | GO:BP | 3.02E-09 |
| regulation of cell population proliferation | GO:0042127 | GO:BP | 3.02E-09 |
| peptide metabolic process | GO:0006518 | GO:BP | 3.18E-09 |
| apoptotic signaling pathway | GO:0097190 | GO:BP | 3.41E-09 |
| regulation of cellular biosynthetic process | GO:0031326 | GO:BP | 3.83E-09 |
| cellular respiration | GO:0045333 | GO:BP | 5.24E-09 |
| cellular catabolic process | GO:0044248 | GO:BP | 7.59E-09 |
| mitochondrial ATP synthesis coupled electron transport | GO:0042775 | GO:BP | 7.59E-09 |
| ATP synthesis coupled electron transport | GO:0042773 | GO:BP | 7.59E-09 |
| cellular lipid metabolic process | GO:0044255 | GO:BP | 8.75E-09 |
| regulation of macromolecule biosynthetic process | GO:0010556 | GO:BP | 8.75E-09 |
| establishment of localization | GO:0051234 | GO:BP | 9.06E-09 |
| cellular response to organic substance | GO:0071310 | GO:BP | 9.23E-09 |
| aerobic electron transport chain | GO:0019646 | GO:BP | 1.03E-08 |
| organonitrogen compound catabolic process | GO:1901565 | GO:BP | 1.28E-08 |
| positive regulation of macromolecule biosynthetic process | GO:0010557 | GO:BP | 1.67E-08 |
| neurogenesis | GO:0022008 | GO:BP | 1.69E-08 |
| purine nucleotide metabolic process | GO:0006163 | GO:BP | 2.09E-08 |
| steroid metabolic process | GO:0008202 | GO:BP | 2.33E-08 |
| organic substance catabolic process | GO:1901575 | GO:BP | 2.34E-08 |
| positive regulation of apoptotic process | GO:0043065 | GO:BP | 2.36E-08 |
| positive regulation of biosynthetic process | GO:0009891 | GO:BP | 2.42E-08 |
| ATP metabolic process | GO:0046034 | GO:BP | 2.57E-08 |
| positive regulation of programmed cell death | GO:0043068 | GO:BP | 2.62E-08 |
| nervous system development | GO:0007399 | GO:BP | 2.63E-08 |
| purine-containing compound metabolic process | GO:0072521 | GO:BP | 2.66E-08 |
| external encapsulating structure organization | GO:0045229 | GO:BP | 2.71E-08 |
| regulation of developmental process | GO:0050793 | GO:BP | 3.24E-08 |
| positive regulation of cellular biosynthetic process | GO:0031328 | GO:BP | 4.58E-08 |
| extracellular structure organization | GO:0043062 | GO:BP | 4.62E-08 |
| nucleobase-containing small molecule metabolic process | GO:0055086 | GO:BP | 4.67E-08 |
| purine nucleoside triphosphate metabolic process | GO:0009144 | GO:BP | 5.34E-08 |
| purine ribonucleoside triphosphate metabolic process | GO:0009205 | GO:BP | 5.35E-08 |
| plasma membrane bounded cell projection organization | GO:0120036 | GO:BP | 6.15E-08 |
| cell projection organization | GO:0030030 | GO:BP | 6.26E-08 |
| alcohol metabolic process | GO:0006066 | GO:BP | 6.43E-08 |
| cell population proliferation | GO:0008283 | GO:BP | 6.47E-08 |
| positive regulation of cell communication | GO:0010647 | GO:BP | 6.59E-08 |
| negative regulation of biological process | GO:0048519 | GO:BP | 6.61E-08 |
| anatomical structure formation involved in morphogenesis | GO:0048646 | GO:BP | 6.93E-08 |
| response to mechanical stimulus | GO:0009612 | GO:BP | 7.08E-08 |
| extracellular matrix organization | GO:0030198 | GO:BP | 7.36E-08 |
| regulation of cytokine production | GO:0001817 | GO:BP | 7.39E-08 |

| | | | |
|---|---|---|---|
| response to chemical | GO:0042221 | GO:BP | 8.17E-08 |
| cell surface receptor signaling pathway | GO:0007166 | GO:BP | 9.17E-08 |
| cytokine production | GO:0001816 | GO:BP | 9.30E-08 |
| respiratory electron transport chain | GO:0022904 | GO:BP | 9.41E-08 |
| positive regulation of signaling | GO:0023056 | GO:BP | 9.55E-08 |
| nucleotide metabolic process | GO:0009117 | GO:BP | 1.10E-07 |
| regulation of response to stress | GO:0080134 | GO:BP | 1.13E-07 |
| proton motive force-driven mitochondrial ATP synthesis | GO:0042776 | GO:BP | 1.15E-07 |
| homeostatic process | GO:0042592 | GO:BP | 1.19E-07 |
| regulation of gene expression | GO:0010468 | GO:BP | 1.20E-07 |
| regulation of immune system process | GO:0002682 | GO:BP | 1.33E-07 |
| protein maturation | GO:0051604 | GO:BP | 1.35E-07 |
| pyridine nucleotide metabolic process | GO:0019362 | GO:BP | 1.62E-07 |
| nicotinamide nucleotide metabolic process | GO:0046496 | GO:BP | 1.62E-07 |
| nucleoside phosphate metabolic process | GO:0006753 | GO:BP | 1.85E-07 |
| regulation of response to external stimulus | GO:0032101 | GO:BP | 1.85E-07 |
| positive regulation of nitrogen compound metabolic process | GO:0051173 | GO:BP | 2.13E-07 |
| regulation of apoptotic signaling pathway | GO:2001233 | GO:BP | 2.13E-07 |
| muscle structure development | GO:0061061 | GO:BP | 2.19E-07 |
| ribonucleoside triphosphate metabolic process | GO:0009199 | GO:BP | 2.30E-07 |
| small molecule catabolic process | GO:0044282 | GO:BP | 2.39E-07 |
| cell motility | GO:0048870 | GO:BP | 2.48E-07 |
| negative regulation of multicellular organismal process | GO:0051241 | GO:BP | 2.62E-07 |
| regulation of intrinsic apoptotic signaling pathway | GO:2001242 | GO:BP | 3.15E-07 |
| pyridine-containing compound metabolic process | GO:0072524 | GO:BP | 3.29E-07 |
| sterol metabolic process | GO:0016125 | GO:BP | 3.34E-07 |
| regulation of cell differentiation | GO:0045595 | GO:BP | 3.58E-07 |
| cell adhesion | GO:0007155 | GO:BP | 3.67E-07 |
| fatty acid metabolic process | GO:0006631 | GO:BP | 3.82E-07 |
| response to cytokine | GO:0034097 | GO:BP | 4.16E-07 |
| amide biosynthetic process | GO:0043604 | GO:BP | 4.29E-07 |
| regulation of molecular function | GO:0065009 | GO:BP | 4.35E-07 |
| negative regulation of response to stimulus | GO:0048585 | GO:BP | 4.47E-07 |
| intracellular signaling cassette | GO:0141124 | GO:BP | 4.71E-07 |
| cholesterol metabolic process | GO:0008203 | GO:BP | 5.03E-07 |
| negative regulation of protein metabolic process | GO:0051248 | GO:BP | 5.03E-07 |
| generation of neurons | GO:0048699 | GO:BP | 5.94E-07 |
| sterol biosynthetic process | GO:0016126 | GO:BP | 6.06E-07 |
| negative regulation of transcription by RNA polymerase II | GO:0000122 | GO:BP | 6.26E-07 |
| nucleoside triphosphate metabolic process | GO:0009141 | GO:BP | 6.49E-07 |
| cilium or flagellum-dependent cell motility | GO:0001539 | GO:BP | 6.57E-07 |
| cilium-dependent cell motility | GO:0060285 | GO:BP | 6.57E-07 |
| secondary alcohol metabolic process | GO:1902652 | GO:BP | 6.82E-07 |
| cellular response to external stimulus | GO:0071496 | GO:BP | 7.60E-07 |
| negative regulation of cell communication | GO:0010648 | GO:BP | 7.94E-07 |
| regulation of multicellular organismal development | GO:2000026 | GO:BP | 8.02E-07 |
| purine ribonucleoside triphosphate biosynthetic process | GO:0009206 | GO:BP | 8.70E-07 |
| negative regulation of programmed cell death | GO:0043069 | GO:BP | 8.99E-07 |

| | | | |
|---|---|---|---|
| muscle cell differentiation | GO:0042692 | GO:BP | 8.99E-07 |
| peptide biosynthetic process | GO:0043043 | GO:BP | 9.95E-07 |
| cilium movement involved in cell motility | GO:0060294 | GO:BP | 1.04E-06 |
| leukocyte activation | GO:0045321 | GO:BP | 1.06E-06 |
| striated muscle cell differentiation | GO:0051146 | GO:BP | 1.06E-06 |
| negative regulation of signaling | GO:0023057 | GO:BP | 1.09E-06 |
| regulation of cellular component organization | GO:0051128 | GO:BP | 1.18E-06 |
| purine nucleoside triphosphate biosynthetic process | GO:0009145 | GO:BP | 1.18E-06 |
| negative regulation of apoptotic process | GO:0043066 | GO:BP | 1.18E-06 |
| response to wounding | GO:0009611 | GO:BP | 1.23E-06 |
| cilium movement | GO:0003341 | GO:BP | 1.29E-06 |
| neuron differentiation | GO:0030182 | GO:BP | 1.37E-06 |
| translation | GO:0006412 | GO:BP | 1.44E-06 |
| positive regulation of signal transduction | GO:0009967 | GO:BP | 1.46E-06 |
| cellular response to abiotic stimulus | GO:0071214 | GO:BP | 1.80E-06 |
| cellular response to environmental stimulus | GO:0104004 | GO:BP | 1.80E-06 |
| cellular response to oxygen-containing compound | GO:1901701 | GO:BP | 1.87E-06 |
| epithelium development | GO:0060429 | GO:BP | 2.55E-06 |
| regulation of inflammatory response | GO:0050727 | GO:BP | 2.55E-06 |
| regulation of proteolysis | GO:0030162 | GO:BP | 2.64E-06 |
| positive regulation of immune system process | GO:0002684 | GO:BP | 3.18E-06 |
| detection of external stimulus | GO:0009581 | GO:BP | 3.58E-06 |
| secretion | GO:0046903 | GO:BP | 3.59E-06 |
| positive regulation of cytokine production | GO:0001819 | GO:BP | 3.59E-06 |
| monosaccharide metabolic process | GO:0005996 | GO:BP | 3.76E-06 |
| response to endogenous stimulus | GO:0009719 | GO:BP | 3.78E-06 |
| negative regulation of cellular process | GO:0048523 | GO:BP | 4.49E-06 |
| maintenance of location | GO:0051235 | GO:BP | 4.91E-06 |
| ribonucleoside triphosphate biosynthetic process | GO:0009201 | GO:BP | 5.70E-06 |
| circulatory system development | GO:0072359 | GO:BP | 6.60E-06 |
| monoatomic cation transport | GO:0006812 | GO:BP | 6.78E-06 |
| phosphorylation | GO:0016310 | GO:BP | 7.62E-06 |
| regulation of cell activation | GO:0050865 | GO:BP | 7.71E-06 |
| detection of abiotic stimulus | GO:0009582 | GO:BP | 7.92E-06 |
| epithelial cell proliferation | GO:0050673 | GO:BP | 8.16E-06 |
| multicellular organismal-level homeostasis | GO:0048871 | GO:BP | 8.99E-06 |
| carbohydrate metabolic process | GO:0005975 | GO:BP | 9.06E-06 |
| intrinsic apoptotic signaling pathway | GO:0097193 | GO:BP | 1.00E-05 |
| glucose metabolic process | GO:0006006 | GO:BP | 1.18E-05 |
| regulation of catalytic activity | GO:0050790 | GO:BP | 1.20E-05 |
| alcohol biosynthetic process | GO:0046165 | GO:BP | 1.20E-05 |
| cellular response to nutrient levels | GO:0031669 | GO:BP | 1.20E-05 |
| proteolysis | GO:0006508 | GO:BP | 1.23E-05 |
| ATP biosynthetic process | GO:0006754 | GO:BP | 1.23E-05 |
| organophosphate biosynthetic process | GO:0090407 | GO:BP | 1.23E-05 |
| cellular response to starvation | GO:0009267 | GO:BP | 1.33E-05 |
| proton motive force-driven ATP synthesis | GO:0015986 | GO:BP | 1.34E-05 |
| secretion by cell | GO:0032940 | GO:BP | 1.35E-05 |

| | | | |
|---|---|---|---|
| negative regulation of DNA-templated transcription | GO:0045892 | GO:BP | 1.46E-05 |
| response to lipid | GO:0033993 | GO:BP | 1.47E-05 |
| purine ribonucleotide metabolic process | GO:0009150 | GO:BP | 1.57E-05 |
| negative regulation of RNA biosynthetic process | GO:1902679 | GO:BP | 1.57E-05 |
| response to hypoxia | GO:0001666 | GO:BP | 1.63E-05 |
| positive regulation of gene expression | GO:0010628 | GO:BP | 1.70E-05 |
| secondary alcohol biosynthetic process | GO:1902653 | GO:BP | 1.89E-05 |
| cholesterol biosynthetic process | GO:0006695 | GO:BP | 1.89E-05 |
| cellular response to extracellular stimulus | GO:0031668 | GO:BP | 1.97E-05 |
| nucleoside triphosphate biosynthetic process | GO:0009142 | GO:BP | 2.07E-05 |
| positive regulation of response to external stimulus | GO:0032103 | GO:BP | 2.09E-05 |
| regulation of phosphate metabolic process | GO:0019220 | GO:BP | 2.11E-05 |
| positive regulation of inflammatory response | GO:0050729 | GO:BP | 2.26E-05 |
| regulation of phosphorus metabolic process | GO:0051174 | GO:BP | 2.28E-05 |
| lymphocyte activation | GO:0046649 | GO:BP | 2.32E-05 |
| ribose phosphate metabolic process | GO:0019693 | GO:BP | 2.32E-05 |
| cellular response to cytokine stimulus | GO:0071345 | GO:BP | 2.75E-05 |
| protein modification process | GO:0036211 | GO:BP | 2.80E-05 |
| export from cell | GO:0140352 | GO:BP | 2.85E-05 |
| response to oxygen levels | GO:0070482 | GO:BP | 2.85E-05 |
| protein folding | GO:0006457 | GO:BP | 2.96E-05 |
| purine-containing compound biosynthetic process | GO:0072522 | GO:BP | 3.41E-05 |
| positive regulation of developmental process | GO:0051094 | GO:BP | 3.50E-05 |
| sarcomere organization | GO:0045214 | GO:BP | 3.54E-05 |
| hexose metabolic process | GO:0019318 | GO:BP | 3.54E-05 |
| response to extracellular stimulus | GO:0009991 | GO:BP | 3.68E-05 |
| response to decreased oxygen levels | GO:0036293 | GO:BP | 3.72E-05 |
| positive regulation of proteolysis | GO:0045862 | GO:BP | 3.80E-05 |
| carboxylic acid catabolic process | GO:0046395 | GO:BP | 3.97E-05 |
| organic acid catabolic process | GO:0016054 | GO:BP | 3.97E-05 |
| cilium organization | GO:0044782 | GO:BP | 4.01E-05 |
| regulation of locomotion | GO:0040012 | GO:BP | 4.01E-05 |
| positive regulation of locomotion | GO:0040017 | GO:BP | 4.16E-05 |
| neuron development | GO:0048666 | GO:BP | 4.19E-05 |
| muscle cell development | GO:0055001 | GO:BP | 4.39E-05 |
| regulation of hormone levels | GO:0010817 | GO:BP | 4.41E-05 |
| inorganic ion transmembrane transport | GO:0098660 | GO:BP | 4.67E-05 |
| ribonucleotide metabolic process | GO:0009259 | GO:BP | 4.69E-05 |
| cell-cell signaling | GO:0007267 | GO:BP | 4.70E-05 |
| regulation of protein modification process | GO:0031399 | GO:BP | 4.70E-05 |
| positive regulation of cell motility | GO:2000147 | GO:BP | 4.70E-05 |
| purine nucleotide biosynthetic process | GO:0006164 | GO:BP | 4.78E-05 |
| negative regulation of signal transduction | GO:0009968 | GO:BP | 4.86E-05 |
| response to nutrient levels | GO:0031667 | GO:BP | 5.17E-05 |
| negative regulation of molecular function | GO:0044092 | GO:BP | 5.45E-05 |
| carbohydrate catabolic process | GO:0016052 | GO:BP | 5.54E-05 |
| response to oxidative stress | GO:0006979 | GO:BP | 5.54E-05 |
| positive regulation of protein metabolic process | GO:0051247 | GO:BP | 5.54E-05 |

| | | | |
|---|---|---|---|
| cellular anatomical entity morphogenesis | GO:0032989 | GO:BP | 5.77E-05 |
| animal organ morphogenesis | GO:0009887 | GO:BP | 6.01E-05 |
| organelle assembly | GO:0070925 | GO:BP | 6.05E-05 |
| regulation of peptidase activity | GO:0052547 | GO:BP | 6.68E-05 |
| supramolecular fiber organization | GO:0097435 | GO:BP | 7.18E-05 |
| positive regulation of intracellular signal transduction | GO:1902533 | GO:BP | 7.18E-05 |
| microtubule-based movement | GO:0007018 | GO:BP | 7.41E-05 |
| monoatomic cation transmembrane transport | GO:0098655 | GO:BP | 7.54E-05 |
| positive regulation of cell migration | GO:0030335 | GO:BP | 7.60E-05 |
| regulation of cell motility | GO:2000145 | GO:BP | 7.99E-05 |
| cellular localization | GO:0051641 | GO:BP | 8.13E-05 |
| myofibril assembly | GO:0030239 | GO:BP | 8.92E-05 |
| wound healing | GO:0042060 | GO:BP | 9.46E-05 |
| inorganic cation transmembrane transport | GO:0098662 | GO:BP | 0.000100291 |
| cellular response to endogenous stimulus | GO:0071495 | GO:BP | 0.000105577 |
| response to osmotic stress | GO:0006970 | GO:BP | 0.000105809 |
| negative regulation of RNA metabolic process | GO:0051253 | GO:BP | 0.000107725 |
| positive regulation of cell population proliferation | GO:0008284 | GO:BP | 0.000112662 |
| regulation of epithelial cell proliferation | GO:0050678 | GO:BP | 0.000112662 |
| negative regulation of cell population proliferation | GO:0008285 | GO:BP | 0.000118296 |
| neuron projection development | GO:0031175 | GO:BP | 0.000119126 |
| negative regulation of nucleobase-containing compound me | GO:0045934 | GO:BP | 0.000122293 |
| regulation of leukocyte activation | GO:0002694 | GO:BP | 0.000124282 |
| response to organonitrogen compound | GO:0010243 | GO:BP | 0.000131375 |
| calcium ion transport | GO:0006816 | GO:BP | 0.000134703 |
| growth | GO:0040007 | GO:BP | 0.000137218 |
| gliogenesis | GO:0042063 | GO:BP | 0.000139051 |
| transmembrane transport | GO:0055085 | GO:BP | 0.000148014 |
| response to nitrogen compound | GO:1901698 | GO:BP | 0.000149557 |
| positive regulation of leukocyte migration | GO:0002687 | GO:BP | 0.000149568 |
| muscle system process | GO:0003012 | GO:BP | 0.000150676 |
| tube development | GO:0035295 | GO:BP | 0.000158861 |
| cellular response to lipid | GO:0071396 | GO:BP | 0.000162409 |
| regulation of immune effector process | GO:0002697 | GO:BP | 0.000166852 |
| positive regulation of transport | GO:0051050 | GO:BP | 0.000168694 |
| regulation of defense response | GO:0031347 | GO:BP | 0.000168694 |
| sperm motility | GO:0097722 | GO:BP | 0.00017072 |
| flagellated sperm motility | GO:0030317 | GO:BP | 0.00017072 |
| purine ribonucleotide biosynthetic process | GO:0009152 | GO:BP | 0.000171638 |
| olefinic compound metabolic process | GO:0120254 | GO:BP | 0.000174831 |
| regulation of cell development | GO:0060284 | GO:BP | 0.000176708 |
| positive regulation of nucleobase-containing compound me | GO:0045935 | GO:BP | 0.000176708 |
| regulation of cell migration | GO:0030334 | GO:BP | 0.000176708 |
| cell-substrate adhesion | GO:0031589 | GO:BP | 0.000177702 |
| pyruvate metabolic process | GO:0006090 | GO:BP | 0.000190127 |
| cellular lipid catabolic process | GO:0044242 | GO:BP | 0.00022124 |
| cell-cell adhesion | GO:0098609 | GO:BP | 0.000231751 |
| macromolecule modification | GO:0043412 | GO:BP | 0.000234416 |

| | | | |
|---|---|---|---|
| maintenance of location in cell | GO:0051651 | GO:BP | 0.000237503 |
| striated muscle cell development | GO:0055002 | GO:BP | 0.000238445 |
| defense response | GO:0006952 | GO:BP | 0.000250504 |
| protein phosphorylation | GO:0006468 | GO:BP | 0.000258678 |
| regulation of synaptic plasticity | GO:0048167 | GO:BP | 0.000273671 |
| monoatomic ion transmembrane transport | GO:0034220 | GO:BP | 0.000281301 |
| organic acid biosynthetic process | GO:0016053 | GO:BP | 0.000286083 |
| ribose phosphate biosynthetic process | GO:0046390 | GO:BP | 0.000287021 |
| positive regulation of cell differentiation | GO:0045597 | GO:BP | 0.000298752 |
| type B pancreatic cell proliferation | GO:0044342 | GO:BP | 0.000302697 |
| cellular response to chemical stress | GO:0062197 | GO:BP | 0.000304238 |
| cellular component assembly involved in morphogenesis | GO:0010927 | GO:BP | 0.000304238 |
| response to reactive oxygen species | GO:0000302 | GO:BP | 0.000304902 |
| sperm flagellum assembly | GO:0120316 | GO:BP | 0.000307716 |
| detection of mechanical stimulus | GO:0050982 | GO:BP | 0.000313629 |
| NADP metabolic process | GO:0006739 | GO:BP | 0.000316114 |
| carboxylic acid biosynthetic process | GO:0046394 | GO:BP | 0.000317892 |
| collagen metabolic process | GO:0032963 | GO:BP | 0.000362731 |
| monocarboxylic acid biosynthetic process | GO:0072330 | GO:BP | 0.000376133 |
| response to starvation | GO:0042594 | GO:BP | 0.000376133 |
| monoatomic ion transport | GO:0006811 | GO:BP | 0.000377334 |
| regulation of lipid biosynthetic process | GO:0046890 | GO:BP | 0.000377901 |
| regulation of phosphorylation | GO:0042325 | GO:BP | 0.000385869 |
| regulation of secretion | GO:0051046 | GO:BP | 0.000405995 |
| cytoskeleton organization | GO:0007010 | GO:BP | 0.000407111 |
| metal ion transport | GO:0030001 | GO:BP | 0.000415052 |
| cilium assembly | GO:0060271 | GO:BP | 0.000428173 |
| nucleotide biosynthetic process | GO:0009165 | GO:BP | 0.000436295 |
| regulation of production of molecular mediator of immune re | GO:0002700 | GO:BP | 0.000440095 |
| sensory perception of mechanical stimulus | GO:0050954 | GO:BP | 0.000440095 |
| cellular modified amino acid metabolic process | GO:0006575 | GO:BP | 0.000449962 |
| positive regulation of apoptotic signaling pathway | GO:2001235 | GO:BP | 0.000451844 |
| positive regulation of peptidase activity | GO:0010952 | GO:BP | 0.000472356 |
| motile cilium assembly | GO:0044458 | GO:BP | 0.00050792 |
| regulation of endopeptidase activity | GO:0052548 | GO:BP | 0.000513256 |
| primary alcohol metabolic process | GO:0034308 | GO:BP | 0.000528503 |
| response to inorganic substance | GO:0010035 | GO:BP | 0.000550669 |
| ribonucleotide biosynthetic process | GO:0009260 | GO:BP | 0.000561245 |
| nucleoside phosphate biosynthetic process | GO:1901293 | GO:BP | 0.000568432 |
| cell-matrix adhesion | GO:0007160 | GO:BP | 0.000574565 |
| negative regulation of intracellular signal transduction | GO:1902532 | GO:BP | 0.000580091 |
| cellular aldehyde metabolic process | GO:0006081 | GO:BP | 0.000584652 |
| actin filament-based process | GO:0030029 | GO:BP | 0.000593581 |
| positive regulation of production of molecular mediator of im | GO:0002702 | GO:BP | 0.000627826 |
| locomotion | GO:0040011 | GO:BP | 0.000639602 |
| regulation of lipid metabolic process | GO:0019216 | GO:BP | 0.0006617 |
| response to organic cyclic compound | GO:0014070 | GO:BP | 0.000669996 |
| mitochondrial electron transport, NADH to ubiquinone | GO:0006120 | GO:BP | 0.000689018 |

| | | | |
|---|---|---|---|
| microtubule-based process | GO:0007017 | GO:BP | 0.000701878 |
| tube morphogenesis | GO:0035239 | GO:BP | 0.000740047 |
| regulation of lymphocyte activation | GO:0051249 | GO:BP | 0.000742501 |
| positive regulation of cell activation | GO:0050867 | GO:BP | 0.000765387 |
| sperm axoneme assembly | GO:0007288 | GO:BP | 0.00077204 |
| NAD metabolic process | GO:0019674 | GO:BP | 0.000794667 |
| monosaccharide biosynthetic process | GO:0046364 | GO:BP | 0.000809932 |
| negative regulation of protein modification process | GO:0031400 | GO:BP | 0.000830594 |
| positive regulation of intrinsic apoptotic signaling pathway | GO:2001244 | GO:BP | 0.000847336 |
| cellular response to oxygen levels | GO:0071453 | GO:BP | 0.00088673 |
| signal transduction by p53 class mediator | GO:0072331 | GO:BP | 0.000901946 |
| mitochondrial electron transport, cytochrome c to oxygen | GO:0006123 | GO:BP | 0.000924502 |
| response to unfolded protein | GO:0006986 | GO:BP | 0.000937872 |
| regulation of protein phosphorylation | GO:0001932 | GO:BP | 0.000955103 |
| positive regulation of defense response | GO:0031349 | GO:BP | 0.000978639 |
| regulation of hemopoiesis | GO:1903706 | GO:BP | 0.001001554 |
| negative regulation of developmental process | GO:0051093 | GO:BP | 0.001004685 |
| protein processing | GO:0016485 | GO:BP | 0.001048055 |
| muscle contraction | GO:0006936 | GO:BP | 0.001097544 |
| positive regulation of endopeptidase activity | GO:0010950 | GO:BP | 0.001144504 |
| cellular response to decreased oxygen levels | GO:0036294 | GO:BP | 0.00115154 |
| terpenoid metabolic process | GO:0006721 | GO:BP | 0.001182079 |
| cell migration | GO:0016477 | GO:BP | 0.001188338 |
| carbohydrate derivative biosynthetic process | GO:1901137 | GO:BP | 0.001188338 |
| negative regulation of growth | GO:0045926 | GO:BP | 0.001203406 |
| cellular response to hypoxia | GO:0071456 | GO:BP | 0.001216375 |
| chemical homeostasis | GO:0048878 | GO:BP | 0.00125678 |
| regulation of adaptive immune response | GO:0002819 | GO:BP | 0.001261298 |
| glucan catabolic process | GO:0009251 | GO:BP | 0.001279718 |
| integrin-mediated signaling pathway | GO:0007229 | GO:BP | 0.001279718 |
| organophosphate catabolic process | GO:0046434 | GO:BP | 0.001335122 |
| regulation of calcium ion transport | GO:0051924 | GO:BP | 0.00135646 |
| developmental growth | GO:0048589 | GO:BP | 0.001359369 |
| reproduction | GO:0000003 | GO:BP | 0.001359369 |
| response to topologically incorrect protein | GO:0035966 | GO:BP | 0.001364073 |
| regulation of cellular response to stress | GO:0080135 | GO:BP | 0.001387864 |
| regulation of response to endoplasmic reticulum stress | GO:1905897 | GO:BP | 0.001473941 |
| angiogenesis | GO:0001525 | GO:BP | 0.001529454 |
| reproductive process | GO:0022414 | GO:BP | 0.00153729 |
| negative regulation of epithelial cell proliferation | GO:0050680 | GO:BP | 0.001543263 |
| photoreceptor cell maintenance | GO:0045494 | GO:BP | 0.001543263 |
| hormone metabolic process | GO:0042445 | GO:BP | 0.00154412 |
| negative regulation of response to endoplasmic reticulum st | GO:1903573 | GO:BP | 0.001552802 |
| bone development | GO:0060348 | GO:BP | 0.001600271 |
| ossification | GO:0001503 | GO:BP | 0.001652397 |
| positive regulation of protein processing | GO:0010954 | GO:BP | 0.001652655 |
| photoreceptor cell outer segment organization | GO:0035845 | GO:BP | 0.001687311 |
| response to molecule of bacterial origin | GO:0002237 | GO:BP | 0.001687311 |

| | | | |
|---|---|---|---|
| blood vessel development | GO:0001568 | GO:BP | 0.001702575 |
| enzyme-linked receptor protein signaling pathway | GO:0007167 | GO:BP | 0.001771849 |
| proton transmembrane transport | GO:1902600 | GO:BP | 0.001791946 |
| regulation of cysteine-type endopeptidase activity | GO:2000116 | GO:BP | 0.001791946 |
| zymogen activation | GO:0031638 | GO:BP | 0.001792743 |
| glial cell differentiation | GO:0010001 | GO:BP | 0.001825565 |
| isoprenoid metabolic process | GO:0006720 | GO:BP | 0.001825565 |
| muscle tissue development | GO:0060537 | GO:BP | 0.001825565 |
| response to endoplasmic reticulum stress | GO:0034976 | GO:BP | 0.001825565 |
| vasculature development | GO:0001944 | GO:BP | 0.001825565 |
| regulation of immune response | GO:0050776 | GO:BP | 0.001825565 |
| positive regulation of cysteine-type endopeptidase activity | GO:2001056 | GO:BP | 0.001885869 |
| negative regulation of immune system process | GO:0002683 | GO:BP | 0.001925677 |
| positive regulation of DNA-templated transcription | GO:0045893 | GO:BP | 0.002010096 |
| axoneme assembly | GO:0035082 | GO:BP | 0.002029482 |
| positive regulation of cell development | GO:0010720 | GO:BP | 0.002034467 |
| response to metal ion | GO:0010038 | GO:BP | 0.002053236 |
| positive regulation of establishment of protein localization | GO:1904951 | GO:BP | 0.00213001 |
| gastro-intestinal system smooth muscle contraction | GO:0014831 | GO:BP | 0.00213001 |
| regulation of small molecule metabolic process | GO:0062012 | GO:BP | 0.002141418 |
| negative regulation of intrinsic apoptotic signaling pathway | GO:2001243 | GO:BP | 0.002193247 |
| NADH metabolic process | GO:0006734 | GO:BP | 0.002193247 |
| positive regulation of CD4-positive, alpha-beta T cell differer | GO:0043372 | GO:BP | 0.002193247 |
| regulation of adaptive immune response based on somatic r | GO:0002822 | GO:BP | 0.00228142 |
| regulation of vesicle-mediated transport | GO:0060627 | GO:BP | 0.002282726 |
| muscle organ development | GO:0007517 | GO:BP | 0.002284644 |
| regulation of secretion by cell | GO:1903530 | GO:BP | 0.002285488 |
| positive regulation of RNA biosynthetic process | GO:1902680 | GO:BP | 0.002289524 |
| regulation of growth | GO:0040008 | GO:BP | 0.002519513 |
| positive regulation of leukocyte activation | GO:0002696 | GO:BP | 0.002570325 |
| male gamete generation | GO:0048232 | GO:BP | 0.002573242 |
| phospholipid catabolic process | GO:0009395 | GO:BP | 0.002588583 |
| positive regulation of immune effector process | GO:0002699 | GO:BP | 0.002589542 |
| response to temperature stimulus | GO:0009266 | GO:BP | 0.002652858 |
| regulation of protein processing | GO:0070613 | GO:BP | 0.002729852 |
| cellular ketone metabolic process | GO:0042180 | GO:BP | 0.002738114 |
| glial cell development | GO:0021782 | GO:BP | 0.002781695 |
| monocarboxylic acid catabolic process | GO:0072329 | GO:BP | 0.002781695 |
| regulation of B cell activation | GO:0050864 | GO:BP | 0.002781695 |
| myeloid cell differentiation | GO:0030099 | GO:BP | 0.00279607 |
| purine ribonucleoside diphosphate catabolic process | GO:0009181 | GO:BP | 0.00279607 |
| purine nucleoside diphosphate catabolic process | GO:0009137 | GO:BP | 0.00279607 |
| tissue remodeling | GO:0048771 | GO:BP | 0.002799515 |
| regulation of MAPK cascade | GO:0043408 | GO:BP | 0.002848422 |
| intraciliary transport | GO:0042073 | GO:BP | 0.002901255 |
| positive regulation of lymphocyte activation | GO:0051251 | GO:BP | 0.002944995 |
| ADP metabolic process | GO:0046031 | GO:BP | 0.003089733 |
| ADP catabolic process | GO:0046032 | GO:BP | 0.003136659 |

| | | | |
|---|---|---|---|
| actin cytoskeleton organization | GO:0030036 | GO:BP | 0.003169565 |
| negative regulation of apoptotic signaling pathway | GO:2001234 | GO:BP | 0.003209818 |
| unsaturated fatty acid metabolic process | GO:0033559 | GO:BP | 0.003254453 |
| cellular response to radiation | GO:0071478 | GO:BP | 0.003362671 |
| regulation of neural precursor cell proliferation | GO:2000177 | GO:BP | 0.00338021 |
| cardiac muscle cell differentiation | GO:0055007 | GO:BP | 0.003501662 |
| hexose biosynthetic process | GO:0019319 | GO:BP | 0.003501662 |
| lipid oxidation | GO:0034440 | GO:BP | 0.003501662 |
| retinoid metabolic process | GO:0001523 | GO:BP | 0.003501662 |
| reactive oxygen species metabolic process | GO:0072593 | GO:BP | 0.003534103 |
| glycogen catabolic process | GO:0005980 | GO:BP | 0.00361363 |
| phosphatidylcholine catabolic process | GO:0034638 | GO:BP | 0.00361363 |
| fat cell differentiation | GO:0045444 | GO:BP | 0.003627737 |
| response to tumor necrosis factor | GO:0034612 | GO:BP | 0.003702764 |
| regulation of hydrolase activity | GO:0051336 | GO:BP | 0.00370414 |
| amino acid metabolic process | GO:0006520 | GO:BP | 0.003734399 |
| chaperone-mediated protein folding | GO:0061077 | GO:BP | 0.003734399 |
| platelet aggregation | GO:0070527 | GO:BP | 0.003734399 |
| gluconeogenesis | GO:0006094 | GO:BP | 0.00390014 |
| regulation of establishment of protein localization | GO:0070201 | GO:BP | 0.003900608 |
| purine ribonucleoside diphosphate metabolic process | GO:0009179 | GO:BP | 0.003906663 |
| purine nucleoside diphosphate metabolic process | GO:0009135 | GO:BP | 0.003906663 |
| lipid localization | GO:0010876 | GO:BP | 0.00410539 |
| microtubule bundle formation | GO:0001578 | GO:BP | 0.004198307 |
| negative regulation of catalytic activity | GO:0043086 | GO:BP | 0.004198307 |
| carbohydrate derivative catabolic process | GO:1901136 | GO:BP | 0.004198307 |
| biological process involved in interspecies interaction betwe | GO:0044419 | GO:BP | 0.004198307 |
| pyridine-containing compound catabolic process | GO:0072526 | GO:BP | 0.00421628 |
| glycolytic process | GO:0006096 | GO:BP | 0.004311966 |
| regulation of catabolic process | GO:0009894 | GO:BP | 0.004325993 |
| regulation of cell adhesion | GO:0030155 | GO:BP | 0.004417429 |
| non-membrane-bounded organelle assembly | GO:0140694 | GO:BP | 0.004429371 |
| liver morphogenesis | GO:0072576 | GO:BP | 0.004449173 |
| glucose catabolic process | GO:0006007 | GO:BP | 0.004449173 |
| positive regulation of protein maturation | GO:1903319 | GO:BP | 0.004449173 |
| regulation of leukocyte differentiation | GO:1902105 | GO:BP | 0.004462204 |
| positive regulation of RNA metabolic process | GO:0051254 | GO:BP | 0.004498924 |
| retina homeostasis | GO:0001895 | GO:BP | 0.004540191 |
| lipid modification | GO:0030258 | GO:BP | 0.004565905 |
| regulation of steroid biosynthetic process | GO:0050810 | GO:BP | 0.0047013 |
| pyridine nucleotide catabolic process | GO:0019364 | GO:BP | 0.0047013 |
| positive regulation of mononuclear cell migration | GO:0071677 | GO:BP | 0.0047013 |
| spermatogenesis | GO:0007283 | GO:BP | 0.0047013 |
| antigen processing and presentation of exogenous peptide a | GO:0002478 | GO:BP | 0.004732954 |
| phospholipid metabolic process | GO:0006644 | GO:BP | 0.004757969 |
| import into cell | GO:0098657 | GO:BP | 0.004816504 |
| polysaccharide catabolic process | GO:0000272 | GO:BP | 0.004818171 |
| regulation of plasminogen activation | GO:0010755 | GO:BP | 0.004818171 |

| | | | |
|---|---|---|---|
| macromolecule catabolic process | GO:0009057 | GO:BP | 0.004848743 |
| blood vessel morphogenesis | GO:0048514 | GO:BP | 0.004851495 |
| regulation of CD4-positive, alpha-beta T cell differentiation | GO:0043370 | GO:BP | 0.004862409 |
| regulation of metal ion transport | GO:0010959 | GO:BP | 0.004876834 |
| negative regulation of cell differentiation | GO:0045596 | GO:BP | 0.004966471 |
| cellular response to tumor necrosis factor | GO:0071356 | GO:BP | 0.005020332 |
| ribonucleoside diphosphate catabolic process | GO:0009191 | GO:BP | 0.005049447 |
| protein catabolic process | GO:0030163 | GO:BP | 0.005143667 |
| cellular response to osmotic stress | GO:0071470 | GO:BP | 0.005251289 |
| positive regulation of B cell activation | GO:0050871 | GO:BP | 0.005282889 |
| intrinsic apoptotic signaling pathway by p53 class mediator | GO:0072332 | GO:BP | 0.005282889 |
| interleukin-6 production | GO:0032635 | GO:BP | 0.005287655 |
| regulation of interleukin-6 production | GO:0032675 | GO:BP | 0.005287655 |
| heart valve morphogenesis | GO:0003179 | GO:BP | 0.005355329 |
| response to radiation | GO:0009314 | GO:BP | 0.005355329 |
| cellular detoxification | GO:1990748 | GO:BP | 0.005521457 |
| cell morphogenesis involved in neuron differentiation | GO:0048667 | GO:BP | 0.005521457 |
| axonogenesis | GO:0007409 | GO:BP | 0.00552298 |
| hemopoiesis | GO:0030097 | GO:BP | 0.00552298 |
| monosaccharide catabolic process | GO:0046365 | GO:BP | 0.005585291 |
| antigen processing and presentation of peptide antigen | GO:0048002 | GO:BP | 0.005677043 |
| calcium ion transmembrane transport | GO:0070588 | GO:BP | 0.005718881 |
| gland morphogenesis | GO:0022612 | GO:BP | 0.005719487 |
| long-term synaptic potentiation | GO:0060291 | GO:BP | 0.005811794 |
| establishment of localization in cell | GO:0051649 | GO:BP | 0.00581712 |
| behavior | GO:0007610 | GO:BP | 0.00581712 |
| positive regulation of immune response | GO:0050778 | GO:BP | 0.005830955 |
| regulation of mitochondrion organization | GO:0010821 | GO:BP | 0.005908966 |
| positive regulation of cellular component organization | GO:0051130 | GO:BP | 0.005908966 |
| negative regulation of cytokine production | GO:0001818 | GO:BP | 0.006282278 |
| MAPK cascade | GO:0000165 | GO:BP | 0.006282278 |
| hepatocyte proliferation | GO:0072574 | GO:BP | 0.006382302 |
| epithelial cell proliferation involved in liver morphogenesis | GO:0072575 | GO:BP | 0.006382302 |
| response to salt stress | GO:0009651 | GO:BP | 0.006382302 |
| cellular response to oxidative stress | GO:0034599 | GO:BP | 0.006382302 |
| cellular response to biotic stimulus | GO:0071216 | GO:BP | 0.006390114 |
| modulation of chemical synaptic transmission | GO:0050804 | GO:BP | 0.006450715 |
| negative regulation of endoplasmic reticulum unfolded prote | GO:1900102 | GO:BP | 0.006813955 |
| positive regulation of interleukin-6 production | GO:0032755 | GO:BP | 0.006850554 |
| regulation of cysteine-type endopeptidase activity involved i | GO:0043281 | GO:BP | 0.006919833 |
| CD4-positive, alpha-beta T cell activation | GO:0035710 | GO:BP | 0.006925445 |
| glycerophospholipid catabolic process | GO:0046475 | GO:BP | 0.00695457 |
| regulation of trans-synaptic signaling | GO:0099177 | GO:BP | 0.006998637 |
| visual perception | GO:0007601 | GO:BP | 0.00702346 |
| positive regulation of synaptic transmission | GO:0050806 | GO:BP | 0.007033098 |
| blood circulation | GO:0008015 | GO:BP | 0.007033098 |
| lipid storage | GO:0019915 | GO:BP | 0.007040293 |
| regulation of protein maturation | GO:1903317 | GO:BP | 0.007126014 |

| | | | |
|---|---|---|---|
| non-motile cilium assembly | GO:1905515 | GO:BP | 0.007126014 |
| regulation of leukocyte mediated immunity | GO:0002703 | GO:BP | 0.007126014 |
| regulation of mononuclear cell migration | GO:0071675 | GO:BP | 0.007206231 |
| regulation of immunoglobulin production | GO:0002637 | GO:BP | 0.007208878 |
| detoxification | GO:0098754 | GO:BP | 0.007208878 |
| cell growth | GO:0016049 | GO:BP | 0.007223874 |
| leukocyte differentiation | GO:0002521 | GO:BP | 0.007369271 |
| response to xenobiotic stimulus | GO:0009410 | GO:BP | 0.007390735 |
| renal system process | GO:0003014 | GO:BP | 0.007515714 |
| CD4-positive, alpha-beta T cell differentiation | GO:0043367 | GO:BP | 0.007597383 |
| multi-organism reproductive process | GO:0044703 | GO:BP | 0.007643729 |
| cellular response to reactive oxygen species | GO:0034614 | GO:BP | 0.007691958 |
| organic substance transport | GO:0071702 | GO:BP | 0.007711507 |
| negative regulation of stress-activated MAPK cascade | GO:0032873 | GO:BP | 0.007711507 |
| daunorubicin metabolic process | GO:0044597 | GO:BP | 0.007711507 |
| protein secretion | GO:0009306 | GO:BP | 0.007725572 |
| positive regulation of B cell proliferation | GO:0030890 | GO:BP | 0.007918401 |
| skeletal muscle contraction | GO:0003009 | GO:BP | 0.007918401 |
| tissue morphogenesis | GO:0048729 | GO:BP | 0.007918401 |
| regulation of transferase activity | GO:0051338 | GO:BP | 0.007980315 |
| response to light stimulus | GO:0009416 | GO:BP | 0.007980315 |
| autophagy | GO:0006914 | GO:BP | 0.007997787 |
| process utilizing autophagic mechanism | GO:0061919 | GO:BP | 0.007997787 |
| signal release | GO:0023061 | GO:BP | 0.007997787 |
| regulation of steroid metabolic process | GO:0019218 | GO:BP | 0.008072842 |
| fatty acid oxidation | GO:0019395 | GO:BP | 0.008130461 |
| chemosensory behavior | GO:0007635 | GO:BP | 0.008140235 |
| cell morphogenesis | GO:0000902 | GO:BP | 0.008274123 |
| morphogenesis of an epithelium | GO:0002009 | GO:BP | 0.00831536 |
| developmental process involved in reproduction | GO:0003006 | GO:BP | 0.008321793 |
| skeletal system development | GO:0001501 | GO:BP | 0.008327809 |
| establishment of protein localization to extracellular region | GO:0035592 | GO:BP | 0.008358521 |
| regulation of fibroblast proliferation | GO:0048145 | GO:BP | 0.008364295 |
| response to vitamin | GO:0033273 | GO:BP | 0.008364295 |
| positive regulation of cysteine-type endopeptidase activity ir | GO:0043280 | GO:BP | 0.008584142 |
| multi-multicellular organism process | GO:0044706 | GO:BP | 0.008594055 |
| regulation of monoatomic ion transport | GO:0043269 | GO:BP | 0.008703551 |
| endoplasmic reticulum unfolded protein response | GO:0030968 | GO:BP | 0.008798969 |
| positive regulation of immunoglobulin production | GO:0002639 | GO:BP | 0.008823512 |
| tertiary alcohol metabolic process | GO:1902644 | GO:BP | 0.009153243 |
| circulatory system process | GO:0003013 | GO:BP | 0.009286848 |
| negative regulation of cell growth | GO:0030308 | GO:BP | 0.009333001 |
| diterpenoid metabolic process | GO:0016101 | GO:BP | 0.009446019 |
| bicarbonate transport | GO:0015701 | GO:BP | 0.009765385 |
| sensory perception of light stimulus | GO:0050953 | GO:BP | 0.009805535 |
| regulation of leukocyte migration | GO:0002685 | GO:BP | 0.009819915 |
| heart valve development | GO:0003170 | GO:BP | 0.009877986 |
| organelle disassembly | GO:1903008 | GO:BP | 0.009969305 |

| | | | |
|---|---|---|---|
| antigen processing and presentation of exogenous peptide a | GO:0019886 | GO:BP | 0.009969305 |
| positive regulation of granulocyte chemotaxis | GO:0071624 | GO:BP | 0.009969305 |
| mitochondrion organization | GO:0007005 | GO:BP | 0.009988275 |
| negative regulation of transport | GO:0051051 | GO:BP | 0.010066107 |
| response to lipopolysaccharide | GO:0032496 | GO:BP | 0.010070645 |
| regulation of protein catabolic process | GO:0042176 | GO:BP | 0.010087264 |
| connective tissue development | GO:0061448 | GO:BP | 0.01018889 |
| nucleoside diphosphate catabolic process | GO:0009134 | GO:BP | 0.010245075 |
| response to fatty acid | GO:0070542 | GO:BP | 0.010398697 |
| positive regulation of hemopoiesis | GO:1903708 | GO:BP | 0.010398697 |
| positive regulation of leukocyte differentiation | GO:1902107 | GO:BP | 0.010398697 |
| antigen processing and presentation of exogenous antigen | GO:0019884 | GO:BP | 0.010398697 |
| regulation of cytokine production involved in immune respor | GO:0002718 | GO:BP | 0.010518006 |
| cytokine production involved in immune response | GO:0002367 | GO:BP | 0.010518006 |
| axon ensheathment | GO:0008366 | GO:BP | 0.01052328 |
| ensheathment of neurons | GO:0007272 | GO:BP | 0.01052328 |
| positive regulation of transcription by RNA polymerase II | GO:0045944 | GO:BP | 0.010588297 |
| cell projection morphogenesis | GO:0048858 | GO:BP | 0.010588297 |
| fatty acid catabolic process | GO:0009062 | GO:BP | 0.010744386 |
| alpha-amino acid metabolic process | GO:1901605 | GO:BP | 0.010759699 |
| regulation of myeloid cell differentiation | GO:0045637 | GO:BP | 0.010759699 |
| transmembrane receptor protein tyrosine kinase signaling pa | GO:0007169 | GO:BP | 0.010838764 |
| ribonucleoside diphosphate metabolic process | GO:0009185 | GO:BP | 0.010885074 |
| hexose catabolic process | GO:0019320 | GO:BP | 0.011161451 |
| mononuclear cell differentiation | GO:1903131 | GO:BP | 0.01122759 |
| response to amino acid starvation | GO:1990928 | GO:BP | 0.011356443 |
| cardiocyte differentiation | GO:0035051 | GO:BP | 0.011605592 |
| exocytosis | GO:0006887 | GO:BP | 0.011620993 |
| regulation of hepatocyte proliferation | GO:2000345 | GO:BP | 0.011620993 |
| MHC protein complex assembly | GO:0002396 | GO:BP | 0.011620993 |
| axon development | GO:0061564 | GO:BP | 0.011620993 |
| peptide antigen assembly with MHC protein complex | GO:0002501 | GO:BP | 0.011620993 |
| cellular response to metal ion | GO:0071248 | GO:BP | 0.011928089 |
| negative regulation of post-translational protein modificatio | GO:1901874 | GO:BP | 0.012063701 |
| regulation of chemotaxis | GO:0050920 | GO:BP | 0.012063701 |
| production of molecular mediator of immune response | GO:0002440 | GO:BP | 0.012063701 |
| negative regulation of phosphate metabolic process | GO:0045936 | GO:BP | 0.012387666 |
| cellular response to amino acid starvation | GO:0034198 | GO:BP | 0.012416928 |
| sensory perception of sound | GO:0007605 | GO:BP | 0.012416928 |
| cellular response to toxic substance | GO:0097237 | GO:BP | 0.012767842 |
| autophagy of mitochondrion | GO:0000422 | GO:BP | 0.012886701 |
| glycolytic process through glucose-6-phosphate | GO:0061620 | GO:BP | 0.013314054 |
| visual system development | GO:0150063 | GO:BP | 0.013358395 |
| rhythmic process | GO:0048511 | GO:BP | 0.013416578 |
| endomembrane system organization | GO:0010256 | GO:BP | 0.013435615 |
| toll-like receptor signaling pathway | GO:0002224 | GO:BP | 0.013462563 |
| negative regulation of phosphorus metabolic process | GO:0010563 | GO:BP | 0.013515671 |
| myelination | GO:0042552 | GO:BP | 0.013553787 |

| | | | |
|---|---|---|---|
| prostaglandin metabolic process | GO:0006693 | GO:BP | 0.013578609 |
| cellular response to UV | GO:0034644 | GO:BP | 0.013578609 |
| antigen processing and presentation of peptide antigen via N | GO:0002495 | GO:BP | 0.014040973 |
| piRNA processing | GO:0034587 | GO:BP | 0.014159816 |
| cellular oxidant detoxification | GO:0098869 | GO:BP | 0.014565811 |
| regulation of cell growth | GO:0001558 | GO:BP | 0.01464917 |
| leukocyte cell-cell adhesion | GO:0007159 | GO:BP | 0.014703348 |
| fatty acid beta-oxidation | GO:0006635 | GO:BP | 0.014802445 |
| positive regulation of muscle contraction | GO:0045933 | GO:BP | 0.014842803 |
| cellular response to light stimulus | GO:0071482 | GO:BP | 0.014883563 |
| positive regulation of lymphocyte mediated immunity | GO:0002708 | GO:BP | 0.014883563 |
| response to purine-containing compound | GO:0014074 | GO:BP | 0.014956157 |
| long-chain fatty acid metabolic process | GO:0001676 | GO:BP | 0.014956157 |
| response to calcium ion | GO:0051592 | GO:BP | 0.014956157 |
| regulation of protein stability | GO:0031647 | GO:BP | 0.014963155 |
| gland development | GO:0048732 | GO:BP | 0.014963155 |
| cellular response to organic cyclic compound | GO:0071407 | GO:BP | 0.014965795 |
| protein localization to extracellular region | GO:0071692 | GO:BP | 0.014974499 |
| macromolecule localization | GO:0033036 | GO:BP | 0.015196466 |
| regulation of nervous system development | GO:0051960 | GO:BP | 0.01556298 |
| cellular component disassembly | GO:0022411 | GO:BP | 0.015939802 |
| positive regulation of CD4-positive, alpha-beta T cell activati | GO:2000516 | GO:BP | 0.016014083 |
| positive regulation of molecular function | GO:0044093 | GO:BP | 0.016052169 |
| cellular response to unfolded protein | GO:0034620 | GO:BP | 0.016304971 |
| regulation of post-translational protein modification | GO:1901873 | GO:BP | 0.016348592 |
| regulation of phosphatidylinositol 3-kinase/protein kinase B | GO:0051896 | GO:BP | 0.016348592 |
| biomineral tissue development | GO:0031214 | GO:BP | 0.016504012 |
| regulation of body fluid levels | GO:0050878 | GO:BP | 0.016571293 |
| ribonucleotide catabolic process | GO:0009261 | GO:BP | 0.016583906 |
| positive regulation of chemotaxis | GO:0050921 | GO:BP | 0.016583906 |
| telomere organization | GO:0032200 | GO:BP | 0.016583906 |
| positive regulation of interleukin-13 production | GO:0032736 | GO:BP | 0.016583906 |
| intrinsic apoptotic signaling pathway in response to DNA dar | GO:0008630 | GO:BP | 0.016583906 |
| regulation of type B pancreatic cell proliferation | GO:0061469 | GO:BP | 0.016583906 |
| regulation of lymphocyte apoptotic process | GO:0070228 | GO:BP | 0.016583906 |
| sulfate transmembrane transport | GO:1902358 | GO:BP | 0.016583906 |
| regulated exocytosis | GO:0045055 | GO:BP | 0.016583906 |
| endoderm formation | GO:0001706 | GO:BP | 0.016583906 |
| induction of positive chemotaxis | GO:0050930 | GO:BP | 0.016583906 |
| intrinsic apoptotic signaling pathway in response to endopla | GO:0070059 | GO:BP | 0.016583906 |
| negative regulation of glycoprotein metabolic process | GO:1903019 | GO:BP | 0.016583906 |
| positive regulation of cell adhesion | GO:0045785 | GO:BP | 0.016780132 |
| plasma membrane bounded cell projection morphogenesis | GO:0120039 | GO:BP | 0.016950888 |
| positive regulation of leukocyte chemotaxis | GO:0002690 | GO:BP | 0.016950888 |
| response to tumor cell | GO:0002347 | GO:BP | 0.016950888 |
| doxorubicin metabolic process | GO:0044598 | GO:BP | 0.01725806 |
| negative regulation of stress-activated protein kinase signali | GO:0070303 | GO:BP | 0.01725806 |
| aminoglycoside antibiotic metabolic process | GO:0030647 | GO:BP | 0.01725806 |

| | | | |
|---|---|---|---|
| polyketide metabolic process | GO:0030638 | GO:BP | 0.01725806 |
| symbiont entry into host cell | GO:0046718 | GO:BP | 0.017267561 |
| prostanoid metabolic process | GO:0006692 | GO:BP | 0.017267561 |
| nucleoside diphosphate metabolic process | GO:0009132 | GO:BP | 0.017321135 |
| regulation of system process | GO:0044057 | GO:BP | 0.017641919 |
| phosphatidylinositol 3-kinase/protein kinase B signal transd | GO:0043491 | GO:BP | 0.017641919 |
| T cell activation | GO:0042110 | GO:BP | 0.017756057 |
| regulation of cellular ketone metabolic process | GO:0010565 | GO:BP | 0.018179313 |
| purine ribonucleotide catabolic process | GO:0009154 | GO:BP | 0.018179313 |
| cellular response to molecule of bacterial origin | GO:0071219 | GO:BP | 0.018276041 |
| response to hormone | GO:0009725 | GO:BP | 0.018276041 |
| activation of cysteine-type endopeptidase activity involved i | GO:0006919 | GO:BP | 0.018276041 |
| neuron projection morphogenesis | GO:0048812 | GO:BP | 0.018301492 |
| eye development | GO:0001654 | GO:BP | 0.018343992 |
| negative regulation of cell activation | GO:0050866 | GO:BP | 0.018395494 |
| regulation of myelination | GO:0031641 | GO:BP | 0.018845309 |
| positive regulation of cytokine production involved in immun | GO:0002720 | GO:BP | 0.018845309 |
| extracellular transport | GO:0006858 | GO:BP | 0.018845309 |
| canonical glycolysis | GO:0061621 | GO:BP | 0.018852968 |
| glucose catabolic process to pyruvate | GO:0061718 | GO:BP | 0.018852968 |
| NADH regeneration | GO:0006735 | GO:BP | 0.018852968 |
| regulation of interleukin-4 production | GO:0032673 | GO:BP | 0.018886125 |
| ligand-gated ion channel signaling pathway | GO:1990806 | GO:BP | 0.018886125 |
| hyaluronan metabolic process | GO:0030212 | GO:BP | 0.018886125 |
| interleukin-4 production | GO:0032633 | GO:BP | 0.018886125 |
| regulation of organelle organization | GO:0033043 | GO:BP | 0.019170984 |
| axon guidance | GO:0007411 | GO:BP | 0.019378652 |
| neuron projection guidance | GO:0097485 | GO:BP | 0.019378652 |
| vesicle-mediated transport | GO:0016192 | GO:BP | 0.019382352 |
| cellular response to mechanical stimulus | GO:0071260 | GO:BP | 0.019473154 |
| regulation of protein localization | GO:0032880 | GO:BP | 0.019798718 |
| retinol metabolic process | GO:0042572 | GO:BP | 0.019798718 |
| hormone biosynthetic process | GO:0042446 | GO:BP | 0.0200291 |
| NAD catabolic process | GO:0019677 | GO:BP | 0.020084694 |
| microtubule-based transport | GO:0099111 | GO:BP | 0.020334801 |
| photoperiodism | GO:0009648 | GO:BP | 0.0203374 |
| plasminogen activation | GO:0031639 | GO:BP | 0.0203374 |
| epithelial cilium movement involved in extracellular fluid mc | GO:0003351 | GO:BP | 0.020357884 |
| negative regulation of proteolysis | GO:0045861 | GO:BP | 0.020357884 |
| sensory system development | GO:0048880 | GO:BP | 0.02039019 |
| ionotropic glutamate receptor signaling pathway | GO:0035235 | GO:BP | 0.020406063 |
| entrainment of circadian clock by photoperiod | GO:0043153 | GO:BP | 0.020406063 |
| regulation of leukocyte proliferation | GO:0070663 | GO:BP | 0.020463354 |
| glycerolipid metabolic process | GO:0046486 | GO:BP | 0.020524317 |
| sequestering of calcium ion | GO:0051208 | GO:BP | 0.020829607 |
| sulfur compound metabolic process | GO:0006790 | GO:BP | 0.020983636 |
| amine metabolic process | GO:0009308 | GO:BP | 0.021026417 |
| regulation of response to biotic stimulus | GO:0002831 | GO:BP | 0.02105783 |

| | | | |
|---|---|---|---|
| positive regulation of leukocyte mediated immunity | GO:0002705 | GO:BP | 0.021099552 |
| sensory organ development | GO:0007423 | GO:BP | 0.021249699 |
| intracellular chemical homeostasis | GO:0055082 | GO:BP | 0.021659055 |
| response to toxic substance | GO:0009636 | GO:BP | 0.021699086 |
| glutamate receptor signaling pathway | GO:0007215 | GO:BP | 0.021702368 |
| fatty acid biosynthetic process | GO:0006633 | GO:BP | 0.021860449 |
| mitophagy | GO:0000423 | GO:BP | 0.022024409 |
| positive regulation of plasminogen activation | GO:0010756 | GO:BP | 0.022182154 |
| regulation of skeletal muscle tissue regeneration | GO:0043416 | GO:BP | 0.022182154 |
| atrioventricular valve formation | GO:0003190 | GO:BP | 0.022182154 |
| keratinocyte proliferation | GO:0043616 | GO:BP | 0.022191453 |
| sensory perception of pain | GO:0019233 | GO:BP | 0.022191453 |
| spermatid differentiation | GO:0048515 | GO:BP | 0.022205074 |
| regulation of lymphocyte mediated immunity | GO:0002706 | GO:BP | 0.022405582 |
| cell projection assembly | GO:0030031 | GO:BP | 0.022405582 |
| response to organophosphorus | GO:0046683 | GO:BP | 0.022405582 |
| plasma membrane bounded cell projection assembly | GO:0120031 | GO:BP | 0.022405582 |
| female pregnancy | GO:0007565 | GO:BP | 0.022405582 |
| negative regulation of response to external stimulus | GO:0032102 | GO:BP | 0.022454435 |
| amino acid catabolic process | GO:0009063 | GO:BP | 0.022777293 |
| homotypic cell-cell adhesion | GO:0034109 | GO:BP | 0.022975074 |
| fat-soluble vitamin biosynthetic process | GO:0042362 | GO:BP | 0.023069752 |
| bone trabecula morphogenesis | GO:0061430 | GO:BP | 0.023069752 |
| positive regulation of striated muscle contraction | GO:0045989 | GO:BP | 0.023069752 |
| CD4-positive, alpha-beta T cell differentiation involved in im | GO:0002294 | GO:BP | 0.023142441 |
| regulation of CD4-positive, alpha-beta T cell activation | GO:2000514 | GO:BP | 0.023142441 |
| serine family amino acid metabolic process | GO:0009069 | GO:BP | 0.023537908 |
| negative regulation of striated muscle cell differentiation | GO:0051154 | GO:BP | 0.023537908 |
| cellular homeostasis | GO:0019725 | GO:BP | 0.023586799 |
| microglial cell activation | GO:0001774 | GO:BP | 0.023603401 |
| intrinsic apoptotic signaling pathway in response to DNA dar | GO:0042771 | GO:BP | 0.023603401 |
| leukocyte proliferation | GO:0070661 | GO:BP | 0.023849138 |
| positive regulation of adaptive immune response based on s | GO:0002824 | GO:BP | 0.02395561 |
| regulation of developmental growth | GO:0048638 | GO:BP | 0.024527056 |
| regulation of extracellular matrix organization | GO:1903053 | GO:BP | 0.024536874 |
| symbiont entry into host | GO:0044409 | GO:BP | 0.024539824 |
| calcium ion import | GO:0070509 | GO:BP | 0.024539824 |
| T cell mediated immunity | GO:0002456 | GO:BP | 0.024539824 |
| biological process involved in interaction with host | GO:0051701 | GO:BP | 0.024608115 |
| response to nutrient | GO:0007584 | GO:BP | 0.024639996 |
| neural precursor cell proliferation | GO:0061351 | GO:BP | 0.024739123 |
| skeletal muscle organ development | GO:0060538 | GO:BP | 0.024912359 |
| response to fluid shear stress | GO:0034405 | GO:BP | 0.025087687 |
| cellular response to interleukin-4 | GO:0071353 | GO:BP | 0.025087687 |
| establishment of skin barrier | GO:0061436 | GO:BP | 0.025087687 |
| regulation of signal transduction by p53 class mediator | GO:1901796 | GO:BP | 0.025433469 |
| membrane organization | GO:0061024 | GO:BP | 0.025526741 |
| glyceraldehyde-3-phosphate biosynthetic process | GO:0046166 | GO:BP | 0.025957277 |

| | | | |
|---|---|---|---|
| telomere assembly | GO:0032202 | GO:BP | 0.025957277 |
| pentose-phosphate shunt, oxidative branch | GO:0009051 | GO:BP | 0.025957277 |
| lipid catabolic process | GO:0016042 | GO:BP | 0.026582996 |
| G1 to G0 transition involved in cell differentiation | GO:0070315 | GO:BP | 0.026582996 |
| regulation of glycogen catabolic process | GO:0005981 | GO:BP | 0.026582996 |
| negative regulation of cellular component organization | GO:0051129 | GO:BP | 0.026674885 |
| regulation of endoplasmic reticulum unfolded protein respo | GO:1900101 | GO:BP | 0.026691541 |
| negative regulation of secretion by cell | GO:1903531 | GO:BP | 0.026698286 |
| short-chain fatty acid metabolic process | GO:0046459 | GO:BP | 0.026698286 |
| sulfate transport | GO:0008272 | GO:BP | 0.026698286 |
| regulation of protein secretion | GO:0050708 | GO:BP | 0.026698286 |
| amide transport | GO:0042886 | GO:BP | 0.026710181 |
| cellular response to glucose starvation | GO:0042149 | GO:BP | 0.026884904 |
| negative regulation of secretion | GO:0051048 | GO:BP | 0.027588302 |
| alpha-beta T cell differentiation involved in immune respons | GO:0002293 | GO:BP | 0.027588302 |
| alpha-beta T cell activation involved in immune response | GO:0002287 | GO:BP | 0.027588302 |
| regulation of T-helper cell differentiation | GO:0045622 | GO:BP | 0.028319649 |
| lymphocyte differentiation | GO:0030098 | GO:BP | 0.028607147 |
| renal tubule development | GO:0061326 | GO:BP | 0.028607147 |
| regulation of supramolecular fiber organization | GO:1902903 | GO:BP | 0.028678949 |
| regulation of lymphocyte proliferation | GO:0050670 | GO:BP | 0.028731684 |
| peripheral nervous system development | GO:0007422 | GO:BP | 0.028800769 |
| positive regulation of neutrophil chemotaxis | GO:0090023 | GO:BP | 0.028895134 |
| vitamin biosynthetic process | GO:0009110 | GO:BP | 0.028895134 |
| decidualization | GO:0046697 | GO:BP | 0.028895134 |
| peptidyl-amino acid modification | GO:0018193 | GO:BP | 0.028903183 |
| regulation of ketone biosynthetic process | GO:0010566 | GO:BP | 0.029235776 |
| fibroblast proliferation | GO:0048144 | GO:BP | 0.029305214 |
| positive regulation of secretion | GO:0051047 | GO:BP | 0.029417063 |
| positive regulation of secretion by cell | GO:1903532 | GO:BP | 0.029482687 |
| endochondral bone morphogenesis | GO:0060350 | GO:BP | 0.029827782 |
| regulation of G protein-coupled receptor signaling pathway | GO:0008277 | GO:BP | 0.029888799 |
| glial cell activation | GO:0061900 | GO:BP | 0.030198452 |
| regulation of neuronal synaptic plasticity | GO:0048168 | GO:BP | 0.030198452 |
| regulation of endocytosis | GO:0030100 | GO:BP | 0.030276462 |
| aortic valve morphogenesis | GO:0003180 | GO:BP | 0.030755067 |
| T-helper cell differentiation | GO:0042093 | GO:BP | 0.03095837 |
| response to activity | GO:0014823 | GO:BP | 0.03095837 |
| positive regulation of lipid metabolic process | GO:0045834 | GO:BP | 0.031409898 |
| regulation of plasma membrane bounded cell projection org | GO:0120035 | GO:BP | 0.031458433 |
| myeloid leukocyte differentiation | GO:0002573 | GO:BP | 0.03146669 |
| regulation of neuron projection development | GO:0010975 | GO:BP | 0.031720364 |
| epithelial cell differentiation | GO:0030855 | GO:BP | 0.031736755 |
| positive regulation of adaptive immune response | GO:0002821 | GO:BP | 0.031894499 |
| regulation of kinase activity | GO:0043549 | GO:BP | 0.032139436 |
| positive regulation of T cell mediated immunity | GO:0002711 | GO:BP | 0.032282102 |
| regulation of B cell proliferation | GO:0030888 | GO:BP | 0.032282102 |
| maintenance of protein location in cell | GO:0032507 | GO:BP | 0.032282102 |

| | | | |
|---|---|---|---|
| positive regulation of reactive oxygen species metabolic pro | GO:2000379 | GO:BP | 0.032282102 |
| glomerular basement membrane development | GO:0032836 | GO:BP | 0.032346832 |
| negative regulation of cardiocyte differentiation | GO:1905208 | GO:BP | 0.032346832 |
| protein folding in endoplasmic reticulum | GO:0034975 | GO:BP | 0.032346832 |
| steroid hormone biosynthetic process | GO:0120178 | GO:BP | 0.03273973 |
| sulfur compound catabolic process | GO:0044273 | GO:BP | 0.03273973 |
| cellular response to lipopolysaccharide | GO:0071222 | GO:BP | 0.032922623 |
| negative regulation of fibroblast proliferation | GO:0048147 | GO:BP | 0.032974598 |
| regulation of sequestering of calcium ion | GO:0051282 | GO:BP | 0.032974598 |
| regulation of interleukin-10 production | GO:0032653 | GO:BP | 0.032974598 |
| musculoskeletal movement | GO:0050881 | GO:BP | 0.032974598 |
| cartilage development involved in endochondral bone morpl | GO:0060351 | GO:BP | 0.032974598 |
| interleukin-10 production | GO:0032613 | GO:BP | 0.032974598 |
| regulation of B cell differentiation | GO:0045577 | GO:BP | 0.032974598 |
| purine nucleotide catabolic process | GO:0006195 | GO:BP | 0.033418592 |
| liver development | GO:0001889 | GO:BP | 0.033418592 |
| neuron apoptotic process | GO:0051402 | GO:BP | 0.033511719 |
| response to biotic stimulus | GO:0009607 | GO:BP | 0.033897043 |
| negative regulation of leukocyte activation | GO:0002695 | GO:BP | 0.033943002 |
| lymphocyte apoptotic process | GO:0070227 | GO:BP | 0.034375315 |
| circadian regulation of gene expression | GO:0032922 | GO:BP | 0.034413975 |
| positive regulation of calcium ion transport | GO:0051928 | GO:BP | 0.034930615 |
| monoatomic cation homeostasis | GO:0055080 | GO:BP | 0.034948424 |
| positive regulation of catalytic activity | GO:0043085 | GO:BP | 0.034971326 |
| positive regulation of response to biotic stimulus | GO:0002833 | GO:BP | 0.035085672 |
| response to alcohol | GO:0097305 | GO:BP | 0.035309158 |
| negative regulation of cell adhesion | GO:0007162 | GO:BP | 0.035514739 |
| aromatic amino acid metabolic process | GO:0009072 | GO:BP | 0.035514739 |
| response to hydrogen peroxide | GO:0042542 | GO:BP | 0.035641127 |
| NADH dehydrogenase complex assembly | GO:0010257 | GO:BP | 0.035807851 |
| maintenance of protein localization in organelle | GO:0072595 | GO:BP | 0.035807851 |
| multicellular organismal reproductive process | GO:0048609 | GO:BP | 0.035807851 |
| cardiac cell development | GO:0055006 | GO:BP | 0.035807851 |
| B cell activation | GO:0042113 | GO:BP | 0.035807851 |
| mitochondrial respiratory chain complex I assembly | GO:0032981 | GO:BP | 0.035807851 |
| protein stabilization | GO:0050821 | GO:BP | 0.035879738 |
| apoptotic mitochondrial changes | GO:0008637 | GO:BP | 0.036558028 |
| regulation of leukocyte apoptotic process | GO:2000106 | GO:BP | 0.037424147 |
| cholesterol homeostasis | GO:0042632 | GO:BP | 0.037424147 |
| negative regulation of protein modification by small protein ( | GO:1903321 | GO:BP | 0.037424147 |
| negative regulation of developmental growth | GO:0048640 | GO:BP | 0.037511667 |
| positive regulation of protein secretion | GO:0050714 | GO:BP | 0.037511667 |
| negative regulation of DNA-binding transcription factor activ | GO:0043433 | GO:BP | 0.037511667 |
| cerebellar Purkinje cell layer development | GO:0021680 | GO:BP | 0.037646864 |
| detection of stimulus involved in sensory perception of pain | GO:0062149 | GO:BP | 0.037646864 |
| purine-containing compound catabolic process | GO:0072523 | GO:BP | 0.037646864 |
| atrioventricular valve development | GO:0003171 | GO:BP | 0.037646864 |
| cellular modified amino acid catabolic process | GO:0042219 | GO:BP | 0.037646864 |

| | | | |
|---|---|---|---|
| negative regulation of hydrolase activity | GO:0051346 | GO:BP | 0.03781621 |
| heart development | GO:0007507 | GO:BP | 0.038015287 |
| muscle filament sliding | GO:0030049 | GO:BP | 0.038015287 |
| positive regulation of autophagy of mitochondrion in respons | GO:1904925 | GO:BP | 0.038015287 |
| lipid droplet formation | GO:0140042 | GO:BP | 0.038015287 |
| cellular response to salt stress | GO:0071472 | GO:BP | 0.038015287 |
| renal system development | GO:0072001 | GO:BP | 0.038140939 |
| extrinsic apoptotic signaling pathway | GO:0097191 | GO:BP | 0.038140939 |
| bone remodeling | GO:0046849 | GO:BP | 0.038390754 |
| regulation of nervous system process | GO:0031644 | GO:BP | 0.038691235 |
| regulation of intrinsic apoptotic signaling pathway in respons | GO:1902229 | GO:BP | 0.038988947 |
| positive regulation of protein transport | GO:0051222 | GO:BP | 0.038999315 |
| atrioventricular valve morphogenesis | GO:0003181 | GO:BP | 0.039625511 |
| positive regulation of interleukin-4 production | GO:0032753 | GO:BP | 0.039625511 |
| lipid transport | GO:0006869 | GO:BP | 0.039669322 |
| multicellular organismal movement | GO:0050879 | GO:BP | 0.039762561 |
| regulation of anatomical structure morphogenesis | GO:0022603 | GO:BP | 0.039766405 |
| regulation of neurogenesis | GO:0050767 | GO:BP | 0.039829673 |
| cell adhesion mediated by integrin | GO:0033627 | GO:BP | 0.039922291 |
| carbohydrate biosynthetic process | GO:0016051 | GO:BP | 0.040131438 |
| regulation of mononuclear cell proliferation | GO:0032944 | GO:BP | 0.040364676 |
| glutathione metabolic process | GO:0006749 | GO:BP | 0.040647064 |
| positive regulation of lymphocyte differentiation | GO:0045621 | GO:BP | 0.040705481 |
| endothelial tube morphogenesis | GO:0061154 | GO:BP | 0.040859773 |
| morphogenesis of an endothelium | GO:0003159 | GO:BP | 0.040859773 |
| negative regulation of sequestering of calcium ion | GO:0051283 | GO:BP | 0.040957554 |
| negative regulation of keratinocyte proliferation | GO:0010839 | GO:BP | 0.040957554 |
| regulation of cellular localization | GO:0060341 | GO:BP | 0.04114124 |
| developmental cell growth | GO:0048588 | GO:BP | 0.041145527 |
| temperature homeostasis | GO:0001659 | GO:BP | 0.041219823 |
| regulation of protein transport | GO:0051223 | GO:BP | 0.041231224 |
| positive regulation of lymphocyte proliferation | GO:0050671 | GO:BP | 0.041365038 |
| sexual reproduction | GO:0019953 | GO:BP | 0.041387713 |
| lactate metabolic process | GO:0006089 | GO:BP | 0.041387713 |
| muscle adaptation | GO:0043500 | GO:BP | 0.041387713 |
| regulation of neuron apoptotic process | GO:0043523 | GO:BP | 0.041387713 |
| positive regulation of leukocyte proliferation | GO:0070665 | GO:BP | 0.041619672 |
| antigen processing and presentation of peptide or polysacch | GO:0002504 | GO:BP | 0.042315471 |
| glycerophospholipid metabolic process | GO:0006650 | GO:BP | 0.042370021 |
| regulation of cell projection organization | GO:0031344 | GO:BP | 0.042428649 |
| epithelial tube morphogenesis | GO:0060562 | GO:BP | 0.042439584 |
| immune effector process | GO:0002252 | GO:BP | 0.042777605 |
| sterol homeostasis | GO:0055092 | GO:BP | 0.042780548 |
| hormone secretion | GO:0046879 | GO:BP | 0.042809558 |
| regulation of lymphocyte differentiation | GO:0045619 | GO:BP | 0.043054654 |
| regulation of protein modification by small protein conjugati | GO:1903320 | GO:BP | 0.043757322 |
| leukocyte apoptotic process | GO:0071887 | GO:BP | 0.043757322 |
| insulin secretion | GO:0030073 | GO:BP | 0.043765755 |

| | | | |
|---|---|---|---|
| circadian rhythm | GO:0007623 | GO:BP | 0.043765755 |
| mononuclear cell migration | GO:0071674 | GO:BP | 0.043765755 |
| hepoxilin biosynthetic process | GO:0051122 | GO:BP | 0.043879632 |
| myeloid leukocyte activation | GO:0002274 | GO:BP | 0.043879632 |
| respiratory burst involved in inflammatory response | GO:0002536 | GO:BP | 0.043879632 |
| regulation of cilium beat frequency involved in ciliary motility | GO:0060296 | GO:BP | 0.043879632 |
| dehydroascorbic acid transport | GO:0070837 | GO:BP | 0.043879632 |
| fructose 1,6-bisphosphate metabolic process | GO:0030388 | GO:BP | 0.043879632 |
| cellular response to growth factor stimulus | GO:0071363 | GO:BP | 0.043879632 |
| glial cell-derived neurotrophic factor receptor signaling path | GO:0035860 | GO:BP | 0.043879632 |
| hepoxilin metabolic process | GO:0051121 | GO:BP | 0.043879632 |
| canonical NF-kappaB signal transduction | GO:0007249 | GO:BP | 0.043879632 |
| aortic valve development | GO:0003176 | GO:BP | 0.044053251 |
| calcium ion transmembrane import into cytosol | GO:0097553 | GO:BP | 0.044053251 |
| regulation of protein targeting | GO:1903533 | GO:BP | 0.044053251 |
| regulation of transmembrane receptor protein serine/threon | GO:0090092 | GO:BP | 0.044059931 |
| memory | GO:0007613 | GO:BP | 0.044228036 |
| positive regulation of cell-substrate adhesion | GO:0010811 | GO:BP | 0.044228036 |
| leukocyte activation involved in inflammatory response | GO:0002269 | GO:BP | 0.044360133 |
| regulation of transmembrane transport | GO:0034762 | GO:BP | 0.045282401 |
| myeloid cell homeostasis | GO:0002262 | GO:BP | 0.045398757 |
| hepaticobiliary system development | GO:0061008 | GO:BP | 0.045928793 |
| monoatomic ion homeostasis | GO:0050801 | GO:BP | 0.045928793 |
| cellular response to topologically incorrect protein | GO:0035967 | GO:BP | 0.045966026 |
| macrophage activation | GO:0042116 | GO:BP | 0.047058361 |
| cellular response to alcohol | GO:0097306 | GO:BP | 0.047058361 |
| positive regulation of non-canonical NF-kappaB signal trans | GO:1901224 | GO:BP | 0.047163864 |
| regulation of carbohydrate catabolic process | GO:0043470 | GO:BP | 0.047163864 |
| photoreceptor cell differentiation | GO:0046530 | GO:BP | 0.047247181 |
| inner ear receptor cell differentiation | GO:0060113 | GO:BP | 0.047247181 |
| regulation of alpha-beta T cell differentiation | GO:0046637 | GO:BP | 0.047247181 |
| positive regulation of protein localization | GO:1903829 | GO:BP | 0.047316527 |
| positive regulation of phosphorus metabolic process | GO:0010562 | GO:BP | 0.048092686 |
| positive regulation of phosphate metabolic process | GO:0045937 | GO:BP | 0.048092686 |
| regulation of autophagy of mitochondrion | GO:1903146 | GO:BP | 0.048840001 |
| intermembrane lipid transfer | GO:0120009 | GO:BP | 0.048927243 |
| negative regulation of transferase activity | GO:0051348 | GO:BP | 0.04896995 |
| excretion | GO:0007588 | GO:BP | 0.049273518 |
| glucose 6-phosphate metabolic process | GO:0051156 | GO:BP | 0.049273518 |
| negative regulation of protein ubiquitination | GO:0031397 | GO:BP | 0.049411495 |
| regulation of T cell mediated cytotoxicity | GO:0001914 | GO:BP | 0.049411495 |
| response to other organism | GO:0051707 | GO:BP | 0.049411495 |
| semi-lunar valve development | GO:1905314 | GO:BP | 0.049411495 |
| regulation of tissue remodeling | GO:0034103 | GO:BP | 0.049411495 |
| regulation of glycoprotein metabolic process | GO:1903018 | GO:BP | 0.049411495 |
| chemical synaptic transmission | GO:0007268 | GO:BP | 0.049569648 |
| anterograde trans-synaptic signaling | GO:0098916 | GO:BP | 0.049569648 |
| regulation of myeloid leukocyte differentiation | GO:0002761 | GO:BP | 0.049950107 |

| | | | |
|---|---|---|---|
| leukocyte migration | GO:0050900 | GO:BP | 0.04997397 |
| collagen fibril organization | GO:0030199 | GO:BP | 0.049995358 |
| taxis | GO:0042330 | GO:BP | 0.049995358 |
| cytoplasm | GO:0005737 | GO:CC | 1.08E-82 |
| extracellular organelle | GO:0043230 | GO:CC | 8.94E-51 |
| extracellular vesicle | GO:1903561 | GO:CC | 8.94E-51 |
| extracellular membrane-bounded organelle | GO:0065010 | GO:CC | 8.94E-51 |
| extracellular exosome | GO:0070062 | GO:CC | 2.22E-50 |
| vesicle | GO:0031982 | GO:CC | 9.40E-47 |
| extracellular space | GO:0005615 | GO:CC | 2.72E-37 |
| extracellular region | GO:0005576 | GO:CC | 9.26E-32 |
| endomembrane system | GO:0012505 | GO:CC | 4.08E-29 |
| membrane | GO:0016020 | GO:CC | 2.84E-23 |
| cytosolic ribosome | GO:0022626 | GO:CC | 4.26E-23 |
| cellular anatomical entity | GO:0110165 | GO:CC | 6.39E-21 |
| endoplasmic reticulum | GO:0005783 | GO:CC | 2.59E-20 |
| cytosol | GO:0005829 | GO:CC | 6.45E-19 |
| membrane-bounded organelle | GO:0043227 | GO:CC | 6.36E-18 |
| cytosolic large ribosomal subunit | GO:0022625 | GO:CC | 7.33E-18 |
| collagen-containing extracellular matrix | GO:0062023 | GO:CC | 1.93E-17 |
| organelle membrane | GO:0031090 | GO:CC | 2.03E-17 |
| ribosomal subunit | GO:0044391 | GO:CC | 5.00E-16 |
| cytoplasmic vesicle | GO:0031410 | GO:CC | 8.51E-16 |
| intracellular vesicle | GO:0097708 | GO:CC | 8.71E-16 |
| cell projection | GO:0042995 | GO:CC | 1.15E-15 |
| plasma membrane bounded cell projection | GO:0120025 | GO:CC | 2.26E-15 |
| endoplasmic reticulum lumen | GO:0005788 | GO:CC | 3.18E-14 |
| cell junction | GO:0030054 | GO:CC | 4.09E-14 |
| organelle | GO:0043226 | GO:CC | 4.40E-14 |
| external encapsulating structure | GO:0030312 | GO:CC | 5.37E-14 |
| intracellular anatomical structure | GO:0005622 | GO:CC | 5.37E-14 |
| cell-substrate junction | GO:0030055 | GO:CC | 8.74E-14 |
| extracellular matrix | GO:0031012 | GO:CC | 8.74E-14 |
| cell periphery | GO:0071944 | GO:CC | 1.68E-13 |
| secretory vesicle | GO:0099503 | GO:CC | 1.93E-13 |
| focal adhesion | GO:0005925 | GO:CC | 2.16E-13 |
| anchoring junction | GO:0070161 | GO:CC | 2.51E-13 |
| secretory granule | GO:0030141 | GO:CC | 6.75E-13 |
| cilium | GO:0005929 | GO:CC | 7.29E-13 |
| mitochondrion | GO:0005739 | GO:CC | 2.14E-12 |
| respiratory chain complex | GO:0098803 | GO:CC | 3.16E-12 |
| cytosolic small ribosomal subunit | GO:0022627 | GO:CC | 5.98E-12 |
| mitochondrial respirasome | GO:0005746 | GO:CC | 1.66E-11 |
| respirasome | GO:0070469 | GO:CC | 5.58E-11 |
| large ribosomal subunit | GO:0015934 | GO:CC | 3.96E-10 |
| motile cilium | GO:0031514 | GO:CC | 5.11E-10 |
| vacuole | GO:0005773 | GO:CC | 6.85E-10 |
| organelle subcompartment | GO:0031984 | GO:CC | 2.56E-09 |

| | | | |
|---|---|---|---|
| bounding membrane of organelle | GO:0098588 | GO:CC | 4.12E-09 |
| lysosome | GO:0005764 | GO:CC | 4.79E-09 |
| lytic vacuole | GO:0000323 | GO:CC | 4.79E-09 |
| oxidoreductase complex | GO:1990204 | GO:CC | 1.18E-08 |
| cytoplasmic vesicle lumen | GO:0060205 | GO:CC | 1.73E-08 |
| secretory granule lumen | GO:0034774 | GO:CC | 1.95E-08 |
| contractile fiber | GO:0043292 | GO:CC | 2.01E-08 |
| vesicle lumen | GO:0031983 | GO:CC | 2.01E-08 |
| inner mitochondrial membrane protein complex | GO:0098800 | GO:CC | 4.17E-08 |
| plasma membrane | GO:0005886 | GO:CC | 4.68E-08 |
| neuron projection | GO:0043005 | GO:CC | 4.89E-08 |
| nuclear outer membrane-endoplasmic reticulum membrane | GO:0042175 | GO:CC | 1.85E-07 |
| mitochondrial envelope | GO:0005740 | GO:CC | 2.26E-07 |
| myofibril | GO:0030016 | GO:CC | 2.58E-07 |
| endoplasmic reticulum subcompartment | GO:0098827 | GO:CC | 2.60E-07 |
| synapse | GO:0045202 | GO:CC | 3.44E-07 |
| 9+2 motile cilium | GO:0097729 | GO:CC | 3.53E-07 |
| endoplasmic reticulum membrane | GO:0005789 | GO:CC | 4.62E-07 |
| sarcomere | GO:0030017 | GO:CC | 4.78E-07 |
| envelope | GO:0031975 | GO:CC | 5.69E-07 |
| organelle envelope | GO:0031967 | GO:CC | 5.69E-07 |
| cytoskeleton | GO:0005856 | GO:CC | 6.75E-07 |
| mitochondrial membrane | GO:0031966 | GO:CC | 8.12E-07 |
| small ribosomal subunit | GO:0015935 | GO:CC | 9.68E-07 |
| supramolecular polymer | GO:0099081 | GO:CC | 1.89E-06 |
| vacuolar lumen | GO:0005775 | GO:CC | 2.64E-06 |
| Z disc | GO:0030018 | GO:CC | 3.25E-06 |
| ficolin-1-rich granule lumen | GO:1904813 | GO:CC | 3.27E-06 |
| I band | GO:0031674 | GO:CC | 4.62E-06 |
| supramolecular fiber | GO:0099512 | GO:CC | 5.88E-06 |
| non-motile cilium | GO:0097730 | GO:CC | 6.74E-06 |
| cytochrome complex | GO:0070069 | GO:CC | 6.80E-06 |
| plasma membrane region | GO:0098590 | GO:CC | 9.57E-06 |
| ficolin-1-rich granule | GO:0101002 | GO:CC | 9.85E-06 |
| membrane protein complex | GO:0098796 | GO:CC | 9.85E-06 |
| organelle inner membrane | GO:0019866 | GO:CC | 1.02E-05 |
| Golgi apparatus | GO:0005794 | GO:CC | 1.02E-05 |
| mitochondrial inner membrane | GO:0005743 | GO:CC | 1.07E-05 |
| NADH dehydrogenase complex | GO:0030964 | GO:CC | 1.61E-05 |
| respiratory chain complex I | GO:0045271 | GO:CC | 1.61E-05 |
| mitochondrial respiratory chain complex I | GO:0005747 | GO:CC | 1.61E-05 |
| microtubule cytoskeleton | GO:0015630 | GO:CC | 1.69E-05 |
| intracellular organelle | GO:0043229 | GO:CC | 2.18E-05 |
| supramolecular complex | GO:0099080 | GO:CC | 2.93E-05 |
| basement membrane | GO:0005604 | GO:CC | 2.97E-05 |
| intracellular membrane-bounded organelle | GO:0043231 | GO:CC | 5.11E-05 |
| vesicle membrane | GO:0012506 | GO:CC | 5.99E-05 |
| mitochondrial respiratory chain complex IV | GO:0005751 | GO:CC | 7.13E-05 |

| | | | |
|---|---|---|---|
| mitochondrial matrix | GO:0005759 | GO:CC | 7.13E-05 |
| cytoplasmic vesicle membrane | GO:0030659 | GO:CC | 7.46E-05 |
| axoneme | GO:0005930 | GO:CC | 8.54E-05 |
| mitochondrial protein-containing complex | GO:0098798 | GO:CC | 8.88E-05 |
| perinuclear region of cytoplasm | GO:0048471 | GO:CC | 9.72E-05 |
| respiratory chain complex IV | GO:0045277 | GO:CC | 9.77E-05 |
| ciliary plasm | GO:0097014 | GO:CC | 0.000101101 |
| lysosomal lumen | GO:0043202 | GO:CC | 0.000110534 |
| ciliary basal body | GO:0036064 | GO:CC | 0.000139959 |
| actin cytoskeleton | GO:0015629 | GO:CC | 0.000140409 |
| lysosomal membrane | GO:0005765 | GO:CC | 0.000145193 |
| cell body | GO:0044297 | GO:CC | 0.000145193 |
| lytic vacuole membrane | GO:0098852 | GO:CC | 0.000145193 |
| 9+0 non-motile cilium | GO:0097731 | GO:CC | 0.000160487 |
| vacuolar membrane | GO:0005774 | GO:CC | 0.000182644 |
| transmembrane transporter complex | GO:1902495 | GO:CC | 0.000203466 |
| sperm flagellum | GO:0036126 | GO:CC | 0.000210186 |
| transporter complex | GO:1990351 | GO:CC | 0.000264571 |
| pigment granule | GO:0048770 | GO:CC | 0.000297378 |
| melanosome | GO:0042470 | GO:CC | 0.000297378 |
| cell surface | GO:0009986 | GO:CC | 0.00037864 |
| stereocilium | GO:0032420 | GO:CC | 0.00038801 |
| axon | GO:0030424 | GO:CC | 0.000403269 |
| microtubule | GO:0005874 | GO:CC | 0.000435729 |
| neuronal cell body | GO:0043025 | GO:CC | 0.000446338 |
| protein complex involved in cell adhesion | GO:0098636 | GO:CC | 0.000508398 |
| photoreceptor cell cilium | GO:0097733 | GO:CC | 0.000562185 |
| integrin complex | GO:0008305 | GO:CC | 0.000596399 |
| transport vesicle | GO:0030133 | GO:CC | 0.000683546 |
| MHC protein complex | GO:0042611 | GO:CC | 0.000700038 |
| dynein complex | GO:0030286 | GO:CC | 0.000807351 |
| endosome | GO:0005768 | GO:CC | 0.00091736 |
| cell-cell junction | GO:0005911 | GO:CC | 0.000991486 |
| tertiary granule lumen | GO:1904724 | GO:CC | 0.001101704 |
| perikaryon | GO:0043204 | GO:CC | 0.001101704 |
| azurophil granule | GO:0042582 | GO:CC | 0.001101704 |
| primary lysosome | GO:0005766 | GO:CC | 0.001101704 |
| MHC class I protein complex | GO:0042612 | GO:CC | 0.001163789 |
| cytoplasmic region | GO:0099568 | GO:CC | 0.001207966 |
| Golgi cis cisterna | GO:0000137 | GO:CC | 0.001332745 |
| basal part of cell | GO:0045178 | GO:CC | 0.001353743 |
| plasma membrane bounded cell projection cytoplasm | GO:0032838 | GO:CC | 0.001535958 |
| basal plasma membrane | GO:0009925 | GO:CC | 0.001535958 |
| catalytic complex | GO:1902494 | GO:CC | 0.001660933 |
| myosin complex | GO:0016459 | GO:CC | 0.002034329 |
| stereocilium bundle | GO:0032421 | GO:CC | 0.00223563 |
| ciliary transition zone | GO:0035869 | GO:CC | 0.002349471 |
| tertiary granule | GO:0070820 | GO:CC | 0.002594944 |

| | | | |
|---|---|---|---|
| complex of collagen trimers | GO:0098644 | GO:CC | 0.002714479 |
| endoplasmic reticulum protein-containing complex | GO:0140534 | GO:CC | 0.00295844 |
| endoplasmic reticulum chaperone complex | GO:0034663 | GO:CC | 0.002998475 |
| microtubule organizing center | GO:0005815 | GO:CC | 0.003024379 |
| secretory granule membrane | GO:0030667 | GO:CC | 0.003232345 |
| ribbon synapse | GO:0097470 | GO:CC | 0.003232345 |
| basolateral plasma membrane | GO:0016323 | GO:CC | 0.003594985 |
| Golgi apparatus subcompartment | GO:0098791 | GO:CC | 0.003938372 |
| apical part of cell | GO:0045177 | GO:CC | 0.004236194 |
| endosome membrane | GO:0010008 | GO:CC | 0.00433217 |
| microtubule associated complex | GO:0005875 | GO:CC | 0.004469651 |
| lumenal side of endoplasmic reticulum membrane | GO:0098553 | GO:CC | 0.004835588 |
| cluster of actin-based cell projections | GO:0098862 | GO:CC | 0.005393885 |
| COPII-coated ER to Golgi transport vesicle | GO:0030134 | GO:CC | 0.005613199 |
| photoreceptor connecting cilium | GO:0032391 | GO:CC | 0.006094885 |
| autophagosome | GO:0005776 | GO:CC | 0.006966049 |
| somatodendritic compartment | GO:0036477 | GO:CC | 0.006966049 |
| side of membrane | GO:0098552 | GO:CC | 0.006971761 |
| endolysosome | GO:0036019 | GO:CC | 0.007446524 |
| myosin II complex | GO:0016460 | GO:CC | 0.007462032 |
| azurophil granule lumen | GO:0035578 | GO:CC | 0.008321953 |
| postsynapse | GO:0098794 | GO:CC | 0.008473015 |
| outer membrane | GO:0019867 | GO:CC | 0.008816887 |
| postsynaptic density | GO:0014069 | GO:CC | 0.008956644 |
| organelle outer membrane | GO:0031968 | GO:CC | 0.011136295 |
| axonemal microtubule | GO:0005879 | GO:CC | 0.011212907 |
| postsynaptic specialization | GO:0099572 | GO:CC | 0.011861982 |
| asymmetric synapse | GO:0032279 | GO:CC | 0.012486902 |
| sarcoplasmic reticulum | GO:0016529 | GO:CC | 0.012947709 |
| coated vesicle membrane | GO:0030662 | GO:CC | 0.0135617 |
| cell projection membrane | GO:0031253 | GO:CC | 0.0135617 |
| platelet alpha granule | GO:0031091 | GO:CC | 0.013633801 |
| eukaryotic translation elongation factor 1 complex | GO:0005853 | GO:CC | 0.015235321 |
| glycogen granule | GO:0042587 | GO:CC | 0.015525432 |
| ionotropic glutamate receptor complex | GO:0008328 | GO:CC | 0.01629409 |
| axonemal dynein complex | GO:0005858 | GO:CC | 0.017103442 |
| perisynaptic extracellular matrix | GO:0098966 | GO:CC | 0.019219666 |
| microbody | GO:0042579 | GO:CC | 0.019219666 |
| peroxisome | GO:0005777 | GO:CC | 0.019219666 |
| apical plasma membrane | GO:0016324 | GO:CC | 0.019219666 |
| site of polarized growth | GO:0030427 | GO:CC | 0.019407869 |
| photoreceptor outer segment | GO:0001750 | GO:CC | 0.019938084 |
| actin-based cell projection | GO:0098858 | GO:CC | 0.019938084 |
| presynapse | GO:0098793 | GO:CC | 0.020692736 |
| neuron to neuron synapse | GO:0098984 | GO:CC | 0.021220874 |
| cytoplasmic microtubule | GO:0005881 | GO:CC | 0.021611132 |
| sarcoplasm | GO:0016528 | GO:CC | 0.022490015 |
| respiratory chain complex III | GO:0045275 | GO:CC | 0.022595961 |

| | | | |
|---|---|---|---|
| dendrite terminus | GO:0044292 | GO:CC | 0.022595961 |
| mitochondrial respiratory chain complex III | GO:0005750 | GO:CC | 0.022595961 |
| distal axon | GO:0150034 | GO:CC | 0.024035022 |
| stereocilium tip | GO:0032426 | GO:CC | 0.025020027 |
| voltage-gated calcium channel complex | GO:0005891 | GO:CC | 0.026478013 |
| growth cone | GO:0030426 | GO:CC | 0.027043869 |
| photoreceptor ribbon synapse | GO:0098684 | GO:CC | 0.027043869 |
| trans-Golgi network | GO:0005802 | GO:CC | 0.027933244 |
| protein-DNA complex | GO:0032993 | GO:CC | 0.028009196 |
| cell leading edge | GO:0031252 | GO:CC | 0.030724877 |
| external side of plasma membrane | GO:0009897 | GO:CC | 0.03074976 |
| presynaptic active zone | GO:0048786 | GO:CC | 0.031286768 |
| coated vesicle | GO:0030135 | GO:CC | 0.03197976 |
| calcium channel complex | GO:0034704 | GO:CC | 0.032347724 |
| synapse-associated extracellular matrix | GO:0099535 | GO:CC | 0.033651736 |
| myosin filament | GO:0032982 | GO:CC | 0.034587733 |
| nucleoplasm | GO:0005654 | GO:CC | 0.034587733 |
| muscle myosin complex | GO:0005859 | GO:CC | 0.036639891 |
| CENP-A containing chromatin | GO:0061638 | GO:CC | 0.037005315 |
| CENP-A containing nucleosome | GO:0043505 | GO:CC | 0.037005315 |
| ciliary tip | GO:0097542 | GO:CC | 0.037284088 |
| photoreceptor inner segment | GO:0001917 | GO:CC | 0.037921742 |
| centriolar satellite | GO:0034451 | GO:CC | 0.037921742 |
| rough endoplasmic reticulum | GO:0005791 | GO:CC | 0.038390548 |
| membrane-enclosed lumen | GO:0031974 | GO:CC | 0.041037743 |
| intracellular organelle lumen | GO:0070013 | GO:CC | 0.041037743 |
| organelle lumen | GO:0043233 | GO:CC | 0.041037743 |
| sarcolemma | GO:0042383 | GO:CC | 0.042614954 |
| intraciliary transport particle | GO:0030990 | GO:CC | 0.047576637 |
| centrosome | GO:0005813 | GO:CC | 0.048433875 |
| transcription export complex 2 | GO:0070390 | GO:CC | 0.048541869 |
| endolysosome lumen | GO:0036021 | GO:CC | 0.048541869 |
| sarcoplasmic reticulum lumen | GO:0033018 | GO:CC | 0.048541869 |
| extrinsic component of membrane | GO:0019898 | GO:CC | 0.048541869 |
| perineuronal net | GO:0072534 | GO:CC | 0.048541869 |
| transcription factor AP-1 complex | GO:0035976 | GO:CC | 0.048541869 |
| GATOR2 complex | GO:0061700 | GO:CC | 0.048541869 |
| insulin-like growth factor binding protein complex | GO:0016942 | GO:CC | 0.048541869 |
| 3M complex | GO:1990393 | GO:CC | 0.048541869 |
| late endosome | GO:0005770 | GO:CC | 0.049063538 |
| ciliary base | GO:0097546 | GO:CC | 0.049970211 |
| neurotransmitter receptor complex | GO:0098878 | GO:CC | 0.049970211 |
| sperm principal piece | GO:0097228 | GO:CC | 0.049970211 |
| protein binding | GO:0005515 | GO:MF | 6.25E-66 |
| small molecule binding | GO:0036094 | GO:MF | 7.70E-29 |
| ion binding | GO:0043167 | GO:MF | 4.75E-27 |
| cation binding | GO:0043169 | GO:MF | 9.17E-21 |
| metal ion binding | GO:0046872 | GO:MF | 5.42E-20 |

| | | | |
|---|---|---|---|
| oxidoreductase activity | GO:0016491 | GO:MF | 1.56E-18 |
| RNA polymerase II transcription regulatory region sequence- | GO:0000977 | GO:MF | 1.70E-17 |
| DNA-binding transcription factor activity | GO:0003700 | GO:MF | 1.70E-17 |
| binding | GO:0005488 | GO:MF | 2.19E-17 |
| transcription cis-regulatory region binding | GO:0000976 | GO:MF | 2.36E-17 |
| transcription regulatory region nucleic acid binding | GO:0001067 | GO:MF | 2.87E-17 |
| DNA-binding transcription factor activity, RNA polymerase II- | GO:0000981 | GO:MF | 6.84E-16 |
| sequence-specific double-stranded DNA binding | GO:1990837 | GO:MF | 8.54E-16 |
| sequence-specific DNA binding | GO:0043565 | GO:MF | 1.16E-15 |
| double-stranded DNA binding | GO:0003690 | GO:MF | 1.42E-15 |
| RNA polymerase II cis-regulatory region sequence-specific D | GO:0000978 | GO:MF | 4.34E-14 |
| cis-regulatory region sequence-specific DNA binding | GO:0000987 | GO:MF | 4.51E-14 |
| transcription regulator activity | GO:0140110 | GO:MF | 4.93E-12 |
| DNA binding | GO:0003677 | GO:MF | 6.62E-09 |
| electron transfer activity | GO:0009055 | GO:MF | 2.38E-08 |
| catalytic activity | GO:0003824 | GO:MF | 4.29E-08 |
| oxidoreductase activity, acting on the CH-OH group of donor | GO:0016616 | GO:MF | 1.23E-07 |
| identical protein binding | GO:0042802 | GO:MF | 2.58E-07 |
| oxidoreductase activity, acting on CH-OH group of donors | GO:0016614 | GO:MF | 4.82E-07 |
| oxidoreduction-driven active transmembrane transporter ac | GO:0015453 | GO:MF | 1.79E-06 |
| DNA-binding transcription repressor activity | GO:0001217 | GO:MF | 1.38E-05 |
| DNA-binding transcription repressor activity, RNA polymeras | GO:0001227 | GO:MF | 1.79E-05 |
| extracellular matrix structural constituent | GO:0005201 | GO:MF | 2.29E-05 |
| growth factor binding | GO:0019838 | GO:MF | 4.46E-05 |
| integrin binding | GO:0005178 | GO:MF | 6.28E-05 |
| oxidoreductase activity, acting on NAD(P)H, quinone or simi | GO:0016655 | GO:MF | 6.44E-05 |
| cell adhesion molecule binding | GO:0050839 | GO:MF | 7.38E-05 |
| oxidoreductase activity, acting on NAD(P)H | GO:0016651 | GO:MF | 0.000136341 |
| calcium ion binding | GO:0005509 | GO:MF | 0.000177597 |
| anion binding | GO:0043168 | GO:MF | 0.000251384 |
| calmodulin binding | GO:0005516 | GO:MF | 0.000291137 |
| minus-end-directed microtubule motor activity | GO:0008569 | GO:MF | 0.000291137 |
| heterocyclic compound binding | GO:1901363 | GO:MF | 0.000325251 |
| unfolded protein binding | GO:0051082 | GO:MF | 0.000325251 |
| NAD(P)H dehydrogenase (quinone) activity | GO:0003955 | GO:MF | 0.000483219 |
| enzyme binding | GO:0019899 | GO:MF | 0.000742393 |
| nucleotide binding | GO:0000166 | GO:MF | 0.000757262 |
| nucleoside phosphate binding | GO:1901265 | GO:MF | 0.000782624 |
| NADH dehydrogenase (ubiquinone) activity | GO:0008137 | GO:MF | 0.000938703 |
| carbohydrate derivative binding | GO:0097367 | GO:MF | 0.001128425 |
| NADH dehydrogenase (quinone) activity | GO:0050136 | GO:MF | 0.001486789 |
| extracellular matrix binding | GO:0050840 | GO:MF | 0.001932992 |
| adenyl nucleotide binding | GO:0030554 | GO:MF | 0.003008963 |
| aldo-keto reductase (NADP) activity | GO:0004033 | GO:MF | 0.003231127 |
| NADH dehydrogenase activity | GO:0003954 | GO:MF | 0.003484914 |
| purine nucleotide binding | GO:0017076 | GO:MF | 0.004077795 |
| collagen binding | GO:0005518 | GO:MF | 0.004650648 |
| primary active transmembrane transporter activity | GO:0015399 | GO:MF | 0.004650648 |

| | | | |
|---|---|---|---|
| lipid binding | GO:0008289 | GO:MF | 0.004650648 |
| oxidoreductase activity, acting on the CH-CH group of donor | GO:0016627 | GO:MF | 0.004738549 |
| protein folding chaperone | GO:0044183 | GO:MF | 0.004920539 |
| protein-containing complex binding | GO:0044877 | GO:MF | 0.005103881 |
| transporter activity | GO:0005215 | GO:MF | 0.006088444 |
| oxidoreductase activity, acting on single donors with incorpc | GO:0016702 | GO:MF | 0.008202869 |
| sulfur compound binding | GO:1901681 | GO:MF | 0.008410204 |
| sugar transmembrane transporter activity | GO:0051119 | GO:MF | 0.011309842 |
| kinase binding | GO:0019900 | GO:MF | 0.011450249 |
| structural constituent of muscle | GO:0008307 | GO:MF | 0.01371739 |
| oxidoreductase activity, acting on single donors with incorpc | GO:0016701 | GO:MF | 0.01371739 |
| cysteine-type endopeptidase activator activity involved in ap | GO:0008656 | GO:MF | 0.014534264 |
| laminin binding | GO:0043236 | GO:MF | 0.014534264 |
| monosaccharide transmembrane transporter activity | GO:0015145 | GO:MF | 0.014534264 |
| cytochrome-c oxidase activity | GO:0004129 | GO:MF | 0.014534264 |
| oxidoreductase activity, acting on a heme group of donors | GO:0016675 | GO:MF | 0.014534264 |
| dynein light intermediate chain binding | GO:0051959 | GO:MF | 0.014534264 |
| structural molecule activity | GO:0005198 | GO:MF | 0.014590227 |
| inorganic molecular entity transmembrane transporter activ | GO:0015318 | GO:MF | 0.016215651 |
| transmembrane transporter activity | GO:0022857 | GO:MF | 0.016336645 |
| peptidase activator activity | GO:0016504 | GO:MF | 0.016336645 |
| NADP binding | GO:0050661 | GO:MF | 0.016336645 |
| peptidase activator activity involved in apoptotic process | GO:0016505 | GO:MF | 0.016589141 |
| alcohol dehydrogenase (NADP+) activity | GO:0008106 | GO:MF | 0.016589141 |
| NADP-retinol dehydrogenase activity | GO:0052650 | GO:MF | 0.016589141 |
| cytoskeletal motor activity | GO:0003774 | GO:MF | 0.016589141 |
| oxidoreductase activity, acting on peroxide as acceptor | GO:0016684 | GO:MF | 0.020391492 |
| FATZ binding | GO:0051373 | GO:MF | 0.021974178 |
| carbohydrate transmembrane transporter activity | GO:0015144 | GO:MF | 0.022937473 |
| cysteine-type endopeptidase regulator activity involved in ap | GO:0043028 | GO:MF | 0.022937473 |
| hydrolase activity | GO:0016787 | GO:MF | 0.023404621 |
| signaling receptor binding | GO:0005102 | GO:MF | 0.024310059 |
| active transmembrane transporter activity | GO:0022804 | GO:MF | 0.025085377 |
| C-acetyltransferase activity | GO:0016453 | GO:MF | 0.025941322 |
| inorganic cation transmembrane transporter activity | GO:0022890 | GO:MF | 0.027371311 |
| steroid dehydrogenase activity, acting on the CH-OH group c | GO:0033764 | GO:MF | 0.029800439 |
| extracellular matrix structural constituent conferring tensile | GO:0030020 | GO:MF | 0.029800439 |
| insulin-like growth factor binding | GO:0005520 | GO:MF | 0.029800439 |
| glycosaminoglycan binding | GO:0005539 | GO:MF | 0.029800439 |
| hydrolase activity, acting on carbon-nitrogen (but not peptid | GO:0016811 | GO:MF | 0.029800439 |
| proteoglycan binding | GO:0043394 | GO:MF | 0.029800439 |
| cytoskeletal protein binding | GO:0008092 | GO:MF | 0.031559509 |
| antioxidant activity | GO:0016209 | GO:MF | 0.037625908 |
| hexose transmembrane transporter activity | GO:0015149 | GO:MF | 0.037625908 |
| glucose transmembrane transporter activity | GO:0005355 | GO:MF | 0.037625908 |
| alpha-actinin binding | GO:0051393 | GO:MF | 0.039637019 |
| adenyl ribonucleotide binding | GO:0032559 | GO:MF | 0.039637019 |
| actinin binding | GO:0042805 | GO:MF | 0.040055921 |

| | | | |
|---|---|---|---|
| phosphatase binding | GO:0019902 | GO:MF | 0.040055921 |
| dynein intermediate chain binding | GO:0045505 | GO:MF | 0.040055921 |
| DNA-binding transcription activator activity, RNA polymeras | GO:0001228 | GO:MF | 0.043607646 |
| misfolded protein binding | GO:0051787 | GO:MF | 0.046432706 |
| ribonucleotide binding | GO:0032553 | GO:MF | 0.046709804 |
| purine ribonucleotide binding | GO:0032555 | GO:MF | 0.046709804 |
| DNA-binding transcription activator activity | GO:0001216 | GO:MF | 0.049323224 |
| proton transmembrane transporter activity | GO:0015078 | GO:MF | 0.049323224 |
| phospholipid binding | GO:0005543 | GO:MF | 0.049323224 |
| Herpes simplex virus 1 infection | KEGG:05168 | KEGG | 7.40E-16 |
| Ribosome | KEGG:03010 | KEGG | 9.16E-15 |
| Coronavirus disease - COVID-19 | KEGG:05171 | KEGG | 4.55E-11 |
| Metabolic pathways | KEGG:01100 | KEGG | 1.64E-07 |
| Parkinson disease | KEGG:05012 | KEGG | 3.00E-06 |
| Diabetic cardiomyopathy | KEGG:05415 | KEGG | 3.00E-06 |
| Oxidative phosphorylation | KEGG:00190 | KEGG | 1.04E-05 |
| Prion disease | KEGG:05020 | KEGG | 1.61E-05 |
| Huntington disease | KEGG:05016 | KEGG | 2.25E-05 |
| Pathways of neurodegeneration - multiple diseases | KEGG:05022 | KEGG | 2.25E-05 |
| Carbon metabolism | KEGG:01200 | KEGG | 5.81E-05 |
| Amyotrophic lateral sclerosis | KEGG:05014 | KEGG | 6.01E-05 |
| Chemical carcinogenesis - reactive oxygen species | KEGG:05208 | KEGG | 7.17E-05 |
| Non-alcoholic fatty liver disease | KEGG:04932 | KEGG | 0.000623863 |
| ECM-receptor interaction | KEGG:04512 | KEGG | 0.000623863 |
| Steroid biosynthesis | KEGG:00100 | KEGG | 0.000936954 |
| Thermogenesis | KEGG:04714 | KEGG | 0.000948545 |
| Glycine, serine and threonine metabolism | KEGG:00260 | KEGG | 0.003340009 |
| Glycolysis / Gluconeogenesis | KEGG:00010 | KEGG | 0.004188454 |
| Cardiac muscle contraction | KEGG:04260 | KEGG | 0.008396422 |
| Alzheimer disease | KEGG:05010 | KEGG | 0.009063232 |
| Fatty acid metabolism | KEGG:01212 | KEGG | 0.014650307 |
| Fatty acid degradation | KEGG:00071 | KEGG | 0.014650307 |
| Pyruvate metabolism | KEGG:00620 | KEGG | 0.021796586 |
| Pentose phosphate pathway | KEGG:00030 | KEGG | 0.025932691 |
| Antigen processing and presentation | KEGG:04612 | KEGG | 0.031664255 |
| Th1 and Th2 cell differentiation | KEGG:04658 | KEGG | 0.044396889 |
| Citrate cycle (TCA cycle) | KEGG:00020 | KEGG | 0.046303816 |
| Biosynthesis of amino acids | KEGG:01230 | KEGG | 0.046303816 |
| Eukaryotic Translation Elongation | REAC:R-HSA-156842 | REAC | 1.45E-31 |
| Peptide chain elongation | REAC:R-HSA-156902 | REAC | 1.07E-29 |
| Formation of a pool of free 40S subunits | REAC:R-HSA-72689 | REAC | 3.20E-28 |
| Selenocysteine synthesis | REAC:R-HSA-2408557 | REAC | 1.18E-27 |
| Eukaryotic Translation Termination | REAC:R-HSA-72764 | REAC | 1.18E-27 |
| Viral mRNA Translation | REAC:R-HSA-192823 | REAC | 1.18E-27 |
| L13a-mediated translational silencing of Ceruloplasmin exp | REAC:R-HSA-156827 | REAC | 1.73E-26 |
| Response of EIF2AK4 (GCN2) to amino acid deficiency | REAC:R-HSA-9633012 | REAC | 1.37E-25 |
| Nonsense Mediated Decay (NMD) independent of the Exon J | REAC:R-HSA-975956 | REAC | 1.37E-25 |
| Selenoamino acid metabolism | REAC:R-HSA-2408522 | REAC | 2.62E-25 |

| | | | |
|---|---|---|---|
| SRP-dependent cotranslational protein targeting to membra | REAC:R-HSA-1799339 | REAC | 2.62E-25 |
| GTP hydrolysis and joining of the 60S ribosomal subunit | REAC:R-HSA-72706 | REAC | 2.62E-25 |
| Eukaryotic Translation Initiation | REAC:R-HSA-72613 | REAC | 4.71E-25 |
| Cap-dependent Translation Initiation | REAC:R-HSA-72737 | REAC | 4.71E-25 |
| Cellular response to starvation | REAC:R-HSA-9711097 | REAC | 2.28E-19 |
| Nonsense-Mediated Decay (NMD) | REAC:R-HSA-927802 | REAC | 1.40E-17 |
| Nonsense Mediated Decay (NMD) enhanced by the Exon Jun | REAC:R-HSA-975957 | REAC | 1.40E-17 |
| Metabolism of amino acids and derivatives | REAC:R-HSA-71291 | REAC | 8.41E-17 |
| Influenza Viral RNA Transcription and Replication | REAC:R-HSA-168273 | REAC | 2.17E-15 |
| Regulation of expression of SLITs and ROBOs | REAC:R-HSA-9010553 | REAC | 2.49E-15 |
| Metabolism | REAC:R-HSA-1430728 | REAC | 3.78E-15 |
| Influenza Infection | REAC:R-HSA-168255 | REAC | 2.87E-13 |
| SARS-CoV-1 modulates host translation machinery | REAC:R-HSA-9735869 | REAC | 4.09E-13 |
| Translation | REAC:R-HSA-72766 | REAC | 8.58E-12 |
| Signaling by ROBO receptors | REAC:R-HSA-376176 | REAC | 2.45E-11 |
| Nervous system development | REAC:R-HSA-9675108 | REAC | 4.86E-11 |
| Activation of the mRNA upon binding of the cap-binding com | REAC:R-HSA-72662 | REAC | 1.16E-10 |
| Axon guidance | REAC:R-HSA-422475 | REAC | 2.02E-10 |
| Formation of the ternary complex, and subsequently, the 43 | REAC:R-HSA-72695 | REAC | 2.02E-10 |
| Translation initiation complex formation | REAC:R-HSA-72649 | REAC | 2.92E-10 |
| Ribosomal scanning and start codon recognition | REAC:R-HSA-72702 | REAC | 1.75E-09 |
| Cellular responses to stress | REAC:R-HSA-2262752 | REAC | 2.21E-09 |
| Cellular responses to stimuli | REAC:R-HSA-8953897 | REAC | 3.84E-09 |
| Respiratory electron transport, ATP synthesis by chemiosmo | REAC:R-HSA-163200 | REAC | 2.49E-07 |
| SARS-CoV-2 modulates host translation machinery | REAC:R-HSA-9754678 | REAC | 6.61E-07 |
| Major pathway of rRNA processing in the nucleolus and cyto | REAC:R-HSA-6791226 | REAC | 6.86E-07 |
| Respiratory electron transport | REAC:R-HSA-611105 | REAC | 9.68E-07 |
| rRNA processing in the nucleus and cytosol | REAC:R-HSA-8868773 | REAC | 2.00E-06 |
| The citric acid (TCA) cycle and respiratory electron transport | REAC:R-HSA-1428517 | REAC | 2.00E-06 |
| Cholesterol biosynthesis | REAC:R-HSA-191273 | REAC | 2.00E-06 |
| Generic Transcription Pathway | REAC:R-HSA-212436 | REAC | 5.60E-06 |
| Extracellular matrix organization | REAC:R-HSA-1474244 | REAC | 6.17E-06 |
| SARS-CoV-1-host interactions | REAC:R-HSA-9692914 | REAC | 8.86E-06 |
| rRNA processing | REAC:R-HSA-72312 | REAC | 1.14E-05 |
| Integrin cell surface interactions | REAC:R-HSA-216083 | REAC | 2.62E-05 |
| Neutrophil degranulation | REAC:R-HSA-6798695 | REAC | 0.00021625 |
| SARS-CoV-1 Infection | REAC:R-HSA-9678108 | REAC | 0.000223585 |
| Collagen formation | REAC:R-HSA-1474290 | REAC | 0.000465535 |
| ECM proteoglycans | REAC:R-HSA-3000178 | REAC | 0.00050135 |
| RNA Polymerase II Transcription | REAC:R-HSA-73857 | REAC | 0.000555689 |
| Gluconeogenesis | REAC:R-HSA-70263 | REAC | 0.000573688 |
| Assembly of collagen fibrils and other multimeric structures | REAC:R-HSA-2022090 | REAC | 0.000925031 |
| NGF-stimulated transcription | REAC:R-HSA-9031628 | REAC | 0.001013879 |
| Signaling by NOTCH1 HD Domain Mutants in Cancer | REAC:R-HSA-2691230 | REAC | 0.004181305 |
| Constitutive Signaling by NOTCH1 HD Domain Mutants | REAC:R-HSA-2691232 | REAC | 0.004181305 |
| Collagen biosynthesis and modifying enzymes | REAC:R-HSA-1650814 | REAC | 0.004194717 |
| NCAM1 interactions | REAC:R-HSA-419037 | REAC | 0.00427255 |
| Trafficking and processing of endosomal TLR | REAC:R-HSA-1679131 | REAC | 0.004359575 |

| | | | |
|---|---|---|---|
| Chaperone Mediated Autophagy | REAC:R-HSA-9613829 | REAC | 0.004717258 |
| Metabolism of proteins | REAC:R-HSA-392499 | REAC | 0.004717258 |
| Complex I biogenesis | REAC:R-HSA-6799198 | REAC | 0.00647502 |
| Laminin interactions | REAC:R-HSA-3000157 | REAC | 0.006752819 |
| Sulfide oxidation to sulfate | REAC:R-HSA-1614517 | REAC | 0.007948274 |
| Collagen degradation | REAC:R-HSA-1442490 | REAC | 0.00946207 |
| Cellular response to chemical stress | REAC:R-HSA-9711123 | REAC | 0.011473133 |
| Metabolism of steroids | REAC:R-HSA-8957322 | REAC | 0.02058163 |
| NOTCH2 Activation and Transmission of Signal to the Nucle | REAC:R-HSA-2979096 | REAC | 0.02076585 |
| Degradation of cysteine and homocysteine | REAC:R-HSA-1614558 | REAC | 0.023306159 |
| Response to elevated platelet cytosolic Ca2+ | REAC:R-HSA-76005 | REAC | 0.023306159 |
| Metabolism of carbohydrates | REAC:R-HSA-71387 | REAC | 0.024530837 |
| Regulation of Insulin-like Growth Factor (IGF) transport and | REAC:R-HSA-381426 | REAC | 0.024969539 |
| Diseases of carbohydrate metabolism | REAC:R-HSA-5663084 | REAC | 0.025839215 |
| Infectious disease | REAC:R-HSA-5663205 | REAC | 0.02695867 |
| NCAM signaling for neurite out-growth | REAC:R-HSA-375165 | REAC | 0.027873632 |
| Platelet degranulation | REAC:R-HSA-114608 | REAC | 0.028894262 |
| Endosomal/Vacuolar pathway | REAC:R-HSA-1236977 | REAC | 0.031812808 |
| The canonical retinoid cycle in rods (twilight vision) | REAC:R-HSA-2453902 | REAC | 0.032141985 |
| Synthesis of 12-eicosatetraenoic acid derivatives | REAC:R-HSA-2142712 | REAC | 0.035317727 |
| Peroxisomal protein import | REAC:R-HSA-9033241 | REAC | 0.035317727 |
| Post-translational protein phosphorylation | REAC:R-HSA-8957275 | REAC | 0.036166107 |
| Activation of gene expression by SREBF (SREBP) | REAC:R-HSA-2426168 | REAC | 0.041102728 |
| Nuclear Events (kinase and transcription factor activation) | REAC:R-HSA-198725 | REAC | 0.04546762 |
| Meiotic recombination | REAC:R-HSA-912446 | REAC | 0.04692868 |

| term_size | intersection_size |
|---|---|
| 5986 | 1570 |
| 4836 | 1306 |
| 12680 | 2962 |
| 4264 | 1167 |
| 5591 | 1459 |
| 5440 | 1417 |
| 12287 | 2857 |
| 4105 | 1101 |
| 4116 | 1103 |
| 6453 | 1621 |
| 4027 | 1079 |
| 7669 | 1883 |
| 3939 | 1057 |
| 156 | 91 |
| 5899 | 1494 |
| 11738 | 2724 |
| 6250 | 1565 |
| 1813 | 546 |
| 3855 | 1028 |
| 4366 | 1141 |
| 4367 | 1141 |
| 5717 | 1435 |
| 4912 | 1256 |
| 3446 | 920 |
| 3464 | 924 |
| 515 | 197 |
| 2567 | 714 |
| 3601 | 952 |
| 4063 | 1057 |
| 4643 | 1187 |
| 3569 | 944 |
| 3443 | 912 |
| 3437 | 910 |
| 2691 | 735 |
| 3763 | 980 |
| 7394 | 1777 |
| 8976 | 2112 |
| 1128 | 354 |
| 2637 | 719 |
| 2596 | 707 |
| 2010 | 568 |
| 2684 | 726 |
| 3010 | 801 |
| 2571 | 698 |
| 2955 | 787 |
| 2819 | 751 |

| | |
|---|---|
| 3973 | 1012 |
| 2683 | 718 |
| 7158 | 1706 |
| 4350 | 1092 |
| 899 | 283 |
| 921 | 287 |
| 592 | 201 |
| 927 | 288 |
| 2723 | 717 |
| 1388 | 402 |
| 2274 | 612 |
| 1988 | 545 |
| 3711 | 940 |
| 1984 | 543 |
| 3039 | 787 |
| 629 | 209 |
| 1463 | 418 |
| 1506 | 428 |
| 3613 | 915 |
| 5520 | 1336 |
| 1637 | 459 |
| 1770 | 489 |
| 2846 | 738 |
| 6537 | 1552 |
| 6673 | 1581 |
| 3346 | 851 |
| 179 | 80 |
| 1663 | 462 |
| 568 | 190 |
| 2668 | 695 |
| 1913 | 520 |
| 2556 | 669 |
| 1766 | 485 |
| 6447 | 1528 |
| 196 | 84 |
| 1761 | 482 |
| 1194 | 346 |
| 6631 | 1564 |
| 2018 | 541 |
| 1092 | 320 |
| 241 | 97 |
| 829 | 254 |
| 1034 | 305 |
| 727 | 228 |
| 145 | 67 |
| 1594 | 440 |
| 335 | 123 |
| 5959 | 1415 |

| | |
|---|---|
| 175 | 76 |
| 3420 | 857 |
| 2202 | 580 |
| 1091 | 316 |
| 2102 | 555 |
| 5813 | 1379 |
| 1683 | 457 |
| 911 | 271 |
| 607 | 194 |
| 5781 | 1371 |
| 244 | 95 |
| 1600 | 435 |
| 100 | 50 |
| 100 | 50 |
| 1005 | 292 |
| 5654 | 1340 |
| 4881 | 1172 |
| 1971 | 521 |
| 92 | 47 |
| 1399 | 386 |
| 2603 | 664 |
| 1742 | 466 |
| 538 | 173 |
| 325 | 116 |
| 2074 | 542 |
| 512 | 166 |
| 2706 | 686 |
| 223 | 87 |
| 532 | 171 |
| 2531 | 646 |
| 567 | 180 |
| 326 | 116 |
| 2455 | 628 |
| 2685 | 679 |
| 325 | 115 |
| 677 | 207 |
| 254 | 95 |
| 247 | 93 |
| 1570 | 422 |
| 1613 | 432 |
| 371 | 127 |
| 2006 | 523 |
| 1777 | 470 |
| 5874 | 1378 |
| 1183 | 330 |
| 217 | 84 |
| 324 | 114 |
| 773 | 230 |

| | |
|---|---|
| 3957 | 960 |
| 2799 | 702 |
| 779 | 231 |
| 122 | 55 |
| 1778 | 469 |
| 601 | 186 |
| 1340 | 366 |
| 66 | 36 |
| 1712 | 453 |
| 5526 | 1300 |
| 1512 | 406 |
| 540 | 170 |
| 166 | 68 |
| 166 | 68 |
| 609 | 187 |
| 1077 | 302 |
| 3001 | 744 |
| 382 | 128 |
| 678 | 204 |
| 254 | 93 |
| 379 | 127 |
| 1709 | 450 |
| 1110 | 309 |
| 182 | 72 |
| 172 | 69 |
| 152 | 63 |
| 1576 | 418 |
| 1512 | 403 |
| 397 | 131 |
| 932 | 265 |
| 887 | 254 |
| 1799 | 469 |
| 1648 | 434 |
| 1861 | 483 |
| 137 | 58 |
| 766 | 224 |
| 1514 | 402 |
| 61 | 33 |
| 986 | 277 |
| 270 | 96 |
| 168 | 67 |
| 168 | 67 |
| 148 | 61 |
| 341 | 115 |
| 1398 | 374 |
| 1411 | 377 |
| 113 | 50 |
| 907 | 257 |

| | |
|---|---|
| 395 | 129 |
| 752 | 219 |
| 163 | 65 |
| 946 | 266 |
| 291 | 101 |
| 1398 | 373 |
| 2425 | 608 |
| 114 | 50 |
| 881 | 250 |
| 557 | 170 |
| 209 | 78 |
| 1431 | 380 |
| 723 | 211 |
| 1561 | 410 |
| 335 | 112 |
| 335 | 112 |
| 1179 | 320 |
| 1225 | 330 |
| 375 | 122 |
| 555 | 168 |
| 1054 | 289 |
| 144 | 58 |
| 963 | 267 |
| 490 | 151 |
| 250 | 88 |
| 1678 | 434 |
| 5541 | 1285 |
| 341 | 112 |
| 119 | 50 |
| 1130 | 305 |
| 1033 | 282 |
| 1579 | 409 |
| 616 | 181 |
| 147 | 58 |
| 441 | 137 |
| 806 | 227 |
| 565 | 168 |
| 315 | 104 |
| 187 | 69 |
| 1227 | 326 |
| 135 | 54 |
| 245 | 85 |
| 1573 | 406 |
| 102 | 44 |
| 600 | 176 |
| 177 | 66 |
| 77 | 36 |
| 827 | 231 |

| | |
|---|---|
| 1333 | 350 |
| 920 | 253 |
| 450 | 138 |
| 1347 | 353 |
| 303 | 100 |
| 1189 | 316 |
| 54 | 28 |
| 54 | 28 |
| 274 | 92 |
| 127 | 51 |
| 609 | 177 |
| 1132 | 302 |
| 141 | 55 |
| 1133 | 302 |
| 783 | 219 |
| 485 | 146 |
| 839 | 232 |
| 3031 | 730 |
| 894 | 245 |
| 345 | 110 |
| 217 | 76 |
| 266 | 89 |
| 1328 | 346 |
| 44 | 24 |
| 229 | 79 |
| 525 | 155 |
| 316 | 102 |
| 320 | 103 |
| 252 | 85 |
| 252 | 85 |
| 414 | 127 |
| 1038 | 278 |
| 583 | 169 |
| 1167 | 308 |
| 190 | 68 |
| 547 | 160 |
| 900 | 245 |
| 471 | 141 |
| 1704 | 431 |
| 1229 | 322 |
| 568 | 165 |
| 257 | 86 |
| 1290 | 336 |
| 496 | 147 |
| 672 | 190 |
| 159 | 59 |
| 393 | 121 |
| 1253 | 327 |

| | |
|---|---|
| 785 | 217 |
| 1035 | 276 |
| 1001 | 268 |
| 244 | 82 |
| 842 | 230 |
| 1033 | 275 |
| 427 | 129 |
| 830 | 227 |
| 540 | 157 |
| 996 | 266 |
| 3621 | 854 |
| 70 | 32 |
| 425 | 128 |
| 808 | 221 |
| 1432 | 366 |
| 80 | 35 |
| 1463 | 373 |
| 962 | 257 |
| 363 | 112 |
| 701 | 195 |
| 1014 | 269 |
| 1588 | 401 |
| 561 | 161 |
| 964 | 257 |
| 436 | 130 |
| 939 | 251 |
| 345 | 107 |
| 1534 | 388 |
| 1060 | 279 |
| 153 | 56 |
| 429 | 128 |
| 1091 | 286 |
| 609 | 172 |
| 378 | 115 |
| 844 | 228 |
| 768 | 210 |
| 143 | 53 |
| 143 | 53 |
| 216 | 73 |
| 161 | 58 |
| 849 | 229 |
| 2052 | 504 |
| 934 | 249 |
| 351 | 108 |
| 119 | 46 |
| 225 | 75 |
| 946 | 251 |
| 3262 | 770 |

| | |
|---|---|
| 233 | 77 |
| 73 | 32 |
| 1791 | 444 |
| 1346 | 343 |
| 215 | 72 |
| 1013 | 266 |
| 327 | 101 |
| 238 | 78 |
| 864 | 231 |
| 29 | 17 |
| 304 | 95 |
| 128 | 48 |
| 197 | 67 |
| 43 | 22 |
| 55 | 26 |
| 46 | 23 |
| 324 | 100 |
| 101 | 40 |
| 217 | 72 |
| 217 | 72 |
| 1256 | 321 |
| 176 | 61 |
| 966 | 254 |
| 619 | 172 |
| 1512 | 379 |
| 877 | 233 |
| 386 | 115 |
| 295 | 92 |
| 188 | 64 |
| 188 | 64 |
| 177 | 61 |
| 137 | 50 |
| 148 | 53 |
| 69 | 30 |
| 219 | 72 |
| 106 | 41 |
| 531 | 150 |
| 231 | 75 |
| 297 | 92 |
| 235 | 76 |
| 544 | 153 |
| 76 | 32 |
| 805 | 215 |
| 135 | 49 |
| 1234 | 314 |
| 310 | 95 |
| 905 | 238 |
| 51 | 24 |

| | |
|---|---|
| 953 | 249 |
| 872 | 230 |
| 497 | 141 |
| 371 | 110 |
| 28 | 16 |
| 64 | 28 |
| 94 | 37 |
| 400 | 117 |
| 61 | 27 |
| 166 | 57 |
| 181 | 61 |
| 23 | 14 |
| 137 | 49 |
| 910 | 238 |
| 471 | 134 |
| 418 | 121 |
| 928 | 242 |
| 243 | 77 |
| 342 | 102 |
| 138 | 49 |
| 149 | 52 |
| 99 | 38 |
| 1496 | 371 |
| 661 | 179 |
| 244 | 77 |
| 142 | 50 |
| 1031 | 265 |
| 202 | 66 |
| 16 | 11 |
| 117 | 43 |
| 237 | 75 |
| 241 | 76 |
| 667 | 180 |
| 1552 | 383 |
| 150 | 52 |
| 512 | 143 |
| 76 | 31 |
| 538 | 149 |
| 1541 | 380 |
| 136 | 48 |
| 44 | 21 |
| 242 | 76 |
| 41 | 20 |
| 219 | 70 |
| 423 | 121 |
| 24 | 14 |
| 14 | 10 |
| 354 | 104 |

| | |
|---|---|
| 721 | 192 |
| 967 | 249 |
| 159 | 54 |
| 174 | 58 |
| 57 | 25 |
| 259 | 80 |
| 137 | 48 |
| 420 | 120 |
| 220 | 70 |
| 752 | 199 |
| 881 | 229 |
| 119 | 43 |
| 495 | 138 |
| 1703 | 415 |
| 98 | 37 |
| 450 | 127 |
| 360 | 105 |
| 328 | 97 |
| 12 | 9 |
| 316 | 94 |
| 109 | 40 |
| 33 | 17 |
| 33 | 17 |
| 187 | 61 |
| 522 | 144 |
| 357 | 104 |
| 564 | 154 |
| 1706 | 415 |
| 620 | 167 |
| 354 | 103 |
| 650 | 174 |
| 55 | 24 |
| 250 | 77 |
| 169 | 56 |
| 65 | 27 |
| 223 | 70 |
| 132 | 46 |
| 132 | 46 |
| 132 | 46 |
| 437 | 123 |
| 96 | 36 |
| 96 | 36 |
| 177 | 58 |
| 643 | 172 |
| 49 | 22 |
| 323 | 95 |
| 100 | 37 |
| 93 | 35 |

| | |
|---|---|
| 717 | 189 |
| 236 | 73 |
| 111 | 40 |
| 182 | 59 |
| 104 | 38 |
| 115 | 41 |
| 90 | 34 |
| 115 | 41 |
| 83 | 32 |
| 225 | 70 |
| 15 | 10 |
| 15 | 10 |
| 237 | 73 |
| 249 | 76 |
| 604 | 162 |
| 297 | 88 |
| 73 | 29 |
| 73 | 29 |
| 87 | 33 |
| 537 | 146 |
| 112 | 40 |
| 112 | 40 |
| 483 | 133 |
| 127 | 44 |
| 396 | 112 |
| 262 | 79 |
| 1724 | 416 |
| 98 | 36 |
| 91 | 34 |
| 1026 | 259 |
| 778 | 202 |
| 409 | 115 |
| 26 | 14 |
| 26 | 14 |
| 26 | 14 |
| 323 | 94 |
| 1852 | 444 |
| 57 | 24 |
| 192 | 61 |
| 74 | 29 |
| 95 | 35 |
| 74 | 29 |
| 633 | 168 |
| 38 | 18 |
| 381 | 108 |
| 923 | 235 |
| 18 | 11 |
| 18 | 11 |

| | |
|---|---|
| 1338 | 329 |
| 625 | 166 |
| 54 | 23 |
| 369 | 105 |
| 681 | 179 |
| 228 | 70 |
| 99 | 36 |
| 968 | 245 |
| 51 | 22 |
| 85 | 32 |
| 85 | 32 |
| 151 | 50 |
| 151 | 50 |
| 61 | 25 |
| 428 | 119 |
| 114 | 40 |
| 597 | 159 |
| 466 | 128 |
| 965 | 244 |
| 48 | 21 |
| 68 | 27 |
| 342 | 98 |
| 129 | 44 |
| 107 | 38 |
| 1972 | 469 |
| 662 | 174 |
| 718 | 187 |
| 148 | 49 |
| 1116 | 278 |
| 290 | 85 |
| 745 | 193 |
| 24 | 13 |
| 24 | 13 |
| 24 | 13 |
| 242 | 73 |
| 250 | 75 |
| 489 | 133 |
| 16 | 10 |
| 97 | 35 |
| 149 | 49 |
| 119 | 41 |
| 33 | 16 |
| 490 | 133 |
| 219 | 67 |
| 176 | 56 |
| 507 | 137 |
| 90 | 33 |
| 69 | 27 |

| | |
|---|---|
| 69 | 27 |
| 243 | 73 |
| 123 | 42 |
| 76 | 29 |
| 138 | 46 |
| 499 | 135 |
| 627 | 165 |
| 432 | 119 |
| 127 | 43 |
| 94 | 34 |
| 204 | 63 |
| 146 | 48 |
| 2358 | 552 |
| 9 | 7 |
| 9 | 7 |
| 366 | 103 |
| 46 | 20 |
| 46 | 20 |
| 611 | 161 |
| 594 | 157 |
| 317 | 91 |
| 577 | 153 |
| 577 | 153 |
| 492 | 133 |
| 98 | 35 |
| 109 | 38 |
| 19 | 11 |
| 986 | 247 |
| 501 | 135 |
| 1017 | 254 |
| 535 | 143 |
| 367 | 103 |
| 91 | 33 |
| 91 | 33 |
| 102 | 36 |
| 213 | 65 |
| 451 | 123 |
| 77 | 29 |
| 53 | 22 |
| 22 | 12 |
| 592 | 156 |
| 190 | 59 |
| 88 | 32 |
| 34 | 16 |
| 222 | 67 |
| 230 | 69 |
| 74 | 28 |
| 171 | 54 |

| | |
|---|---|
| 28 | 14 |
| 28 | 14 |
| 495 | 133 |
| 415 | 114 |
| 336 | 95 |
| 365 | 102 |
| 291 | 84 |
| 103 | 36 |
| 57 | 23 |
| 187 | 58 |
| 187 | 58 |
| 47 | 20 |
| 118 | 40 |
| 118 | 40 |
| 156 | 50 |
| 156 | 50 |
| 1253 | 306 |
| 676 | 175 |
| 107 | 37 |
| 215 | 65 |
| 215 | 65 |
| 616 | 161 |
| 122 | 41 |
| 44 | 19 |
| 484 | 130 |
| 54 | 22 |
| 149 | 48 |
| 346 | 97 |
| 17 | 10 |
| 17 | 10 |
| 527 | 140 |
| 17 | 10 |
| 200 | 61 |
| 93 | 33 |
| 224 | 67 |
| 220 | 66 |
| 355 | 99 |
| 51 | 21 |
| 165 | 52 |
| 123 | 41 |
| 97 | 34 |
| 20 | 11 |
| 406 | 111 |
| 298 | 85 |
| 602 | 157 |
| 72 | 27 |
| 356 | 99 |
| 154 | 49 |

| | |
|---|---|
| 48 | 20 |
| 90 | 32 |
| 32 | 15 |
| 23 | 12 |
| 94 | 33 |
| 424 | 115 |
| 403 | 110 |
| 76 | 28 |
| 45 | 19 |
| 124 | 41 |
| 124 | 41 |
| 147 | 47 |
| 109 | 37 |
| 147 | 47 |
| 324 | 91 |
| 454 | 122 |
| 535 | 141 |
| 374 | 103 |
| 3210 | 732 |
| 463 | 124 |
| 523 | 138 |
| 42 | 18 |
| 1097 | 269 |
| 91 | 32 |
| 247 | 72 |
| 247 | 72 |
| 179 | 55 |
| 371 | 102 |
| 144 | 46 |
| 144 | 46 |
| 191 | 58 |
| 15 | 9 |
| 106 | 36 |
| 15 | 9 |
| 59 | 23 |
| 15 | 9 |
| 227 | 67 |
| 59 | 23 |
| 15 | 9 |
| 66 | 25 |
| 15 | 9 |
| 477 | 127 |
| 671 | 172 |
| 95 | 33 |
| 39 | 17 |
| 10 | 7 |
| 10 | 7 |
| 10 | 7 |

| | |
|---|---|
| 10 | 7 |
| 156 | 49 |
| 49 | 20 |
| 129 | 42 |
| 542 | 142 |
| 293 | 83 |
| 555 | 145 |
| 137 | 44 |
| 137 | 44 |
| 224 | 66 |
| 874 | 218 |
| 63 | 24 |
| 655 | 168 |
| 402 | 109 |
| 208 | 62 |
| 46 | 19 |
| 81 | 29 |
| 46 | 19 |
| 18 | 10 |
| 18 | 10 |
| 18 | 10 |
| 33 | 15 |
| 33 | 15 |
| 33 | 15 |
| 33 | 15 |
| 1173 | 285 |
| 249 | 72 |
| 249 | 72 |
| 1547 | 368 |
| 74 | 27 |
| 889 | 221 |
| 53 | 21 |
| 67 | 25 |
| 21 | 11 |
| 213 | 63 |
| 27 | 13 |
| 27 | 13 |
| 43 | 18 |
| 193 | 58 |
| 412 | 111 |
| 24 | 12 |
| 24 | 12 |
| 266 | 76 |
| 391 | 106 |
| 142 | 45 |
| 320 | 89 |
| 119 | 39 |
| 502 | 132 |

| | |
|---|---|
| 146 | 46 |
| 610 | 157 |
| 728 | 184 |
| 238 | 69 |
| 50 | 20 |
| 158 | 49 |
| 40 | 17 |
| 8 | 6 |
| 8 | 6 |
| 8 | 6 |
| 57 | 22 |
| 82 | 29 |
| 218 | 64 |
| 182 | 55 |
| 598 | 154 |
| 131 | 42 |
| 585 | 151 |
| 182 | 55 |
| 405 | 109 |
| 112 | 37 |
| 97 | 33 |
| 13 | 8 |
| 13 | 8 |
| 13 | 8 |
| 75 | 27 |
| 75 | 27 |
| 37 | 16 |
| 37 | 16 |
| 831 | 207 |
| 47 | 19 |
| 47 | 19 |
| 351 | 96 |
| 120 | 39 |
| 322 | 89 |
| 61 | 23 |
| 163 | 50 |
| 90 | 31 |
| 124 | 40 |
| 211 | 62 |
| 167 | 51 |
| 171 | 52 |
| 183 | 55 |
| 34 | 15 |
| 34 | 15 |
| 34 | 15 |
| 109 | 36 |
| 815 | 203 |
| 4 | 4 |

| | |
|---|---|
| 4 | 4 |
| 4 | 4 |
| 344 | 94 |
| 6 | 5 |
| 6 | 5 |
| 719 | 181 |
| 31 | 14 |
| 148 | 46 |
| 16 | 9 |
| 16 | 9 |
| 265 | 75 |
| 319 | 88 |
| 51 | 20 |
| 168 | 51 |
| 76 | 27 |
| 76 | 27 |
| 41 | 17 |
| 434 | 115 |
| 106 | 35 |
| 383 | 103 |
| 237 | 68 |
| 91 | 31 |
| 25 | 12 |
| 25 | 12 |
| 25 | 12 |
| 747 | 187 |
| 22 | 11 |
| 110 | 36 |
| 316 | 87 |
| 291 | 81 |
| 62 | 23 |
| 145 | 45 |
| 55 | 21 |
| 55 | 21 |
| 283 | 79 |
| 38 | 16 |
| 73 | 26 |
| 73 | 26 |
| 122 | 39 |
| 652 | 165 |
| 238 | 68 |
| 457 | 120 |
| 740 | 185 |
| 126 | 40 |
| 483 | 126 |
| 66 | 24 |
| 66 | 24 |
| 66 | 24 |

| | |
|---|---|
| 66 | 24 |
| 11 | 7 |
| 11 | 7 |
| 11 | 7 |
| 45 | 18 |
| 45 | 18 |
| 214 | 62 |
| 35 | 15 |
| 138 | 43 |
| 59 | 22 |
| 59 | 22 |
| 35 | 15 |
| 59 | 22 |
| 35 | 15 |
| 146 | 45 |
| 146 | 45 |
| 276 | 77 |
| 1586 | 373 |
| 186 | 55 |
| 81 | 28 |
| 70 | 25 |
| 119 | 38 |
| 606 | 154 |
| 773 | 192 |
| 369 | 99 |
| 260 | 73 |
| 285 | 79 |
| 32 | 14 |
| 100 | 33 |
| 63 | 23 |
| 42 | 17 |
| 929 | 227 |
| 85 | 29 |
| 281 | 78 |
| 63 | 23 |
| 215 | 62 |
| 104 | 34 |
| 89 | 30 |
| 89 | 30 |
| 89 | 30 |
| 108 | 35 |
| 147 | 45 |
| 147 | 45 |
| 29 | 13 |
| 29 | 13 |
| 151 | 46 |
| 29 | 13 |
| 29 | 13 |

| | |
|---|---|
| 167 | 50 |
| 599 | 152 |
| 14 | 8 |
| 14 | 8 |
| 14 | 8 |
| 14 | 8 |
| 328 | 89 |
| 228 | 65 |
| 93 | 31 |
| 116 | 37 |
| 39 | 16 |
| 253 | 71 |
| 26 | 12 |
| 26 | 12 |
| 435 | 114 |
| 60 | 22 |
| 838 | 206 |
| 388 | 103 |
| 82 | 28 |
| 204 | 59 |
| 241 | 68 |
| 53 | 20 |
| 132 | 41 |
| 17 | 9 |
| 17 | 9 |
| 136 | 42 |
| 23 | 11 |
| 995 | 241 |
| 233 | 66 |
| 184 | 54 |
| 440 | 115 |
| 144 | 44 |
| 1094 | 263 |
| 20 | 10 |
| 105 | 34 |
| 229 | 65 |
| 164 | 49 |
| 36 | 15 |
| 300 | 82 |
| 667 | 167 |
| 338 | 91 |
| 698 | 174 |
| 90 | 30 |
| 317 | 86 |
| 213 | 61 |
| 242 | 68 |
| 117 | 37 |
| 205 | 59 |

| | |
|---|---|
| 205 | 59 |
| 205 | 59 |
| 9 | 6 |
| 238 | 67 |
| 9 | 6 |
| 9 | 6 |
| 9 | 6 |
| 9 | 6 |
| 681 | 170 |
| 9 | 6 |
| 9 | 6 |
| 309 | 84 |
| 43 | 17 |
| 197 | 57 |
| 79 | 27 |
| 263 | 73 |
| 125 | 39 |
| 125 | 39 |
| 50 | 19 |
| 446 | 116 |
| 177 | 52 |
| 149 | 45 |
| 616 | 155 |
| 102 | 33 |
| 106 | 34 |
| 106 | 34 |
| 61 | 22 |
| 61 | 22 |
| 72 | 25 |
| 72 | 25 |
| 72 | 25 |
| 490 | 126 |
| 705 | 175 |
| 705 | 175 |
| 40 | 16 |
| 54 | 20 |
| 194 | 56 |
| 30 | 13 |
| 30 | 13 |
| 76 | 26 |
| 47 | 18 |
| 1548 | 362 |
| 47 | 18 |
| 76 | 26 |
| 47 | 18 |
| 750 | 185 |
| 750 | 185 |
| 126 | 39 |

| | |
|---|---|
| 396 | 104 |
| 65 | 23 |
| 465 | 120 |
| 12345 | 3127 |
| 2134 | 729 |
| 2133 | 729 |
| 2134 | 729 |
| 2109 | 721 |
| 4004 | 1179 |
| 3303 | 974 |
| 4213 | 1164 |
| 4777 | 1280 |
| 9864 | 2347 |
| 117 | 76 |
| 21548 | 4529 |
| 2055 | 600 |
| 5487 | 1379 |
| 15671 | 3479 |
| 57 | 44 |
| 425 | 168 |
| 3770 | 985 |
| 192 | 93 |
| 2513 | 686 |
| 2518 | 687 |
| 2389 | 656 |
| 2277 | 628 |
| 313 | 127 |
| 2230 | 610 |
| 16808 | 3671 |
| 556 | 195 |
| 17834 | 3866 |
| 432 | 160 |
| 555 | 194 |
| 6228 | 1499 |
| 1069 | 326 |
| 422 | 156 |
| 905 | 284 |
| 896 | 280 |
| 768 | 247 |
| 1689 | 472 |
| 93 | 52 |
| 44 | 32 |
| 96 | 52 |
| 104 | 54 |
| 117 | 57 |
| 275 | 105 |
| 855 | 257 |
| 1521 | 415 |

| | |
|---|---|
| 2203 | 572 |
| 762 | 230 |
| 762 | 230 |
| 132 | 59 |
| 323 | 114 |
| 320 | 113 |
| 252 | 94 |
| 324 | 114 |
| 162 | 67 |
| 5742 | 1343 |
| 1334 | 363 |
| 1217 | 332 |
| 831 | 239 |
| 242 | 88 |
| 1200 | 327 |
| 1468 | 389 |
| 187 | 72 |
| 1194 | 324 |
| 220 | 81 |
| 1299 | 348 |
| 1299 | 348 |
| 2430 | 606 |
| 782 | 224 |
| 79 | 38 |
| 1062 | 289 |
| 178 | 67 |
| 134 | 54 |
| 124 | 51 |
| 149 | 58 |
| 1054 | 284 |
| 168 | 63 |
| 40 | 23 |
| 1266 | 332 |
| 184 | 67 |
| 1401 | 363 |
| 568 | 166 |
| 1629 | 415 |
| 512 | 152 |
| 53 | 27 |
| 53 | 27 |
| 53 | 27 |
| 1405 | 362 |
| 16018 | 3432 |
| 1426 | 365 |
| 92 | 39 |
| 14605 | 3143 |
| 1240 | 320 |
| 23 | 15 |

| | |
|---|---|
| 496 | 144 |
| 1226 | 316 |
| 173 | 61 |
| 306 | 96 |
| 736 | 201 |
| 26 | 16 |
| 174 | 61 |
| 100 | 40 |
| 172 | 60 |
| 515 | 147 |
| 446 | 130 |
| 565 | 159 |
| 446 | 130 |
| 133 | 49 |
| 489 | 140 |
| 445 | 129 |
| 167 | 58 |
| 472 | 135 |
| 111 | 42 |
| 111 | 42 |
| 903 | 236 |
| 55 | 25 |
| 656 | 178 |
| 477 | 135 |
| 498 | 140 |
| 59 | 26 |
| 121 | 44 |
| 32 | 17 |
| 432 | 123 |
| 24 | 14 |
| 54 | 24 |
| 1060 | 269 |
| 519 | 143 |
| 55 | 24 |
| 154 | 52 |
| 154 | 52 |
| 154 | 52 |
| 8 | 7 |
| 299 | 89 |
| 28 | 15 |
| 300 | 89 |
| 261 | 79 |
| 281 | 84 |
| 1776 | 427 |
| 57 | 24 |
| 64 | 26 |
| 71 | 28 |
| 163 | 53 |

| | |
|---|---|
| 21 | 12 |
| 126 | 43 |
| 11 | 8 |
| 865 | 220 |
| 316 | 91 |
| 16 | 10 |
| 248 | 74 |
| 384 | 107 |
| 473 | 128 |
| 567 | 150 |
| 159 | 51 |
| 25 | 13 |
| 168 | 53 |
| 89 | 32 |
| 41 | 18 |
| 116 | 39 |
| 848 | 213 |
| 721 | 184 |
| 29 | 14 |
| 26 | 13 |
| 91 | 32 |
| 650 | 167 |
| 256 | 74 |
| 318 | 89 |
| 254 | 73 |
| 43 | 18 |
| 346 | 95 |
| 334 | 92 |
| 75 | 27 |
| 199 | 59 |
| 356 | 97 |
| 90 | 31 |
| 4 | 4 |
| 6 | 5 |
| 41 | 17 |
| 22 | 11 |
| 11 | 7 |
| 146 | 45 |
| 146 | 45 |
| 407 | 108 |
| 174 | 52 |
| 96 | 32 |
| 223 | 64 |
| 564 | 144 |
| 366 | 98 |
| 108 | 35 |
| 93 | 31 |
| 14 | 8 |

| | |
|---|---|
| 14 | 8 |
| 14 | 8 |
| 279 | 77 |
| 20 | 10 |
| 57 | 21 |
| 169 | 50 |
| 9 | 6 |
| 264 | 73 |
| 1436 | 337 |
| 427 | 111 |
| 384 | 101 |
| 80 | 27 |
| 312 | 84 |
| 84 | 28 |
| 12 | 7 |
| 24 | 11 |
| 4220 | 933 |
| 15 | 8 |
| 18 | 9 |
| 18 | 9 |
| 48 | 18 |
| 70 | 24 |
| 128 | 39 |
| 59 | 21 |
| 6657 | 1444 |
| 6657 | 1444 |
| 6657 | 1444 |
| 141 | 42 |
| 25 | 11 |
| 726 | 177 |
| 5 | 4 |
| 5 | 4 |
| 10 | 6 |
| 212 | 59 |
| 10 | 6 |
| 5 | 4 |
| 10 | 6 |
| 5 | 4 |
| 5 | 4 |
| 314 | 83 |
| 46 | 17 |
| 46 | 17 |
| 32 | 13 |
| 14838 | 3775 |
| 6357 | 1754 |
| 6146 | 1693 |
| 4444 | 1247 |
| 4354 | 1220 |

| | |
|---|---|
| 742 | 278 |
| 1383 | 453 |
| 1439 | 468 |
| 18370 | 4271 |
| 1480 | 478 |
| 1482 | 478 |
| 1351 | 437 |
| 1541 | 487 |
| 1654 | 516 |
| 1637 | 511 |
| 1180 | 383 |
| 1203 | 389 |
| 1952 | 574 |
| 2532 | 699 |
| 115 | 57 |
| 5758 | 1460 |
| 128 | 60 |
| 2162 | 596 |
| 138 | 62 |
| 66 | 36 |
| 332 | 116 |
| 323 | 113 |
| 167 | 67 |
| 134 | 56 |
| 158 | 63 |
| 57 | 30 |
| 561 | 175 |
| 88 | 40 |
| 726 | 216 |
| 2452 | 640 |
| 206 | 75 |
| 17 | 13 |
| 2325 | 608 |
| 113 | 47 |
| 45 | 24 |
| 2103 | 551 |
| 2187 | 571 |
| 2188 | 571 |
| 41 | 22 |
| 2304 | 597 |
| 42 | 22 |
| 60 | 28 |
| 1661 | 438 |
| 30 | 17 |
| 44 | 22 |
| 2021 | 523 |
| 69 | 30 |
| 166 | 59 |

| | |
|---|---|
| 841 | 235 |
| 66 | 29 |
| 60 | 27 |
| 1752 | 457 |
| 1239 | 332 |
| 24 | 14 |
| 268 | 86 |
| 30 | 16 |
| 790 | 219 |
| 42 | 20 |
| 25 | 14 |
| 20 | 12 |
| 28 | 15 |
| 28 | 15 |
| 15 | 10 |
| 15 | 10 |
| 28 | 15 |
| 1100 | 294 |
| 692 | 193 |
| 1127 | 300 |
| 52 | 23 |
| 52 | 23 |
| 23 | 13 |
| 23 | 13 |
| 13 | 9 |
| 115 | 42 |
| 56 | 24 |
| 5 | 5 |
| 41 | 19 |
| 41 | 19 |
| 2478 | 620 |
| 1501 | 388 |
| 453 | 131 |
| 7 | 6 |
| 569 | 160 |
| 33 | 16 |
| 45 | 20 |
| 19 | 11 |
| 247 | 77 |
| 74 | 29 |
| 36 | 17 |
| 1002 | 266 |
| 82 | 31 |
| 25 | 13 |
| 25 | 13 |
| 28 | 14 |
| 1560 | 399 |
| 37 | 17 |

| | |
|---|---|
| 190 | 61 |
| 37 | 17 |
| 473 | 134 |
| 20 | 11 |
| 1936 | 487 |
| 1919 | 483 |
| 479 | 135 |
| 144 | 48 |
| 483 | 136 |
| 506 | 218 |
| 153 | 88 |
| 231 | 110 |
| 1537 | 490 |
| 265 | 108 |
| 202 | 87 |
| 134 | 62 |
| 271 | 107 |
| 305 | 117 |
| 474 | 169 |
| 115 | 53 |
| 363 | 133 |
| 221 | 88 |
| 154 | 63 |
| 89 | 41 |
| 20 | 14 |
| 232 | 87 |
| 39 | 21 |
| 67 | 31 |
| 87 | 37 |
| 382 | 127 |
| 57 | 26 |
| 43 | 21 |
| 47 | 22 |
| 31 | 16 |
| 69 | 29 |
| 89 | 35 |
| 30 | 15 |
| 74 | 30 |
| 94 | 80 |
| 90 | 76 |
| 102 | 81 |
| 94 | 76 |
| 94 | 76 |
| 90 | 74 |
| 112 | 84 |
| 102 | 78 |
| 96 | 75 |
| 117 | 85 |

| | |
|---|---|
| 113 | 83 |
| 113 | 83 |
| 120 | 86 |
| 120 | 86 |
| 157 | 95 |
| 116 | 75 |
| 116 | 75 |
| 370 | 169 |
| 133 | 79 |
| 171 | 94 |
| 2074 | 667 |
| 152 | 83 |
| 37 | 32 |
| 292 | 130 |
| 217 | 103 |
| 574 | 218 |
| 60 | 41 |
| 549 | 208 |
| 52 | 37 |
| 59 | 40 |
| 59 | 39 |
| 780 | 273 |
| 794 | 276 |
| 127 | 62 |
| 51 | 32 |
| 182 | 80 |
| 103 | 52 |
| 192 | 82 |
| 177 | 77 |
| 26 | 20 |
| 1234 | 383 |
| 297 | 114 |
| 95 | 47 |
| 202 | 83 |
| 84 | 42 |
| 476 | 161 |
| 140 | 59 |
| 89 | 41 |
| 75 | 36 |
| 1355 | 401 |
| 33 | 20 |
| 60 | 30 |
| 39 | 22 |
| 15 | 11 |
| 15 | 11 |
| 67 | 31 |
| 42 | 22 |
| 13 | 10 |

| | |
|---|---|
| 22 | 14 |
| 1946 | 548 |
| 57 | 27 |
| 30 | 17 |
| 6 | 6 |
| 64 | 29 |
| 195 | 70 |
| 152 | 56 |
| 22 | 13 |
| 15 | 10 |
| 130 | 49 |
| 286 | 95 |
| 124 | 47 |
| 33 | 17 |
| 999 | 290 |
| 59 | 26 |
| 125 | 47 |
| 11 | 8 |
| 23 | 13 |
| 7 | 6 |
| 60 | 26 |
| 107 | 41 |
| 40 | 19 |
| 61 | 26 |
| 86 | 34 |
