## Supplementary file S3 for "Resveratrol targets G-quadruplexes to exert its pharmacological effects"

**Supplementary file 3. Functional enrichment analysis of downregulated DEGs**

| term_name | term_id | source | p_value |
|---|---|---|---|
| regulation of nitrogen compound metabolic process | GO:0051171 | GO:BP | 9.78E-128 |
| regulation of primary metabolic process | GO:0080090 | GO:BP | 1.23E-126 |
| macromolecule modification | GO:0043412 | GO:BP | 2.02E-117 |
| cell cycle | GO:0007049 | GO:BP | 3.37E-113 |
| organelle organization | GO:0006996 | GO:BP | 7.75E-102 |
| regulation of nucleobase-containing compound metabo | GO:0019219 | GO:BP | 1.50E-99 |
| cellular nitrogen compound biosynthetic process | GO:0044271 | GO:BP | 1.26E-95 |
| protein modification process | GO:0036211 | GO:BP | 1.26E-95 |
| cell cycle process | GO:0022402 | GO:BP | 3.80E-93 |
| positive regulation of biological process | GO:0048518 | GO:BP | 4.31E-93 |
| positive regulation of nitrogen compound metabolic pro | GO:0051173 | GO:BP | 2.75E-92 |
| positive regulation of cellular process | GO:0048522 | GO:BP | 1.28E-90 |
| protein metabolic process | GO:0019538 | GO:BP | 2.04E-86 |
| regulation of RNA metabolic process | GO:0051252 | GO:BP | 1.99E-85 |
| mitotic cell cycle | GO:0000278 | GO:BP | 1.40E-83 |
| positive regulation of macromolecule metabolic proces | GO:0010604 | GO:BP | 5.04E-82 |
| organonitrogen compound metabolic process | GO:1901564 | GO:BP | 3.36E-81 |
| regulation of metabolic process | GO:0019222 | GO:BP | 1.25E-79 |
| heterocycle biosynthetic process | GO:0018130 | GO:BP | 3.67E-79 |
| positive regulation of nucleobase-containing compounc | GO:0045935 | GO:BP | 4.27E-79 |
| aromatic compound biosynthetic process | GO:0019438 | GO:BP | 1.78E-78 |
| positive regulation of metabolic process | GO:0009893 | GO:BP | 4.32E-78 |
| nucleobase-containing compound biosynthetic process | GO:0034654 | GO:BP | 1.39E-77 |
| regulation of macromolecule metabolic process | GO:0060255 | GO:BP | 6.96E-77 |
| mitotic cell cycle process | GO:1903047 | GO:BP | 1.03E-73 |
| organic cyclic compound biosynthetic process | GO:1901362 | GO:BP | 3.42E-73 |
| biological regulation | GO:0065007 | GO:BP | 1.06E-72 |
| regulation of biological process | GO:0050789 | GO:BP | 4.86E-72 |
| positive regulation of RNA metabolic process | GO:0051254 | GO:BP | 1.56E-71 |
| RNA biosynthetic process | GO:0032774 | GO:BP | 1.60E-71 |
| DNA-templated transcription | GO:0006351 | GO:BP | 1.50E-70 |
| regulation of cellular process | GO:0050794 | GO:BP | 3.31E-70 |
| regulation of cellular metabolic process | GO:0031323 | GO:BP | 3.31E-70 |
| DNA metabolic process | GO:0006259 | GO:BP | 4.46E-70 |
| ncRNA metabolic process | GO:0034660 | GO:BP | 1.37E-67 |
| cellular response to stress | GO:0033554 | GO:BP | 1.40E-67 |
| positive regulation of cellular metabolic process | GO:0031325 | GO:BP | 1.93E-67 |
| DNA damage response | GO:0006974 | GO:BP | 8.73E-67 |
| regulation of RNA biosynthetic process | GO:2001141 | GO:BP | 1.31E-64 |
| regulation of DNA-templated transcription | GO:0006355 | GO:BP | 1.78E-64 |
| intracellular transport | GO:0046907 | GO:BP | 8.39E-62 |
| regulation of cellular component organization | GO:0051128 | GO:BP | 1.05E-61 |
| regulation of cell cycle | GO:0051726 | GO:BP | 1.93E-60 |
| positive regulation of RNA biosynthetic process | GO:1902680 | GO:BP | 5.71E-59 |
| establishment of localization in cell | GO:0051649 | GO:BP | 2.17E-58 |
| positive regulation of DNA-templated transcription | GO:0045893 | GO:BP | 3.10E-58 |

| | | | |
|---|---|---|---|
| cell division | GO:0051301 | GO:BP | 1.62E-55 |
| nitrogen compound metabolic process | GO:0006807 | GO:BP | 2.31E-55 |
| cellular localization | GO:0051641 | GO:BP | 5.14E-55 |
| DNA repair | GO:0006281 | GO:BP | 8.79E-55 |
| chromosome organization | GO:0051276 | GO:BP | 1.84E-53 |
| ncRNA processing | GO:0034470 | GO:BP | 3.20E-53 |
| positive regulation of cellular biosynthetic process | GO:0031328 | GO:BP | 8.81E-53 |
| positive regulation of macromolecule biosynthetic proc | GO:0010557 | GO:BP | 1.08E-52 |
| protein-DNA complex organization | GO:0071824 | GO:BP | 2.58E-52 |
| positive regulation of biosynthetic process | GO:0009891 | GO:BP | 4.84E-52 |
| regulation of cell cycle process | GO:0010564 | GO:BP | 1.48E-50 |
| post-translational protein modification | GO:0043687 | GO:BP | 7.08E-50 |
| primary metabolic process | GO:0044238 | GO:BP | 1.10E-49 |
| macromolecule catabolic process | GO:0009057 | GO:BP | 3.87E-49 |
| negative regulation of nitrogen compound metabolic pr | GO:0051172 | GO:BP | 5.42E-49 |
| cell cycle phase transition | GO:0044770 | GO:BP | 2.19E-48 |
| intracellular signal transduction | GO:0035556 | GO:BP | 5.99E-48 |
| chromosome segregation | GO:0007059 | GO:BP | 4.46E-47 |
| transcription by RNA polymerase II | GO:0006366 | GO:BP | 4.55E-47 |
| regulation of biosynthetic process | GO:0009889 | GO:BP | 1.23E-46 |
| protein modification by small protein conjugation or ren | GO:0070647 | GO:BP | 1.47E-46 |
| regulation of cellular biosynthetic process | GO:0031326 | GO:BP | 3.21E-46 |
| regulation of organelle organization | GO:0033043 | GO:BP | 5.38E-46 |
| regulation of macromolecule biosynthetic process | GO:0010556 | GO:BP | 5.57E-46 |
| protein modification by small protein conjugation | GO:0032446 | GO:BP | 5.70E-45 |
| macromolecule localization | GO:0033036 | GO:BP | 2.90E-44 |
| regulation of protein metabolic process | GO:0051246 | GO:BP | 4.39E-44 |
| negative regulation of cellular process | GO:0048523 | GO:BP | 4.80E-44 |
| regulation of gene expression | GO:0010468 | GO:BP | 3.02E-43 |
| mitotic cell cycle phase transition | GO:0044772 | GO:BP | 7.44E-43 |
| ribosome biogenesis | GO:0042254 | GO:BP | 1.86E-42 |
| chromatin organization | GO:0006325 | GO:BP | 1.99E-42 |
| regulation of transcription by RNA polymerase II | GO:0006357 | GO:BP | 3.18E-42 |
| cellular macromolecule localization | GO:0070727 | GO:BP | 5.22E-42 |
| protein localization | GO:0008104 | GO:BP | 2.58E-41 |
| negative regulation of biological process | GO:0048519 | GO:BP | 3.32E-41 |
| cellular component organization or biogenesis | GO:0071840 | GO:BP | 6.82E-41 |
| negative regulation of nucleobase-containing compoun | GO:0045934 | GO:BP | 1.53E-40 |
| regulation of DNA metabolic process | GO:0051052 | GO:BP | 2.95E-39 |
| rRNA metabolic process | GO:0016072 | GO:BP | 5.73E-39 |
| nitrogen compound transport | GO:0071705 | GO:BP | 2.25E-38 |
| protein ubiquitination | GO:0016567 | GO:BP | 3.82E-38 |
| localization | GO:0051179 | GO:BP | 6.85E-38 |
| protein transport | GO:0015031 | GO:BP | 1.53E-37 |
| sister chromatid segregation | GO:0000819 | GO:BP | 3.67E-35 |
| negative regulation of RNA metabolic process | GO:0051253 | GO:BP | 4.89E-35 |
| regulation of mitotic cell cycle | GO:0007346 | GO:BP | 2.00E-34 |
| positive regulation of transcription by RNA polymerase I | GO:0045944 | GO:BP | 2.16E-34 |

| | | | |
|---|---|---|---|
| catabolic process | GO:0009056 | GO:BP | 2.75E-34 |
| regulation of cell cycle phase transition | GO:1901987 | GO:BP | 3.98E-34 |
| chromatin remodeling | GO:0006338 | GO:BP | 1.03E-33 |
| multicellular organism development | GO:0007275 | GO:BP | 1.16E-33 |
| developmental process | GO:0032502 | GO:BP | 2.04E-33 |
| regulation of signal transduction | GO:0009966 | GO:BP | 2.07E-33 |
| rRNA processing | GO:0006364 | GO:BP | 9.37E-33 |
| regulation of catabolic process | GO:0009894 | GO:BP | 2.31E-32 |
| regulation of chromosome organization | GO:0033044 | GO:BP | 2.44E-32 |
| DNA replication | GO:0006260 | GO:BP | 3.15E-32 |
| regulation of response to stimulus | GO:0048583 | GO:BP | 3.75E-32 |
| mitotic nuclear division | GO:0140014 | GO:BP | 3.88E-32 |
| phosphorus metabolic process | GO:0006793 | GO:BP | 4.15E-32 |
| nuclear chromosome segregation | GO:0098813 | GO:BP | 5.75E-32 |
| anatomical structure development | GO:0048856 | GO:BP | 1.01E-31 |
| phosphate-containing compound metabolic process | GO:0006796 | GO:BP | 1.55E-31 |
| cytoskeleton organization | GO:0007010 | GO:BP | 3.42E-31 |
| organelle localization | GO:0051640 | GO:BP | 1.09E-30 |
| double-strand break repair | GO:0006302 | GO:BP | 1.35E-30 |
| regulation of cellular response to stress | GO:0080135 | GO:BP | 1.49E-30 |
| organic substance transport | GO:0071702 | GO:BP | 5.16E-30 |
| modification-dependent macromolecule catabolic proc | GO:0043632 | GO:BP | 7.71E-30 |
| regulation of mitotic cell cycle phase transition | GO:1901990 | GO:BP | 8.15E-30 |
| mitotic sister chromatid segregation | GO:0000070 | GO:BP | 1.10E-29 |
| organelle fission | GO:0048285 | GO:BP | 2.57E-29 |
| nuclear transport | GO:0051169 | GO:BP | 8.36E-29 |
| nucleocytoplasmic transport | GO:0006913 | GO:BP | 8.36E-29 |
| organic substance catabolic process | GO:1901575 | GO:BP | 9.21E-29 |
| regulation of signaling | GO:0023051 | GO:BP | 9.88E-29 |
| intracellular protein transport | GO:0006886 | GO:BP | 2.09E-28 |
| ubiquitin-dependent protein catabolic process | GO:0006511 | GO:BP | 2.10E-28 |
| cellular response to stimulus | GO:0051716 | GO:BP | 2.21E-28 |
| regulation of mRNA metabolic process | GO:1903311 | GO:BP | 1.33E-27 |
| negative regulation of DNA-templated transcription | GO:0045892 | GO:BP | 1.44E-27 |
| cellular component organization | GO:0016043 | GO:BP | 1.90E-27 |
| regulation of cell communication | GO:0010646 | GO:BP | 2.15E-27 |
| modification-dependent protein catabolic process | GO:0019941 | GO:BP | 2.45E-27 |
| phosphorylation | GO:0016310 | GO:BP | 3.24E-27 |
| negative regulation of RNA biosynthetic process | GO:1902679 | GO:BP | 3.80E-27 |
| protein catabolic process | GO:0030163 | GO:BP | 4.02E-27 |
| proteolysis involved in protein catabolic process | GO:0051603 | GO:BP | 4.06E-27 |
| nuclear division | GO:0000280 | GO:BP | 5.57E-27 |
| regulation of cellular component biogenesis | GO:0044087 | GO:BP | 8.50E-27 |
| vesicle-mediated transport | GO:0016192 | GO:BP | 1.03E-26 |
| microtubule cytoskeleton organization | GO:0000226 | GO:BP | 6.99E-26 |
| organonitrogen compound biosynthetic process | GO:1901566 | GO:BP | 3.33E-25 |
| establishment of protein localization | GO:0045184 | GO:BP | 3.69E-25 |
| protein polyubiquitination | GO:0000209 | GO:BP | 4.15E-25 |

| | | | |
|---|---|---|---|
| positive regulation of cellular component organization | GO:0051130 | GO:BP | 5.44E-25 |
| macromolecule metabolic process | GO:0043170 | GO:BP | 5.44E-25 |
| establishment of organelle localization | GO:0051656 | GO:BP | 5.52E-25 |
| chordate embryonic development | GO:0043009 | GO:BP | 8.22E-25 |
| embryo development ending in birth or egg hatching | GO:0009792 | GO:BP | 1.04E-24 |
| DNA-templated DNA replication | GO:0006261 | GO:BP | 1.07E-24 |
| metabolic process | GO:0008152 | GO:BP | 1.59E-24 |
| regulation of molecular function | GO:0065009 | GO:BP | 2.58E-24 |
| negative regulation of cell cycle process | GO:0010948 | GO:BP | 2.62E-24 |
| negative regulation of cell cycle | GO:0045786 | GO:BP | 3.07E-24 |
| RNA catabolic process | GO:0006401 | GO:BP | 5.29E-24 |
| establishment of localization | GO:0051234 | GO:BP | 5.72E-24 |
| transport | GO:0006810 | GO:BP | 1.44E-23 |
| regulation of developmental process | GO:0050793 | GO:BP | 1.51E-23 |
| regulation of protein modification process | GO:0031399 | GO:BP | 1.94E-23 |
| system development | GO:0048731 | GO:BP | 2.00E-23 |
| response to stress | GO:0006950 | GO:BP | 2.24E-23 |
| embryo development | GO:0009790 | GO:BP | 2.76E-23 |
| RNA localization | GO:0006403 | GO:BP | 4.30E-23 |
| regulation of DNA repair | GO:0006282 | GO:BP | 4.35E-23 |
| negative regulation of cell cycle phase transition | GO:1901988 | GO:BP | 4.45E-23 |
| organic substance metabolic process | GO:0071704 | GO:BP | 5.49E-23 |
| negative regulation of metabolic process | GO:0009892 | GO:BP | 7.89E-23 |
| regulation of cellular localization | GO:0060341 | GO:BP | 9.30E-23 |
| regulation of chromosome segregation | GO:0051983 | GO:BP | 1.47E-22 |
| nuclear export | GO:0051168 | GO:BP | 1.69E-22 |
| organelle assembly | GO:0070925 | GO:BP | 3.57E-22 |
| cell cycle checkpoint signaling | GO:0000075 | GO:BP | 7.91E-22 |
| regulation of intracellular signal transduction | GO:1902531 | GO:BP | 8.84E-22 |
| apoptotic process | GO:0006915 | GO:BP | 1.32E-21 |
| non-membrane-bounded organelle assembly | GO:0140694 | GO:BP | 1.34E-21 |
| nervous system development | GO:0007399 | GO:BP | 1.97E-21 |
| positive regulation of organelle organization | GO:0010638 | GO:BP | 2.41E-21 |
| endosomal transport | GO:0016197 | GO:BP | 2.57E-21 |
| regulation of cellular catabolic process | GO:0031329 | GO:BP | 2.60E-21 |
| negative regulation of cellular metabolic process | GO:0031324 | GO:BP | 2.98E-21 |
| negative regulation of macromolecule metabolic proces | GO:0010605 | GO:BP | 3.02E-21 |
| mRNA catabolic process | GO:0006402 | GO:BP | 3.29E-21 |
| double-strand break repair via homologous recombinat | GO:0000724 | GO:BP | 3.75E-21 |
| positive regulation of DNA metabolic process | GO:0051054 | GO:BP | 4.10E-21 |
| recombinational repair | GO:0000725 | GO:BP | 4.54E-21 |
| spindle organization | GO:0007051 | GO:BP | 4.99E-21 |
| nucleic acid transport | GO:0050657 | GO:BP | 5.03E-21 |
| RNA transport | GO:0050658 | GO:BP | 5.03E-21 |
| positive regulation of cell cycle process | GO:0090068 | GO:BP | 5.15E-21 |
| establishment of RNA localization | GO:0051236 | GO:BP | 8.15E-21 |
| programmed cell death | GO:0012501 | GO:BP | 1.15E-20 |
| DNA recombination | GO:0006310 | GO:BP | 1.16E-20 |

| | | | |
|---|---|---|---|
| cell death | GO:0008219 | GO:BP | 1.26E-20 |
| cellular nitrogen compound metabolic process | GO:0034641 | GO:BP | 2.49E-20 |
| tRNA metabolic process | GO:0006399 | GO:BP | 3.39E-20 |
| in utero embryonic development | GO:0001701 | GO:BP | 3.55E-20 |
| microtubule cytoskeleton organization involved in mitos | GO:1902850 | GO:BP | 3.82E-20 |
| protein phosphorylation | GO:0006468 | GO:BP | 4.42E-20 |
| nucleobase-containing compound catabolic process | GO:0034655 | GO:BP | 4.59E-20 |
| regulation of amide metabolic process | GO:0034248 | GO:BP | 5.45E-20 |
| cellular metabolic process | GO:0044237 | GO:BP | 5.45E-20 |
| positive regulation of cell cycle | GO:0045787 | GO:BP | 6.32E-20 |
| regulation of translation | GO:0006417 | GO:BP | 1.30E-19 |
| Golgi vesicle transport | GO:0048193 | GO:BP | 1.80E-19 |
| protein localization to organelle | GO:0033365 | GO:BP | 2.77E-19 |
| organonitrogen compound catabolic process | GO:1901565 | GO:BP | 3.08E-19 |
| heterocycle catabolic process | GO:0046700 | GO:BP | 3.95E-19 |
| proteasomal protein catabolic process | GO:0010498 | GO:BP | 3.99E-19 |
| regulation of protein localization | GO:0032880 | GO:BP | 4.05E-19 |
| small GTPase-mediated signal transduction | GO:0007264 | GO:BP | 6.11E-19 |
| positive regulation of protein metabolic process | GO:0051247 | GO:BP | 7.22E-19 |
| positive regulation of catabolic process | GO:0009896 | GO:BP | 8.41E-19 |
| macromolecule methylation | GO:0043414 | GO:BP | 9.26E-19 |
| regulation of catalytic activity | GO:0050790 | GO:BP | 1.17E-18 |
| microtubule-based process | GO:0007017 | GO:BP | 1.22E-18 |
| nucleobase-containing compound transport | GO:0015931 | GO:BP | 2.16E-18 |
| negative regulation of signal transduction | GO:0009968 | GO:BP | 3.00E-18 |
| cellular nitrogen compound catabolic process | GO:0044270 | GO:BP | 3.34E-18 |
| intracellular signaling cassette | GO:0141124 | GO:BP | 3.54E-18 |
| methylation | GO:0032259 | GO:BP | 8.17E-18 |
| RNA modification | GO:0009451 | GO:BP | 8.98E-18 |
| cellular catabolic process | GO:0044248 | GO:BP | 9.95E-18 |
| mRNA transport | GO:0051028 | GO:BP | 9.95E-18 |
| negative regulation of transcription by RNA polymerase | GO:0000122 | GO:BP | 1.14E-17 |
| aromatic compound catabolic process | GO:0019439 | GO:BP | 1.37E-17 |
| chromosome localization | GO:0050000 | GO:BP | 1.43E-17 |
| proteasome-mediated ubiquitin-dependent protein cat | GO:0043161 | GO:BP | 1.46E-17 |
| cytokinesis | GO:0000910 | GO:BP | 2.74E-17 |
| negative regulation of cellular component organization | GO:0051129 | GO:BP | 3.05E-17 |
| membrane organization | GO:0061024 | GO:BP | 4.01E-17 |
| protein localization to nucleus | GO:0034504 | GO:BP | 4.54E-17 |
| regulation of sister chromatid segregation | GO:0033045 | GO:BP | 7.34E-17 |
| regulation of cytoskeleton organization | GO:0051493 | GO:BP | 9.59E-17 |
| positive regulation of mRNA metabolic process | GO:1903313 | GO:BP | 9.92E-17 |
| DNA-templated transcription initiation | GO:0006352 | GO:BP | 9.92E-17 |
| signal transduction in response to DNA damage | GO:0042770 | GO:BP | 1.28E-16 |
| cell development | GO:0048468 | GO:BP | 1.37E-16 |
| DNA conformation change | GO:0071103 | GO:BP | 1.38E-16 |
| regulation of response to stress | GO:0080134 | GO:BP | 1.60E-16 |
| heterocycle metabolic process | GO:0046483 | GO:BP | 2.08E-16 |

| | | | |
|---|---|---|---|
| cell cycle G2/M phase transition | GO:0044839 | GO:BP | 2.44E-16 |
| regulation of RNA stability | GO:0043487 | GO:BP | 4.55E-16 |
| regulation of mRNA processing | GO:0050684 | GO:BP | 5.51E-16 |
| nucleobase-containing compound metabolic process | GO:0006139 | GO:BP | 6.00E-16 |
| establishment of chromosome localization | GO:0051303 | GO:BP | 7.38E-16 |
| regulation of DNA replication | GO:0006275 | GO:BP | 8.12E-16 |
| metaphase/anaphase transition of mitotic cell cycle | GO:0007091 | GO:BP | 8.32E-16 |
| process utilizing autophagic mechanism | GO:0061919 | GO:BP | 9.44E-16 |
| autophagy | GO:0006914 | GO:BP | 9.44E-16 |
| organic cyclic compound catabolic process | GO:1901361 | GO:BP | 9.49E-16 |
| negative regulation of mitotic cell cycle phase transitior | GO:1901991 | GO:BP | 9.49E-16 |
| DNA-templated transcription elongation | GO:0006354 | GO:BP | 1.01E-15 |
| neurogenesis | GO:0022008 | GO:BP | 1.01E-15 |
| translation | GO:0006412 | GO:BP | 1.10E-15 |
| nuclear-transcribed mRNA catabolic process | GO:0000956 | GO:BP | 1.55E-15 |
| amide biosynthetic process | GO:0043604 | GO:BP | 1.86E-15 |
| anatomical structure morphogenesis | GO:0009653 | GO:BP | 1.87E-15 |
| response to stimulus | GO:0050896 | GO:BP | 1.94E-15 |
| G1/S transition of mitotic cell cycle | GO:0000082 | GO:BP | 1.94E-15 |
| RNA export from nucleus | GO:0006405 | GO:BP | 2.39E-15 |
| metaphase chromosome alignment | GO:0051310 | GO:BP | 2.47E-15 |
| growth | GO:0040007 | GO:BP | 2.47E-15 |
| peptide biosynthetic process | GO:0043043 | GO:BP | 2.75E-15 |
| regulation of mRNA catabolic process | GO:0061013 | GO:BP | 3.10E-15 |
| positive regulation of molecular function | GO:0044093 | GO:BP | 3.40E-15 |
| cellular process | GO:0009987 | GO:BP | 4.37E-15 |
| mitotic spindle organization | GO:0007052 | GO:BP | 5.22E-15 |
| spindle assembly | GO:0051225 | GO:BP | 5.22E-15 |
| G2/M transition of mitotic cell cycle | GO:0000086 | GO:BP | 6.03E-15 |
| cellular aromatic compound metabolic process | GO:0006725 | GO:BP | 6.04E-15 |
| negative regulation of signaling | GO:0023057 | GO:BP | 7.30E-15 |
| negative regulation of mitotic cell cycle | GO:0045930 | GO:BP | 8.04E-15 |
| metaphase/anaphase transition of cell cycle | GO:0044784 | GO:BP | 8.43E-15 |
| cell projection organization | GO:0030030 | GO:BP | 1.10E-14 |
| plasma membrane bounded cell projection organizatioı | GO:0120036 | GO:BP | 1.14E-14 |
| negative regulation of cell communication | GO:0010648 | GO:BP | 1.17E-14 |
| peptidyl-amino acid modification | GO:0018193 | GO:BP | 1.23E-14 |
| transcription elongation by RNA polymerase II | GO:0006368 | GO:BP | 1.25E-14 |
| regulation of GTPase activity | GO:0043087 | GO:BP | 1.56E-14 |
| positive regulation of cellular component biogenesis | GO:0044089 | GO:BP | 1.95E-14 |
| nucleus organization | GO:0006997 | GO:BP | 2.06E-14 |
| negative regulation of response to stimulus | GO:0048585 | GO:BP | 2.10E-14 |
| regulation of double-strand break repair | GO:2000779 | GO:BP | 2.12E-14 |
| cytoskeleton-dependent cytokinesis | GO:0061640 | GO:BP | 2.36E-14 |
| nucleic acid metabolic process | GO:0090304 | GO:BP | 2.46E-14 |
| regulation of mitotic metaphase/anaphase transition | GO:0030071 | GO:BP | 3.35E-14 |
| epigenetic regulation of gene expression | GO:0040029 | GO:BP | 3.41E-14 |
| cell cycle G1/S phase transition | GO:0044843 | GO:BP | 3.46E-14 |

| | | | |
|---|---|---|---|
| mitotic cell cycle checkpoint signaling | GO:0007093 | GO:BP | 3.52E-14 |
| signal transduction | GO:0007165 | GO:BP | 4.41E-14 |
| DNA geometric change | GO:0032392 | GO:BP | 4.45E-14 |
| regulation of intracellular transport | GO:0032386 | GO:BP | 4.51E-14 |
| regulation of mRNA stability | GO:0043488 | GO:BP | 4.81E-14 |
| microtubule organizing center organization | GO:0031023 | GO:BP | 4.86E-14 |
| tRNA processing | GO:0008033 | GO:BP | 5.20E-14 |
| cellular developmental process | GO:0048869 | GO:BP | 8.66E-14 |
| mitochondrial gene expression | GO:0140053 | GO:BP | 1.06E-13 |
| cell differentiation | GO:0030154 | GO:BP | 1.07E-13 |
| stem cell population maintenance | GO:0019827 | GO:BP | 1.18E-13 |
| amide metabolic process | GO:0043603 | GO:BP | 1.19E-13 |
| maintenance of cell number | GO:0098727 | GO:BP | 2.26E-13 |
| positive regulation of cellular catabolic process | GO:0031331 | GO:BP | 2.32E-13 |
| RNA destabilization | GO:0050779 | GO:BP | 2.36E-13 |
| negative regulation of chromosome organization | GO:2001251 | GO:BP | 2.58E-13 |
| negative regulation of organelle organization | GO:0010639 | GO:BP | 2.67E-13 |
| DNA integrity checkpoint signaling | GO:0031570 | GO:BP | 2.71E-13 |
| generation of neurons | GO:0048699 | GO:BP | 2.78E-13 |
| negative regulation of cellular biosynthetic process | GO:0031327 | GO:BP | 2.78E-13 |
| signaling | GO:0023052 | GO:BP | 3.02E-13 |
| cell morphogenesis | GO:0000902 | GO:BP | 3.03E-13 |
| regulation of metaphase/anaphase transition of cell cyc | GO:1902099 | GO:BP | 3.04E-13 |
| centrosome cycle | GO:0007098 | GO:BP | 3.41E-13 |
| regulation of RNA splicing | GO:0043484 | GO:BP | 3.43E-13 |
| regulation of programmed cell death | GO:0043067 | GO:BP | 3.56E-13 |
| DNA duplex unwinding | GO:0032508 | GO:BP | 3.76E-13 |
| negative regulation of biosynthetic process | GO:0009890 | GO:BP | 3.83E-13 |
| cell communication | GO:0007154 | GO:BP | 4.16E-13 |
| positive regulation of mRNA catabolic process | GO:0061014 | GO:BP | 4.40E-13 |
| regulation of DNA-templated transcription elongation | GO:0032784 | GO:BP | 5.42E-13 |
| regulation of chromosome separation | GO:1905818 | GO:BP | 5.52E-13 |
| organic cyclic compound metabolic process | GO:1901360 | GO:BP | 6.54E-13 |
| RNA 3'-end processing | GO:0031123 | GO:BP | 6.73E-13 |
| transcription by RNA polymerase I | GO:0006360 | GO:BP | 6.85E-13 |
| regulation of transferase activity | GO:0051338 | GO:BP | 6.93E-13 |
| cellular response to endogenous stimulus | GO:0071495 | GO:BP | 7.84E-13 |
| chromosome separation | GO:0051304 | GO:BP | 8.12E-13 |
| regulation of protein modification by small protein conju | GO:1903320 | GO:BP | 1.02E-12 |
| negative regulation of macromolecule biosynthetic proc | GO:0010558 | GO:BP | 1.20E-12 |
| peptide metabolic process | GO:0006518 | GO:BP | 1.22E-12 |
| cytosolic transport | GO:0016482 | GO:BP | 1.23E-12 |
| regulation of localization | GO:0032879 | GO:BP | 1.30E-12 |
| regulation of apoptotic process | GO:0042981 | GO:BP | 1.37E-12 |
| positive regulation of DNA repair | GO:0045739 | GO:BP | 1.50E-12 |
| DNA damage checkpoint signaling | GO:0000077 | GO:BP | 1.50E-12 |
| negative regulation of protein metabolic process | GO:0051248 | GO:BP | 1.52E-12 |
| mRNA destabilization | GO:0061157 | GO:BP | 1.53E-12 |

| | | | |
|---|---|---|---|
| mitochondrion organization | GO:0007005 | GO:BP | 1.77E-12 |
| regulation of mRNA splicing, via spliceosome | GO:0048024 | GO:BP | 1.79E-12 |
| developmental growth | GO:0048589 | GO:BP | 1.93E-12 |
| negative regulation of gene expression, epigenetic | GO:0045814 | GO:BP | 2.02E-12 |
| positive regulation of protein modification process | GO:0031401 | GO:BP | 2.17E-12 |
| regulation of post-translational protein modification | GO:1901873 | GO:BP | 2.24E-12 |
| regulation of transcription elongation by RNA polymeras | GO:0034243 | GO:BP | 2.50E-12 |
| positive regulation of mitotic cell cycle | GO:0045931 | GO:BP | 2.83E-12 |
| macroautophagy | GO:0016236 | GO:BP | 4.55E-12 |
| cell-cell signaling by wnt | GO:0198738 | GO:BP | 4.55E-12 |
| establishment of cell polarity | GO:0030010 | GO:BP | 5.05E-12 |
| transcription initiation at RNA polymerase II promoter | GO:0006367 | GO:BP | 5.27E-12 |
| ncRNA transcription | GO:0098781 | GO:BP | 5.53E-12 |
| Wnt signaling pathway | GO:0016055 | GO:BP | 5.75E-12 |
| regulation of small GTPase mediated signal transductio | GO:0051056 | GO:BP | 6.07E-12 |
| protein-containing complex disassembly | GO:0032984 | GO:BP | 6.33E-12 |
| viral process | GO:0016032 | GO:BP | 7.08E-12 |
| regulation of mitotic sister chromatid segregation | GO:0033047 | GO:BP | 7.77E-12 |
| establishment or maintenance of cell polarity | GO:0007163 | GO:BP | 8.97E-12 |
| positive regulation of developmental process | GO:0051094 | GO:BP | 9.78E-12 |
| protein K48-linked ubiquitination | GO:0070936 | GO:BP | 1.04E-11 |
| ribosomal small subunit biogenesis | GO:0042274 | GO:BP | 1.24E-11 |
| endomembrane system organization | GO:0010256 | GO:BP | 1.27E-11 |
| neuron differentiation | GO:0030182 | GO:BP | 1.38E-11 |
| mitotic cytokinesis | GO:0000281 | GO:BP | 1.93E-11 |
| attachment of spindle microtubules to kinetochore | GO:0008608 | GO:BP | 2.17E-11 |
| response to endogenous stimulus | GO:0009719 | GO:BP | 2.17E-11 |
| tRNA modification | GO:0006400 | GO:BP | 2.40E-11 |
| telomere maintenance | GO:0000723 | GO:BP | 3.03E-11 |
| cellular component disassembly | GO:0022411 | GO:BP | 3.05E-11 |
| DNA biosynthetic process | GO:0071897 | GO:BP | 3.44E-11 |
| positive regulation of catalytic activity | GO:0043085 | GO:BP | 4.86E-11 |
| mRNA export from nucleus | GO:0006406 | GO:BP | 5.12E-11 |
| head development | GO:0060322 | GO:BP | 5.96E-11 |
| actin cytoskeleton organization | GO:0030036 | GO:BP | 6.08E-11 |
| regulation of DNA-templated DNA replication | GO:0090329 | GO:BP | 6.30E-11 |
| regulation of cell cycle G2/M phase transition | GO:1902749 | GO:BP | 6.56E-11 |
| regulation of cell projection organization | GO:0031344 | GO:BP | 6.75E-11 |
| RNA methylation | GO:0001510 | GO:BP | 6.89E-11 |
| intracellular receptor signaling pathway | GO:0030522 | GO:BP | 8.82E-11 |
| regulation of Wnt signaling pathway | GO:0030111 | GO:BP | 9.19E-11 |
| regulation of organelle assembly | GO:1902115 | GO:BP | 9.48E-11 |
| import into nucleus | GO:0051170 | GO:BP | 9.67E-11 |
| neuron projection development | GO:0031175 | GO:BP | 9.71E-11 |
| hemopoiesis | GO:0030097 | GO:BP | 9.83E-11 |
| regulation of microtubule cytoskeleton organization | GO:0070507 | GO:BP | 1.07E-10 |
| protein import into nucleus | GO:0006606 | GO:BP | 1.07E-10 |
| positive regulation of chromosome segregation | GO:0051984 | GO:BP | 1.08E-10 |

| | | | |
|---|---|---|---|
| positive regulation of chromosome organization | GO:2001252 | GO:BP | 1.35E-10 |
| regulation of plasma membrane bounded cell projectio | GO:0120035 | GO:BP | 1.36E-10 |
| positive regulation of DNA-templated transcription initia | GO:2000144 | GO:BP | 1.48E-10 |
| regulation of G2/M transition of mitotic cell cycle | GO:0010389 | GO:BP | 1.60E-10 |
| cell cycle DNA replication | GO:0044786 | GO:BP | 1.75E-10 |
| central nervous system development | GO:0007417 | GO:BP | 1.84E-10 |
| response to growth factor | GO:0070848 | GO:BP | 1.91E-10 |
| regulation of phosphate metabolic process | GO:0019220 | GO:BP | 2.15E-10 |
| transcription by RNA polymerase III | GO:0006383 | GO:BP | 2.30E-10 |
| mitotic sister chromatid separation | GO:0051306 | GO:BP | 2.30E-10 |
| positive regulation of transcription initiation by RNA pol | GO:0060261 | GO:BP | 2.30E-10 |
| regulation of DNA-templated transcription initiation | GO:2000142 | GO:BP | 2.39E-10 |
| regulation of phosphorus metabolic process | GO:0051174 | GO:BP | 2.44E-10 |
| positive regulation of protein modification by small prot | GO:1903322 | GO:BP | 2.70E-10 |
| response to organic substance | GO:0010033 | GO:BP | 2.87E-10 |
| positive regulation of GTPase activity | GO:0043547 | GO:BP | 3.06E-10 |
| negative regulation of intracellular signal transduction | GO:1902532 | GO:BP | 3.28E-10 |
| proteolysis | GO:0006508 | GO:BP | 3.28E-10 |
| snRNA metabolic process | GO:0016073 | GO:BP | 3.32E-10 |
| protein dephosphorylation | GO:0006470 | GO:BP | 3.97E-10 |
| maturation of SSU-rRNA | GO:0030490 | GO:BP | 4.03E-10 |
| neuron development | GO:0048666 | GO:BP | 4.12E-10 |
| positive regulation of post-translational protein modific | GO:1901875 | GO:BP | 4.22E-10 |
| positive regulation of response to stimulus | GO:0048584 | GO:BP | 4.64E-10 |
| regulation of transcription initiation by RNA polymerase | GO:0060260 | GO:BP | 5.19E-10 |
| regulation of microtubule-based process | GO:0032886 | GO:BP | 5.21E-10 |
| mitotic metaphase chromosome alignment | GO:0007080 | GO:BP | 5.80E-10 |
| viral gene expression | GO:0019080 | GO:BP | 5.92E-10 |
| vesicle organization | GO:0016050 | GO:BP | 6.13E-10 |
| regulation of mitotic sister chromatid separation | GO:0010965 | GO:BP | 6.67E-10 |
| cellular response to growth factor stimulus | GO:0071363 | GO:BP | 7.63E-10 |
| regulation of nucleocytoplasmic transport | GO:0046822 | GO:BP | 7.95E-10 |
| positive regulation of protein localization | GO:1903829 | GO:BP | 8.53E-10 |
| regulation of protein stability | GO:0031647 | GO:BP | 8.86E-10 |
| actin filament-based process | GO:0030029 | GO:BP | 1.02E-09 |
| protein-DNA complex assembly | GO:0065004 | GO:BP | 1.09E-09 |
| regulation of G1/S transition of mitotic cell cycle | GO:2000045 | GO:BP | 1.15E-09 |
| positive regulation of signal transduction | GO:0009967 | GO:BP | 1.19E-09 |
| enzyme-linked receptor protein signaling pathway | GO:0007167 | GO:BP | 1.32E-09 |
| brain development | GO:0007420 | GO:BP | 1.41E-09 |
| cellular response to insulin stimulus | GO:0032869 | GO:BP | 1.60E-09 |
| regulation of anatomical structure morphogenesis | GO:0022603 | GO:BP | 1.79E-09 |
| regulation of phosphorylation | GO:0042325 | GO:BP | 1.82E-09 |
| retrograde transport, endosome to Golgi | GO:0042147 | GO:BP | 2.09E-09 |
| cellular response to organic substance | GO:0071310 | GO:BP | 2.38E-09 |
| mitochondrial RNA metabolic process | GO:0000959 | GO:BP | 2.54E-09 |
| canonical Wnt signaling pathway | GO:0060070 | GO:BP | 2.86E-09 |
| regulation of cell differentiation | GO:0045595 | GO:BP | 2.86E-09 |

| | | | |
|---|---|---|---|
| regulation of protein ubiquitination | GO:0031396 | GO:BP | 2.90E-09 |
| negative regulation of mitotic sister chromatid segregat | GO:0033048 | GO:BP | 3.09E-09 |
| negative regulation of mitotic metaphase/anaphase trai | GO:0045841 | GO:BP | 3.09E-09 |
| negative regulation of sister chromatid segregation | GO:0033046 | GO:BP | 3.09E-09 |
| negative regulation of mitotic sister chromatid separatic | GO:2000816 | GO:BP | 3.09E-09 |
| regulation of transcription by RNA polymerase I | GO:0006356 | GO:BP | 3.09E-09 |
| regulation of supramolecular fiber organization | GO:1902903 | GO:BP | 3.10E-09 |
| positive regulation of mitotic cell cycle phase transition | GO:1901992 | GO:BP | 3.11E-09 |
| establishment of vesicle localization | GO:0051650 | GO:BP | 3.29E-09 |
| actin filament organization | GO:0007015 | GO:BP | 3.43E-09 |
| miRNA processing | GO:0035196 | GO:BP | 3.55E-09 |
| TOR signaling | GO:0031929 | GO:BP | 3.97E-09 |
| protein localization to microtubule cytoskeleton | GO:0072698 | GO:BP | 4.94E-09 |
| snRNA processing | GO:0016180 | GO:BP | 5.55E-09 |
| regulation of protein-containing complex assembly | GO:0043254 | GO:BP | 6.75E-09 |
| snRNA 3'-end processing | GO:0034472 | GO:BP | 7.30E-09 |
| protein acylation | GO:0043543 | GO:BP | 7.30E-09 |
| cell migration | GO:0016477 | GO:BP | 7.30E-09 |
| regulation of autophagy | GO:0010506 | GO:BP | 8.06E-09 |
| miRNA metabolic process | GO:0010586 | GO:BP | 9.48E-09 |
| microtubule organizing center localization | GO:0061842 | GO:BP | 9.81E-09 |
| centrosome localization | GO:0051642 | GO:BP | 9.81E-09 |
| nuclear DNA replication | GO:0033260 | GO:BP | 1.03E-08 |
| DNA strand elongation | GO:0022616 | GO:BP | 1.07E-08 |
| regulation of protein phosphorylation | GO:0001932 | GO:BP | 1.10E-08 |
| mitotic chromosome condensation | GO:0007076 | GO:BP | 1.16E-08 |
| protein polymerization | GO:0051258 | GO:BP | 1.17E-08 |
| regulation of protein catabolic process | GO:0042176 | GO:BP | 1.24E-08 |
| positive regulation of cytoskeleton organization | GO:0051495 | GO:BP | 1.25E-08 |
| glycoprotein biosynthetic process | GO:0009101 | GO:BP | 1.26E-08 |
| heterochromatin formation | GO:0031507 | GO:BP | 1.43E-08 |
| positive regulation of cell cycle phase transition | GO:1901989 | GO:BP | 1.43E-08 |
| vesicle localization | GO:0051648 | GO:BP | 1.48E-08 |
| regulation of canonical Wnt signaling pathway | GO:0060828 | GO:BP | 1.51E-08 |
| regulation of ncRNA transcription | GO:0140747 | GO:BP | 1.57E-08 |
| negative regulation of chromosome separation | GO:1905819 | GO:BP | 1.65E-08 |
| negative regulation of metaphase/anaphase transition ( | GO:1902100 | GO:BP | 1.65E-08 |
| negative regulation of chromosome segregation | GO:0051985 | GO:BP | 1.65E-08 |
| negative regulation of protein modification process | GO:0031400 | GO:BP | 1.68E-08 |
| protein localization to cytoskeleton | GO:0044380 | GO:BP | 1.81E-08 |
| mitotic spindle assembly | GO:0090307 | GO:BP | 1.93E-08 |
| telomere organization | GO:0032200 | GO:BP | 1.96E-08 |
| cellular response to peptide hormone stimulus | GO:0071375 | GO:BP | 1.99E-08 |
| regulation of mitotic nuclear division | GO:0007088 | GO:BP | 2.04E-08 |
| mitotic spindle assembly checkpoint signaling | GO:0007094 | GO:BP | 2.09E-08 |
| mitotic spindle checkpoint signaling | GO:0071174 | GO:BP | 2.09E-08 |
| spindle assembly checkpoint signaling | GO:0071173 | GO:BP | 2.09E-08 |
| cell projection morphogenesis | GO:0048858 | GO:BP | 2.09E-08 |

| | | | |
|---|---|---|---|
| response to insulin | GO:0032868 | GO:BP | 2.14E-08 |
| regulation of multicellular organismal development | GO:2000026 | GO:BP | 2.19E-08 |
| hippo signaling | GO:0035329 | GO:BP | 2.32E-08 |
| regulation of binding | GO:0051098 | GO:BP | 2.36E-08 |
| negative regulation of catabolic process | GO:0009895 | GO:BP | 2.46E-08 |
| plasma membrane bounded cell projection morphogen | GO:0120039 | GO:BP | 2.61E-08 |
| peptidyl-lysine modification | GO:0018205 | GO:BP | 2.88E-08 |
| cell population proliferation | GO:0008283 | GO:BP | 2.91E-08 |
| maturation of 5.8S rRNA | GO:0000460 | GO:BP | 2.93E-08 |
| positive regulation of double-strand break repair | GO:2000781 | GO:BP | 3.16E-08 |
| regulation of cell cycle G1/S phase transition | GO:1902806 | GO:BP | 3.22E-08 |
| microtubule polymerization or depolymerization | GO:0031109 | GO:BP | 3.28E-08 |
| centrosome duplication | GO:0051298 | GO:BP | 3.41E-08 |
| regulation of double-strand break repair via homologous | GO:0010569 | GO:BP | 3.41E-08 |
| regulation of intracellular protein transport | GO:0033157 | GO:BP | 3.41E-08 |
| mitochondrial translation | GO:0032543 | GO:BP | 3.72E-08 |
| cellular response to hormone stimulus | GO:0032870 | GO:BP | 3.78E-08 |
| dephosphorylation | GO:0016311 | GO:BP | 3.95E-08 |
| regulation of protein kinase activity | GO:0045859 | GO:BP | 4.05E-08 |
| negative regulation of programmed cell death | GO:0043069 | GO:BP | 4.05E-08 |
| mitochondrial transport | GO:0006839 | GO:BP | 4.11E-08 |
| alternative mRNA splicing, via spliceosome | GO:0000380 | GO:BP | 4.24E-08 |
| phospholipid biosynthetic process | GO:0008654 | GO:BP | 4.26E-08 |
| regulation of kinase activity | GO:0043549 | GO:BP | 4.57E-08 |
| spindle checkpoint signaling | GO:0031577 | GO:BP | 4.66E-08 |
| meiotic cell cycle | GO:0051321 | GO:BP | 4.91E-08 |
| negative regulation of mitotic nuclear division | GO:0045839 | GO:BP | 5.11E-08 |
| regulation of cytokinesis | GO:0032465 | GO:BP | 5.23E-08 |
| regulation of telomere maintenance | GO:0032204 | GO:BP | 6.01E-08 |
| DNA modification | GO:0006304 | GO:BP | 6.37E-08 |
| focal adhesion assembly | GO:0048041 | GO:BP | 6.69E-08 |
| positive regulation of DNA-templated transcription, elon | GO:0032786 | GO:BP | 6.77E-08 |
| heterochromatin organization | GO:0070828 | GO:BP | 6.98E-08 |
| DNA replication initiation | GO:0006270 | GO:BP | 7.72E-08 |
| regulation of proteolysis involved in protein catabolic pr | GO:1903050 | GO:BP | 7.94E-08 |
| transcription preinitiation complex assembly | GO:0070897 | GO:BP | 8.54E-08 |
| positive regulation of protein ubiquitination | GO:0031398 | GO:BP | 8.58E-08 |
| post-Golgi vesicle-mediated transport | GO:0006892 | GO:BP | 9.35E-08 |
| vacuolar transport | GO:0007034 | GO:BP | 1.00E-07 |
| sister chromatid cohesion | GO:0007062 | GO:BP | 1.01E-07 |
| regulation of cell cycle checkpoint | GO:1901976 | GO:BP | 1.02E-07 |
| carbohydrate derivative biosynthetic process | GO:1901137 | GO:BP | 1.06E-07 |
| cell surface receptor signaling pathway | GO:0007166 | GO:BP | 1.15E-07 |
| nuclear-transcribed mRNA catabolic process, deadenyl | GO:0000288 | GO:BP | 1.19E-07 |
| regulation of attachment of spindle microtubules to kin | GO:0051988 | GO:BP | 1.21E-07 |
| positive regulation of signaling | GO:0023056 | GO:BP | 1.21E-07 |
| regulation of TOR signaling | GO:0032006 | GO:BP | 1.36E-07 |
| nucleotide-excision repair | GO:0006289 | GO:BP | 1.42E-07 |

| | | | |
|---|---|---|---|
| positive regulation of intracellular transport | GO:0032388 | GO:BP | 1.45E-07 |
| response to ionizing radiation | GO:0010212 | GO:BP | 1.45E-07 |
| Golgi organization | GO:0007030 | GO:BP | 1.61E-07 |
| regulation of stem cell population maintenance | GO:2000036 | GO:BP | 1.66E-07 |
| glycosylation | GO:0070085 | GO:BP | 1.78E-07 |
| neuron projection morphogenesis | GO:0048812 | GO:BP | 1.85E-07 |
| autophagosome assembly | GO:0000045 | GO:BP | 1.97E-07 |
| positive regulation of cell projection organization | GO:0031346 | GO:BP | 2.03E-07 |
| regulation of growth | GO:0040008 | GO:BP | 2.05E-07 |
| response to organonitrogen compound | GO:0010243 | GO:BP | 2.16E-07 |
| mRNA modification | GO:0016556 | GO:BP | 2.20E-07 |
| ribosomal large subunit biogenesis | GO:0042273 | GO:BP | 2.24E-07 |
| regulation of establishment of protein localization | GO:0070201 | GO:BP | 2.32E-07 |
| negative regulation of apoptotic process | GO:0043066 | GO:BP | 2.32E-07 |
| negative regulation of nuclear division | GO:0051784 | GO:BP | 2.33E-07 |
| negative regulation of cell cycle G2/M phase transition | GO:1902750 | GO:BP | 2.64E-07 |
| regulation of cell-substrate junction organization | GO:0150116 | GO:BP | 2.64E-07 |
| peptidyl-threonine modification | GO:0018210 | GO:BP | 2.77E-07 |
| regulation of centrosome cycle | GO:0046605 | GO:BP | 2.90E-07 |
| animal organ development | GO:0048513 | GO:BP | 2.98E-07 |
| endoplasmic reticulum to Golgi vesicle-mediated trans | GO:0006888 | GO:BP | 3.14E-07 |
| vesicle targeting | GO:0006903 | GO:BP | 3.33E-07 |
| regulation of DNA biosynthetic process | GO:2000278 | GO:BP | 3.34E-07 |
| regulation of hydrolase activity | GO:0051336 | GO:BP | 3.42E-07 |
| establishment of mitotic spindle localization | GO:0040001 | GO:BP | 3.42E-07 |
| protein localization to cell periphery | GO:1990778 | GO:BP | 3.67E-07 |
| regulation of neuron projection development | GO:0010975 | GO:BP | 3.67E-07 |
| insulin receptor signaling pathway | GO:0008286 | GO:BP | 3.75E-07 |
| autophagosome organization | GO:1905037 | GO:BP | 3.75E-07 |
| response to nitrogen compound | GO:1901698 | GO:BP | 3.75E-07 |
| mitochondrial RNA processing | GO:0000963 | GO:BP | 3.77E-07 |
| regulation of cell morphogenesis | GO:0022604 | GO:BP | 3.78E-07 |
| cell junction organization | GO:0034330 | GO:BP | 3.92E-07 |
| cell growth | GO:0016049 | GO:BP | 4.09E-07 |
| protein K63-linked ubiquitination | GO:0070534 | GO:BP | 4.13E-07 |
| regulation of DNA recombination | GO:0000018 | GO:BP | 4.43E-07 |
| canonical NF-kappaB signal transduction | GO:0007249 | GO:BP | 4.64E-07 |
| regulation of protein localization to nucleus | GO:1900180 | GO:BP | 4.78E-07 |
| rRNA transcription | GO:0009303 | GO:BP | 4.98E-07 |
| cellular response to nitrogen compound | GO:1901699 | GO:BP | 5.04E-07 |
| regulation of biological quality | GO:0065008 | GO:BP | 5.11E-07 |
| positive regulation of transcription by RNA polymerase I | GO:0045943 | GO:BP | 5.46E-07 |
| maturation of SSU-rRNA from tricistronic rRNA transcrip | GO:0000462 | GO:BP | 5.46E-07 |
| macromolecule glycosylation | GO:0043413 | GO:BP | 5.62E-07 |
| protein glycosylation | GO:0006486 | GO:BP | 5.62E-07 |
| tube development | GO:0035295 | GO:BP | 5.64E-07 |
| vesicle budding from membrane | GO:0006900 | GO:BP | 6.06E-07 |
| N-terminal protein amino acid modification | GO:0031365 | GO:BP | 6.07E-07 |

| | | | |
|---|---|---|---|
| mitotic sister chromatid cohesion | GO:0007064 | GO:BP | 6.14E-07 |
| positive regulation of transferase activity | GO:0051347 | GO:BP | 6.36E-07 |
| positive regulation of cell communication | GO:0010647 | GO:BP | 6.55E-07 |
| microtubule polymerization | GO:0046785 | GO:BP | 6.55E-07 |
| ncRNA catabolic process | GO:0034661 | GO:BP | 6.65E-07 |
| cell-substrate junction organization | GO:0150115 | GO:BP | 6.98E-07 |
| regulation of miRNA metabolic process | GO:2000628 | GO:BP | 7.05E-07 |
| spindle localization | GO:0051653 | GO:BP | 7.17E-07 |
| positive regulation of Wnt signaling pathway | GO:0030177 | GO:BP | 7.30E-07 |
| response to hormone | GO:0009725 | GO:BP | 7.33E-07 |
| cellular response to organic cyclic compound | GO:0071407 | GO:BP | 7.51E-07 |
| regulation of actin filament organization | GO:0110053 | GO:BP | 7.73E-07 |
| positive regulation of supramolecular fiber organization | GO:1902905 | GO:BP | 7.84E-07 |
| carbohydrate derivative metabolic process | GO:1901135 | GO:BP | 7.99E-07 |
| mRNA methylation | GO:0080009 | GO:BP | 8.05E-07 |
| cellular anatomical entity morphogenesis | GO:0032989 | GO:BP | 8.38E-07 |
| regulation of nuclear division | GO:0051783 | GO:BP | 8.38E-07 |
| supramolecular fiber organization | GO:0097435 | GO:BP | 8.67E-07 |
| regulation of multicellular organismal process | GO:0051239 | GO:BP | 8.67E-07 |
| regulation of focal adhesion assembly | GO:0051893 | GO:BP | 8.96E-07 |
| regulation of cell-substrate junction assembly | GO:0090109 | GO:BP | 8.96E-07 |
| protein autophosphorylation | GO:0046777 | GO:BP | 9.02E-07 |
| negative regulation of phosphorylation | GO:0042326 | GO:BP | 9.25E-07 |
| peptidyl-threonine phosphorylation | GO:0018107 | GO:BP | 9.48E-07 |
| replication fork processing | GO:0031297 | GO:BP | 9.80E-07 |
| nuclear envelope organization | GO:0006998 | GO:BP | 9.82E-07 |
| DNA-templated DNA replication maintenance of fidelity | GO:0045005 | GO:BP | 9.82E-07 |
| cell-substrate junction assembly | GO:0007044 | GO:BP | 9.82E-07 |
| negative regulation of phosphorus metabolic process | GO:0010563 | GO:BP | 9.88E-07 |
| regulation of ubiquitin-dependent protein catabolic pro | GO:2000058 | GO:BP | 9.89E-07 |
| erythrocyte homeostasis | GO:0034101 | GO:BP | 9.94E-07 |
| cellular response to external stimulus | GO:0071496 | GO:BP | 1.09E-06 |
| response to peptide hormone | GO:0043434 | GO:BP | 1.14E-06 |
| negative regulation of molecular function | GO:0044092 | GO:BP | 1.20E-06 |
| regulation of chromatin organization | GO:1902275 | GO:BP | 1.25E-06 |
| apoptotic signaling pathway | GO:0097190 | GO:BP | 1.30E-06 |
| circulatory system development | GO:0072359 | GO:BP | 1.31E-06 |
| endoplasmic reticulum organization | GO:0007029 | GO:BP | 1.32E-06 |
| cellular response to peptide | GO:1901653 | GO:BP | 1.38E-06 |
| protein localization to plasma membrane | GO:0072659 | GO:BP | 1.39E-06 |
| negative regulation of phosphate metabolic process | GO:0045936 | GO:BP | 1.47E-06 |
| centrosome separation | GO:0051299 | GO:BP | 1.47E-06 |
| DNA strand elongation involved in DNA replication | GO:0006271 | GO:BP | 1.47E-06 |
| regulation of proteasomal protein catabolic process | GO:0061136 | GO:BP | 1.59E-06 |
| attachment of mitotic spindle microtubules to kinetoch | GO:0051315 | GO:BP | 1.60E-06 |
| protein K11-linked ubiquitination | GO:0070979 | GO:BP | 1.63E-06 |
| regulation of actin cytoskeleton organization | GO:0032956 | GO:BP | 1.70E-06 |
| transmembrane receptor protein tyrosine kinase signali | GO:0007169 | GO:BP | 1.82E-06 |

| | | | |
|---|---|---|---|
| blastocyst development | GO:0001824 | GO:BP | 1.82E-06 |
| tube morphogenesis | GO:0035239 | GO:BP | 1.83E-06 |
| peptidyl-serine modification | GO:0018209 | GO:BP | 1.84E-06 |
| negative regulation of protein phosphorylation | GO:0001933 | GO:BP | 1.84E-06 |
| negative regulation of cellular catabolic process | GO:0031330 | GO:BP | 1.84E-06 |
| signal transduction by p53 class mediator | GO:0072331 | GO:BP | 1.85E-06 |
| cellular response to organonitrogen compound | GO:0071417 | GO:BP | 1.95E-06 |
| erythrocyte differentiation | GO:0030218 | GO:BP | 1.97E-06 |
| double-strand break repair via nonhomologous end join | GO:0006303 | GO:BP | 2.02E-06 |
| regulation of cellular component size | GO:0032535 | GO:BP | 2.20E-06 |
| glycerophospholipid biosynthetic process | GO:0046474 | GO:BP | 2.24E-06 |
| positive regulation of stem cell population maintenance | GO:1902459 | GO:BP | 2.26E-06 |
| mRNA transcription by RNA polymerase II | GO:0042789 | GO:BP | 2.26E-06 |
| cytoplasmic pattern recognition receptor signaling path | GO:0002753 | GO:BP | 2.33E-06 |
| vacuole organization | GO:0007033 | GO:BP | 2.37E-06 |
| protein autoubiquitination | GO:0051865 | GO:BP | 2.44E-06 |
| regulation of protein transport | GO:0051223 | GO:BP | 2.45E-06 |
| negative regulation of mRNA metabolic process | GO:1903312 | GO:BP | 2.50E-06 |
| negative regulation of G2/M transition of mitotic cell cyc | GO:0010972 | GO:BP | 2.56E-06 |
| positive regulation of viral process | GO:0048524 | GO:BP | 2.56E-06 |
| heart development | GO:0007507 | GO:BP | 2.59E-06 |
| viral transcription | GO:0019083 | GO:BP | 2.72E-06 |
| mRNA transcription | GO:0009299 | GO:BP | 2.72E-06 |
| vesicle-mediated transport to the plasma membrane | GO:0098876 | GO:BP | 2.83E-06 |
| protein localization to microtubule organizing center | GO:1905508 | GO:BP | 3.17E-06 |
| postreplication repair | GO:0006301 | GO:BP | 3.17E-06 |
| protein sumoylation | GO:0016925 | GO:BP | 3.33E-06 |
| chromosome condensation | GO:0030261 | GO:BP | 3.35E-06 |
| cytokinetic process | GO:0032506 | GO:BP | 3.35E-06 |
| plasma membrane bounded cell projection assembly | GO:0120031 | GO:BP | 3.60E-06 |
| peptidyl-serine phosphorylation | GO:0018105 | GO:BP | 3.65E-06 |
| kinetochore organization | GO:0051383 | GO:BP | 4.19E-06 |
| nucleolar large rRNA transcription by RNA polymerase I | GO:0042790 | GO:BP | 4.19E-06 |
| negative regulation of protein localization | GO:1903828 | GO:BP | 4.67E-06 |
| response to radiation | GO:0009314 | GO:BP | 4.67E-06 |
| maturation of 5.8S rRNA from tricistronic rRNA transcrip | GO:0000466 | GO:BP | 4.72E-06 |
| interstrand cross-link repair | GO:0036297 | GO:BP | 4.91E-06 |
| mitotic centrosome separation | GO:0007100 | GO:BP | 4.93E-06 |
| positive regulation of DNA-templated DNA replication | GO:2000105 | GO:BP | 4.93E-06 |
| regulation of centrosome duplication | GO:0010824 | GO:BP | 5.26E-06 |
| regulation of protein-containing complex disassembly | GO:0043244 | GO:BP | 5.47E-06 |
| cell projection assembly | GO:0030031 | GO:BP | 5.54E-06 |
| pallium development | GO:0021543 | GO:BP | 5.62E-06 |
| negative regulation of DNA metabolic process | GO:0051053 | GO:BP | 5.74E-06 |
| positive regulation of DNA replication | GO:0045740 | GO:BP | 5.83E-06 |
| negative regulation of translation | GO:0017148 | GO:BP | 5.84E-06 |
| regulation of actin filament-based process | GO:0032970 | GO:BP | 6.61E-06 |
| regulation of chromosome condensation | GO:0060623 | GO:BP | 6.74E-06 |

| | | | |
|---|---|---|---|
| protein acetylation | GO:0006473 | GO:BP | 6.74E-06 |
| regulation of protein serine/threonine kinase activity | GO:0071900 | GO:BP | 6.83E-06 |
| myeloid cell differentiation | GO:0030099 | GO:BP | 7.03E-06 |
| positive regulation of canonical NF-kappaB signal trans | GO:0043123 | GO:BP | 7.44E-06 |
| regulation of stem cell differentiation | GO:2000736 | GO:BP | 7.58E-06 |
| positive regulation of transcription elongation by RNA p | GO:0032968 | GO:BP | 7.72E-06 |
| positive regulation of amide metabolic process | GO:0034250 | GO:BP | 7.77E-06 |
| centriole replication | GO:0007099 | GO:BP | 7.92E-06 |
| tRNA methylation | GO:0030488 | GO:BP | 7.92E-06 |
| protein localization to vacuole | GO:0072665 | GO:BP | 7.94E-06 |
| protein localization to centrosome | GO:0071539 | GO:BP | 7.99E-06 |
| regulation of canonical NF-kappaB signal transduction | GO:0043122 | GO:BP | 7.99E-06 |
| endocytosis | GO:0006897 | GO:BP | 8.27E-06 |
| positive regulation of DNA recombination | GO:0045911 | GO:BP | 8.63E-06 |
| Golgi to plasma membrane transport | GO:0006893 | GO:BP | 8.86E-06 |
| establishment of spindle localization | GO:0051293 | GO:BP | 8.86E-06 |
| glycerolipid biosynthetic process | GO:0045017 | GO:BP | 8.91E-06 |
| mitotic DNA integrity checkpoint signaling | GO:0044774 | GO:BP | 9.05E-06 |
| hematopoietic progenitor cell differentiation | GO:0002244 | GO:BP | 9.41E-06 |
| centriole assembly | GO:0098534 | GO:BP | 9.56E-06 |
| glycoprotein metabolic process | GO:0009100 | GO:BP | 9.71E-06 |
| positive regulation of cell differentiation | GO:0045597 | GO:BP | 1.02E-05 |
| positive regulation of chromosome separation | GO:1905820 | GO:BP | 1.04E-05 |
| mitotic G2/M transition checkpoint | GO:0044818 | GO:BP | 1.07E-05 |
| response to abiotic stimulus | GO:0009628 | GO:BP | 1.08E-05 |
| organophosphate biosynthetic process | GO:0090407 | GO:BP | 1.10E-05 |
| regulation of cell growth | GO:0001558 | GO:BP | 1.12E-05 |
| centromere complex assembly | GO:0034508 | GO:BP | 1.15E-05 |
| telencephalon development | GO:0021537 | GO:BP | 1.16E-05 |
| protein stabilization | GO:0050821 | GO:BP | 1.17E-05 |
| DNA alkylation | GO:0006305 | GO:BP | 1.19E-05 |
| DNA methylation | GO:0006306 | GO:BP | 1.19E-05 |
| regulation of cell development | GO:0060284 | GO:BP | 1.25E-05 |
| miRNA transcription | GO:0061614 | GO:BP | 1.25E-05 |
| ameboidal-type cell migration | GO:0001667 | GO:BP | 1.26E-05 |
| viral life cycle | GO:0019058 | GO:BP | 1.28E-05 |
| positive regulation of organelle assembly | GO:1902117 | GO:BP | 1.32E-05 |
| organophosphate metabolic process | GO:0019637 | GO:BP | 1.34E-05 |
| myeloid cell homeostasis | GO:0002262 | GO:BP | 1.37E-05 |
| vasculature development | GO:0001944 | GO:BP | 1.37E-05 |
| establishment of mitotic spindle orientation | GO:0000132 | GO:BP | 1.39E-05 |
| response to UV | GO:0009411 | GO:BP | 1.46E-05 |
| negative regulation of amide metabolic process | GO:0034249 | GO:BP | 1.46E-05 |
| regulation of cell projection assembly | GO:0060491 | GO:BP | 1.47E-05 |
| protein depolymerization | GO:0051261 | GO:BP | 1.48E-05 |
| regulation of metaphase plate congression | GO:0090235 | GO:BP | 1.63E-05 |
| negative regulation of double-strand break repair | GO:2000780 | GO:BP | 1.66E-05 |
| positive regulation of translation | GO:0045727 | GO:BP | 1.67E-05 |

| | | | |
|---|---|---|---|
| forebrain development | GO:0030900 | GO:BP | 1.67E-05 |
| dendrite development | GO:0016358 | GO:BP | 1.73E-05 |
| cellular response to chemical stimulus | GO:0070887 | GO:BP | 1.73E-05 |
| regulation of cell migration | GO:0030334 | GO:BP | 1.74E-05 |
| regulation of plasma membrane bounded cell projectio | GO:0120032 | GO:BP | 1.84E-05 |
| actin filament bundle assembly | GO:0051017 | GO:BP | 2.02E-05 |
| actin filament bundle organization | GO:0061572 | GO:BP | 2.06E-05 |
| stem cell differentiation | GO:0048863 | GO:BP | 2.08E-05 |
| positive regulation of cell growth | GO:0030307 | GO:BP | 2.10E-05 |
| protein targeting to vacuole | GO:0006623 | GO:BP | 2.12E-05 |
| response to transforming growth factor beta | GO:0071559 | GO:BP | 2.28E-05 |
| ERBB signaling pathway | GO:0038127 | GO:BP | 2.29E-05 |
| regulation of transcription by RNA polymerase III | GO:0006359 | GO:BP | 2.30E-05 |
| positive regulation of growth | GO:0045927 | GO:BP | 2.40E-05 |
| RNA stabilization | GO:0043489 | GO:BP | 2.45E-05 |
| regulation of miRNA transcription | GO:1902893 | GO:BP | 2.45E-05 |
| protein localization to condensed chromosome | GO:1903083 | GO:BP | 2.52E-05 |
| regulation of DNA-templated DNA replication initiation | GO:0030174 | GO:BP | 2.52E-05 |
| protein localization to kinetochore | GO:0034501 | GO:BP | 2.52E-05 |
| organelle fusion | GO:0048284 | GO:BP | 2.56E-05 |
| cellular response to extracellular stimulus | GO:0031668 | GO:BP | 2.64E-05 |
| microtubule nucleation | GO:0007020 | GO:BP | 2.65E-05 |
| RNA polymerase II preinitiation complex assembly | GO:0051123 | GO:BP | 2.67E-05 |
| positive regulation of canonical Wnt signaling pathway | GO:0090263 | GO:BP | 2.74E-05 |
| establishment of protein localization to vacuole | GO:0072666 | GO:BP | 2.86E-05 |
| mitochondrial transmembrane transport | GO:1990542 | GO:BP | 3.00E-05 |
| regulation of alternative mRNA splicing, via spliceosom | GO:0000381 | GO:BP | 3.02E-05 |
| RNA surveillance | GO:0071025 | GO:BP | 3.16E-05 |
| nucleolus organization | GO:0007000 | GO:BP | 3.16E-05 |
| positive regulation of intracellular signal transduction | GO:1902533 | GO:BP | 3.20E-05 |
| viral genome replication | GO:0019079 | GO:BP | 3.22E-05 |
| regulation of DNA damage checkpoint | GO:2000001 | GO:BP | 3.28E-05 |
| activation of GTPase activity | GO:0090630 | GO:BP | 3.31E-05 |
| membrane fission | GO:0090148 | GO:BP | 3.31E-05 |
| negative regulation of catalytic activity | GO:0043086 | GO:BP | 3.45E-05 |
| response to endoplasmic reticulum stress | GO:0034976 | GO:BP | 3.47E-05 |
| regulation of centriole replication | GO:0046599 | GO:BP | 3.50E-05 |
| positive regulation of cell cycle checkpoint | GO:1901978 | GO:BP | 3.50E-05 |
| roof of mouth development | GO:0060021 | GO:BP | 3.50E-05 |
| protein export from nucleus | GO:0006611 | GO:BP | 3.67E-05 |
| mRNA 3'-end processing | GO:0031124 | GO:BP | 3.67E-05 |
| regulation of microtubule polymerization or depolymeri | GO:0031110 | GO:BP | 3.85E-05 |
| embryonic morphogenesis | GO:0048598 | GO:BP | 3.89E-05 |
| response to peptide | GO:1901652 | GO:BP | 3.93E-05 |
| mitotic DNA damage checkpoint signaling | GO:0044773 | GO:BP | 4.09E-05 |
| TORC1 signaling | GO:0038202 | GO:BP | 4.20E-05 |
| positive regulation of gene expression | GO:0010628 | GO:BP | 4.37E-05 |
| regulation of proteolysis | GO:0030162 | GO:BP | 4.52E-05 |

| | | | |
|---|---|---|---|
| lamellipodium organization | GO:0097581 | GO:BP | 4.54E-05 |
| actin polymerization or depolymerization | GO:0008154 | GO:BP | 4.54E-05 |
| regulation of cell division | GO:0051302 | GO:BP | 4.56E-05 |
| import into the mitochondrion | GO:0170036 | GO:BP | 4.62E-05 |
| blastocyst formation | GO:0001825 | GO:BP | 4.62E-05 |
| base-excision repair | GO:0006284 | GO:BP | 4.62E-05 |
| facultative heterochromatin formation | GO:0140718 | GO:BP | 4.62E-05 |
| striated muscle cell proliferation | GO:0014855 | GO:BP | 4.69E-05 |
| exit from mitosis | GO:0010458 | GO:BP | 4.74E-05 |
| steroid hormone mediated signaling pathway | GO:0043401 | GO:BP | 4.85E-05 |
| protein deacylation | GO:0035601 | GO:BP | 4.85E-05 |
| cortical cytoskeleton organization | GO:0030865 | GO:BP | 4.85E-05 |
| embryonic skeletal system morphogenesis | GO:0048704 | GO:BP | 4.93E-05 |
| protein localization to chromosome | GO:0034502 | GO:BP | 5.00E-05 |
| vesicle coating | GO:0006901 | GO:BP | 5.05E-05 |
| positive regulation of gene expression, epigenetic | GO:0141137 | GO:BP | 5.05E-05 |
| anterior/posterior pattern specification | GO:0009952 | GO:BP | 5.13E-05 |
| intracellular steroid hormone receptor signaling pathwa | GO:0030518 | GO:BP | 5.21E-05 |
| somatic diversification of immune receptors | GO:0002200 | GO:BP | 5.34E-05 |
| positive regulation of protein localization to nucleus | GO:1900182 | GO:BP | 5.35E-05 |
| regulation of macroautophagy | GO:0016241 | GO:BP | 5.40E-05 |
| cerebral cortex development | GO:0021987 | GO:BP | 5.62E-05 |
| negative regulation of RNA catabolic process | GO:1902369 | GO:BP | 5.78E-05 |
| establishment of spindle orientation | GO:0051294 | GO:BP | 5.78E-05 |
| organelle transport along microtubule | GO:0072384 | GO:BP | 5.78E-05 |
| mitochondrial membrane organization | GO:0007006 | GO:BP | 5.78E-05 |
| regulation of proteasomal ubiquitin-dependent protein | GO:0032434 | GO:BP | 5.96E-05 |
| regulation of protein localization to cell periphery | GO:1904375 | GO:BP | 6.13E-05 |
| late endosome to vacuole transport | GO:0045324 | GO:BP | 6.20E-05 |
| transcription initiation-coupled chromatin remodeling | GO:0045815 | GO:BP | 6.20E-05 |
| negative regulation of DNA repair | GO:0045738 | GO:BP | 6.20E-05 |
| positive regulation of double-strand break repair via hor | GO:1905168 | GO:BP | 6.20E-05 |
| epithelial cell migration | GO:0010631 | GO:BP | 6.22E-05 |
| cellular response to nutrient levels | GO:0031669 | GO:BP | 6.88E-05 |
| embryonic skeletal system development | GO:0048706 | GO:BP | 7.17E-05 |
| early endosome to late endosome transport | GO:0045022 | GO:BP | 7.26E-05 |
| cellular response to epidermal growth factor stimulus | GO:0071364 | GO:BP | 7.26E-05 |
| ribonucleoside monophosphate metabolic process | GO:0009161 | GO:BP | 7.26E-05 |
| cellular response to transforming growth factor beta sti | GO:0071560 | GO:BP | 7.46E-05 |
| homeostasis of number of cells | GO:0048872 | GO:BP | 7.54E-05 |
| rRNA modification | GO:0000154 | GO:BP | 7.63E-05 |
| mitochondrial tRNA processing | GO:0090646 | GO:BP | 7.65E-05 |
| nuclear RNA surveillance | GO:0071027 | GO:BP | 7.65E-05 |
| regulatory ncRNA processing | GO:0070918 | GO:BP | 7.74E-05 |
| vesicle-mediated transport between endosomal compa | GO:0098927 | GO:BP | 7.78E-05 |
| blood vessel development | GO:0001568 | GO:BP | 7.85E-05 |
| protein monoubiquitination | GO:0006513 | GO:BP | 7.88E-05 |
| cellular response to steroid hormone stimulus | GO:0071383 | GO:BP | 8.05E-05 |

| | | | |
|---|---|---|---|
| 7-methylguanosine RNA capping | GO:0009452 | GO:BP | 8.16E-05 |
| positive regulation of protein catabolic process | GO:0045732 | GO:BP | 8.33E-05 |
| transforming growth factor beta receptor signaling pathw | GO:0007179 | GO:BP | 8.33E-05 |
| cell morphogenesis involved in neuron differentiation | GO:0048667 | GO:BP | 8.38E-05 |
| multicellular organism growth | GO:0035264 | GO:BP | 8.54E-05 |
| positive regulation of miRNA metabolic process | GO:2000630 | GO:BP | 8.54E-05 |
| regulation of cytokine-mediated signaling pathway | GO:0001959 | GO:BP | 8.62E-05 |
| regulation of embryonic development | GO:0045995 | GO:BP | 8.80E-05 |
| morphogenesis of an epithelium | GO:0002009 | GO:BP | 8.87E-05 |
| positive regulation of mRNA processing | GO:0050685 | GO:BP | 9.02E-05 |
| pre-miRNA processing | GO:0031054 | GO:BP | 9.09E-05 |
| tRNA catabolic process | GO:0016078 | GO:BP | 9.09E-05 |
| phosphatidylinositol-3-phosphate biosynthetic process | GO:0036092 | GO:BP | 9.12E-05 |
| regulation of epithelial cell migration | GO:0010632 | GO:BP | 9.17E-05 |
| vesicle targeting, to, from or within Golgi | GO:0048199 | GO:BP | 9.19E-05 |
| autophagosome maturation | GO:0097352 | GO:BP | 9.22E-05 |
| telomere maintenance via telomere lengthening | GO:0010833 | GO:BP | 9.48E-05 |
| transmembrane receptor protein serine/threonine kinas | GO:0007178 | GO:BP | 9.59E-05 |
| meiotic cell cycle process | GO:1903046 | GO:BP | 9.60E-05 |
| tRNA wobble base modification | GO:0002097 | GO:BP | 9.60E-05 |
| regulation of nucleobase-containing compound transpc | GO:0032239 | GO:BP | 9.63E-05 |
| regulation of clathrin-dependent endocytosis | GO:2000369 | GO:BP | 9.63E-05 |
| miRNA-mediated gene silencing by inhibition of translat | GO:0035278 | GO:BP | 9.63E-05 |
| nuclear membrane organization | GO:0071763 | GO:BP | 9.69E-05 |
| negative regulation of organelle assembly | GO:1902116 | GO:BP | 9.69E-05 |
| macromolecule deacylation | GO:0098732 | GO:BP | 9.69E-05 |
| epithelium migration | GO:0090132 | GO:BP | 0.00010076 |
| pattern recognition receptor signaling pathway | GO:0002221 | GO:BP | 0.00010199 |
| anatomical structure formation involved in morphogene | GO:0048646 | GO:BP | 0.0001075 |
| positive regulation of nucleocytoplasmic transport | GO:0046824 | GO:BP | 0.00010894 |
| cell motility | GO:0048870 | GO:BP | 0.00010941 |
| positive regulation of cell development | GO:0010720 | GO:BP | 0.00010967 |
| regulation of cell-matrix adhesion | GO:0001952 | GO:BP | 0.00011002 |
| stress granule assembly | GO:0034063 | GO:BP | 0.00011027 |
| positive regulation of phosphorylation | GO:0042327 | GO:BP | 0.00011299 |
| stress fiber assembly | GO:0043149 | GO:BP | 0.00011935 |
| contractile actin filament bundle assembly | GO:0030038 | GO:BP | 0.00011935 |
| positive regulation of phosphate metabolic process | GO:0045937 | GO:BP | 0.00012107 |
| positive regulation of phosphorus metabolic process | GO:0010562 | GO:BP | 0.00012107 |
| COPII-coated vesicle budding | GO:0090114 | GO:BP | 0.00012281 |
| regulation of spindle organization | GO:0090224 | GO:BP | 0.00012675 |
| positive regulation of protein-containing complex assen | GO:0031334 | GO:BP | 0.00012817 |
| positive regulation of intracellular protein transport | GO:0090316 | GO:BP | 0.00012904 |
| appendage development | GO:0048736 | GO:BP | 0.00013199 |
| limb development | GO:0060173 | GO:BP | 0.00013199 |
| negative regulation of protein localization to cell periphe | GO:1904376 | GO:BP | 0.00013217 |
| response to organic cyclic compound | GO:0014070 | GO:BP | 0.00014082 |
| type I interferon production | GO:0032606 | GO:BP | 0.00014488 |

| | | | |
|---|---|---|---|
| regulation of type I interferon production | GO:0032479 | GO:BP | 0.00014488 |
| cell junction assembly | GO:0034329 | GO:BP | 0.00014796 |
| positive regulation of miRNA transcription | GO:1902895 | GO:BP | 0.00015189 |
| positive regulation of cell cycle G1/S phase transition | GO:1902808 | GO:BP | 0.00015189 |
| translational initiation | GO:0006413 | GO:BP | 0.00015258 |
| positive regulation of G1/S transition of mitotic cell cycl | GO:1900087 | GO:BP | 0.00015523 |
| endosome transport via multivesicular body sorting patl | GO:0032509 | GO:BP | 0.00015523 |
| regulation of anatomical structure size | GO:0090066 | GO:BP | 0.0001563 |
| late endosome to vacuole transport via multivesicular b | GO:0032511 | GO:BP | 0.00015658 |
| negative regulation of protein localization to plasma me | GO:1903077 | GO:BP | 0.00015658 |
| negative regulation of transferase activity | GO:0051348 | GO:BP | 0.00016055 |
| response to epidermal growth factor | GO:0070849 | GO:BP | 0.00016107 |
| positive regulation of protein phosphorylation | GO:0001934 | GO:BP | 0.00016155 |
| RNA metabolic process | GO:0016070 | GO:BP | 0.00016261 |
| positive regulation of attachment of spindle microtubul | GO:0051987 | GO:BP | 0.0001656 |
| regulation of chromatin binding | GO:0035561 | GO:BP | 0.0001656 |
| transcription initiation at RNA polymerase I promoter | GO:0006361 | GO:BP | 0.0001656 |
| positive regulation of hydrolase activity | GO:0051345 | GO:BP | 0.00016807 |
| positive regulation of autophagy | GO:0010508 | GO:BP | 0.00016889 |
| JNK cascade | GO:0007254 | GO:BP | 0.00016898 |
| regulation of translational initiation | GO:0006446 | GO:BP | 0.00016951 |
| nuclear-transcribed mRNA poly(A) tail shortening | GO:0000289 | GO:BP | 0.0001699 |
| rhythmic process | GO:0048511 | GO:BP | 0.00018305 |
| regulation of protein polymerization | GO:0032271 | GO:BP | 0.00018801 |
| epidermal growth factor receptor signaling pathway | GO:0007173 | GO:BP | 0.00019471 |
| regulation of innate immune response | GO:0045088 | GO:BP | 0.00019487 |
| regulation of cell size | GO:0008361 | GO:BP | 0.00019619 |
| regulation of protein binding | GO:0043393 | GO:BP | 0.00020365 |
| multivesicular body sorting pathway | GO:0071985 | GO:BP | 0.00020365 |
| positive regulation of myoblast differentiation | GO:0045663 | GO:BP | 0.00020485 |
| protein targeting to lysosome | GO:0006622 | GO:BP | 0.00021061 |
| regulation of signal transduction by p53 class mediator | GO:1901796 | GO:BP | 0.00021156 |
| regulation of TORC1 signaling | GO:1903432 | GO:BP | 0.00021164 |
| positive regulation of epithelial cell migration | GO:0010634 | GO:BP | 0.00021227 |
| negative regulation of protein kinase activity | GO:0006469 | GO:BP | 0.00021227 |
| response to virus | GO:0009615 | GO:BP | 0.00021721 |
| MAPK cascade | GO:0000165 | GO:BP | 0.00021733 |
| regulation of cell junction assembly | GO:1901888 | GO:BP | 0.0002198 |
| cytoskeleton-dependent intracellular transport | GO:0030705 | GO:BP | 0.0002198 |
| mitochondrial DNA replication | GO:0006264 | GO:BP | 0.00022639 |
| negative regulation of hippo signaling | GO:0035331 | GO:BP | 0.00022639 |
| negative regulation of small GTPase mediated signal tra | GO:0051058 | GO:BP | 0.00023071 |
| regulation of developmental growth | GO:0048638 | GO:BP | 0.00023244 |
| vasculogenesis | GO:0001570 | GO:BP | 0.00023635 |
| epithelium development | GO:0060429 | GO:BP | 0.00023738 |
| Ras protein signal transduction | GO:0007265 | GO:BP | 0.00023747 |
| endothelial cell development | GO:0001885 | GO:BP | 0.00023833 |
| phosphatidylinositol phosphate biosynthetic process | GO:0046854 | GO:BP | 0.00023833 |

| | | | |
|---|---|---|---|
| axonal transport | GO:0098930 | GO:BP | 0.00023833 |
| positive regulation of innate immune response | GO:0045089 | GO:BP | 0.00024556 |
| positive regulation of transcription by RNA polymerase I | GO:0045945 | GO:BP | 0.00025439 |
| tissue migration | GO:0090130 | GO:BP | 0.00025586 |
| regulation of RNA export from nucleus | GO:0046831 | GO:BP | 0.00025662 |
| N-terminal protein amino acid acetylation | GO:0006474 | GO:BP | 0.00025662 |
| regulation of transcription of nucleolar large rRNA by RN | GO:1901836 | GO:BP | 0.00025662 |
| cortical actin cytoskeleton organization | GO:0030866 | GO:BP | 0.00025758 |
| DNA synthesis involved in DNA repair | GO:0000731 | GO:BP | 0.00025758 |
| clathrin-dependent endocytosis | GO:0072583 | GO:BP | 0.00025758 |
| regulation of response to cytokine stimulus | GO:0060759 | GO:BP | 0.00026081 |
| tissue morphogenesis | GO:0048729 | GO:BP | 0.00026194 |
| epithelial to mesenchymal transition | GO:0001837 | GO:BP | 0.00026194 |
| kinetochore assembly | GO:0051382 | GO:BP | 0.00026212 |
| axon development | GO:0061564 | GO:BP | 0.00026381 |
| phosphatidylinositol metabolic process | GO:0046488 | GO:BP | 0.00026381 |
| regulation of cell motility | GO:2000145 | GO:BP | 0.00027032 |
| tRNA surveillance | GO:0106354 | GO:BP | 0.00027362 |
| positive regulation of chromosome condensation | GO:1905821 | GO:BP | 0.00027362 |
| 7-methylguanosine mRNA capping | GO:0006370 | GO:BP | 0.00027362 |
| positive regulation of attachment of mitotic spindle mic | GO:1902425 | GO:BP | 0.00027362 |
| regulation of attachment of mitotic spindle microtubule | GO:1902423 | GO:BP | 0.00027362 |
| nucleoside monophosphate metabolic process | GO:0009123 | GO:BP | 0.0002767 |
| endothelial cell differentiation | GO:0045446 | GO:BP | 0.0002825 |
| somatic cell DNA recombination | GO:0016444 | GO:BP | 0.00029729 |
| somatic diversification of immune receptors via germlir | GO:0002562 | GO:BP | 0.00029729 |
| regulation of telomere maintenance via telomere length | GO:1904356 | GO:BP | 0.00030509 |
| activation of innate immune response | GO:0002218 | GO:BP | 0.00030599 |
| maturation of LSU-rRNA | GO:0000470 | GO:BP | 0.00030724 |
| regulation of nucleotide-excision repair | GO:2000819 | GO:BP | 0.00030724 |
| protein localization to mitochondrion | GO:0070585 | GO:BP | 0.0003091 |
| neuron apoptotic process | GO:0051402 | GO:BP | 0.00031472 |
| cellular biosynthetic process | GO:0044249 | GO:BP | 0.00031705 |
| regulation of vesicle-mediated transport | GO:0060627 | GO:BP | 0.00031856 |
| lysosomal transport | GO:0007041 | GO:BP | 0.0003212 |
| negative regulation of cytoskeleton organization | GO:0051494 | GO:BP | 0.000326 |
| regulation of cell shape | GO:0008360 | GO:BP | 0.000326 |
| actomyosin structure organization | GO:0031032 | GO:BP | 0.00033412 |
| regulation of cytoplasmic pattern recognition receptor s | GO:0039531 | GO:BP | 0.00033689 |
| intrinsic apoptotic signaling pathway | GO:0097193 | GO:BP | 0.00034024 |
| regulation of ERBB signaling pathway | GO:1901184 | GO:BP | 0.00034273 |
| mesenchymal cell differentiation | GO:0048762 | GO:BP | 0.00034393 |
| axo-dendritic transport | GO:0008088 | GO:BP | 0.00035587 |
| embryonic organ development | GO:0048568 | GO:BP | 0.00036068 |
| regulation of hippo signaling | GO:0035330 | GO:BP | 0.00036879 |
| vesicle targeting, rough ER to cis-Golgi | GO:0048207 | GO:BP | 0.00036879 |
| COPII vesicle coating | GO:0048208 | GO:BP | 0.00036879 |
| animal organ morphogenesis | GO:0009887 | GO:BP | 0.00037395 |

| | | | |
|---|---|---|---|
| innate immune response-activating signaling pathway | GO:0002758 | GO:BP | 0.00037395 |
| establishment of protein localization to mitochondrion | GO:0072655 | GO:BP | 0.00037547 |
| blood vessel morphogenesis | GO:0048514 | GO:BP | 0.00038656 |
| phospholipid metabolic process | GO:0006644 | GO:BP | 0.00038665 |
| regulation of protein import into nucleus | GO:0042306 | GO:BP | 0.00039097 |
| tissue development | GO:0009888 | GO:BP | 0.00039968 |
| phosphatidylinositol biosynthetic process | GO:0006661 | GO:BP | 0.00039968 |
| glycerophospholipid metabolic process | GO:0006650 | GO:BP | 0.00040016 |
| nuclear-transcribed mRNA catabolic process, nonsens | GO:0000184 | GO:BP | 0.00040182 |
| Rac protein signal transduction | GO:0016601 | GO:BP | 0.00040182 |
| negative regulation of kinase activity | GO:0033673 | GO:BP | 0.00040465 |
| regulation of neurogenesis | GO:0050767 | GO:BP | 0.00040792 |
| protein modification by small protein removal | GO:0070646 | GO:BP | 0.00040867 |
| protein deacetylation | GO:0006476 | GO:BP | 0.00041536 |
| regulation of protein localization to plasma membrane | GO:1903076 | GO:BP | 0.00045559 |
| post-embryonic development | GO:0009791 | GO:BP | 0.00046299 |
| positive regulation of viral genome replication | GO:0045070 | GO:BP | 0.00046765 |
| mitochondrial genome maintenance | GO:0000002 | GO:BP | 0.00046765 |
| regulation of viral process | GO:0050792 | GO:BP | 0.00048943 |
| somatic diversification of immunoglobulins | GO:0016445 | GO:BP | 0.00049303 |
| positive regulation of proteolysis involved in protein cat | GO:1903052 | GO:BP | 0.00051433 |
| establishment of sister chromatid cohesion | GO:0034085 | GO:BP | 0.00051728 |
| positive regulation of transcription of nucleolar large rI | GO:1901838 | GO:BP | 0.00051728 |
| nuclear mRNA surveillance | GO:0071028 | GO:BP | 0.00051728 |
| nucleolar chromatin organization | GO:1990700 | GO:BP | 0.00051728 |
| DNA topological change | GO:0006265 | GO:BP | 0.00051728 |
| hematopoietic stem cell differentiation | GO:0060218 | GO:BP | 0.00052019 |
| regulation of nervous system development | GO:0051960 | GO:BP | 0.00053404 |
| positive regulation of telomere maintenance | GO:0032206 | GO:BP | 0.00054154 |
| regulation of actomyosin structure organization | GO:0110020 | GO:BP | 0.00054601 |
| cellular senescence | GO:0090398 | GO:BP | 0.00054601 |
| positive regulation of blood vessel endothelial cell migr | GO:0043536 | GO:BP | 0.00056797 |
| intracellular estrogen receptor signaling pathway | GO:0030520 | GO:BP | 0.00056797 |
| actin nucleation | GO:0045010 | GO:BP | 0.00057021 |
| negative regulation of ERBB signaling pathway | GO:1901185 | GO:BP | 0.00057888 |
| positive regulation of G2/M transition of mitotic cell cyc | GO:0010971 | GO:BP | 0.00057888 |
| developmental growth involved in morphogenesis | GO:0060560 | GO:BP | 0.00058099 |
| RNA-templated DNA biosynthetic process | GO:0006278 | GO:BP | 0.00058745 |
| positive regulation of TOR signaling | GO:0032008 | GO:BP | 0.00058745 |
| endoplasmic reticulum tubular network organization | GO:0071786 | GO:BP | 0.00058847 |
| regulation of locomotion | GO:0040012 | GO:BP | 0.00061741 |
| establishment of endothelial barrier | GO:0061028 | GO:BP | 0.00062355 |
| response to decreased oxygen levels | GO:0036293 | GO:BP | 0.00062717 |
| endothelium development | GO:0003158 | GO:BP | 0.00064582 |
| response to oxygen-containing compound | GO:1901700 | GO:BP | 0.00065843 |
| positive regulation by host of viral transcription | GO:0043923 | GO:BP | 0.00065901 |
| primary miRNA processing | GO:0031053 | GO:BP | 0.00065901 |
| regulation of mRNA 3'-end processing | GO:0031440 | GO:BP | 0.00066158 |

| | | | |
|---|---|---|---|
| mitotic spindle elongation | GO:0000022 | GO:BP | 0.00066158 |
| regulation of helicase activity | GO:0051095 | GO:BP | 0.00066158 |
| regulation of mitotic spindle organization | GO:0060236 | GO:BP | 0.00066297 |
| DNA damage response, signal transduction by p53 clas | GO:0030330 | GO:BP | 0.00067605 |
| telomere capping | GO:0016233 | GO:BP | 0.00067889 |
| activation of NF-kappaB-inducing kinase activity | GO:0007250 | GO:BP | 0.00070622 |
| regulation of rRNA processing | GO:2000232 | GO:BP | 0.00070622 |
| maturation of LSU-rRNA from tricistronic rRNA transcrip | GO:0000463 | GO:BP | 0.00070622 |
| spindle midzone assembly | GO:0051255 | GO:BP | 0.00071556 |
| spindle elongation | GO:0051231 | GO:BP | 0.00071556 |
| protein targeting to mitochondrion | GO:0006626 | GO:BP | 0.00073142 |
| membrane assembly | GO:0071709 | GO:BP | 0.00073243 |
| actin filament polymerization | GO:0030041 | GO:BP | 0.00073673 |
| cardiac septum development | GO:0003279 | GO:BP | 0.00074625 |
| organ growth | GO:0035265 | GO:BP | 0.00074625 |
| endosome to lysosome transport | GO:0008333 | GO:BP | 0.00075265 |
| telomere maintenance via telomerase | GO:0007004 | GO:BP | 0.00075265 |
| biosynthetic process | GO:0009058 | GO:BP | 0.00075531 |
| appendage morphogenesis | GO:0035107 | GO:BP | 0.00077964 |
| limb morphogenesis | GO:0035108 | GO:BP | 0.00077964 |
| positive regulation of binding | GO:0051099 | GO:BP | 0.00079566 |
| regulation of JNK cascade | GO:0046328 | GO:BP | 0.00080341 |
| positive regulation of cell cycle G2/M phase transition | GO:1902751 | GO:BP | 0.00082312 |
| vesicle tethering | GO:0099022 | GO:BP | 0.00082312 |
| regulation of stress fiber assembly | GO:0051492 | GO:BP | 0.00082312 |
| import into cell | GO:0098657 | GO:BP | 0.00082546 |
| organic substance biosynthetic process | GO:1901576 | GO:BP | 0.00083662 |
| positive regulation of establishment of protein localizati | GO:1904951 | GO:BP | 0.0008487 |
| negative regulation of Ras protein signal transduction | GO:0046580 | GO:BP | 0.00086296 |
| negative regulation of mRNA processing | GO:0050686 | GO:BP | 0.00086296 |
| mitotic G2 DNA damage checkpoint signaling | GO:0007095 | GO:BP | 0.00086296 |
| positive regulation of cardiac muscle hypertrophy | GO:0010613 | GO:BP | 0.00086296 |
| defense response to virus | GO:0051607 | GO:BP | 0.00088407 |
| protein O-linked glycosylation | GO:0006493 | GO:BP | 0.0009021 |
| meiotic nuclear division | GO:0140013 | GO:BP | 0.0009021 |
| protein lipidation | GO:0006497 | GO:BP | 0.00091467 |
| regulation of actin filament bundle assembly | GO:0032231 | GO:BP | 0.0009234 |
| establishment of mitotic sister chromatid cohesion | GO:0034087 | GO:BP | 0.00093029 |
| positive regulation of nuclear cell cycle DNA replication | GO:0010571 | GO:BP | 0.00093029 |
| negative regulation of centriole replication | GO:0046600 | GO:BP | 0.00093029 |
| mitochondrion localization | GO:0051646 | GO:BP | 0.00093029 |
| nuclear ncRNA surveillance | GO:0071029 | GO:BP | 0.00093029 |
| nuclear polyadenylation-dependent tRNA catabolic pro | GO:0071038 | GO:BP | 0.00093029 |
| nuclear polyadenylation-dependent rRNA catabolic pro | GO:0071035 | GO:BP | 0.00093029 |
| nuclear polyadenylation-dependent ncRNA catabolic p | GO:0071046 | GO:BP | 0.00093029 |
| placenta development | GO:0001890 | GO:BP | 0.00096289 |
| regulation of response to biotic stimulus | GO:0002831 | GO:BP | 0.00096766 |
| mesenchyme development | GO:0060485 | GO:BP | 0.00097371 |

| | | | |
|---|---|---|---|
| modulation by host of symbiont process | GO:0051851 | GO:BP | 0.00100231 |
| telomere maintenance in response to DNA damage | GO:0043247 | GO:BP | 0.00102985 |
| receptor metabolic process | GO:0043112 | GO:BP | 0.00103629 |
| regulation of heterochromatin formation | GO:0031445 | GO:BP | 0.00104045 |
| regulation of heterochromatin organization | GO:0120261 | GO:BP | 0.00104045 |
| positive regulation of TORC1 signaling | GO:1904263 | GO:BP | 0.00107321 |
| negative regulation of apoptotic signaling pathway | GO:2001234 | GO:BP | 0.00108004 |
| negative regulation of Wnt signaling pathway | GO:0030178 | GO:BP | 0.00108016 |
| axonogenesis | GO:0007409 | GO:BP | 0.00108068 |
| regulation of ubiquitin-protein transferase activity | GO:0051438 | GO:BP | 0.001081 |
| positive regulation of macroautophagy | GO:0016239 | GO:BP | 0.00110233 |
| developmental cell growth | GO:0048588 | GO:BP | 0.00111045 |
| regulation of cell population proliferation | GO:0042127 | GO:BP | 0.00111229 |
| Arp2/3 complex-mediated actin nucleation | GO:0034314 | GO:BP | 0.00120625 |
| positive regulation of RNA splicing | GO:0033120 | GO:BP | 0.00120625 |
| response to leukemia inhibitory factor | GO:1990823 | GO:BP | 0.00120904 |
| negative regulation of developmental process | GO:0051093 | GO:BP | 0.00121061 |
| ERAD pathway | GO:0036503 | GO:BP | 0.00121061 |
| Rho protein signal transduction | GO:0007266 | GO:BP | 0.00122095 |
| DNA unwinding involved in DNA replication | GO:0006268 | GO:BP | 0.00124397 |
| rRNA catabolic process | GO:0016075 | GO:BP | 0.00124397 |
| organelle membrane fusion | GO:0090174 | GO:BP | 0.00124926 |
| skeletal system development | GO:0001501 | GO:BP | 0.00126273 |
| positive regulation of nervous system development | GO:0051962 | GO:BP | 0.00129749 |
| cellular response to oxygen-containing compound | GO:1901701 | GO:BP | 0.00130843 |
| positive regulation of DNA biosynthetic process | GO:2000573 | GO:BP | 0.00133095 |
| negative regulation of TOR signaling | GO:0032007 | GO:BP | 0.00133095 |
| protein localization to lysosome | GO:0061462 | GO:BP | 0.00134397 |
| synapse organization | GO:0050808 | GO:BP | 0.00136799 |
| membrane docking | GO:0022406 | GO:BP | 0.00137435 |
| anterograde axonal transport | GO:0008089 | GO:BP | 0.00138397 |
| positive regulation of neurogenesis | GO:0050769 | GO:BP | 0.00138606 |
| G0 to G1 transition | GO:0045023 | GO:BP | 0.00138722 |
| positive regulation of multicellular organismal process | GO:0051240 | GO:BP | 0.00142231 |
| regulation of transport | GO:0051049 | GO:BP | 0.00142764 |
| regulation of protein localization to membrane | GO:1905475 | GO:BP | 0.00144036 |
| regulation of telomere maintenance in response to DNA | GO:1904505 | GO:BP | 0.00146112 |
| somatic recombination of immunoglobulin gene segme | GO:0016447 | GO:BP | 0.0014697 |
| positive regulation of actin filament bundle assembly | GO:0032233 | GO:BP | 0.0014697 |
| positive regulation of cell migration | GO:0030335 | GO:BP | 0.00148074 |
| regulation of autophagosome assembly | GO:2000785 | GO:BP | 0.0015524 |
| positive regulation of cytokinesis | GO:0032467 | GO:BP | 0.0015524 |
| positive regulation of proteasomal protein catabolic pro | GO:1901800 | GO:BP | 0.00155345 |
| regulation of telomere maintenance via telomerase | GO:0032210 | GO:BP | 0.00155951 |
| blood vessel endothelial cell migration | GO:0043534 | GO:BP | 0.00156588 |
| vesicle fusion | GO:0006906 | GO:BP | 0.00156588 |
| transport along microtubule | GO:0010970 | GO:BP | 0.00156851 |
| protein hexamerization | GO:0034214 | GO:BP | 0.00158356 |

| | | | |
|---|---|---|---|
| telomere maintenance via semi-conservative replicatio | GO:0032201 | GO:BP | 0.00158356 |
| membrane biogenesis | GO:0044091 | GO:BP | 0.00158356 |
| cardiac chamber development | GO:0003205 | GO:BP | 0.00158356 |
| rRNA methylation | GO:0031167 | GO:BP | 0.00158356 |
| anaphase-promoting complex-dependent catabolic pro | GO:0031145 | GO:BP | 0.00158356 |
| positive regulation of muscle hypertrophy | GO:0014742 | GO:BP | 0.00158356 |
| translesion synthesis | GO:0019985 | GO:BP | 0.00158356 |
| positive regulation of chromatin binding | GO:0035563 | GO:BP | 0.00158356 |
| rDNA heterochromatin formation | GO:0000183 | GO:BP | 0.00158356 |
| regulation of mitochondrial translation | GO:0070129 | GO:BP | 0.00158356 |
| regulation of telomere capping | GO:1904353 | GO:BP | 0.00158356 |
| regulation of mitotic centrosome separation | GO:0046602 | GO:BP | 0.00158356 |
| RNA polymerase I preinitiation complex assembly | GO:0001188 | GO:BP | 0.00158356 |
| tRNA wobble uridine modification | GO:0002098 | GO:BP | 0.00165638 |
| skeletal muscle satellite cell proliferation | GO:0014841 | GO:BP | 0.00165638 |
| endonucleolytic cleavage of tricistronic rRNA transcript | GO:0000479 | GO:BP | 0.00165638 |
| regulation of hematopoietic stem cell differentiation | GO:1902036 | GO:BP | 0.00165638 |
| mitochondrial fission | GO:0000266 | GO:BP | 0.00167018 |
| positive regulation of protein import into nucleus | GO:0042307 | GO:BP | 0.00167018 |
| vesicle-mediated transport in synapse | GO:0099003 | GO:BP | 0.00170041 |
| regulation of protein depolymerization | GO:1901879 | GO:BP | 0.00172485 |
| regulation of mitochondrial gene expression | GO:0062125 | GO:BP | 0.00174186 |
| positive regulation of cell-substrate junction organizatic | GO:0150117 | GO:BP | 0.00174186 |
| viral protein processing | GO:0019082 | GO:BP | 0.00174186 |
| regulation of G0 to G1 transition | GO:0070316 | GO:BP | 0.00175063 |
| negative regulation of protein localization to membrane | GO:1905476 | GO:BP | 0.00175063 |
| ribonucleoside monophosphate biosynthetic process | GO:0009156 | GO:BP | 0.00175063 |
| lamellipodium assembly | GO:0030032 | GO:BP | 0.00178564 |
| vesicle cytoskeletal trafficking | GO:0099518 | GO:BP | 0.00178564 |
| cellular response to cytokine stimulus | GO:0071345 | GO:BP | 0.00179359 |
| epithelial cell development | GO:0002064 | GO:BP | 0.00179798 |
| positive regulation of protein kinase activity | GO:0045860 | GO:BP | 0.00180783 |
| nuclear pore organization | GO:0006999 | GO:BP | 0.00183184 |
| protein localization to cell cortex | GO:0072697 | GO:BP | 0.00188442 |
| mitotic DNA replication checkpoint signaling | GO:0033314 | GO:BP | 0.00188442 |
| mitotic spindle midzone assembly | GO:0051256 | GO:BP | 0.00188442 |
| transforming growth factor beta receptor superfamily si | GO:0141091 | GO:BP | 0.00190717 |
| positive regulation of ubiquitin-dependent protein catal | GO:2000060 | GO:BP | 0.00190717 |
| negative regulation of cell projection organization | GO:0031345 | GO:BP | 0.00190717 |
| positive regulation of spindle checkpoint | GO:0090232 | GO:BP | 0.00193208 |
| Rap protein signal transduction | GO:0032486 | GO:BP | 0.00193208 |
| positive regulation of stress-activated protein kinase sig | GO:0070304 | GO:BP | 0.00194622 |
| negative regulation of epidermal growth factor receptor | GO:0042059 | GO:BP | 0.00194622 |
| muscle cell proliferation | GO:0033002 | GO:BP | 0.00195713 |
| positive regulation of stress fiber assembly | GO:0051496 | GO:BP | 0.00197168 |
| positive regulation of kinase activity | GO:0033674 | GO:BP | 0.00198307 |
| ribosome assembly | GO:0042255 | GO:BP | 0.00200705 |
| negative regulation of neuron apoptotic process | GO:0043524 | GO:BP | 0.00200746 |

| | | | |
|---|---|---|---|
| morphogenesis of a branching structure | GO:0001763 | GO:BP | 0.00201091 |
| organelle localization by membrane tethering | GO:0140056 | GO:BP | 0.00203918 |
| cellular response to abiotic stimulus | GO:0071214 | GO:BP | 0.00204711 |
| cellular response to environmental stimulus | GO:0104004 | GO:BP | 0.00204711 |
| pattern specification process | GO:0007389 | GO:BP | 0.00205387 |
| response to oxygen levels | GO:0070482 | GO:BP | 0.0020631 |
| response to cytokine | GO:0034097 | GO:BP | 0.0020631 |
| regulation of actin nucleation | GO:0051125 | GO:BP | 0.00212804 |
| Golgi to plasma membrane protein transport | GO:0043001 | GO:BP | 0.00212804 |
| muscle structure development | GO:0061061 | GO:BP | 0.0021395 |
| mRNA stabilization | GO:0048255 | GO:BP | 0.0021395 |
| regulation of protein deacetylation | GO:0090311 | GO:BP | 0.00218608 |
| cellular response to angiotensin | GO:1904385 | GO:BP | 0.00222681 |
| positive regulation of response to biotic stimulus | GO:0002833 | GO:BP | 0.00226125 |
| dendritic spine development | GO:0060996 | GO:BP | 0.00234977 |
| RNA 5'-end processing | GO:0000966 | GO:BP | 0.00239105 |
| negative regulation of DNA-templated transcription, elo | GO:0032785 | GO:BP | 0.00239105 |
| regulation of spindle checkpoint | GO:0090231 | GO:BP | 0.00239105 |
| neural tube development | GO:0021915 | GO:BP | 0.00241731 |
| regulation of protein dephosphorylation | GO:0035304 | GO:BP | 0.00251286 |
| hormone-mediated signaling pathway | GO:0009755 | GO:BP | 0.00251464 |
| regulation of microtubule polymerization | GO:0031113 | GO:BP | 0.0025906 |
| regulation of neuron apoptotic process | GO:0043523 | GO:BP | 0.00260317 |
| outflow tract morphogenesis | GO:0003151 | GO:BP | 0.00261037 |
| multicellular organismal process | GO:0032501 | GO:BP | 0.00276184 |
| regulation of protein polyubiquitination | GO:1902914 | GO:BP | 0.00278613 |
| nuclear membrane reassembly | GO:0031468 | GO:BP | 0.00278613 |
| protein K63-linked deubiquitination | GO:0070536 | GO:BP | 0.00278613 |
| immune response-activating signaling pathway | GO:0002757 | GO:BP | 0.0027935 |
| mismatch repair | GO:0006298 | GO:BP | 0.0028721 |
| negative regulation of intracellular protein transport | GO:0090317 | GO:BP | 0.0028721 |
| negative regulation of cell migration | GO:0030336 | GO:BP | 0.0029042 |
| paraxial mesoderm development | GO:0048339 | GO:BP | 0.00292801 |
| glial cell differentiation | GO:0010001 | GO:BP | 0.00295815 |
| immune system development | GO:0002520 | GO:BP | 0.00296292 |
| endosome organization | GO:0007032 | GO:BP | 0.00297711 |
| negative regulation of binding | GO:0051100 | GO:BP | 0.00303594 |
| positive regulation of plasma membrane bounded cell p | GO:0120034 | GO:BP | 0.00303594 |
| cellular response to leukemia inhibitory factor | GO:1990830 | GO:BP | 0.00303594 |
| myelination | GO:0042552 | GO:BP | 0.00310994 |
| meiotic chromosome condensation | GO:0010032 | GO:BP | 0.00313333 |
| regulation of ribosome biogenesis | GO:0090069 | GO:BP | 0.00313333 |
| oxidative RNA demethylation | GO:0035513 | GO:BP | 0.00313333 |
| snRNA transport | GO:0051030 | GO:BP | 0.00313333 |
| positive regulation of mRNA 3'-end processing | GO:0031442 | GO:BP | 0.00313333 |
| regulation of endoplasmic reticulum tubular network or | GO:1903371 | GO:BP | 0.00313333 |
| regulation of pattern recognition receptor signaling path | GO:0062207 | GO:BP | 0.00316155 |
| ventricular septum development | GO:0003281 | GO:BP | 0.00320755 |

| | | | |
|---|---|---|---|
| sexual reproduction | GO:0019953 | GO:BP | 0.00327748 |
| negative regulation of mRNA catabolic process | GO:1902373 | GO:BP | 0.00332958 |
| regulation of epidermal growth factor receptor signaling | GO:0042058 | GO:BP | 0.00332958 |
| dendrite morphogenesis | GO:0048813 | GO:BP | 0.00334949 |
| circadian rhythm | GO:0007623 | GO:BP | 0.00335462 |
| positive regulation of programmed cell death | GO:0043068 | GO:BP | 0.00348294 |
| skeletal muscle cell proliferation | GO:0014856 | GO:BP | 0.00350822 |
| regulation of early endosome to late endosome transpo | GO:2000641 | GO:BP | 0.00351231 |
| negative regulation of protein-containing complex disas | GO:0043242 | GO:BP | 0.00352195 |
| response to oxidative stress | GO:0006979 | GO:BP | 0.00353798 |
| nucleus localization | GO:0051647 | GO:BP | 0.00354048 |
| positive regulation of mRNA splicing, via spliceosome | GO:0048026 | GO:BP | 0.00354048 |
| stress-activated protein kinase signaling cascade | GO:0031098 | GO:BP | 0.00354473 |
| hindbrain development | GO:0030902 | GO:BP | 0.00354539 |
| pyrimidine-containing compound metabolic process | GO:0072527 | GO:BP | 0.00358685 |
| extrinsic apoptotic signaling pathway | GO:0097191 | GO:BP | 0.00360639 |
| positive regulation of apoptotic process | GO:0043065 | GO:BP | 0.00364214 |
| protein localization to Golgi apparatus | GO:0034067 | GO:BP | 0.00365124 |
| regulation of blood vessel endothelial cell migration | GO:0043535 | GO:BP | 0.00369792 |
| cellular response to starvation | GO:0009267 | GO:BP | 0.00370522 |
| nucleoside monophosphate biosynthetic process | GO:0009124 | GO:BP | 0.00375867 |
| regulation of type I interferon-mediated signaling pathw | GO:0060338 | GO:BP | 0.00375867 |
| response to steroid hormone | GO:0048545 | GO:BP | 0.00390682 |
| somatic diversification of immunoglobulins involved in i | GO:0002208 | GO:BP | 0.00396956 |
| somatic recombination of immunoglobulin genes involv | GO:0002204 | GO:BP | 0.00396956 |
| isotype switching | GO:0045190 | GO:BP | 0.00396956 |
| response to leucine | GO:0043201 | GO:BP | 0.00414177 |
| RNA capping | GO:0036260 | GO:BP | 0.00414177 |
| negative regulation of cell motility | GO:2000146 | GO:BP | 0.00415694 |
| synaptic vesicle cycle | GO:0099504 | GO:BP | 0.00417166 |
| ubiquitin-dependent protein catabolic process via the n | GO:0043162 | GO:BP | 0.00421606 |
| protein transmembrane import into intracellular organe | GO:0044743 | GO:BP | 0.00421606 |
| vesicle transport along microtubule | GO:0047496 | GO:BP | 0.00424807 |
| negative regulation of mRNA splicing, via spliceosome | GO:0048025 | GO:BP | 0.00438758 |
| response to platelet-derived growth factor | GO:0036119 | GO:BP | 0.00438758 |
| regulation of establishment of cell polarity | GO:2000114 | GO:BP | 0.00438758 |
| epithelial tube morphogenesis | GO:0060562 | GO:BP | 0.00438758 |
| mitochondrial DNA metabolic process | GO:0032042 | GO:BP | 0.00438758 |
| ensheathment of neurons | GO:0007272 | GO:BP | 0.00443968 |
| axon ensheathment | GO:0008366 | GO:BP | 0.00443968 |
| gliogenesis | GO:0042063 | GO:BP | 0.00455572 |
| lipoprotein biosynthetic process | GO:0042158 | GO:BP | 0.00462123 |
| myeloid cell development | GO:0061515 | GO:BP | 0.00462123 |
| response to topologically incorrect protein | GO:0035966 | GO:BP | 0.00465769 |
| regulation of circadian rhythm | GO:0042752 | GO:BP | 0.00466802 |
| protein localization to endosome | GO:0036010 | GO:BP | 0.00466802 |
| protein methylation | GO:0006479 | GO:BP | 0.00468477 |
| protein alkylation | GO:0008213 | GO:BP | 0.00468477 |

| | | | |
|---|---|---|---|
| response to extracellular stimulus | GO:0009991 | GO:BP | 0.00468477 |
| regulation of dephosphorylation | GO:0035303 | GO:BP | 0.0047192 |
| positive regulation of actin nucleation | GO:0051127 | GO:BP | 0.00472186 |
| ribosomal subunit export from nucleus | GO:0000054 | GO:BP | 0.00472186 |
| ribosome localization | GO:0033750 | GO:BP | 0.00472186 |
| protein K6-linked ubiquitination | GO:0085020 | GO:BP | 0.00472186 |
| regulation of mRNA export from nucleus | GO:0010793 | GO:BP | 0.00476027 |
| U4 snRNA 3'-end processing | GO:0034475 | GO:BP | 0.00476027 |
| endonucleolytic cleavage in 5'-ETS of tricistronic rRNA t | GO:0000480 | GO:BP | 0.00476027 |
| positive regulation of early endosome to late endosome | GO:2000643 | GO:BP | 0.00476027 |
| positive regulation of Arp2/3 complex-mediated actin nu | GO:2000601 | GO:BP | 0.00476027 |
| monoubiquitinated protein deubiquitination | GO:0035520 | GO:BP | 0.00476027 |
| polyadenylation-dependent RNA catabolic process | GO:0043633 | GO:BP | 0.00476027 |
| random inactivation of X chromosome | GO:0060816 | GO:BP | 0.00476027 |
| positive regulation of mitotic cytokinesis | GO:1903490 | GO:BP | 0.00476027 |
| embryonic cleavage | GO:0040016 | GO:BP | 0.00476027 |
| establishment of protein localization to plasma membra | GO:0061951 | GO:BP | 0.00476027 |
| transcription initiation at RNA polymerase III promoter | GO:0006384 | GO:BP | 0.00476027 |
| copper ion transmembrane transport | GO:0035434 | GO:BP | 0.00476027 |
| polyadenylation-dependent ncRNA catabolic process | GO:0043634 | GO:BP | 0.00476027 |
| negative regulation of Schwann cell proliferation | GO:0010626 | GO:BP | 0.00476027 |
| regulation by virus of viral protein levels in host cell | GO:0046719 | GO:BP | 0.00476027 |
| 2'-deoxyribonucleotide metabolic process | GO:0009394 | GO:BP | 0.00477396 |
| regulation of cellular response to transforming growth fa | GO:1903844 | GO:BP | 0.00477396 |
| cytochrome complex assembly | GO:0017004 | GO:BP | 0.00477396 |
| heart morphogenesis | GO:0003007 | GO:BP | 0.00477396 |
| neuron migration | GO:0001764 | GO:BP | 0.00492166 |
| protein transmembrane transport | GO:0071806 | GO:BP | 0.00493961 |
| gland development | GO:0048732 | GO:BP | 0.00496605 |
| negative regulation of locomotion | GO:0040013 | GO:BP | 0.00502559 |
| retrograde vesicle-mediated transport, Golgi to endopla | GO:0006890 | GO:BP | 0.00502559 |
| meiotic spindle assembly | GO:0090306 | GO:BP | 0.00512247 |
| double-strand break repair via break-induced replicatio | GO:0000727 | GO:BP | 0.00512247 |
| negative regulation of transcription by RNA polymerase | GO:0016479 | GO:BP | 0.00512247 |
| positive regulation of mitotic cell cycle spindle assembl | GO:0090267 | GO:BP | 0.00512247 |
| progesterone receptor signaling pathway | GO:0050847 | GO:BP | 0.00512247 |
| negative regulation of centrosome cycle | GO:0046606 | GO:BP | 0.00512247 |
| negative regulation of centrosome duplication | GO:0010826 | GO:BP | 0.00512247 |
| immune response-regulating signaling pathway | GO:0002764 | GO:BP | 0.00516497 |
| response to hypoxia | GO:0001666 | GO:BP | 0.00518586 |
| mitochondrial RNA modification | GO:1900864 | GO:BP | 0.00524296 |
| positive regulation of ubiquitin protein ligase activity | GO:1904668 | GO:BP | 0.00524296 |
| nuclear pore complex assembly | GO:0051292 | GO:BP | 0.00524296 |
| actomyosin contractile ring organization | GO:0044837 | GO:BP | 0.00524296 |
| mitochondrial tRNA modification | GO:0070900 | GO:BP | 0.00524296 |
| chromatin remodeling at centromere | GO:0031055 | GO:BP | 0.00524296 |
| CENP-A containing chromatin assembly | GO:0034080 | GO:BP | 0.00524296 |
| cranial skeletal system development | GO:1904888 | GO:BP | 0.00525394 |

| | | | |
|---|---|---|---|
| regulation of actin polymerization or depolymerization | GO:0008064 | GO:BP | 0.00528596 |
| ncRNA 5'-end processing | GO:0034471 | GO:BP | 0.00532269 |
| positive regulation of neuron migration | GO:2001224 | GO:BP | 0.00532269 |
| regulation of mitotic spindle checkpoint | GO:1903504 | GO:BP | 0.00532269 |
| cellular response to platelet-derived growth factor stimu | GO:0036120 | GO:BP | 0.00532269 |
| regulation of mitotic cell cycle spindle assembly checkp | GO:0090266 | GO:BP | 0.00532269 |
| embryonic cranial skeleton morphogenesis | GO:0048701 | GO:BP | 0.00532285 |
| pyrimidine nucleotide metabolic process | GO:0006220 | GO:BP | 0.0053339 |
| cardiac muscle tissue growth | GO:0055017 | GO:BP | 0.00537209 |
| translational elongation | GO:0006414 | GO:BP | 0.00549729 |
| regulation of cell-substrate adhesion | GO:0010810 | GO:BP | 0.00555675 |
| negative regulation of protein serine/threonine kinase a | GO:0071901 | GO:BP | 0.00559427 |
| regionalization | GO:0003002 | GO:BP | 0.00564321 |
| viral release from host cell | GO:0019076 | GO:BP | 0.00569108 |
| exit from host cell | GO:0035891 | GO:BP | 0.00569108 |
| reproduction | GO:0000003 | GO:BP | 0.00573032 |
| negative regulation of autophagy | GO:0010507 | GO:BP | 0.00575999 |
| meiotic chromosome segregation | GO:0045132 | GO:BP | 0.00577778 |
| neurotrophin TRK receptor signaling pathway | GO:0048011 | GO:BP | 0.00577778 |
| angiogenesis | GO:0001525 | GO:BP | 0.00578625 |
| microtubule depolymerization | GO:0007019 | GO:BP | 0.00587338 |
| regulation of telomerase activity | GO:0051972 | GO:BP | 0.00587338 |
| negative regulation of canonical Wnt signaling pathway | GO:0090090 | GO:BP | 0.00604808 |
| response to angiotensin | GO:1990776 | GO:BP | 0.0061064 |
| regulation of cellular response to growth factor stimulus | GO:0090287 | GO:BP | 0.0062898 |
| cardiac muscle cell proliferation | GO:0060038 | GO:BP | 0.00642102 |
| cardiac ventricle development | GO:0003231 | GO:BP | 0.00643328 |
| homologous recombination | GO:0035825 | GO:BP | 0.00648792 |
| positive regulation of mitotic sister chromatid separatio | GO:1901970 | GO:BP | 0.00658496 |
| membrane lipid biosynthetic process | GO:0046467 | GO:BP | 0.00658496 |
| meiotic spindle organization | GO:0000212 | GO:BP | 0.00658496 |
| negative regulation of cyclin-dependent protein serine/t | GO:0045736 | GO:BP | 0.00658496 |
| negative regulation of transcription elongation by RNA p | GO:0034244 | GO:BP | 0.00658496 |
| protein localization to nucleolus | GO:1902570 | GO:BP | 0.00658496 |
| protein localization to microtubule | GO:0035372 | GO:BP | 0.00658496 |
| pore complex assembly | GO:0046931 | GO:BP | 0.00658496 |
| cellular response to decreased oxygen levels | GO:0036294 | GO:BP | 0.00658496 |
| cellular response to ionizing radiation | GO:0071479 | GO:BP | 0.00665474 |
| regulation of spindle assembly | GO:0090169 | GO:BP | 0.00681884 |
| positive regulation of type I interferon production | GO:0032481 | GO:BP | 0.00682073 |
| presynaptic endocytosis | GO:0140238 | GO:BP | 0.00682073 |
| reproductive process | GO:0022414 | GO:BP | 0.00682349 |
| face development | GO:0060324 | GO:BP | 0.00684965 |
| deoxyribonucleotide metabolic process | GO:0009262 | GO:BP | 0.00685445 |
| deoxyribose phosphate metabolic process | GO:0019692 | GO:BP | 0.00685445 |
| positive regulation of cell motility | GO:2000147 | GO:BP | 0.00710218 |
| positive regulation of protein polymerization | GO:0032273 | GO:BP | 0.00726237 |
| negative regulation of miRNA metabolic process | GO:2000629 | GO:BP | 0.0073175 |

| | | | |
|---|---|---|---|
| regulation of Arp2/3 complex-mediated actin nucleatioı | GO:0034315 | GO:BP | 0.0073175 |
| positive regulation of protein transport | GO:0051222 | GO:BP | 0.00735181 |
| histone mRNA metabolic process | GO:0008334 | GO:BP | 0.00735181 |
| regulation of translation in response to stress | GO:0043555 | GO:BP | 0.00735181 |
| response to nutrient levels | GO:0031667 | GO:BP | 0.00736238 |
| response to testosterone | GO:0033574 | GO:BP | 0.00759217 |
| positive regulation of protein binding | GO:0032092 | GO:BP | 0.00764348 |
| immunoglobulin production involved in immunoglobulir | GO:0002381 | GO:BP | 0.00764348 |
| glycerolipid metabolic process | GO:0046486 | GO:BP | 0.00805979 |
| stem cell proliferation | GO:0072089 | GO:BP | 0.00807183 |
| magnesium ion transmembrane transport | GO:1903830 | GO:BP | 0.00808614 |
| positive regulation of telomere capping | GO:1904355 | GO:BP | 0.00808614 |
| mitotic DNA replication | GO:1902969 | GO:BP | 0.00808614 |
| skeletal system morphogenesis | GO:0048705 | GO:BP | 0.00808614 |
| regulation of sister chromatid cohesion | GO:0007063 | GO:BP | 0.00808614 |
| positive regulation of protein polyubiquitination | GO:1902916 | GO:BP | 0.00808614 |
| regulation of DNA strand elongation | GO:0060382 | GO:BP | 0.00808614 |
| protein-containing complex localization | GO:0031503 | GO:BP | 0.00812781 |
| cellular response to nerve growth factor stimulus | GO:1990090 | GO:BP | 0.0082266 |
| negative regulation of intracellular transport | GO:0032387 | GO:BP | 0.0082266 |
| regulation of transforming growth factor beta receptor s | GO:0017015 | GO:BP | 0.00845491 |
| sphingolipid biosynthetic process | GO:0030148 | GO:BP | 0.00859978 |
| regulation of epithelial cell differentiation | GO:0030856 | GO:BP | 0.0087522 |
| regulation of actin filament length | GO:0030832 | GO:BP | 0.0087522 |
| neural tube formation | GO:0001841 | GO:BP | 0.00879353 |
| response to arsenic-containing substance | GO:0046685 | GO:BP | 0.00879353 |
| regulation of Ras protein signal transduction | GO:0046578 | GO:BP | 0.00879353 |
| regulation of microtubule depolymerization | GO:0031114 | GO:BP | 0.00879353 |
| regulation of intracellular steroid hormone receptor sigı | GO:0033143 | GO:BP | 0.00879353 |
| regulation of lamellipodium organization | GO:1902743 | GO:BP | 0.00879353 |
| Fc receptor mediated stimulatory signaling pathway | GO:0002431 | GO:BP | 0.00879353 |
| multivesicular body assembly | GO:0036258 | GO:BP | 0.00879353 |
| negative regulation of extrinsic apoptotic signaling path | GO:2001237 | GO:BP | 0.0089538 |
| positive regulation of axonogenesis | GO:0050772 | GO:BP | 0.00899286 |
| hippocampus development | GO:0021766 | GO:BP | 0.00900664 |
| heart growth | GO:0060419 | GO:BP | 0.00903233 |
| synaptic vesicle recycling | GO:0036465 | GO:BP | 0.00903233 |
| embryonic placenta development | GO:0001892 | GO:BP | 0.00904955 |
| positive regulation of proteasomal ubiquitin-dependent | GO:0032436 | GO:BP | 0.00904955 |
| blastocyst growth | GO:0001832 | GO:BP | 0.0092865 |
| DNA repair-dependent chromatin remodeling | GO:0140861 | GO:BP | 0.0092865 |
| positive regulation of ubiquitin-protein transferase activ | GO:0051443 | GO:BP | 0.0092865 |
| lymph vessel morphogenesis | GO:0036303 | GO:BP | 0.0092865 |
| regulation of mitochondrial membrane permeability | GO:0046902 | GO:BP | 0.0092865 |
| positive regulation of stress-activated MAPK cascade | GO:0032874 | GO:BP | 0.0092865 |
| multicellular organismal-level homeostasis | GO:0048871 | GO:BP | 0.00929167 |
| phosphatidylinositol dephosphorylation | GO:0046856 | GO:BP | 0.00932166 |
| regulation of establishment or maintenance of cell pola | GO:0032878 | GO:BP | 0.00932166 |

| | | | |
|---|---|---|---|
| establishment of protein localization to organelle | GO:0072594 | GO:BP | 0.00932166 |
| positive regulation of telomere maintenance in respons | GO:1904507 | GO:BP | 0.00973963 |
| regulation of peptidyl-threonine phosphorylation | GO:0010799 | GO:BP | 0.00973963 |
| hair follicle maturation | GO:0048820 | GO:BP | 0.00973963 |
| nucleobase-containing small molecule metabolic proce | GO:0055086 | GO:BP | 0.01008107 |
| regulation of intracellular estrogen receptor signaling pa | GO:0033146 | GO:BP | 0.0101487 |
| nucleobase metabolic process | GO:0009112 | GO:BP | 0.0101487 |
| biological process involved in symbiotic interaction | GO:0044403 | GO:BP | 0.01015552 |
| regulation of lymphangiogenesis | GO:1901490 | GO:BP | 0.01019358 |
| nuclear cell cycle DNA replication initiation | GO:1902315 | GO:BP | 0.01019358 |
| cell cycle DNA replication initiation | GO:1902292 | GO:BP | 0.01019358 |
| mitotic DNA replication initiation | GO:1902975 | GO:BP | 0.01019358 |
| regulation of axonogenesis | GO:0050770 | GO:BP | 0.01019358 |
| regulation of membrane tubulation | GO:1903525 | GO:BP | 0.01019358 |
| rRNA 5'-end processing | GO:0000967 | GO:BP | 0.01019358 |
| positive regulation of DNA damage checkpoint | GO:2000003 | GO:BP | 0.01019358 |
| endonucleolytic cleavage to generate mature 5'-end of S | GO:0000472 | GO:BP | 0.01019358 |
| CUT catabolic process | GO:0071034 | GO:BP | 0.01019358 |
| CUT metabolic process | GO:0071043 | GO:BP | 0.01019358 |
| positive regulation of locomotion | GO:0040017 | GO:BP | 0.01019633 |
| negative regulation of cell differentiation | GO:0045596 | GO:BP | 0.01037533 |
| regulation of DNA-binding transcription factor activity | GO:0051090 | GO:BP | 0.01039136 |
| regulation of actin filament polymerization | GO:0030833 | GO:BP | 0.01044604 |
| organelle disassembly | GO:1903008 | GO:BP | 0.01046414 |
| tRNA aminoacylation | GO:0043039 | GO:BP | 0.01047494 |
| regulation of cellular response to insulin stimulus | GO:1900076 | GO:BP | 0.0106354 |
| artery development | GO:0060840 | GO:BP | 0.01079982 |
| cellular response to topologically incorrect protein | GO:0035967 | GO:BP | 0.01079982 |
| negative regulation of protein depolymerization | GO:1901880 | GO:BP | 0.01102442 |
| synaptic vesicle endocytosis | GO:0048488 | GO:BP | 0.01102442 |
| regulation of endocytosis | GO:0030100 | GO:BP | 0.01108305 |
| positive regulation of protein serine/threonine kinase ac | GO:0071902 | GO:BP | 0.01108305 |
| regulation of gene silencing by regulatory ncRNA | GO:0060966 | GO:BP | 0.01123299 |
| positive regulation of axon extension | GO:0045773 | GO:BP | 0.01135357 |
| exonucleolytic catabolism of deadenylated mRNA | GO:0043928 | GO:BP | 0.01146557 |
| endoplasmic reticulum membrane organization | GO:0090158 | GO:BP | 0.01146557 |
| CDP-diacylglycerol biosynthetic process | GO:0016024 | GO:BP | 0.01146557 |
| positive regulation of cell size | GO:0045793 | GO:BP | 0.01146557 |
| positive regulation of protein sumoylation | GO:0033235 | GO:BP | 0.01146557 |
| CDP-diacylglycerol metabolic process | GO:0046341 | GO:BP | 0.01146557 |
| regulation of protein K63-linked ubiquitination | GO:1900044 | GO:BP | 0.01146557 |
| regulation of protein neddylation | GO:2000434 | GO:BP | 0.0116162 |
| nuclear-transcribed mRNA catabolic process, exonucle | GO:0000291 | GO:BP | 0.0116162 |
| response to light stimulus | GO:0009416 | GO:BP | 0.0116162 |
| regulation of viral transcription | GO:0046782 | GO:BP | 0.0116162 |
| negative regulation of GTPase activity | GO:0034260 | GO:BP | 0.0116162 |
| protein import into mitochondrial matrix | GO:0030150 | GO:BP | 0.0116162 |
| positive regulation of NLRP3 inflammasome complex as | GO:1900227 | GO:BP | 0.0116162 |

| | | | |
|---|---|---|---|
| sex-chromosome dosage compensation | GO:0007549 | GO:BP | 0.0116162 |
| negative regulation of neuron projection development | GO:0010977 | GO:BP | 0.01165309 |
| Fc-epsilon receptor signaling pathway | GO:0038095 | GO:BP | 0.01187335 |
| cell-substrate adhesion | GO:0031589 | GO:BP | 0.01190227 |
| positive regulation of DNA-binding transcription factor a | GO:0051091 | GO:BP | 0.01214338 |
| carbohydrate transport | GO:0008643 | GO:BP | 0.01219706 |
| cellular response to chemical stress | GO:0062197 | GO:BP | 0.01220802 |
| postsynapse organization | GO:0099173 | GO:BP | 0.01223698 |
| response to unfolded protein | GO:0006986 | GO:BP | 0.01232557 |
| ventricular septum morphogenesis | GO:0060412 | GO:BP | 0.01242233 |
| myoblast differentiation | GO:0045445 | GO:BP | 0.0124435 |
| sulfur compound biosynthetic process | GO:0044272 | GO:BP | 0.01250445 |
| cellular response to radiation | GO:0071478 | GO:BP | 0.01266815 |
| ceramide metabolic process | GO:0006672 | GO:BP | 0.01292465 |
| cellular response to oxygen levels | GO:0071453 | GO:BP | 0.01293903 |
| multivesicular body organization | GO:0036257 | GO:BP | 0.01295405 |
| protein insertion into ER membrane | GO:0045048 | GO:BP | 0.01295405 |
| astral microtubule organization | GO:0030953 | GO:BP | 0.0130726 |
| regulation of mitotic cytokinesis | GO:1902412 | GO:BP | 0.0130726 |
| regulation of Fc receptor mediated stimulatory signaling | GO:0060368 | GO:BP | 0.0130726 |
| paraxial mesoderm morphogenesis | GO:0048340 | GO:BP | 0.0130726 |
| histone mRNA catabolic process | GO:0071044 | GO:BP | 0.0130726 |
| manganese ion transport | GO:0006828 | GO:BP | 0.0130726 |
| sphingomyelin biosynthetic process | GO:0006686 | GO:BP | 0.0130726 |
| tRNA transcription | GO:0009304 | GO:BP | 0.0130726 |
| negative regulation of cellular response to growth factor | GO:0090288 | GO:BP | 0.0130726 |
| response to hydrogen peroxide | GO:0042542 | GO:BP | 0.01326054 |
| negative regulation of supramolecular fiber organization | GO:1902904 | GO:BP | 0.01329416 |
| regulation of insulin receptor signaling pathway | GO:0046626 | GO:BP | 0.01334887 |
| regulation of erythrocyte differentiation | GO:0045646 | GO:BP | 0.01334887 |
| regulation of stress-activated protein kinase signaling c | GO:0070302 | GO:BP | 0.01334887 |
| purine nucleoside monophosphate metabolic process | GO:0009126 | GO:BP | 0.01334887 |
| macromolecule biosynthetic process | GO:0009059 | GO:BP | 0.01338507 |
| primary neural tube formation | GO:0014020 | GO:BP | 0.01339608 |
| embryonic limb morphogenesis | GO:0030326 | GO:BP | 0.01354573 |
| protein deubiquitination | GO:0016579 | GO:BP | 0.01354573 |
| proteoglycan biosynthetic process | GO:0030166 | GO:BP | 0.01354573 |
| embryonic appendage morphogenesis | GO:0035113 | GO:BP | 0.01354573 |
| negative regulation of transforming growth factor beta re | GO:0030512 | GO:BP | 0.01354573 |
| transcription-dependent tethering of RNA polymerase II | GO:0000972 | GO:BP | 0.01366872 |
| regulation of RNA polymerase II transcription preinitiatio | GO:0045898 | GO:BP | 0.01366872 |
| ribosomal large subunit export from nucleus | GO:0000055 | GO:BP | 0.01366872 |
| chromosome movement towards spindle pole | GO:0051305 | GO:BP | 0.01366872 |
| peptidyl-lysine monomethylation | GO:0018026 | GO:BP | 0.01366872 |
| negative regulation of gene expression via CpG island m | GO:0044027 | GO:BP | 0.01366872 |
| cardiac septum morphogenesis | GO:0060411 | GO:BP | 0.01366872 |
| ncRNA export from nucleus | GO:0097064 | GO:BP | 0.01366872 |
| cellular response to progesterone stimulus | GO:0071393 | GO:BP | 0.01366872 |

| | | | |
|---|---|---|---|
| regulation of mitochondrial mRNA stability | GO:0044528 | GO:BP | 0.01366872 |
| negative regulation of double-strand break repair via no | GO:2001033 | GO:BP | 0.01366872 |
| positive regulation of helicase activity | GO:0051096 | GO:BP | 0.01366872 |
| neural tube closure | GO:0001843 | GO:BP | 0.01366872 |
| negative regulation of G1/S transition of mitotic cell cyc | GO:2000134 | GO:BP | 0.01374142 |
| regulation of viral genome replication | GO:0045069 | GO:BP | 0.01375078 |
| regulation of cyclin-dependent protein serine/threonine | GO:0000079 | GO:BP | 0.01375078 |
| execution phase of apoptosis | GO:0097194 | GO:BP | 0.01377135 |
| cardiac muscle hypertrophy | GO:0003300 | GO:BP | 0.01377135 |
| extrinsic apoptotic signaling pathway via death domain | GO:0008625 | GO:BP | 0.01377135 |
| cellular response to type I interferon | GO:0071357 | GO:BP | 0.01377135 |
| exonucleolytic trimming to generate mature 3'-end of 5. | GO:0000467 | GO:BP | 0.01381927 |
| negative regulation of retrograde protein transport, ER t | GO:1904153 | GO:BP | 0.01381927 |
| regulation of ribonucleoprotein complex localization | GO:2000197 | GO:BP | 0.01381927 |
| regulation of DNA-directed DNA polymerase activity | GO:1900262 | GO:BP | 0.01381927 |
| cardiac pacemaker cell development | GO:0060926 | GO:BP | 0.01381927 |
| pyrimidine nucleoside diphosphate metabolic process | GO:0009138 | GO:BP | 0.01381927 |
| spindle assembly involved in female meiosis | GO:0007056 | GO:BP | 0.01381927 |
| Schwann cell proliferation | GO:0014010 | GO:BP | 0.01381927 |
| positive regulation of centriole replication | GO:0046601 | GO:BP | 0.01381927 |
| aggrephagy | GO:0035973 | GO:BP | 0.01381927 |
| regulation of wound healing, spreading of epidermal cel | GO:1903689 | GO:BP | 0.01381927 |
| RNA secondary structure unwinding | GO:0010501 | GO:BP | 0.01381927 |
| peptidyl-methionine modification | GO:0018206 | GO:BP | 0.01381927 |
| positive regulation of sister chromatid cohesion | GO:0045876 | GO:BP | 0.01381927 |
| regulation of Schwann cell proliferation | GO:0010624 | GO:BP | 0.01381927 |
| positive regulation of DNA-directed DNA polymerase ac | GO:1900264 | GO:BP | 0.01381927 |
| actomyosin contractile ring assembly | GO:0000915 | GO:BP | 0.01381927 |
| regulation of glomerular filtration | GO:0003093 | GO:BP | 0.01381927 |
| regulation of protein localization to cell cortex | GO:1904776 | GO:BP | 0.01381927 |
| mRNA pseudouridine synthesis | GO:1990481 | GO:BP | 0.01381927 |
| assembly of actomyosin apparatus involved in cytokine | GO:0000912 | GO:BP | 0.01381927 |
| DNA dealkylation involved in DNA repair | GO:0006307 | GO:BP | 0.01381927 |
| amino acid activation | GO:0043038 | GO:BP | 0.01385762 |
| regulation of post-transcriptional gene silencing by regu | GO:1900368 | GO:BP | 0.01392876 |
| negative regulation of microtubule depolymerization | GO:0007026 | GO:BP | 0.01392876 |
| mitotic cytokinetic process | GO:1902410 | GO:BP | 0.01392876 |
| negative regulation of RNA splicing | GO:0033119 | GO:BP | 0.01392876 |
| magnesium ion transport | GO:0015693 | GO:BP | 0.01409083 |
| regulation of protein export from nucleus | GO:0046825 | GO:BP | 0.01409083 |
| mitochondrion distribution | GO:0048311 | GO:BP | 0.01409083 |
| dosage compensation by inactivation of X chromosome | GO:0009048 | GO:BP | 0.01409083 |
| nucleobase biosynthetic process | GO:0046112 | GO:BP | 0.01409083 |
| regulation of DNA methylation-dependent heterochrom | GO:0090308 | GO:BP | 0.01409083 |
| membrane fusion | GO:0061025 | GO:BP | 0.01409083 |
| pseudouridine synthesis | GO:0001522 | GO:BP | 0.01409083 |
| lamellipodium morphogenesis | GO:0072673 | GO:BP | 0.01409083 |
| regulation of extent of cell growth | GO:0061387 | GO:BP | 0.01445103 |

| | | | |
|---|---|---|---|
| regulation of endothelial cell migration | GO:0010594 | GO:BP | 0.01445324 |
| response to nerve growth factor | GO:1990089 | GO:BP | 0.01451809 |
| body morphogenesis | GO:0010171 | GO:BP | 0.01451809 |
| immune response-regulating cell surface receptor signa | GO:0002433 | GO:BP | 0.01465903 |
| negative regulation of double-strand break repair via ho | GO:2000042 | GO:BP | 0.01465903 |
| Fc-gamma receptor signaling pathway involved in phag | GO:0038096 | GO:BP | 0.01465903 |
| regulation of receptor-mediated endocytosis | GO:0048259 | GO:BP | 0.01471545 |
| negative regulation of protein catabolic process | GO:0042177 | GO:BP | 0.01471545 |
| negative regulation of protein-containing complex asse | GO:0031333 | GO:BP | 0.0150564 |
| fibroblast migration | GO:0010761 | GO:BP | 0.01513242 |
| stress-activated MAPK cascade | GO:0051403 | GO:BP | 0.01513242 |
| response to antibiotic | GO:0046677 | GO:BP | 0.01542685 |
| regulation of cell migration involved in sprouting angiog | GO:0090049 | GO:BP | 0.01542685 |
| female meiotic nuclear division | GO:0007143 | GO:BP | 0.01542685 |
| Fc receptor signaling pathway | GO:0038093 | GO:BP | 0.01566717 |
| cellular response to hypoxia | GO:0071456 | GO:BP | 0.0160421 |
| negative regulation of cell-substrate adhesion | GO:0010812 | GO:BP | 0.016103 |
| pigment biosynthetic process | GO:0046148 | GO:BP | 0.016103 |
| mitochondrial fusion | GO:0008053 | GO:BP | 0.01614443 |
| regulation of neuron projection regeneration | GO:0070570 | GO:BP | 0.01614443 |
| SMAD protein signal transduction | GO:0060395 | GO:BP | 0.01633299 |
| tube closure | GO:0060606 | GO:BP | 0.01633299 |
| vesicle docking | GO:0048278 | GO:BP | 0.0164174 |
| protein folding | GO:0006457 | GO:BP | 0.01660791 |
| erythrocyte development | GO:0048821 | GO:BP | 0.01665349 |
| purine ribonucleoside monophosphate metabolic proce | GO:0009167 | GO:BP | 0.01665349 |
| positive regulation of pattern recognition receptor signa | GO:0062208 | GO:BP | 0.01665349 |
| endocytic recycling | GO:0032456 | GO:BP | 0.01667336 |
| localization within membrane | GO:0051668 | GO:BP | 0.01674234 |
| ceramide biosynthetic process | GO:0046513 | GO:BP | 0.01681066 |
| regulation of DNA binding | GO:0051101 | GO:BP | 0.0168993 |
| aminoglycan biosynthetic process | GO:0006023 | GO:BP | 0.01692591 |
| glucose import | GO:0046323 | GO:BP | 0.01692591 |
| protein N-linked glycosylation | GO:0006487 | GO:BP | 0.01692591 |
| cellular response to mechanical stimulus | GO:0071260 | GO:BP | 0.01692591 |
| positive regulation of developmental growth | GO:0048639 | GO:BP | 0.01726453 |
| regulation of secondary metabolic process | GO:0043455 | GO:BP | 0.01759192 |
| negative regulation of cell-substrate junction organizati | GO:0150118 | GO:BP | 0.01759192 |
| sno(s)RNA metabolic process | GO:0016074 | GO:BP | 0.01759192 |
| negative regulation of focal adhesion assembly | GO:0051895 | GO:BP | 0.01759192 |
| regulation of secondary metabolite biosynthetic proces | GO:1900376 | GO:BP | 0.01759192 |
| regulation of melanin biosynthetic process | GO:0048021 | GO:BP | 0.01759192 |
| regulation of DNA methylation | GO:0044030 | GO:BP | 0.01759192 |
| connective tissue development | GO:0061448 | GO:BP | 0.01766897 |
| microtubule anchoring | GO:0034453 | GO:BP | 0.01782691 |
| positive regulation of focal adhesion assembly | GO:0051894 | GO:BP | 0.01782691 |
| positive regulation of protein autophosphorylation | GO:0031954 | GO:BP | 0.01782691 |
| sphingolipid metabolic process | GO:0006665 | GO:BP | 0.01782691 |

| | | | |
|---|---|---|---|
| secondary palate development | GO:0062009 | GO:BP | 0.01782691 |
| stem cell division | GO:0017145 | GO:BP | 0.01804843 |
| negative regulation of signal transduction by p53 class r | GO:1901797 | GO:BP | 0.01804843 |
| positive regulation of endothelial cell migration | GO:0010595 | GO:BP | 0.0184162 |
| negative regulation of establishment of protein localizat | GO:1904950 | GO:BP | 0.01842971 |
| membrane lipid metabolic process | GO:0006643 | GO:BP | 0.01847819 |
| receptor recycling | GO:0001881 | GO:BP | 0.01850584 |
| positive regulation of inflammasome-mediated signalin | GO:0141087 | GO:BP | 0.01869261 |
| negative regulation of cyclin-dependent protein kinase a | GO:1904030 | GO:BP | 0.01869261 |
| peptidyl-lysine acetylation | GO:0018394 | GO:BP | 0.01869261 |
| P-body assembly | GO:0033962 | GO:BP | 0.01869261 |
| regulation of hemopoiesis | GO:1903706 | GO:BP | 0.01907289 |
| locomotion | GO:0040011 | GO:BP | 0.0192211 |
| regulation of axon extension | GO:0030516 | GO:BP | 0.01951922 |
| tetrapyrrole biosynthetic process | GO:0033014 | GO:BP | 0.01982457 |
| negative regulation of telomere maintenance | GO:0032205 | GO:BP | 0.01982457 |
| negative regulation of cellular response to insulin stimu | GO:1900077 | GO:BP | 0.01982457 |
| nucleotide-sugar metabolic process | GO:0009225 | GO:BP | 0.01982457 |
| porphyrin-containing compound biosynthetic process | GO:0006779 | GO:BP | 0.01982457 |
| morphogenesis of a branching epithelium | GO:0061138 | GO:BP | 0.02034472 |
| regulation of synapse organization | GO:0050807 | GO:BP | 0.02044984 |
| cytoplasmic microtubule organization | GO:0031122 | GO:BP | 0.02075347 |
| response to X-ray | GO:0010165 | GO:BP | 0.02081004 |
| regulation of post-transcriptional gene silencing | GO:0060147 | GO:BP | 0.02081004 |
| rescue of stalled ribosome | GO:0072344 | GO:BP | 0.02081004 |
| activation of protein kinase activity | GO:0032147 | GO:BP | 0.02088391 |
| type I interferon-mediated signaling pathway | GO:0060337 | GO:BP | 0.02088391 |
| negative regulation of transmembrane receptor protein | GO:0090101 | GO:BP | 0.02105718 |
| endothelial cell migration | GO:0043542 | GO:BP | 0.02112962 |
| regulation of membrane permeability | GO:0090559 | GO:BP | 0.02112962 |
| regulation of synapse structure or activity | GO:0050803 | GO:BP | 0.02115931 |
| somitogenesis | GO:0001756 | GO:BP | 0.02118046 |
| glycosaminoglycan biosynthetic process | GO:0006024 | GO:BP | 0.02118046 |
| neurotrophin signaling pathway | GO:0038179 | GO:BP | 0.02127865 |
| response to progesterone | GO:0032570 | GO:BP | 0.02127865 |
| 'de novo' protein folding | GO:0006458 | GO:BP | 0.02127865 |
| negative regulation of microtubule polymerization or de | GO:0031111 | GO:BP | 0.02127865 |
| regulation of JUN kinase activity | GO:0043506 | GO:BP | 0.02127865 |
| positive regulation of proteolysis | GO:0045862 | GO:BP | 0.02141636 |
| positive regulation of protein acetylation | GO:1901985 | GO:BP | 0.02164259 |
| microtubule anchoring at microtubule organizing center | GO:0072393 | GO:BP | 0.02164259 |
| regulation of nuclear cell cycle DNA replication | GO:0033262 | GO:BP | 0.02164259 |
| positive regulation of post-transcriptional gene silencin | GO:0060148 | GO:BP | 0.02164259 |
| inner cell mass cell proliferation | GO:0001833 | GO:BP | 0.02164259 |
| positive regulation of post-transcriptional gene silencin | GO:1900370 | GO:BP | 0.02164259 |
| negative regulation of substrate adhesion-dependent ce | GO:1900025 | GO:BP | 0.02164259 |
| positive regulation of miRNA-mediated gene silencing | GO:2000637 | GO:BP | 0.02164259 |
| intracellular mRNA localization | GO:0008298 | GO:BP | 0.02164259 |

| | | | |
|---|---|---|---|
| positive regulation of protein localization to endosome | GO:1905668 | GO:BP | 0.02164259 |
| interferon-mediated signaling pathway | GO:0140888 | GO:BP | 0.02193276 |
| tRNA aminoacylation for protein translation | GO:0006418 | GO:BP | 0.02252658 |
| pyrimidine-containing compound biosynthetic process | GO:0072528 | GO:BP | 0.02252658 |
| neuron projection extension | GO:1990138 | GO:BP | 0.02254111 |
| lipoprotein metabolic process | GO:0042157 | GO:BP | 0.02290818 |
| outflow tract septum morphogenesis | GO:0003148 | GO:BP | 0.02293018 |
| neuronal stem cell population maintenance | GO:0097150 | GO:BP | 0.02293018 |
| positive regulation of microtubule polymerization | GO:0031116 | GO:BP | 0.02324914 |
| cellular component disassembly involved in execution p | GO:0006921 | GO:BP | 0.02324914 |
| regulation of cellular senescence | GO:2000772 | GO:BP | 0.02360962 |
| ER-nucleus signaling pathway | GO:0006984 | GO:BP | 0.02360962 |
| response to hepatocyte growth factor | GO:0035728 | GO:BP | 0.0238033 |
| trophectodermal cell differentiation | GO:0001829 | GO:BP | 0.0238033 |
| regulation of vesicle size | GO:0097494 | GO:BP | 0.0238033 |
| plasma membrane tubulation | GO:0097320 | GO:BP | 0.0238033 |
| DNA replication checkpoint signaling | GO:0000076 | GO:BP | 0.0238033 |
| nucleosome disassembly | GO:0006337 | GO:BP | 0.0238033 |
| mitochondrial transcription | GO:0006390 | GO:BP | 0.0238033 |
| leukocyte differentiation | GO:0002521 | GO:BP | 0.02421426 |
| negative regulation of cysteine-type endopeptidase acti | GO:0043154 | GO:BP | 0.02445241 |
| striated muscle tissue development | GO:0014706 | GO:BP | 0.02445983 |
| response to starvation | GO:0042594 | GO:BP | 0.02459732 |
| response to type I interferon | GO:0034340 | GO:BP | 0.02477235 |
| protein complex oligomerization | GO:0051259 | GO:BP | 0.02501106 |
| regulation of viral life cycle | GO:1903900 | GO:BP | 0.02510072 |
| striated muscle hypertrophy | GO:0014897 | GO:BP | 0.02516246 |
| regulation of immune system process | GO:0002682 | GO:BP | 0.02524681 |
| positive regulation of neuron projection development | GO:0010976 | GO:BP | 0.02524681 |
| regulation of endothelial cell differentiation | GO:0045601 | GO:BP | 0.02544227 |
| cell migration involved in sprouting angiogenesis | GO:0002042 | GO:BP | 0.02566317 |
| protein maturation | GO:0051604 | GO:BP | 0.02570177 |
| negative regulation of cell cycle G1/S phase transition | GO:1902807 | GO:BP | 0.02581057 |
| negative regulation of protein ubiquitination | GO:0031397 | GO:BP | 0.02608442 |
| nuclear-transcribed mRNA catabolic process, exonucle | GO:0034427 | GO:BP | 0.02643987 |
| error-prone translesion synthesis | GO:0042276 | GO:BP | 0.02643987 |
| sulfur amino acid transport | GO:0000101 | GO:BP | 0.02643987 |
| maintenance of organelle location | GO:0051657 | GO:BP | 0.02643987 |
| regulation of protein localization to centrosome | GO:1904779 | GO:BP | 0.02643987 |
| ubiquitin-dependent protein catabolic process via the C | GO:0140627 | GO:BP | 0.02643987 |
| somatic stem cell population maintenance | GO:0035019 | GO:BP | 0.02643987 |
| regulation of skeletal muscle satellite cell proliferation | GO:0014842 | GO:BP | 0.02643987 |
| pyrimidine ribonucleoside monophosphate biosynthetic | GO:0009174 | GO:BP | 0.02643987 |
| branching morphogenesis of a nerve | GO:0048755 | GO:BP | 0.02643987 |
| regulation of double-strand break repair via nonhomolo | GO:2001032 | GO:BP | 0.02660723 |
| protein K48-linked deubiquitination | GO:0071108 | GO:BP | 0.02660723 |
| regulation of miRNA-mediated gene silencing | GO:0060964 | GO:BP | 0.02660723 |
| negative regulation of miRNA transcription | GO:1902894 | GO:BP | 0.02660723 |

| | | | |
|---|---|---|---|
| regulation of glial cell proliferation | GO:0060251 | GO:BP | 0.02725053 |
| nucleotide metabolic process | GO:0009117 | GO:BP | 0.02811892 |
| ossification | GO:0001503 | GO:BP | 0.02895809 |
| regulation of DNA damage response, signal transductio | GO:0043516 | GO:BP | 0.02895809 |
| cellular response to oxidative stress | GO:0034599 | GO:BP | 0.02965892 |
| regulation of ARF protein signal transduction | GO:0032012 | GO:BP | 0.02970544 |
| ARF protein signal transduction | GO:0032011 | GO:BP | 0.02970544 |
| nucleoside phosphate metabolic process | GO:0006753 | GO:BP | 0.03011391 |
| regulation of cyclin-dependent protein kinase activity | GO:1904029 | GO:BP | 0.03011749 |
| regulation of organ growth | GO:0046620 | GO:BP | 0.03011749 |
| positive regulation of translational initiation | GO:0045948 | GO:BP | 0.03011772 |
| lymph vessel development | GO:0001945 | GO:BP | 0.03011772 |
| regulation of apoptotic signaling pathway | GO:2001233 | GO:BP | 0.03019929 |
| regulation of endothelial cell development | GO:1901550 | GO:BP | 0.0303797 |
| intestinal epithelial cell development | GO:0060576 | GO:BP | 0.0303797 |
| nucleophagy | GO:0044804 | GO:BP | 0.0303797 |
| pyrimidine ribonucleoside monophosphate metabolic p | GO:0009173 | GO:BP | 0.0303797 |
| regulation of establishment of endothelial barrier | GO:1903140 | GO:BP | 0.0303797 |
| lymphangiogenesis | GO:0001946 | GO:BP | 0.0303797 |
| cellular response to hepatocyte growth factor stimulus | GO:0035729 | GO:BP | 0.0303797 |
| negative regulation of smooth muscle cell differentiatio | GO:0051151 | GO:BP | 0.0303797 |
| regulation of autophagosome maturation | GO:1901096 | GO:BP | 0.0303797 |
| Golgi to endosome transport | GO:0006895 | GO:BP | 0.0303797 |
| response to amino acid | GO:0043200 | GO:BP | 0.03043701 |
| female gamete generation | GO:0007292 | GO:BP | 0.03084428 |
| meiosis I cell cycle process | GO:0061982 | GO:BP | 0.03102535 |
| cytosol to Golgi apparatus transport | GO:0140820 | GO:BP | 0.03121964 |
| ER to Golgi ceramide transport | GO:0035621 | GO:BP | 0.03121964 |
| negative regulation of ribosome biogenesis | GO:0090071 | GO:BP | 0.03121964 |
| regulation of transmembrane receptor protein serine/th | GO:0090092 | GO:BP | 0.03121964 |
| regulation of mitochondrial DNA replication | GO:0090296 | GO:BP | 0.03121964 |
| negative regulation of excitatory postsynaptic potential | GO:0090394 | GO:BP | 0.03121964 |
| positive regulation by virus of viral protein levels in host | GO:0046726 | GO:BP | 0.03121964 |
| telomerase RNA stabilization | GO:0090669 | GO:BP | 0.03121964 |
| presynaptic membrane assembly | GO:0097105 | GO:BP | 0.03121964 |
| negative regulation of cellular response to oxidative stre | GO:1900408 | GO:BP | 0.03121964 |
| endonucleolytic cleavage in ITS1 to separate SSU-rRNA | GO:0000447 | GO:BP | 0.03121964 |
| negative regulation of cytokine-mediated signaling path | GO:0001960 | GO:BP | 0.03121964 |
| N-terminal peptidyl-lysine acetylation | GO:0018076 | GO:BP | 0.03121964 |
| negative regulation of protein kinase C signaling | GO:0090038 | GO:BP | 0.03121964 |
| mitotic nuclear membrane organization | GO:0101024 | GO:BP | 0.03121964 |
| Golgi vesicle budding | GO:0048194 | GO:BP | 0.03121964 |
| DNA endoreduplication | GO:0042023 | GO:BP | 0.03121964 |
| regulation of angiotensin-activated signaling pathway | GO:0110061 | GO:BP | 0.03121964 |
| rRNA 3'-end processing | GO:0031125 | GO:BP | 0.03121964 |
| GCN2-mediated signaling | GO:0140469 | GO:BP | 0.03121964 |
| constitutive heterochromatin formation | GO:0140719 | GO:BP | 0.03121964 |
| protein deneddylation | GO:0000338 | GO:BP | 0.03121964 |

| | | | |
|---|---|---|---|
| protein myristoylation | GO:0018377 | GO:BP | 0.03121964 |
| viral penetration into host nucleus | GO:0075732 | GO:BP | 0.03121964 |
| minus-end-directed organelle transport along microtub | GO:0072385 | GO:BP | 0.03121964 |
| positive regulation of stress granule assembly | GO:0062029 | GO:BP | 0.03121964 |
| mitotic nuclear membrane reassembly | GO:0007084 | GO:BP | 0.03121964 |
| spindle assembly involved in female meiosis I | GO:0007057 | GO:BP | 0.03121964 |
| regulation of vitamin D receptor signaling pathway | GO:0070562 | GO:BP | 0.03121964 |
| negative regulation of protein exit from endoplasmic ret | GO:0070862 | GO:BP | 0.03121964 |
| vitellogenesis | GO:0007296 | GO:BP | 0.03121964 |
| RISC complex assembly | GO:0070922 | GO:BP | 0.03121964 |
| regulation of ER to Golgi vesicle-mediated transport | GO:0060628 | GO:BP | 0.03121964 |
| protein dealkylation | GO:0008214 | GO:BP | 0.03121964 |
| meiotic cell cycle checkpoint signaling | GO:0033313 | GO:BP | 0.03121964 |
| demethylation | GO:0070988 | GO:BP | 0.03121964 |
| phenylalanine transport | GO:0015823 | GO:BP | 0.03121964 |
| regulation of transcription by glucose | GO:0046015 | GO:BP | 0.03121964 |
| polyadenylation-dependent mRNA catabolic process | GO:0071047 | GO:BP | 0.03121964 |
| ribonucleoprotein complex localization | GO:0071166 | GO:BP | 0.03121964 |
| N-terminal protein myristoylation | GO:0006499 | GO:BP | 0.03121964 |
| CMP biosynthetic process | GO:0009224 | GO:BP | 0.03121964 |
| manganese ion transmembrane transport | GO:0071421 | GO:BP | 0.03121964 |
| snRNA export from nucleus | GO:0006408 | GO:BP | 0.03121964 |
| termination of RNA polymerase I transcription | GO:0006363 | GO:BP | 0.03121964 |
| establishment of centrosome localization | GO:0051660 | GO:BP | 0.03121964 |
| lagging strand elongation | GO:0006273 | GO:BP | 0.03121964 |
| leading strand elongation | GO:0006272 | GO:BP | 0.03121964 |
| oxidative single-stranded RNA demethylation | GO:0035553 | GO:BP | 0.03121964 |
| nuclear polyadenylation-dependent mRNA catabolic pr | GO:0071042 | GO:BP | 0.03121964 |
| protein demethylation | GO:0006482 | GO:BP | 0.03121964 |
| tRNA transcription by RNA polymerase III | GO:0042797 | GO:BP | 0.03121964 |
| negative regulation of DNA duplex unwinding | GO:1905463 | GO:BP | 0.03121964 |
| negative regulation of wound healing, spreading of epid | GO:1903690 | GO:BP | 0.03121964 |
| positive regulation of endoplasmic reticulum tubular ne | GO:1903373 | GO:BP | 0.03121964 |
| positive regulation of rRNA processing | GO:2000234 | GO:BP | 0.03121964 |
| regulation of protein localization to early endosome | GO:1902965 | GO:BP | 0.03121964 |
| positive regulation of protein localization to early endos | GO:1902966 | GO:BP | 0.03121964 |
| positive regulation of clathrin-dependent endocytosis | GO:2000370 | GO:BP | 0.03121964 |
| regulation of cardiac muscle hypertrophy | GO:0010611 | GO:BP | 0.03154952 |
| artery morphogenesis | GO:0048844 | GO:BP | 0.03178903 |
| regulation of cilium assembly | GO:1902017 | GO:BP | 0.03178903 |
| positive regulation of establishment of protein localizati | GO:1903749 | GO:BP | 0.03179373 |
| positive regulation of telomerase activity | GO:0051973 | GO:BP | 0.03179373 |
| DNA methylation-dependent heterochromatin formatio | GO:0006346 | GO:BP | 0.03179373 |
| positive regulation of protein targeting to mitochondrior | GO:1903955 | GO:BP | 0.03179373 |
| ruffle organization | GO:0031529 | GO:BP | 0.03182567 |
| negative regulation of cysteine-type endopeptidase acti | GO:2000117 | GO:BP | 0.03182567 |
| positive regulation of cytokine-mediated signaling pathv | GO:0001961 | GO:BP | 0.03197791 |
| regulation of interferon-beta production | GO:0032648 | GO:BP | 0.03197791 |

| | | | |
|---|---|---|---|
| interferon-beta production | GO:0032608 | GO:BP | 0.03197791 |
| neutral amino acid transport | GO:0015804 | GO:BP | 0.03197791 |
| regulation of dendritic spine development | GO:0060998 | GO:BP | 0.03197791 |
| regulation of dendrite development | GO:0050773 | GO:BP | 0.03249745 |
| kidney development | GO:0001822 | GO:BP | 0.0327066 |
| regulation of cell adhesion | GO:0030155 | GO:BP | 0.03321702 |
| regulation of glucose metabolic process | GO:0010906 | GO:BP | 0.03336288 |
| regulation of protein sumoylation | GO:0033233 | GO:BP | 0.03343446 |
| mitotic recombination | GO:0006312 | GO:BP | 0.03343446 |
| regulation of Rac protein signal transduction | GO:0035020 | GO:BP | 0.03343446 |
| nuclear migration | GO:0007097 | GO:BP | 0.03343446 |
| nucleotide-binding domain, leucine rich repeat containi | GO:0035872 | GO:BP | 0.03343446 |
| respiratory system development | GO:0060541 | GO:BP | 0.03357468 |
| negative regulation of insulin receptor signaling pathwa | GO:0046627 | GO:BP | 0.03388656 |
| ribosome disassembly | GO:0032790 | GO:BP | 0.03388656 |
| 'de novo' post-translational protein folding | GO:0051084 | GO:BP | 0.03388656 |
| positive regulation of microtubule polymerization or dep | GO:0031112 | GO:BP | 0.03388656 |
| positive regulation of lamellipodium organization | GO:1902745 | GO:BP | 0.03388656 |
| regulation of smooth muscle cell differentiation | GO:0051150 | GO:BP | 0.03388656 |
| peptidyl-cysteine modification | GO:0018198 | GO:BP | 0.03388656 |
| negative regulation of protein transport | GO:0051224 | GO:BP | 0.03433062 |
| morphogenesis of embryonic epithelium | GO:0016331 | GO:BP | 0.03446454 |
| embryonic organ morphogenesis | GO:0048562 | GO:BP | 0.03477694 |
| peptidyl-histidine modification | GO:0018202 | GO:BP | 0.03498484 |
| vesicle uncoating | GO:0072319 | GO:BP | 0.03498484 |
| regulation of stress granule assembly | GO:0062028 | GO:BP | 0.03498484 |
| pyrimidine nucleobase biosynthetic process | GO:0019856 | GO:BP | 0.03498484 |
| DNA strand resection involved in replication fork proces | GO:0110025 | GO:BP | 0.03498484 |
| positive regulation of spindle assembly | GO:1905832 | GO:BP | 0.03498484 |
| polyadenylation-dependent snoRNA 3'-end processing | GO:0071051 | GO:BP | 0.03498484 |
| trans-synaptic signaling by endocannabinoid | GO:0099542 | GO:BP | 0.03498484 |
| positive regulation of protein K63-linked ubiquitination | GO:1902523 | GO:BP | 0.03498484 |
| negative regulation of erythrocyte differentiation | GO:0045647 | GO:BP | 0.03498484 |
| trans-synaptic signaling by lipid | GO:0099541 | GO:BP | 0.03498484 |
| negative regulation of response to oxidative stress | GO:1902883 | GO:BP | 0.03498484 |
| meiotic cytokinesis | GO:0033206 | GO:BP | 0.03498484 |
| positive regulation of translation in response to stress | GO:0032056 | GO:BP | 0.03498484 |
| activation of immune response | GO:0002253 | GO:BP | 0.03533329 |
| muscle hypertrophy | GO:0014896 | GO:BP | 0.03541395 |
| negative regulation of protein localization to nucleus | GO:1900181 | GO:BP | 0.03553819 |
| DNA dealkylation | GO:0035510 | GO:BP | 0.03553819 |
| mammary gland development | GO:0030879 | GO:BP | 0.0356046 |
| gastrulation | GO:0007369 | GO:BP | 0.03617033 |
| cerebellum development | GO:0021549 | GO:BP | 0.03620675 |
| Mo-molybdopterin cofactor biosynthetic process | GO:0006777 | GO:BP | 0.03641429 |
| detection of virus | GO:0009597 | GO:BP | 0.03641429 |
| pyrimidine nucleoside diphosphate biosynthetic proces | GO:0009139 | GO:BP | 0.03641429 |
| R-loop processing | GO:0062176 | GO:BP | 0.03641429 |

| | | | |
|---|---|---|---|
| regulation of ubiquitin protein ligase activity | GO:1904666 | GO:BP | 0.03641429 |
| positive regulation of protein localization to cell cortex | GO:1904778 | GO:BP | 0.03641429 |
| regulation of DNA duplex unwinding | GO:1905462 | GO:BP | 0.03641429 |
| maintenance of centrosome location | GO:0051661 | GO:BP | 0.03641429 |
| ribosome-associated ubiquitin-dependent protein catal | GO:1990116 | GO:BP | 0.03641429 |
| abscission | GO:0009838 | GO:BP | 0.03641429 |
| positive regulation of cytoplasmic mRNA processing bo | GO:0010606 | GO:BP | 0.03641429 |
| peptidyl-lysine methylation | GO:0018022 | GO:BP | 0.03641429 |
| prosthetic group metabolic process | GO:0051189 | GO:BP | 0.03641429 |
| Mo-molybdopterin cofactor metabolic process | GO:0019720 | GO:BP | 0.03641429 |
| response to L-phenylalanine derivative | GO:1904386 | GO:BP | 0.03641429 |
| ubiquinone metabolic process | GO:0006743 | GO:BP | 0.03641429 |
| sphingomyelin metabolic process | GO:0006684 | GO:BP | 0.03641429 |
| regulation of translation at postsynapse | GO:0140245 | GO:BP | 0.03641429 |
| regulation of focal adhesion disassembly | GO:0120182 | GO:BP | 0.03641429 |
| positive regulation of focal adhesion disassembly | GO:0120183 | GO:BP | 0.03641429 |
| protein mannosylation | GO:0035268 | GO:BP | 0.03641429 |
| mitochondrial RNA 3'-end processing | GO:0000965 | GO:BP | 0.03641429 |
| apoptotic process involved in heart morphogenesis | GO:0003278 | GO:BP | 0.03641429 |
| regulation of translation at synapse | GO:0140243 | GO:BP | 0.03641429 |
| DNA replication-dependent chromatin assembly | GO:0006335 | GO:BP | 0.03641429 |
| molybdopterin cofactor biosynthetic process | GO:0032324 | GO:BP | 0.03641429 |
| positive regulation of nucleobase-containing compounc | GO:0032241 | GO:BP | 0.03641429 |
| regulation of protein monoubiquitination | GO:1902525 | GO:BP | 0.03641429 |
| molybdopterin cofactor metabolic process | GO:0043545 | GO:BP | 0.03641429 |
| 5S class rRNA transcription by RNA polymerase III | GO:0042791 | GO:BP | 0.03641429 |
| regulation of nuclease activity | GO:0032069 | GO:BP | 0.03641429 |
| N-terminal protein lipidation | GO:0006498 | GO:BP | 0.03641429 |
| homeostatic process | GO:0042592 | GO:BP | 0.03641429 |
| DNA strand invasion | GO:0042148 | GO:BP | 0.03641429 |
| meiotic cell cycle phase transition | GO:0044771 | GO:BP | 0.03651537 |
| t-circle formation | GO:0090656 | GO:BP | 0.03651537 |
| regulation of protein localization to endosome | GO:1905666 | GO:BP | 0.03651537 |
| formation of extrachromosomal circular DNA | GO:0001325 | GO:BP | 0.03651537 |
| pyrimidine nucleoside monophosphate biosynthetic pro | GO:0009130 | GO:BP | 0.03651537 |
| telomere maintenance via telomere trimming | GO:0090737 | GO:BP | 0.03651537 |
| centriole-centriole cohesion | GO:0010457 | GO:BP | 0.03651537 |
| GMP metabolic process | GO:0046037 | GO:BP | 0.03653592 |
| lung cell differentiation | GO:0060479 | GO:BP | 0.03653592 |
| lung epithelial cell differentiation | GO:0060487 | GO:BP | 0.03653592 |
| positive regulation of peptidyl-threonine phosphorylatio | GO:0010800 | GO:BP | 0.03653592 |
| regulation of glucose transmembrane transport | GO:0010827 | GO:BP | 0.03742346 |
| positive regulation of protein localization to cell periphe | GO:1904377 | GO:BP | 0.03742346 |
| hair cycle process | GO:0022405 | GO:BP | 0.0375435 |
| regulation of epithelial to mesenchymal transition | GO:0010717 | GO:BP | 0.0375435 |
| molting cycle process | GO:0022404 | GO:BP | 0.0375435 |
| metencephalon development | GO:0022037 | GO:BP | 0.03807582 |
| regulation of meiotic cell cycle | GO:0051445 | GO:BP | 0.03845774 |

| | | | |
|---|---|---|---|
| antiviral innate immune response | GO:0140374 | GO:BP | 0.03880508 |
| androgen receptor signaling pathway | GO:0030521 | GO:BP | 0.03880508 |
| positive regulation of response to cytokine stimulus | GO:0060760 | GO:BP | 0.03882453 |
| cellular response to heat | GO:0034605 | GO:BP | 0.03911666 |
| response to amino acid starvation | GO:1990928 | GO:BP | 0.03912376 |
| adherens junction organization | GO:0034332 | GO:BP | 0.03918618 |
| cellular component biogenesis | GO:0044085 | GO:BP | 0.03918618 |
| regulation of vacuole organization | GO:0044088 | GO:BP | 0.03918618 |
| water-soluble vitamin metabolic process | GO:0006767 | GO:BP | 0.03918618 |
| vascular endothelial growth factor receptor signaling pa | GO:0048010 | GO:BP | 0.03918618 |
| chaperone cofactor-dependent protein refolding | GO:0051085 | GO:BP | 0.03981775 |
| negative regulation of DNA binding | GO:0043392 | GO:BP | 0.03981775 |
| viral RNA genome replication | GO:0039694 | GO:BP | 0.03981775 |
| establishment of mitochondrion localization | GO:0051654 | GO:BP | 0.03981775 |
| negative regulation of cell junction assembly | GO:1901889 | GO:BP | 0.03981775 |
| respiratory chain complex IV assembly | GO:0008535 | GO:BP | 0.03981775 |
| limbic system development | GO:0021761 | GO:BP | 0.04001309 |
| fibroblast proliferation | GO:0048144 | GO:BP | 0.04045793 |
| cellular response to unfolded protein | GO:0034620 | GO:BP | 0.04056666 |
| mucopolysaccharide metabolic process | GO:1903510 | GO:BP | 0.04056666 |
| sex differentiation | GO:0007548 | GO:BP | 0.04122568 |
| cardiocyte differentiation | GO:0035051 | GO:BP | 0.04134794 |
| positive regulation of cell population proliferation | GO:0008284 | GO:BP | 0.04137326 |
| B cell activation involved in immune response | GO:0002312 | GO:BP | 0.04162252 |
| negative regulation of nucleocytoplasmic transport | GO:0046823 | GO:BP | 0.04212131 |
| renal system development | GO:0072001 | GO:BP | 0.04212131 |
| pulmonary valve development | GO:0003177 | GO:BP | 0.04212131 |
| tubulin deacetylation | GO:0090042 | GO:BP | 0.04212131 |
| regulation of protein acetylation | GO:1901983 | GO:BP | 0.04212131 |
| negative regulation of intrinsic apoptotic signaling pathv | GO:1902254 | GO:BP | 0.04212131 |
| hematopoietic stem cell proliferation | GO:0071425 | GO:BP | 0.04254013 |
| cardiac cell development | GO:0055006 | GO:BP | 0.04256831 |
| negative regulation of transport | GO:0051051 | GO:BP | 0.04333855 |
| ribose phosphate metabolic process | GO:0019693 | GO:BP | 0.04333855 |
| carboxylic acid transmembrane transport | GO:1905039 | GO:BP | 0.04379606 |
| substrate adhesion-dependent cell spreading | GO:0034446 | GO:BP | 0.04440099 |
| positive regulation of telomere maintenance via telome | GO:1904358 | GO:BP | 0.04482844 |
| prenylation | GO:0097354 | GO:BP | 0.04627124 |
| positive regulation of heterochromatin organization | GO:0120263 | GO:BP | 0.04627124 |
| adherens junction assembly | GO:0034333 | GO:BP | 0.04627124 |
| protein localization to nuclear envelope | GO:0090435 | GO:BP | 0.04627124 |
| somatic diversification of immune receptors via somati | GO:0002566 | GO:BP | 0.04627124 |
| pigment granule organization | GO:0048753 | GO:BP | 0.04627124 |
| positive regulation of dendritic spine development | GO:0060999 | GO:BP | 0.04627124 |
| protein prenylation | GO:0018342 | GO:BP | 0.04627124 |
| positive regulation of heterochromatin formation | GO:0031453 | GO:BP | 0.04627124 |
| ubiquitin recycling | GO:0010992 | GO:BP | 0.04627124 |
| regulation of ERAD pathway | GO:1904292 | GO:BP | 0.04627124 |

| | | | |
|---|---|---|---|
| regulation of retrograde protein transport, ER to cytosol | GO:1904152 | GO:BP | 0.04627124 |
| tRNA splicing, via endonucleolytic cleavage and ligation | GO:0006388 | GO:BP | 0.04627124 |
| regulation of protein autophosphorylation | GO:0031952 | GO:BP | 0.04627124 |
| cellular response to leucine | GO:0071233 | GO:BP | 0.04627124 |
| phospholipid dephosphorylation | GO:0046839 | GO:BP | 0.04627124 |
| regulation of hematopoietic progenitor cell differentiatio | GO:1901532 | GO:BP | 0.04627124 |
| protein localization to chromosome, centromeric region | GO:0071459 | GO:BP | 0.04627124 |
| positive regulation of gluconeogenesis | GO:0045722 | GO:BP | 0.04634951 |
| regulation of carbohydrate biosynthetic process | GO:0043255 | GO:BP | 0.04687966 |
| tube formation | GO:0035148 | GO:BP | 0.0471018 |
| negative regulation of stem cell differentiation | GO:2000737 | GO:BP | 0.04725797 |
| protein palmitoylation | GO:0018345 | GO:BP | 0.04725797 |
| negative regulation of type I interferon production | GO:0032480 | GO:BP | 0.04748558 |
| lactation | GO:0007595 | GO:BP | 0.04748558 |
| negative regulation of ERK1 and ERK2 cascade | GO:0070373 | GO:BP | 0.04757964 |
| negative regulation of proteolysis involved in protein cat | GO:1903051 | GO:BP | 0.04757964 |
| axon extension | GO:0048675 | GO:BP | 0.04777995 |
| regulation of MAP kinase activity | GO:0043405 | GO:BP | 0.04777995 |
| positive regulation of NF-kappaB transcription factor ac | GO:0051092 | GO:BP | 0.04777995 |
| endolysosomal toll-like receptor signaling pathway | GO:0140894 | GO:BP | 0.04830393 |
| embryonic digit morphogenesis | GO:0042733 | GO:BP | 0.04866038 |
| cerebral cortex cell migration | GO:0021795 | GO:BP | 0.04883626 |
| regulation of gluconeogenesis | GO:0006111 | GO:BP | 0.04883626 |
| porphyrin-containing compound metabolic process | GO:0006778 | GO:BP | 0.04914759 |
| negative regulation of protein modification by small pro | GO:1903321 | GO:BP | 0.04947961 |
| organic acid transmembrane transport | GO:1903825 | GO:BP | 0.04955285 |
| branching morphogenesis of an epithelial tube | GO:0048754 | GO:BP | 0.04955285 |
| nucleoplasm | GO:0005654 | GO:CC | 2.6756978114 |
| cytoplasm | GO:0005737 | GO:CC | 1.05E-240 |
| cytosol | GO:0005829 | GO:CC | 2.63E-195 |
| nuclear lumen | GO:0031981 | GO:CC | 7.17E-185 |
| intracellular anatomical structure | GO:0005622 | GO:CC | 1.09E-167 |
| membrane-enclosed lumen | GO:0031974 | GO:CC | 6.52E-157 |
| intracellular organelle lumen | GO:0070013 | GO:CC | 6.52E-157 |
| organelle lumen | GO:0043233 | GO:CC | 6.52E-157 |
| intracellular membrane-bounded organelle | GO:0043231 | GO:CC | 6.66E-152 |
| intracellular organelle | GO:0043229 | GO:CC | 8.41E-138 |
| membrane-bounded organelle | GO:0043227 | GO:CC | 1.50E-130 |
| organelle | GO:0043226 | GO:CC | 8.94E-124 |
| catalytic complex | GO:1902494 | GO:CC | 1.07E-87 |
| chromosome | GO:0005694 | GO:CC | 1.22E-86 |
| nucleus | GO:0005634 | GO:CC | 9.58E-77 |
| transferase complex | GO:1990234 | GO:CC | 2.87E-68 |
| intracellular protein-containing complex | GO:0140535 | GO:CC | 2.87E-68 |
| nuclear body | GO:0016604 | GO:CC | 3.61E-65 |
| intracellular non-membrane-bounded organelle | GO:0043232 | GO:CC | 2.32E-54 |
| non-membrane-bounded organelle | GO:0043228 | GO:CC | 2.54E-54 |
| organelle membrane | GO:0031090 | GO:CC | 7.11E-49 |

| | | | |
|---|---|---|---|
| chromosomal region | GO:0098687 | GO:CC | 2.04E-45 |
| microtubule cytoskeleton | GO:0015630 | GO:CC | 4.61E-44 |
| nuclear speck | GO:0016607 | GO:CC | 2.35E-39 |
| chromosome, centromeric region | GO:0000775 | GO:CC | 1.06E-38 |
| centrosome | GO:0005813 | GO:CC | 4.37E-38 |
| protein-DNA complex | GO:0032993 | GO:CC | 1.30E-35 |
| bounding membrane of organelle | GO:0098588 | GO:CC | 3.42E-33 |
| chromatin | GO:0000785 | GO:CC | 4.59E-33 |
| microtubule organizing center | GO:0005815 | GO:CC | 6.99E-33 |
| cytoskeleton | GO:0005856 | GO:CC | 7.97E-33 |
| endomembrane system | GO:0012505 | GO:CC | 9.68E-33 |
| condensed chromosome, centromeric region | GO:0000779 | GO:CC | 1.55E-32 |
| spindle | GO:0005819 | GO:CC | 1.74E-32 |
| condensed chromosome | GO:0000793 | GO:CC | 4.82E-32 |
| cellular anatomical entity | GO:0110165 | GO:CC | 1.13E-29 |
| kinetochore | GO:0000776 | GO:CC | 1.94E-29 |
| envelope | GO:0031975 | GO:CC | 1.35E-28 |
| organelle envelope | GO:0031967 | GO:CC | 1.35E-28 |
| Golgi apparatus | GO:0005794 | GO:CC | 8.91E-27 |
| ubiquitin ligase complex | GO:0000151 | GO:CC | 3.68E-25 |
| mitochondrion | GO:0005739 | GO:CC | 3.96E-24 |
| spliceosomal complex | GO:0005681 | GO:CC | 2.54E-22 |
| nuclear envelope | GO:0005635 | GO:CC | 3.42E-22 |
| Golgi membrane | GO:0000139 | GO:CC | 7.82E-21 |
| transcription regulator complex | GO:0005667 | GO:CC | 1.75E-20 |
| nuclear chromosome | GO:0000228 | GO:CC | 1.05E-19 |
| spindle pole | GO:0000922 | GO:CC | 1.31E-18 |
| midbody | GO:0030496 | GO:CC | 3.21E-18 |
| fibrillar center | GO:0001650 | GO:CC | 3.89E-18 |
| nuclear membrane | GO:0031965 | GO:CC | 6.35E-18 |
| nuclear periphery | GO:0034399 | GO:CC | 2.44E-17 |
| mitotic spindle | GO:0072686 | GO:CC | 9.65E-17 |
| endosome | GO:0005768 | GO:CC | 1.51E-16 |
| transferase complex, transferring phosphorus-containi | GO:0061695 | GO:CC | 1.84E-16 |
| preribosome | GO:0030684 | GO:CC | 3.32E-16 |
| methyltransferase complex | GO:0034708 | GO:CC | 5.71E-16 |
| RNA polymerase II transcription regulator complex | GO:0090575 | GO:CC | 8.93E-16 |
| organelle subcompartment | GO:0031984 | GO:CC | 2.80E-15 |
| cell leading edge | GO:0031252 | GO:CC | 4.73E-15 |
| acetyltransferase complex | GO:1902493 | GO:CC | 6.12E-15 |
| protein acetyltransferase complex | GO:0031248 | GO:CC | 1.52E-14 |
| nuclear matrix | GO:0016363 | GO:CC | 1.68E-14 |
| membrane | GO:0016020 | GO:CC | 2.26E-14 |
| ruffle | GO:0001726 | GO:CC | 3.50E-14 |
| early endosome | GO:0005769 | GO:CC | 3.94E-14 |
| ribonucleoprotein granule | GO:0035770 | GO:CC | 5.61E-14 |
| protein-containing complex | GO:0032991 | GO:CC | 9.82E-14 |
| perinuclear region of cytoplasm | GO:0048471 | GO:CC | 1.40E-13 |

| | | | |
|---|---|---|---|
| U2-type spliceosomal complex | GO:0005684 | GO:CC | 9.88E-13 |
| supramolecular complex | GO:0099080 | GO:CC | 1.43E-12 |
| vesicle membrane | GO:0012506 | GO:CC | 3.35E-12 |
| site of DNA damage | GO:0090734 | GO:CC | 3.67E-12 |
| histone acetyltransferase complex | GO:0000123 | GO:CC | 4.58E-12 |
| heterochromatin | GO:0000792 | GO:CC | 5.33E-12 |
| nuclear pore | GO:0005643 | GO:CC | 5.42E-12 |
| SWI/SNF superfamily-type complex | GO:0070603 | GO:CC | 6.76E-12 |
| endosome membrane | GO:0010008 | GO:CC | 6.97E-12 |
| cytoplasmic vesicle membrane | GO:0030659 | GO:CC | 7.18E-12 |
| cell junction | GO:0030054 | GO:CC | 7.33E-12 |
| cytoplasmic ribonucleoprotein granule | GO:0036464 | GO:CC | 1.46E-11 |
| replication fork | GO:0005657 | GO:CC | 2.75E-11 |
| histone methyltransferase complex | GO:0035097 | GO:CC | 3.29E-11 |
| microtubule | GO:0005874 | GO:CC | 1.93E-10 |
| intracellular vesicle | GO:0097708 | GO:CC | 2.22E-10 |
| Golgi apparatus subcompartment | GO:0098791 | GO:CC | 3.21E-10 |
| cytoplasmic stress granule | GO:0010494 | GO:CC | 3.48E-10 |
| cytoplasmic vesicle | GO:0031410 | GO:CC | 3.57E-10 |
| lamellipodium | GO:0030027 | GO:CC | 7.22E-10 |
| catalytic step 2 spliceosome | GO:0071013 | GO:CC | 1.00E-09 |
| trans-Golgi network | GO:0005802 | GO:CC | 1.25E-09 |
| small-subunit processome | GO:0032040 | GO:CC | 1.56E-09 |
| mitochondrial envelope | GO:0005740 | GO:CC | 1.63E-09 |
| spindle microtubule | GO:0005876 | GO:CC | 2.05E-09 |
| spindle midzone | GO:0051233 | GO:CC | 4.37E-09 |
| PML body | GO:0016605 | GO:CC | 4.65E-09 |
| site of double-strand break | GO:0035861 | GO:CC | 4.90E-09 |
| chromosome, telomeric region | GO:0000781 | GO:CC | 1.13E-08 |
| mitotic spindle pole | GO:0097431 | GO:CC | 1.20E-08 |
| mitochondrial matrix | GO:0005759 | GO:CC | 1.38E-08 |
| anchoring junction | GO:0070161 | GO:CC | 1.73E-08 |
| mitochondrial membrane | GO:0031966 | GO:CC | 1.91E-08 |
| nuclear DNA-directed RNA polymerase complex | GO:0055029 | GO:CC | 2.11E-08 |
| protein kinase complex | GO:1902911 | GO:CC | 2.11E-08 |
| exoribonuclease complex | GO:1905354 | GO:CC | 2.34E-08 |
| RNA polymerase complex | GO:0030880 | GO:CC | 3.26E-08 |
| nuclear outer membrane-endoplasmic reticulum meml | GO:0042175 | GO:CC | 4.59E-08 |
| cullin-RING ubiquitin ligase complex | GO:0031461 | GO:CC | 4.89E-08 |
| dendrite | GO:0030425 | GO:CC | 6.46E-08 |
| endoplasmic reticulum subcompartment | GO:0098827 | GO:CC | 6.74E-08 |
| actin cytoskeleton | GO:0015629 | GO:CC | 6.76E-08 |
| exosome (RNase complex) | GO:0000178 | GO:CC | 7.21E-08 |
| endoplasmic reticulum membrane | GO:0005789 | GO:CC | 7.26E-08 |
| DNA-directed RNA polymerase complex | GO:0000428 | GO:CC | 8.05E-08 |
| dendritic tree | GO:0097447 | GO:CC | 8.37E-08 |
| PcG protein complex | GO:0031519 | GO:CC | 9.85E-08 |
| histone deacetylase complex | GO:0000118 | GO:CC | 1.75E-07 |

| | | | |
|---|---|---|---|
| recycling endosome | GO:0055037 | GO:CC | 2.21E-07 |
| mitochondrial outer membrane | GO:0005741 | GO:CC | 2.44E-07 |
| trans-Golgi network membrane | GO:0032588 | GO:CC | 4.15E-07 |
| vacuolar membrane | GO:0005774 | GO:CC | 5.53E-07 |
| cell cortex | GO:0005938 | GO:CC | 5.71E-07 |
| focal adhesion | GO:0005925 | GO:CC | 6.24E-07 |
| ATPase complex | GO:1904949 | GO:CC | 7.21E-07 |
| serine/threonine protein kinase complex | GO:1902554 | GO:CC | 7.57E-07 |
| 90S preribosome | GO:0030686 | GO:CC | 7.96E-07 |
| kinesin complex | GO:0005871 | GO:CC | 9.79E-07 |
| early endosome membrane | GO:0031901 | GO:CC | 1.01E-06 |
| somatodendritic compartment | GO:0036477 | GO:CC | 1.08E-06 |
| cell-substrate junction | GO:0030055 | GO:CC | 1.11E-06 |
| organelle outer membrane | GO:0031968 | GO:CC | 1.56E-06 |
| P-body | GO:0000932 | GO:CC | 1.76E-06 |
| exon-exon junction complex | GO:0035145 | GO:CC | 2.06E-06 |
| nuclear ubiquitin ligase complex | GO:0000152 | GO:CC | 2.28E-06 |
| outer membrane | GO:0019867 | GO:CC | 2.38E-06 |
| vesicle | GO:0031982 | GO:CC | 3.48E-06 |
| cytoplasmic exosome (RNase complex) | GO:0000177 | GO:CC | 4.15E-06 |
| nuclear exosome (RNase complex) | GO:0000176 | GO:CC | 4.15E-06 |
| MLL1/2 complex | GO:0044665 | GO:CC | 4.30E-06 |
| sex chromosome | GO:0000803 | GO:CC | 4.92E-06 |
| SAGA-type complex | GO:0070461 | GO:CC | 5.31E-06 |
| transcription elongation factor complex | GO:0008023 | GO:CC | 5.89E-06 |
| lysosomal membrane | GO:0005765 | GO:CC | 5.94E-06 |
| lytic vacuole membrane | GO:0098852 | GO:CC | 5.94E-06 |
| synapse | GO:0045202 | GO:CC | 6.89E-06 |
| precatalytic spliceosome | GO:0071011 | GO:CC | 1.02E-05 |
| mediator complex | GO:0016592 | GO:CC | 1.03E-05 |
| MLL1 complex | GO:0071339 | GO:CC | 1.09E-05 |
| U2-type precatalytic spliceosome | GO:0071005 | GO:CC | 1.30E-05 |
| microtubule end | GO:1990752 | GO:CC | 1.57E-05 |
| outer kinetochore | GO:0000940 | GO:CC | 1.84E-05 |
| microtubule associated complex | GO:0005875 | GO:CC | 1.92E-05 |
| organelle inner membrane | GO:0019866 | GO:CC | 2.07E-05 |
| axon cytoplasm | GO:1904115 | GO:CC | 2.50E-05 |
| adherens junction | GO:0005912 | GO:CC | 3.06E-05 |
| centriole | GO:0005814 | GO:CC | 3.17E-05 |
| INO80-type complex | GO:0097346 | GO:CC | 3.28E-05 |
| neuron projection cytoplasm | GO:0120111 | GO:CC | 3.36E-05 |
| organellar ribosome | GO:0000313 | GO:CC | 3.36E-05 |
| mitochondrial ribosome | GO:0005761 | GO:CC | 3.36E-05 |
| mitotic spindle midzone | GO:1990023 | GO:CC | 4.11E-05 |
| ISWI-type complex | GO:0031010 | GO:CC | 4.11E-05 |
| RSC-type complex | GO:0016586 | GO:CC | 4.89E-05 |
| endoplasmic reticulum | GO:0005783 | GO:CC | 5.13E-05 |
| Cul4-RING E3 ubiquitin ligase complex | GO:0080008 | GO:CC | 5.47E-05 |

| | | | |
|---|---|---|---|
| pericentric heterochromatin | GO:0005721 | GO:CC | 5.55E-05 |
| recycling endosome membrane | GO:0055038 | GO:CC | 6.30E-05 |
| nuclear replication fork | GO:0043596 | GO:CC | 7.64E-05 |
| ruffle membrane | GO:0032587 | GO:CC | 7.83E-05 |
| MCM complex | GO:0042555 | GO:CC | 9.47E-05 |
| growth cone | GO:0030426 | GO:CC | 9.99E-05 |
| cell projection | GO:0042995 | GO:CC | 0.00010342 |
| cell division site | GO:0032153 | GO:CC | 0.00010621 |
| cell-cell junction | GO:0005911 | GO:CC | 0.00012685 |
| endoplasmic reticulum tubular network | GO:0071782 | GO:CC | 0.00013399 |
| preribosome, large subunit precursor | GO:0030687 | GO:CC | 0.00013932 |
| Set1C/COMPASS complex | GO:0048188 | GO:CC | 0.00014615 |
| site of polarized growth | GO:0030427 | GO:CC | 0.00015024 |
| condensin complex | GO:0000796 | GO:CC | 0.00016226 |
| RNA polymerase II, holoenzyme | GO:0016591 | GO:CC | 0.00021276 |
| H4 histone acetyltransferase complex | GO:1902562 | GO:CC | 0.00021463 |
| endoplasmic reticulum-Golgi intermediate compartmen | GO:0005793 | GO:CC | 0.00021472 |
| cyclin-dependent protein kinase holoenzyme complex | GO:0000307 | GO:CC | 0.00024916 |
| plasma membrane bounded cell projection | GO:0120025 | GO:CC | 0.00025292 |
| mitochondrial nucleoid | GO:0042645 | GO:CC | 0.00026094 |
| nucleoid | GO:0009295 | GO:CC | 0.00026094 |
| axon | GO:0030424 | GO:CC | 0.00026094 |
| euchromatin | GO:0000791 | GO:CC | 0.00027496 |
| contractile ring | GO:0070938 | GO:CC | 0.00030309 |
| DNA polymerase complex | GO:0042575 | GO:CC | 0.00032636 |
| integrator complex | GO:0032039 | GO:CC | 0.00032636 |
| Cul3-RING ubiquitin ligase complex | GO:0031463 | GO:CC | 0.00035003 |
| endoribonuclease complex | GO:1902555 | GO:CC | 0.00035003 |
| RNA polymerase III complex | GO:0005666 | GO:CC | 0.00036658 |
| catalytic step 1 spliceosome | GO:0071012 | GO:CC | 0.00037648 |
| nucleocytoplasmic transport complex | GO:0031074 | GO:CC | 0.00037648 |
| U2-type catalytic step 1 spliceosome | GO:0071006 | GO:CC | 0.00037648 |
| npBAF complex | GO:0071564 | GO:CC | 0.00040155 |
| Cul4A-RING E3 ubiquitin ligase complex | GO:0031464 | GO:CC | 0.00040155 |
| protein serine/threonine phosphatase complex | GO:0008287 | GO:CC | 0.00040327 |
| phosphatase complex | GO:1903293 | GO:CC | 0.00040327 |
| vesicle tethering complex | GO:0099023 | GO:CC | 0.00045531 |
| endonuclease complex | GO:1905348 | GO:CC | 0.00051745 |
| filopodium | GO:0030175 | GO:CC | 0.00053238 |
| U2-type catalytic step 2 spliceosome | GO:0071007 | GO:CC | 0.00054353 |
| actomyosin | GO:0042641 | GO:CC | 0.00064744 |
| mitochondrial inner membrane | GO:0005743 | GO:CC | 0.00078721 |
| filopodium tip | GO:0032433 | GO:CC | 0.00082728 |
| core mediator complex | GO:0070847 | GO:CC | 0.00088071 |
| microtubule plus-end | GO:0035371 | GO:CC | 0.00088071 |
| SMAD protein complex | GO:0071141 | GO:CC | 0.00095204 |
| RNA N6-methyladenosine methyltransferase complex | GO:0036396 | GO:CC | 0.00095204 |
| centriolar satellite | GO:0034451 | GO:CC | 0.00106682 |

| | | | |
|---|---|---|---|
| preribosome, small subunit precursor | GO:0030688 | GO:CC | 0.00106849 |
| pronucleus | GO:0045120 | GO:CC | 0.00106849 |
| replisome | GO:0030894 | GO:CC | 0.00109064 |
| actin filament bundle | GO:0032432 | GO:CC | 0.00109064 |
| mitochondrial large ribosomal subunit | GO:0005762 | GO:CC | 0.00110608 |
| organellar large ribosomal subunit | GO:0000315 | GO:CC | 0.00110608 |
| CMG complex | GO:0071162 | GO:CC | 0.00110608 |
| beta-catenin-TCF complex | GO:1990907 | GO:CC | 0.0011121 |
| Prp19 complex | GO:0000974 | GO:CC | 0.0011121 |
| inclusion body | GO:0016234 | GO:CC | 0.00116006 |
| Flemming body | GO:0090543 | GO:CC | 0.00118376 |
| glutamatergic synapse | GO:0098978 | GO:CC | 0.00120124 |
| endoplasmic reticulum-Golgi intermediate compartmer | GO:0033116 | GO:CC | 0.00127534 |
| cytoplasmic side of membrane | GO:0098562 | GO:CC | 0.00145229 |
| leading edge membrane | GO:0031256 | GO:CC | 0.00148408 |
| clathrin-coated pit | GO:0005905 | GO:CC | 0.00161141 |
| mitotic cohesin complex | GO:0030892 | GO:CC | 0.00192056 |
| autophagosome | GO:0005776 | GO:CC | 0.00214767 |
| transcription repressor complex | GO:0017053 | GO:CC | 0.00215216 |
| contractile actin filament bundle | GO:0097517 | GO:CC | 0.00223632 |
| stress fiber | GO:0001725 | GO:CC | 0.00223632 |
| intercellular bridge | GO:0045171 | GO:CC | 0.00225285 |
| CCR4-NOT complex | GO:0030014 | GO:CC | 0.00236963 |
| ESC/E(Z) complex | GO:0035098 | GO:CC | 0.00236963 |
| PRC1 complex | GO:0035102 | GO:CC | 0.00236963 |
| polymeric cytoskeletal fiber | GO:0099513 | GO:CC | 0.00271626 |
| ATAC complex | GO:0140672 | GO:CC | 0.00276803 |
| heteromeric SMAD protein complex | GO:0071144 | GO:CC | 0.00290454 |
| H3 histone acetyltransferase complex | GO:0070775 | GO:CC | 0.00290454 |
| MLL3/4 complex | GO:0044666 | GO:CC | 0.00307764 |
| NLS-dependent protein nuclear import complex | GO:0042564 | GO:CC | 0.00315801 |
| nucleolar exosome (RNase complex) | GO:0101019 | GO:CC | 0.00315801 |
| N-terminal protein acetyltransferase complex | GO:0031414 | GO:CC | 0.00315801 |
| nuclear pore outer ring | GO:0031080 | GO:CC | 0.00315801 |
| sno(s)RNA-containing ribonucleoprotein complex | GO:0005732 | GO:CC | 0.00327815 |
| coated vesicle | GO:0030135 | GO:CC | 0.00382473 |
| cytoplasmic side of plasma membrane | GO:0009898 | GO:CC | 0.00391452 |
| mRNA cleavage factor complex | GO:0005849 | GO:CC | 0.00417004 |
| neuron projection | GO:0043005 | GO:CC | 0.00419605 |
| mRNA cleavage and polyadenylation specificity factor c | GO:0005847 | GO:CC | 0.00423984 |
| postsynapse | GO:0098794 | GO:CC | 0.00429258 |
| Ino80 complex | GO:0031011 | GO:CC | 0.0047923 |
| late endosome | GO:0005770 | GO:CC | 0.0047923 |
| nuclear inner membrane | GO:0005637 | GO:CC | 0.00523513 |
| DNA repair complex | GO:1990391 | GO:CC | 0.00545001 |
| coated membrane | GO:0048475 | GO:CC | 0.00545001 |
| membrane coat | GO:0030117 | GO:CC | 0.00545001 |
| nuclear replisome | GO:0043601 | GO:CC | 0.00545001 |

| | | | |
|---|---|---|---|
| germ cell nucleus | GO:0043073 | GO:CC | 0.00570101 |
| spindle pole centrosome | GO:0031616 | GO:CC | 0.00580884 |
| XY body | GO:0001741 | GO:CC | 0.00580884 |
| nucleolus | GO:0005730 | GO:CC | 0.0060163 |
| cortical microtubule cytoskeleton | GO:0030981 | GO:CC | 0.00637896 |
| interchromatin granule | GO:0035061 | GO:CC | 0.00637896 |
| DNA replication factor C complex | GO:0005663 | GO:CC | 0.00637896 |
| annulate lamellae | GO:0005642 | GO:CC | 0.00637896 |
| SWI/SNF complex | GO:0016514 | GO:CC | 0.00637896 |
| protein folding chaperone complex | GO:0101031 | GO:CC | 0.00681845 |
| DNA replication preinitiation complex | GO:0031261 | GO:CC | 0.00681845 |
| apical junction complex | GO:0043296 | GO:CC | 0.00681845 |
| chromatin silencing complex | GO:0005677 | GO:CC | 0.00681845 |
| Swr1 complex | GO:0000812 | GO:CC | 0.00681845 |
| cell projection membrane | GO:0031253 | GO:CC | 0.00746236 |
| cortical cytoskeleton | GO:0030863 | GO:CC | 0.00767524 |
| CD40 receptor complex | GO:0035631 | GO:CC | 0.00790425 |
| SMN complex | GO:0032797 | GO:CC | 0.00790425 |
| post-mRNA release spliceosomal complex | GO:0071014 | GO:CC | 0.00790425 |
| ESCRT complex | GO:0036452 | GO:CC | 0.00818046 |
| mitochondrial protein-containing complex | GO:0098798 | GO:CC | 0.00851742 |
| protein phosphatase type 2A complex | GO:0000159 | GO:CC | 0.00851742 |
| MOZ/MORF histone acetyltransferase complex | GO:0070776 | GO:CC | 0.00851742 |
| transcription factor TFTC complex | GO:0033276 | GO:CC | 0.00851742 |
| RNA polymerase III transcription regulator complex | GO:0090576 | GO:CC | 0.00858962 |
| telomerase holoenzyme complex | GO:0005697 | GO:CC | 0.00866976 |
| phagocytic vesicle | GO:0045335 | GO:CC | 0.00901305 |
| vacuole | GO:0005773 | GO:CC | 0.00943666 |
| neuron spine | GO:0044309 | GO:CC | 0.00959429 |
| enzyme activator complex | GO:0150005 | GO:CC | 0.01071248 |
| mRNA editing complex | GO:0045293 | GO:CC | 0.01071248 |
| TIM23 mitochondrial import inner membrane transloca: | GO:0005744 | GO:CC | 0.01071248 |
| chromocenter | GO:0010369 | GO:CC | 0.01071248 |
| cleavage furrow | GO:0032154 | GO:CC | 0.01099938 |
| NuA4 histone acetyltransferase complex | GO:0035267 | GO:CC | 0.01225647 |
| H4/H2A histone acetyltransferase complex | GO:0043189 | GO:CC | 0.01225647 |
| brahma complex | GO:0035060 | GO:CC | 0.01360543 |
| mitochondrial intermembrane space | GO:0005758 | GO:CC | 0.01370543 |
| condensed nuclear chromosome | GO:0000794 | GO:CC | 0.01370543 |
| neuron to neuron synapse | GO:0098984 | GO:CC | 0.01370543 |
| caveola | GO:0005901 | GO:CC | 0.01370543 |
| anaphase-promoting complex | GO:0005680 | GO:CC | 0.01370557 |
| dendritic spine | GO:0043197 | GO:CC | 0.01370557 |
| cortical actin cytoskeleton | GO:0030864 | GO:CC | 0.01422459 |
| male germ cell nucleus | GO:0001673 | GO:CC | 0.01429256 |
| Golgi stack | GO:0005795 | GO:CC | 0.01600737 |
| nuclear inclusion body | GO:0042405 | GO:CC | 0.01649354 |
| actin filament | GO:0005884 | GO:CC | 0.01649354 |

| | | | |
|---|---|---|---|
| SCAR complex | GO:0031209 | GO:CC | 0.01649354 |
| TRAPPIII protein complex | GO:1990072 | GO:CC | 0.01649354 |
| Gemini of coiled bodies | GO:0097504 | GO:CC | 0.01649354 |
| asymmetric synapse | GO:0032279 | GO:CC | 0.01753977 |
| mitochondria-associated endoplasmic reticulum meml | GO:0044233 | GO:CC | 0.01775037 |
| kinetochore microtubule | GO:0005828 | GO:CC | 0.01775037 |
| phagophore assembly site membrane | GO:0034045 | GO:CC | 0.01775037 |
| SNARE complex | GO:0031201 | GO:CC | 0.01796042 |
| RNA cap binding complex | GO:0034518 | GO:CC | 0.01842532 |
| nuclear cyclin-dependent protein kinase holoenzyme c | GO:0019908 | GO:CC | 0.01842532 |
| TRAPP complex | GO:0030008 | GO:CC | 0.01842532 |
| bBAF complex | GO:0140092 | GO:CC | 0.01977858 |
| female pronucleus | GO:0001939 | GO:CC | 0.01977858 |
| CHRAC | GO:0008623 | GO:CC | 0.02028556 |
| BRCA1-B complex | GO:0070532 | GO:CC | 0.02028556 |
| mitotic checkpoint complex | GO:0033597 | GO:CC | 0.02028556 |
| AP-5 adaptor complex | GO:0044599 | GO:CC | 0.02028556 |
| Ndc80 complex | GO:0031262 | GO:CC | 0.02028556 |
| cytoplasmic side of early endosome membrane | GO:0098559 | GO:CC | 0.02028556 |
| actomyosin contractile ring | GO:0005826 | GO:CC | 0.02028556 |
| Pwp2p-containing subcomplex of 90S preribosome | GO:0034388 | GO:CC | 0.02028556 |
| FAR/SIN/STRIPAK complex | GO:0090443 | GO:CC | 0.02028556 |
| phagophore assembly site | GO:0000407 | GO:CC | 0.02087117 |
| RISC-loading complex | GO:0070578 | GO:CC | 0.02231083 |
| ubiquitin conjugating enzyme complex | GO:0031371 | GO:CC | 0.02231083 |
| Ctf18 RFC-like complex | GO:0031390 | GO:CC | 0.02231083 |
| B-WICH complex | GO:0110016 | GO:CC | 0.02231083 |
| X chromosome | GO:0000805 | GO:CC | 0.02231083 |
| gamma-tubulin ring complex | GO:0000931 | GO:CC | 0.02231083 |
| extrinsic component of synaptic membrane | GO:0099243 | GO:CC | 0.02311428 |
| perichromatin fibrils | GO:0005726 | GO:CC | 0.02334849 |
| procentriole replication complex | GO:0120099 | GO:CC | 0.02334849 |
| Atg12-Atg5-Atg16 complex | GO:0034274 | GO:CC | 0.02334849 |
| external side of apical plasma membrane | GO:0098591 | GO:CC | 0.02334849 |
| transcription factor TFIIIC complex | GO:0000127 | GO:CC | 0.02334849 |
| elongator holoenzyme complex | GO:0033588 | GO:CC | 0.02334849 |
| BRCA1-C complex | GO:0070533 | GO:CC | 0.02334849 |
| chromosome passenger complex | GO:0032133 | GO:CC | 0.02334849 |
| CCR4-NOT core complex | GO:0030015 | GO:CC | 0.02334849 |
| distal axon | GO:0150034 | GO:CC | 0.02378343 |
| endocytic vesicle | GO:0030139 | GO:CC | 0.02595758 |
| organelle envelope lumen | GO:0031970 | GO:CC | 0.02613036 |
| autophagosome membrane | GO:0000421 | GO:CC | 0.02842287 |
| SMN-Sm protein complex | GO:0034719 | GO:CC | 0.02899795 |
| DNA replication factor A complex | GO:0005662 | GO:CC | 0.02914005 |
| TOR complex | GO:0038201 | GO:CC | 0.02914005 |
| RNA polymerase I complex | GO:0005736 | GO:CC | 0.02914005 |
| retromer complex | GO:0030904 | GO:CC | 0.03430929 |

| | | | |
|---|---|---|---|
| aggresome | GO:0016235 | GO:CC | 0.03470151 |
| COP9 signalosome | GO:0008180 | GO:CC | 0.03470151 |
| postsynaptic density | GO:0014069 | GO:CC | 0.0354651 |
| phagocytic vesicle membrane | GO:0030670 | GO:CC | 0.03589549 |
| transcription factor TFIID complex | GO:0005669 | GO:CC | 0.03663035 |
| NSL complex | GO:0044545 | GO:CC | 0.0373722 |
| astral microtubule | GO:0000235 | GO:CC | 0.0373722 |
| aster | GO:0005818 | GO:CC | 0.0373722 |
| nBAF complex | GO:0071565 | GO:CC | 0.03747657 |
| COPII vesicle coat | GO:0030127 | GO:CC | 0.03747657 |
| SAGA complex | GO:0000124 | GO:CC | 0.03747657 |
| guanyl-nucleotide exchange factor complex | GO:0032045 | GO:CC | 0.03747657 |
| outer mitochondrial membrane protein complex | GO:0098799 | GO:CC | 0.03747657 |
| Golgi cisterna membrane | GO:0032580 | GO:CC | 0.03778155 |
| clathrin-coated vesicle | GO:0030136 | GO:CC | 0.03879644 |
| tight junction | GO:0070160 | GO:CC | 0.04009376 |
| organellar small ribosomal subunit | GO:0000314 | GO:CC | 0.04460754 |
| mitochondrial small ribosomal subunit | GO:0005763 | GO:CC | 0.04460754 |
| Sin3-type complex | GO:0070822 | GO:CC | 0.04648272 |
| origin recognition complex | GO:0000808 | GO:CC | 0.04648272 |
| nuclear origin of replication recognition complex | GO:0005664 | GO:CC | 0.04648272 |
| zonula adherens | GO:0005915 | GO:CC | 0.04648272 |
| extrinsic component of postsynaptic membrane | GO:0098890 | GO:CC | 0.04648272 |
| host cell | GO:0043657 | GO:CC | 0.04648272 |
| mitotic spindle microtubule | GO:1990498 | GO:CC | 0.04889579 |
| GBAF complex | GO:0140288 | GO:CC | 0.04889579 |
| lysosome | GO:0005764 | GO:CC | 0.04889579 |
| ER membrane insertion complex | GO:0072379 | GO:CC | 0.04889579 |
| lamellipodium membrane | GO:0031258 | GO:CC | 0.04889579 |
| pericentriolar material | GO:0000242 | GO:CC | 0.04889579 |
| lytic vacuole | GO:0000323 | GO:CC | 0.04889579 |
| protein binding | GO:0005515 | GO:MF | 1.78E-199 |
| transferase activity | GO:0016740 | GO:MF | 6.82E-60 |
| binding | GO:0005488 | GO:MF | 2.89E-56 |
| purine ribonucleoside triphosphate binding | GO:0035639 | GO:MF | 3.58E-53 |
| catalytic activity | GO:0003824 | GO:MF | 3.78E-52 |
| purine ribonucleotide binding | GO:0032555 | GO:MF | 6.02E-50 |
| ribonucleotide binding | GO:0032553 | GO:MF | 1.02E-49 |
| ATP binding | GO:0005524 | GO:MF | 1.68E-48 |
| purine nucleotide binding | GO:0017076 | GO:MF | 4.13E-48 |
| nucleotide binding | GO:0000166 | GO:MF | 5.98E-48 |
| nucleoside phosphate binding | GO:1901265 | GO:MF | 6.91E-48 |
| adenyl ribonucleotide binding | GO:0032559 | GO:MF | 2.24E-45 |
| heterocyclic compound binding | GO:1901363 | GO:MF | 6.59E-45 |
| enzyme binding | GO:0019899 | GO:MF | 2.18E-43 |
| adenyl nucleotide binding | GO:0030554 | GO:MF | 2.62E-43 |
| ion binding | GO:0043167 | GO:MF | 9.35E-43 |
| molecular adaptor activity | GO:0060090 | GO:MF | 2.25E-42 |

| | | | |
|---|---|---|---|
| catalytic activity, acting on a nucleic acid | GO:0140640 | GO:MF | 2.30E-42 |
| small molecule binding | GO:0036094 | GO:MF | 1.86E-40 |
| protein-macromolecule adaptor activity | GO:0030674 | GO:MF | 1.96E-38 |
| anion binding | GO:0043168 | GO:MF | 1.25E-37 |
| catalytic activity, acting on a protein | GO:0140096 | GO:MF | 2.58E-36 |
| chromatin binding | GO:0003682 | GO:MF | 1.69E-33 |
| carbohydrate derivative binding | GO:0097367 | GO:MF | 1.83E-31 |
| DNA binding | GO:0003677 | GO:MF | 1.35E-29 |
| protein-containing complex binding | GO:0044877 | GO:MF | 2.85E-29 |
| transcription coregulator activity | GO:0003712 | GO:MF | 3.02E-29 |
| transferase activity, transferring phosphorus-containing | GO:0016772 | GO:MF | 1.78E-28 |
| catalytic activity, acting on RNA | GO:0140098 | GO:MF | 2.89E-27 |
| organic cyclic compound binding | GO:0097159 | GO:MF | 5.77E-26 |
| acyltransferase activity | GO:0016746 | GO:MF | 1.81E-23 |
| pyrophosphatase activity | GO:0016462 | GO:MF | 3.85E-23 |
| hydrolase activity, acting on acid anhydrides, in phosph | GO:0016818 | GO:MF | 4.78E-23 |
| hydrolase activity, acting on acid anhydrides | GO:0016817 | GO:MF | 4.78E-23 |
| ribonucleoside triphosphate phosphatase activity | GO:0017111 | GO:MF | 1.07E-22 |
| ubiquitin-like protein transferase activity | GO:0019787 | GO:MF | 9.43E-22 |
| protein domain specific binding | GO:0019904 | GO:MF | 1.88E-21 |
| ubiquitin-protein transferase activity | GO:0004842 | GO:MF | 1.85E-20 |
| catalytic activity, acting on DNA | GO:0140097 | GO:MF | 3.46E-20 |
| ATP hydrolysis activity | GO:0016887 | GO:MF | 5.72E-20 |
| ATP-dependent activity | GO:0140657 | GO:MF | 6.01E-20 |
| kinase activity | GO:0016301 | GO:MF | 7.58E-20 |
| transcription factor binding | GO:0008134 | GO:MF | 7.58E-20 |
| aminoacyltransferase activity | GO:0016755 | GO:MF | 1.09E-19 |
| histone binding | GO:0042393 | GO:MF | 1.42E-19 |
| metal ion binding | GO:0046872 | GO:MF | 2.11E-19 |
| helicase activity | GO:0004386 | GO:MF | 3.52E-19 |
| histone modifying activity | GO:0140993 | GO:MF | 1.01E-18 |
| phosphotransferase activity, alcohol group as acceptor | GO:0016773 | GO:MF | 2.86E-18 |
| transcription coactivator activity | GO:0003713 | GO:MF | 3.75E-18 |
| cation binding | GO:0043169 | GO:MF | 6.03E-18 |
| protein serine kinase activity | GO:0106310 | GO:MF | 1.13E-17 |
| nucleoside-triphosphatase regulator activity | GO:0060589 | GO:MF | 2.79E-17 |
| GTPase regulator activity | GO:0030695 | GO:MF | 2.79E-17 |
| protein serine/threonine kinase activity | GO:0004674 | GO:MF | 4.18E-17 |
| small GTPase binding | GO:0031267 | GO:MF | 8.76E-17 |
| GTPase binding | GO:0051020 | GO:MF | 1.56E-16 |
| cytoskeletal protein binding | GO:0008092 | GO:MF | 9.11E-16 |
| ATP-dependent activity, acting on DNA | GO:0008094 | GO:MF | 3.87E-15 |
| protein kinase activity | GO:0004672 | GO:MF | 2.29E-14 |
| DNA-binding transcription factor binding | GO:0140297 | GO:MF | 4.15E-14 |
| transcription regulator activity | GO:0140110 | GO:MF | 1.27E-13 |
| microtubule binding | GO:0008017 | GO:MF | 3.47E-13 |
| modification-dependent protein binding | GO:0140030 | GO:MF | 4.39E-13 |
| ubiquitin-like protein ligase activity | GO:0061659 | GO:MF | 4.52E-13 |

| | | | |
|---|---|---|---|
| tubulin binding | GO:0015631 | GO:MF | 2.73E-12 |
| ubiquitin protein ligase activity | GO:0061630 | GO:MF | 7.19E-12 |
| transcription corepressor activity | GO:0003714 | GO:MF | 9.90E-12 |
| ubiquitin-like protein ligase binding | GO:0044389 | GO:MF | 1.24E-11 |
| enzyme activator activity | GO:0008047 | GO:MF | 1.32E-11 |
| ubiquitin protein ligase binding | GO:0031625 | GO:MF | 1.57E-11 |
| cadherin binding | GO:0045296 | GO:MF | 1.57E-11 |
| mRNA binding | GO:0003729 | GO:MF | 4.68E-11 |
| DNA helicase activity | GO:0003678 | GO:MF | 5.11E-11 |
| enzyme regulator activity | GO:0030234 | GO:MF | 5.44E-11 |
| nucleic acid binding | GO:0003676 | GO:MF | 5.45E-11 |
| identical protein binding | GO:0042802 | GO:MF | 1.00E-10 |
| single-stranded DNA binding | GO:0003697 | GO:MF | 2.68E-10 |
| GTPase activator activity | GO:0005096 | GO:MF | 5.48E-10 |
| ubiquitin-like protein binding | GO:0032182 | GO:MF | 6.66E-10 |
| RNA polymerase II-specific DNA-binding transcription fa | GO:0061629 | GO:MF | 7.63E-10 |
| hydrolase activity | GO:0016787 | GO:MF | 8.96E-10 |
| ATP-dependent activity, acting on RNA | GO:0008186 | GO:MF | 1.44E-09 |
| methylation-dependent protein binding | GO:0140034 | GO:MF | 2.07E-09 |
| RNA helicase activity | GO:0003724 | GO:MF | 7.43E-09 |
| ubiquitin conjugating enzyme activity | GO:0061631 | GO:MF | 7.80E-09 |
| methylated histone binding | GO:0035064 | GO:MF | 1.01E-08 |
| nucleotidyltransferase activity | GO:0016779 | GO:MF | 1.25E-08 |
| ribonucleoprotein complex binding | GO:0043021 | GO:MF | 1.25E-08 |
| kinase binding | GO:0019900 | GO:MF | 2.06E-08 |
| magnesium ion binding | GO:0000287 | GO:MF | 4.74E-08 |
| double-stranded RNA binding | GO:0003725 | GO:MF | 7.74E-08 |
| methyltransferase activity | GO:0008168 | GO:MF | 7.74E-08 |
| damaged DNA binding | GO:0003684 | GO:MF | 7.74E-08 |
| ubiquitin-like protein conjugating enzyme activity | GO:0061650 | GO:MF | 8.99E-08 |
| p53 binding | GO:0002039 | GO:MF | 1.05E-07 |
| transferase activity, transferring one-carbon groups | GO:0016741 | GO:MF | 1.28E-07 |
| RNA methyltransferase activity | GO:0008173 | GO:MF | 1.66E-07 |
| zinc ion binding | GO:0008270 | GO:MF | 2.28E-07 |
| catalytic activity, acting on a tRNA | GO:0140101 | GO:MF | 2.78E-07 |
| peptide N-acetyltransferase activity | GO:0034212 | GO:MF | 3.23E-07 |
| exonuclease activity | GO:0004527 | GO:MF | 4.05E-07 |
| acetyltransferase activity | GO:0016407 | GO:MF | 4.05E-07 |
| protein kinase binding | GO:0019901 | GO:MF | 4.05E-07 |
| basal transcription machinery binding | GO:0001098 | GO:MF | 4.10E-07 |
| basal RNA polymerase II transcription machinery bindin | GO:0001099 | GO:MF | 4.10E-07 |
| exonuclease activity, active with either ribo- or deoxyrib | GO:0016796 | GO:MF | 7.86E-07 |
| ubiquitin binding | GO:0043130 | GO:MF | 8.17E-07 |
| peptide-lysine-N-acetyltransferase activity | GO:0061733 | GO:MF | 9.10E-07 |
| nuclear receptor coactivator activity | GO:0030374 | GO:MF | 1.06E-06 |
| histone acetyltransferase activity | GO:0004402 | GO:MF | 1.10E-06 |
| snoRNA binding | GO:0030515 | GO:MF | 1.38E-06 |
| nucleocytoplasmic carrier activity | GO:0140142 | GO:MF | 1.38E-06 |

| | | | |
|---|---|---|---|
| general transcription initiation factor binding | GO:0140296 | GO:MF | 1.38E-06 |
| sequence-specific DNA binding | GO:0043565 | GO:MF | 1.46E-06 |
| guanyl-nucleotide exchange factor activity | GO:0005085 | GO:MF | 1.56E-06 |
| nuclear receptor binding | GO:0016922 | GO:MF | 1.60E-06 |
| phosphoprotein phosphatase activity | GO:0004721 | GO:MF | 1.94E-06 |
| nuclease activity | GO:0004518 | GO:MF | 2.96E-06 |
| N-acetyltransferase activity | GO:0008080 | GO:MF | 2.96E-06 |
| general transcription initiation factor activity | GO:0140223 | GO:MF | 3.01E-06 |
| S-adenosylmethionine-dependent methyltransferase a | GO:0008757 | GO:MF | 3.42E-06 |
| phosphatidylinositol binding | GO:0035091 | GO:MF | 3.62E-06 |
| GTP binding | GO:0005525 | GO:MF | 3.83E-06 |
| double-stranded DNA binding | GO:0003690 | GO:MF | 4.17E-06 |
| RNA exonuclease activity, producing 5'-phosphomonoe | GO:0016896 | GO:MF | 6.46E-06 |
| promoter-specific chromatin binding | GO:1990841 | GO:MF | 7.22E-06 |
| 3'-5'-RNA exonuclease activity | GO:0000175 | GO:MF | 7.66E-06 |
| guanyl nucleotide binding | GO:0019001 | GO:MF | 1.13E-05 |
| guanyl ribonucleotide binding | GO:0032561 | GO:MF | 1.13E-05 |
| cell adhesion molecule binding | GO:0050839 | GO:MF | 1.16E-05 |
| mRNA 3'-UTR binding | GO:0003730 | GO:MF | 1.24E-05 |
| histone deacetylase binding | GO:0042826 | GO:MF | 1.37E-05 |
| 3'-5' exonuclease activity | GO:0008408 | GO:MF | 1.49E-05 |
| RNA exonuclease activity | GO:0004532 | GO:MF | 2.65E-05 |
| DNA secondary structure binding | GO:0000217 | GO:MF | 3.04E-05 |
| protein homodimerization activity | GO:0042803 | GO:MF | 3.11E-05 |
| GTPase activity | GO:0003924 | GO:MF | 3.48E-05 |
| four-way junction DNA binding | GO:0000400 | GO:MF | 4.00E-05 |
| kinase regulator activity | GO:0019207 | GO:MF | 4.69E-05 |
| phosphatase activity | GO:0016791 | GO:MF | 5.06E-05 |
| MAP kinase kinase kinase activity | GO:0004709 | GO:MF | 5.61E-05 |
| rRNA binding | GO:0019843 | GO:MF | 6.20E-05 |
| protein kinase regulator activity | GO:0019887 | GO:MF | 6.34E-05 |
| nucleosome binding | GO:0031491 | GO:MF | 7.87E-05 |
| RNA polymerase binding | GO:0070063 | GO:MF | 8.65E-05 |
| sequence-specific double-stranded DNA binding | GO:1990837 | GO:MF | 9.61E-05 |
| histone H3 methyltransferase activity | GO:0140938 | GO:MF | 0.0001075 |
| single-stranded RNA binding | GO:0003727 | GO:MF | 0.00010987 |
| lysine-acetylated histone binding | GO:0070577 | GO:MF | 0.00013763 |
| structural constituent of nuclear pore | GO:0017056 | GO:MF | 0.00013763 |
| acetylation-dependent protein binding | GO:0140033 | GO:MF | 0.00013763 |
| acyltransferase activity, transferring groups other than a | GO:0016747 | GO:MF | 0.00013829 |
| single-stranded DNA helicase activity | GO:0017116 | GO:MF | 0.00013829 |
| ribosome binding | GO:0043022 | GO:MF | 0.00013829 |
| ATP-dependent chromatin remodeler activity | GO:0140658 | GO:MF | 0.00014318 |
| 4 iron, 4 sulfur cluster binding | GO:0051539 | GO:MF | 0.00015543 |
| tRNA binding | GO:0000049 | GO:MF | 0.00020365 |
| iron-sulfur cluster binding | GO:0051536 | GO:MF | 0.0002119 |
| protein serine/threonine kinase activator activity | GO:0043539 | GO:MF | 0.0002127 |
| demethylase activity | GO:0032451 | GO:MF | 0.00022057 |

| | | | |
|---|---|---|---|
| protein methyltransferase activity | GO:0008276 | GO:MF | 0.00022794 |
| transcription coregulator binding | GO:0001221 | GO:MF | 0.00027196 |
| N-acyltransferase activity | GO:0016410 | GO:MF | 0.000282 |
| plus-end-directed microtubule motor activity | GO:0008574 | GO:MF | 0.00029692 |
| metal cluster binding | GO:0051540 | GO:MF | 0.00030439 |
| nuclear localization sequence binding | GO:0008139 | GO:MF | 0.00031889 |
| hydrolase activity, acting on ester bonds | GO:0016788 | GO:MF | 0.00031889 |
| 2-oxoglutarate-dependent dioxygenase activity | GO:0016706 | GO:MF | 0.00031889 |
| histone methyltransferase activity | GO:0042054 | GO:MF | 0.00031889 |
| histone H4 acetyltransferase activity | GO:0010485 | GO:MF | 0.00033298 |
| RNA polymerase II complex binding | GO:0000993 | GO:MF | 0.00035109 |
| phosphoric ester hydrolase activity | GO:0042578 | GO:MF | 0.00042614 |
| histone ubiquitin ligase activity | GO:0140852 | GO:MF | 0.00046009 |
| N-methyltransferase activity | GO:0008170 | GO:MF | 0.00052335 |
| RNA nuclease activity | GO:0004540 | GO:MF | 0.00052806 |
| kinase activator activity | GO:0019209 | GO:MF | 0.00059454 |
| SMAD binding | GO:0046332 | GO:MF | 0.00061362 |
| nuclear vitamin D receptor binding | GO:0042809 | GO:MF | 0.0007976 |
| signal sequence binding | GO:0005048 | GO:MF | 0.00085792 |
| 5'-3' exonuclease activity | GO:0008409 | GO:MF | 0.00085792 |
| enzyme-substrate adaptor activity | GO:0140767 | GO:MF | 0.00086113 |
| protein-lysine N-methyltransferase activity | GO:0016279 | GO:MF | 0.0008919 |
| ubiquitin-like protein peptidase activity | GO:0019783 | GO:MF | 0.00096976 |
| actin binding | GO:0003779 | GO:MF | 0.00096976 |
| miRNA binding | GO:0035198 | GO:MF | 0.00110592 |
| ubiquitin-like protein conjugating enzyme binding | GO:0044390 | GO:MF | 0.00110592 |
| transition metal ion binding | GO:0046914 | GO:MF | 0.00117029 |
| DNA-binding transcription activator activity | GO:0001216 | GO:MF | 0.00117029 |
| mRNA 3'-UTR AU-rich region binding | GO:0035925 | GO:MF | 0.00119539 |
| tRNA methyltransferase activity | GO:0008175 | GO:MF | 0.00119539 |
| lysine N-methyltransferase activity | GO:0016278 | GO:MF | 0.00128594 |
| RNA polymerase core enzyme binding | GO:0043175 | GO:MF | 0.00150889 |
| RNA polymerase II general transcription initiation factor | GO:0016251 | GO:MF | 0.00150889 |
| chromatin DNA binding | GO:0031490 | GO:MF | 0.0015684 |
| protein dimerization activity | GO:0046983 | GO:MF | 0.00175089 |
| RNA polymerase III general transcription initiation facto | GO:0000995 | GO:MF | 0.00180869 |
| nuclear export signal receptor activity | GO:0005049 | GO:MF | 0.00180869 |
| ubiquitin conjugating enzyme binding | GO:0031624 | GO:MF | 0.00185545 |
| transcription regulatory region nucleic acid binding | GO:0001067 | GO:MF | 0.00186935 |
| nuclear import signal receptor activity | GO:0061608 | GO:MF | 0.00189713 |
| RNA cap binding | GO:0000339 | GO:MF | 0.00189713 |
| protein kinase activator activity | GO:0030295 | GO:MF | 0.00195814 |
| C2H2 zinc finger domain binding | GO:0070742 | GO:MF | 0.00199058 |
| mRNA methyltransferase activity | GO:0008174 | GO:MF | 0.00199058 |
| histone H3 acetyltransferase activity | GO:0010484 | GO:MF | 0.00199058 |
| SUMO binding | GO:0032183 | GO:MF | 0.00204884 |
| core promoter sequence-specific DNA binding | GO:0001046 | GO:MF | 0.00235589 |
| molecular carrier activity | GO:0140104 | GO:MF | 0.00238107 |

| | | | |
|---|---|---|---|
| DNA-binding transcription activator activity, RNA polym | GO:0001228 | GO:MF | 0.00238107 |
| transcription cis-regulatory region binding | GO:0000976 | GO:MF | 0.00243652 |
| RNA polymerase activity | GO:0097747 | GO:MF | 0.00256554 |
| 5'-3' RNA polymerase activity | GO:0034062 | GO:MF | 0.00256554 |
| heat shock protein binding | GO:0031072 | GO:MF | 0.00258912 |
| glycosyltransferase activity | GO:0016757 | GO:MF | 0.00258912 |
| ubiquitin ligase-substrate adaptor activity | GO:1990756 | GO:MF | 0.00284722 |
| RNA polymerase II general transcription initiation factor | GO:0001091 | GO:MF | 0.00289402 |
| DNA-directed 5'-3' RNA polymerase activity | GO:0003899 | GO:MF | 0.00307269 |
| protein tyrosine phosphatase activity | GO:0004725 | GO:MF | 0.0032562 |
| transcription coactivator binding | GO:0001223 | GO:MF | 0.00362889 |
| 2-acylglycerol-3-phosphate O-acyltransferase activity | GO:0047144 | GO:MF | 0.00375534 |
| anaphase-promoting complex binding | GO:0010997 | GO:MF | 0.00375534 |
| 5'-3' DNA helicase activity | GO:0043139 | GO:MF | 0.00375534 |
| 1-acylglycerol-3-phosphate O-acyltransferase activity | GO:0003841 | GO:MF | 0.00378542 |
| microtubule plus-end binding | GO:0051010 | GO:MF | 0.00378542 |
| microtubule motor activity | GO:0003777 | GO:MF | 0.00388761 |
| nuclear thyroid hormone receptor binding | GO:0046966 | GO:MF | 0.00438972 |
| lncRNA binding | GO:0106222 | GO:MF | 0.00470057 |
| K63-linked deubiquitinase activity | GO:0061578 | GO:MF | 0.00500044 |
| poly-purine tract binding | GO:0070717 | GO:MF | 0.00506178 |
| catalytic activity, acting on a rRNA | GO:0140102 | GO:MF | 0.00522605 |
| cis-regulatory region sequence-specific DNA binding | GO:0000987 | GO:MF | 0.00536397 |
| protein demethylase activity | GO:0140457 | GO:MF | 0.00615687 |
| histone demethylase activity | GO:0032452 | GO:MF | 0.00615687 |
| rRNA methyltransferase activity | GO:0008649 | GO:MF | 0.00618556 |
| chromo shadow domain binding | GO:0070087 | GO:MF | 0.00660764 |
| C-methyltransferase activity | GO:0008169 | GO:MF | 0.00660764 |
| ligase activity, forming phosphoric ester bonds | GO:0016886 | GO:MF | 0.00660764 |
| lipid kinase activity | GO:0001727 | GO:MF | 0.00661996 |
| deubiquitinase activity | GO:0101005 | GO:MF | 0.00669004 |
| protein serine/threonine phosphatase activity | GO:0004722 | GO:MF | 0.00690133 |
| lysophosphatidic acid acyltransferase activity | GO:0042171 | GO:MF | 0.00711649 |
| lysophospholipid acyltransferase activity | GO:0071617 | GO:MF | 0.00711649 |
| phosphatase regulator activity | GO:0019208 | GO:MF | 0.00711649 |
| protein phosphatase 2A binding | GO:0051721 | GO:MF | 0.00726269 |
| regulatory RNA binding | GO:0061980 | GO:MF | 0.00746662 |
| GDP binding | GO:0019003 | GO:MF | 0.00756943 |
| RNA endonuclease activity, producing 5'-phosphomono | GO:0016891 | GO:MF | 0.00784957 |
| phosphatidylinositol phosphate binding | GO:1901981 | GO:MF | 0.00820116 |
| pentosyltransferase activity | GO:0016763 | GO:MF | 0.00856622 |
| translation factor activity, RNA binding | GO:0008135 | GO:MF | 0.00885001 |
| phosphatase binding | GO:0019902 | GO:MF | 0.00967015 |
| transition metal ion transmembrane transporter activity | GO:0046915 | GO:MF | 0.00967015 |
| histone H3K4 monomethyltransferase activity | GO:0140945 | GO:MF | 0.01039388 |
| TBP-class protein binding | GO:0017025 | GO:MF | 0.01039388 |
| histone kinase activity | GO:0035173 | GO:MF | 0.01039388 |
| RNA polymerase I transcription regulatory region seque | GO:0001163 | GO:MF | 0.01039388 |

| | | | |
|---|---|---|---|
| K63-linked polyubiquitin modification-dependent prote | GO:0070530 | GO:MF | 0.01039388 |
| RNA polymerase I core promoter sequence-specific DN | GO:0001164 | GO:MF | 0.01039388 |
| GDP-dissociation inhibitor activity | GO:0005092 | GO:MF | 0.01046467 |
| diacylglycerol-dependent serine/threonine kinase activ | GO:0004697 | GO:MF | 0.01046467 |
| protein phosphatase binding | GO:0019903 | GO:MF | 0.01058625 |
| phosphatidylinositol phosphate phosphatase activity | GO:0052866 | GO:MF | 0.01149919 |
| DNA replication origin binding | GO:0003688 | GO:MF | 0.01149919 |
| RNA polymerase II cis-regulatory region sequence-spec | GO:0000978 | GO:MF | 0.01183616 |
| Tat protein binding | GO:0030957 | GO:MF | 0.01183616 |
| histone H4K12 acetyltransferase activity | GO:0043997 | GO:MF | 0.01196978 |
| RNA 7-methylguanosine cap binding | GO:0000340 | GO:MF | 0.01196978 |
| beta-catenin binding | GO:0008013 | GO:MF | 0.01263296 |
| cyclin-dependent protein serine/threonine kinase regul | GO:0016538 | GO:MF | 0.01263296 |
| endonuclease activity, active with either ribo- or deoxyri | GO:0016893 | GO:MF | 0.01361756 |
| SH3 domain binding | GO:0017124 | GO:MF | 0.01380387 |
| actin filament binding | GO:0051015 | GO:MF | 0.01436705 |
| gamma-tubulin binding | GO:0043015 | GO:MF | 0.01442676 |
| DNA nuclease activity | GO:0004536 | GO:MF | 0.01540647 |
| RNA polymerase II CTD heptapeptide repeat modifying a | GO:0140994 | GO:MF | 0.01702793 |
| phosphatidylinositol bisphosphate phosphatase activity | GO:0034593 | GO:MF | 0.01702793 |
| histone H3K4 methyltransferase activity | GO:0042800 | GO:MF | 0.01732297 |
| DNA binding, bending | GO:0008301 | GO:MF | 0.01732297 |
| transcription corepressor binding | GO:0001222 | GO:MF | 0.01756104 |
| myosin phosphatase activity | GO:0017018 | GO:MF | 0.01861677 |
| DNA demethylase activity | GO:0035514 | GO:MF | 0.01937589 |
| neuropilin binding | GO:0038191 | GO:MF | 0.01937589 |
| histone H2A ubiquitin ligase activity | GO:0141053 | GO:MF | 0.01937589 |
| oxidative RNA demethylase activity | GO:0035515 | GO:MF | 0.01937589 |
| histone H4 demethylase activity | GO:0141058 | GO:MF | 0.01937589 |
| N-acetyllactosamine synthase activity | GO:0003945 | GO:MF | 0.01937589 |
| cysteine-type deubiquitinase activity | GO:0004843 | GO:MF | 0.01937589 |
| broad specificity oxidative DNA demethylase activity | GO:0035516 | GO:MF | 0.01937589 |
| ribosomal small subunit binding | GO:0043024 | GO:MF | 0.01990959 |
| translation regulator activity | GO:0045182 | GO:MF | 0.02067592 |
| histone H3 demethylase activity | GO:0141052 | GO:MF | 0.02142633 |
| adenyl-nucleotide exchange factor activity | GO:0000774 | GO:MF | 0.02303456 |
| phosphatidylinositol-3,5-bisphosphate phosphatase ac | GO:0106018 | GO:MF | 0.02303456 |
| protein-containing complex destabilizing activity | GO:0140776 | GO:MF | 0.02303456 |
| telomerase RNA binding | GO:0070034 | GO:MF | 0.02459118 |
| poly(A)-specific ribonuclease activity | GO:0004535 | GO:MF | 0.02589332 |
| microtubule minus-end binding | GO:0051011 | GO:MF | 0.02589332 |
| insulin receptor substrate binding | GO:0043560 | GO:MF | 0.02589332 |
| phosphatidylinositol-3-phosphate phosphatase activity | GO:0004438 | GO:MF | 0.02589332 |
| protein kinase A binding | GO:0051018 | GO:MF | 0.02589332 |
| histone H3K36 methyltransferase activity | GO:0046975 | GO:MF | 0.02589332 |
| non-membrane spanning protein tyrosine kinase activit | GO:0004715 | GO:MF | 0.02589711 |
| signaling adaptor activity | GO:0035591 | GO:MF | 0.02591855 |
| calcium-dependent protein serine/threonine phosphata | GO:0004723 | GO:MF | 0.02713868 |

| | | | |
|---|---|---|---|
| histone H3K9me2 methyltransferase activity | GO:0140947 | GO:MF | 0.02713868 |
| deSUMOylase activity | GO:0016929 | GO:MF | 0.02713868 |
| copper ion transmembrane transporter activity | GO:0005375 | GO:MF | 0.02713868 |
| UDP-xylosyltransferase activity | GO:0035252 | GO:MF | 0.02756429 |
| histone H3K36 demethylase activity | GO:0051864 | GO:MF | 0.02756429 |
| UDP-glycosyltransferase activity | GO:0008194 | GO:MF | 0.02756429 |
| xylosyltransferase activity | GO:0042285 | GO:MF | 0.02756429 |
| sulfur amino acid transmembrane transporter activity | GO:0000099 | GO:MF | 0.02756429 |
| RNA polymerase II CTD heptapeptide repeat kinase acti | GO:0008353 | GO:MF | 0.02756429 |
| single-stranded DNA endodeoxyribonuclease activity | GO:0000014 | GO:MF | 0.02756429 |
| ATPase regulator activity | GO:0060590 | GO:MF | 0.02780412 |
| histone H3K14 acetyltransferase activity | GO:0036408 | GO:MF | 0.02822741 |
| primary miRNA binding | GO:0070878 | GO:MF | 0.02822741 |
| cysteine-type peptidase activity | GO:0008234 | GO:MF | 0.02822741 |
| tRNA (cytidine) methyltransferase activity | GO:0016427 | GO:MF | 0.02822741 |
| nucleoside kinase activity | GO:0019206 | GO:MF | 0.02822741 |
| DNA exonuclease activity | GO:0004529 | GO:MF | 0.03076323 |
| S-acyltransferase activity | GO:0016417 | GO:MF | 0.03076561 |
| translation regulator activity, nucleic acid binding | GO:0090079 | GO:MF | 0.03145047 |
| prenyltransferase activity | GO:0004659 | GO:MF | 0.0337038 |
| protein-RNA adaptor activity | GO:0140517 | GO:MF | 0.0337038 |
| telomeric DNA binding | GO:0042162 | GO:MF | 0.0337038 |
| phospholipid binding | GO:0005543 | GO:MF | 0.0337038 |
| inositol phosphate phosphatase activity | GO:0052745 | GO:MF | 0.0337038 |
| phosphoprotein binding | GO:0051219 | GO:MF | 0.03437675 |
| protein tyrosine kinase activity | GO:0004713 | GO:MF | 0.03704923 |
| phosphotransferase activity, for other substituted phos | GO:0016780 | GO:MF | 0.03749246 |
| endonuclease activity | GO:0004519 | GO:MF | 0.03749246 |
| poly(A) binding | GO:0008143 | GO:MF | 0.03768648 |
| manganese ion binding | GO:0030145 | GO:MF | 0.04024925 |
| magnesium ion transmembrane transporter activity | GO:0015095 | GO:MF | 0.04026228 |
| lamin binding | GO:0005521 | GO:MF | 0.04026228 |
| G protein activity | GO:0003925 | GO:MF | 0.04147642 |
| hexosyltransferase activity | GO:0016758 | GO:MF | 0.04157125 |
| molecular function regulator activity | GO:0098772 | GO:MF | 0.04301377 |
| polyubiquitin modification-dependent protein binding | GO:0031593 | GO:MF | 0.04445677 |
| chromatin-protein adaptor activity | GO:0140463 | GO:MF | 0.0456876 |
| DNA polymerase binding | GO:0070182 | GO:MF | 0.0456876 |
| protein-cysteine S-palmitoyltransferase activity | GO:0019706 | GO:MF | 0.04612607 |
| protein-cysteine S-acyltransferase activity | GO:0019707 | GO:MF | 0.04612607 |
| nuclear retinoic acid receptor binding | GO:0042974 | GO:MF | 0.04612607 |
| RNA polymerase II transcription regulatory region seque | GO:0000977 | GO:MF | 0.0465078 |
| Arp2/3 complex binding | GO:0071933 | GO:MF | 0.04698331 |
| 1-phosphatidylinositol binding | GO:0005545 | GO:MF | 0.04698331 |
| phosphatidylinositol monophosphate phosphatase acti | GO:0052744 | GO:MF | 0.04698331 |
| protein phosphatase regulator activity | GO:0019888 | GO:MF | 0.04962326 |
| Cell cycle | KEGG:04110 | KEGG | 5.77E-18 |
| Ubiquitin mediated proteolysis | KEGG:04120 | KEGG | 4.81E-15 |

| | | | |
|---|---|---|---|
| Nucleocytoplasmic transport | KEGG:03013 | KEGG | 7.91E-15 |
| Polycomb repressive complex | KEGG:03083 | KEGG | 5.65E-12 |
| Endocytosis | KEGG:04144 | KEGG | 2.62E-08 |
| mRNA surveillance pathway | KEGG:03015 | KEGG | 8.08E-07 |
| Shigellosis | KEGG:05131 | KEGG | 2.12E-06 |
| DNA replication | KEGG:03030 | KEGG | 2.12E-06 |
| Chronic myeloid leukemia | KEGG:05220 | KEGG | 2.23E-06 |
| Renal cell carcinoma | KEGG:05211 | KEGG | 3.16E-06 |
| Adherens junction | KEGG:04520 | KEGG | 3.16E-06 |
| Spliceosome | KEGG:03040 | KEGG | 3.36E-06 |
| Cellular senescence | KEGG:04218 | KEGG | 4.65E-06 |
| Oocyte meiosis | KEGG:04114 | KEGG | 5.92E-06 |
| EGFR tyrosine kinase inhibitor resistance | KEGG:01521 | KEGG | 6.82E-06 |
| ATP-dependent chromatin remodeling | KEGG:03082 | KEGG | 1.20E-05 |
| Pancreatic cancer | KEGG:05212 | KEGG | 1.28E-05 |
| Phosphatidylinositol signaling system | KEGG:04070 | KEGG | 1.28E-05 |
| RNA degradation | KEGG:03018 | KEGG | 1.71E-05 |
| Autophagy - animal | KEGG:04140 | KEGG | 3.32E-05 |
| Progesterone-mediated oocyte maturation | KEGG:04914 | KEGG | 7.99E-05 |
| Salmonella infection | KEGG:05132 | KEGG | 0.00016127 |
| Hepatocellular carcinoma | KEGG:05225 | KEGG | 0.00016963 |
| Nucleotide excision repair | KEGG:03420 | KEGG | 0.00017541 |
| Hepatitis B | KEGG:05161 | KEGG | 0.00020984 |
| Human immunodeficiency virus 1 infection | KEGG:05170 | KEGG | 0.00020984 |
| Neurotrophin signaling pathway | KEGG:04722 | KEGG | 0.00020984 |
| Colorectal cancer | KEGG:05210 | KEGG | 0.00024215 |
| mTOR signaling pathway | KEGG:04150 | KEGG | 0.00024215 |
| Human T-cell leukemia virus 1 infection | KEGG:05166 | KEGG | 0.00031884 |
| Hedgehog signaling pathway | KEGG:04340 | KEGG | 0.00034683 |
| Lysine degradation | KEGG:00310 | KEGG | 0.00034683 |
| Fanconi anemia pathway | KEGG:03460 | KEGG | 0.00039425 |
| Thyroid hormone signaling pathway | KEGG:04919 | KEGG | 0.0004249 |
| Basal transcription factors | KEGG:03022 | KEGG | 0.00046277 |
| Sphingolipid signaling pathway | KEGG:04071 | KEGG | 0.0006396 |
| Hippo signaling pathway | KEGG:04390 | KEGG | 0.0006396 |
| Yersinia infection | KEGG:05135 | KEGG | 0.00066231 |
| Base excision repair | KEGG:03410 | KEGG | 0.00072156 |
| Protein processing in endoplasmic reticulum | KEGG:04141 | KEGG | 0.00119118 |
| Viral life cycle - HIV-1 | KEGG:03250 | KEGG | 0.00136893 |
| Glioma | KEGG:05214 | KEGG | 0.00171553 |
| Gap junction | KEGG:04540 | KEGG | 0.00188223 |
| Longevity regulating pathway - multiple species | KEGG:04213 | KEGG | 0.00226984 |
| PD-L1 expression and PD-1 checkpoint pathway in canc | KEGG:05235 | KEGG | 0.00246342 |
| ErbB signaling pathway | KEGG:04012 | KEGG | 0.00251284 |
| Prostate cancer | KEGG:05215 | KEGG | 0.0025319 |
| Human cytomegalovirus infection | KEGG:05163 | KEGG | 0.00286355 |
| Non-small cell lung cancer | KEGG:05223 | KEGG | 0.00298317 |
| Proteoglycans in cancer | KEGG:05205 | KEGG | 0.00372659 |

| | | | |
|---|---|---|---|
| RNA polymerase | KEGG:03020 | KEGG | 0.00393389 |
| Inositol phosphate metabolism | KEGG:00562 | KEGG | 0.00393389 |
| Mismatch repair | KEGG:03430 | KEGG | 0.0041578 |
| Longevity regulating pathway | KEGG:04211 | KEGG | 0.00445667 |
| Choline metabolism in cancer | KEGG:05231 | KEGG | 0.00594526 |
| Long-term potentiation | KEGG:04720 | KEGG | 0.00615884 |
| Hepatitis C | KEGG:05160 | KEGG | 0.00615884 |
| MAPK signaling pathway | KEGG:04010 | KEGG | 0.00660583 |
| Pathways in cancer | KEGG:05200 | KEGG | 0.00735218 |
| Regulation of actin cytoskeleton | KEGG:04810 | KEGG | 0.0073798 |
| Apelin signaling pathway | KEGG:04371 | KEGG | 0.0087317 |
| Pathogenic Escherichia coli infection | KEGG:05130 | KEGG | 0.00889622 |
| Homologous recombination | KEGG:03440 | KEGG | 0.00934485 |
| Endocrine resistance | KEGG:01522 | KEGG | 0.00934485 |
| Bacterial invasion of epithelial cells | KEGG:05100 | KEGG | 0.01067301 |
| Rap1 signaling pathway | KEGG:04015 | KEGG | 0.01090211 |
| Apoptosis - multiple species | KEGG:04215 | KEGG | 0.01154216 |
| TGF-beta signaling pathway | KEGG:04350 | KEGG | 0.01179765 |
| Mannose type O-glycan biosynthesis | KEGG:00515 | KEGG | 0.0127916 |
| Other types of O-glycan biosynthesis | KEGG:00514 | KEGG | 0.01292318 |
| Mitophagy - animal | KEGG:04137 | KEGG | 0.01571917 |
| Gastric cancer | KEGG:05226 | KEGG | 0.01755763 |
| Fc gamma R-mediated phagocytosis | KEGG:04666 | KEGG | 0.01999045 |
| Acute myeloid leukemia | KEGG:05221 | KEGG | 0.0227209 |
| Hippo signaling pathway - multiple species | KEGG:04392 | KEGG | 0.023314 |
| Cushing syndrome | KEGG:04934 | KEGG | 0.02428956 |
| Aminoacyl-tRNA biosynthesis | KEGG:00970 | KEGG | 0.02428956 |
| Ribosome biogenesis in eukaryotes | KEGG:03008 | KEGG | 0.02447992 |
| Kaposi sarcoma-associated herpesvirus infection | KEGG:05167 | KEGG | 0.02657273 |
| Nucleotide metabolism | KEGG:01232 | KEGG | 0.02657273 |
| Notch signaling pathway | KEGG:04330 | KEGG | 0.02681581 |
| Autophagy - other | KEGG:04136 | KEGG | 0.02731366 |
| Apoptosis | KEGG:04210 | KEGG | 0.03088746 |
| Measles | KEGG:05162 | KEGG | 0.03140664 |
| T cell receptor signaling pathway | KEGG:04660 | KEGG | 0.03579635 |
| Insulin signaling pathway | KEGG:04910 | KEGG | 0.0417812 |
| Growth hormone synthesis, secretion and action | KEGG:04935 | KEGG | 0.0417812 |
| Small cell lung cancer | KEGG:05222 | KEGG | 0.0417812 |
| Sphingolipid metabolism | KEGG:00600 | KEGG | 0.04188214 |
| Chagas disease | KEGG:05142 | KEGG | 0.04633312 |
| Tight junction | KEGG:04530 | KEGG | 0.04847197 |
| Metabolism of RNA | REAC:R-HSA-8 | REAC | 4.91E-39 |
| Cell Cycle | REAC:R-HSA-1 | REAC | 4.62E-34 |
| Cell Cycle, Mitotic | REAC:R-HSA-6 | REAC | 4.06E-29 |
| Processing of Capped Intron-Containing Pre-mRNA | REAC:R-HSA-7 | REAC | 4.08E-26 |
| Gene expression (Transcription) | REAC:R-HSA-7 | REAC | 1.66E-23 |
| Mitotic Prometaphase | REAC:R-HSA-6 | REAC | 6.69E-20 |
| mRNA Splicing - Major Pathway | REAC:R-HSA-7 | REAC | 1.11E-19 |

| | | | |
|---|---|---|---|
| mRNA Splicing | REAC:R-HSA-7 | REAC | 1.70E-19 |
| Signaling by Rho GTPases, Miro GTPases and RHOBTB3 | REAC:R-HSA-9 | REAC | 4.29E-19 |
| Signaling by Rho GTPases | REAC:R-HSA-1 | REAC | 7.30E-18 |
| Resolution of Sister Chromatid Cohesion | REAC:R-HSA-2 | REAC | 1.18E-15 |
| Membrane Trafficking | REAC:R-HSA-1 | REAC | 6.82E-15 |
| Mitotic Spindle Checkpoint | REAC:R-HSA-6 | REAC | 1.55E-14 |
| Amplification of signal from unattached kinetochores v | REAC:R-HSA-1 | REAC | 1.55E-14 |
| Amplification of signal from the kinetochores | REAC:R-HSA-1 | REAC | 1.55E-14 |
| Mitotic Metaphase and Anaphase | REAC:R-HSA-2 | REAC | 3.09E-14 |
| M Phase | REAC:R-HSA-6 | REAC | 5.03E-14 |
| Mitotic Anaphase | REAC:R-HSA-6 | REAC | 5.03E-14 |
| RHO GTPase cycle | REAC:R-HSA-9 | REAC | 9.11E-13 |
| DNA Repair | REAC:R-HSA-7 | REAC | 9.53E-13 |
| EML4 and NUDC in mitotic spindle formation | REAC:R-HSA-9 | REAC | 1.32E-12 |
| RNA Polymerase II Transcription | REAC:R-HSA-7 | REAC | 2.81E-12 |
| Transcriptional Regulation by TP53 | REAC:R-HSA-3 | REAC | 3.29E-12 |
| SUMOylation | REAC:R-HSA-2 | REAC | 4.80E-12 |
| tRNA processing | REAC:R-HSA-7 | REAC | 4.80E-12 |
| RHO GTPases Activate Formins | REAC:R-HSA-5 | REAC | 6.17E-12 |
| Cell Cycle Checkpoints | REAC:R-HSA-6 | REAC | 1.00E-11 |
| Transport of Mature Transcript to Cytoplasm | REAC:R-HSA-7 | REAC | 2.77E-11 |
| HIV Life Cycle | REAC:R-HSA-1 | REAC | 2.77E-11 |
| SUMO E3 ligases SUMOylate target proteins | REAC:R-HSA-3 | REAC | 5.13E-11 |
| Late Phase of HIV Life Cycle | REAC:R-HSA-1 | REAC | 7.74E-11 |
| Transport of Mature mRNA derived from an Intron-Conta | REAC:R-HSA-1 | REAC | 2.14E-10 |
| SUMOylation of DNA replication proteins | REAC:R-HSA-4 | REAC | 4.89E-10 |
| SUMOylation of DNA damage response and repair prote | REAC:R-HSA-3 | REAC | 4.94E-10 |
| Regulation of TP53 Activity | REAC:R-HSA-5 | REAC | 6.76E-10 |
| Nucleotide Excision Repair | REAC:R-HSA-5 | REAC | 9.32E-10 |
| Separation of Sister Chromatids | REAC:R-HSA-2 | REAC | 2.89E-09 |
| rRNA modification in the nucleus and cytosol | REAC:R-HSA-6 | REAC | 3.10E-09 |
| Chromatin modifying enzymes | REAC:R-HSA-3 | REAC | 2.63E-08 |
| Chromatin organization | REAC:R-HSA-4 | REAC | 2.63E-08 |
| mRNA 3'-end processing | REAC:R-HSA-7 | REAC | 3.42E-08 |
| Global Genome Nucleotide Excision Repair (GG-NER) | REAC:R-HSA-5 | REAC | 4.10E-08 |
| S Phase | REAC:R-HSA-6 | REAC | 8.54E-08 |
| RNA Polymerase II Transcription Termination | REAC:R-HSA-7 | REAC | 8.95E-08 |
| Post-translational protein modification | REAC:R-HSA-5 | REAC | 8.95E-08 |
| Intra-Golgi and retrograde Golgi-to-ER traffic | REAC:R-HSA-6 | REAC | 9.53E-08 |
| tRNA processing in the nucleus | REAC:R-HSA-6 | REAC | 3.54E-07 |
| Vesicle-mediated transport | REAC:R-HSA-5 | REAC | 6.40E-07 |
| Antigen processing: Ubiquitination & Proteasome degra | REAC:R-HSA-9 | REAC | 1.17E-06 |
| Regulation of TP53 Activity through Phosphorylation | REAC:R-HSA-6 | REAC | 1.19E-06 |
| CDC42 GTPase cycle | REAC:R-HSA-9 | REAC | 1.19E-06 |
| Transcriptional Regulation by E2F6 | REAC:R-HSA-8 | REAC | 1.26E-06 |
| Metabolism of non-coding RNA | REAC:R-HSA-1 | REAC | 1.41E-06 |
| snRNP Assembly | REAC:R-HSA-1 | REAC | 1.41E-06 |
| RAC1 GTPase cycle | REAC:R-HSA-9 | REAC | 1.44E-06 |

| | | | |
|---|---|---|---|
| NS1 Mediated Effects on Host Pathways | REAC:R-HSA-1 | REAC | 1.77E-06 |
| Rab regulation of trafficking | REAC:R-HSA-9 | REAC | 2.13E-06 |
| RHOC GTPase cycle | REAC:R-HSA-9 | REAC | 2.13E-06 |
| Nuclear Pore Complex (NPC) Disassembly | REAC:R-HSA-3 | REAC | 2.71E-06 |
| SUMOylation of ubiquitinylation proteins | REAC:R-HSA-3 | REAC | 3.28E-06 |
| Transcription-Coupled Nucleotide Excision Repair (TC-N | REAC:R-HSA-6 | REAC | 3.38E-06 |
| RAB GEFs exchange GTP for GDP on RABs | REAC:R-HSA-8 | REAC | 3.79E-06 |
| Mitotic G1 phase and G1/S transition | REAC:R-HSA-4 | REAC | 3.92E-06 |
| RHO GTPase Effectors | REAC:R-HSA-1 | REAC | 4.04E-06 |
| Transport of Mature mRNAs Derived from Intronless Tra | REAC:R-HSA-1 | REAC | 4.07E-06 |
| RNA Polymerase III Abortive And Retractive Initiation | REAC:R-HSA-7 | REAC | 4.24E-06 |
| RNA Polymerase III Transcription | REAC:R-HSA-7 | REAC | 4.24E-06 |
| Formation of WDR5-containing histone-modifying comp | REAC:R-HSA-9 | REAC | 4.24E-06 |
| Regulation of HSF1-mediated heat shock response | REAC:R-HSA-3 | REAC | 4.24E-06 |
| RNA Polymerase II Pre-transcription Events | REAC:R-HSA-6 | REAC | 4.80E-06 |
| Generic Transcription Pathway | REAC:R-HSA-2 | REAC | 4.97E-06 |
| Transport of the SLBP Dependant Mature mRNA | REAC:R-HSA-1 | REAC | 5.23E-06 |
| DNA strand elongation | REAC:R-HSA-6 | REAC | 5.23E-06 |
| Rev-mediated nuclear export of HIV RNA | REAC:R-HSA-1 | REAC | 5.23E-06 |
| Cellular response to heat stress | REAC:R-HSA-3 | REAC | 5.83E-06 |
| Interactions of Rev with host cellular proteins | REAC:R-HSA-1 | REAC | 6.14E-06 |
| Transport of Mature mRNA Derived from an Intronless T | REAC:R-HSA-1 | REAC | 7.86E-06 |
| Transport of Ribonucleoproteins into the Host Nucleus | REAC:R-HSA-1 | REAC | 9.50E-06 |
| NEP/NS2 Interacts with the Cellular Export Machinery | REAC:R-HSA-1 | REAC | 9.50E-06 |
| Transport of the SLBP independent Mature mRNA | REAC:R-HSA-1 | REAC | 1.16E-05 |
| Postmitotic nuclear pore complex (NPC) reformation | REAC:R-HSA-9 | REAC | 1.27E-05 |
| Activation of the pre-replicative complex | REAC:R-HSA-6 | REAC | 1.34E-05 |
| Mitotic G2-G2/M phases | REAC:R-HSA-4 | REAC | 1.35E-05 |
| Epigenetic regulation of gene expression | REAC:R-HSA-2 | REAC | 1.41E-05 |
| Antiviral mechanism by IFN-stimulated genes | REAC:R-HSA-1 | REAC | 1.41E-05 |
| Activation of ATR in response to replication stress | REAC:R-HSA-1 | REAC | 1.58E-05 |
| HDR through Single Strand Annealing (SSA) | REAC:R-HSA-5 | REAC | 1.58E-05 |
| HIV Infection | REAC:R-HSA-1 | REAC | 1.63E-05 |
| Regulation of PLK1 Activity at G2/M Transition | REAC:R-HSA-2 | REAC | 1.65E-05 |
| RHOJ GTPase cycle | REAC:R-HSA-9 | REAC | 1.65E-05 |
| SUMOylation of RNA binding proteins | REAC:R-HSA-4 | REAC | 1.75E-05 |
| Nuclear Envelope Breakdown | REAC:R-HSA-2 | REAC | 1.76E-05 |
| Nuclear Envelope (NE) Reassembly | REAC:R-HSA-2 | REAC | 1.82E-05 |
| ISG15 antiviral mechanism | REAC:R-HSA-1 | REAC | 1.82E-05 |
| Dual incision in TC-NER | REAC:R-HSA-6 | REAC | 2.28E-05 |
| Diseases of signal transduction by growth factor recept | REAC:R-HSA-5 | REAC | 2.28E-05 |
| Export of Viral Ribonucleoproteins from Nucleus | REAC:R-HSA-1 | REAC | 2.28E-05 |
| SUMOylation of SUMOylation proteins | REAC:R-HSA-4 | REAC | 2.61E-05 |
| G2/M Transition | REAC:R-HSA-6 | REAC | 2.68E-05 |
| RNA polymerase II transcribes snRNA genes | REAC:R-HSA-6 | REAC | 2.70E-05 |
| Deadenylation-dependent mRNA decay | REAC:R-HSA-4 | REAC | 2.79E-05 |
| RNA Polymerase III Transcription Initiation | REAC:R-HSA-7 | REAC | 3.02E-05 |
| Transcription of the HIV genome | REAC:R-HSA-1 | REAC | 3.12E-05 |

| | | | |
|---|---|---|---|
| Diseases of DNA repair | REAC:R-HSA-9 | REAC | 3.12E-05 |
| Presynaptic phase of homologous DNA pairing and strai | REAC:R-HSA-5 | REAC | 3.18E-05 |
| Viral Messenger RNA Synthesis | REAC:R-HSA-1 | REAC | 3.18E-05 |
| Cyclin A/B1/B2 associated events during G2/M transitio | REAC:R-HSA-6 | REAC | 3.27E-05 |
| Class I MHC mediated antigen processing & presentatic | REAC:R-HSA-9 | REAC | 3.27E-05 |
| Gap-filling DNA repair synthesis and ligation in TC-NER | REAC:R-HSA-6 | REAC | 3.80E-05 |
| MAP kinase activation | REAC:R-HSA-4 | REAC | 3.80E-05 |
| RAC3 GTPase cycle | REAC:R-HSA-9 | REAC | 3.87E-05 |
| RNA Polymerase III Transcription Initiation From Type 2 | REAC:R-HSA-7 | REAC | 3.95E-05 |
| RHOQ GTPase cycle | REAC:R-HSA-9 | REAC | 4.32E-05 |
| DNA Double-Strand Break Repair | REAC:R-HSA-5 | REAC | 4.81E-05 |
| Nuclear import of Rev protein | REAC:R-HSA-1 | REAC | 5.05E-05 |
| Vpr-mediated nuclear import of PICs | REAC:R-HSA-1 | REAC | 5.05E-05 |
| Signal Transduction | REAC:R-HSA-1 | REAC | 5.05E-05 |
| RNA Polymerase I Transcription Initiation | REAC:R-HSA-7 | REAC | 5.80E-05 |
| Homologous DNA Pairing and Strand Exchange | REAC:R-HSA-5 | REAC | 5.98E-05 |
| SUMOylation of transcription cofactors | REAC:R-HSA-3 | REAC | 5.98E-05 |
| Diseases of DNA Double-Strand Break Repair | REAC:R-HSA-9 | REAC | 5.98E-05 |
| Defective homologous recombination repair (HRR) due | REAC:R-HSA-9 | REAC | 5.98E-05 |
| Golgi-to-ER retrograde transport | REAC:R-HSA-8 | REAC | 6.67E-05 |
| Establishment of Sister Chromatid Cohesion | REAC:R-HSA-2 | REAC | 7.97E-05 |
| G0 and Early G1 | REAC:R-HSA-1 | REAC | 8.46E-05 |
| RNA Polymerase III Transcription Initiation From Type 1 | REAC:R-HSA-7 | REAC | 9.45E-05 |
| Defective TPR may confer susceptibility towards thyroid | REAC:R-HSA-5 | REAC | 9.45E-05 |
| Regulation of Glucokinase by Glucokinase Regulatory P | REAC:R-HSA-1 | REAC | 9.45E-05 |
| Signaling by Hippo | REAC:R-HSA-2 | REAC | 9.45E-05 |
| DNA Damage Bypass | REAC:R-HSA-7 | REAC | 0.00010205 |
| Synthesis of active ubiquitin: roles of E1 and E2 enzyme | REAC:R-HSA-8 | REAC | 0.00010297 |
| RHOA GTPase cycle | REAC:R-HSA-8 | REAC | 0.00010297 |
| G1/S Transition | REAC:R-HSA-6 | REAC | 0.00011346 |
| RUNX1 interacts with co-factors whose precise effect o | REAC:R-HSA-8 | REAC | 0.00011524 |
| RAC2 GTPase cycle | REAC:R-HSA-9 | REAC | 0.00012326 |
| RHOB GTPase cycle | REAC:R-HSA-9 | REAC | 0.0001249 |
| HDR through Homologous Recombination (HRR) | REAC:R-HSA-5 | REAC | 0.00013797 |
| Extension of Telomeres | REAC:R-HSA-1 | REAC | 0.00016724 |
| Toll Like Receptor 3 (TLR3) Cascade | REAC:R-HSA-1 | REAC | 0.0001846 |
| Recruitment of mitotic centrosome proteins and compl | REAC:R-HSA-3 | REAC | 0.00019238 |
| Centrosome maturation | REAC:R-HSA-3 | REAC | 0.00019238 |
| Interleukin-17 signaling | REAC:R-HSA-4 | REAC | 0.00019241 |
| AURKA Activation by TPX2 | REAC:R-HSA-8 | REAC | 0.00019241 |
| Mitotic Telophase/Cytokinesis | REAC:R-HSA-6 | REAC | 0.00020406 |
| Downregulation of SMAD2/3:SMAD4 transcriptional act | REAC:R-HSA-2 | REAC | 0.00021023 |
| tRNA modification in the nucleus and cytosol | REAC:R-HSA-6 | REAC | 0.00021189 |
| Impaired BRCA2 binding to RAD51 | REAC:R-HSA-9 | REAC | 0.00022505 |
| rRNA processing | REAC:R-HSA-7 | REAC | 0.00023268 |
| Transcriptional activity of SMAD2/SMAD3:SMAD4 heter | REAC:R-HSA-2 | REAC | 0.00030049 |
| TRIF(TICAM1)-mediated TLR4 signaling | REAC:R-HSA-9 | REAC | 0.00030917 |
| MyD88-independent TLR4 cascade | REAC:R-HSA-1 | REAC | 0.00030917 |

| | | | |
|---|---|---|---|
| Signaling by EGFR | REAC:R-HSA-1 | REAC | 0.00031892 |
| Polo-like kinase mediated events | REAC:R-HSA-1 | REAC | 0.00035312 |
| Loss of Nlp from mitotic centrosomes | REAC:R-HSA-3 | REAC | 0.00035658 |
| Loss of proteins required for interphase microtubule org | REAC:R-HSA-3 | REAC | 0.00035658 |
| RNA Polymerase II Transcription Elongation | REAC:R-HSA-7 | REAC | 0.00037063 |
| Impaired BRCA2 binding to PALB2 | REAC:R-HSA-9 | REAC | 0.00037063 |
| Formation of RNA Pol II elongation complex | REAC:R-HSA-1 | REAC | 0.00037063 |
| RNA Polymerase III Transcription Initiation From Type 3 | REAC:R-HSA-7 | REAC | 0.00042803 |
| Interactions of Vpr with host cellular proteins | REAC:R-HSA-1 | REAC | 0.00043771 |
| RHOG GTPase cycle | REAC:R-HSA-9 | REAC | 0.00065186 |
| RNA Polymerase III Transcription Termination | REAC:R-HSA-7 | REAC | 0.00083539 |
| Defective HDR through Homologous Recombination Re | REAC:R-HSA-9 | REAC | 0.00086702 |
| Deadenylation of mRNA | REAC:R-HSA-4 | REAC | 0.00086702 |
| Defective HDR through Homologous Recombination Re | REAC:R-HSA-9 | REAC | 0.00086702 |
| Defective homologous recombination repair (HRR) due | REAC:R-HSA-9 | REAC | 0.00086702 |
| Defective homologous recombination repair (HRR) due | REAC:R-HSA-9 | REAC | 0.00086702 |
| RAF activation | REAC:R-HSA-5 | REAC | 0.00087124 |
| Resolution of D-loop Structures through Synthesis-Depe | REAC:R-HSA-5 | REAC | 0.00088311 |
| RNA Polymerase I Transcription Termination | REAC:R-HSA-7 | REAC | 0.00088311 |
| Recruitment of NuMA to mitotic centrosomes | REAC:R-HSA-3 | REAC | 0.00099396 |
| HIV Transcription Initiation | REAC:R-HSA-1 | REAC | 0.00101151 |
| RNA Polymerase II Transcription Initiation And Promoter | REAC:R-HSA-7 | REAC | 0.00101151 |
| RNA Polymerase II Transcription Initiation | REAC:R-HSA-7 | REAC | 0.00101151 |
| RNA Polymerase II Promoter Escape | REAC:R-HSA-7 | REAC | 0.00101151 |
| RNA Polymerase II HIV Promoter Escape | REAC:R-HSA-1 | REAC | 0.00101151 |
| RNA Polymerase II Transcription Pre-Initiation And Prom | REAC:R-HSA-7 | REAC | 0.00101151 |
| Synthesis of DNA | REAC:R-HSA-6 | REAC | 0.00107832 |
| Protein ubiquitination | REAC:R-HSA-8 | REAC | 0.00115607 |
| rRNA processing in the nucleus and cytosol | REAC:R-HSA-8 | REAC | 0.00123921 |
| Condensation of Prometaphase Chromosomes | REAC:R-HSA-2 | REAC | 0.00128663 |
| Transcription of E2F targets under negative control by D | REAC:R-HSA-1 | REAC | 0.00136604 |
| RUNX3 regulates p14-ARF | REAC:R-HSA-8 | REAC | 0.00153262 |
| Lagging Strand Synthesis | REAC:R-HSA-6 | REAC | 0.00154628 |
| Formation of the HIV-1 Early Elongation Complex | REAC:R-HSA-1 | REAC | 0.00162235 |
| Formation of the Early Elongation Complex | REAC:R-HSA-1 | REAC | 0.00162235 |
| Recognition of DNA damage by PCNA-containing replica | REAC:R-HSA-1 | REAC | 0.00168126 |
| Transport to the Golgi and subsequent modification | REAC:R-HSA-9 | REAC | 0.00174582 |
| MyD88 cascade initiated on plasma membrane | REAC:R-HSA-9 | REAC | 0.00174818 |
| Anchoring of the basal body to the plasma membrane | REAC:R-HSA-5 | REAC | 0.00174818 |
| Toll Like Receptor 10 (TLR10) Cascade | REAC:R-HSA-1 | REAC | 0.00174818 |
| Toll Like Receptor 5 (TLR5) Cascade | REAC:R-HSA-1 | REAC | 0.00174818 |
| Formation of Incision Complex in GG-NER | REAC:R-HSA-5 | REAC | 0.00193261 |
| Metabolism of proteins | REAC:R-HSA-3 | REAC | 0.00193635 |
| TP53 Regulates Transcription of DNA Repair Genes | REAC:R-HSA-6 | REAC | 0.00198146 |
| Synthesis of PIPs at the plasma membrane | REAC:R-HSA-1 | REAC | 0.00203245 |
| Formation of TC-NER Pre-Incision Complex | REAC:R-HSA-6 | REAC | 0.00203245 |
| COPI-dependent Golgi-to-ER retrograde traffic | REAC:R-HSA-6 | REAC | 0.00204497 |
| Dual Incision in GG-NER | REAC:R-HSA-5 | REAC | 0.00206304 |

| | | | |
|---|---|---|---|
| SUMOylation of chromatin organization proteins | REAC:R-HSA-4 | REAC | 0.00226695 |
| Maturation of nucleoprotein | REAC:R-HSA-9 | REAC | 0.00248519 |
| Transcription of E2F targets under negative control by p | REAC:R-HSA-1 | REAC | 0.00248519 |
| PI Metabolism | REAC:R-HSA-1 | REAC | 0.00267299 |
| Regulation of RUNX1 Expression and Activity | REAC:R-HSA-8 | REAC | 0.00289198 |
| Butyrate Response Factor 1 (BRF1) binds and destabiliz | REAC:R-HSA-4 | REAC | 0.00289198 |
| Tristetraprolin (TTP, ZFP36) binds and destabilizes mRN | REAC:R-HSA-4 | REAC | 0.00289198 |
| Asparagine N-linked glycosylation | REAC:R-HSA-4 | REAC | 0.00291392 |
| Intra-Golgi traffic | REAC:R-HSA-6 | REAC | 0.00302111 |
| Death Receptor Signaling | REAC:R-HSA-7 | REAC | 0.00303196 |
| Intracellular signaling by second messengers | REAC:R-HSA-9 | REAC | 0.00306836 |
| Cohesin Loading onto Chromatin | REAC:R-HSA-2 | REAC | 0.00307283 |
| Homology Directed Repair | REAC:R-HSA-5 | REAC | 0.00316101 |
| G1/S-Specific Transcription | REAC:R-HSA-6 | REAC | 0.00318318 |
| Resolution of AP sites via the multiple-nucleotide patch | REAC:R-HSA-1 | REAC | 0.00326114 |
| MicroRNA (miRNA) biogenesis | REAC:R-HSA-2 | REAC | 0.00327607 |
| TRAF6 mediated induction of NFkB and MAP kinases up | REAC:R-HSA-9 | REAC | 0.00347814 |
| Toll Like Receptor 9 (TLR9) Cascade | REAC:R-HSA-1 | REAC | 0.00353692 |
| Retrograde transport at the Trans-Golgi-Network | REAC:R-HSA-6 | REAC | 0.00371151 |
| Diseases of mitotic cell cycle | REAC:R-HSA-9 | REAC | 0.00390403 |
| Reversal of alkylation damage by DNA dioxygenases | REAC:R-HSA-7 | REAC | 0.00390403 |
| Oncogenic MAPK signaling | REAC:R-HSA-6 | REAC | 0.0039543 |
| Unwinding of DNA | REAC:R-HSA-1 | REAC | 0.00442477 |
| MyD88 dependent cascade initiated on endosome | REAC:R-HSA-9 | REAC | 0.00452434 |
| Signaling by TGF-beta Receptor Complex | REAC:R-HSA-1 | REAC | 0.00491414 |
| RHOD GTPase cycle | REAC:R-HSA-9 | REAC | 0.00496116 |
| Ca2+ pathway | REAC:R-HSA-4 | REAC | 0.00505856 |
| SARS-CoV-2 activates/modulates innate and adaptive i | REAC:R-HSA-9 | REAC | 0.00515556 |
| Major pathway of rRNA processing in the nucleolus and | REAC:R-HSA-6 | REAC | 0.00564948 |
| mRNA Capping | REAC:R-HSA-7 | REAC | 0.00575151 |
| Toll Like Receptor 7/8 (TLR7/8) Cascade | REAC:R-HSA-1 | REAC | 0.00583774 |
| Translesion synthesis by Y family DNA polymerases byp | REAC:R-HSA-1 | REAC | 0.00626251 |
| RNA Polymerase III Chain Elongation | REAC:R-HSA-7 | REAC | 0.00645812 |
| Abortive elongation of HIV-1 transcript in the absence o | REAC:R-HSA-1 | REAC | 0.00648015 |
| E2F mediated regulation of DNA replication | REAC:R-HSA-1 | REAC | 0.00648015 |
| RHOV GTPase cycle | REAC:R-HSA-9 | REAC | 0.00684062 |
| RHOH GTPase cycle | REAC:R-HSA-9 | REAC | 0.00684062 |
| Macroautophagy | REAC:R-HSA-1 | REAC | 0.00687006 |
| Transcriptional activation of mitochondrial biogenesis | REAC:R-HSA-2 | REAC | 0.00713495 |
| Opioid Signalling | REAC:R-HSA-1 | REAC | 0.00716804 |
| Signaling by TGFB family members | REAC:R-HSA-9 | REAC | 0.00721065 |
| Translocation of SLC2A4 (GLUT4) to the plasma membr | REAC:R-HSA-1 | REAC | 0.00721175 |
| Resolution of D-Loop Structures | REAC:R-HSA-5 | REAC | 0.00736072 |
| RHOBTB GTPase Cycle | REAC:R-HSA-9 | REAC | 0.00736072 |
| Small interfering RNA (siRNA) biogenesis | REAC:R-HSA-4 | REAC | 0.00736072 |
| Formation of HIV elongation complex in the absence of | REAC:R-HSA-1 | REAC | 0.00752339 |
| PKMTs methylate histone lysines | REAC:R-HSA-3 | REAC | 0.00811077 |
| Tat-mediated elongation of the HIV-1 transcript | REAC:R-HSA-1 | REAC | 0.00830348 |

| | | | |
|---|---|---|---|
| HIV Transcription Elongation | REAC:R-HSA-1 | REAC | 0.00830348 |
| Formation of HIV-1 elongation complex containing HIV-1 | REAC:R-HSA-1 | REAC | 0.00830348 |
| RHOF GTPase cycle | REAC:R-HSA-9 | REAC | 0.00830348 |
| Deactivation of the beta-catenin transactivating comple | REAC:R-HSA-3 | REAC | 0.00830348 |
| COPII-mediated vesicle transport | REAC:R-HSA-2 | REAC | 0.00909117 |
| G-protein mediated events | REAC:R-HSA-1 | REAC | 0.00991305 |
| Maturation of nucleoprotein | REAC:R-HSA-9 | REAC | 0.00991305 |
| Intrinsic Pathway for Apoptosis | REAC:R-HSA-1 | REAC | 0.00991305 |
| Phosphorylation of Emi1 | REAC:R-HSA-1 | REAC | 0.01018092 |
| Autophagy | REAC:R-HSA-9 | REAC | 0.01024664 |
| DNA Damage Recognition in GG-NER | REAC:R-HSA-5 | REAC | 0.01028372 |
| Clathrin-mediated endocytosis | REAC:R-HSA-8 | REAC | 0.01057546 |
| Regulation of PTEN gene transcription | REAC:R-HSA-8 | REAC | 0.01076803 |
| Resolution of Abasic Sites (AP sites) | REAC:R-HSA-7 | REAC | 0.01135257 |
| Aberrant regulation of mitotic cell cycle due to RB1 defe | REAC:R-HSA-9 | REAC | 0.01135257 |
| Cytochrome c-mediated apoptotic response | REAC:R-HSA-1 | REAC | 0.01135257 |
| Golgi Cisternae Pericentriolar Stack Reorganization | REAC:R-HSA-1 | REAC | 0.01135257 |
| RHOBTB1 GTPase cycle | REAC:R-HSA-9 | REAC | 0.01146739 |
| ER to Golgi Anterograde Transport | REAC:R-HSA-1 | REAC | 0.01170647 |
| PCNA-Dependent Long Patch Base Excision Repair | REAC:R-HSA-5 | REAC | 0.01200452 |
| Translesion Synthesis by POLH | REAC:R-HSA-1 | REAC | 0.01228543 |
| Telomere C-strand (Lagging Strand) Synthesis | REAC:R-HSA-1 | REAC | 0.01228543 |
| HDR through Homologous Recombination (HRR) or Sing | REAC:R-HSA-5 | REAC | 0.01228543 |
| Processing of Intronless Pre-mRNAs | REAC:R-HSA-7 | REAC | 0.01228543 |
| KSRP (KHSRP) binds and destabilizes mRNA | REAC:R-HSA-4 | REAC | 0.01228543 |
| Resolution of D-loop Structures through Holliday Juncti | REAC:R-HSA-5 | REAC | 0.01228543 |
| mRNA Splicing - Minor Pathway | REAC:R-HSA-7 | REAC | 0.01383041 |
| Kinesins | REAC:R-HSA-9 | REAC | 0.01451168 |
| RUNX2 regulates bone development | REAC:R-HSA-8 | REAC | 0.0151193 |
| Regulation of MECP2 expression and activity | REAC:R-HSA-9 | REAC | 0.0151193 |
| Transcriptional Regulation by VENTX | REAC:R-HSA-8 | REAC | 0.01516054 |
| G1 Phase | REAC:R-HSA-6 | REAC | 0.01548656 |
| Cyclin D associated events in G1 | REAC:R-HSA-6 | REAC | 0.01548656 |
| Processing of Capped Intronless Pre-mRNA | REAC:R-HSA-7 | REAC | 0.01650559 |
| Notch-HLH transcription pathway | REAC:R-HSA-3 | REAC | 0.01650559 |
| HuR (ELAVL1) binds and stabilizes mRNA | REAC:R-HSA-4 | REAC | 0.01705316 |
| Miro GTPase Cycle | REAC:R-HSA-9 | REAC | 0.01705316 |
| DNA Damage Reversal | REAC:R-HSA-7 | REAC | 0.01705316 |
| Apoptotic execution phase | REAC:R-HSA-7 | REAC | 0.01705316 |
| RNA Pol II CTD phosphorylation and interaction with CE | REAC:R-HSA-7 | REAC | 0.0178708 |
| RNA Pol II CTD phosphorylation and interaction with CE | REAC:R-HSA-1 | REAC | 0.0178708 |
| RHO GTPases Activate WASPs and WAVEs | REAC:R-HSA-5 | REAC | 0.01877617 |
| TP53 Regulates Transcription of Cell Cycle Genes | REAC:R-HSA-6 | REAC | 0.01916855 |
| PLC beta mediated events | REAC:R-HSA-1 | REAC | 0.01916855 |
| Metabolism of nucleotides | REAC:R-HSA-1 | REAC | 0.02053465 |
| JNK (c-Jun kinases) phosphorylation and activation med | REAC:R-HSA-4 | REAC | 0.02102625 |
| Processing and activation of SUMO | REAC:R-HSA-3 | REAC | 0.02152026 |
| MASTL Facilitates Mitotic Progression | REAC:R-HSA-2 | REAC | 0.02152026 |

| | | | |
|---|---|---|---|
| EPHB-mediated forward signaling | REAC:R-HSA-3 | REAC | 0.0219238 |
| G2/M Checkpoints | REAC:R-HSA-6 | REAC | 0.02303372 |
| Toll Like Receptor TLR6:TLR2 Cascade | REAC:R-HSA-1 | REAC | 0.02303372 |
| Chromosome Maintenance | REAC:R-HSA-7 | REAC | 0.02303372 |
| MyD88:MAL(TIRAP) cascade initiated on plasma memb | REAC:R-HSA-1 | REAC | 0.02303372 |
| MAPK targets/ Nuclear events mediated by MAP kinases | REAC:R-HSA-4 | REAC | 0.0230741 |
| Disease | REAC:R-HSA-1 | REAC | 0.02335228 |
| APC truncation mutants have impaired AXIN binding | REAC:R-HSA-5 | REAC | 0.02378749 |
| AXIN missense mutants destabilize the destruction con | REAC:R-HSA-5 | REAC | 0.02378749 |
| Truncations of AMER1 destabilize the destruction comp | REAC:R-HSA-5 | REAC | 0.02378749 |
| Signaling by AMER1 mutants | REAC:R-HSA-4 | REAC | 0.02378749 |
| Signaling by APC mutants | REAC:R-HSA-4 | REAC | 0.02378749 |
| Leading Strand Synthesis | REAC:R-HSA-6 | REAC | 0.02378749 |
| Synthesis of IP2, IP, and Ins in the cytosol | REAC:R-HSA-1 | REAC | 0.02378749 |
| Signaling by AXIN mutants | REAC:R-HSA-4 | REAC | 0.02378749 |
| Polymerase switching | REAC:R-HSA-6 | REAC | 0.02378749 |
| p75 NTR receptor-mediated signalling | REAC:R-HSA-1 | REAC | 0.02422248 |
| Mitochondrial translation elongation | REAC:R-HSA-5 | REAC | 0.02516106 |
| G2/M DNA replication checkpoint | REAC:R-HSA-6 | REAC | 0.02521527 |
| N-glycan trimming and elongation in the cis-Golgi | REAC:R-HSA-9 | REAC | 0.02521527 |
| Transcriptional regulation of white adipocyte differentia | REAC:R-HSA-3 | REAC | 0.02523687 |
| SMAD2/SMAD3:SMAD4 heterotrimer regulates transcrip | REAC:R-HSA-2 | REAC | 0.02674232 |
| PPARA activates gene expression | REAC:R-HSA-1 | REAC | 0.02711073 |
| PIP3 activates AKT signaling | REAC:R-HSA-1 | REAC | 0.02711073 |
| Neddylation | REAC:R-HSA-8 | REAC | 0.02776167 |
| DAG and IP3 signaling | REAC:R-HSA-1 | REAC | 0.03015376 |
| MTOR signalling | REAC:R-HSA-1 | REAC | 0.03015376 |
| Gap-filling DNA repair synthesis and ligation in GG-NER | REAC:R-HSA-5 | REAC | 0.03057029 |
| Deubiquitination | REAC:R-HSA-5 | REAC | 0.0307881 |
| Glycolysis | REAC:R-HSA-7 | REAC | 0.03257509 |
| Regulation of lipid metabolism by PPARalpha | REAC:R-HSA-4 | REAC | 0.03282991 |
| Phospholipid metabolism | REAC:R-HSA-1 | REAC | 0.03320773 |
| Termination of translesion DNA synthesis | REAC:R-HSA-5 | REAC | 0.03358604 |
| Signaling by FGFR1 in disease | REAC:R-HSA-5 | REAC | 0.03358604 |
| RHOU GTPase cycle | REAC:R-HSA-9 | REAC | 0.0338081 |
| PTEN Regulation | REAC:R-HSA-6 | REAC | 0.03493639 |
| Signaling by FGFR in disease | REAC:R-HSA-1 | REAC | 0.03552916 |
| Inactivation of APC/C via direct inhibition of the APC/C c | REAC:R-HSA-1 | REAC | 0.03658415 |
| Inhibition of the proteolytic activity of APC/C required fo | REAC:R-HSA-1 | REAC | 0.03658415 |
| Regulation of TP53 Activity through Acetylation | REAC:R-HSA-6 | REAC | 0.03761154 |
| RHOT2 GTPase cycle | REAC:R-HSA-9 | REAC | 0.03834453 |
| SMAC(DIABLO)-mediated dissociation of IAP:caspase c | REAC:R-HSA-1 | REAC | 0.03834453 |
| SMAC (DIABLO) binds to IAPs | REAC:R-HSA-1 | REAC | 0.03834453 |
| CREB phosphorylation | REAC:R-HSA-1 | REAC | 0.03834453 |
| Post-transcriptional silencing by small RNAs | REAC:R-HSA-4 | REAC | 0.03834453 |
| SMAC, XIAP-regulated apoptotic response | REAC:R-HSA-1 | REAC | 0.03834453 |
| Activation of NIMA Kinases NEK9, NEK6, NEK7 | REAC:R-HSA-2 | REAC | 0.03834453 |
| Toll Like Receptor 2 (TLR2) Cascade | REAC:R-HSA-1 | REAC | 0.03836227 |

| | | | |
|---|---|---|---|
| Toll Like Receptor TLR1:TLR2 Cascade | REAC:R-HSA-1 | REAC | 0.03836227 |
| Signaling by BRAF and RAF1 fusions | REAC:R-HSA-6 | REAC | 0.04053063 |
| Cytosolic sensors of pathogen-associated DNA | REAC:R-HSA-1 | REAC | 0.04053063 |
| Negative regulation of MAPK pathway | REAC:R-HSA-5 | REAC | 0.04077161 |
| Signaling by BMP | REAC:R-HSA-2 | REAC | 0.04123807 |
| Cellular responses to stress | REAC:R-HSA-2 | REAC | 0.041461 |
| Translesion synthesis by POLK | REAC:R-HSA-5 | REAC | 0.04180876 |
| Mitochondrial translation termination | REAC:R-HSA-5 | REAC | 0.04212386 |
| Heme signaling | REAC:R-HSA-9 | REAC | 0.04285283 |
| CTNNB1 S45 mutants aren't phosphorylated | REAC:R-HSA-5 | REAC | 0.04309907 |
| Signaling by GSK3beta mutants | REAC:R-HSA-5 | REAC | 0.04309907 |
| CTNNB1 S37 mutants aren't phosphorylated | REAC:R-HSA-5 | REAC | 0.04309907 |
| MAPK3 (ERK1) activation | REAC:R-HSA-1 | REAC | 0.04309907 |
| MET receptor recycling | REAC:R-HSA-8 | REAC | 0.04309907 |
| CTNNB1 S33 mutants aren't phosphorylated | REAC:R-HSA-5 | REAC | 0.04309907 |
| E2F-enabled inhibition of pre-replication complex forma | REAC:R-HSA-1 | REAC | 0.04309907 |
| CTNNB1 T41 mutants aren't phosphorylated | REAC:R-HSA-5 | REAC | 0.04309907 |
| Signaling by CTNNB1 phospho-site mutants | REAC:R-HSA-4 | REAC | 0.04309907 |
| Regulation of PTEN mRNA translation | REAC:R-HSA-8 | REAC | 0.04309907 |
| SARS-CoV-1 targets host intracellular signalling and reg | REAC:R-HSA-9 | REAC | 0.04309907 |
| Processive synthesis on the lagging strand | REAC:R-HSA-6 | REAC | 0.04309907 |
| Translation of Structural Proteins | REAC:R-HSA-9 | REAC | 0.04391614 |
| Constitutive Signaling by EGFRvIII | REAC:R-HSA-5 | REAC | 0.04448215 |
| Cytosolic iron-sulfur cluster assembly | REAC:R-HSA-2 | REAC | 0.04448215 |
| Regulation of NPAS4 gene expression | REAC:R-HSA-9 | REAC | 0.04448215 |
| Signaling by EGFRvIII in Cancer | REAC:R-HSA-5 | REAC | 0.04448215 |
| Formation of apoptosome | REAC:R-HSA-1 | REAC | 0.044524 |
| Interleukin-6 signaling | REAC:R-HSA-1 | REAC | 0.044524 |
| Regulation of the apoptosome activity | REAC:R-HSA-9 | REAC | 0.044524 |
| ESR-mediated signaling | REAC:R-HSA-8 | REAC | 0.04706812 |
| Toll Like Receptor 4 (TLR4) Cascade | REAC:R-HSA-1 | REAC | 0.04743621 |
| Mitochondrial translation | REAC:R-HSA-5 | REAC | 0.04784331 |
| NOTCH3 Intracellular Domain Regulates Transcription | REAC:R-HSA-9 | REAC | 0.04900606 |
| SUMOylation of intracellular receptors | REAC:R-HSA-4 | REAC | 0.04900606 |
| Estrogen-dependent nuclear events downstream of ESF | REAC:R-HSA-9 | REAC | 0.04900606 |

| term_size | intersection_size |
|---|---|
| 5440 | 2119 |
| 5591 | 2160 |
| 3262 | 1414 |
| 1819 | 911 |
| 3613 | 1486 |
| 4063 | 1620 |
| 4836 | 1843 |
| 3031 | 1283 |
| 1276 | 671 |
| 6250 | 2248 |
| 3001 | 1264 |
| 5717 | 2086 |
| 4912 | 1836 |
| 3763 | 1485 |
| 895 | 506 |
| 3420 | 1368 |
| 5986 | 2131 |
| 7158 | 2456 |
| 4105 | 1570 |
| 2052 | 916 |
| 4116 | 1571 |
| 3711 | 1446 |
| 4027 | 1541 |
| 6631 | 2298 |
| 745 | 429 |
| 4264 | 1598 |
| 12680 | 3875 |
| 12287 | 3775 |
| 1852 | 829 |
| 3601 | 1391 |
| 3569 | 1378 |
| 11738 | 3630 |
| 6673 | 2284 |
| 1040 | 537 |
| 640 | 376 |
| 1770 | 791 |
| 3346 | 1299 |
| 896 | 476 |
| 3464 | 1326 |
| 3446 | 1320 |
| 1371 | 639 |
| 2425 | 991 |
| 1105 | 541 |
| 1706 | 747 |
| 1972 | 834 |
| 1703 | 744 |

| | |
|---|---|
| 656 | 362 |
| 11995 | 3632 |
| 3621 | 1340 |
| 604 | 340 |
| 635 | 350 |
| 438 | 269 |
| 2685 | 1045 |
| 2603 | 1019 |
| 915 | 454 |
| 2706 | 1049 |
| 724 | 379 |
| 1021 | 487 |
| 12505 | 3734 |
| 1338 | 596 |
| 2202 | 880 |
| 537 | 302 |
| 2668 | 1023 |
| 425 | 254 |
| 2691 | 1027 |
| 5813 | 1942 |
| 970 | 461 |
| 5781 | 1931 |
| 1173 | 531 |
| 5654 | 1894 |
| 858 | 417 |
| 3210 | 1175 |
| 2102 | 832 |
| 5541 | 1853 |
| 5526 | 1845 |
| 432 | 250 |
| 322 | 203 |
| 816 | 396 |
| 2567 | 971 |
| 2742 | 1024 |
| 2731 | 1018 |
| 5874 | 1933 |
| 8361 | 2613 |
| 1588 | 655 |
| 521 | 279 |
| 263 | 172 |
| 1911 | 752 |
| 774 | 371 |
| 5520 | 1817 |
| 1424 | 592 |
| 230 | 152 |
| 1463 | 597 |
| 496 | 260 |
| 1253 | 526 |

| | |
|---|---|
| 2556 | 937 |
| 433 | 235 |
| 664 | 321 |
| 4643 | 1547 |
| 6453 | 2053 |
| 3010 | 1070 |
| 224 | 146 |
| 1026 | 444 |
| 248 | 156 |
| 281 | 170 |
| 3939 | 1337 |
| 279 | 169 |
| 2596 | 940 |
| 318 | 185 |
| 5899 | 1890 |
| 2571 | 930 |
| 1512 | 600 |
| 620 | 298 |
| 309 | 179 |
| 512 | 258 |
| 2358 | 859 |
| 620 | 296 |
| 332 | 187 |
| 189 | 126 |
| 495 | 249 |
| 334 | 186 |
| 334 | 186 |
| 2074 | 767 |
| 3437 | 1174 |
| 686 | 316 |
| 599 | 285 |
| 7394 | 2280 |
| 302 | 171 |
| 1333 | 530 |
| 8151 | 2478 |
| 3443 | 1169 |
| 609 | 286 |
| 1579 | 607 |
| 1347 | 533 |
| 968 | 409 |
| 730 | 328 |
| 447 | 226 |
| 971 | 409 |
| 1547 | 595 |
| 659 | 300 |
| 1761 | 656 |
| 1924 | 706 |
| 261 | 150 |

| | |
|---|---|
| 1116 | 451 |
| 12170 | 3512 |
| 486 | 236 |
| 670 | 301 |
| 691 | 308 |
| 162 | 107 |
| 14261 | 4035 |
| 1799 | 664 |
| 302 | 165 |
| 388 | 198 |
| 306 | 166 |
| 4881 | 1560 |
| 4350 | 1408 |
| 2455 | 859 |
| 1229 | 482 |
| 3973 | 1300 |
| 3855 | 1266 |
| 1128 | 449 |
| 197 | 120 |
| 216 | 128 |
| 260 | 146 |
| 13768 | 3902 |
| 3455 | 1148 |
| 995 | 404 |
| 129 | 89 |
| 169 | 107 |
| 1001 | 404 |
| 197 | 118 |
| 1766 | 643 |
| 1913 | 687 |
| 409 | 200 |
| 2531 | 871 |
| 505 | 234 |
| 286 | 153 |
| 687 | 297 |
| 3226 | 1073 |
| 3240 | 1077 |
| 251 | 139 |
| 176 | 108 |
| 300 | 158 |
| 181 | 110 |
| 205 | 120 |
| 158 | 100 |
| 158 | 100 |
| 257 | 141 |
| 161 | 101 |
| 1984 | 704 |
| 334 | 170 |

| | |
|---|---|
| 1988 | 705 |
| 8692 | 2572 |
| 206 | 119 |
| 410 | 197 |
| 168 | 103 |
| 1346 | 506 |
| 526 | 238 |
| 439 | 207 |
| 12881 | 3657 |
| 349 | 174 |
| 378 | 184 |
| 301 | 155 |
| 1184 | 452 |
| 1399 | 519 |
| 568 | 250 |
| 496 | 225 |
| 889 | 357 |
| 506 | 228 |
| 1253 | 472 |
| 533 | 237 |
| 178 | 105 |
| 1227 | 463 |
| 953 | 376 |
| 229 | 125 |
| 1290 | 481 |
| 567 | 247 |
| 1861 | 655 |
| 268 | 139 |
| 182 | 105 |
| 1600 | 574 |
| 128 | 82 |
| 986 | 383 |
| 587 | 252 |
| 124 | 80 |
| 400 | 187 |
| 189 | 107 |
| 719 | 295 |
| 815 | 326 |
| 315 | 155 |
| 104 | 70 |
| 515 | 225 |
| 143 | 87 |
| 189 | 106 |
| 197 | 109 |
| 2819 | 928 |
| 94 | 65 |
| 1340 | 489 |
| 8228 | 2417 |

| | |
|---|---|
| 154 | 91 |
| 187 | 104 |
| 134 | 82 |
| 8055 | 2367 |
| 116 | 74 |
| 123 | 77 |
| 94 | 64 |
| 577 | 243 |
| 577 | 243 |
| 616 | 256 |
| 181 | 101 |
| 135 | 82 |
| 1742 | 607 |
| 723 | 291 |
| 131 | 80 |
| 887 | 343 |
| 2684 | 882 |
| 8976 | 2607 |
| 213 | 113 |
| 91 | 62 |
| 102 | 67 |
| 939 | 359 |
| 752 | 299 |
| 191 | 104 |
| 1097 | 408 |
| 19708 | 5268 |
| 133 | 80 |
| 133 | 80 |
| 138 | 82 |
| 8280 | 2418 |
| 1398 | 499 |
| 232 | 119 |
| 97 | 64 |
| 1613 | 563 |
| 1570 | 550 |
| 1398 | 498 |
| 747 | 295 |
| 118 | 73 |
| 210 | 110 |
| 509 | 216 |
| 145 | 84 |
| 1648 | 572 |
| 133 | 79 |
| 119 | 73 |
| 7499 | 2204 |
| 90 | 60 |
| 225 | 115 |
| 241 | 121 |

| | |
|---|---|
| 146 | 84 |
| 5959 | 1786 |
| 86 | 58 |
| 266 | 130 |
| 174 | 95 |
| 154 | 87 |
| 137 | 80 |
| 4367 | 1346 |
| 163 | 90 |
| 4366 | 1345 |
| 181 | 97 |
| 1194 | 431 |
| 185 | 98 |
| 332 | 152 |
| 104 | 65 |
| 95 | 61 |
| 361 | 162 |
| 140 | 80 |
| 1514 | 526 |
| 2824 | 908 |
| 6447 | 1911 |
| 986 | 365 |
| 93 | 60 |
| 138 | 79 |
| 186 | 98 |
| 1506 | 523 |
| 80 | 54 |
| 2838 | 911 |
| 6537 | 1934 |
| 105 | 65 |
| 103 | 64 |
| 74 | 51 |
| 8525 | 2464 |
| 108 | 66 |
| 70 | 49 |
| 594 | 239 |
| 1432 | 499 |
| 81 | 54 |
| 242 | 118 |
| 2773 | 889 |
| 911 | 339 |
| 138 | 78 |
| 2018 | 671 |
| 1463 | 507 |
| 131 | 75 |
| 131 | 75 |
| 766 | 293 |
| 100 | 62 |

| | |
|---|---|
| 495 | 205 |
| 112 | 67 |
| 667 | 261 |
| 139 | 78 |
| 787 | 299 |
| 247 | 119 |
| 87 | 56 |
| 120 | 70 |
| 325 | 146 |
| 454 | 190 |
| 156 | 84 |
| 151 | 82 |
| 141 | 78 |
| 452 | 189 |
| 306 | 139 |
| 250 | 119 |
| 429 | 181 |
| 54 | 40 |
| 229 | 111 |
| 1328 | 462 |
| 80 | 52 |
| 101 | 61 |
| 602 | 237 |
| 1431 | 492 |
| 90 | 56 |
| 53 | 39 |
| 1678 | 564 |
| 95 | 58 |
| 160 | 84 |
| 523 | 210 |
| 192 | 96 |
| 773 | 289 |
| 71 | 47 |
| 787 | 293 |
| 717 | 271 |
| 48 | 36 |
| 116 | 66 |
| 667 | 255 |
| 92 | 56 |
| 362 | 155 |
| 327 | 143 |
| 222 | 106 |
| 168 | 86 |
| 1014 | 362 |
| 965 | 347 |
| 163 | 84 |
| 163 | 84 |
| 28 | 25 |

| | |
|---|---|
| 110 | 63 |
| 652 | 249 |
| 68 | 45 |
| 103 | 60 |
| 45 | 34 |
| 1035 | 367 |
| 711 | 267 |
| 1132 | 396 |
| 62 | 42 |
| 62 | 42 |
| 62 | 42 |
| 80 | 50 |
| 1133 | 396 |
| 134 | 72 |
| 2637 | 833 |
| 142 | 75 |
| 544 | 213 |
| 1573 | 526 |
| 56 | 39 |
| 218 | 103 |
| 52 | 37 |
| 1167 | 405 |
| 135 | 72 |
| 2274 | 728 |
| 72 | 46 |
| 264 | 119 |
| 61 | 41 |
| 103 | 59 |
| 364 | 153 |
| 59 | 40 |
| 681 | 255 |
| 116 | 64 |
| 490 | 194 |
| 324 | 139 |
| 805 | 293 |
| 238 | 109 |
| 161 | 81 |
| 1561 | 519 |
| 967 | 342 |
| 736 | 271 |
| 200 | 95 |
| 838 | 302 |
| 966 | 341 |
| 98 | 56 |
| 1971 | 636 |
| 52 | 36 |
| 305 | 131 |
| 1576 | 521 |

| | |
|---|---|
| 199 | 94 |
| 48 | 34 |
| 48 | 34 |
| 48 | 34 |
| 48 | 34 |
| 48 | 34 |
| 383 | 157 |
| 89 | 52 |
| 205 | 96 |
| 463 | 183 |
| 44 | 32 |
| 167 | 82 |
| 55 | 37 |
| 27 | 23 |
| 417 | 167 |
| 22 | 20 |
| 93 | 53 |
| 1496 | 495 |
| 357 | 147 |
| 106 | 58 |
| 33 | 26 |
| 33 | 26 |
| 41 | 30 |
| 37 | 28 |
| 910 | 320 |
| 19 | 18 |
| 287 | 123 |
| 365 | 149 |
| 184 | 87 |
| 311 | 131 |
| 112 | 60 |
| 112 | 60 |
| 224 | 101 |
| 250 | 110 |
| 97 | 54 |
| 50 | 34 |
| 50 | 34 |
| 50 | 34 |
| 400 | 160 |
| 59 | 38 |
| 73 | 44 |
| 191 | 89 |
| 304 | 128 |
| 118 | 62 |
| 46 | 32 |
| 46 | 32 |
| 46 | 32 |
| 676 | 247 |

| | |
|---|---|
| 263 | 114 |
| 1411 | 467 |
| 44 | 31 |
| 243 | 107 |
| 347 | 142 |
| 671 | 245 |
| 140 | 70 |
| 2006 | 638 |
| 36 | 27 |
| 86 | 49 |
| 187 | 87 |
| 135 | 68 |
| 74 | 44 |
| 74 | 44 |
| 154 | 75 |
| 130 | 66 |
| 609 | 225 |
| 298 | 125 |
| 451 | 175 |
| 907 | 316 |
| 185 | 86 |
| 72 | 43 |
| 254 | 110 |
| 483 | 185 |
| 47 | 32 |
| 290 | 122 |
| 56 | 36 |
| 92 | 51 |
| 105 | 56 |
| 100 | 54 |
| 85 | 48 |
| 61 | 38 |
| 121 | 62 |
| 37 | 27 |
| 233 | 102 |
| 83 | 47 |
| 111 | 58 |
| 106 | 56 |
| 171 | 80 |
| 57 | 36 |
| 48 | 32 |
| 661 | 239 |
| 2799 | 857 |
| 69 | 41 |
| 22 | 19 |
| 1778 | 568 |
| 147 | 71 |
| 84 | 47 |

| | |
|---|---|
| 139 | 68 |
| 139 | 68 |
| 153 | 73 |
| 72 | 42 |
| 239 | 103 |
| 655 | 236 |
| 113 | 58 |
| 357 | 142 |
| 620 | 225 |
| 964 | 329 |
| 34 | 25 |
| 75 | 43 |
| 537 | 199 |
| 881 | 304 |
| 63 | 38 |
| 68 | 40 |
| 68 | 40 |
| 93 | 50 |
| 54 | 34 |
| 3039 | 920 |
| 133 | 65 |
| 66 | 39 |
| 117 | 59 |
| 604 | 219 |
| 43 | 29 |
| 360 | 142 |
| 457 | 173 |
| 120 | 60 |
| 120 | 60 |
| 1060 | 356 |
| 21 | 18 |
| 245 | 104 |
| 759 | 266 |
| 499 | 186 |
| 64 | 38 |
| 134 | 65 |
| 309 | 125 |
| 148 | 70 |
| 37 | 26 |
| 672 | 239 |
| 2846 | 864 |
| 35 | 25 |
| 35 | 25 |
| 223 | 96 |
| 223 | 96 |
| 1091 | 364 |
| 82 | 45 |
| 27 | 21 |

| | |
|---|---|
| 31 | 23 |
| 366 | 143 |
| 1777 | 562 |
| 95 | 50 |
| 53 | 33 |
| 103 | 53 |
| 90 | 48 |
| 65 | 38 |
| 141 | 67 |
| 874 | 299 |
| 535 | 196 |
| 269 | 111 |
| 175 | 79 |
| 1091 | 363 |
| 18 | 16 |
| 785 | 272 |
| 147 | 69 |
| 842 | 289 |
| 2955 | 892 |
| 63 | 37 |
| 63 | 37 |
| 187 | 83 |
| 288 | 117 |
| 83 | 45 |
| 49 | 31 |
| 56 | 34 |
| 56 | 34 |
| 96 | 50 |
| 356 | 139 |
| 173 | 78 |
| 142 | 67 |
| 341 | 134 |
| 429 | 162 |
| 672 | 237 |
| 36 | 25 |
| 607 | 217 |
| 1130 | 373 |
| 102 | 52 |
| 364 | 141 |
| 293 | 118 |
| 355 | 138 |
| 15 | 14 |
| 15 | 14 |
| 192 | 84 |
| 24 | 19 |
| 30 | 22 |
| 334 | 131 |
| 616 | 219 |

| | |
|---|---|
| 119 | 58 |
| 872 | 296 |
| 222 | 94 |
| 270 | 110 |
| 222 | 94 |
| 181 | 80 |
| 610 | 217 |
| 133 | 63 |
| 67 | 38 |
| 376 | 144 |
| 208 | 89 |
| 48 | 30 |
| 48 | 30 |
| 156 | 71 |
| 232 | 97 |
| 80 | 43 |
| 440 | 164 |
| 101 | 51 |
| 65 | 37 |
| 65 | 37 |
| 599 | 213 |
| 53 | 32 |
| 53 | 32 |
| 151 | 69 |
| 35 | 24 |
| 35 | 24 |
| 68 | 38 |
| 44 | 28 |
| 44 | 28 |
| 585 | 208 |
| 210 | 89 |
| 21 | 17 |
| 21 | 17 |
| 211 | 89 |
| 428 | 159 |
| 25 | 19 |
| 40 | 26 |
| 14 | 13 |
| 14 | 13 |
| 47 | 29 |
| 125 | 59 |
| 598 | 211 |
| 194 | 83 |
| 142 | 65 |
| 38 | 25 |
| 128 | 60 |
| 376 | 142 |
| 11 | 11 |

| | |
|---|---|
| 57 | 33 |
| 264 | 106 |
| 437 | 161 |
| 207 | 87 |
| 80 | 42 |
| 50 | 30 |
| 166 | 73 |
| 43 | 27 |
| 43 | 27 |
| 88 | 45 |
| 34 | 23 |
| 274 | 109 |
| 704 | 242 |
| 70 | 38 |
| 60 | 34 |
| 60 | 34 |
| 247 | 100 |
| 91 | 46 |
| 138 | 63 |
| 48 | 29 |
| 372 | 140 |
| 864 | 289 |
| 30 | 21 |
| 53 | 31 |
| 1128 | 366 |
| 600 | 210 |
| 424 | 156 |
| 28 | 20 |
| 279 | 110 |
| 215 | 89 |
| 46 | 28 |
| 46 | 28 |
| 849 | 284 |
| 76 | 40 |
| 428 | 157 |
| 320 | 123 |
| 84 | 43 |
| 1034 | 338 |
| 177 | 76 |
| 752 | 255 |
| 37 | 24 |
| 151 | 67 |
| 151 | 67 |
| 204 | 85 |
| 117 | 55 |
| 13 | 12 |
| 35 | 23 |
| 140 | 63 |

| | |
|---|---|
| 417 | 153 |
| 238 | 96 |
| 2683 | 803 |
| 934 | 308 |
| 202 | 84 |
| 158 | 69 |
| 161 | 70 |
| 245 | 98 |
| 164 | 71 |
| 47 | 28 |
| 267 | 105 |
| 124 | 57 |
| 31 | 21 |
| 258 | 102 |
| 75 | 39 |
| 75 | 39 |
| 15 | 13 |
| 15 | 13 |
| 15 | 13 |
| 156 | 68 |
| 274 | 107 |
| 45 | 27 |
| 70 | 37 |
| 105 | 50 |
| 65 | 35 |
| 97 | 47 |
| 55 | 31 |
| 17 | 14 |
| 17 | 14 |
| 1033 | 335 |
| 128 | 58 |
| 25 | 18 |
| 36 | 23 |
| 43 | 26 |
| 396 | 145 |
| 220 | 89 |
| 23 | 17 |
| 19 | 15 |
| 92 | 45 |
| 63 | 34 |
| 63 | 34 |
| 95 | 46 |
| 617 | 212 |
| 511 | 180 |
| 87 | 43 |
| 98 | 47 |
| 1189 | 379 |
| 555 | 193 |

| | |
|---|---|
| 90 | 44 |
| 197 | 81 |
| 185 | 77 |
| 46 | 27 |
| 46 | 27 |
| 46 | 27 |
| 46 | 27 |
| 74 | 38 |
| 32 | 21 |
| 141 | 62 |
| 51 | 29 |
| 51 | 29 |
| 93 | 45 |
| 121 | 55 |
| 39 | 24 |
| 39 | 24 |
| 225 | 90 |
| 124 | 56 |
| 77 | 39 |
| 96 | 46 |
| 159 | 68 |
| 130 | 58 |
| 88 | 43 |
| 44 | 26 |
| 88 | 43 |
| 113 | 52 |
| 136 | 60 |
| 139 | 61 |
| 37 | 23 |
| 37 | 23 |
| 37 | 23 |
| 37 | 23 |
| 304 | 115 |
| 245 | 96 |
| 128 | 57 |
| 42 | 25 |
| 42 | 25 |
| 62 | 33 |
| 261 | 101 |
| 321 | 120 |
| 35 | 22 |
| 14 | 12 |
| 14 | 12 |
| 70 | 36 |
| 47 | 27 |
| 721 | 241 |
| 52 | 29 |
| 215 | 86 |

| | |
|---|---|
| 9 | 9 |
| 206 | 83 |
| 206 | 83 |
| 597 | 204 |
| 155 | 66 |
| 65 | 34 |
| 158 | 67 |
| 92 | 44 |
| 501 | 175 |
| 40 | 24 |
| 16 | 13 |
| 16 | 13 |
| 22 | 16 |
| 228 | 90 |
| 33 | 21 |
| 60 | 32 |
| 84 | 41 |
| 374 | 136 |
| 219 | 87 |
| 20 | 15 |
| 18 | 14 |
| 18 | 14 |
| 18 | 14 |
| 45 | 26 |
| 45 | 26 |
| 55 | 30 |
| 307 | 115 |
| 241 | 94 |
| 1183 | 374 |
| 63 | 33 |
| 1709 | 523 |
| 450 | 159 |
| 121 | 54 |
| 31 | 20 |
| 630 | 213 |
| 107 | 49 |
| 107 | 49 |
| 705 | 235 |
| 705 | 235 |
| 43 | 25 |
| 48 | 27 |
| 199 | 80 |
| 96 | 45 |
| 190 | 77 |
| 190 | 77 |
| 29 | 19 |
| 905 | 293 |
| 122 | 54 |

| | |
|---|---|
| 122 | 54 |
| 449 | 158 |
| 56 | 30 |
| 56 | 30 |
| 128 | 56 |
| 41 | 24 |
| 41 | 24 |
| 519 | 179 |
| 27 | 18 |
| 27 | 18 |
| 194 | 78 |
| 46 | 26 |
| 603 | 204 |
| 6946 | 1941 |
| 11 | 10 |
| 11 | 10 |
| 11 | 10 |
| 378 | 136 |
| 152 | 64 |
| 173 | 71 |
| 83 | 40 |
| 34 | 21 |
| 298 | 111 |
| 204 | 81 |
| 103 | 47 |
| 415 | 147 |
| 195 | 78 |
| 135 | 58 |
| 49 | 27 |
| 44 | 25 |
| 32 | 20 |
| 109 | 49 |
| 92 | 43 |
| 150 | 63 |
| 150 | 63 |
| 406 | 144 |
| 745 | 245 |
| 211 | 83 |
| 211 | 83 |
| 13 | 11 |
| 13 | 11 |
| 57 | 30 |
| 322 | 118 |
| 84 | 40 |
| 1225 | 383 |
| 121 | 53 |
| 65 | 33 |
| 65 | 33 |

| | |
|---|---|
| 65 | 33 |
| 345 | 125 |
| 19 | 14 |
| 313 | 115 |
| 15 | 12 |
| 15 | 12 |
| 15 | 12 |
| 42 | 24 |
| 42 | 24 |
| 47 | 26 |
| 169 | 69 |
| 611 | 205 |
| 166 | 68 |
| 17 | 13 |
| 527 | 180 |
| 157 | 65 |
| 996 | 317 |
| 8 | 8 |
| 8 | 8 |
| 8 | 8 |
| 8 | 8 |
| 8 | 8 |
| 79 | 38 |
| 113 | 50 |
| 71 | 35 |
| 71 | 35 |
| 63 | 32 |
| 279 | 104 |
| 28 | 18 |
| 28 | 18 |
| 125 | 54 |
| 276 | 103 |
| 10565 | 2888 |
| 522 | 178 |
| 134 | 57 |
| 161 | 66 |
| 140 | 59 |
| 204 | 80 |
| 105 | 47 |
| 315 | 115 |
| 66 | 33 |
| 248 | 94 |
| 77 | 37 |
| 463 | 160 |
| 26 | 17 |
| 26 | 17 |
| 26 | 17 |
| 1035 | 327 |

| | |
|---|---|
| 258 | 97 |
| 117 | 51 |
| 625 | 208 |
| 381 | 135 |
| 61 | 31 |
| 2010 | 601 |
| 129 | 55 |
| 300 | 110 |
| 48 | 26 |
| 48 | 26 |
| 159 | 65 |
| 388 | 137 |
| 141 | 59 |
| 43 | 24 |
| 109 | 48 |
| 89 | 41 |
| 31 | 19 |
| 31 | 19 |
| 169 | 68 |
| 67 | 33 |
| 133 | 56 |
| 10 | 9 |
| 10 | 9 |
| 10 | 9 |
| 10 | 9 |
| 10 | 9 |
| 36 | 21 |
| 463 | 159 |
| 70 | 34 |
| 101 | 45 |
| 101 | 45 |
| 54 | 28 |
| 54 | 28 |
| 62 | 31 |
| 29 | 18 |
| 29 | 18 |
| 248 | 93 |
| 73 | 35 |
| 73 | 35 |
| 20 | 14 |
| 1038 | 326 |
| 49 | 26 |
| 316 | 114 |
| 134 | 56 |
| 1663 | 502 |
| 18 | 13 |
| 18 | 13 |
| 12 | 10 |

| | |
|---|---|
| 12 | 10 |
| 12 | 10 |
| 44 | 24 |
| 79 | 37 |
| 39 | 22 |
| 16 | 12 |
| 16 | 12 |
| 16 | 12 |
| 14 | 11 |
| 14 | 11 |
| 105 | 46 |
| 60 | 30 |
| 168 | 67 |
| 108 | 47 |
| 165 | 66 |
| 71 | 34 |
| 71 | 34 |
| 10704 | 2917 |
| 153 | 62 |
| 153 | 62 |
| 123 | 52 |
| 138 | 57 |
| 32 | 19 |
| 32 | 19 |
| 91 | 41 |
| 923 | 292 |
| 10646 | 2901 |
| 328 | 117 |
| 25 | 16 |
| 25 | 16 |
| 37 | 21 |
| 25 | 16 |
| 312 | 112 |
| 100 | 44 |
| 197 | 76 |
| 80 | 37 |
| 103 | 45 |
| 7 | 7 |
| 7 | 7 |
| 7 | 7 |
| 50 | 26 |
| 7 | 7 |
| 7 | 7 |
| 7 | 7 |
| 7 | 7 |
| 151 | 61 |
| 502 | 169 |
| 316 | 113 |

| | |
|---|---|
| 86 | 39 |
| 30 | 18 |
| 72 | 34 |
| 23 | 15 |
| 23 | 15 |
| 53 | 27 |
| 236 | 88 |
| 170 | 67 |
| 466 | 158 |
| 40 | 22 |
| 75 | 35 |
| 233 | 87 |
| 1683 | 505 |
| 48 | 25 |
| 48 | 25 |
| 107 | 46 |
| 928 | 292 |
| 56 | 28 |
| 143 | 58 |
| 21 | 14 |
| 21 | 14 |
| 119 | 50 |
| 535 | 178 |
| 292 | 105 |
| 1179 | 363 |
| 70 | 33 |
| 70 | 33 |
| 59 | 29 |
| 485 | 163 |
| 90 | 40 |
| 51 | 26 |
| 244 | 90 |
| 38 | 21 |
| 1637 | 491 |
| 1594 | 479 |
| 184 | 71 |
| 19 | 13 |
| 62 | 30 |
| 62 | 30 |
| 540 | 179 |
| 46 | 24 |
| 46 | 24 |
| 111 | 47 |
| 54 | 27 |
| 117 | 49 |
| 117 | 49 |
| 172 | 67 |
| 9 | 8 |

| | |
|---|---|
| 9 | 8 |
| 65 | 31 |
| 169 | 66 |
| 26 | 16 |
| 26 | 16 |
| 26 | 16 |
| 26 | 16 |
| 9 | 8 |
| 9 | 8 |
| 26 | 16 |
| 26 | 16 |
| 9 | 8 |
| 9 | 8 |
| 17 | 12 |
| 17 | 12 |
| 17 | 12 |
| 17 | 12 |
| 41 | 22 |
| 41 | 22 |
| 223 | 83 |
| 88 | 39 |
| 31 | 18 |
| 31 | 18 |
| 31 | 18 |
| 36 | 20 |
| 36 | 20 |
| 36 | 20 |
| 71 | 33 |
| 71 | 33 |
| 839 | 265 |
| 217 | 81 |
| 275 | 99 |
| 15 | 11 |
| 11 | 9 |
| 11 | 9 |
| 11 | 9 |
| 341 | 119 |
| 106 | 45 |
| 192 | 73 |
| 13 | 10 |
| 13 | 10 |
| 24 | 15 |
| 24 | 15 |
| 208 | 78 |
| 52 | 26 |
| 292 | 104 |
| 63 | 30 |
| 161 | 63 |

| | |
|---|---|
| 205 | 77 |
| 80 | 36 |
| 335 | 117 |
| 335 | 117 |
| 479 | 160 |
| 345 | 120 |
| 932 | 291 |
| 39 | 21 |
| 39 | 21 |
| 678 | 218 |
| 66 | 31 |
| 29 | 17 |
| 34 | 19 |
| 369 | 127 |
| 98 | 42 |
| 22 | 14 |
| 22 | 14 |
| 22 | 14 |
| 162 | 63 |
| 75 | 34 |
| 200 | 75 |
| 61 | 29 |
| 229 | 84 |
| 78 | 35 |
| 7669 | 2109 |
| 27 | 16 |
| 27 | 16 |
| 27 | 16 |
| 465 | 155 |
| 32 | 18 |
| 32 | 18 |
| 285 | 101 |
| 20 | 13 |
| 259 | 93 |
| 201 | 75 |
| 96 | 41 |
| 111 | 46 |
| 111 | 46 |
| 105 | 44 |
| 154 | 60 |
| 6 | 6 |
| 6 | 6 |
| 6 | 6 |
| 6 | 6 |
| 6 | 6 |
| 6 | 6 |
| 151 | 59 |
| 73 | 33 |

| | |
|---|---|
| 1094 | 335 |
| 76 | 34 |
| 59 | 28 |
| 142 | 56 |
| 205 | 76 |
| 532 | 174 |
| 25 | 15 |
| 18 | 12 |
| 82 | 36 |
| 393 | 133 |
| 35 | 19 |
| 35 | 19 |
| 62 | 29 |
| 161 | 62 |
| 85 | 37 |
| 228 | 83 |
| 512 | 168 |
| 30 | 17 |
| 91 | 39 |
| 177 | 67 |
| 43 | 22 |
| 43 | 22 |
| 337 | 116 |
| 54 | 26 |
| 54 | 26 |
| 54 | 26 |
| 16 | 11 |
| 16 | 11 |
| 301 | 105 |
| 200 | 74 |
| 38 | 20 |
| 38 | 20 |
| 46 | 23 |
| 23 | 14 |
| 23 | 14 |
| 23 | 14 |
| 338 | 116 |
| 23 | 14 |
| 156 | 60 |
| 156 | 60 |
| 345 | 118 |
| 86 | 37 |
| 86 | 37 |
| 150 | 58 |
| 113 | 46 |
| 28 | 16 |
| 49 | 24 |
| 49 | 24 |

| | |
|---|---|
| 525 | 171 |
| 107 | 44 |
| 14 | 10 |
| 14 | 10 |
| 14 | 10 |
| 14 | 10 |
| 8 | 7 |
| 8 | 7 |
| 8 | 7 |
| 8 | 7 |
| 8 | 7 |
| 8 | 7 |
| 8 | 7 |
| 8 | 7 |
| 8 | 7 |
| 8 | 7 |
| 63 | 29 |
| 8 | 7 |
| 8 | 7 |
| 8 | 7 |
| 8 | 7 |
| 8 | 7 |
| 41 | 21 |
| 144 | 56 |
| 41 | 21 |
| 266 | 94 |
| 182 | 68 |
| 66 | 30 |
| 454 | 150 |
| 336 | 115 |
| 52 | 25 |
| 12 | 9 |
| 12 | 9 |
| 12 | 9 |
| 12 | 9 |
| 12 | 9 |
| 12 | 9 |
| 12 | 9 |
| 492 | 161 |
| 303 | 105 |
| 10 | 8 |
| 10 | 8 |
| 10 | 8 |
| 10 | 8 |
| 10 | 8 |
| 10 | 8 |
| 10 | 8 |
| 72 | 32 |

| | |
|---|---|
| 154 | 59 |
| 21 | 13 |
| 21 | 13 |
| 21 | 13 |
| 21 | 13 |
| 21 | 13 |
| 44 | 22 |
| 55 | 26 |
| 75 | 33 |
| 78 | 34 |
| 215 | 78 |
| 81 | 35 |
| 435 | 144 |
| 31 | 17 |
| 31 | 17 |
| 1552 | 460 |
| 90 | 38 |
| 93 | 39 |
| 26 | 15 |
| 538 | 174 |
| 47 | 23 |
| 47 | 23 |
| 136 | 53 |
| 39 | 20 |
| 318 | 109 |
| 50 | 24 |
| 127 | 50 |
| 67 | 30 |
| 19 | 12 |
| 149 | 57 |
| 19 | 12 |
| 19 | 12 |
| 19 | 12 |
| 19 | 12 |
| 19 | 12 |
| 19 | 12 |
| 149 | 57 |
| 70 | 31 |
| 34 | 18 |
| 73 | 32 |
| 73 | 32 |
| 1541 | 456 |
| 53 | 25 |
| 42 | 21 |
| 42 | 21 |
| 568 | 182 |
| 91 | 38 |
| 29 | 16 |

| | |
|---|---|
| 29 | 16 |
| 253 | 89 |
| 24 | 14 |
| 24 | 14 |
| 496 | 161 |
| 45 | 22 |
| 59 | 27 |
| 59 | 27 |
| 391 | 130 |
| 122 | 48 |
| 17 | 11 |
| 17 | 11 |
| 17 | 11 |
| 234 | 83 |
| 17 | 11 |
| 17 | 11 |
| 17 | 11 |
| 195 | 71 |
| 48 | 23 |
| 48 | 23 |
| 141 | 54 |
| 110 | 44 |
| 157 | 59 |
| 157 | 59 |
| 104 | 42 |
| 32 | 17 |
| 51 | 24 |
| 32 | 17 |
| 74 | 32 |
| 51 | 24 |
| 32 | 17 |
| 32 | 17 |
| 98 | 40 |
| 80 | 34 |
| 95 | 39 |
| 83 | 35 |
| 83 | 35 |
| 89 | 37 |
| 86 | 36 |
| 22 | 13 |
| 22 | 13 |
| 22 | 13 |
| 22 | 13 |
| 54 | 25 |
| 22 | 13 |
| 806 | 249 |
| 27 | 15 |
| 27 | 15 |

| | |
|---|---|
| 669 | 210 |
| 15 | 10 |
| 43 | 21 |
| 15 | 10 |
| 677 | 212 |
| 35 | 18 |
| 35 | 18 |
| 319 | 108 |
| 5 | 5 |
| 5 | 5 |
| 5 | 5 |
| 5 | 5 |
| 158 | 59 |
| 5 | 5 |
| 5 | 5 |
| 5 | 5 |
| 5 | 5 |
| 5 | 5 |
| 5 | 5 |
| 583 | 185 |
| 681 | 213 |
| 353 | 118 |
| 136 | 52 |
| 171 | 63 |
| 46 | 22 |
| 66 | 29 |
| 102 | 41 |
| 102 | 41 |
| 72 | 31 |
| 72 | 31 |
| 283 | 97 |
| 149 | 56 |
| 30 | 16 |
| 38 | 19 |
| 13 | 9 |
| 13 | 9 |
| 13 | 9 |
| 13 | 9 |
| 13 | 9 |
| 13 | 9 |
| 13 | 9 |
| 20 | 12 |
| 20 | 12 |
| 317 | 107 |
| 20 | 12 |
| 20 | 12 |
| 20 | 12 |
| 20 | 12 |

| | |
|---|---|
| 20 | 12 |
| 143 | 54 |
| 25 | 14 |
| 351 | 117 |
| 211 | 75 |
| 156 | 58 |
| 304 | 103 |
| 198 | 71 |
| 137 | 52 |
| 41 | 20 |
| 112 | 44 |
| 169 | 62 |
| 182 | 66 |
| 106 | 42 |
| 166 | 61 |
| 33 | 17 |
| 33 | 17 |
| 11 | 8 |
| 11 | 8 |
| 11 | 8 |
| 11 | 8 |
| 11 | 8 |
| 11 | 8 |
| 11 | 8 |
| 11 | 8 |
| 103 | 41 |
| 100 | 40 |
| 163 | 60 |
| 64 | 28 |
| 44 | 21 |
| 44 | 21 |
| 44 | 21 |
| 9691 | 2624 |
| 97 | 39 |
| 125 | 48 |
| 125 | 48 |
| 67 | 29 |
| 125 | 48 |
| 94 | 38 |
| 7 | 6 |
| 7 | 6 |
| 7 | 6 |
| 7 | 6 |
| 7 | 6 |
| 7 | 6 |
| 70 | 30 |
| 7 | 6 |
| 7 | 6 |

| | |
|---|---|
| 7 | 6 |
| 7 | 6 |
| 7 | 6 |
| 91 | 37 |
| 73 | 31 |
| 85 | 35 |
| 85 | 35 |
| 76 | 32 |
| 82 | 34 |
| 82 | 34 |
| 82 | 34 |
| 9 | 7 |
| 9 | 7 |
| 9 | 7 |
| 9 | 7 |
| 9 | 7 |
| 9 | 7 |
| 9 | 7 |
| 9 | 7 |
| 9 | 7 |
| 9 | 7 |
| 9 | 7 |
| 9 | 7 |
| 9 | 7 |
| 9 | 7 |
| 9 | 7 |
| 9 | 7 |
| 9 | 7 |
| 9 | 7 |
| 9 | 7 |
| 9 | 7 |
| 9 | 7 |
| 9 | 7 |
| 47 | 22 |
| 28 | 15 |
| 28 | 15 |
| 28 | 15 |
| 28 | 15 |
| 18 | 11 |
| 36 | 18 |
| 18 | 11 |
| 18 | 11 |
| 18 | 11 |
| 18 | 11 |
| 183 | 66 |
| 18 | 11 |
| 18 | 11 |
| 113 | 44 |

| | |
|---|---|
| 167 | 61 |
| 50 | 23 |
| 50 | 23 |
| 23 | 13 |
| 23 | 13 |
| 23 | 13 |
| 110 | 43 |
| 110 | 43 |
| 148 | 55 |
| 53 | 24 |
| 53 | 24 |
| 39 | 19 |
| 39 | 19 |
| 39 | 19 |
| 56 | 25 |
| 142 | 53 |
| 59 | 26 |
| 59 | 26 |
| 31 | 16 |
| 31 | 16 |
| 92 | 37 |
| 92 | 37 |
| 62 | 27 |
| 217 | 76 |
| 42 | 20 |
| 42 | 20 |
| 65 | 28 |
| 86 | 35 |
| 919 | 278 |
| 68 | 29 |
| 80 | 33 |
| 74 | 31 |
| 74 | 31 |
| 74 | 31 |
| 74 | 31 |
| 165 | 60 |
| 16 | 10 |
| 16 | 10 |
| 16 | 10 |
| 16 | 10 |
| 16 | 10 |
| 16 | 10 |
| 16 | 10 |
| 291 | 98 |
| 26 | 14 |
| 26 | 14 |
| 26 | 14 |
| 162 | 59 |

| | |
|---|---|
| 26 | 14 |
| 34 | 17 |
| 34 | 17 |
| 105 | 41 |
| 127 | 48 |
| 208 | 73 |
| 48 | 22 |
| 21 | 12 |
| 21 | 12 |
| 21 | 12 |
| 21 | 12 |
| 418 | 135 |
| 1234 | 365 |
| 96 | 38 |
| 37 | 18 |
| 37 | 18 |
| 37 | 18 |
| 37 | 18 |
| 37 | 18 |
| 189 | 67 |
| 242 | 83 |
| 60 | 26 |
| 29 | 15 |
| 29 | 15 |
| 29 | 15 |
| 81 | 33 |
| 81 | 33 |
| 147 | 54 |
| 219 | 76 |
| 66 | 28 |
| 249 | 85 |
| 69 | 29 |
| 72 | 30 |
| 40 | 19 |
| 40 | 19 |
| 40 | 19 |
| 40 | 19 |
| 40 | 19 |
| 320 | 106 |
| 14 | 9 |
| 14 | 9 |
| 14 | 9 |
| 14 | 9 |
| 14 | 9 |
| 14 | 9 |
| 14 | 9 |
| 14 | 9 |
| 14 | 9 |

| | |
|---|---|
| 14 | 9 |
| 106 | 41 |
| 43 | 20 |
| 43 | 20 |
| 180 | 64 |
| 122 | 46 |
| 24 | 13 |
| 24 | 13 |
| 32 | 16 |
| 32 | 16 |
| 46 | 21 |
| 46 | 21 |
| 19 | 11 |
| 19 | 11 |
| 19 | 11 |
| 19 | 11 |
| 19 | 11 |
| 19 | 11 |
| 19 | 11 |
| 627 | 194 |
| 49 | 22 |
| 247 | 84 |
| 217 | 75 |
| 88 | 35 |
| 264 | 89 |
| 145 | 53 |
| 85 | 34 |
| 1512 | 440 |
| 158 | 57 |
| 35 | 17 |
| 55 | 24 |
| 540 | 169 |
| 79 | 32 |
| 76 | 31 |
| 12 | 8 |
| 12 | 8 |
| 12 | 8 |
| 12 | 8 |
| 12 | 8 |
| 12 | 8 |
| 67 | 28 |
| 12 | 8 |
| 12 | 8 |
| 12 | 8 |
| 27 | 14 |
| 27 | 14 |
| 27 | 14 |
| 27 | 14 |

| | |
|---|---|
| 38 | 18 |
| 601 | 186 |
| 423 | 135 |
| 41 | 19 |
| 242 | 82 |
| 22 | 12 |
| 22 | 12 |
| 609 | 188 |
| 89 | 35 |
| 89 | 35 |
| 30 | 15 |
| 30 | 15 |
| 382 | 123 |
| 17 | 10 |
| 17 | 10 |
| 17 | 10 |
| 17 | 10 |
| 17 | 10 |
| 17 | 10 |
| 17 | 10 |
| 17 | 10 |
| 17 | 10 |
| 17 | 10 |
| 127 | 47 |
| 166 | 59 |
| 140 | 51 |
| 4 | 4 |
| 4 | 4 |
| 4 | 4 |
| 263 | 88 |
| 4 | 4 |
| 10 | 7 |
| 4 | 4 |
| 4 | 4 |
| 10 | 7 |
| 4 | 4 |
| 10 | 7 |
| 77 | 31 |
| 4 | 4 |
| 4 | 4 |
| 10 | 7 |
| 10 | 7 |
| 4 | 4 |
| 4 | 4 |
| 10 | 7 |
| 4 | 4 |
| 10 | 7 |
| 10 | 7 |

| | |
|---|---|
| 4 | 4 |
| 4 | 4 |
| 4 | 4 |
| 4 | 4 |
| 10 | 7 |
| 4 | 4 |
| 10 | 7 |
| 10 | 7 |
| 4 | 4 |
| 10 | 7 |
| 10 | 7 |
| 4 | 4 |
| 10 | 7 |
| 50 | 22 |
| 4 | 4 |
| 10 | 7 |
| 4 | 4 |
| 10 | 7 |
| 4 | 4 |
| 4 | 4 |
| 10 | 7 |
| 4 | 4 |
| 4 | 4 |
| 10 | 7 |
| 4 | 4 |
| 4 | 4 |
| 4 | 4 |
| 4 | 4 |
| 4 | 4 |
| 10 | 7 |
| 4 | 4 |
| 4 | 4 |
| 4 | 4 |
| 10 | 7 |
| 10 | 7 |
| 10 | 7 |
| 4 | 4 |
| 53 | 23 |
| 71 | 29 |
| 71 | 29 |
| 33 | 16 |
| 33 | 16 |
| 33 | 16 |
| 33 | 16 |
| 56 | 24 |
| 56 | 24 |
| 59 | 25 |
| 59 | 25 |

| | |
|---|---|
| 59 | 25 |
| 59 | 25 |
| 59 | 25 |
| 99 | 38 |
| 318 | 104 |
| 778 | 235 |
| 96 | 37 |
| 25 | 13 |
| 25 | 13 |
| 25 | 13 |
| 25 | 13 |
| 25 | 13 |
| 220 | 75 |
| 36 | 17 |
| 36 | 17 |
| 36 | 17 |
| 36 | 17 |
| 36 | 17 |
| 36 | 17 |
| 36 | 17 |
| 112 | 42 |
| 154 | 55 |
| 305 | 100 |
| 8 | 6 |
| 8 | 6 |
| 8 | 6 |
| 8 | 6 |
| 8 | 6 |
| 8 | 6 |
| 8 | 6 |
| 8 | 6 |
| 8 | 6 |
| 8 | 6 |
| 8 | 6 |
| 8 | 6 |
| 8 | 6 |
| 8 | 6 |
| 531 | 165 |
| 87 | 34 |
| 39 | 18 |
| 39 | 18 |
| 138 | 50 |
| 194 | 67 |
| 106 | 40 |
| 6 | 5 |
| 6 | 5 |
| 6 | 5 |
| 6 | 5 |

| | |
|---|---|
| 20 | 11 |
| 6 | 5 |
| 6 | 5 |
| 6 | 5 |
| 6 | 5 |
| 6 | 5 |
| 6 | 5 |
| 20 | 11 |
| 6 | 5 |
| 6 | 5 |
| 6 | 5 |
| 20 | 11 |
| 20 | 11 |
| 6 | 5 |
| 6 | 5 |
| 6 | 5 |
| 20 | 11 |
| 6 | 5 |
| 6 | 5 |
| 6 | 5 |
| 6 | 5 |
| 6 | 5 |
| 6 | 5 |
| 6 | 5 |
| 6 | 5 |
| 6 | 5 |
| 20 | 11 |
| 6 | 5 |
| 1712 | 492 |
| 6 | 5 |
| 15 | 9 |
| 15 | 9 |
| 15 | 9 |
| 15 | 9 |
| 15 | 9 |
| 15 | 9 |
| 15 | 9 |
| 28 | 14 |
| 28 | 14 |
| 28 | 14 |
| 28 | 14 |
| 75 | 30 |
| 75 | 30 |
| 100 | 38 |
| 100 | 38 |
| 100 | 38 |
| 116 | 43 |
| 69 | 28 |

| | |
|---|---|
| 51 | 22 |
| 51 | 22 |
| 66 | 27 |
| 63 | 26 |
| 54 | 23 |
| 57 | 24 |
| 4955 | 1362 |
| 57 | 24 |
| 60 | 25 |
| 57 | 24 |
| 31 | 15 |
| 31 | 15 |
| 31 | 15 |
| 31 | 15 |
| 31 | 15 |
| 31 | 15 |
| 126 | 46 |
| 110 | 41 |
| 91 | 35 |
| 91 | 35 |
| 290 | 95 |
| 149 | 53 |
| 962 | 285 |
| 88 | 34 |
| 23 | 12 |
| 328 | 106 |
| 23 | 12 |
| 23 | 12 |
| 23 | 12 |
| 23 | 12 |
| 34 | 16 |
| 85 | 33 |
| 415 | 131 |
| 485 | 151 |
| 156 | 55 |
| 101 | 38 |
| 37 | 17 |
| 13 | 8 |
| 13 | 8 |
| 13 | 8 |
| 13 | 8 |
| 13 | 8 |
| 40 | 18 |
| 40 | 18 |
| 13 | 8 |
| 13 | 8 |
| 13 | 8 |
| 13 | 8 |

| | |
|---|---|
| 13 | 8 |
| 13 | 8 |
| 40 | 18 |
| 13 | 8 |
| 40 | 18 |
| 40 | 18 |
| 40 | 18 |
| 18 | 10 |
| 95 | 36 |
| 150 | 53 |
| 26 | 13 |
| 26 | 13 |
| 43 | 19 |
| 43 | 19 |
| 67 | 27 |
| 67 | 27 |
| 124 | 45 |
| 124 | 45 |
| 124 | 45 |
| 46 | 20 |
| 61 | 25 |
| 49 | 21 |
| 49 | 21 |
| 55 | 23 |
| 89 | 34 |
| 157 | 55 |
| 157 | 55 |
| 4220 | 2104 |
| 12345 | 4221 |
| 5487 | 2277 |
| 5521 | 2263 |
| 17834 | 5238 |
| 6657 | 2526 |
| 6657 | 2526 |
| 6657 | 2526 |
| 14605 | 4536 |
| 16018 | 4811 |
| 15671 | 4718 |
| 16808 | 4944 |
| 1776 | 831 |
| 1949 | 888 |
| 10074 | 3193 |
| 890 | 471 |
| 951 | 494 |
| 895 | 467 |
| 6591 | 2167 |
| 6592 | 2167 |
| 3770 | 1343 |

| | |
|---|---|
| 402 | 239 |
| 1405 | 597 |
| 419 | 236 |
| 259 | 168 |
| 726 | 349 |
| 1436 | 582 |
| 2203 | 813 |
| 1368 | 552 |
| 865 | 385 |
| 2430 | 880 |
| 4777 | 1558 |
| 182 | 125 |
| 436 | 230 |
| 282 | 168 |
| 21548 | 5634 |
| 171 | 116 |
| 1299 | 515 |
| 1299 | 515 |
| 1629 | 612 |
| 319 | 171 |
| 1689 | 620 |
| 206 | 121 |
| 497 | 230 |
| 671 | 286 |
| 518 | 233 |
| 222 | 123 |
| 177 | 103 |
| 206 | 114 |
| 150 | 91 |
| 313 | 154 |
| 153 | 91 |
| 185 | 103 |
| 1060 | 396 |
| 325 | 155 |
| 108 | 70 |
| 102 | 67 |
| 258 | 129 |
| 1521 | 530 |
| 427 | 187 |
| 103 | 66 |
| 102 | 65 |
| 128 | 76 |
| 9864 | 2789 |
| 182 | 97 |
| 429 | 185 |
| 272 | 130 |
| 8588 | 2451 |
| 736 | 283 |

| | |
|---|---|
| 92 | 58 |
| 1426 | 488 |
| 1240 | 431 |
| 118 | 68 |
| 92 | 57 |
| 83 | 53 |
| 97 | 59 |
| 95 | 58 |
| 567 | 223 |
| 1226 | 425 |
| 2230 | 716 |
| 255 | 118 |
| 74 | 48 |
| 72 | 47 |
| 477 | 189 |
| 2518 | 787 |
| 384 | 158 |
| 92 | 54 |
| 2513 | 784 |
| 202 | 95 |
| 89 | 52 |
| 264 | 116 |
| 73 | 45 |
| 831 | 295 |
| 83 | 49 |
| 37 | 28 |
| 107 | 58 |
| 87 | 50 |
| 171 | 81 |
| 38 | 28 |
| 496 | 188 |
| 905 | 312 |
| 782 | 275 |
| 113 | 59 |
| 159 | 76 |
| 25 | 21 |
| 122 | 62 |
| 1217 | 401 |
| 187 | 85 |
| 622 | 224 |
| 1200 | 395 |
| 515 | 191 |
| 24 | 20 |
| 1194 | 393 |
| 119 | 60 |
| 624 | 224 |
| 47 | 31 |
| 74 | 42 |

| | |
|---|---|
| 201 | 88 |
| 225 | 96 |
| 107 | 54 |
| 489 | 179 |
| 314 | 124 |
| 422 | 158 |
| 153 | 70 |
| 139 | 65 |
| 26 | 20 |
| 48 | 30 |
| 192 | 83 |
| 848 | 285 |
| 432 | 160 |
| 254 | 103 |
| 100 | 50 |
| 21 | 17 |
| 47 | 29 |
| 256 | 103 |
| 4004 | 1154 |
| 16 | 14 |
| 16 | 14 |
| 32 | 22 |
| 30 | 21 |
| 39 | 25 |
| 51 | 30 |
| 446 | 161 |
| 446 | 161 |
| 1468 | 457 |
| 52 | 30 |
| 40 | 25 |
| 31 | 21 |
| 50 | 29 |
| 36 | 23 |
| 21 | 16 |
| 159 | 68 |
| 568 | 195 |
| 64 | 34 |
| 188 | 77 |
| 158 | 67 |
| 28 | 19 |
| 89 | 43 |
| 89 | 43 |
| 89 | 43 |
| 14 | 12 |
| 14 | 12 |
| 16 | 13 |
| 2055 | 612 |
| 38 | 23 |

| | |
|---|---|
| 31 | 20 |
| 105 | 48 |
| 41 | 24 |
| 100 | 46 |
| 11 | 10 |
| 169 | 69 |
| 2389 | 700 |
| 73 | 36 |
| 519 | 176 |
| 30 | 19 |
| 19 | 14 |
| 17 | 13 |
| 174 | 70 |
| 8 | 8 |
| 89 | 41 |
| 43 | 24 |
| 136 | 57 |
| 59 | 30 |
| 2277 | 665 |
| 46 | 25 |
| 46 | 25 |
| 656 | 214 |
| 62 | 31 |
| 10 | 9 |
| 20 | 14 |
| 20 | 14 |
| 39 | 22 |
| 39 | 22 |
| 18 | 13 |
| 12 | 10 |
| 12 | 10 |
| 12 | 10 |
| 14 | 11 |
| 14 | 11 |
| 55 | 28 |
| 55 | 28 |
| 69 | 33 |
| 45 | 24 |
| 107 | 46 |
| 30 | 18 |
| 93 | 41 |
| 512 | 169 |
| 19 | 13 |
| 26 | 16 |
| 26 | 16 |
| 9 | 8 |
| 9 | 8 |
| 128 | 52 |

| | |
|---|---|
| 15 | 11 |
| 15 | 11 |
| 24 | 15 |
| 95 | 41 |
| 55 | 27 |
| 55 | 27 |
| 11 | 9 |
| 13 | 10 |
| 13 | 10 |
| 72 | 33 |
| 34 | 19 |
| 424 | 142 |
| 81 | 36 |
| 212 | 78 |
| 180 | 68 |
| 76 | 34 |
| 6 | 6 |
| 116 | 47 |
| 77 | 34 |
| 86 | 37 |
| 86 | 37 |
| 95 | 40 |
| 16 | 11 |
| 16 | 11 |
| 16 | 11 |
| 807 | 249 |
| 14 | 10 |
| 8 | 7 |
| 8 | 7 |
| 12 | 9 |
| 10 | 8 |
| 10 | 8 |
| 10 | 8 |
| 10 | 8 |
| 26 | 15 |
| 312 | 106 |
| 173 | 64 |
| 29 | 16 |
| 1334 | 393 |
| 24 | 14 |
| 650 | 203 |
| 17 | 11 |
| 314 | 106 |
| 57 | 26 |
| 22 | 13 |
| 99 | 40 |
| 99 | 40 |
| 22 | 13 |

| | |
|---|---|
| 69 | 30 |
| 15 | 10 |
| 15 | 10 |
| 2001 | 572 |
| 5 | 5 |
| 5 | 5 |
| 5 | 5 |
| 5 | 5 |
| 30 | 16 |
| 41 | 20 |
| 13 | 9 |
| 154 | 57 |
| 13 | 9 |
| 13 | 9 |
| 356 | 117 |
| 107 | 42 |
| 11 | 8 |
| 11 | 8 |
| 11 | 8 |
| 28 | 15 |
| 306 | 102 |
| 18 | 11 |
| 7 | 6 |
| 18 | 11 |
| 9 | 7 |
| 23 | 13 |
| 143 | 53 |
| 855 | 257 |
| 176 | 63 |
| 16 | 10 |
| 16 | 10 |
| 16 | 10 |
| 16 | 10 |
| 54 | 24 |
| 29 | 15 |
| 29 | 15 |
| 14 | 9 |
| 82 | 33 |
| 82 | 33 |
| 366 | 118 |
| 82 | 33 |
| 24 | 13 |
| 172 | 61 |
| 73 | 30 |
| 55 | 24 |
| 150 | 54 |
| 12 | 8 |
| 121 | 45 |

| | |
|---|---|
| 12 | 8 |
| 12 | 8 |
| 12 | 8 |
| 334 | 108 |
| 22 | 12 |
| 22 | 12 |
| 22 | 12 |
| 50 | 22 |
| 17 | 10 |
| 17 | 10 |
| 17 | 10 |
| 10 | 7 |
| 10 | 7 |
| 4 | 4 |
| 4 | 4 |
| 4 | 4 |
| 4 | 4 |
| 4 | 4 |
| 4 | 4 |
| 4 | 4 |
| 4 | 4 |
| 4 | 4 |
| 39 | 18 |
| 8 | 6 |
| 8 | 6 |
| 8 | 6 |
| 8 | 6 |
| 8 | 6 |
| 8 | 6 |
| 15 | 9 |
| 6 | 5 |
| 6 | 5 |
| 6 | 5 |
| 6 | 5 |
| 6 | 5 |
| 6 | 5 |
| 6 | 5 |
| 6 | 5 |
| 6 | 5 |
| 279 | 91 |
| 346 | 110 |
| 92 | 35 |
| 55 | 23 |
| 18 | 10 |
| 13 | 8 |
| 13 | 8 |
| 13 | 8 |
| 21 | 11 |

| | |
|---|---|
| 35 | 16 |
| 35 | 16 |
| 318 | 101 |
| 81 | 31 |
| 47 | 20 |
| 11 | 7 |
| 11 | 7 |
| 11 | 7 |
| 16 | 9 |
| 16 | 9 |
| 24 | 12 |
| 24 | 12 |
| 24 | 12 |
| 91 | 34 |
| 215 | 71 |
| 134 | 47 |
| 33 | 15 |
| 33 | 15 |
| 9 | 6 |
| 9 | 6 |
| 9 | 6 |
| 9 | 6 |
| 9 | 6 |
| 9 | 6 |
| 14 | 8 |
| 14 | 8 |
| 762 | 223 |
| 14 | 8 |
| 22 | 11 |
| 22 | 11 |
| 762 | 223 |
| 14838 | 4984 |
| 2350 | 1016 |
| 18370 | 5441 |
| 1853 | 823 |
| 5758 | 2074 |
| 1919 | 836 |
| 1936 | 841 |
| 1500 | 684 |
| 2021 | 865 |
| 2187 | 921 |
| 2188 | 921 |
| 1560 | 696 |
| 2325 | 957 |
| 2103 | 877 |
| 1661 | 725 |
| 6146 | 2145 |
| 1136 | 535 |

| | |
|---|---|
| 623 | 339 |
| 6357 | 2196 |
| 947 | 455 |
| 2452 | 972 |
| 2355 | 935 |
| 602 | 312 |
| 2304 | 899 |
| 2532 | 965 |
| 1752 | 708 |
| 524 | 272 |
| 959 | 432 |
| 399 | 218 |
| 7546 | 2460 |
| 733 | 336 |
| 794 | 357 |
| 795 | 357 |
| 795 | 357 |
| 734 | 334 |
| 484 | 239 |
| 653 | 301 |
| 453 | 224 |
| 248 | 142 |
| 404 | 204 |
| 580 | 270 |
| 787 | 344 |
| 600 | 277 |
| 501 | 240 |
| 251 | 142 |
| 4354 | 1474 |
| 160 | 102 |
| 212 | 124 |
| 697 | 307 |
| 290 | 155 |
| 4444 | 1491 |
| 359 | 181 |
| 492 | 230 |
| 492 | 230 |
| 434 | 208 |
| 285 | 150 |
| 317 | 162 |
| 1002 | 403 |
| 123 | 79 |
| 586 | 255 |
| 490 | 220 |
| 1952 | 700 |
| 278 | 139 |
| 185 | 102 |
| 372 | 174 |

| | |
|---|---|
| 389 | 178 |
| 354 | 164 |
| 197 | 104 |
| 326 | 153 |
| 585 | 245 |
| 308 | 146 |
| 335 | 156 |
| 369 | 167 |
| 59 | 43 |
| 1296 | 477 |
| 5491 | 1744 |
| 2162 | 747 |
| 121 | 70 |
| 271 | 128 |
| 118 | 68 |
| 349 | 156 |
| 2478 | 837 |
| 78 | 50 |
| 83 | 52 |
| 76 | 48 |
| 33 | 27 |
| 81 | 50 |
| 149 | 78 |
| 172 | 87 |
| 790 | 300 |
| 229 | 107 |
| 73 | 45 |
| 228 | 106 |
| 73 | 45 |
| 39 | 29 |
| 69 | 43 |
| 238 | 109 |
| 63 | 40 |
| 847 | 313 |
| 134 | 69 |
| 53 | 35 |
| 83 | 48 |
| 95 | 53 |
| 706 | 266 |
| 60 | 38 |
| 60 | 38 |
| 61 | 38 |
| 99 | 54 |
| 48 | 32 |
| 57 | 36 |
| 44 | 30 |
| 32 | 24 |
| 32 | 24 |

| | |
|---|---|
| 53 | 34 |
| 1654 | 560 |
| 231 | 103 |
| 139 | 69 |
| 185 | 86 |
| 214 | 96 |
| 68 | 40 |
| 52 | 33 |
| 198 | 90 |
| 277 | 118 |
| 391 | 157 |
| 1637 | 551 |
| 40 | 27 |
| 65 | 38 |
| 36 | 25 |
| 412 | 162 |
| 412 | 162 |
| 561 | 211 |
| 111 | 56 |
| 127 | 62 |
| 57 | 34 |
| 42 | 27 |
| 40 | 26 |
| 724 | 261 |
| 335 | 134 |
| 18 | 15 |
| 266 | 110 |
| 281 | 115 |
| 26 | 19 |
| 67 | 37 |
| 227 | 96 |
| 70 | 38 |
| 58 | 33 |
| 1541 | 510 |
| 42 | 26 |
| 86 | 44 |
| 25 | 18 |
| 25 | 18 |
| 25 | 18 |
| 225 | 94 |
| 23 | 17 |
| 113 | 54 |
| 38 | 24 |
| 45 | 27 |
| 80 | 41 |
| 75 | 39 |
| 60 | 33 |
| 41 | 25 |

| | |
|---|---|
| 96 | 47 |
| 118 | 55 |
| 94 | 46 |
| 16 | 13 |
| 76 | 39 |
| 24 | 17 |
| 768 | 268 |
| 61 | 33 |
| 66 | 35 |
| 22 | 16 |
| 35 | 22 |
| 372 | 141 |
| 11 | 10 |
| 96 | 46 |
| 126 | 57 |
| 132 | 59 |
| 78 | 39 |
| 15 | 12 |
| 46 | 26 |
| 19 | 14 |
| 66 | 34 |
| 61 | 32 |
| 134 | 59 |
| 443 | 162 |
| 37 | 22 |
| 37 | 22 |
| 1124 | 373 |
| 479 | 173 |
| 28 | 18 |
| 28 | 18 |
| 62 | 32 |
| 40 | 23 |
| 40 | 23 |
| 122 | 54 |
| 1113 | 368 |
| 12 | 10 |
| 12 | 10 |
| 31 | 19 |
| 1482 | 479 |
| 20 | 14 |
| 20 | 14 |
| 123 | 54 |
| 14 | 11 |
| 14 | 11 |
| 14 | 11 |
| 16 | 12 |
| 41 | 23 |
| 96 | 44 |

| | |
|---|---|
| 473 | 169 |
| 1480 | 477 |
| 34 | 20 |
| 34 | 20 |
| 130 | 56 |
| 275 | 105 |
| 54 | 28 |
| 25 | 16 |
| 32 | 19 |
| 100 | 45 |
| 47 | 25 |
| 9 | 8 |
| 9 | 8 |
| 9 | 8 |
| 21 | 14 |
| 21 | 14 |
| 68 | 33 |
| 28 | 17 |
| 17 | 12 |
| 15 | 11 |
| 33 | 19 |
| 26 | 16 |
| 1203 | 390 |
| 31 | 18 |
| 31 | 18 |
| 24 | 15 |
| 6 | 6 |
| 6 | 6 |
| 6 | 6 |
| 41 | 22 |
| 123 | 52 |
| 106 | 46 |
| 22 | 14 |
| 22 | 14 |
| 109 | 47 |
| 29 | 17 |
| 49 | 25 |
| 73 | 34 |
| 34 | 19 |
| 180 | 71 |
| 52 | 26 |
| 93 | 41 |
| 190 | 74 |
| 42 | 22 |
| 8 | 7 |
| 25 | 15 |
| 25 | 15 |
| 8 | 7 |

| | |
|---|---|
| 25 | 15 |
| 8 | 7 |
| 16 | 11 |
| 16 | 11 |
| 143 | 58 |
| 30 | 17 |
| 14 | 10 |
| 1180 | 379 |
| 10 | 8 |
| 12 | 9 |
| 12 | 9 |
| 89 | 39 |
| 48 | 24 |
| 43 | 22 |
| 121 | 50 |
| 208 | 79 |
| 33 | 18 |
| 62 | 29 |
| 26 | 15 |
| 26 | 15 |
| 19 | 12 |
| 19 | 12 |
| 49 | 24 |
| 85 | 37 |
| 5 | 5 |
| 5 | 5 |
| 5 | 5 |
| 5 | 5 |
| 5 | 5 |
| 5 | 5 |
| 117 | 48 |
| 5 | 5 |
| 17 | 11 |
| 147 | 58 |
| 29 | 16 |
| 15 | 10 |
| 15 | 10 |
| 15 | 10 |
| 22 | 13 |
| 13 | 9 |
| 13 | 9 |
| 13 | 9 |
| 13 | 9 |
| 53 | 25 |
| 13 | 9 |
| 45 | 22 |
| 104 | 43 |
| 7 | 6 |

| | |
|---|---|
| 7 | 6 |
| 7 | 6 |
| 7 | 6 |
| 11 | 8 |
| 11 | 8 |
| 146 | 57 |
| 11 | 8 |
| 11 | 8 |
| 11 | 8 |
| 11 | 8 |
| 48 | 23 |
| 9 | 7 |
| 9 | 7 |
| 198 | 74 |
| 9 | 7 |
| 9 | 7 |
| 25 | 14 |
| 30 | 16 |
| 117 | 47 |
| 18 | 11 |
| 18 | 11 |
| 38 | 19 |
| 483 | 163 |
| 18 | 11 |
| 94 | 39 |
| 148 | 57 |
| 23 | 13 |
| 130 | 51 |
| 28 | 15 |
| 66 | 29 |
| 16 | 10 |
| 16 | 10 |
| 44 | 21 |
| 198 | 73 |
| 2063 | 631 |
| 58 | 26 |
| 21 | 12 |
| 21 | 12 |
| 26 | 14 |
| 26 | 14 |
| 26 | 14 |
| 1383 | 431 |
| 14 | 9 |
| 14 | 9 |
| 14 | 9 |
| 96 | 39 |
| 157 | 104 |
| 140 | 91 |

| | |
|---|---|
| 108 | 75 |
| 84 | 59 |
| 248 | 122 |
| 96 | 56 |
| 246 | 115 |
| 36 | 27 |
| 76 | 46 |
| 68 | 42 |
| 93 | 53 |
| 174 | 86 |
| 155 | 78 |
| 131 | 68 |
| 79 | 46 |
| 116 | 61 |
| 76 | 44 |
| 97 | 53 |
| 79 | 45 |
| 165 | 79 |
| 102 | 53 |
| 249 | 108 |
| 166 | 77 |
| 61 | 35 |
| 162 | 75 |
| 210 | 93 |
| 118 | 58 |
| 86 | 45 |
| 155 | 72 |
| 218 | 95 |
| 56 | 32 |
| 63 | 35 |
| 54 | 31 |
| 121 | 58 |
| 43 | 26 |
| 120 | 57 |
| 157 | 71 |
| 136 | 63 |
| 44 | 26 |
| 168 | 74 |
| 62 | 33 |
| 75 | 38 |
| 88 | 43 |
| 61 | 32 |
| 89 | 43 |
| 84 | 41 |
| 97 | 46 |
| 223 | 92 |
| 72 | 36 |
| 205 | 85 |

| | |
|---|---|
| 34 | 20 |
| 73 | 36 |
| 23 | 15 |
| 89 | 42 |
| 98 | 45 |
| 67 | 33 |
| 158 | 67 |
| 301 | 117 |
| 527 | 193 |
| 227 | 91 |
| 138 | 59 |
| 197 | 80 |
| 41 | 22 |
| 95 | 43 |
| 77 | 36 |
| 210 | 84 |
| 32 | 18 |
| 107 | 47 |
| 23 | 14 |
| 47 | 24 |
| 103 | 45 |
| 148 | 61 |
| 96 | 42 |
| 67 | 31 |
| 29 | 16 |
| 153 | 62 |
| 44 | 22 |
| 139 | 57 |
| 194 | 76 |
| 84 | 37 |
| 60 | 28 |
| 32 | 17 |
| 135 | 55 |
| 138 | 56 |
| 119 | 49 |
| 137 | 55 |
| 120 | 49 |
| 92 | 39 |
| 54 | 25 |
| 101 | 42 |
| 170 | 66 |
| 701 | 400 |
| 677 | 378 |
| 547 | 310 |
| 276 | 180 |
| 1518 | 681 |
| 199 | 132 |
| 202 | 133 |

| | |
|---|---|
| 209 | 136 |
| 717 | 354 |
| 702 | 344 |
| 123 | 87 |
| 624 | 303 |
| 109 | 78 |
| 92 | 69 |
| 92 | 69 |
| 229 | 135 |
| 406 | 211 |
| 228 | 134 |
| 449 | 225 |
| 329 | 175 |
| 115 | 78 |
| 1355 | 569 |
| 359 | 186 |
| 176 | 106 |
| 103 | 71 |
| 136 | 87 |
| 289 | 155 |
| 80 | 58 |
| 145 | 90 |
| 170 | 101 |
| 132 | 83 |
| 71 | 52 |
| 43 | 36 |
| 72 | 52 |
| 160 | 94 |
| 109 | 70 |
| 188 | 105 |
| 59 | 44 |
| 272 | 138 |
| 272 | 138 |
| 56 | 41 |
| 84 | 55 |
| 162 | 90 |
| 65 | 45 |
| 1419 | 565 |
| 202 | 107 |
| 55 | 39 |
| 718 | 305 |
| 308 | 147 |
| 92 | 56 |
| 155 | 84 |
| 34 | 27 |
| 51 | 36 |
| 51 | 36 |
| 184 | 96 |

| | |
|---|---|
| 38 | 29 |
| 120 | 68 |
| 74 | 47 |
| 33 | 26 |
| 35 | 27 |
| 77 | 48 |
| 88 | 53 |
| 147 | 79 |
| 319 | 149 |
| 39 | 29 |
| 41 | 30 |
| 41 | 30 |
| 41 | 30 |
| 67 | 43 |
| 80 | 49 |
| 1234 | 486 |
| 32 | 25 |
| 32 | 25 |
| 32 | 25 |
| 87 | 52 |
| 34 | 26 |
| 38 | 28 |
| 29 | 23 |
| 29 | 23 |
| 31 | 24 |
| 24 | 20 |
| 33 | 25 |
| 197 | 98 |
| 185 | 93 |
| 78 | 47 |
| 37 | 27 |
| 37 | 27 |
| 225 | 109 |
| 85 | 50 |
| 55 | 36 |
| 43 | 30 |
| 47 | 32 |
| 70 | 43 |
| 70 | 43 |
| 64 | 40 |
| 417 | 183 |
| 30 | 23 |
| 32 | 24 |
| 195 | 96 |
| 73 | 44 |
| 56 | 36 |
| 36 | 26 |
| 69 | 42 |

| | |
|---|---|
| 50 | 33 |
| 40 | 28 |
| 40 | 28 |
| 25 | 20 |
| 380 | 168 |
| 63 | 39 |
| 63 | 39 |
| 94 | 53 |
| 27 | 21 |
| 59 | 37 |
| 168 | 84 |
| 31 | 23 |
| 31 | 23 |
| 2545 | 931 |
| 47 | 31 |
| 43 | 29 |
| 41 | 28 |
| 41 | 28 |
| 41 | 28 |
| 133 | 69 |
| 11 | 11 |
| 26 | 20 |
| 28 | 21 |
| 28 | 21 |
| 28 | 21 |
| 19 | 16 |
| 48 | 31 |
| 30 | 22 |
| 149 | 75 |
| 130 | 67 |
| 36 | 25 |
| 88 | 49 |
| 70 | 41 |
| 68 | 40 |
| 51 | 32 |
| 103 | 55 |
| 80 | 45 |
| 80 | 45 |
| 71 | 41 |
| 71 | 41 |
| 13 | 12 |
| 29 | 21 |
| 41 | 27 |
| 35 | 24 |
| 202 | 95 |
| 50 | 31 |
| 107 | 56 |
| 107 | 56 |

| | |
|---|---|
| 48 | 30 |
| 15 | 13 |
| 68 | 39 |
| 68 | 39 |
| 57 | 34 |
| 24 | 18 |
| 57 | 34 |
| 28 | 20 |
| 34 | 23 |
| 74 | 41 |
| 23 | 17 |
| 25 | 18 |
| 25 | 18 |
| 25 | 18 |
| 25 | 18 |
| 25 | 18 |
| 33 | 22 |
| 27 | 19 |
| 31 | 21 |
| 94 | 49 |
| 46 | 28 |
| 46 | 28 |
| 46 | 28 |
| 46 | 28 |
| 46 | 28 |
| 46 | 28 |
| 121 | 60 |
| 78 | 42 |
| 192 | 88 |
| 11 | 10 |
| 18 | 14 |
| 8 | 8 |
| 20 | 15 |
| 32 | 21 |
| 32 | 21 |
| 30 | 20 |
| 186 | 85 |
| 96 | 49 |
| 96 | 49 |
| 96 | 49 |
| 96 | 49 |
| 43 | 26 |
| 1946 | 707 |
| 61 | 34 |
| 52 | 30 |
| 52 | 30 |
| 99 | 50 |
| 41 | 25 |

| | |
|---|---|
| 66 | 36 |
| 15 | 12 |
| 15 | 12 |
| 83 | 43 |
| 17 | 13 |
| 17 | 13 |
| 17 | 13 |
| 305 | 129 |
| 44 | 26 |
| 153 | 71 |
| 292 | 124 |
| 10 | 9 |
| 138 | 65 |
| 27 | 18 |
| 25 | 17 |
| 23 | 16 |
| 101 | 50 |
| 106 | 52 |
| 49 | 28 |
| 38 | 23 |
| 7 | 7 |
| 82 | 42 |
| 12 | 10 |
| 102 | 50 |
| 90 | 45 |
| 52 | 29 |
| 59 | 32 |
| 120 | 57 |
| 182 | 81 |
| 28 | 18 |
| 103 | 50 |
| 39 | 23 |
| 18 | 13 |
| 22 | 15 |
| 22 | 15 |
| 37 | 22 |
| 37 | 22 |
| 134 | 62 |
| 53 | 29 |
| 89 | 44 |
| 119 | 56 |
| 72 | 37 |
| 35 | 21 |
| 35 | 21 |
| 9 | 8 |
| 44 | 25 |
| 70 | 36 |
| 42 | 24 |

| | |
|---|---|
| 42 | 24 |
| 42 | 24 |
| 42 | 24 |
| 42 | 24 |
| 68 | 35 |
| 54 | 29 |
| 11 | 9 |
| 54 | 29 |
| 6 | 6 |
| 149 | 67 |
| 38 | 22 |
| 144 | 65 |
| 59 | 31 |
| 36 | 21 |
| 36 | 21 |
| 13 | 10 |
| 13 | 10 |
| 23 | 15 |
| 155 | 69 |
| 21 | 14 |
| 19 | 13 |
| 34 | 20 |
| 132 | 60 |
| 19 | 13 |
| 17 | 12 |
| 34 | 20 |
| 48 | 26 |
| 60 | 31 |
| 30 | 18 |
| 30 | 18 |
| 39 | 22 |
| 46 | 25 |
| 46 | 25 |
| 28 | 17 |
| 28 | 17 |
| 8 | 7 |
| 8 | 7 |
| 8 | 7 |
| 51 | 27 |
| 26 | 16 |
| 26 | 16 |
| 35 | 20 |
| 49 | 26 |
| 49 | 26 |
| 91 | 43 |
| 22 | 14 |
| 10 | 8 |
| 10 | 8 |

| | |
|---|---|
| 40 | 22 |
| 167 | 72 |
| 112 | 51 |
| 138 | 61 |
| 112 | 51 |
| 31 | 18 |
| 1767 | 630 |
| 14 | 10 |
| 14 | 10 |
| 14 | 10 |
| 14 | 10 |
| 14 | 10 |
| 14 | 10 |
| 14 | 10 |
| 14 | 10 |
| 14 | 10 |
| 97 | 45 |
| 87 | 41 |
| 5 | 5 |
| 5 | 5 |
| 82 | 39 |
| 36 | 20 |
| 113 | 51 |
| 252 | 103 |
| 244 | 100 |
| 41 | 22 |
| 41 | 22 |
| 25 | 15 |
| 297 | 119 |
| 68 | 33 |
| 114 | 51 |
| 210 | 87 |
| 32 | 18 |
| 32 | 18 |
| 39 | 21 |
| 138 | 60 |
| 56 | 28 |
| 21 | 13 |
| 21 | 13 |
| 30 | 17 |
| 7 | 6 |
| 7 | 6 |
| 7 | 6 |
| 7 | 6 |
| 7 | 6 |
| 7 | 6 |
| 7 | 6 |
| 115 | 51 |

| | |
|---|---|
| 115 | 51 |
| 64 | 31 |
| 64 | 31 |
| 42 | 22 |
| 28 | 16 |
| 780 | 288 |
| 17 | 11 |
| 87 | 40 |
| 47 | 24 |
| 15 | 10 |
| 15 | 10 |
| 15 | 10 |
| 9 | 7 |
| 9 | 7 |
| 15 | 10 |
| 9 | 7 |
| 15 | 10 |
| 15 | 10 |
| 9 | 7 |
| 15 | 10 |
| 15 | 10 |
| 57 | 28 |
| 13 | 9 |
| 13 | 9 |
| 13 | 9 |
| 13 | 9 |
| 11 | 8 |
| 11 | 8 |
| 11 | 8 |
| 216 | 88 |
| 143 | 61 |
| 93 | 42 |
| 24 | 14 |
| 24 | 14 |
| 24 | 14 |
